# Evolution and systematics of the Aculeata and kin (Hymenoptera), with emphasis on the ants (Formicoidea: †@@@idae fam. nov., Formicidae)

**DOI:** 10.1101/2022.02.20.480183

**Authors:** Brendon E. Boudinot, Ziad Khouri, Adrian Richter, Zachary H. Griebenow, Thomas van de Kamp, Vincent Perrichot, Phillip Barden

## Abstract

Fossils provide unique opportunity to understand the tempo and mode of evolution and are essential for modeling the history of lineage diversification. Here, we interrogate the Mesozoic fossil record of the Aculeata, with emphasis on the ants (Formicidae), and conduct an extended series of ancestral state estimation exercises on distributions of tip-dated combined-evidence phylogenies. We developed and illustrated from ground-up a series of 576 morphological characters which we scored for 144 extant and 431 fossil taxa, including all families of Aculeata, Trigonaloidea, Evanioidea, and †Ephialtitoidea. We used average posterior probability support to guide composition of a target matrix of 303 taxa, for which we integrated strongly filtered ultraconserved element (UCE) data for 115 living species. We also implemented reversible jump MCMC (rjMCMC) and hidden state methods to model complex behavioral characters to test hypotheses about the pathway to obligate eusociality. In addition to revising the higher classification of all sampled groups to family or subfamily level using estimated character polarities to diagnose nodes across the phylogeny, we find that the mid-Cretaceous genera †*Camelomecia* and †*Camelosphecia* form a clade which is robustly supported as sister to all living and fossil Formicidae. For this reason, we name this extinct clade as †@@@idae **fam. nov.** and provide a definition for the expanded Formicoidea. Based on our results, we recognize three major phases in the early evolution of the ants: (1) origin of Formicoidea as ground-adapted huntresses during the Late Jurassic in the “stinging aggressor” guild (Aculeata) among various lineages of “sneaking parasitoids” (non-aculeate Vespina); (2) the first formicoid radiation during the Early Cretaceous, by the end of which all major extant linages originated; and (3) turnover of the Formicoidea at the end-Cretaceous leading to the second formicoid radiation. We conclude with a concentrated series of considerations for future directions of study with this dataset and beyond.

## Introduction

Ants (Formicidae) and other Aculeata are among the most familiar of insects. Advances in the classification and systematics of these groups and the Hymenoptera more broadly have proceeded over the past several hundred years in waves following new logical and technological innovations. These innovations range from the nomenclature of Linnaeus (1758) to the cladistics of Hennig (1981), and from the naked eyes to optical and scanning electron microscopes, with the ramifications of micro-computed tomography (µ-CT) yet to be realized. The most profound transformation in recent memory has been the analysis of nucleotide sequence data, which has become essentially industrial in infrastructural scale and massive in data scale. While early phylogenetic studies of insects using Sanger sequences were met with varying degrees of resistance (Kjer *et al*. 2016a), the scope of phylogenomic data have basically rendered the question of relationship among organisms a statistical problem (*e.g.*, Kjer *et al*. 2016b, Young & Gillung 2019). Two key insights arising from the phylogenomic revolution include the sistergroup relationship between ants and Apoidea (Johnson *et al*. 2013) and, indeed, the near total resolution of the backbone phylogeny of the Hymenoptera (Peters *et al*. 2017, Branstetter *et al*. 2017b). As the field of systematics and evolutionary biology has settled into the groove of “big data”, it has become increasingly apparent that a key ingredient is missing: Geochronological evidence for organismal diversification, transformation, and turnover from the fossil record (Marshall 2017, Louca & Pennell 2020).

In the present study, we interrogate the fossil record of Aculeata and kin, integrate morphological and genomic data for combined analysis, and model the transformations of a series of traits, following the independent leads of Rasnitsyn (1975), Brothers (1975), Bolton (2003), Pagel *et al*. (2004), Ronquist *et al*. (2013), and Branstetter *et al*. (2017b). Our objectives, most broadly, were to: (1) apply tip-dating phylogenetic methods to a sample of ants, other Aculeata, and outgroups; (2) reconstruct the paleomorphological evolution of these groups; (3) evaluate the informativeness of fossils for the origins of eusociality; and (4) systematically revise the classification of the total clade Aculeata and their sampled relatives. Toward these ends, we developed and illustrated 576 morphological characters which we scored for a comprehensive sample of Mesozoic Aculeata and kin (totaling over 300,000 point-observations across all families). We combined these observations with stringently filtered ultraconserved element (UCE) data to estimate the topology, timing, and ancestral states for all characters and nodes in our sampling. We also conducted an extended series of modeling exercises on a smaller set of behavioral characters, implementing reversible jump MCMC (rjMCMC, *e.g.*, Pagel & Meade 2006) and hidden state models (*e.g.*, Beaulieu *et al*. 2013). Based on our study and analyses, we provide a review of the fossil record of Hymenoptera focused on ants and other Aculeata, a revised classification of these groups based on estimated character polarities, and a series of new and revised evolutionary hypotheses for ants in the context of the total clade Aculeata. Our results are divided into five parts, with part- and section-specific introductions and discussions provided therein.

### Materials and methods

#### Overview

Our study began with two workflows: **(1)** Generation and analysis of morphological data (Fig. 1, steps 1–3), and **(2)** generation and collation of genomic data for combined phylogenetic analysis (Fig. 1, step 4). From this point, our workflow divides and becomes more complex. In brief overview, the main effort partitions are as follows: **(3)** Scaffolded combined-evidence tip-dating topology searches using a matrix with “diversified” taxon sampling (*sensu* Zhang *et al*. 2016; Fig. 1, step 5), and **(4)** time-calibrated ancestral state estimation (ASE) using a matrix with denser taxon sampling (Fig. 1, steps 6–10). For the ASE branch of this study, we analyzed the morphological data under tip-dating **(5)** (Fig. 1, step 6) and extant-only conditions **(6)** (Fig. 1, steps 8, 10); we also conducted ASE for behavioral characters under these two regimes due to results arising from our collective topology searches **(7, 8)**.

**Fig. 1.**
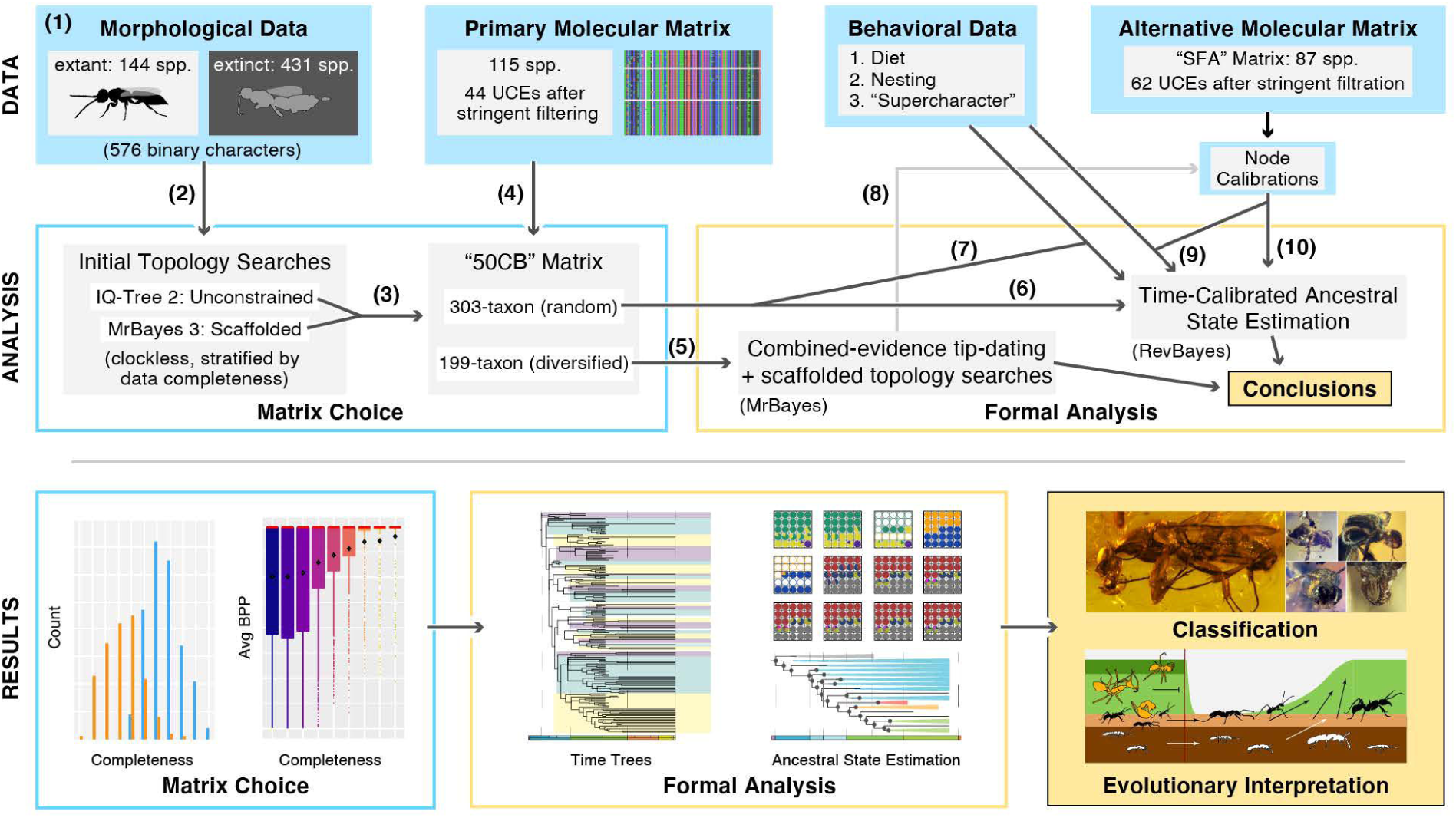
Study design graphical overview. **(1)** We generated a morphological matrix ***de novo*** of 576 binary-characters for 575 taxa; **(2)** after analyzing these data stratified by completeness per taxon, **(3)** we constructed the “50CB” matrix which included all taxa which were ≥ 50% complete plus the most complete †Bethylonymidae; **(4)** we combined these data with 44 stringently filtered UCEs resulting in a 303-taxon matrix with “random” taxon sampling; **(5)** for topology searches and tip-dating, we pruned this combined matrix down to 199 taxa, resulting in a “diversified” taxon sampling; **(6)** for time-calibrated ancestral state estimation, we employed the denser (303-taxon 50CB) sampling for all 576 morphological characters plus (7) a series of behavioral data across several transition models. **(8)** To further explore the effect of our fossil data on dating and transformation series inference, we used calibrations from our combined-evidence tip-dating results for **(9, 10)** divergence dating and time-calibrated ancestral state estimation analyses using an extant-only matrix (SFA). From these collective results we revised the classification of the Aculeata, Trigonaloidea, Evanioidea, and †Ephialtitoidea, and provided hypotheses of Mesozoic evolution for the Hymenoptera, emphasizing the Formicoidea.

#### Section 1: Specimen-based methods

##### Specimens

For extended information on data source by terminal, see the raw morphological data matrix (File X, doi: XXX on Zenodo). For extant taxa, the majority of non-ant aculeates were scored from the UCDC, while all ant specimens were scored from the UCDC with supplemental image vouchering via AntWeb (https://www.antweb.org/). In contrast, the majority of Mesozoic non-ant aculeates were scored from the literature, supplemented by material from the AMNH, BEBC, CELC, and CNUC. All literature used for scoring is cited in the data matrix and included in the references below. Other listed collections were extensively used during the early character development stages. Sources for the image panels used to illustrate the character matrix are provided in Supplemental File S1.

##### Museum abbreviations

Specimens were directly examined from the following museum collections:

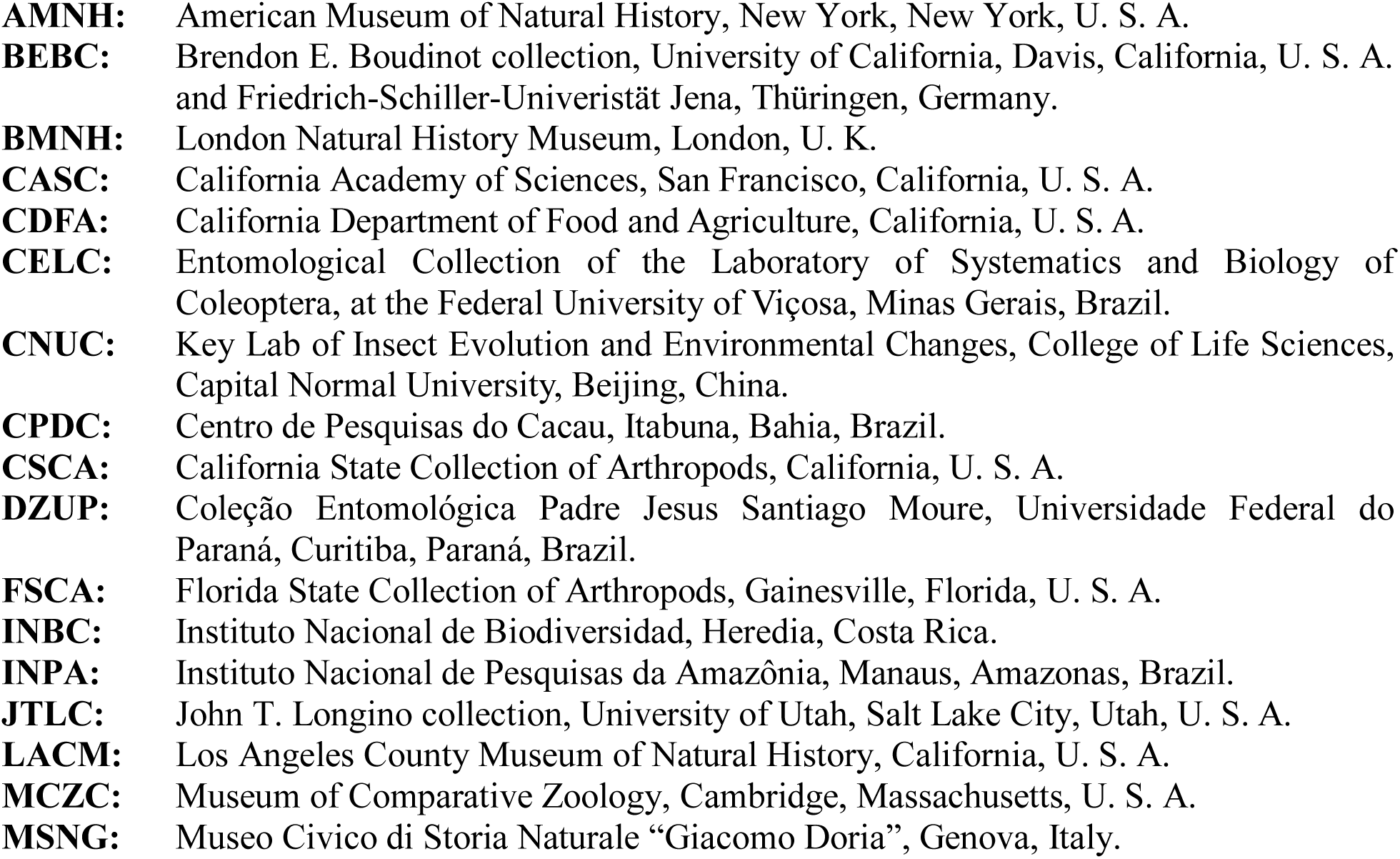

|  |  |
| --- | --- |
| **AMNH:** | American Museum of Natural History, New York, New York, U. S. A. |
| **BEBC:** | Brendon E. Boudinot collection, University of California, Davis, California, U. S. A. and Friedrich-Schiller-Universität Jena, Thüringen, Germany. |
| **BMNH:** | London Natural History Museum, London, U. K. |
| **CASC:** | California Academy of Sciences, San Francisco, California, U. S. A. |
| **CDFA:** | California Department of Food and Agriculture, California, U. S. A. |
| **CELC:** | Entomological Collection of the Laboratory of Systematics and Biology of Coleoptera, at the Federal University of Viçosa, Minas Gerais, Brazil. |
| **CNUC:** | Key Lab of Insect Evolution and Environmental Changes, College of Life Sciences, Capital Normal University, Beijing, China. |
| **CPDC:** | Centro de Pesquisas do Cacau, Itabuna, Bahia, Brazil. |
| **CSCA:** | California State Collection of Arthropods, California, U. S. A. |
| **DZUP:** | Coleção Entomológica Padre Jesus Santiago Moure, Universidade Federal do Paraná, Curitiba, Paraná, Brazil. |
| **FSCA:** | Florida State Collection of Arthropods, Gainesville, Florida, U. S. A. |
| **INBC:** | Instituto Nacional de Biodiversidad, Heredia, Costa Rica. |
| **INPA:** | Instituto Nacional de Pesquisas da Amazônia, Manaus, Amazonas, Brazil. |
| **JTLC:** | John T. Longino collection, University of Utah, Salt Lake City, Utah, U. S. A. |
| **LACM:** | Los Angeles County Museum of Natural History, California, U. S. A. |
| **MCZC:** | Museum of Comparative Zoology, Cambridge, Massachusetts, U. S. A. |
| **MSNG:** | Museo Civico di Storia Naturale “Giacomo Doria”, Genova, Italy. |

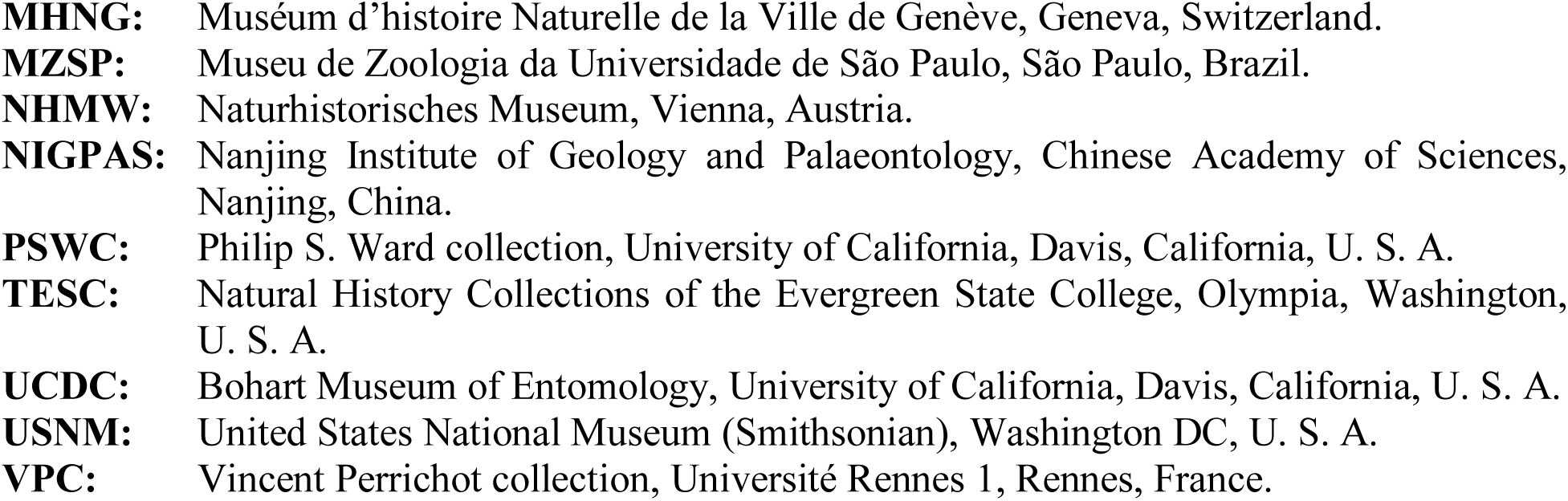

##### Micro-CT scanning and processin

Synchrotron microtomographic scans were performed at the imaging cluster of the KIT light source using a parallel polychromatic X-ray beam produced by a 1.5 T bending magnet. The beam was spectrally filtered by 0.7 mm aluminum with a spectrum peak at about 15 keV. We employed a fast indirect detector system, consisting of a 12 µm LSO:Tb scintillator (Cecilia *et al*., 2011), and a diffraction limited optical microscope (Optique Peter) (Douissard *et al*., 2012) coupled with a 12bit pco.dimax high speed camera with 2016 x 2016 pixels (dos Santos Rolo *et al*., 2014).

Scans were done by taking 3,000 projections at 70 fps and optical magnifications of 5x and 10X, resulting in effective pixel sizes of 2.44 µm and 1.22 µm, respectively. The control system concert (Vogelgesang *et al*., 2016) was used for automated data acquisition and online reconstruction of tomographic slices for data quality assurance. Online and final data processing including tomographic reconstruction were done by the UFO framework (Vogelgesang *et al*., 2012). Before tomographic reconstruction, we performed flat-field correction of the projections and applied phase retrieval based on the transport of intensity equation (Paganin *et al*., 2002).

Microtomographic image sequences were imported to Amira 6.0 (Visage Imaging GmbH, Berlin, Germany) and cropped to the relevant body parts for easier data handling. Individual structures were segmented using the brush and magic wand tools of Amira and extracted using the arithmetic function. They were exported as TIFF image series and imported to VG Studio Max 2.0 (Volume Graphics GmbH, Heidelberg, Germany) to visualize as volume renders. Imaris 6.2.1 (Bitplane AG, Zürich, Switzerland) was used to create smoothed surface renders of selected structures. Surfaces of these structures were exported as inventor scenes (.iv files) which were then transformed into Wavefront-files (.obj) with Amira 6.0. Additional segmentations and digital surface reconstructions were conducted using Dragonfly 4.1 (Object Research Systems (ORS) Inc., Montreal, Canada).

##### Morphological terminology

In general, we employed terminology from the following sources: Richter *et al*. (2019, 2020, 2021) for the head; Goulet & Huber (1993) and Boudinot (2015) for the mesosoma and its appendages; Bolton (1990a,b) and Goulet & Huber (1993) for the metasoma. Concepts from these sources were checked against and supplemented by the Hymenoptera Anatomy Ontology (Yoder *et al*., 2010; Hymenoptera Anatomy Consortium, 2021), with comparative reference to the *Drosophila* Anatomy Ontology (Costa *et al*., 2013; OLS, 2021). A number of modifications were implemented in order to obtain maximum specificity in the character state definitions; these are indicated in the notes sections of the affected state definitions. To improve repeatability of character state scoring, we provide here an outline of our orientation system, the venational system, and a set of new morphological concepts which were employed in the work.

*Orientation*. Body symmetry and positional specification are conserved processes of development (Lipshitz, 1991; Nüsslein-Volhard, 1991). We recognize the anteroposterior (“a-p”), the dorsoventral (“d-v”), the lateromedial (“l-m”), the oral-aboral (“o-a”), and the proximodistal (“p-d”) axes, which in combination form the sagittal (a-p, d-v), the frontal (a-p, l-m), and the transverse (d-v, l-m) planes (Fig. 2). We refer to the exact sagittal midline of the body as the “bilateral symmetry line”, or “bilateral line”. Due to preservational variation in appendage orientation, we recognized specific coordinate systems for the antennae, legs, and wings. The antennae were described as if they were oriented in the sagittal plane, directly away from the cranium, thus having defined medial, lateral, oral, and aboral surfaces (Fig. 3). The legs were described as if they were positioned in the transverse plane, directly away from the mesosoma, thus having defined anterior, posterior, dorsal, and ventral surfaces (Fig. 3). Although the fore legs of ants are modified such that they are more-or-less oriented in the sagittal plane, they are described as if they were in the transverse plane. The wings were described as if they were oriented in the frontal plane, thus having defined dorsal and ventral surfaces, and anterior (costal), distal (apical), and posterior (anal) margins (Fig. 4). Because the heads of the sampled Hymenoptera vary in mandibular orientation (*i.e.*, hypognathy and prognathy), we described this tagma as a strongly indivuated body region, *i.e.*, as if it were separate from the postcephalic soma. The coordinate system used for the heads recognized the oral-aboral, the dorsoventral, and the lateromedial axes (Fig. 2A, B). We recognize the frontal and abfrontal surfaces and areas of the head (Fig. 2C, D); the former refers to the anatomical area of the cranial sclerite that includes the clypeus and antennal foramina, and the latter refers to the anatomical area on the head opposite of the frontal surface and that includes the hypostoma and posterior tentorial pits. This results in the recognition of the frontoabfrontal axis of the head.

**Fig. 2.**
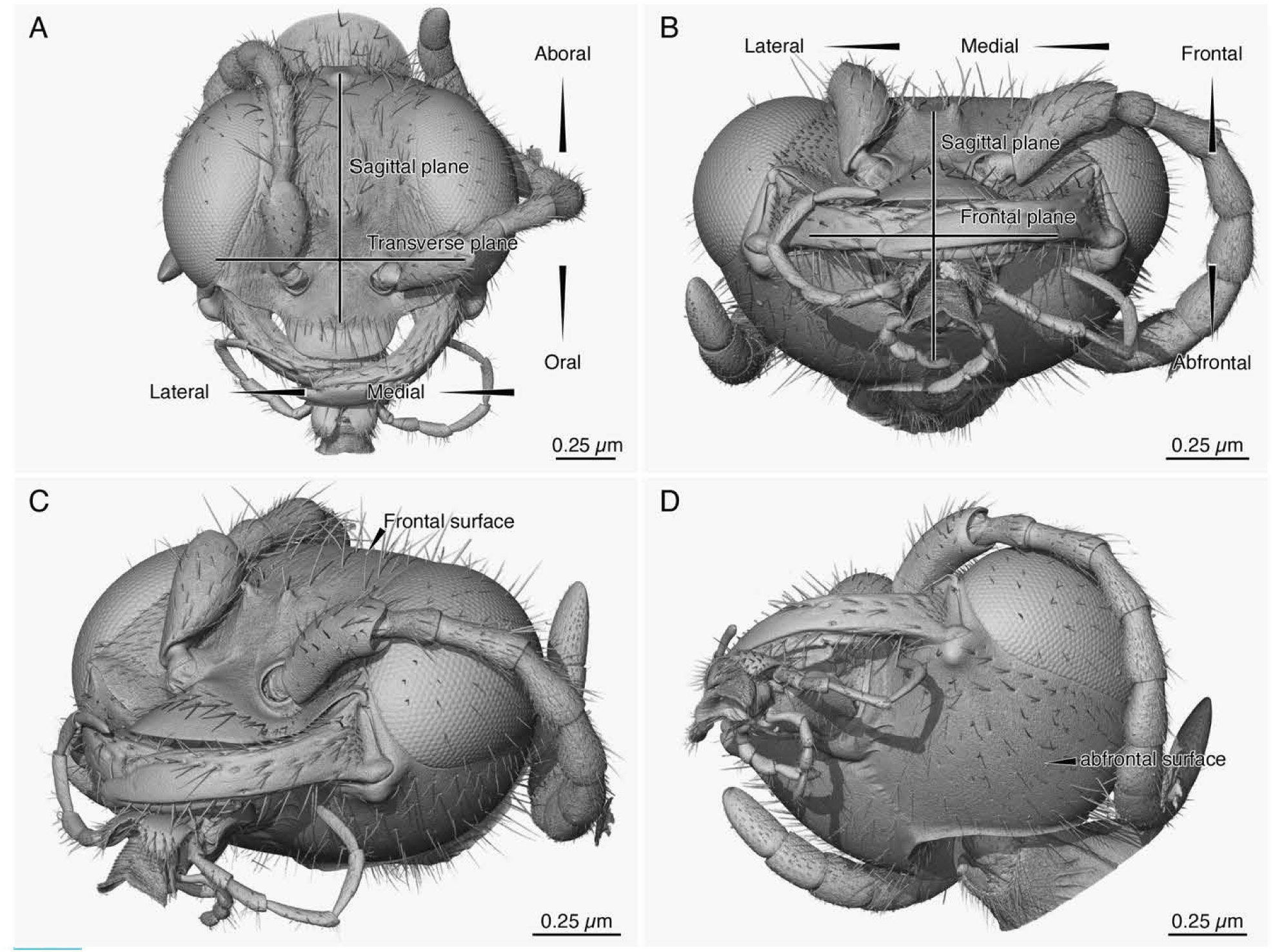
Overview of the body axes of the head. **(A)** Head in facial view, *i.e.*, viewed in the frontal plane, with the oral-aboral axis aligned in the sagittal plane and the lateromedial axis aligned in the transverse plane. **(B)** Head in oral view, *i.e.*, viewed in the transverse plane, with the frontal-abfrontal aligned in the sagittal plane and the lateromedial axis aligned in the frontal plane. **(C)** Dorsally oblique oral-lateral view, emphasizing the frontal surface of the head. **(D)** Ventrally oblique oral-lateral view, emphasizing the abfrontal surface of the head. Taxon: *Methoca* (*Dryinopsis*) nr. *simpliceps* (CASENT0796990).

**Fig. 3.**
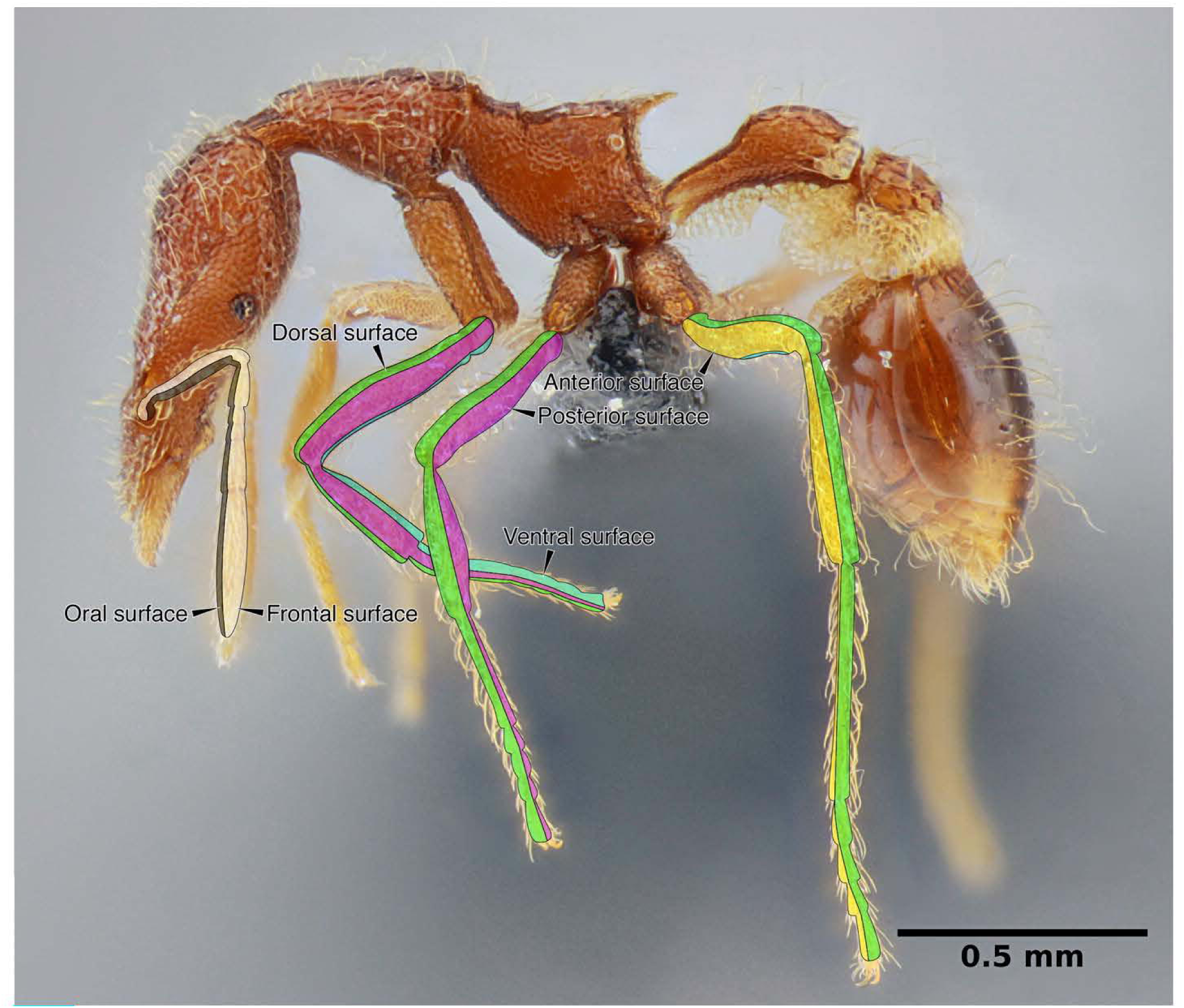
Overview of antenna and leg surface orientation. Black = oral, white = frontal, green = dorsal, magenta = posterior, cyan = ventral, anterior = yellow. Taxon: *Strumigenys* ufv-11, imaged by Júlio Chaul.

**Fig. 4.**
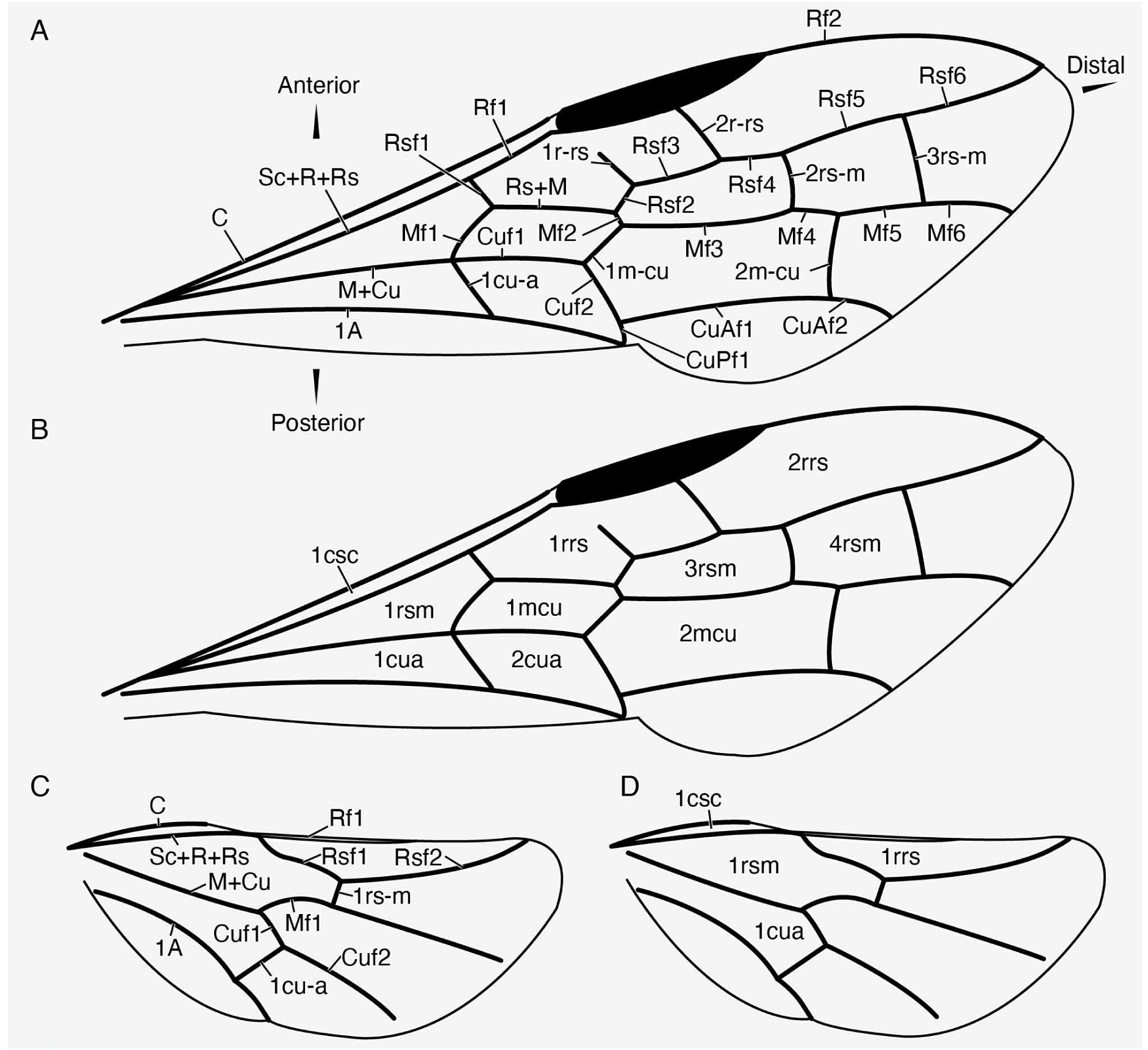
Demonstration of the abscissa-based wing venation system used in the present work. **(A, B)** †*Stephanogaster pristinus* Rasnitsyn & Zhang, 2010 (after Rasnitsyn & Zhang 2010), showing a “generalized” fore wing pattern. **(C, D)** †*Praeproapocritus flexus* Li *et al.,* 2013 (after Ren *et al.* 2019), showing a “generalized” hind wing pattern. **(A, B)** Venation labels. **(B, D)** Cell labels. ***Longitudinal vein labels*:** C = costal vein; Sc+R+Rs = subcostal, radial, plus sectoral composite vein; Rf1, Rf2 = first and second free radial abscissae; Rsf1-6 = first through sixth free sectorial abscissae; Rs+M = sectorial plus medial composite abscissa; M+Cu = medial plus cubital composite vein; Mf1-6 = first through sixth free medial abscissae; Cuf1, 2 = first and second free cubital abscissae; CuAfl, 2 = first and second free abscissae of the anterior cubital branch; CuPf1 = first free abscissa of posterior cubital branch; 1A = first anal vein. ***Crossvein labels*:** 1r-rs, 2r-rs = first and second radial-sectorial crossveins; 2rs-m, 3rs-m = second and third sectorial-medial crossveins (*note: 1rs-m obliterated by fusion of Rs and M*); lm-cu, 2m-cu = first and second medial-cubital crossveins; 1cu-a = first cubital-anal crossvein. ***Cell labels*:** 1csc = costal cell; 1rrs, 2rrs = first and second radial cells (“first submarginal” and “marginal” cells); lrsm, 3rsm, 4rsm = first, third, and fourth sectorial cells (“basal”, “second and third submarginal” cells) (*note: 2rsm obliterated by Rs+M*); 1mcu, 2mcu = first and second medial cells (“first and second discal” cells); 1cua, 2cua = first and second cubital cells (“subbasal” and “subdiscal” cells).

*Venation*. Venational terminology is that of the Comstock-Needham-Ross system (Ross 1936), with certain modifications proposed by Hamilton (1972b), and application of the Brown-Nutting abscissal nomenclature for consistent vein segments (Brown & Nutting 1949); see Fig. 4 for an example of the system employed here. Unfortunately, we were unable to apply the differentiating criteria of Schubnel *et al*. (2020) for recognition of the postcubital veins. For maximal clarity, cell (“domain”, Salcedo *et al*., 2019) names are described using both the Comstock-Needham system and the homology-neutral terminology of Michener (1944) as employed by Gauld & Bolton (1988).

*Contact surfaces*. Contact surfaces are those external, complementary areas on two opposing sclerites which come into frequent contact due to motion and are very often margined with a distinct rim or carina. The surface receiving the moving structure is often depressed relative to surrounding cuticle, thus forming a “scrobe”. Given the widespread occurrence of these features throughout the Hymenoptera, and because most of these have not been recognized in Formicidae or other groups, we provide terms for specific contact surfaces. Terms were constructed with the word root for the mobile structure forming the first part, and the word root for the static surface forming the second. Contact surfaces and their rims, when developed, were observed to occur in the following areas, but also exist elsewhere (Figs. 5, 6).

**Fig. 5.**
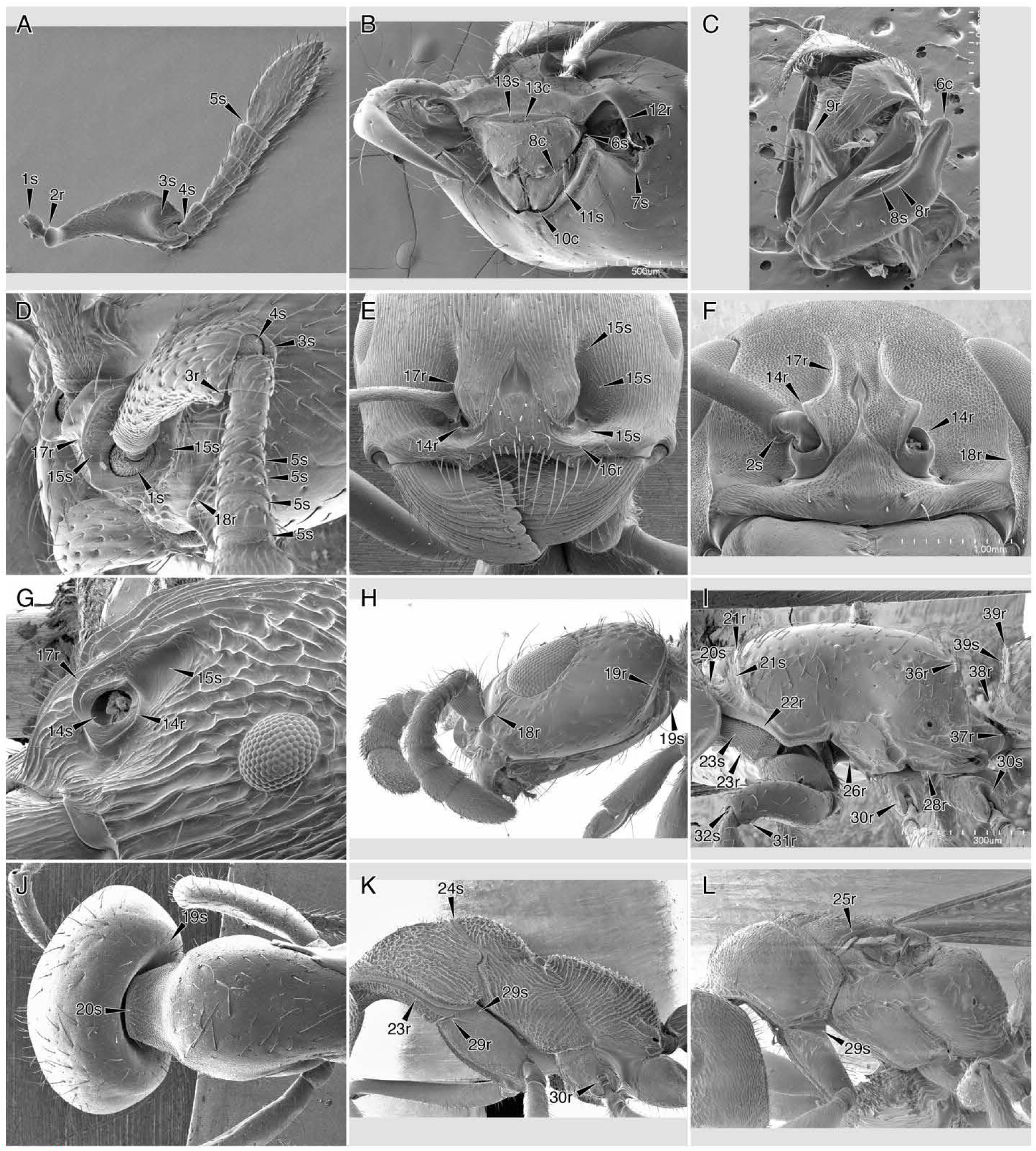
Contact surfaces and rims of the head and mesosoma. Numbers refer to indices in the text; s = contact surface; r = contact rim; c = complementary structures. Taxa: (A) *Tatuidris tatusia,* ANTWEB1008593; (B, J) *Cheliomyrmex morosus,* ANTWEB 1008512; (C) *Paraponera clavata,* ANTWEB1008572; (D, I) *Parasyscia nitidula,* ANTWEB1008511; (E) *Pogonomyrmex barbatus,* ANTWEB 1008577; (F) *Neoponera apicalis,* ANTWEB1008561; (G) *Rhytidoponera confusa,* ANTWEB 1008587; (H) *Simopone schoutedeni,* ANTWEB1008590; (K) *Myrmecia nigris capa,* ANTWEB1008552; (L) *Fulakora mystriops,* ANTWEB1008500. (All images by Roberto Keller, AntWeb.)

**Fig. 6.**
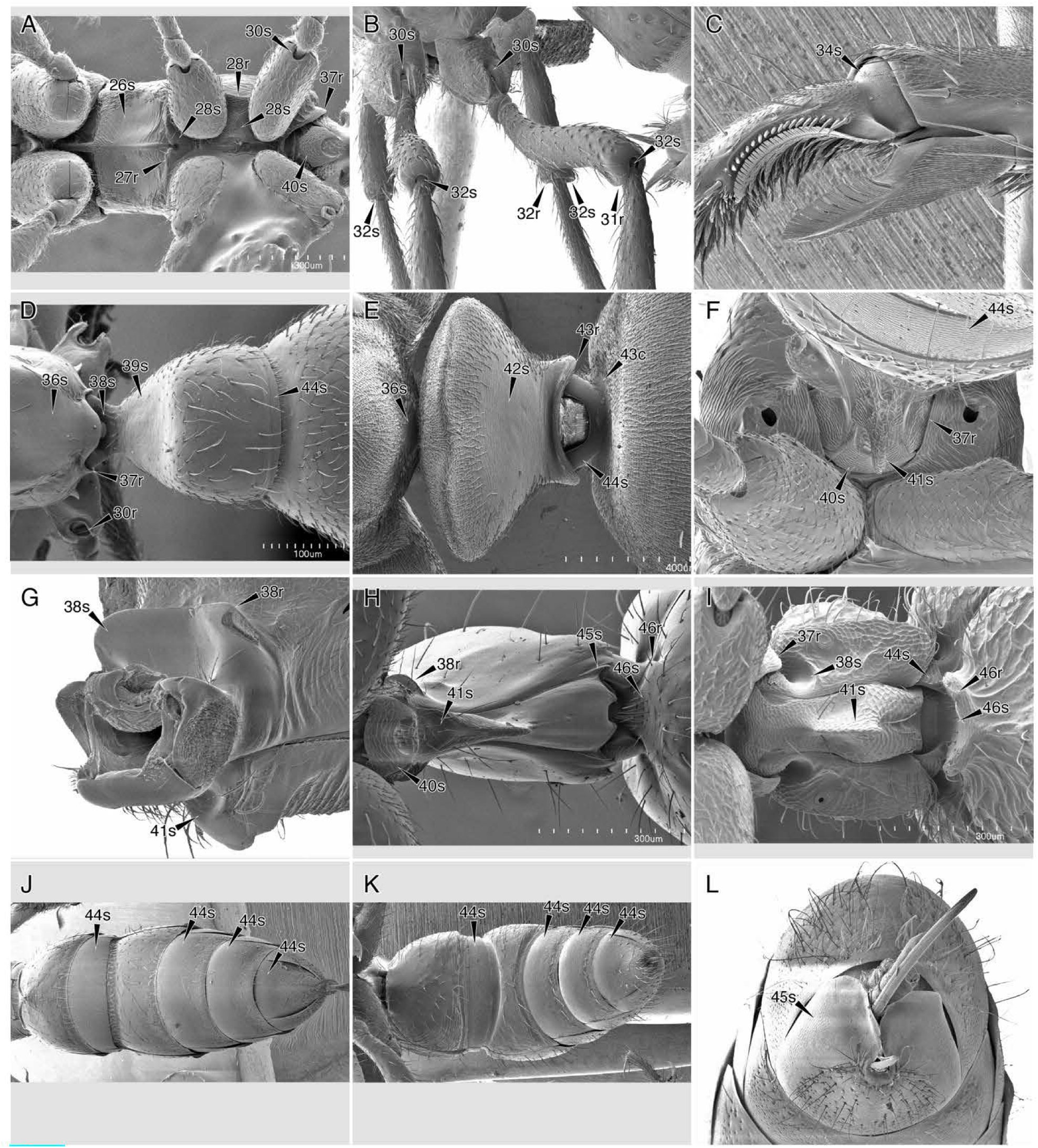
Contact surfaces and rims of the mesosoma and metasoma. Numbers refer to indices in the text; s = contact surface; r = contact rim; c = complementary structures. Taxa: (A) *Fulakora chilensis,* ANTWEB1008496; (B) *Gnamptogenys simulans,* ANTWEB1008520; (C) *Neoponera villosa,* ANTWEB1008571; (D) *Prionopelta concerta,* ANTWEB1008513; (E) *Formica fusca,* ANTWEB1008526; (F, J) *Amblyopone australis,* ANTWEB1008497; (G) *Neoponera apicalis,* ANTWEB1008561; (H) *Centromyrmex brachycola,* ANTWEB1008508; (I) *Proceratium croceum,* ANTWEB1008583; (K) *Ectatomma tuberculatum,* ANTWEB1008524; (L) *Stigmatomma pallipes,* ANTWEB1008501. (All images by Roberto Keller, AntWeb.)

**(I) *Antennal***: (1) the bulbus (*cranioantennal*, Figs. 5A, D); (2) the scape base (margin of the bulbus neck, *antennocranial*, Figs. 5A, F); (3) the scape apex (*pedicelloscapal*, Figs. 5A, D); (4) the pedicellar base (*scapopedicellar*, Figs. 5A, D); (5) the flagellomeral bases (*flagelloflagellomeral*, Figs. 5A, D).

**(II) *Mouth appendages***: (6) the labrum (“proximolateral labral processes”, *stipitolabral*, Figs. 5B, C); (7) the mandible (“mandibular acetabulum” or the anterior/dorsal mandibular articulation, *craniomandibular*, Figs. 5B); (8) the maxillary stipes (“transverse stipital ridge”, *labrostipital*, Figs. 5C); (9) the labium (“distal premental ridge”, *labrolabial*, Figs. 5C).

**(III) *Cranial***: (10) the medialmost region of the hypostoma (*cardinohypostomal*, Figs. 5B); (11) the lateral regions of the hypostoma (*stipitohypostomal*, Figs. 5C); (12) the pleurostoma (“cranial acetabulum” or the posterior/ventral mandibular articulation, *mandibulocranial*, Figs. 5B); (13) the anterior clypeal margin (*labroclypeal*, Figs. 5B); (14) the antennal foramen (“torulus” and its “antennal acetabulum”, *antennotorular*, Figs. 5E– G); (15) the cranial surfaces surrounding the torulus (antennal contact surface, *antennocranial*, Figs. 5D, E, G); (16) the clypeus, anterad the toruli (“clypeal shelf” as in *Protanilla* or *Tetramorium*, *antennoclypeal*, Figs. 5E); (17) the face, mediad the toruli (“frontal carinae”, *medial antennocranial*, Figs. 5D,– G); (18) the face, laterad the toruli (“parafrontal carinae” as in many Dorylinae, or the “malar carinae” as in *Neoponera*, *lateral antennocranial*, Figs. 5D, F, H); (19) the occiput (“occipital carina”, *craniothoracic*, Figs. 5H).

**(IV) *Thoracic***: (20) the anterior pronotal margin (*craniothoracic*, Figs. 5I, J); (21) the pronotal disc (“pronotal flange” as in some Dorylinae, *craniothoracic*, Figs. 5I); (22) the ventrolateral margin of the pronotum (the contact surface is concealed, *propleuropronotal*, Figs. 5I); (23) the ventrolateral margin of the propleuron (*propleuronotal*, Figs. 5I, K); (24) the anterior mesoscutum or mesonotum (*promesonotal*, Figs. 5K); (25) the lateral margin of the mesoscutum dorsad the fore wing foramen (“parascutal carina”, *alarothoracic*, Figs. 5L); (26) the ventral paramedian surfaces and ventrolateral margins of the mesopectus, sometimes continuous with that of the promesonotal articulation (“epicnemium”, *coxothoracic*, Figs. 5I); (27) the ventromedial margin of the mesopectus (ventromedian longitudinal or Y-shaped carina, *coxothoracic*, Figs. 6A); (28) the ventrolateral margins of the meso- and metapecta (*coxothoracic*, Figs. 5I, 5A).

**(V) *Leg***: (29) the coxal bases (“basicoxa” and “basicosta”, *coxothoracic*, Figs. 5I, 6A– D); (30) the distal meso- and metacoxal foramina (*trochanterocoxal*, Figs. 5I, K, 6B); (31) the femoral apices (*tibiofemoral*, Figs. 5I, 6B); (32) the tibial bases (*femorotibial*, Figs. 5I, 6B); (33) the tibial apices (*tarsotibial*, Figs. 6C); (34) the basitarsal base (*tibiotarsal*, Figs. 6C); (35) the tarsomeral apices (*tarsotarsomeral*).

**(VI) *Propodeal***: (36) the posterior propodeal surface (*petiolopropodeal*, Figs. 5I, 6D, E); (37) the propodeal foramen (“foraminal rim”, “propodeal lobes”, *pedunculopropodeal*, Figs. 5I, 6A, D, F, I).

**(VII) *Metasomal***: (38) the anterodorsal petiolar neck and corners (*pedunculopropodeal dorsal*, Figs. 6D, E); (39) the anterior surface of the petiolar tergum (*propodeopetiolar*, Figs. 6D, G–I); (40) the anteroventral petiolar sternum, anterior to the proprioceptor zone (*pedunculopropodeal anterior*, Figs. 6A, F, H); (41) the anteroventral petiolar sternum, anterior and lateral (“subpetiolar process”, *sternocoxal*, Figs. 6F–I); (42) the posterior surface of the petiolar tergum (*postpetiolopetiolar*, Figs. 6E); (43) the posterior petiolar articulation (“posterior petiolar collar”, *helciopetiolar*, Figs. 6E); (44) the presclerites or prescleritic areas of abdominal segments III–VII (*postscleritopresclerital*, Figs. 6D–F, I–K); (45) the dorsolateral surfaces of the abdominal sterna (*tergosternal*, Figs. 6H, L); (46) the abdominal sternum III (“prora”, *petiolosternal*, Figs. 6H, I).

That such rims are nearly ubiquitous as the defining margins of contact surfaces suggests a common developmental mechanism for specifying location and form, perhaps in a sequential manner between the convex and concave surfaces as observed in the development of condylar articulations in *Drosophila* (Tajiri *et al*. 2010). Usually, one or more of these carinae are absent or poorly developed, and loss of several contact surface rims in the omnivorous Dolichoderinae and Formicinae suggest nutritional limits on the developmental capacity for these structures. Other forms of margination may be present, particularly in genitalic structures, as well as the longitudinal dorsolateral margination of the mesosoma and metasoma as observed in Dorylinae and various Camponotini, for example. The non-genitalic examples of the non-contact ridges are due to some undefined process of specified cuticular folding, and their functions are much less certain.

#### Section 2: Phenotypic data

##### Character and state concepts

In the present study, phenotypic characters were not treated as Hennigian transformation series (*e.g.*, Hennig, 1966, Farris *et al*., 1970). Rather, we follow Wagner (Wagner, 2007, 2014; McKenna *et al*., 2021) in conceiving of the physical features of organisms as **characters** that are developmentally specified phenotypic objects, object systems, or regions which have specific **states** that are genetically specified geometric or functional variants of the phenotypic object, object system, or region. Our preference for this conceptualization derives from the combination of the developmental causality of phenotype and the capacity for developmental genetic experimentation on anatomy and physiology (*e.g.*, see Gilbert 2021 and references therein). Previously we had been roughly working from the distinction of “anatomy” and “morphology” as defined by Snodgrass (1935)—namely that the former refers to physical “fact” and the latter to “theory”—without the theoretical consideration of the developmental causes of the evaluated features.

During the initial phases of this work, we had conceived of a Boolean “observation character” approach, where specific features were observed in a specimen or group of specimens, compared to other groups, described, and iteratively scored for all sampled taxa. Now we conceive of the process of phenotypic quality determination at three levels: (1) at the level of developmental character (developmental specification, effectively a linguistic Noun); (2) the level of developmental state (developmental variation, effectively a linguistic Adjective describing a Noun); and (3) the level of observational criteria for evaluating whether the observed specimen or specimens *do* or *do not* display the character and state (effectively a heuristic natural language formulation to guide TRUE/FALSE judgment). A key distinction caused by espousal of the “developmental character” concept is that evolutionary novelty is considered the *de novo* origin of new characters, and innovation as the modification of previous characters for functional purposes (see the cited works of Wagner). Consequently, when we refer to what would be traditional Hennigian characters, we use the “Wagnerian” term and concept “character state transformations”; when specifying the traditional notion of “character” we employ the compound noun “character-state”; and finally, when specifying the traditional notion of “state” we employ the compound noun “character state” or simply the noun “state”.

Regarding the morphological data themselves, we recognize that scoring discrete observations is a lossy information compression process that is also prone to subjective bias. For this reason, we have proactively addressed a number of known issues (see “Scoring procedure” below for specific detail). We have marked in the character state definition list each instance where an arbitrary choice has been made in token (0 or 1) designation due to logical equivalency, such as the states “head broader” or “narrower than long”. Such token designation of arbitrariness has previously been used as reasoning for excluding unequal-rate transition models for morphological data (*e.g.*, Lewis, 2001; Ronquist *et al*., 2011); the modeling consequences of this inconsistency remain unknown. Through our binary approach we have tried to limit the number of composite character states (“composite characters”, *e.g.*, Strong & Lipscomb, 1999; Brazeau, 2011); where such composites are defined, the rationale is explicitly outlined. Because non-additive binary coding has been criticized as possibly resulting in illogical ancestral state reconstruction (Jenner, 2002; Brazeau, 2011), we constructed sets of separate additive character states where necessary. To avoid the problem of “repeated absence” weighting (Brazeau, 2011), we reductively scored as inapplicable certain character states for terminals that lack a given feature (Strong & Lipscomb, 1999). In some cases, such as position and form of wing crossveins, we have scored additive character states reductively, resulting in “additive reductive” scoring, effectively creating hierarchical characters in the sense of Hopkins & St. John (2021). In situations where the scoring of two similar character states was found to be identical (*i.e.*, to have duplicate information content), one of those characters was deleted and a note provided in the other. Finally, given the critique that “absence” is “empirically empty” (Jenner, 2002), we ran preliminary analyses where “not observed” was scored as inapplicable (“-”); these were not computable, and we point out that observing absence of phenotypic expression in a specific specimen or set of specimens is a process of the empirical research program.

##### Homology hypotheses

We differentiated between primary and secondary homology hypotheses, following de Pinna (1991). The former in our sense refers to the initial definition and scoring of a phenotypic feature (the raw data), and the latter the resultant polarities after the process of phylogenetic analysis. We attempted to be as precise as possible with specific primary homology hypotheses in order to facilitate replication and possible developmental experimentation. When putative serial homologs (“paramorphs” versus “homomorphs” *sensu* Wagner) displayed noticeable individuation, these were separated as distinct characters. Our method for testing the primary homology hypothesis was probabilistic ancestral state estimation, which we conducted using a posterior distribution of trees rather than a single “optimized” tree, as we found the latter to result in overinflated support values. We considered the resultant non-falsified primary hypotheses to be our set of secondary homology hypotheses, which we formally present in the extended transformation series (Results Part IV).

We employed five of the criteria of Remane (1956) for positing primary homology hypotheses, with attention to the principle of connection: (1) phenotypic features are probably homologous if they have similar positional relationship relative to other features (“criterion of position”); (2) complex phenotypic features are probably homologous if they have similar qualities, material composition, and function, although of course these may vary considerably (“criterion of special quality”); (3) extreme forms are probably homologous to other forms if there is continuous linkage through an observable transformation series; (4) simple phenotypic features are probably homologous if found with frequency in closely related individuals and/or species; and (5) phenotypic features observed in distantly related clades without intermediates are probably not homologous. Features which were judged to fail the fifth criterion were scored as “taxic”, or taxon-specific, states (taxic concept from Carine & Scotland, 1999). Justification for taxic definitions, noted in the definition list, is based on extensive morphological comparisons in combination with the general congruence between independent transcriptomic and ultraconserved element phylogenomic analyses of Peters *et al*. (2017) and Branstetter *et al*. (2017b).

##### Scoring procedure

Pragmatically, all numbered phenotypic character states were scored based on direct observation as binary Boolean values, *i.e.*, as TRUE or FALSE (1 or 0), given the comparison-derived character and state definitions. When the application of FALSE requires explanation, this is indicated parenthetically. In a number of cases, inapplicability was lumped with FALSE; when inapplicability must be explicit for downweighting, it was indicated using dashes ("-") instead of a Boolean value. Uncertainty is expressed as question mark tokens ("?"); this state is scored for many fossils because of either evolutionary uncertainty (*e.g.*, †Falsiformicidae), or impossibility of observation. In complex cases, we distinguished between “additive”, “reductive”, “additive reductive”, and “reductive taxic” character states; these coding classes is explicitly stated in the character state definitions.

*Additive scoring.* For transformation series of a given structure, the series was categorically divided and broken into separate binary character states. In most cases where such a series is obvious (*e.g.*, linear variation), these are treated as additive; in other cases, such as three-dimensional form variation without obvious intermediates, these were not additively scored. Relative scape length is an example of an additive set of character states. When a scape was observed to be “elongate”, *i.e.*, longer than at least half the head length (char. 155), the observational criteria of “intermediate” (scape length ≥ 4x width, char. 154) and “somewhat short” (length > 2 x width, char. 153) were also met, and thus all were scored as TRUE. In practice, there was no ambiguity in interpreting the results of ancestral state estimation for these additive characters, *i.e.*, there were no instances where a node was reconstructed as having an “elongate” but not a “slightly elongate” scape. Thus, the estimation of the “ancestral grotesque monster” of Jenner (2002) was avoided.

*Reductive scoring.* Reductive scoring was employed in the matrix in cases where the form or position of a structure was dependent on its presence, and where primary homology of presence/absence of the given structure was uncertain. This application of reductive scoring applies to several character state sets, but particularly wing venation, as rampant parallelism in reduction is known to occur (Klopfstein *et al*., 2015). For example, if the third radial sector-medial crossvein (3rs-m) is observed to be absent (scored as FALSE; char. 517), then its shape (char. 518), location relative to the distal wing margin (char. 519), and position relative to 2rs-m (char. 520) are inapplicable and scored as “-”. Our intent with using inapplicable coding for dependent states was to avoid overweighting these conditions. Presence or absence of 3rs-m would be a “controlling primary” character in the hierarchical character nomenclature of Hopkins & St. John (2021).

*Additive-reductive scoring*. To avoid overweighting extreme conditions, certain additive states were reductively scored. Primary examples from our work are the form and degree of constriction of the articulatory surfaces of abdominal segment III (metasomal II). The tergum and sternum of abdominal segment II may be separated into “pre-” and “postsclerites” by the presence of a transverse line of variable impression (char. sets 401–404); when the line is present, the “presternite”, for example, could project below the overlapping tergite (Bolton,

1990a,b), or the “presternal” region may be strongly narrowed relative to the remainder of the segment, a condition known to be homoplastic among Formicidae (*e.g.*, Bolton, 2003; Branstetter, 2017a).

*Taxic scoring*. In some cases, it was clear that a condition was not due to shared descent. An example for this is the optimization of the mesometanotal articulation for a high degree of flexibility in apterous females of Plumariidae, Bethylidae (Pristocerinae), and Thynnoidea, which were scored as independent characters (chars. 172–174). Such conditions were scored as inapplicable (“-”) for any taxon not diagnosable as those or near those families. Note that this particular set of analogous conditions is understudied, for example, so refined characterization will be possible via explicit anatomical analysis.

*Taxic-reductive scoring*. This scoring was employed in the context of very few phylogenetically restricted states which are dependent on the occurrence of apterous adults in a given species. An example of this scoring approach is the relative position of the reduced compound eye of Formicidae (chars. 38, 39). Scoring of this character depends on the presence of female winged-wingless polyphenism and was thus additive. Given that it is highly unlikely based on comparisons of all major extant clades to assume that this polyphenism was ancestral to the entire Aculeata, the location of the compound eye was reductively scored for the Formicidae. Because previous phylogenetic studies have independently recovered wingless female aculeates with metapleural glands as stem Formicidae (Barden & Grimaldi, 2016; Barden *et al*., 2020), we were confident in scoring certain extinct terminals as TRUE for these conditions; for those where we were not confident, we scored the terminals as uncertain, “?”. In some cases, the taxic-reductive condition had a dependent additive state (*e.g.*, char. 39, extreme posterior location of compound eye in female Formicidae).

##### Matrix construction

All morphological data were scored *de novo*, *i.e.*, every cell in the was filled by direct observation rather than from scorings from prior studies, and all characters were newly defined based on the observational experience. Prior to character definition and scoring, specimens were compared for identification purposes and in order to digest character definitions from the taxonomic references used, especially the systematic works of Brothers (1975, 1976), Bohart & Menke (1976), Königsmann (1978), Michener (1981), Rasnitsyn (1988), Gauld & Bolton (1988), Johnson (1988), Kimsey & Bohart (1990), Kimsey (1991), Baroni Urbani *et al*. (1992), Goulet & Huber (1993), Brothers & Carpenter (1993), Gibson (1993), Grimaldi *et al*. (1997), Prentice (1998), Carmean & Kimsey (1998), Ronquist *et al*. (1999, 2012), Vilhelmsen (2001, 2010, 2011), Bolton (2003), Turrisi & Vilhelmsen (2010), Keller (2011), Sharkey *et al*. (2012), Li *et al*. (2015), Boudinot (2015), Zimmerman & Vilhelmsen (2016), and Barden & Grimaldi (2016). Numerous other references are cited and discussed as needed, particularly where prior character descriptions were specifically relied upon. Because of our reliance on fossils for which we do not have µ-CT data, our character set strictly comprises external features of the body although our work on amber †*Gerontoformica* (Boudinot *et al*. 2022, Richter *et al*. submitted) shows that it may be possible to develop a character set akin to and directly interlinked with that of Ronquist *et al*. (2012) and other key studies. We emphasize that the character definitions and matrix were constructed from the ground up based on one of the authors’ observational experience, thus character-by-character tracings through the literature was not possible for all characters. Character definitions are most broadly organized by tagma, anterior to posterior; generally, within tagma, they are reported from anterior to posterior, dorsal to ventral, and proximal to distal.

Matrix construction took place in three phases, which are referred to in some of the character-specific notes: (1) character development, in which raw comparative observations were described as observational characters, and in which previously-described characters were translated for the purposes and into the format of the present study; (2) character scoring, in which at least one representative of each terminal was critically examined and data were recorded for all characters; and (3) quality control, in which every character description was revised for clarity, pertinent notes on state distributions, marginal cases, and other considerations were recorded, every column was checked for typos, and characters were fragmented, aggregated, or deleted based on an evaluation of their informativeness and consistency with the remainder of the matrix (for example, characters with duplicate information were deleted). We note that this process was not perfect; we have, however, executed this process to the best of our ability, and have provided statements of uncertainty as seen fit.

The development and scoring phases were intermingled, as each taxon or set of taxa added to the matrix was or were critically examined for shared phenotypic states, and as discovery of new conditions often required wholesale reexamination of previously scored taxa; during quality control, some character states were scored afresh based on expanded understanding of variation. To attain maximal consistency all character states were developed and scored by one of the authors (B.E.B.). Beyond definitional and observational error, it is expected that there is some degree of bias in this approach, as (1) virtually all of the character states developed were qualitative, (2) evaluation required the experience of one individual (augmented by observations of others), and (3) the scorer’s species-level experience is largely restricted to one family (the Formicidae). The majority of fossil taxa were scored based on graphical representations or descriptions in the literature, thus a number of characters or states could simply not be evaluated for these terminals outside of those captured by prior authorities.

#### Section 3: Data acquisition and processing

##### Morphological data

To evaluate phylogeny and time-calibrated transformation sequences, we developed and scored *de novo* a list of 576 binary morphological character states for 575 species from directly examined specimens and the literature (illustrated in Figs. C1–66). In total, we scored 431 fossil taxa representing a comprehensive sample of Mesozoic Aculeata and outgroups (Rasnitsyn, 1988; Peters *et al*., 2017; Branstetter *et al*., 2017b), including †Bethylonymoidea, Trigonaloidea, Evanioidea, and the more distantly related and possibly ancestral †Ephialtitoidea. Morphological data were scored for 144 extant taxa. Altogether, the matrix was 55.8% complete, with 3.9% of the incompleteness due to inapplicability (“-” scores) and 40.3% due to uncertainty (“?” scores). Most of the uncertainty was attributable to the fossils, which were collectively 43.3% complete; the extant terminals were 93.2% complete. In general, character states were scored for females unless otherwise indicated. We recommend the inclusion of male genitalic and internal anatomical features in future study.

Although state reversal may inflate divergence dating estimates (Klopfstein *et al*., 2019), cases of probable or known reversals were retained for the purposes of ancestral state reconstruction. Autapomorphies were scored for a number of terminals; it is not known if these specific features in this dataset are clocklike (*e.g.*, Matzke & Irmis, 2018). Although crucial to the general evolutionary conception of the Aculeata, the character state "ovipositor modified as sting" was deleted from this matrix as substantial anatomical study is necessary to structurally define the functional sting (see the notes for char. 433). Overall, we found that data completeness by taxon was the most important variable determining analysis quality. See “Phylogenetic analysis” below for our methods and considerations of choosing morphological matrices for further analysis. For character state definitions, literature references, and taxon sampling, see Results Part V below.

##### Genomic data

For combined morphological and molecular analyses, ultraconserved element (UCE) (Faircloth *et al*., 2015) genomic data were available for 115 of 144 of the extant taxa and were harvested from multiple sources (Branstetter *et al*., 2017b; Branstetter *et al*., 2017a; Blaimer *et al*., 2015; Borowiec, 2019; P. S. Ward unpubl.). Since a small minority of these taxa were represented by the Hymenoptera v. 2 probe set (Branstetter *et al*., 2017a), we filtered our UCE loci to those present in the Hym. v. 1 set (Faircloth *et al*., 2015). In cases for which untrimmed UCE sequences were not publicly available, we extracted UCE sequences from the available assemblies using the “assembly match contigs to probes”, “assembly get match counts”, and “get fastas from match counts” scripts from the PHYLUCE (v. 1.6.7) package (Faircloth, 2016). The combined UCE dataset contained sequences from 1391 UCE loci and 115 taxa.

We aligned the sequences with MAFFT v7.407 (Katoh *et al*., 2002; Katoh *et al*., 2013) using the E-INS-i algorithm. The resulting alignments were edge-trimmed using the “align get trimmed alignments from untrimmed” script with the following settings: max_divergence 1.0, min_length 200, proportion 0.8, window 80, and the rest left at default values. After trimming, 345 loci remained. Using the program AMAS (Borowiec, 2016), we added taxa with missing data to each alignment so that all alignments contained all taxa and calculated basic summary statistics. We then selected the 71 alignments that had less than 30% missing data at the site level.

To filter loci whose evolution was likely not stationary, reversible, and homogeneous, we concatenated the alignments and performed matched-pairs tests of symmetry (Jermiin *et al*., 2017; Naser *et al*., 2019) on each partition (corresponding to one UCE locus) as implemented in the program IQ-Tree (Minh *et al*., 2020; Chernomor *et al*., 2016). Using a p-value cutoff of 0.05, 27 partitions/loci were rejected. We then checked the 44-locus alignment by eye in AliView (Larsson, 2014), deleting gap-only columns plus ambiguously aligned sections, and evaluated the phylogenetic information content via topology estimation and AMAS.

##### Alternative matrices

To assess support for divergence dating and ancestral state estimation results from the total-evidence data, we performed a set of alternative analyses with the phylogenetic scope narrowed to the Scolioidea, Formicoidea, and Apoidea with assemblies from additional sources (Bossert *et al*., 2019 and Gillung unpubl.; hereafter “SFA” matrix). For these, we extracted UCE sequences from the assemblies using PHYLUCE as outlined above. In an attempt to increase the number of included loci, we used the Hym. v. 2 probe set. As assemblies were not available for some taxa from Branstetter *et al*., (2017b), we used the untrimmed, unaligned UCE sequences instead. We compared the number of UCEs recovered from alternative assemblies for some species as well as assemblies from closely related species. For each taxon, we kept the data from the assembly that yielded the greater number of UCEs. At this stage, the dataset comprised 87 taxa and 2590 loci.

Sequence alignment and edge-trimming were conducted as above, except that min_length was set to 300. After trimming, 931 loci remained. Given that gaps are treated as missing data (*i.e.*, indels are not explicitly modeled), we removed unique indels using the del.colgapsonly() function from the ape (Paradis & Schliep, 2019) package in R (R Core Team, 2020). Using AMAS, we added taxa with missing data to each alignment. We then concatenated the alignments and performed matched-pairs tests of symmetry on each partition as above, after which 350 UCEs remained. We sorted the remaining alignments by increasing percent of missing data and selected the most complete alignments while keeping the cumulative fraction of missing data less than 25% at the site level. After visual inspection of the chosen alignments, we eliminated those that appeared to have long, ambiguously aligned regions, resulting in a final count of 62 alignments. For UCE source per taxon for all analyses, see the XXX on Zenodo (doi: [XXX]).

#### Section 4: Phylogenetic analysis, part 1

*Note*: This section outlines our methods for topology searches, divergence dating, and ancestral state estimation (ASE) for morphological data. For behavioral data ASE, see Section 5.

##### Morphology-only topology searches

The topology searches using morphological data only were conducted using the maximum likelihood (ML) program IQ-Tree 2.0 (Minh *et al.,* 2020) and the Bayesian program MrBayes 3.2.6 (Ronquist *et al*., 2012) with BEAGLE (Ayres *et al*., 2012) using the CIPRES Science Gateway (Miller *et al*., 2010).

All IQ-Tree Modelfinder (Kalyaanamoorthy *et al*., 2017) tests favored two-state GTR (Tavaré, 1986) with estimated state frequencies and multiple rate categories; because morphological models in MrBayes are limited, we chose Mk with or without ascertainment bias (Lewis, 2001) plus gamma (Yang, 1991) to control for among character (state) rate variation (ACRV).

ML tree searches, without constraints, initialized 1000 trees and ran 1000 ultrafast bootstrap replicates (Hoang *et al*., 2018). Values set for the ML analyses were as follows: `iqtree -ninit 98 -bb 1000 -st BIN -m GTR2+FO+R5 -t BIONJ`.

Soft phylogenetic constraints for Bayesian analysis (Ronquist *et al*., 2012; Fikáček *et al*., 2020; Field *et al*., 2020) were placed on extant taxa to guide placement of fossil terminals based on congruent results from prior genomic studies (Branstetter *et al*., 2017b; Branstetter *et al*., 2017a; Blaimer *et al*., 2015; Sann *et al*., 2018). See the Nexus files provided in the supplemental material for constraint sets. MCMC was performed for 25 million generations, sampling every 5 thousandth, using 3 heated and 1 cold chain with 4 runs each, temp set at 0.01 to increase mixing, and burnin determined by manual examination of trace files using Tracer 1.7.1 (Rambaut *et al*., 2018). Convergence was assessed via average standard deviation of split frequencies (ASDSF), potential scale reduction factors (PSRF), effective sample sizes (ESS), and through among-run distribution overlap as plotted in Tracer.

To evaluate the effect of missing morphological data, we ran 10 analyses in each program under these settings, stratified by matrix completeness: all taxa included, only taxa with 90% phenotypic data completeness, 80%, and so on to 10%, 5%, and 0% (all terminals included). See Fig. 7 for distribution of fossil terminals by completeness and fossil source (*i.e.*, amber versus compression). Using AMAS (Borowiec 2016) to calculate average support across these analyses, we observed that with the stratified addition of more incomplete taxa, mean BPP (Bayesian posterior probability) decreased while the standard deviation increased (Fig. 8). We could not retroactively run replicate analyses to further evaluate convergence due to the loss of funding for CIPRES and the shift of the server to a subscription-based service.

**Fig. 7.**
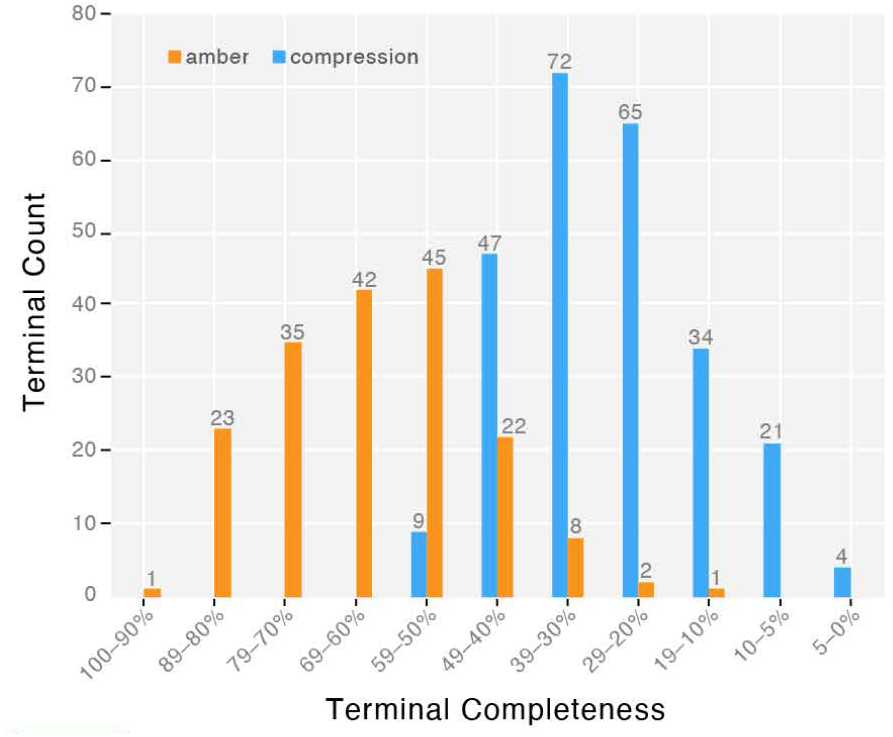
Fossil terminal stratification by data completeness and fossilization matrix. With the addition of those †Bethylonymidae with > 40% completeness, we used a minimum of 50% completeness as the cutoff for our preferred dating matrix (“50CB”).

**Fig. 8.**
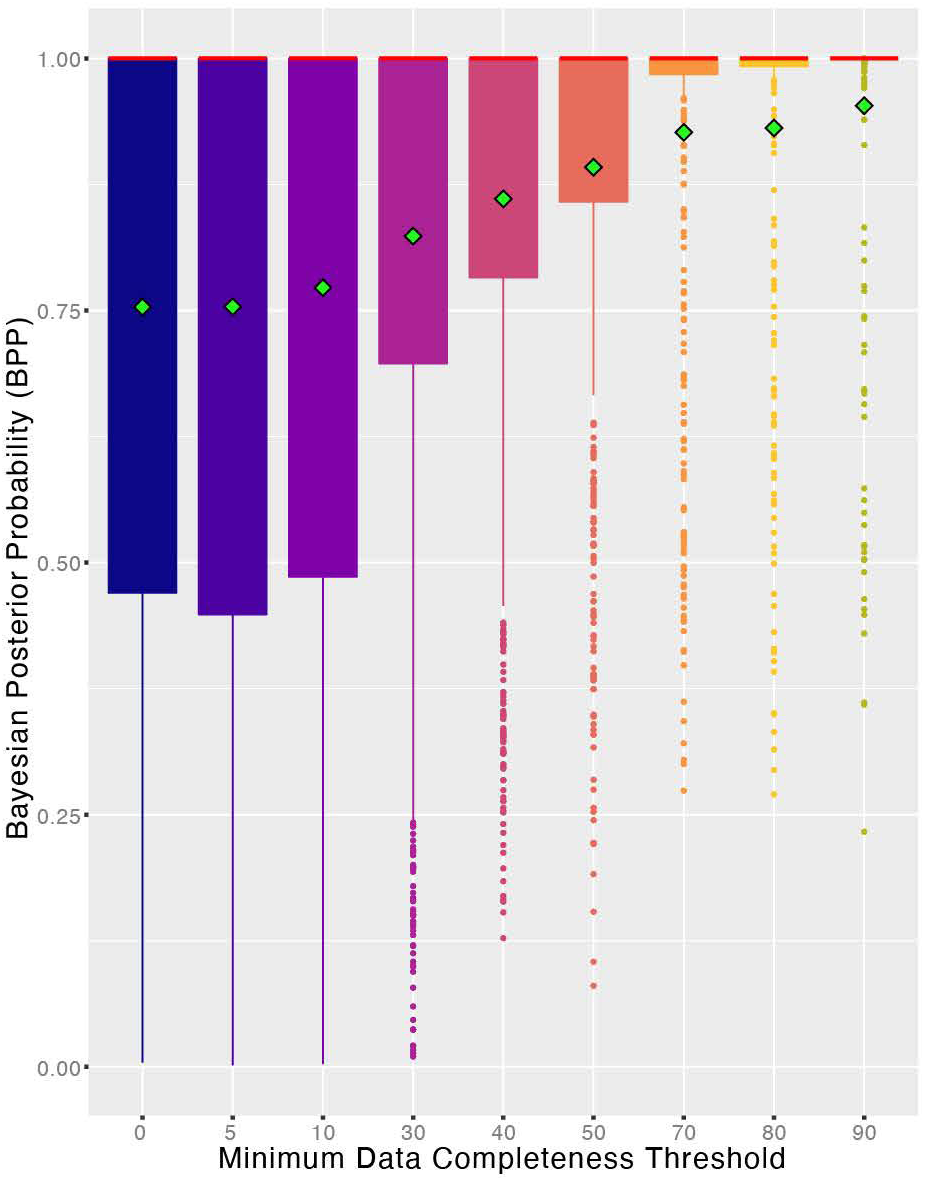
Box-and-whisker plot showing support values by minimum data completeness threshold. Values from morphology-only soft-constrained Bayesian analysis; green diamonds indicate mean BPP.

##### Combined evidence tip-dating

Tip-dating analyses combining morphological and molecular data were conducted in MrBayes on CIPRES. Morphological data were modeled as above; sequence data were included in a single partition with GTR+Γ. All parameters between the two partitions were unlinked. Soft constraints on extant taxa were implemented as above, with the exception that fossil relationships with > 0.90 BPP were also constrained. For this reason, we do not report the support values from the tip-dating analysis, although simulation study indicates that ultrametric clock analyses may have higher topological accuracy (Mongiardino Koch *et al*. 2021); we leave this exploration to future study.

For subsequent analyses, we chose the 50% complete matrix (299 taxa total, ∼48% of terminals extant) and added to it the four most complete †Bethylonymidae fossils (> 40% but < 50% complete) resulting in a 303-taxon matrix with “random” taxon sampling (hereafter “303t-50CB” matrix). We then deleted extant terminals pruned to exclude splits younger than 66 Ma, in accord with the assumptions of diversified sampling (Zhang *et al*., 2016); to increase crown to stem balance, we thinned stem Formicidae to the most disparate exemplars, resulting in a matrix with 199 taxa, of which 67 were extant (∼34%). We refer to this sampling scheme as the “199t-50CB” matrix.

Preliminary analyses revealed that, probably due to rapid transition rates of some character states in conjunction with model mismatch, inclusion of the morphological partition biased estimates toward unrealistically ancient ages, as predicted in simulation studies (Püschel *et al*., 2020; Klopfstein *et al*., 2019). We therefore constrained the root node to 200–165 Ma and the aculeate node to 175–160 Ma, the former representing the range wherein the first apocritan fossils are recovered (*e.g.*, Rasnitsyn, 1988), and both by the consistent placement of the Oxfordian-aged †*Bethylonymus curtipes* and †*Bethylonymellus euryurus* with or within the Aculeata.

##### Fossil dates and clock model

Fossil tips were provided with uniform age priors based on stratigraphic range as recorded in PaleoDB (Behrensmeyer & Turner, 2013) and as recommended by O’Reilly *et al*. (2015) and Barido *et al*. (2020). We implemented the fossilized birth-death model (Heath *et al*., 2014). The clock rate prior was realistically diffuse, set as lognormal(-5,1.2) or an expectation of 0.014 changes per million years with a standard deviation of 0.025. Strict clock assumptions were relaxed using a “white noise” or IGR model (Lepage *et al*., 2007), with the clock variation prior set to exp(37), based on previous values for the Hymenoptera (Ronquist *et al*., 2012). The branch length prior was set to fossilization, with sample stratification set as “diversity” which assumes maximized diversity of sampled extant taxa and random sampling of fossil taxa. The recovery– sampling probability was set to 0.0015 based on the ∼80,000 species described for extant Aculeata, Trigonaloidea, and Evanioidea as accounted from Aguiar *et al*. (2013) and Bolton (2020), with the expectation that this number is an underestimate of the total species presently extant. Speciation, extinction, and fossilization priors were left at their diffuse defaults, while the tree age prior was set to an exponential distribution with a mean of 175 Ma and an offset of 165 Ma, given the broad taxonomic sampling. Final MCMC and burnin were conducted as above, but the results of two replicated analyses were combined in order to attain sufficiently large ESSs.

##### Alternative dating

We used the Scolioidea-Formicoidea-Apoidea (SFA) matrix (see “alternative matrices” above for description of this dataset) to corroborate age estimates for the Formicoidea, Formicapoidina, and Scolioida, molecular-only dating analyses were conducted in RevBayes (Höhna *et al*. 2014; Höhna *et al*. 2016) version 1.1.1 (commit 20534c1f3e7ca1cdcb77ca73e5822efc82cdca53 development branch). The dating model which we employed had the following components: (1) a birth-death process (Feller, 1939; Rannala, 1996) prior on topologies and node ages with exponential priors on birth rate and death rate; (2) monophyly constraints on Scolioidea and Formicoidea + Apoidea respectively; (3) root and node age calibrations based on fossil occurrence times (see below); (4) an uncorrelated log-normal (UCLN) relaxed clock (Drummond *et al*., 2006) with exponential hyperpriors on the mean and standard deviation (in log-space) of the clock rate; (5) locus-specific clock rate multipliers with a flat Dirichlet prior (scaled so the sum of multipliers equals the number of loci); (6) a GTR substitution matrix for each locus; and (7) a discrete-gamma model, as above. We performed two independent runs to ease convergence assessment.

Nine fossil-based lognormal node calibrations plus two alternative prior distributions on the root node age were employed for the alternative dating analyses. In choosing fossils for calibration, we were sensitive to the fact that inclusion of a clock model often resulted in “stemward slip” of fossils, possibly due to missing data and despite compelling morphological evidence. Statements about support for fossil placement from present analyses refer to clockless analyses (see Results Part II below). Below are the parameters and justifications for these ten priors (specific information on construction of priors can be found in the .rev files uploaded to Zenodo):

1. Root (uniform): offset 125.5 Ma (†*Protoscolia normalis* Zhang *et al*., 2002; Yixian formation; sister to crown Scoliidae based on present analyses); hard maximum 174.1 Ma (†*Bethylonymus* and †*Bethylonymellus* in aculeate polytomy).
2. Root (lognormal), offset 125.5 Ma (as above), expected age set to 140.0 Ma (†Angarosphecidae and †*Iwestia provecta* Rasnitsyn & Jarzembowski, 1998; Lulworth formation, England; clusters with stem Pemphredonidae [Apoidea] in Bayesian analysis with soft constraints on extant taxa).
3. Scolioidea crown: offset 125.5 Ma (†*Protoscolia normalis*, as above).
4. Anthophila stem: offset 97.0 Ma (†*Melittosphex burmensis* Poinar & Danforth, 2006; maximally supported as sister to or nested within Anthophila in present analyses).
5. Melittidae crown: offset 52.0 Ma (†*Palaeomacropis eocenicus* Michez *et al*., 2007, Le Quesnoy amber; calibration after Branstetter *et al*., 2017b).
6. Halictidae crown: offset 36.0 Ma (†*Electrolictus antiquus* Engel, 2001, Baltic ambers; calibration after Branstetter *et al*., 2017b).
7. *Apis* (Apidae) stem: offset 68.0 Ma (†*Cretotrigona prisca* Engel, 2000; New Jersey amber; maximally supported as Apidae in present morphology-only analyses).
8. Ponerinae crown: offset 97.0 Ma (Ponerinae indet.; Burmese amber; maximally supported as Ponerini in present analyses).
9. Formicinae crown: offset 97.0 Ma ([1] Formicinae indet.; Burmese amber; maximally supported as Formicinae in present analyses; [2] †*Kyromyrma neffi* Grimaldi *et al*., 2000; Raritan amber; maximally supported as crown Formicidae in present analyses).
10. *Oecophylla* (Formicinae) stem: 44.0 Ma (†*Oecophylla eckfeldiana* Dlussky *et al*., 2008, Eifel formation; shared autapomorphies with *Oecophylla*; sister to extant species in combined-evidence tip-dating analysis of Boudinot *et al*. 2021).
11. Crematogastrini (Myrmicinae) crown: 36.0 Ma (†*Pristomyrmex rasnitsyni* Dlussky & Radchenko, 2011, Baltic ambers; shared autapomorphies with *Pristomyrmex*).

##### Ancestral state estimation (ASE)

Using the program RevBayes and following the insights of Pagel *et al*. (2004), we performed Bayesian ancestral state estimation (ASE) on dated topologies. For ASE, we used the un-thinned (303-taxon) 50CB matrix on all 576 characters (**576C**) and the extant-only Scolioides matrix (SFA) for a select character subset (see “Alternative ASE” below). We also analyzed behavioral characters, namely diet (**D3**) and nesting (**N2**) on the 303t-50CB and SFA matrices, and a complex “supercharacter” with twelve superstates modeling rate transition dependencies among the substates of eusociality, diet, and nesting (**SC**) on the 303t-50CB matrix (see “Section 5” below for explanation). The analyses relying on the 303t-50CB matrix relied on the same posterior distribution of time trees (or the same MCC tree thereof).

The tip-dating model for the base 303t-50CB analysis had the following components: (1) a distribution of tree topologies and node ages taken from the MrBayes tip-dating posterior; (2) an uncorrelated log-normal (UCLN) relaxed clock (Drummond *et al*. 2006) with exponential hyperpriors on the mean and standard deviation (in log-space) of the clock rate; (3) an Mk substitution matrix (Lewis 2001); and (4) a discrete gamma model (Yang 1991) for accommodating ACRV with 4 rate categories, using quadrature (Felsenstein 2001), with an exponential prior on the variance of the gamma distribution. For the 576C trials, all character states were analyzed as a single partition, thus linked by one UCLN clock and Mk+G transformation matrix, and two independent, replicate MCMC runs were performed while sampling ancestral character states

Note that we did not explore parsimony optimization as our preliminary attempts to reconstruct the topology of the dataset using the Maximum Parsimony (MP) criterion using TNT (Goloboff *et al*. 2008) was unsuccessful due to the scale of the present dataset. Further, we preferred to let the uncertainties of character state identity and branch length inform our analyses as previously recommended (*e.g.*, Duchêne & Lanfear 2015, Wright *et al*. 2015).

Support values for all ancestral state estimates in this study were interpreted as follows: 0.50–0.74 Bayesian posterior probability (BPP) for an estimated state at a given node is “no” to “low” support, or “equivocal”; 0.75–0.89 BPP is “weakly” to “well” supported; and 0.90–1.00 BPP is “strongly” supported. See Section 4 of Part I for an explanation of how we came to use this system.

##### Alternative ASE

To evaluate the sensitivity of our ancestral state estimations, we chose 43 morphological characters which transform along the formicoid stem to model in RevBayes using the posterior distribution of SFA timetrees generated in the above analysis. The posterior distribution of dated trees used for this analysis were derived from the “Alternative dating” analyses (above). In contrast to the 50CB analysis, we performed Bayesian ancestral state estimation separately for each morphological character of interest using the posterior distribution of timetrees generated in the above analysis. We ran analyses under three different models while logging ancestral character states: (1) UCLN clock and F81 substitution matrix, (2) strict clock and F81, and (3) strict clock and Mk. In cases where estimates changed substantially with the model, we tested relative model fit by estimating model marginal likelihoods with stepping stone sampling (Xie et al. 2011).

#### Section 5: Phylogenetic analysis, part 2

*Note*: This section outlines our methods for behavioral character ASE.

##### Data and character state coding

Because of our discovery of †*Camelosphecia* (*e.g.*, Boudinot *et al*. 2020) and recovery of †@@@idae **fam. nov.** as the most distant relative of Formicidae (present study), we were compelled to ask if these fossils and this topology could inform the question of eusocial origins in Hymenoptera. Our aim was to reconstruct the evolution of three behavioral characters: diet, nesting, and eusociality, which we treated as discrete. We ran independent analyses on diet and nesting (D3, N2), and then conducted a set of analyses on all three characters combined using alternative transition models (S##).

For the independent analyses of **diet** (**D3**), we recognized (0) parasitoidy, (1) predation, and omnivory, while for **nesting** (**N2**), we recognized (0) non-nesting and (1) nesting. We consider nesting behaviors to include any host relocation behavior enacted by the wasp as this represents a potential transition point from host to habitat modification or manipulation for the optimization of larval development. For the combined (“supercharacter”, **S##**) analyses, we modified **diet** (subcharacter 1) to (0) parasitoidy, (1) provisioning, and (2) kleptoparasitism, we used the two-state formulation of **nesting** (subcharacter 2), and binned **eusociality** (subcharacter 3) into two states, (0) non-eusocial or (1) eusocial. These three subcharacters were combined together, allowing us to create alternative transition models among substate transitions, for which we provide a detailed explanation in Section 4 of Results Part I and especially Results Part V. Regarding “eusociality”, we employ the “obligate eusociality” concept of Crespi & Yanega (1995) as discussed by Boomsma & Gawne (2018). See Eggleton & Belshaw (1992) and Melo *et al*. (2011) for foundational considerations of diet and nesting.

Since we expect diet, nesting, and eusociality to evolve in a non-independent manner, we combined these into a single “supercharacter” with 12 “superstates” (**12S**). We excluded the superstates corresponding to “provisioning, non-nesting, non-eusocial” and “provisioning, non-nesting, eusocial” as they are internally inconsistent (*i.e.*, an animal cannot provision without a nest), leaving a base supercharacter with 10 superstates (**10S**). We also conducted an extreme analysis with only 5 superstates (**5S**), which excluded the two eusocial parasitoid superstates and three eusocial kleptoparasite states as these conditions were not observed in our dataset (*e.g.*, we did not sample socially parasitic Formicidae). The D3, N2, 5S, and 10S data were subjected to an array of modeling tests, which we summarize in Tables 1 and 2. The 5S analysis implemented two rate matrices, with reversible-jump MCMC (rjMCMC) sampling among the two in order to allow for a “skips” or “jumps” to eusociality. Because the 5S and 10S analysis variants represent alternative pathways for the evolution of eusociality, we list them in a supercharacter hypothesis table (Table 3). Note that we use the term “transition” for “substitution” in reference to our morphological rate (Q) matrices.

**Table 1.**
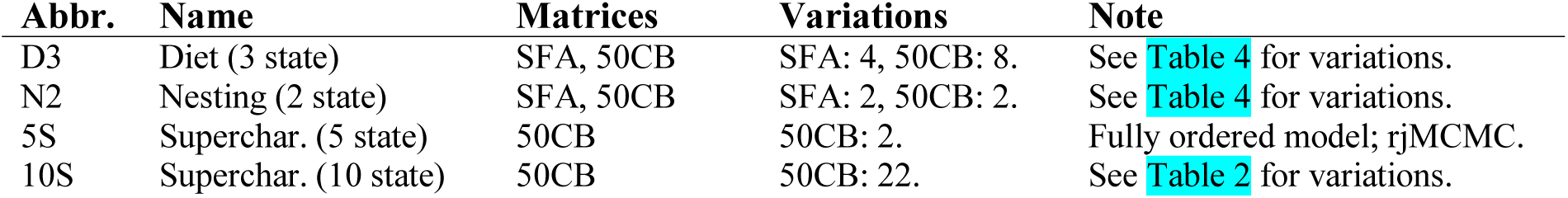
Coarse overview of the 40 behavioral reconstructions conducted in the present study. . SFA = Scolioides extant-only matrix (86-taxon), 50CB = 303-taxon 50CB matrix (*i.e.*, with random rather than diversified sampling), rjMCMC = model averaging via reversible jump MCMC. The two alternative analyses for **5S** were estimation across a single tree or a posterior distribution of trees; both of these analyses were effectively nonstationary as the states at the root of the tree were allowed to vary, *i.e.*, were independent.

**Table 2.**
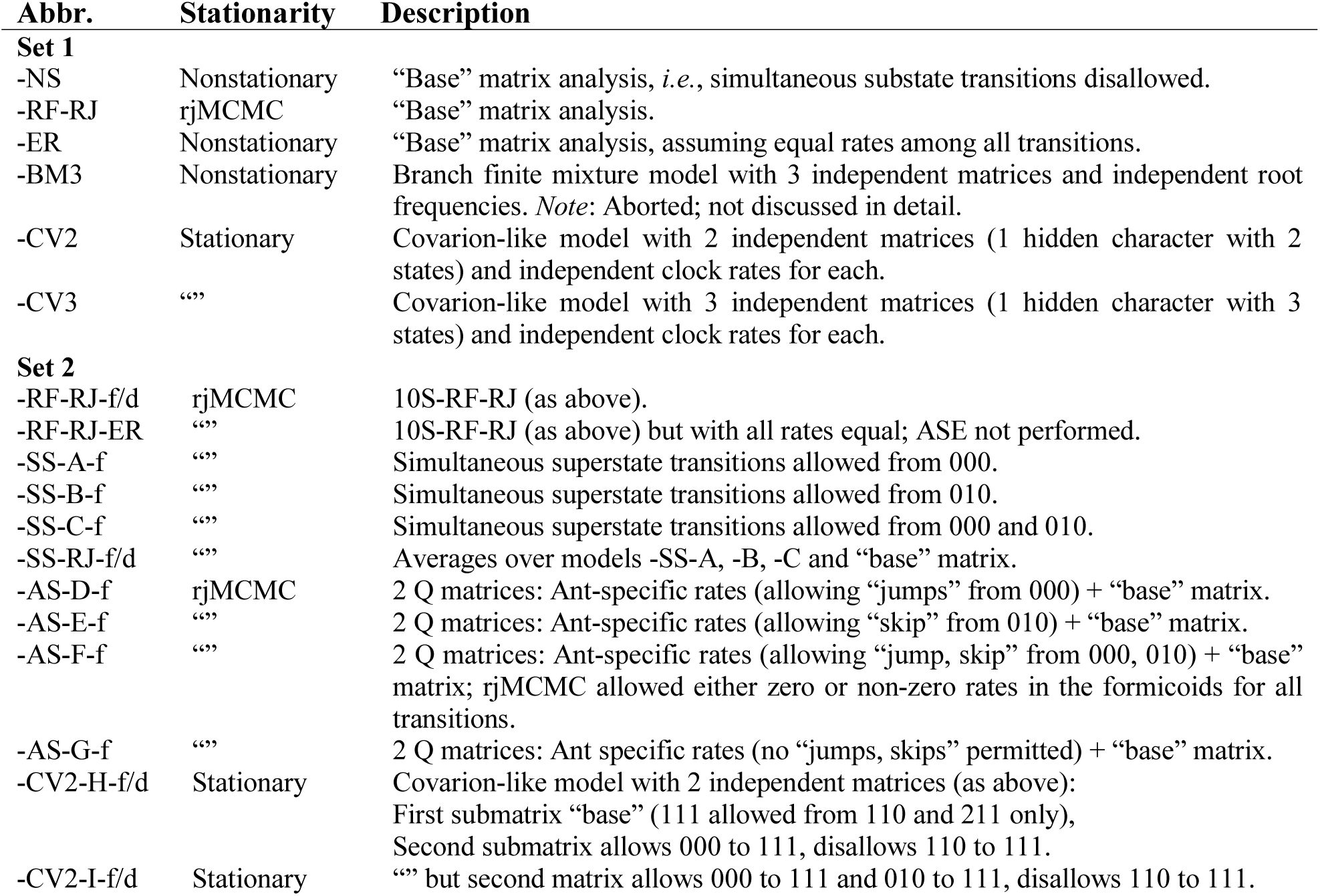
Overview of all 22 10-superstate supercharacter analyses (10S); these were executed in two batches (“Sets 1 and 2”) . Set 1 disallowed simultaneous changes of multiple superstates, and estimates were conducted using a fixed tree; these experiments tested the effects of stationarity or nonstationarity on our estimations (-NS, -RF-RJ, -BM3) and whether the data are better modeled when “jumps” are allowed (-CV2, 3). Set 2 extended these models and allowed or disallowed simultaneous substate transitions. *Additional abbreviations*: -f = estimation across a fixed tree, -d = estimation across a posterior distribution of trees, rjMCMC = reversible-jump MCMC implemented to average over dependent (stationary) and independent (nonstationary) root frequency models. *Superstate combinations*: 000 = parasitoid non-nesting non-eusocial, 010 = parasitoid nesting non-eusocial, 110 = provisioning nesting non-eusocial, 111 = provisioning nesting eusocial, 211 = socially parasitic eusocial species.

| Abbr. | Stationarity | Description |
| --- | --- | --- |
| **Set 1** |  |  |
| -NS | Nonstationary | “Base” matrix analysis, *i.e.*, simultaneous substate transitions disallowed. |
| -RF-RJ | rjMCMC | “Base” matrix analysis. |
| -ER | Nonstationary | “Base” matrix analysis, assuming equal rates among all transitions. |
| -BM3 | Nonstationary | Branch finite mixture model with 3 independent matrices and independent root frequencies. *Note:* Aborted; not discussed in detail. |
| -CV2 | Stationary | Covarian-like model with 2 independent matrices (1 hidden character with 2 states) and independent clock rates for each. |
| -CV3 | “” | Covarian-like model with 3 independent matrices (1 hidden character with 3 states) and independent clock rates for each. |
| **Set 2** |  |  |
| -RF-RJ-f/d | rjMCMC | 10S-RF-RJ (as above). |
| -RF-RJ-ER | “” | 10S-RF-RJ (as above) but with all rates equal; ASE not performed. |
| -SS-A-f | “” | Simultaneous superstate transitions allowed from 000. |
| -SS-B-f | “” | Simultaneous superstate transitions allowed from 010. |
| -SS-C-f | “” | Simultaneous superstate transitions allowed from 000 and 010. |
| -SS-RJ-f/d | “” | Averages over models -SS-A, -B, -C and “base” matrix. |
| -AS-D-f | rjMCMC | 2 Q matrices: Ant-specific rates (allowing “jumps” from 000) + “base” matrix. |
| -AS-E-f | “” | 2 Q matrices: Ant-specific rates (allowing “skip” from 010) + “base” matrix. |
| -AS-F-f | “” | 2 Q matrices: Ant-specific rates (allowing “jump, skip” from 000, 010) + “base” matrix; rjMCMC allowed either zero or non-zero rates in the formicoids for all transitions. |
| -AS-G-f | “” | 2 Q matrices: Ant specific rates (no “jumps, skips” permitted) + “base” matrix. |
| -CV2-H-f/d | Stationary | Covarian-like model with 2 independent matrices (as above):<br>First submatrix “base” (111 allowed from 110 and 211 only),<br>Second submatrix allows 000 to 111, disallows 110 to 111. |
| -CV2-I-f/d | Stationary | “” but second matrix allows 000 to 111 and 010 to 111, disallows 110 to 111. |

**Table 3.** The alternative hypotheses for the evolutionary pathway to eusociality in Hymenoptera as represented by all supercharacter analysis variations (5S and 10S). Analysis labels for 10S are the same as those presented in Table 2. Pathway labels are as follows: “Strict stepwise” = the null hypothesis that parasitoid Hymenoptera derive nesting, provisioning, and eusociality in a strictly stepwise manner (*i.e.*, without “jumps” or simultaneous substate changes); “Provisioning skip” = the alternative hypothesis that eusociality (111) and provisioning (110) can be simultaneously derived from (010); “Total jump” = the alternative hypothesis that eusociality (111) can be derived from non-nesting parasitoidy (000) or from various intermediates depending on the specific model implemented.

| Pathway | Matrix |
| --- | --- |
| Strict stepwise | 10: **5S**, **10S**-NS, -RF-RJ, -ER, -BM3, -CV2, -CV3, -RF-RJ-d/f, -AS-G |
| Provisioning skip | 4: **10S**-SS-B, -AS-E, -CV2-I-f/d |
| Total jump | 9: **10S**, -SS-A, -SS-C, -SS-RJ-f/d, AS-D, AS-F, -CV2-H-f/d |

Our scoring for extant taxa is explained in Tables 6 and 7 of Part V. Scoring of fossils was as follows: all non-ants were categorically scored as uncertain (“?”); †@@@idae **fam. nov.** were scored as uncertain; the formicid groups †Armaniinae, †*Baikuris*, †*Dlusskyidris*, †*Sphecomyrma*, and †*Brownimecia* were scored as uncertain; “eusociality” was scored for †Zigrasimeciinae, †Haidomyrmecinae, and †*Gerontoformica* as both winged and wingless females are known and as a wingless †*Ceratomyrmex* has been preserved holding prey (Perrichot *et al*. 2020); “eusociality” was also scored for †*Gerontoformica* given the indication of this condition from wingless females that have been preserved in combat (Barden & Grimaldi 2016) and, more recently, moving brood (Boudinot *et al*. 2022b). For the diet-only analysis, we scored the †Zigrasimeciinae and †Haidomyrmecinae as “predatory” in the sense that they are reasonably inferred to be provisioning due to the evidence that they are obligately eusocial and as there is no obvious manner to determine whether they also relied on honeydew or plant secretions as a carbon source; †*Gerontoformica* and other stem lineages were scored as uncertain. The reproductive condition of wingless female stem ants remains unknown but can be evaluated using µ-CT technology (Boudinot *et al*. 2022b); this would be highly desirable to inform coding in future analyses of this problem.

**Table 4.** Marginal log likelihoods for ancestral state estimation analyses of diet and nesting. Abbreviations: Diet-fu = diet, stem Formicidae scored as unknown; Diet-f = diet, stem Formicidae scored as predators; SFA = 86-taxon Scolioida matrix; 50CB = 303-taxon 50% complete + †Bethylonymidae matrix.

| Data | Variable | Model | LnL |
| --- | --- | --- | --- |
| SFA | Diet | Ordered, 0 → 1 reversible | -37.50 |
| “” | “” | **Ordered, 0 → 1 irreversible** | **-36.65** |
| “” | “” | Unordered, F81 | -39.17 |
| “” | “” | Unordered, JC | -39.92 |
| “” | Nesting | F81 (variable rates) | -7.84 |
| “” | “” | JC (equal rates) | -7.78 |
| 50CB | Diet-fu | Ordered, 0 → 1 reversible | -45.32 |
| “” | “” | **Ordered, 0 → 1 irreversible** | **-42.22** |
| “” | “” | Unordered, F81 | -47.14 |
| “” | “” | Unordered, JC | -48.96 |
| “” | Diet-f | Ordered, 0 → 1 reversible | -50.17 |
| “” | “” | **Ordered, 0 → 1 irreversible** | **-47.24** |
| “” | “” | Unordered, F81 | -51.86 |
| “” | “” | Unordered, JC | -53.00 |
| “” | Nesting | **F81 (variable rates)** | **-17.76** |
| “” | “” | JC (equal rates) | -19.30 |

**Table 5.** Marginal log likelihoods for the 10S (set 2) analyses, sorted by lowest to highest fit, and support for key nodes. Likelihood estimation method abbreviations: SS = stepping stone, PS = path sampling; n.c. = not calculated. Model abbreviations follow Table 2.

| Model | Hypothesis | SS | PS |
| --- | --- | --- | --- |
| -AS-G | Strict stepwise | -68.77 | -68.79 |
| -CV2-H | Potential for jump (from 000) | -68.57 | -68.60 |
| -CV2-I | Potential for jump (from 000) and skip (010) | -54.22 | -55.33 |
| -RF-RJ | Strict stepwise. | -68.09 | -68.11 |
| -SS-A | Total jump (from 000). | -65.03 | -65.06 |
| -SS-C | Jump (from 000) or skip (from 010) | -64.64 | -64.66 |
| -SS-B | Skip from 010 | -64.49 | -64.55 |
| -SS-RJ | Averaging over -A, -B, -C | (n.c.) | (n.c.) |
| -AS-E | Ant-specific skip (from 010) | -64.29 | -64.31 |
| -AS-F | Ant-specific jump (from 000) or skip (010) | -63.43 | -63.44 |
| -AS-D | Ant-specific jump (from 000) | -63.14 | -63.15 |

**Table 6.** Species-level diversity of obligately eusocial insects. The count of non-eusocial species is only intended to highlight the absence of intermediates in crown Formicidae. Note that social parasites and inquilines are not counted among Formicidae as non-eusocial species here as both conditions are clearly derived within the crown clade. Sources: Formicidae: Bolton (2022), Dictyoptera: Beccaloni & Eggleton (2013), Apidae: Ascher (2022), Vespidae Piekarski *et al*. (2018).

| Clade | Non-eusoc spp. | Eusoc. spp. | Proportion Eusocial |
| --- | --- | --- | --- |
| Crown Formicidae | 0 | > 14,000 | 100% |
| Dictyoptera | ~4,600 | ~3,000 | 65% |
| Apidae | ~6,000 | ~530 | 9% |
| Vespidae (Vespinae) | ~5,300 | ~70 | 1% |

##### Analysis overview

Regardless of model choice, all behavioral ASE analyses were conducted in a time-calibrated manner in RevBayes 1.1.1, using either a single summary chronogram or a posterior distribution of chronograms. These chronograms were obtained from analysis of the 303-taxon 50CB (“random sampling”) and 86-taxon SFA (Scolioides) matrices as described in the “Ancestral state estimation ASE” and “Alternative ASE” sections above. At least two independent chains were run for all analyses. For each 10S (set 2) analysis, we included one run under the prior. Diagnosis for these analyses were conducted as described above, using Tracer (Rambaut *et al*. 2018) to visually assess convergence and the bonsai (May & Moore 2017) R (Core R Team, 2020) package to calculate convergence statistics. In both 5S and a subset of the 10S analyses we averaged over alternative models using rjMCMC, which we describe below. Where applicable, we use the Kass & Raftery (1995) interpretation of untransformed Bayes factor values (e.g., Bergsten et al. 2013): 1–3 = “not worth more than a bare mention”, 3–20 = “positive”, 20–150 = “strong”, > 150 = “very strong”.

*Output visualization*. We used the following R packages for visualizing the results from these analyses: RevGadgets (Tribble *et al*. 2021), phytools (Revell 2012), plotrix (Lemon 2006), ape (Paradis & Schliep 2019), ggplot2 (Wickham 2016), gridExtra (Auguie & Antonov 2017), RColorBrewer (Neuwirth 2014).

##### Models

To further explore the support for alternative scenarios of behavioral character evolution, we employed six sets of models to analyze our 10S behavioral data. These models effectively ask: Is there a single rate of transition to eusociality from non-nesting parasitoidy? In other words, do the data indicate a lock-step path of first gaining nesting then provisioning before a transition to eusociality can be made, or are some intermediates temporally unstable and therefore can be skipped, or can all logical intermediates be jumped over to derive eusociality (and if so, does this only apply to Formicoidea)? Below we outline the alternative models which we also summarize in Table 2. A seventh analytical approach was attempted but abandoned; we include this for the sake of clarity.

*1. The “base” matrix*. We used a "base" transition matrix with 10 superstates (**10S-NS**) which allows stepwise transition to nesting-provisioning-eusociality from non-nesting-parasitoid-non-eusociality and disallows any simultaneous transition of more than one subcharacter (*i.e.*, the corresponding rates are fixed to zero). This matrix was subsequently modified and extended to test different evolutionary hypotheses. In all cases, we used a strict clock with a lognormal prior on the rate. We used an exponential (*lambda* = 10) hyperprior on the mean rate and an exponential (*lambda* = 0.587405) hyperprior on sigma (the standard deviation of the logarithm of the rate). Depending on the analysis, we either sampled from the posterior distribution of time trees inferred from the 303t-50CB tip-dating analysis or conditioned on the Maximum Clade Credibility (MCC) chronogram therefrom (see “Ancestral state estimation” above). For most models, we ran power posterior analyses and estimated model marginal likelihoods using stepping stone sampling (Xie *et al*. 2011, Fan *et al*. 2011) and path sampling (Lartillot & Philippe, 2006).
*2. Base model stationary/non-stationary averaging*. To average over a stationary model and a simple non-stationary model in a manner analogous to the approach of Klopfstein *et al*. (2015), we used rjMCMC (Green 1995) on the base matrix described above (**10S**-**RF-RJ**, -**RF-RJ-d/f**, **-RF-RJ-ER**) as well as for modifications thereof (see below). Root frequencies either matched the stationary frequencies of the substitution matrix or were an independent stochastic variable with a flat Dirichlet prior. We specified equal prior weights for the two models.
*3. Equal rates*. We used rjMCMC to test whether the base model where each rate is drawn from an exponential (lambda = 1) prior fit the data better than a simpler model were all rates have equal, fixed values (**10S-RF-RJ-ER**).
*4. Simultaneous substate transitions*. We tested models that allowed provisioning-nesting-eusociality to evolve directly from a non-nesting parasitoid state (**10S-SS-A**), a nesting parasitoid state (**10S-SS-B**), and both the previous states (**10S-SS-C**). All models still allowed all the pathways to and from provisioning-nesting-eusociality present in the base model. Additionally, we ran analyses (on a fixed tree and on the posterior distribution of trees) using rjMCMC to average over models A, B, C, and the base model that does not allow simultaneous substitutions (**10S-SSC-RJ-f/d**).
*5. Formicoidea-specific matrices*. We also tested models where Formicoidea were assigned a different transition matrix, while all other branches used the matrix from the base model. We allowed provisioning-nesting-eusociality (111) in the Formicoidea to evolve directly from a non-nesting parasitoid superstate (000) for model **10S-AS-D** and from a nesting parasitoid (010) state for model **10S-AS-E**. In both cases, the evolution of eusociality from a provisioning-nesting-non-eusocial (110) superstate was disallowed for formicoids. In a third analysis (model **10S-AS-F**), the rates associated with the following transition were either fixed to zero or drawn from an exponential (*lambda* = 1) prior using rjMCMC: parasitoid-non-nesting-non-eusocial (000) to provisioning-nesting-eusocial (111), parasitoid-nesting-non-eusocial (010) to provisioning-nesting-eusocial (111), and provisioning-nesting-non-eusocial (110) to provisioning-nesting-eusocial (111). Note that the latter rate was independent from the rate of the corresponding transition in the non-formicoid matrix. Finally, model **10S-AS-G** disallowed simultaneous transition of more than one sub-character on all branches but had formicoid-specific (independent) rates associated with the following transitions: parasitoid-non-nesting-non-eusocial (000) to parasitoid-nesting-non-eusocial (010), parasitoid-nesting-non-eusocial (010) to provisioning-nesting-non-eusocial (110), and provisioning-nesting-non-eusocial (110) to provisioning-nesting-eusocial (111).
*6. Covarion-like models*. As an alternative to the invariable assignment of a distinct rate matrix to the Formicoidea as for the 10S-AS series (“5. Formicoidea-specific matrices” above), we implemented a covarion-like model to allow for flexible matrix usage by any given branch. In other words, this approach used models that were similar to the covarion model (Tuffley & Steel, 1998) to allow the substitution process to change over the tree without rigidly assigning matrices to specific branches as above. In the first analysis set of the 10S matrix we employed two (**10S-CV2**) and three (**10S-CV3**) hidden states, respectively, while we only implemented two hidden states in the second analysis set. Switch rates between the two "submatrices" were independent stochastic variables with an exponential (*lambda* = 1) prior. Clock rates for the two submatrices were fixed to being equal. For model **10S-CV2-H**, the first submatrix (corresponding to hidden state 0) was equivalent to the base model above, while the second submatrix disallowed the provisioning nesting non-eusocial (110) to provisioning nesting eusocial (111) transition while allowing the parasitoid non-nesting non-eusocial (000) to provisioning nesting eusocial substitution (111). Model **10S-CV2-I** was similar to -CV2-H but the second matrix allowed both the parasitoid non-nesting non-eusocial (000) to provisioning nesting eusocial (111) and the parasitoid nesting non-eusocial (010) to provisioning nesting eusocial (111) transitions while also disallowing the provisioning nesting non-eusocial (110) to provisioning nesting eusocial (111) transition. Note that first analysis set differs from the second in that all rates of all submatrices are independent. As the first set is highly overparameterized (70 rates for 10S-CV2, 108 for -CV3) and the models are non-identifiable, we rely on the results of the second analysis set. Additionally, the models of the first set were insufficiently constrained for hypothesis testing.
*7. Exploratory finite branch mixture model*. This model implemented three transition matrices in analogy to a gamma model for among character rate variation (**10S-BM3**). During analysis, likelihood was calculated for a given branch across each of the three matrices. In other words, each branch could use one of the matrices as its transition model. We aborted this approach as it was prohibitively computationally expensive, it mixed poorly in our designated implementation, and as there were potential non-identifiability issues among the three matrices as they were symmetrical, in contrast to the second set of covarion-like matrices.

#### Section 6: Character and state definitions

The definitions of all characters and states are reported and figured in Results Part V due to the length of this section. In this part of the manuscript, we provide 66 figures illustrating the majority of the 576 morphological characters; we provide the permissions and sources where necessary for these images in Table S1.

#### Section 7: Mesozoic Hymenoptera fossil occurrences

Species occurrences of Mesozoic Hymenoptera by genus, family, deposit, and formation were downloaded in January 2020 and on 31 December 2021 from PaleoDB (2021). Further data were added by hand, including higher classification, geological ages, taphonomy, depositional environment, and lifestyle (Table S2). With this information, several patterns were graphed and are presented in Section 2 of Results Part I.

## RESULTS & DISCUSSION

### Outline

We have divided our results into five parts. In Part I, we provide a review of the fossil record of the Hymenoptera from the perspective of our phylogenetic results (Section 1), an overview of key topology and dating results (Section 2), a discussion of our morphological and behavioral ancestral state estimation analyses (Sections 3, 4), and a synthetic hypothesis for the evolution of the ants and Aculeata (Section 5). We conclude Part I with a brief summary discussion in which we highlight directions for future study (Section 6). We detail our topological results in Part II and provide a taxonomic synopsis of the Aculeata and kin in the light of previous work and our study in Part III. This synopsis represents a revision of the total clade Aculeata based on our character polarity estimates which are presented as an extended transformation series with diagnoses for most nodes in Part IV. Our character definitions and illustrations are presented in Part V. Logically, Part I follows Parts II–V, but we provide this first as it contains our focal results and conclusions. Note that the taxonomic nomenclature which we use is derived from our systematic revision outlined in Parts III and IV; key clade names are indicated in Figs. 14–17.

## Part I: Fossils, Phylogeny, and Evolution

### Section 1: The Fossil Record of Hymenoptera

As of January 1^st^, 2022, there were ∼ 1515 valid fossil species of Hymenoptera described from 93 Mesozoic deposits, representing 90 families and 505 genera (Table S2; Fig. 9). A single putative stem species, †*Avioxyela gallica* Nel *et al*., 2013b (†Avioxyelidae), has been described from the Moscovian Age of the Carboniferous (314–307 Mya), being associated with the Hymenoptera via the enlarged pterostigmal cell and the compound Rs+M abscissa. The first crown Hymenoptera are Xyelidae recorded from the Madygen and other formations dated to the Ladinian through Norian Ages of the Mid and Late Triassic (242–209 Mya), respectively (Rasnitsyn 1964, 1969; Schlüter 2000; Kopylov 2014a; Oyama & Maeda 2020, Lara *et al*. 2014), with only a few records from near the End Triassic and earliest Jurassic (Riek 1955; Rasnitsyn 1966; Engel 2005b), including one pamphiloid (†*Sogutia*, †Xyelydidae; Rasnitsyn 1977) and one cephoid (genus cf. †*Sepulenia*, †Sepulcidae; Barth *et al*. 2014). Having survived the Carnian Pluvial and the end-Triassic episodes (see Del Corso *et al*. 2020, Vajda & Bercovici 2014, also Fraser & Sues 2011, Tanner *et al*. 2004, Lucas & Tanner 2015), the hymenopteran crown clade experienced substantial diversification via the entomophagous Vespina during the Jurassic and into the Cretaceous, with the potential of more than one transition to “wasp-waistedness” suggested by the paraphyletic “proto” vespinan superfamily †Ephialtitoidea (Li *et al*. 2015). Although the phylogenetic topology of the “parasitica” or parasitican grade remain incompletely resolved (Branstetter *et al*. 2017b; Peters *et al*. 2017), the Stephanoidea, Evanioidea, and Trigonaloidea are indeed supported as close relatives of the Aculeata.

**Fig. 9.**
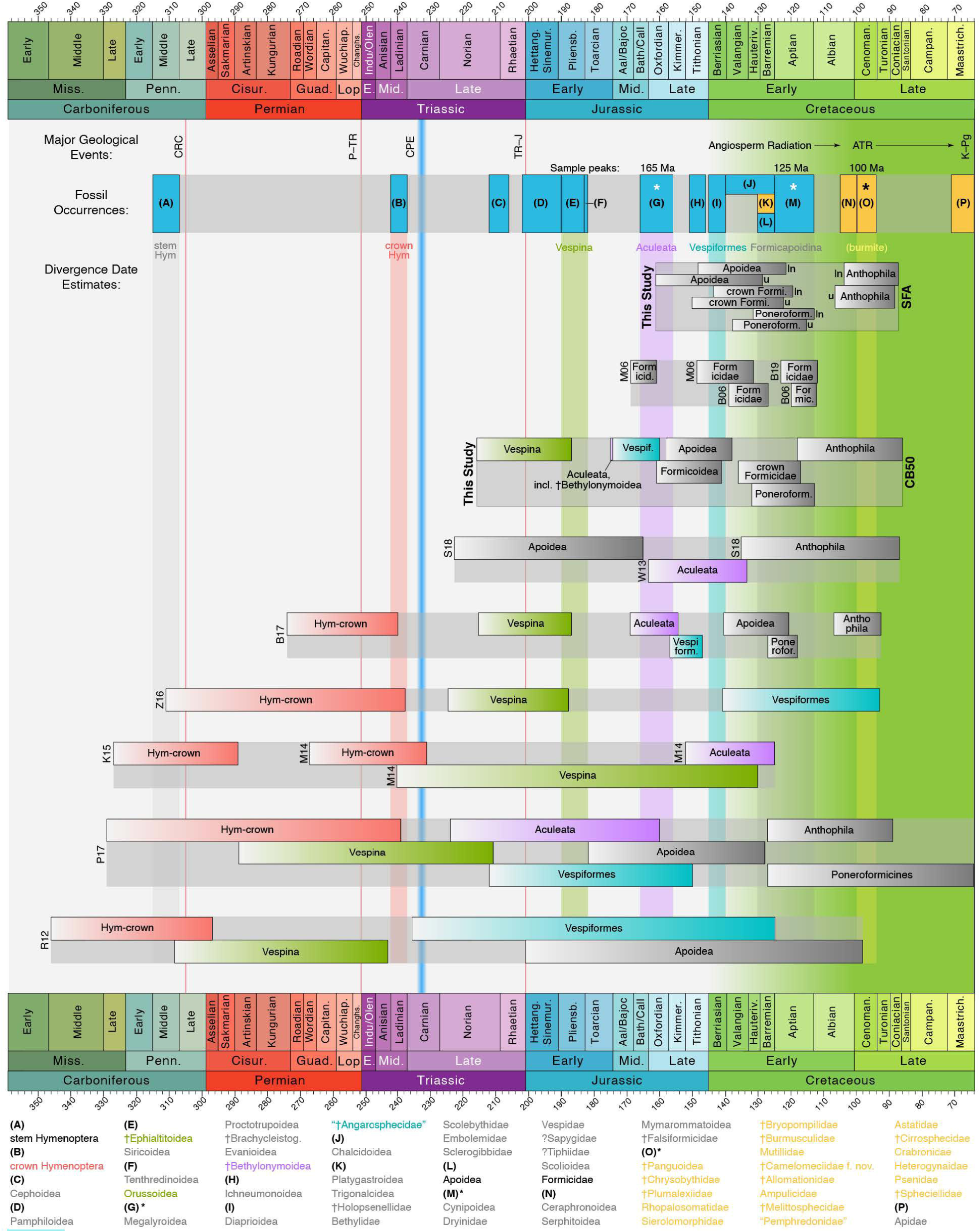
Summary of first appearances of key hymenopteran groups in the fossil record and select divergence dating estimates. Dating estimates are represented as 95% highest probability densities (HPDs). Abbreviations for ““geological events”: CRC = Carboniferous Forest Collapse (*e.g.*, Sahney *et al.* 2010); P–Tr = Permian-Triassic Extinction Event (*e.g.*, Mundil *et al.* 2004; Benton 2005; Winguth *et al.* 2015); CPE = Carnian Pluvial Event (*e.g.,* Del Corso *et al.* 2020); Tr–J = Triassic-Jurassic Extinction Event (*e.g.,* Tanner *et al.* 2004); ATR = the Angiosperm Terrestrial Revolution (*e.g.,* Benton *et al.* 2021); K–Pg = End-Cretaceous Extinction Event (*e.g.,* Schulte *et al.* 2010, Hull *et al.* 2020). Boxes in “fossil occurrences” = stratigraphic ranges of compression (blue) and amber (orange) fossil deposits; letters therein refer to the groups, indicated at the bottom of the panel; asterisks (*) indicate the “big three” sampling peaks (see Fig. 11). Abbreviations for “divergence dates”: CB50 / This Study = diversified-sampling tip-dating analysis; SFA / This Study = Scolioides analysis (u = uniform prior, ln = lognormal prior); B06 = Brady *et al.* (2006); B17 = Branstetter *et al.* (2017b); B19 = Borowiec *et al.* (2019); K15 = Klopfstein *et al.* (2015); M06 = Moreau *et al.* (2006); M14 = Misof *et al.* (2014); P17 = Peters *et al.* (2017); R12 = Ronquist *et al.* (2012); S18 = Sann *et al.* (2018); W13 = Wilson *et al.* (2013); Z16 = diversified sampling tree from Zhang *et al.* (2016). Note that the Vespina and Aculeata in the present study were age constrained following the results of Branstetter *et al.* (2017b), and that the Anthophila 95% HPD range from this study spans both the total and crown clade nodes, *i.e.,* including or excluding †*Melittosphex.* First geological records are as follows (Fm = Formation): **(A)** Stem Hymenoptera: †*Avioxyela gallica* Nel *et al.,* 2013 (Avion Fm); **(B)** crown Hymenoptera: †*Asioxyela paurura* Rasnitsyn, 1969 (Madygen Fm); **(C)** Cephoidea: †*Sepulenia* sp. (Barth *et al.* 2014; Langenberg Formation:); **(D)** Pamphiloidea: †*Sogutia liassica* Rasnitsyn, 1977 (Dzil F,); **(E)** †Ephialtitoidea: †*Proapocritus praecursor* Rasnitsyn, 1975 (Sagul Fm), †*Liasirex sogdianus* Rasnitsyn, 1968 (Sagul Fm); **(F)** Tenthredinoidea: †*Pseudoxyelocerus bascharagensis* Nel *et al.,* 2004b (Schistes Carton Fm); Orussoidea: †*Grimmaratavites mirabilis* Rasnitsyn *et al.,* 2006 (Green Series Fm); **(G)** Megalyroidea: †*Cleistogaster,* other genera (Rasnitsyn 1975; Karabastau Fm); Proctotrupoidea: Heloridae, †Mesoserphidae (*e.g.,* Shih *et al.* 2011; Daohugou Fm), Pelecinidae (*e.g.,* Shih *et al.* 2009; Daohugou Fm), Roprionidae (*e.g.,* Wang 1987; Haifanggou Fm); Brachycleistogastromorpha: †Brachycleistogastridae (*e.g.,* Rasnitsyn 1975; Karabastau Fm); Evanioidea: several families (*e.g.,* Rasnitsyn 1975); †Bethylonymidae: †*Bethylonymellus,* †*Bethylonymus* (Rasnitsyn, 1975; Karabastau Fm); **(H)** Ichneumonoidea: †*Cretobraconus maculatus* Rasnitsyn & Sharkey 1988 (Ulugei Fm); **(I)** Diaprioidea: †*Coramia minuta* Rasnitsyn & Jarzembowski, 1998 (Durlston Fm); †Angarosphecidae, “Pemphredonidae”: see Rasnitsyn *et al.* (1998); **(J)** Chalcidoidea: †*Khutelchalcis gobiensis* Rasnitsyn *et al.,* 2004 (Tsagaantsav Fm); **(K)** Platygastroidea: Scelionidae (*e.g.,* Nel & Azar 2005; Lebanite); Trigonaloidea: †Maimetshidae (*e.g.,* Perrichot *et al.* 2011; Lebanite); †Holopsenellidae: †*Holopsenella* (Lepeco & Melo 2022); Bethylidae: †*Lancepyris opertus* Azevedo & Azar, 2012 (*e.g.,* Engel *et al.* 2016; Lebanite); Embolemidae: †*Embolemopsis maryannae* Olmi *et al.,* 2021 (Perkovsky *et al.* 2021; Wealden amber); Sclerogibbidae: †*Sclerogibbodes embioleia* Engel & Grimaldi, 2006 (Lebanite); **(L)** Apoidea: †*Apisphex penyaleveri* (Rasnitsyn & Martínez-Delclòs, 2000; La Pedrera de Rúbies fm); **(M)** Cynipoidea: †Archaeocynipidae (*e.g.,* Rasnitsyn & Kovalev 1988, Kopylov 2014b, multiple Fm); Dryinidae: †*Deinodryinus aptianus* Olmi *et al.,* 2010 (Dzun-Bain Fm); Vespidae: two genera (*e.g.,* Carpenter & Rasnitsyn 1990, multiple Fm); ?Sapygidae: †*Cretofedtschenkia santanensis* Osten, 2007 (Crato Fm); ?Tiphiidae: †*Architiphia rasnitsyni* Darling, 1990 (Darling & Sharkey 1990; Crato Fm); Formicidae: †*Cariridris bipetiolata* Brandão & Martins-Neto, 1990 (Crato Fm); **(N)** Ceraphronoidea: †*Conostigmus dolicharthrus* Alekseev & Rasnitsyn, 1981 (Taimyr amber); Serphitoidea: Serphitidae (Ortega-Blanco *et al.* 2011a; Escucha Fm); Mymarommatoidea: two families (Ortega-Blanco *et al.* 2011b, Escucha Fm); †Falsiformicidae: undescribed species (suppl, fig. 6 in LaPolla *et al.* 2013; Charentes amber); **(O)** all from Kachin amber: †Chrysobythidae: several genera (Melo & Lucena 2019); †Plumalexiidae: †*Plumalexius ohmkuhnlei* Rasnitsyn & Brothers, 2020; Rhopalosomatidae: two genera (*e.g.,* Lohrmann *et al.* 2020); Sierolomorphidae: specimen CASENT0844589, this study; †Bryopompilidae: †*Bryopompilus interfector* Engel & Grimaldi, 2006; †Burmusculidae: †*Burmusculus* spp. (*e.g.,* Li *et al.* 2020); Mutillidae: specimen CASENT0844575, this study; †@@@idae **fam. nov.:** Boudinot *et al.* (2020), this study; †Allommationidae: †*Allommotion* spp. (Rosa & Melo 2021); Ampulicidae: specimen CASENT0844571, this study (other taxa uncertain); †Melittosphecidae: †*Melittosphex burmensis* Poinar & Danforth, 2006; “Pemphredonidae”: several taxa, including specimen CASENT0844582, this study; Astatidae (uncertain), †Cirrosphecidae, Crabronidae (uncertain), Heterogynaidae (uncertain), Psenidae (uncertain), †Spheciellidae: several taxa (Rosa & Melo 2021); **(P)** Apidae: †*Cretotrigona prisca* Michener & Grimaldi, 1988.

The first indication of stinging wasps in the fossil record is of the †Bethylonymidae (†Bethylonymoidea, Apocrita *incertae sedis*), described from the Mid to Late Jurassic Karabastau formation (Callovian–Kimmeridgian, 166–152 Mya) and suggested to be ancestral to the Aculeata, albeit directly derived from †Ephialtitoidea and perhaps close to Stephanoidea (Rasnitsyn 1975). These wasps are small, apparently prognathous, and have an elongated, muscular pronotum; overall, they roughly resemble members of the extant lineages Bethylidae or Sierolomorphidae, with the primary distinctions of the fossils from the first family being the complete wing venation and, relative to the second, the arcuate second free Rs abscissa (*i.e.*, the first after Rs+M) and the unpetiolated metasoma. Based on the gestalt of the †Bethylonymidae, Rasnitsyn (1975) inferred that these wasps were active hunters of the soil and perhaps leaf litter, seeking hosts or prey in sheltered places. Two additional taxa have been attributed to the bethylonymid group, both from the Early Cretaceous: †*Bethylonymellus feltoni* Rasnitsyn & Jarzembowski, 1998, a wing from the Durlston formation of England (Berriasian, 145–140 Mya), and the large-eyed †*Meiagaster cretaceus* Rasnitsyn & Ansorge, 2000 from the La Pedrera de Rúbies Formation of Spain (Hauterivian–Barremian, 133–125 Mya).

True, or definitive crown Aculeata are undoubtedly present by the midpoint of the Early Cretaceous, as families of the traditional Chrysidoidea—Bethylidae, Scolebythidae, Dryinidae, and Sclerogibbidae, plus the enigmatic †Holopsenellidae—are recorded from Lebanese amber (Hauterivian–Barremian; Prentice *et al*. 1996; Olmi 2000; Engel & Grimaldi 2006a, 2007; Azevedo & Azar 2012; Engel *et al*. 2016a), with Embolemidae very recently recovered from the coeval Wealden amber (Perkovsky *et al*. 2021). Earlier fossils attributed to Aculeata are much more difficult to evaluate, as all are compression fossils and are either wings with poorly preserved bodies or wings only. The oldest are all putative Apoidea from the coeval Durlston and Lulworth Formations, both aged to the earliest Cretaceous and together including the species †*Pompilopterus difficilis* Rasnitsyn & Jarzembowski, 1998, †*P. wimbledoni* Rasnitsyn & Jarzembowski, 1998, †*Iwestia provecta* Rasnitsyn & Jarzembowski, 1998 (Rasnitsyn *et al*. 1998), plus the slightly younger †*Archisphex crowsoni* Evans, 1969 from the Wadhurst Formation (Valangian 140–133 Mya; Evans 1969). With the exception of †*Iwestia*, which is stated to belong to Pemphredoninae (Rasnitsyn *et al*. 1998), all of these earliest putative Aculeata are attributed to the family †Angarosphecidae Rasnitsyn, 1975. Note that our analyses support these fossils as Vespaculeata *i.e.*, all Aculeata with the exception of traditional Chrysidoidea, but are equivocal about superfamilial placement; see Section 2 of below for our treatment of these taxa. A profusion of †Angarosphecidae are described from deposits throughout the Valangian and Albian (for synopsis, see table 1 of Zheng *et al*. 2021 or Table S2 here; see also “On fossil Apoidea and the oldest Vespaculeata” in Section 2 below), and two genera preserved in Kachin amber have recently been attributed to this plesiomorphically defined group, †*Burmasphex* Melo & Rosa, 2018 and †*Decasphex* Zheng *et al*., 2021. Because †Angarosphecidae in general and the Lulworth and Durlston fossils in particular represent not only the earliest potential crown Aculeata, but also potential Apoidea, they are of very high interest for the evolution of the stinging wasps.

Other definitive vespiform Aculeata appear in the fossil record during the Valangian– Barremian (136–125 Mya) interval of the Early Cretaceous. These fossils represent further “†Angarosphecidae”, two genera of putative Scoliidae from the La Huérgina and La Pedrera de Rúbies Formations of Spain, †*Archaeoscolia* Rasnitsyn & Martínez-Delclòs, 1999 and †*Cretoscolia* Rasnitsyn & Martínez-Delclòs, 1999 (see also Rasnitsyn & Martínez-Delclòs 2000), both attributed to the subfamily †Archaeoscoliinae Rasnitsyn, 1993. The former is strongly supported as a true scolioid in our study, but the latter is effectively unplaceable. The next geological interval, that of the Aptian Age (125–113 Mya), includes several more putative Scoliidae, definitive Vespidae (†*Curiosivespa* Rasnitsyn, 1975, †*Priorvespa* Carpenter & Rasnitsyn, 1990), putative Tiphiidae (†*Architiphia* Darling, 1990 in Darling & Sharkey 1990), putative Sapygidae (†*Cretofedtschenkia* Osten, 2007), and a potential crown ant based on our present analyses (†*Cariridris* Brandão & Martins-Neto, 1990, see also Ohl 2004). These Aptian fossils have been recovered from several localities, most notably the Laiyang (*e.g.*, Zhou *et al*. 2020), Yixian (*e.g.*, Chang *et al*. 2017), Khurilt (= Dzun-Bain, Kopylov *et al*. 2020), and Crato Formations (*e.g.*, Martill *et al*. 2009). No other first appearances of vespiform Aculeata occurs before the Kachin amber deposit, although definitive stem Formicidae are recovered from the late Albian Charentese and Baikura Formations, including the unplaceable male-based genus †*Baikuris* Dlussky, 1987, the sphecomyrmine †*Gerontoformica cretacica* Nel & Perrault, 2004, and the haidomyrmecine †*Haidomyrmodes mammuthus* Perrichot *et al*., 2008.

Undoubtedly the most important fossil deposit for understanding the paleodiversity of the Aculeata is that of the mid Cretaceous Kachin (Burmese) amber (∼ 99 Mya: Shi *et al*. 2012, Mao *et al*. 2018; see also Balashov 2021). While there was about a three-quarter century gap in the study of insects from this amber (Zherikhin & Ross 2000; Grimaldi *et al*. 2002), there has been a veritable explosion of interest and controversy in the past decade, with 230 hymenopteran species described in 107 articles published between 2011–2021 (Fig. 10). As of the first day of 2022, there were 3 families and 9 genera of “symphyta” described from Kachin amber, 31 families and 74 genera of Vespina excluding Aculeata, 29 families and 87 genera of Aculeata, and one superfamily—†Panguoidea—which is potentially sister to the Aculeata including

**Fig. 10.**
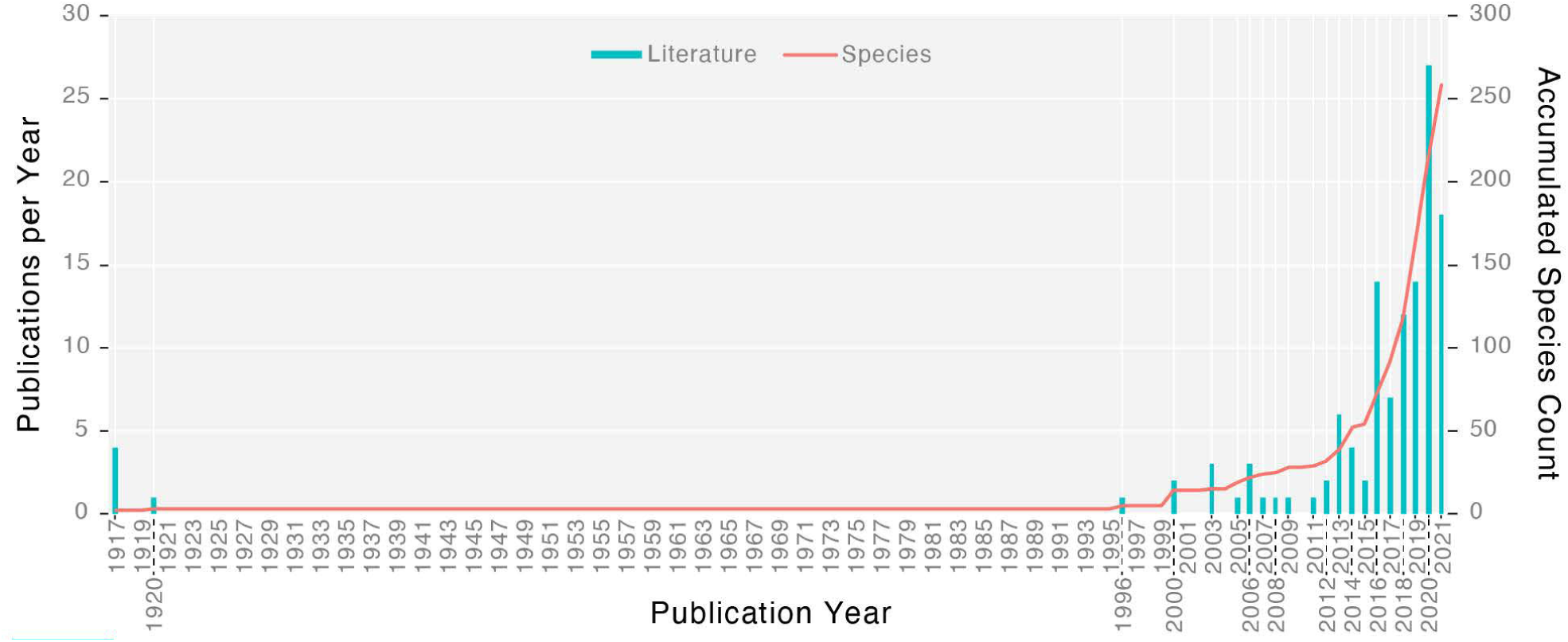
Accumulation of species described from Kachin amber and count of publications per year.

†Bethylonymidae. When species occurrences per deposit are plotted against deposit base age (Fig. 11), it is apparent that mid Cretaceous amber comprises one of the “big three” sampling peaks of Mesozoic Hymenoptera; when these species sampling peaks are subdivided by relative contribution per deposit, it becomes clear that Kachin amber is the most significant record of aculeate biodiversity before the End Cretaceous crisis (Fig. 12). Further, a trend forms when family and genus occurrences per “big three” plus Triassic sampling are compared (Fig. 13): Simultaneous decline of phytophagous “symphyta” and rise of Aculeata during the Early Cretaceous, the epoch of angiosperm radiation. At present, it is not totally clear if this pattern is due to alpha taxonomic bias favoring description of “parasitica” and Aculeata, or whether “symphyta” have indeed suffered extinctions during the Early Cretaceous.

**Fig. 11.**
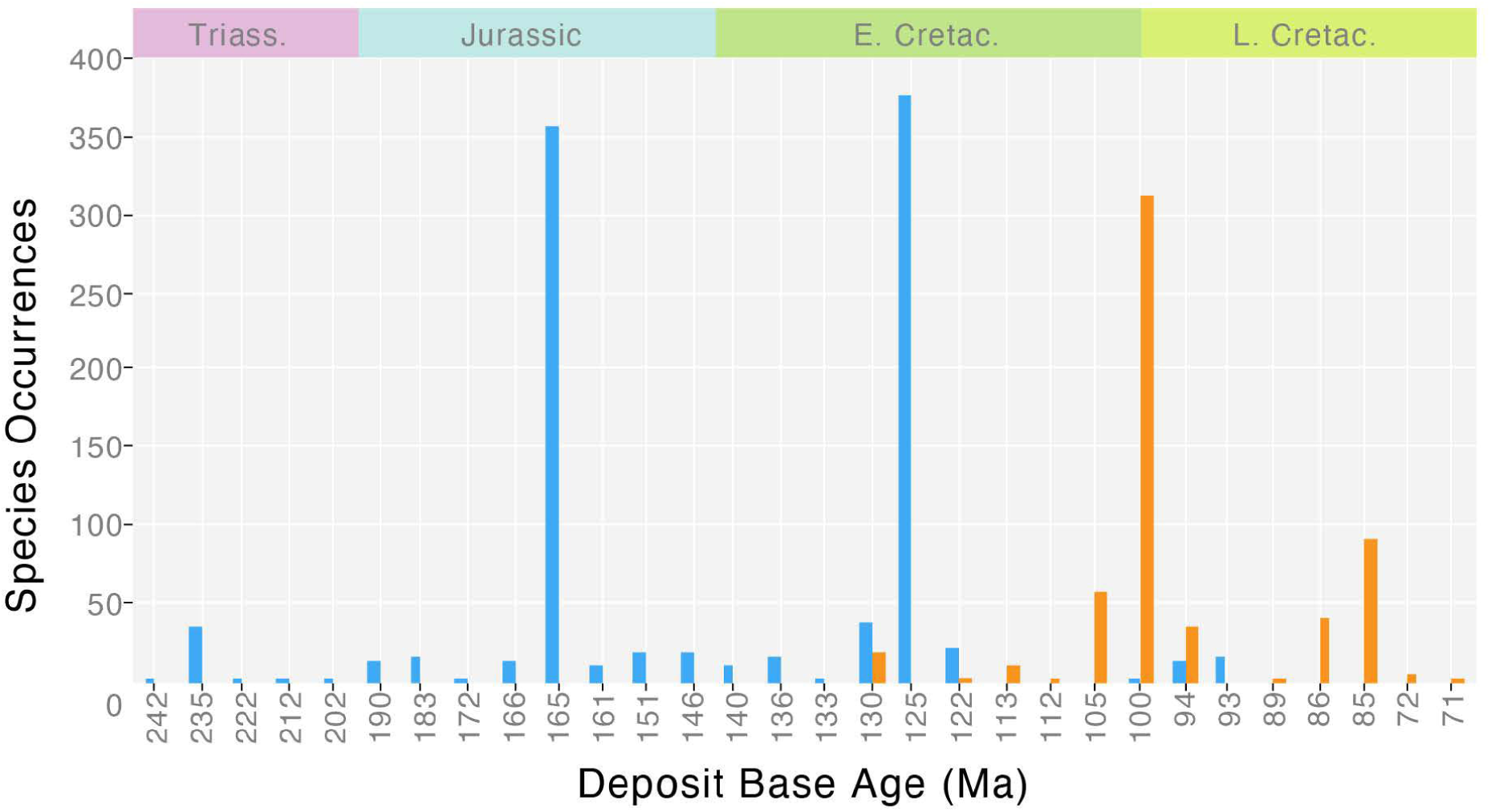
Species occurrences by deposit base age and taphonomy. Blue = compression fossils, orange = amber.

**Fig. 12.**
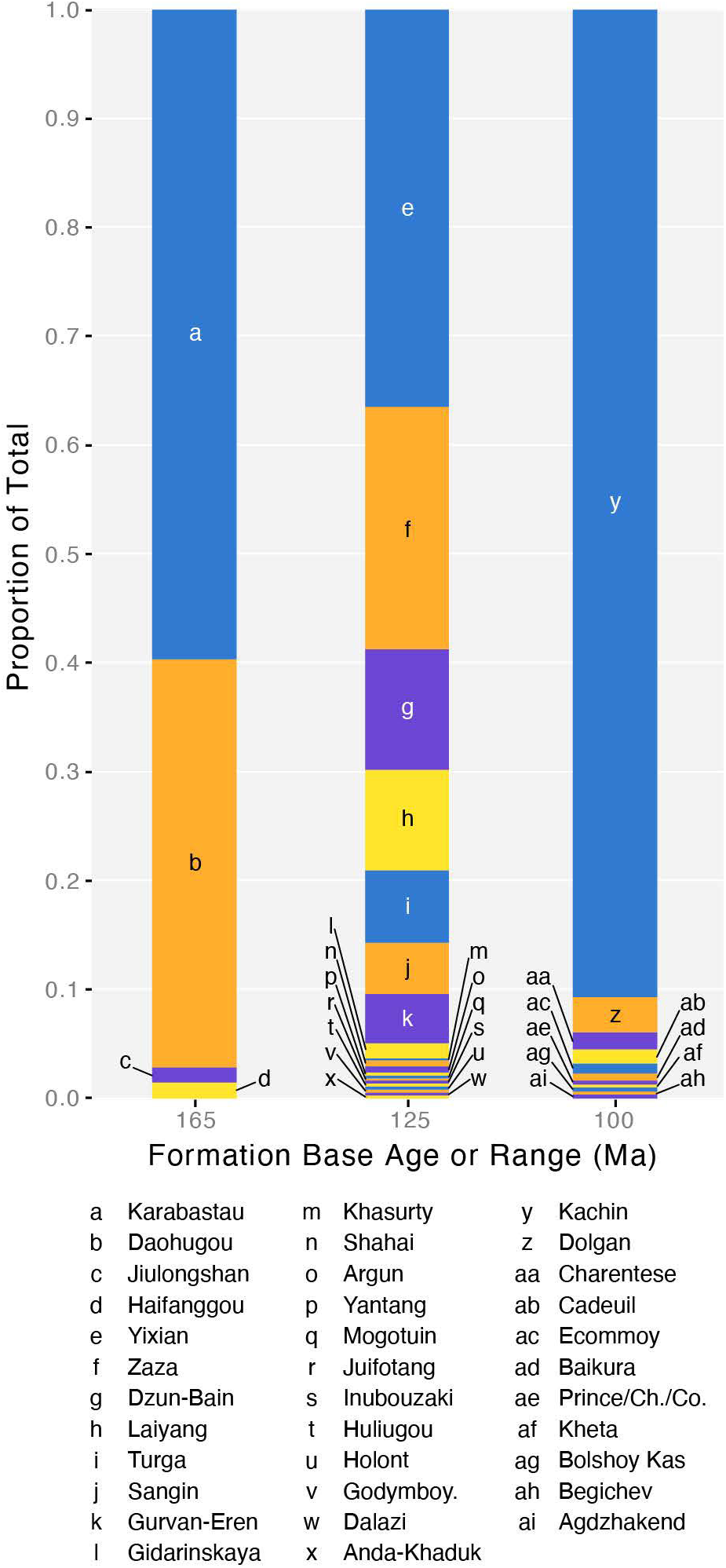
Depositional composition of the three major species sampling peaks of for the Mesozoic Hymenoptera.

**Fig. 13.**
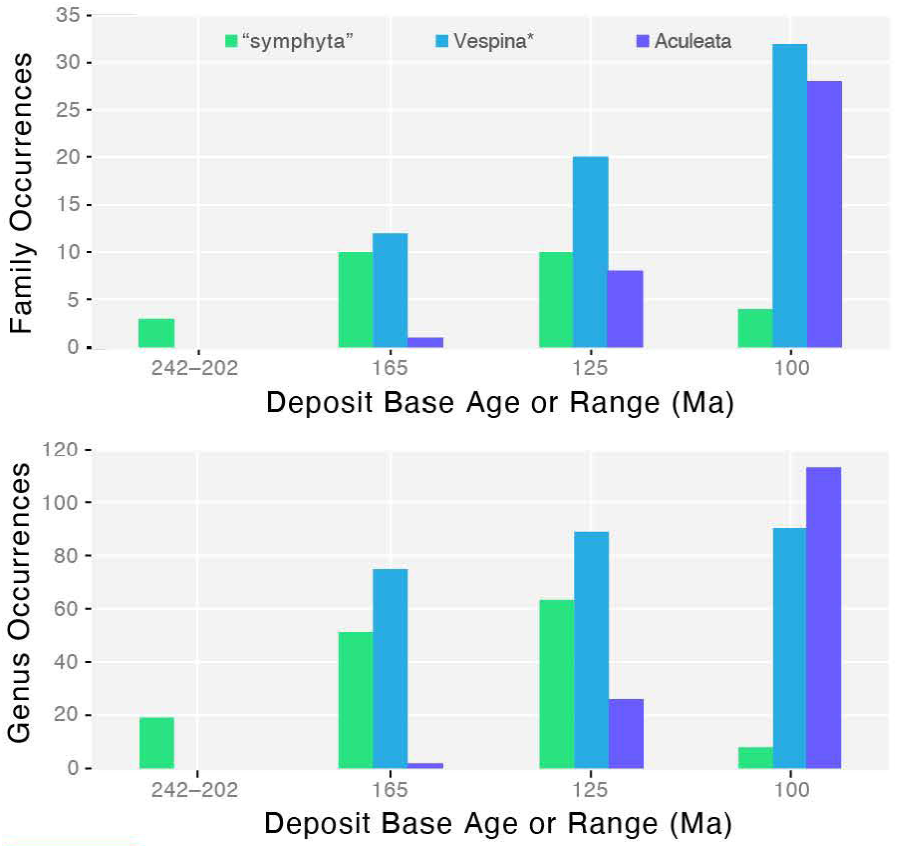
Distribution of family (top) and genus (bottom) occurrences across the three major Mesozoic Hymenoptera sampling peaks plus the summed sampling of the Triassic. “Symphyta” = all non-Vespina Hymenoptera; Vespina* = all Vespina excluding Aculeata; Aculeata = all Aculeata, including †Bethylonymidae.

**Fig. 14.**
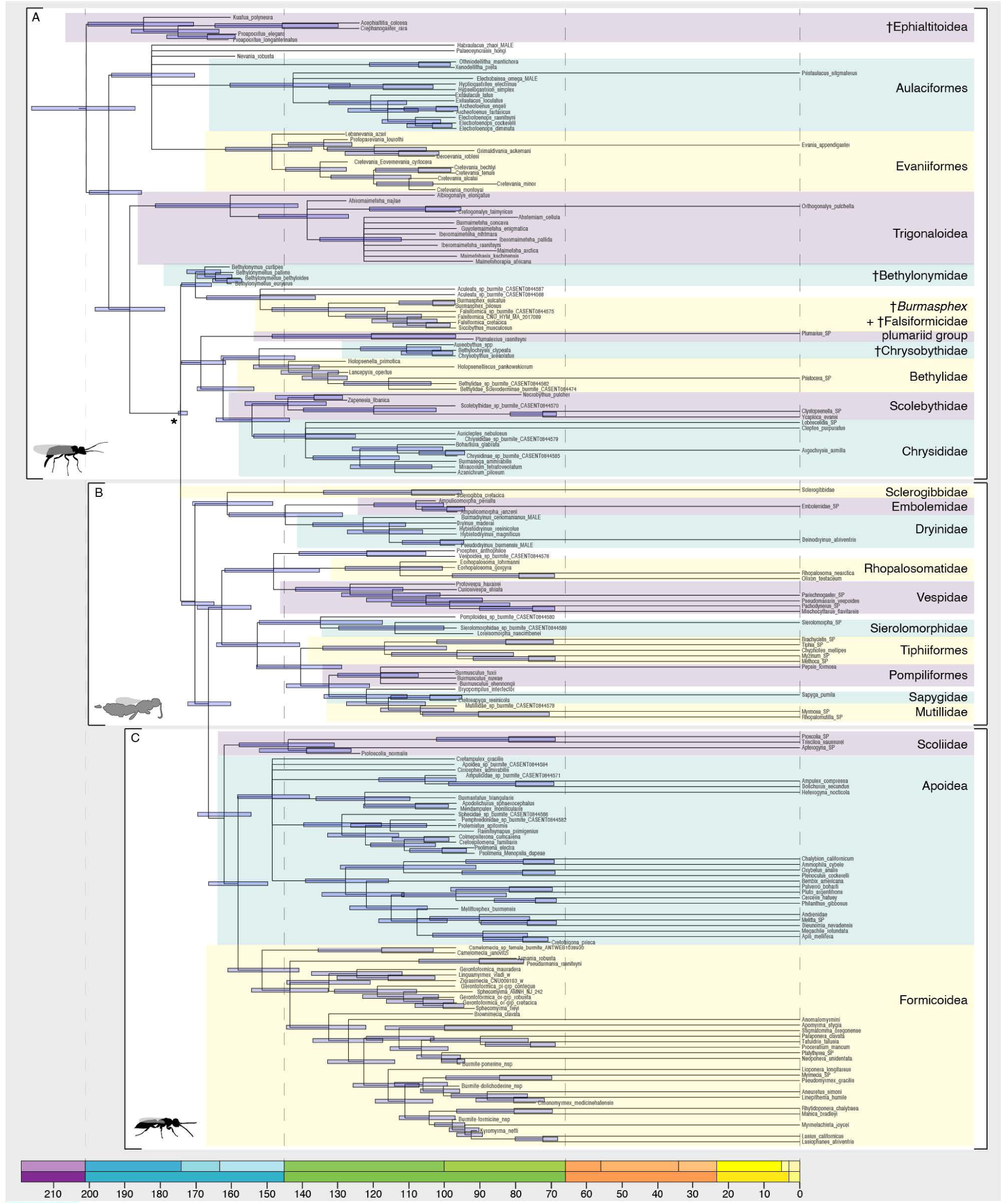
Overview of the diversified 199-50CB fossilized birth-death tip-dating combined-evidence analysis. Although we scored 575 terminals overall, we took an “artisanal data” approach by excluding extant taxa derived from splits < 65 Ma to fit the assumptions of diversified sampling, and we selected only those fossil terminals which were > 50% complete, with the addition of the most complete †Bethylonymidae. Panels **A–C** outline the tree sections focused on in the next three figures; the colored highlight boxes indicate clades which are compressed for clarity in the figures below. The asterisk in panel **A** indicates the Aculeata node, which was provided with an upper age limit.

Among the aculeate Kachin biota are representatives of 11 extinct and 19 extant families across all but one of the extant superfamilies (see Results Part II for the 8-superfamily 34-family classification), with the exception being Scolioidea. At the genus level, all but five of the 87 are extinct, with the extant genera belonging to Bethylidae, Dryinidae, and Sclerogibbidae. These extinct/extant proportions across ranks strongly indicate that the Kachin biota is transitional and captures the formation of infrafamilial diversity via the generic lineages which did not survive to the Cenozoic. The extinct families are of special interest as they provide meaningful information for understanding the paleoecology and evolutionary morphology of their extant relatives. From this perspective, two families are of particular note: †Melittosphecidae, a monospecific family of putative Anthophila, and †@@@idae **fam. nov.**, a lineage found to be sister to all other ants in the present study. With the present classification, the remaining families are attributed to Apoidea (“†Angarosphecidae”, †Allommationidae, †Cirrosphecidae, †Spheciellidae), Pompiloidea (†Bryopompilidae, †Burmusculidae), Chrysidoidea *sensu stricto* (†Chrysobythidae, †Holopsenellidae, †Plumalexiidae), or unplaced (†Falsiformicidae). Additionally, the recently described superfamily †Panguoidea may represent a lineage of stem Aculeata. That the species accumulation curve by literature year is nearly vertical at the time of writing (Fig. 10) indicates that our collective knowledge of the Hymenoptera from Kachin amber is not yet exhaustive; we should expect many new discoveries across phylogenetic depths in the coming years.

The final range of the aculeate fossil record to address is the post-Kachin Late Cretaceous. The 70 fossils described from this interval increasingly show the modernization of the Mesozoic aculeate fauna, including crown Formicidae (Formicinae: †*Kyromyrma*; Dolichoderinae: †*Chronomyrmex*) and Apidae (putative Meliponini: †*Cretotrigona*). Notably, only one of the aculeate families recorded from this time span is extinct (†Plumalexiidae); the remaining 10 are definitive members of extant families. This pattern weakly suggests broad-scale turnover of the aculeate fauna, but the necessity of further sampling is obvious from the documented diversity of this span relative to Kachin amber (Figs. 11, 12) and the limited record at the family level (∼1/3 extant families sampled). Formicidae are the best represented family from this interval, yet only those fossils from amber are placeable with confidence (Boudinot *et al*. 2020; Results Part II). Two of these ant groups, †Brownimeciinae and “†Armaniinae/†Armaniidae” are of substantial phylogenetic importance, as the former have been reported as the sistergroup to the crown Formicidae (Barden & Grimaldi 2016) and the latter suggested to be a “scolioid–formicoid” transitional lineage (Dlussky 1983), albeit of uncertain relationship to ants or other Aculeata (*e.g.*, Wilson 1987b, Dlussky & Fedoseeva 1988, Ward 2007, Borysenko 2017, Barden & Engel 2019). Remarkably, the only recorded hymenopteran from the Maastrichtian is the sweat bee †*Cretotrigona prisca* (Michener & Grimaldi, 1988) (see also: Engel 2000); no other fossil Hymenoptera have been recorded during the Maastrichtian. The Cretaceous–Paleogene turnover will be discussed in the present work with respect to Formicoidea (Sections 4, 5).

### Section 2: Topology and Dating

#### Summary of key topological results

Our topology searches were morphology-only, clockless, stratified by data completeness by taxon, and either provided with “molecular scaffolds” on extant taxa (*i.e.*, soft constraints) or not. Support decayed with the addition of less complete taxa (Fig. 8); support values are mapped onto our 50CB-matrix combined-evidence chronogram in Figs. 14–16 for key nodes. As the topology searches included up to 575 taxa when all fossils are included, the consensus tree presented in these figures is lumped roughly at the family level, with wildcards (“rogue taxa”) and paraphyletic fossil groups pruned. For all output trees please see the online supplementary files. We encourage readers to see the detailed and taxon-by-taxon account of support for fossil placement which we provide in Results Part II. We further emphasize the need to include the recently described †Panguoidea and to expand the taxon sampling to the entire Apocrita, which was beyond the scope of the present study. Our most salient results for ingroup taxa are as follows.

The oldest putative family of Aculeata, †Bethylonymidae, consistently cluster with Aculeata under constrained (Bayesian) and unconstrained (ML) conditions. This result is robust to the inclusion of the less well-preserved bethylonymid fossils (*i.e.*, those with < 40% completeness), albeit the family is only weakly cohesive and may very well be paraphyletic, although this in itself is uncertain. Surprisingly, †Falsiformicidae is supported as a member of the basal polytomy of Aculeata, with some support for a close relationship with the putative “angarosphecid” †*Burmasphex*. Because of the likely polyphyly of Chrysidoidea *sensu lato* (Branstetter *et al*. 2017b), the bethylonymid and falsiformicid results imply that the Aculeata passed through a “chrysidoid-like” ancestor.

Within the Chrysidoidea there is limited support for interfamilial relationships, with the exception that †Chrysobythidae consistently cluster with Bethylidae and †Holopsenellidae. Note that the key holopsenelline features recently revealed by Lepeco & Melo (2022) could not be discerned during the scoring phase of the present study; it is possible these fossils represent another lineage attributable to the basal aculeate polytomy. Interestingly, support for the total clade Scolebythidae and †Plumalexiidae + Plumariidae is higher in unconstrained than in constrained analyses. It is possible that this is due to the use of a more complex model in the ML runs, but these specific results were not interrogated further. Within Dryinoidea, the total clades of Embolemidae and Dryinidae are strongly to well-supported, while the core Dryinoidea (Embolemidae + Dryinidae) is best supported in unconstrained analyses with a high proportion of missing data.

With the exception of Aculeata itself, the deepest nodes of the Aculeata are never recovered in unconstrained analysis. In constrained analyses, the Dryinaculeata, Vespaculeata, Vespoides, Scolioides, and Formicapoidina are supported to some degree until the ∼ 50% completeness threshold, beneath which compression fossils are added *en masse*, and support disintegrates. We expect that this pattern is due to the combination of two factors: inclusion of “†Angarosphecidae” fossils and certain wildcard taxa, such as the putative mutillid “†*Mesomutilla*”, which is positively unplaceable due to poor preservation. It is possible that the “†Angarosphecidae” represent the ancestral form of the Vespaculeata and grade into other superfamilies in addition to the Apoidea. For these reasons, we consider analysis of wing-only characters, or geometric morphometric plotting of wing morphospace to be highly desirable. We did not pursue these avenues as our focus was on the origin and evolution of the Formicidae, for which our results were more robust.

Among the Vespoides, superfamily Vespoidea was better supported than Pompiloidea. The total clades of Vespidae and Rhopalosomatidae were very highly supported. The loss of wings in Mutillidae, Tiphiiformes, and various Dryinoidea resulted in obviously false clustering with the Formicoidea in unconstrained analyses. We suggest reanalyzing the present dataset in the ML framework with soft constraints, which we did not do in the present study. A number of fossils clustered reasonably with Sierolomorphidae, Pompilidae, and Mutillidae. Among Scolioides, the present morphological data recovered Apoidea until compression fossils dominated the taxon sampling. The putative anthophilan †*Melittosphex* strongly clustered with Anthophila. See the extended results below for more detail on the vespoidean and scolioidean relationships touched on here, plus those of Scolioidea.

From the myrmecological perspective, and driving the development of the present study, the critical result is the robust support for the sistergroup relationship of the †*Camelomecia* genus group (*e.g.*, Boudinot *et al*. 2020) with the Formicidae. At first glance, and upon deeper examination, these fossils appear highly dissimilar to ants as generally conceived, yet the sistergroup relationship is virtually unambiguously supported under both constrained and unconstrained conditions. For these reasons, we recognize the †*Camelomecia* genus group as the family †@@@idae **fam. nov.**; synapomorphies of the Formicoidea and diagnoses of the two constituent families of this superfamily are provided and discussed below. Within the Formicidae, we find that the erstwhile †Armaniidae are supported in a basal polytomy of the family, that there is some support for monophyly of the putative †Sphecomyrmines recognized here, and †Brownimeciinae are the likeliest sistergroup to the crown Formicidae, as had been previously suggested (Barden & Grimaldi 2016). Further results are outlined in the extended results presented in Results Parts II and IV.

#### Summary of key dating results

In order to obtain detailed dating estimates for the Formicidae and kin, we included a comprehensive sample of fossils from the total clade Aculeata plus select vespinan outgroups. We conducted two sets of dating analyses: (1) combined-evidence analysis including all sampled groups and fossils < 40% complete pruned, with the exception of the most complete †Bethylonymidae (“50CB”) (Figs. 7, 9, 14–17), and (2) molecular-only estimates focused on the Scolioides, *i.e.*, Scolioidea + Formicoidea + Apoidea (“SFA”) (Figs. 9, 17). The SFA analyses implemented either a lognormal or normal prior, and results are presented for both. Both analysis sets implemented the fossilized birth-death (FBD) process with diversified sampling of extant taxa, following the recommendations of Zhang *et al*. (2016). To reduce computation time for the 50CB analysis, we implemented hard constraints on clades with high support from morphology-only analysis. Fig. 9 compares our dating estimates to those of other studies. Before proceeding, it is critical to point out that in our 50CB dating analyses, we constrained the upper ages of the Vespina and Aculeata given the results of Branstetter *et al*. (2017b). This potential biasing decision aside, when fossils are comprehensively sampled and in absence of definitive fossil evidence to inform deeper nodes in our phylogeny, we generally prefer to interpret toward the younger end of our overlapping 95% highest probability densities, aligned with the “minimum age tree” reasoning of Klopfstein (2021).

While the oldest and only putative stem hymenopteran fossil is from the Carboniferous, the first definitive crown hymenopteran is known from the Middle Triassic, representing a gap of nearly 60 million years. All of the age estimates for the crown Hymenoptera recorded in Fig. 9 fall within this gap; the lower ends of the 95% HPDs for these nodes either fall around the Mid Triassic (M14, B17, P17), or around the beginning of the Permian (R12, K15). Notably, the data from Ronquist *et al*. (2012) are congruent with the Mid Triassic estimates when analyzed under the FBD process with diversified sampling (Zhang *et al*. 2016). Given the apparent consensus of the younger estimates, we consider the time span around the Permo-Triassic boundary to be a reasonable period for the crown group age of the Hymenoptera. In contrast to these estimates, the HPDs for the Vespina overlap in the Late Triassic around the Norian Age, with the exception of Ronquist *et al*. (2012). This congruence suggests that the Vespina may have originated during the Triassic, after the composition of the global paleocommunity was altered by the Carnian Pluvial Event.

Among the studies which have provided an age estimate for the Aculeata, the 95% HPDs are either very broad (> 50 Ma span) or comparatively precise (< 30 Ma). Ignoring the studies of Misof *et al*. (2014) and Wilson *et al*. (2013) for which the younger end of the Aculeata HPDs exceed the Berriasian base age of the vespiform Aculeata, the estimates of Branstetter *et al*. (2017b) and Peters *et al*. (2017) overlap in the Mid Jurassic, the geological period from which †Bethylonymidae have been recovered from the Karabastau formation. Our time-constrained analysis strongly indicates that Aculeata may be older. However, our estimate for this node is highly precise and very likely inaccurate, as the date range was “crammed” to the older end of our constraint. However, the HPD range for Vespina from our analysis is as broad as that of Branstetter *et al*. (2017b), suggesting that it is only the aculeatan node which is compressed. Either way, we strongly encourage reanalysis of our data which: (1) removes the age constraints, selectively deletes the characters which are highly homoplastic (high rate) characters, and (3) carefully chooses a diversified sample of well-preserved and old fossils to include.

With respect to the Vespaculeata, our estimate spans the Mid Jurassic and is encompassed by the less precise but perhaps more accurate HPD of Peters *et al*. (2017), as well as the very broad estimate of Ronquist *et al*. (2012) and the random sampling estimate of Zhang *et al*. (2016; not mapped). While our HPD range for this node is older than that of Branstetter *et al*. (2017b) and especially the diversified sampling scheme of Zhang *et al*. (2016), we note that the taxon sampling of the latter study did not include many aculeates and excluded the Early Cretaceous fossils, and that the variable placement of †Bethylonymidae in our analysis likely influenced the age of the Vespaculeata. Remarkably, the height of the vespaculeate node is the least congruous among the studies addressed so far.

Across the Aculeata, our 50CB analysis supports the Mid to Late Jurassic as the epochs of origin for superfamilies, and the Early Cretaceous for the total clades of most modern families (Figs. 14–17). This pattern also stands for the Apoidea and the Formicoidea (Figs. 9, 17). The date ranges for Apoidea, crown Formicidae, Poneroformicines, and Anthophila are congruent among our 50CB and SFA analyses, with the primary overlap for crown Formicidae around the Valangian to Barremian Ages. These ranges also overlap with the younger estimate of Moreau *et al*. (2006), the older estimate of Brady *et al*. (2006), and the consensus estimate of Borowiec *et al*. (2019). This congruence compellingly suggests that the crown Formicidae originated sometime during the first half of the Early Cretaceous (∼145–125 Mya), antedating the first potential crown formicid from the fossil record, the Aptian-aged Crato fossil †*Cariridris bipetiolata*. Within the crown Formicidae, the crown clades of the major groups Poneria and Formicae are supported as having originated during the second half of the Early Cretaceous, certainly before the deposition of Kachin amber from which we report new, undescribed crown ants (Figs. S17–19).

#### On fossil Apoidea and the oldest Vespaculeata

The oldest previously hypothesized vespiform Aculeata—and supported by our analyses—are four fossils attributed to the Apoidea: †*Pompilopterus difficilis*, †*Pompilopterus wimbledoni*, †*Archisphex crowsoni*, and †*Iwestia provecta*. These taxa are represented by wing impressions, with the †*Pompilopterus* species plus †*Iwestia* recovered from the Berriasian Lulworth formation (fm) fossils (145–140 Ma), and †*Ar. crowsoni* from the Valangian Wadhurst Clay fm (140–133 Ma). Rasnitsyn & Jarzembowski (1998) placed †*Iw. provecta* in Pemphredoninae and the three other species in “†Angarosphecidae”. While these fossils are supported as Vespaculeata, they are not consistently recovered with Apoidea: †*P. wimbledoni* weakly clusters with core Pompiloidea, †*Ar. crowsoni* is not meaningfully supported with Apoidea, and the Bayesian and ML results for †*P. difficilis* and †*Iw. provecta* are conflicting. Whereas †*P. difficilis* receives some degree of support in the ML analyses, with an ultrafast bootstrap value of 94, †*Iw. provecta* is supported close to other putative pemphredonines in the Bayesian analysis, with ∼ 0.60 BPP. Based on these results, we conclude that there is limited justification to use these fossils as node calibrations for the crown Apoidea, albeit they are reasonable to use for the crown Vespaculeata. The next candidates for oldest Apoidea are seven species from the lower member of the Weald Clay (136–130 Ma): †*Angarosphex bleachi*, *†An. consensus*, †*Archisphex boothi*, †*Ar. proximus*, †*Pompilopterus keymerensis*, †*P. leei*, and †*P. worssami*. In brief, none of these fossil species cluster with Apoidea in our present analyses, and there is limited to no meaningful support for placement elsewhere in the Vespaculeata. Among the numerous other fossils attributed to the Apoidea prior to Kachin amber, only five species cluster with or are otherwise strongly supported as Apoidea: †*Angarosphex penyaleveri* (130–125 Ma, La Pedrera de Rúbies fm), †*An. baektoensus* (130–113 Ma, Sinuiju fm), †*An. saxosus* (125–123 Ma, Yixian fm), †*An. lithodes* (125–113 Ma, Laiyang fm), and †*Cretosphecium lobatum* (125–113 Ma, Dzun-Bain fm). To capture the phylogenetic information from our present analyses, we designate †*An. penyaleveri* as the type species of a new genus, forming †*Apisphex penyaleveri* **gen. nov. comb.nov.**; †*Apisphex* **gen. nov.** thus represents the oldest definitive apoid with the narrowest stratigraphic range. Therefore, for calibration of the Apoidea crown node, †*Ap. penyaleveri* is the strongest choices based on analyses across all levels of data completeness.

We emphasize that further study is necessary, particularly for those compression and amber fossils attributed to the “Pemphredoninae”, given the rupture of this group under the weight of phylogenomic analysis (Branstetter *et al*. 2017b; Sann *et al*. 2018). Revised analyses and codings may improve placement of other putative Apoidea of the Early Cretaceous.

### Section 3: Ancestral State Estimation: Morphology

As described in the Methods, we conducted ancestral state estimation (ASE) across all characters (576C), diet and nesting (D3, N2), the eusociality supercharacter (10S, 5S), and on 43 key morphological traits (AASE). The results for the morphological estimates are addressed below and are outlined in detail in Part IV, and those for the behavioral estimates are addressed in Section 4 below. We synthesize the results from our ASE analyses in Section 5.

#### Brief Results & Discussion

To explicitly account for topological, branch length, and state identity uncertainty, all of our ASE analyses were conducted using a posterior distribution of chronograms in a Bayesian framework. Because we were interested in the total evolution of the Aculeata, we estimated morphological ancestral states for all 576 characters across all nodes of the 303 taxon 50CB FBD tip-dating analysis (576C) and for 43 key characters across the 86 taxon SFA node-dated BD analyses (AASE). Comparing the two analysis sets, we observe that the polarity of certain characters is sensitive to fossil inclusion or exclusion, that support for a given state at a given node may become misleadingly high in the extant-only analyses, that state information can be lost without fossils, and that stepwise transitions may be estimated as saltational jumps in fossil-less analysis (Fig. 18). Such observations are not new but further highlight the importance of fossil inclusion in the estimation of ancestral states (*e.g.*, Puttick 2016, Mongiardino Koch *et al*. 2021) and phylogeny (Louca & Pennell 2020, Mongiardino Koch & Parry 2020, Parry 2021), particularly in the context of the present study.

**Fig. 15.**
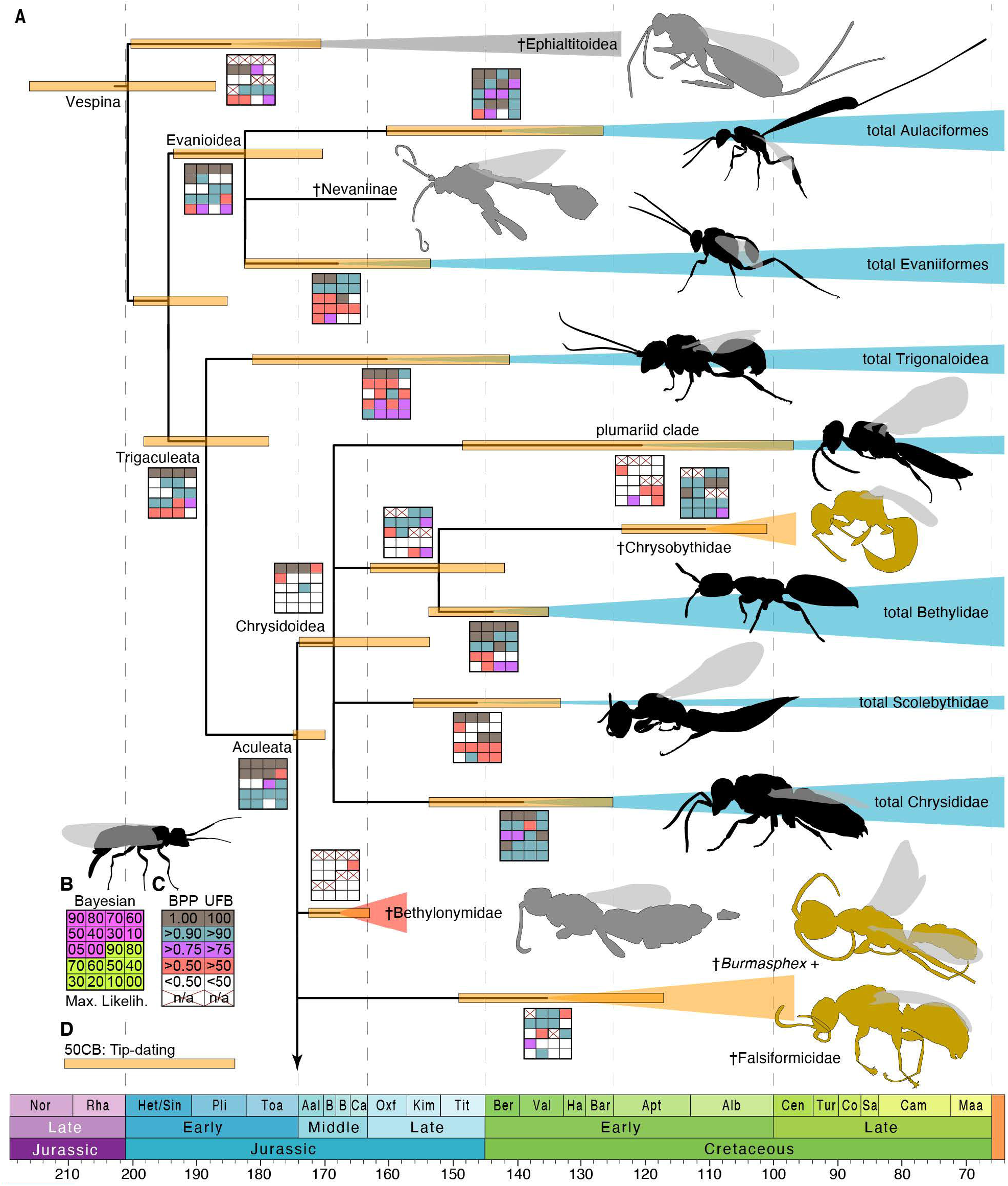
Summary of topological support and dating estimates for Aculeata and sampled outgroups. **(A)** Simplified chronogram of Aculeata and outgroups. **(B, C)** Topological support legend: “Bayesian”, clockless morphological analyses in MrBayes constrained by a molecular scaffold, “Max. Likelih.”, unconstrained IQTree analyses; boxed numbers in **(B)** indicate data completeness threshold; “BPP”, Bayesian posterior probability, “UFB”, ultrafast bootstrap support. The xyelid silhouette above the support legends roughly suggests the ancestral form of the clade under consideration. **(D)** Color scheme for 95% highest probability densities: “50CB”, 50% complete plus †Bethylonymidae matrix with diversified sampling (199 taxa).

**Fig. 16.**
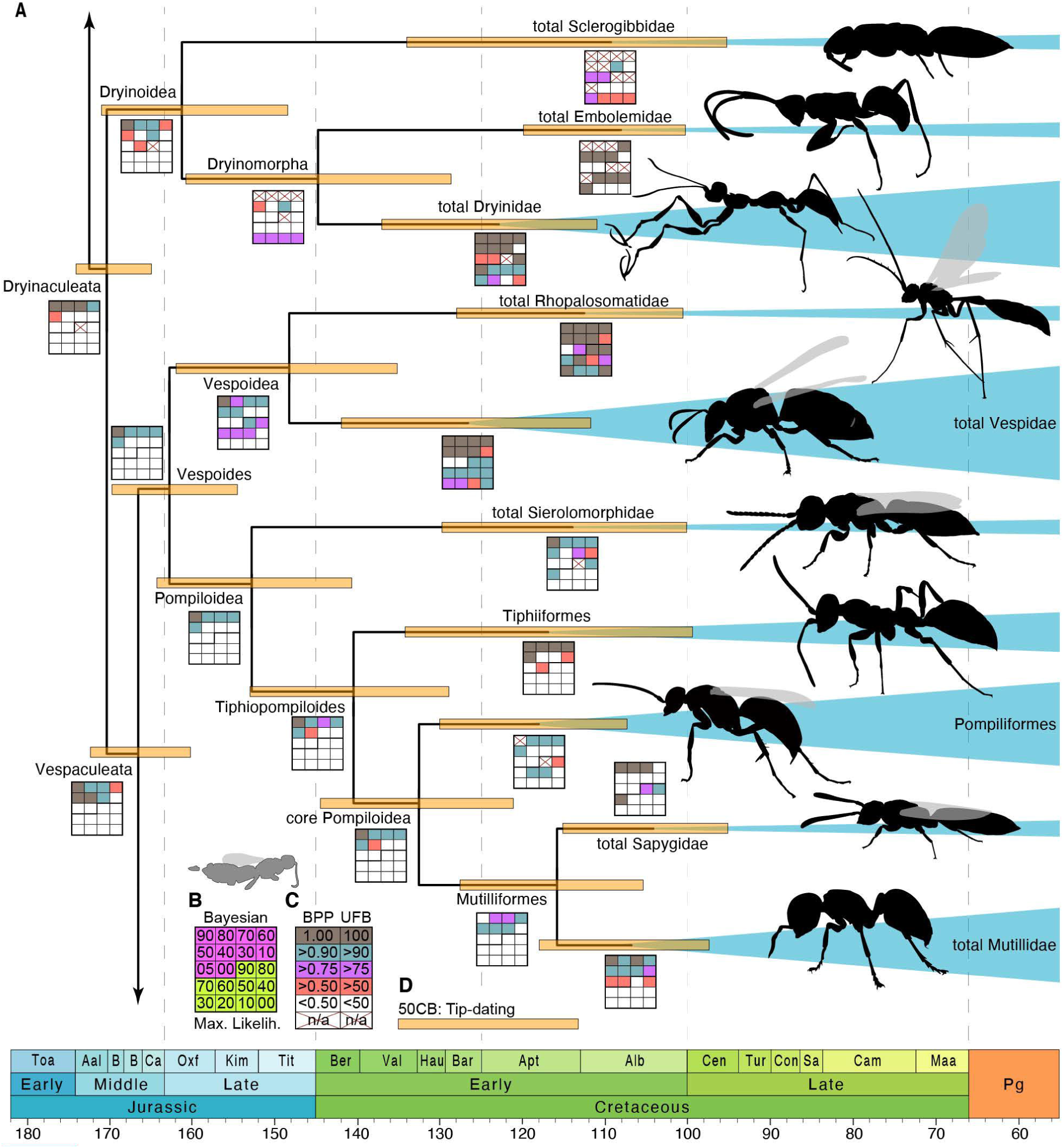
Summary of topological support and dating estimates for clade Dryinaculeata, in part. **(A)** Simplified chronogram of Dryinaculeata, excluding clade Scolioides. **(B, C)** Topological support legend: “Bayesian”, clockless morphological analyses in MrBayes constrained by a molecular scaffold, “Max. Likelih.”, unconstrained IQTree analyses; boxed numbers in **(B)** indicate data completeness threshold; “BPP”, Bayesian posterior probability, “UFB”, ultrafast bootstrap support. The bethylonymid silhouette over the support legends roughly indicates the ancestral form of the Aculeata, based on our present analyses. **(D)** Color scheme for 95% highest probability densities: “50CB”, 50% complete plus †Bethylonymidae matrix with diversified sampling (199 taxa).

**Fig. 17.**
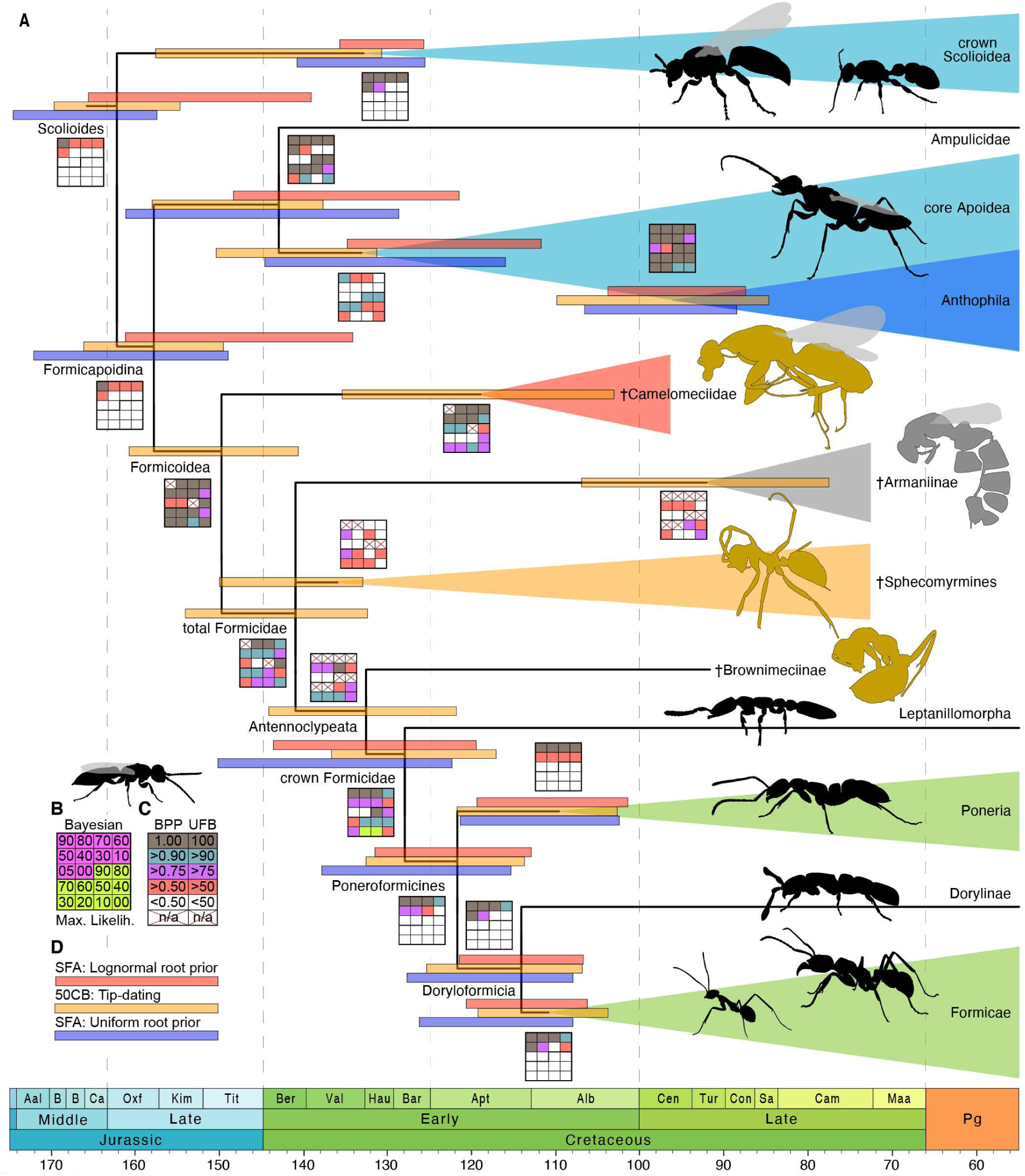
Summary of topological support and dating estimates for clade Scolioides. **(A)** Simplified chronogram of Formicoidea and closest outgroups. **(B, C)** Topological support legend: “Bayesian”, clockless morphological analyses in MrBayes constrained by a molecular scaffold, “Max. Likelih.”, unconstrained IQTree analyses; boxed numbers in **(B)** indicate data completeness threshold; “BPP”, Bayesian posterior probability, “UFB”, ultrafast bootstrap support. The sierolomorphid silhouette above the support legends roughly indicates the ancestral form of the Vespaculeata and Scolioides, based on our present analyses. **(D)** Color scheme for 95% highest probability densities: “SFA”, molecular-only Scolioidea-Fonnicoidea-Apoidea matrix; “50CB”, 50% complete plus †Bethylonymidae matrix with diversified sampling (199 taxa).

**Fig. 18.**
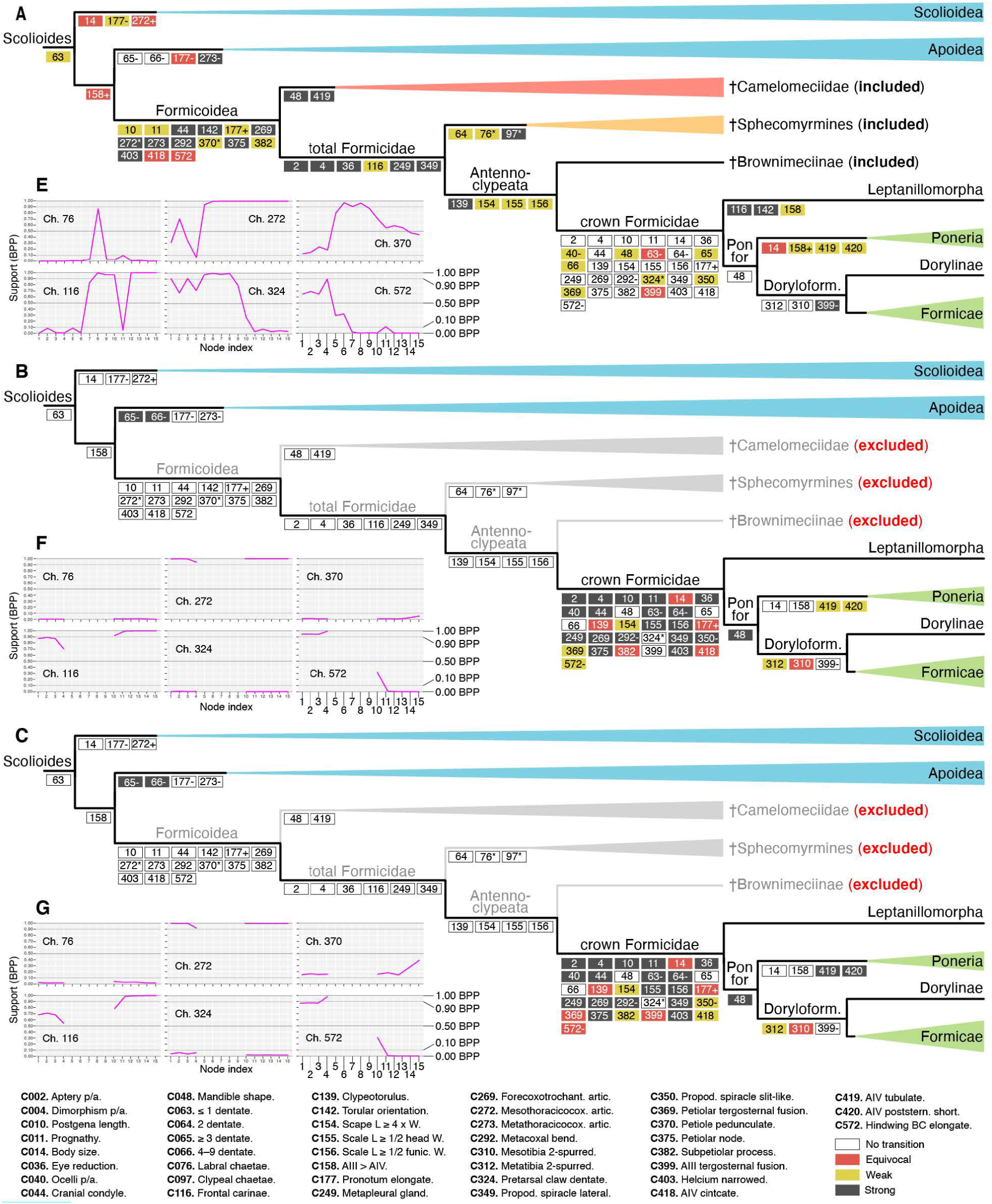
Alternative ancestral state estimation analyses demonstrating importance of fossil terminals. When fossils are excluded the polarity of certain characters can invert (*e.g.*, char. 116, **E-G),** support can become misleading (*e.g.*, char. 370, **E-G),** certain state information is completely lost (*e.g.*, char. 76, **E-G),** and stepwise transitions **(A)** can be estimated as “jumps” **(B, C). (A–C)** Trees with mapped transitions; note that stabilization of support at shallower nodes is not indicated; see character definitions, extended transformation series, and the supplementary material for more detail. **(E–G)** Bayesian posterior probabilities (BPP) for select characters across the depicted nodes. Support values range from 0.00 (= maximal support for state 0) to 1.00 (maximal support for state 1); values of 0.50 are completely equivocal; values > 0.90 or < 0.10 are interpreted as “strongly” supported. Node indices: 1 = Scolioides, 2 = Scolioidea, 3 = Formicapoidina, 4 = Apoidea, 5 = Formicoidea, 6 = †@@@idae, 7 = total Formicidae, 8 = †Sphecomyrmines, 9 = Antennoclypeata, 10 = crown Formicidae, 11 = Leptanillomorpha, 12 = Poneroformicines, 13 = Poneria, 14 = Doryloformicia, 15 = Formicae. **(A)** Tip-dating 303-50CB analysis; average BPP across the depicted nodes = 0.94. **(B)** Extant-only SFA analysis implementing a strict clock model; average BPP across depicted nodes = 0.92; favored by stepping-stone integration relative to the relaxed clock model (BF > 2). **(C)** Extant-only SFA analysis implementing an uncorrelated lognormal (UCLN) relaxed clock model; average BPP across depicted nodes = 0.89; disfavored by stepping-stone integration. State transitions for the Leptanillomorpha could not be estimated for the SFA analyses due to the availability of nucleotide data for only a single terminal of this clade.

Here, we detail our approach to large-scale statistical character polarity testing as our guide studies which including fossils and several hundred (to thousands) of morphological characters relied on parsimony reconstructions for transition series (O’Leary *et al*. 2013, Field *et al*. 2020). Procedurally, a custom Python code was used to extract estimated states and Bayesian posterior probabilities (BPPs) for each indexed node from the summary tree. These “harvested” estimates were then placed in a color-coded spreadsheet, and node-by-node (column-by-column) comparisons were manually made for each character. As all states across all characters and nodes included in the “harvest” spreadsheet were highly supported on average (∼0.98 BPP), fluctuation in BPP for a given character from a parent to a daughter node indicated either a potential state transition, variability within a daughter clade, or statistical uncertainty due to lack of information about a character state among sampled terminals (*e.g.*, development of the jugal lobe in fossil clades, a challenging condition to evaluate which was often scored as uncertain, “?”). Because we had no template studies to work from, we developed the following scheme for interpreting these potential transitions.

**(1)** We interpret transitions to be *unequivocal* if a state changes from a deeper to a shallower node with > 0.90 BPP for both nodes. For example, char. 410 transitions from 0 to 1 from the Amblyoponomorpha to the Amblyoponinae node with maximal support. Thus, as state 0 = helcium axial or infraaxial and 1 = helcium supraaxial, we interpret the supraaxial condition to be an unequivocal synapomorphy of Amblyoponinae, despite previous hypotheses of the plesiomorphy of this condition. **(2)** In many cases, the BPP of a given state at a deeper node will drop below 0.90 but remain above 0.75 before switching to another state with > 0.90 BPP; we would interpret such a pattern as *strong* evidence for a transition, but not unequivocal. For this reason, we also developed the “state support interpretation” scheme outlined in the Methods (*i.e.*, x > 0.90 = strong, 0.90 > x > 0.75 = weak, 0.75 > x = equivocal). **(3)** In other cases, the BPP of a given state will be weak to equivocal for a derived state at a shallower node and would *stabilize* across one or more nested nodes. An example of this is the derived condition of having abdominal segment IV (metasomal III) constricted (char. 418). This transition is equivocally supported for the Formicoidea (BPP = 0.74), weakly supported for the total clade Formicidae (BPP = 0.84), and maximally supported at the Antennoclypeata node (BPP = 1.0). Such stabilizations are frequent and may reflect state uncertainty among fossils or transitional variability before fixation within a given clade. **(4)** Often, the BPPs of both the deep and shallow node with differing states would be < 0.90 but > 0.75, thus there is *weak* evidence for a transition. **(5)** Likewise, the potential of multiple transitions may be indicated by an unequivocal state at a parent node and weak to equivocal support at a daughter node, although this is also often associated with variability within the shallower clade. For example, elongation of antennomere III in the Poneria displays this pattern (char. 158), where deeper BPPs are weak, BPPs for Proceratiinae and Ponerinae stabilize on the plesiomorphic condition, and BPPs in amblyoponomorphan nodes are equivocal. **(6)** In cases where a fossil has an uncertain state, but a potential transition occurs from the parent to daughter nodes, the support for the state of the deeper and shallower nodes would be strong to unequivocal, but the fossil node would be equivocal, *e.g.*, char. 423 transitioning from 0 to 1 from the Mutilliformes to crown Mutillidae through the total clade Mutillidae, as stridulitra presence or absence could not be confidently evaluated in the stem mutillid and was scored as uncertain (“?”). **(7)** The final scenario of note is uniform uncertainty among clades for which a given condition is inapplicable, such as the shape of the second mediocubital crossvein (2m-cu) in crown ants. In these circumstances, we would mark the deepest node for which the inapplicability applies as inapplicable (“-”), then ignore the variable among shallower nodes.

Using this schema, we interpreted and transcribed all potential transitions in the extended transformation series (Part IV) via node-by-node comparisons for all characters (note that our “harvest” spreadsheet is included in the supplementary material). For each potential transition we indicated the statistical support for the polarity of the daughter state and discussed sources of uncertainty or ambiguity of interpretation. In future study, we highly recommend programming this process for efficiency; if this were to be done, a matrix summary of plesiomorphic or uncertain states for a given node would also be of diagnostic value. Ideally, diagnostic combinations could be output with characters sorted by incisiveness (uniqueness) and polarity. The main elements of the procedure are the states and their support at the root node (*i.e.*, the global set of symplesiomorphies), the state differences and their respective posterior probabilities between parent and daughter nodes, and whether or not the character is applicable to that node. With an index or dictionary of character definitions, it will be possible to automatically produce clade-by-clade diagnoses of estimated synapomorphies, with explicitly stated “equivocal” to “unequivocal” support phrases. We did not pursue this here given the experimental nature of our approach; however, we note that this approach is also amenable to modeling exercises and statistical hypothesis testing. For paleoecological hypotheses arising from our large-scale ancestral state estimation analysis, see below.

### Section 4: Ancestral State Estimation: Behavior

#### Background

The primary factor which has arguably led to the ecological success of the Formicidae is reproductive altruism, labeled “eusociality” since the prior midcentury. Before proceeding, it is necessary to clarify our definition of this term. “Eusociality” was coined by for bees in which daughters are recruited by nest foundresses and, after formalization by Michener (1969), the term was defined by Wilson (1971) as the condition of having overlapping generations, cooperative brood care, and reproductive division of labor. Boomsma & Gawne (2018) have recently argued for a return to the “superorganismality” concept of Wheeler (1911), emphasizing the developmental differentiation of morphologically discrete castes over the continuous behavioral castes of Wilson (1971, 1975, 1985) and the irreversibility of this “major transition” in the sense of Maynard Smith & Szathmáry (1995). Drawing on Crespi & Yanega’s (1995) discussion of this transition and its associated terminology, Boomsma & Gawne (2018) recognized the distinction between the reversible social organization with some degree of overlapping generations and cooperative brood care and the irreversible expression of morphologically distinct workers as “facultative” and “obligate eusociality”, respectively. We follow this conceptual clarification and employ these terms here and later in the text.

Considering the importance of superorganismality as one of the major transitions in the organization of life, our question is *to what degree can fossils inform our understanding of eusocial origins*? With the definition of obligate eusociality provided above, the only groups comparable to ants are Dictyoptera, Vespidae, and Apidae, as these clades contain lineages that have morphologically defined worker castes. While we cannot address the role of kin selection and monogamy from fossils, we can capitalize on two findings to address the problem of the apparent singularity of eusocial origins in the Formicoidea. The first is that certain stem Formicidae display winged-wingless polyphenism, de-alation, and cooperative brood care (*e.g.*, Perrichot *et al*. 2008, Barden & Grimaldi 2012, Perrichot *et al*. 2020, Barden *et al*. 2020, Boudinot *et al*. 2020, 2022), which is the best available evidence for obligate eusociality in these extinct clades. The second is our unexpected finding that @@@idae **fam. nov.** is the sistergroup to the total clade Formicidae, which provides an additional node for estimating ancestral states while also accounting for the uncertain presence or absence of derived conditions in these fossils. Additionally, we took this opportunity to explore complex models for the evolution of dependent discrete characters, including the construction of alternative rate matrices, and the implementation of reversible jump MCMC (rjMCMC) (*e.g.*, Pagel & Meade 2006, Klopfstein *et al*. 2015) and hidden state models (*e.g.*, Beaulieu *et al*. 2013, Beaulieu & O’Meara 2016).

We chose to focus on nesting, diet, and degree of eusociality as all known obligately eusocial species maintain nests or defensible spaces (even if simple) and all forage or otherwise provision (even if within an enclosed space); it is therefore reasonable to infer that these conditions are also true for fossils with evidence of morphology-associated reproductive division of labor. Nesting and nest reuse is a fundamental feature of social groups, as it leads to the formation of groups themselves, it provides physical structure for (reproductive) dominance interactions, and it facilitates the care of offspring by non-ovipositing females (West-Eberhard 1989, Bonabeau *et al*. 1997). Further, nests provide defensible spaces and superorganismal infrastructure (Starr 1985, Hansell 1993, Crespi 1996, Bonabeau *et al*. 1997, Duarte *et al*. 2011, Tschinkel 2013; also Andersson 1984 and references therein), as well as a continuous food source in some cases (Crespi 1996, Chouvenc *et al*. 2021). We also posit that in the context of the Vespina, the transition from non-nesting to nesting represents a shift from manipulating a host to modifying the environment for the optimization of offspring development. Similarly, derivation of nesting behavior may serve as a necessary intermediate condition for multiple or continuous (progressive) provisioning (Evans, 1977), thus providing a food source that is not limited to the size of a single host body. Further, a transition to progressive provisioning may perhaps easily cause uneven nutrient distribution among brood, resulting in reduced flight and reproductive potential in underfed individuals (Trible & Kronauer, 2017) which could then have acted as ground-based foragers (Evans, 1977; Peeters et al., 2020) and perhaps as nest-defenders (although see Kronauer & Libbrecht, 2018).

*Results: Diet (D3) and nesting (N2)*.

We conducted 16 alternative analyses of diet and nesting as independent behavioral characters (D3, N2) to evaluate their basic phylogenetic signal and to serve as a contrast to the considerably more complex supercharacter analyses. Table 4 presents the marginal log likelihoods for all analysis variants, from which we have chosen our focal analyses to discuss. The only equivocal set of variants was that of the SFA matrix nesting analyses, from which we focus arbitrarily on the F81 test, as the results are more-or-less identical.

Parasitoidy is robustly supported as ancestral to the Vespina sampled in the 303t-50CB matrix (Fig. 19A.1–2). When the diet of stem Formicoidea is treated as unknown, the posterior probability of predation is fractional and increases to dominance at the Antennoclypeata node (20 in Fig. 19; for definition of this clade see Boudinot *et al*. 2020 and its diagnosis in Results Part IV). When lineages of the †Sphecomyrmines are scored as “predatory” and †@@@idae **fam. nov.**is scored as uncertain, the posterior probabilities for parasitoidy and predation are about evenly split at the Formicoidea and †@@@idae nodes and strongly support predation at the total Formicidae node (18 in Fig. 19). We emphasize that we scored †*Gerontoformica* as uncertain, and that there is no obvious manner in which to determine whether †Sphecomyrmines were omnivorous with the exception that they do not have large seed-crushing mandibles, thus the “predatory” scoring is an approximation for the provisioning condition. Transition to omnivory is supported as occurring well within the crown Formicidae (Formicae, 25 in Fig. 19) and the Apoidea (Anthophila; not shown). Because omnivores are also provisioners, we excluded this state from the supercharacter analyses. Exclusion of fossil tips results in a jump from parasitoidy in the ancestor of Formicapoidina to predation in the crown Formicidae, effectively removing evidence for stepwise transitions. See our taxonomic synopsis and extended transformation series below for definition of the clade Formicapoidina.

**Fig. 19.**
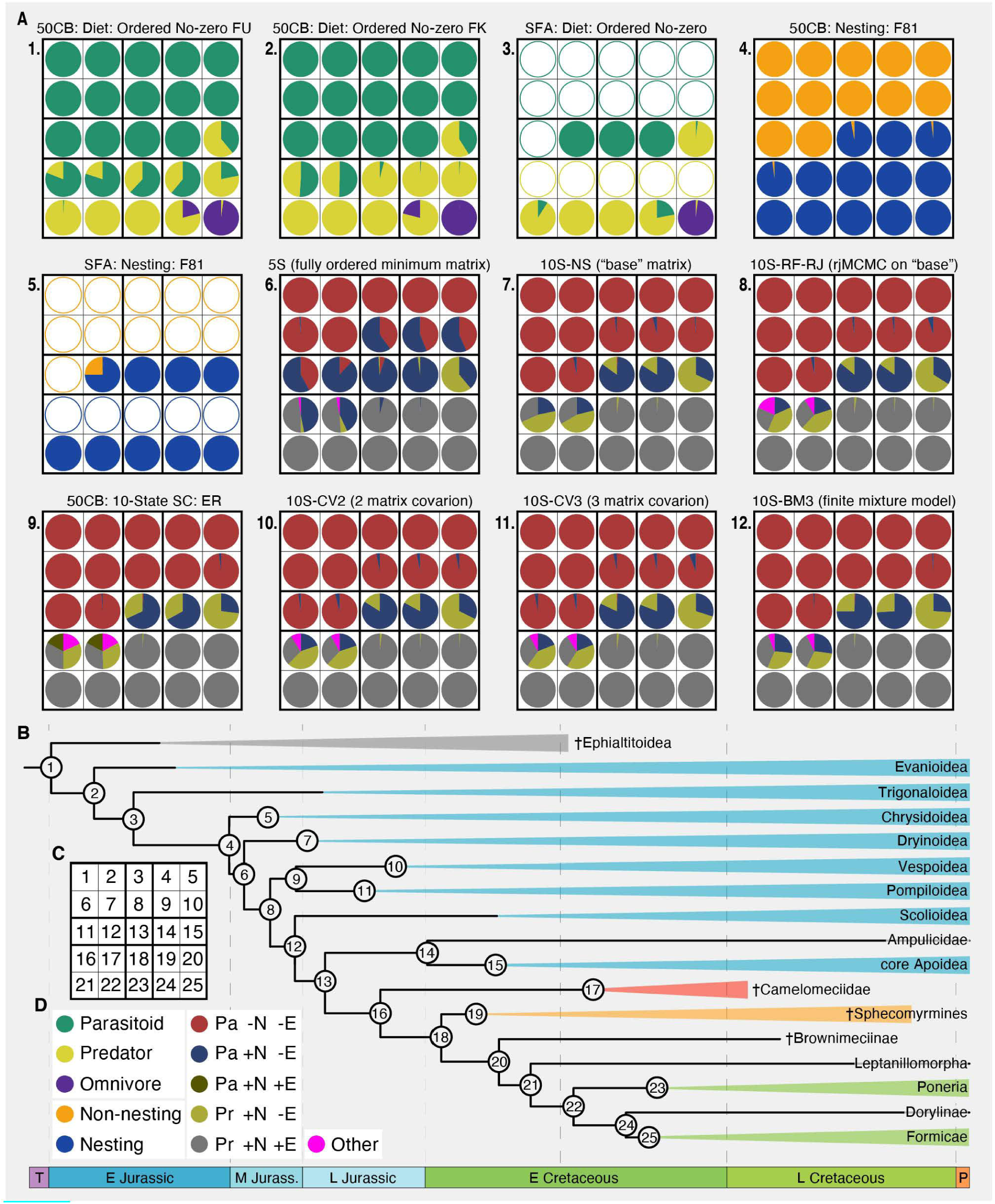
Results of the behavioral reconstruction analyses (D3, N2, S10 set 1). Diet (D3) analyses in quilts 1-3 (green, yellow, purple); nesting (N2) analyses in quilts 4 and 5 (orange, blue); 5- and 10-state supercharacter analyses in quilts 6-12 (6-tone). Labels for the supercharacter analyses are provided in Table 2. (A) Node-by-node support for estimated states. **(B)** Simplified summary tree with corresponding node indices. **(C)** Legend for node index grids. **(D)** Legend for state identities. Abbreviations in A: 50CB = 303-taxon 50% complete + †Bethylonymidae matrix; SFA = Scolioida matrix; FU = diet states of stem Formicidae scored as uncertain; FK = diet states of stem Formicidae scored as “predator’; F81 = Felsenstein 1981 model applied to morphology, *i.e.,* allowing state frequencies to vary. Abbreviations in **D:** Pa = parasitoid; Pr = provisioner (predator); -N non-nesting; +n = nesting; -E = non-social or facultatively eusocial; +E = obligately eusocial.

Similar to parasitoidy, non-nesting was also strongly supported as ancestral to the Vespina, despite the lack of information for the fossil group †Ephialtitoidea (Fig. 19A.4). Multiple transitions are estimated across the Aculeata but are restricted to the Vespaculeata, *i.e.*, all Aculeata with the exclusion of the superfamilies Chrysidoidea and Dryinoidea (results not shown; see the Supplementary Material for more information). The Scolioides are nearly maximally supported as having the derived nesting state in both tip-dating and extant-only analyses (Fig. 19A.4, 5), and nesting is supported as a consistent feature among the descendent nodes of Apoidea and Formicoidea.

#### Results: 5S

The **5S** supercharacter analyses evaluated state-dependent transitions among the five combinations of three substates that were observed among our sampled taxa; this analysis represented the “extreme stepwise” pathway, which we treat as the null hypothesis for our more-complex analyses. Among our focal nodes the 5S matrix provided support for nesting parasitoidy at the Vespaculeata and daughter nodes (Fig. 19A.6), with nesting becoming the dominant condition at the Scolioides node. These two results stand in contrast to all 10S analyses, wherein nesting is not meaningfully supported until the Formicapoidina. Notably, support for nesting at the Scolioides node is only recovered in the SFA N2 analysis. In 5S, the transition to non-eusocial provisioning is supported to some degree for the core Apoidea, with support increasing therein (results not shown). For the Formicoidea and †@@@idae **fam. nov.**, the probabilities are more-or-less evenly split between non-nesting parasitoidy and total eusociality, with eusociality becoming nearly unequivocal at the total Formicidae node, despite uncertainty among certain groups. This transition from uncertainty at the Formicoidea node to maximal or near-maximal support for eusociality at the total Formicidae node is consistent across all of the more-complex analyses, strongly indicating that eusociality was a fixed inherited trait of the Formicidae.

#### Results: 10S

*1. “Base” matrix.* The nonstationary 10S “base” matrix (**10S-NS**) represents the most stringent “stepwise” pathway (Fig. 19A.7), disallowing simultaneous transitions of two or more subcharacters. All subsequent analyses are variants of this model thus it stands as the primary reference point. Similar to the 50CB N2 analysis (Fig. 19A.4) and in contrast to that of the 5S matrix (Fig. 19A.6), nesting is nearly maximally supported as a derived condition of the Formicapoidina, with probability for this state distributed between non-eusocial parasitoidy (> 0.75 Bayesian posterior probability [BPP]) and non-eusocial provisioning (< 0.25 BPP). This also contrasts with the diet only (D3) analysis, wherein the parasitoid condition formicapoidinan node is maximally supported. Also dissimilar to the 5S analysis, non-eusociality comprises the majority of the probability mass for the Formicoidea node, although eusociality has a value of ∼0.29 BPP. †@@@idae **fam. nov.** retain this uncertainty, and a transition to obligate eusociality is supported for the total Formicidae node. Note that the supplementary materials include maximum *a posteriori* (MAP) graphs for all of the 10S (set 2) analyses.

*2. Base model stationary/non-stationary averaging*. These analyses used rjMCMC to average over stationary versus non-stationary root frequencies for all supercharacters (**10S**-**RF-RJ**: Fig. 19A.8, -**RF-RJ-d/f**: Fig. 20.A.4). In the 10S (set 2) analyses, the posterior probability of the stationary model was approximately 0.4, both when using a fixed tree and the posterior distribution of trees, corresponding to a Bayes factor of about 1.5 in favor of the non-stationary model. Given that neither root frequency model was strongly supported over the other, we averaged over both models for all subsequent analyses (Fig. 20.A). The averaged model (including reversible jump over the stationary and non-stationary versions) has an estimated log marginal likelihood of -68.1 when used in conjunction with the fixed tree. Overall, this analyses largely agrees with the “base” matrix analysis (10S-NS), with the only notable distinctions being that the Formicoidea and †@@@idae **fam. nov.** nodes are further split by a small minority of other, unspecified conditions.

**Fig. 20.**
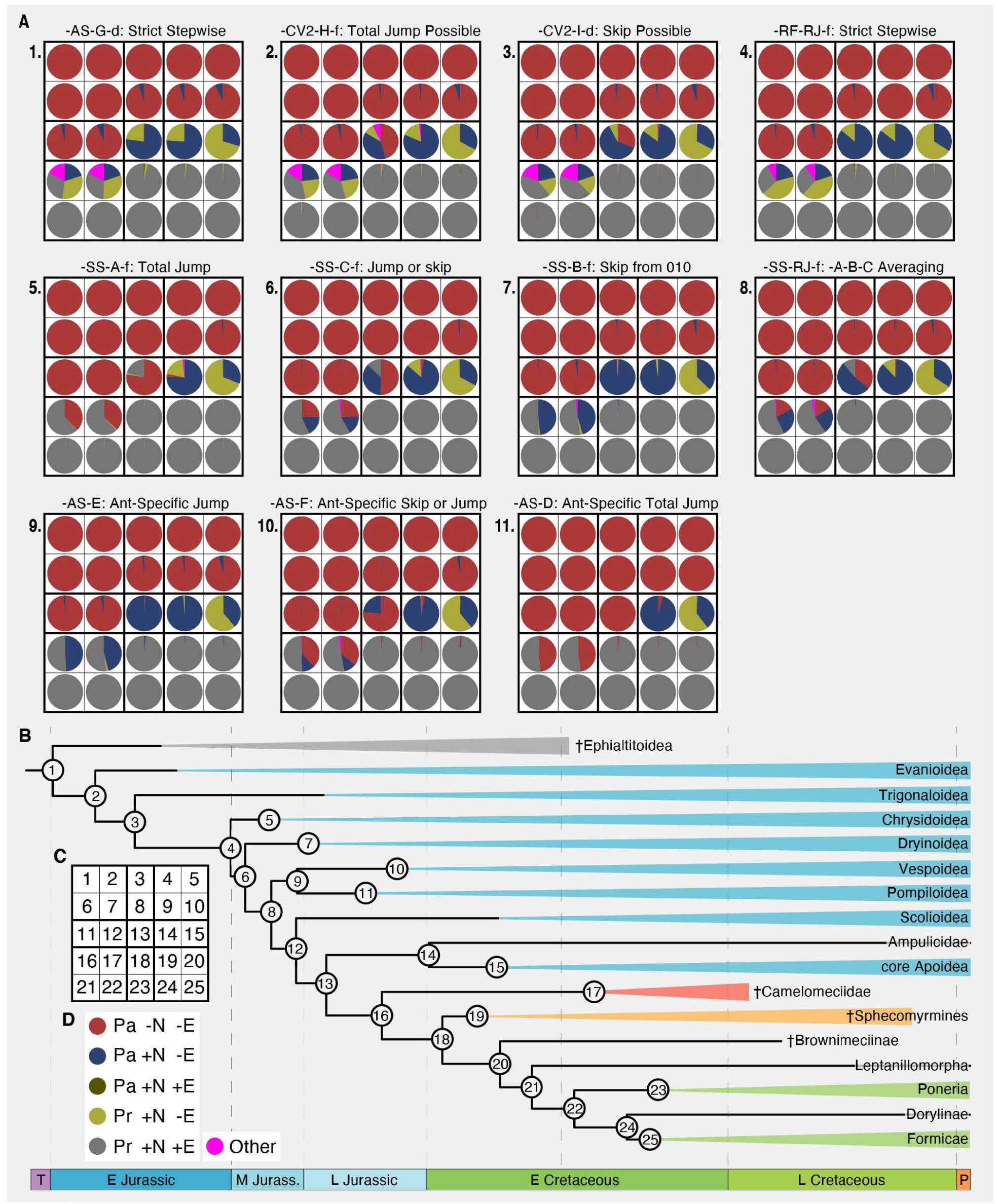
Results of the behavioral reconstruction analyses (S10 set 2). Note that choice of representing results from the -f (fixed tree) and -d (tree distribution) analyses was arbitrary, as both methods provided nearly identical results. **(A)** Node-by-node support for estimated states. **(B)** Simplified summary tree with corresponding node indices. **(C)** Legend for node index grids. (D) Legend for state identities. Model labels in A correspond to those in Table 2; results sorted left-to-right top-to-bottom from lowest log likelihood to highest. Abbreviations in D: Pa = parasitoid; Pr = provisioner (predator); -N = non-nesting; +n = nesting; -E = non-social or facultatively eusocial; +E = obligately eusocial.

*3. Equal rates*. The posterior probability of the equal-rates model (**10S-RF-RJ-ER**) was only 0.06, corresponding to a Bayes factor of about 15.3 in favor of the less constrained model where substitution rates are free parameters. Therefore, with this positive result, we did not pursue ancestral state estimation with this model in the 10S (set 2) analyses. The results from the 10S set 1 analysis for focal nodes are presented in Fig. 19A.9 for comparison to the “base” analysis (10S-NS). The major difference between these two is greater support for provisioning at the Formicapoidina and Apoidea nodes, and the highly equivocal reconstruction of the Formicoidea and †@@@idae **fam. nov.** nodes, for which support is roughly evenly split among provisioning eusociality, non-eusocial provisioning, “eusocial nesting parasitoidy”, and “other”. No other analysis supported eusocial nesting parasitoidy among our focal nodes.

*4. Simultaneous superstate transitions*. These models (**10S-SS-A**: Fig. 20A.5, **-SS-B**: Fig. 20A.7, **-SS-C**: Fig. 20A.6) generally performed better than the “base” and independent root frequencies models (10S-NS, -RF-RJ), despite having more free parameters (Table 5). In addition to the stepwise sequence allowed by the “base” matrix, these models allow eusociality to evolve directly from non-nesting parasitoidy (10S-SS-A), or from nesting parasitoidy but not non-nesting parasitoidy (10S-SS-B), or from both (10S-SS-C). Bayes factors calculated from these marginal likelihoods matched those calculated using rjMCMC across all models (10S-SS-RJ). Our confidence in choice among these models is low because of the limited difference among the marginal likelihoods of these analyses, for which reason we emphasize the results of the -SS-RJ analysis.

Each model agrees with the “base” analysis (10S-NS) in supporting transition from non-nesting parasitoidy at the Scolioides node and maximal support for eusociality at the total clade Formicidae node. Their primary distinctions are the necessity of nesting as an intermediate condition for eusociality, with almost no probability mass for nesting parasitoidy at the Formicoidea node (-SS-A) to almost absolutely favoring nesting (-SS-B). While the support for a nesting-parasitoid intermediate increases for the Formicoidea node from -SS-A to -SS-B through -SS-C, the invariable element is that provisioning and eusociality are simultaneously derived in each, with only very limited support for a non-eusocial provisioning intermediate; this is also the case in the runs that average over these models (10S-SS-RJ). Uniquely, the -SS-A, -SS-B, and -SS-RJ analyses have non-negligible support for obligate eusociality at the Formicapoidina node. Overall, the results of the model averaging analysis (-SS-RJ) most closely matches the “jump or skip” model (-SS-C), with the primary difference being higher support for nesting as an intermediate condition. It is important to recall that these analyses employ a single matrix that still allows for stepwise transitions, and that †@@@idae **fam. nov.** was scored as uncertain for all conditions.

*5. Formicoidea-specific matrices*. Across all 10S (set 2) models, the best performing analyses were a set of those implementing two rate matrices for which one is specific to the Formicoidea (Table 5). We implemented four versions of this matrix: (1) **10S-AS-D** (Fig. 20A.11) allowed the ant specific matrix to “jump” from non-nesting parasitoidy, (2) **10S-AS-E** (Fig. 20A.9) allowed a “skip” from nesting parasitoidy, (3) **10S-AS-F** (Fig. 20A.10), allowed a jump and a skip, and (4) **10S-AS-G** (Fig. 20A.1) did not allow jumps or skips, *i.e.*, the ant-specific matrix was “base” but had independent rates for each of the stepwise transitions. This fourth analysis was the poorest performing of all 10S (set 2) models. The pairwise Bayes factor associated with -AS-D versus - AS-E as calculated based on stepping-stone-estimated marginal likelihoods (3.1) is similar to that estimated by rjMCMC under -AS-F (3.0). However, analysis under -AS-F, which averages over all possible submodels, had the highest posterior probability (0.275) associated with a formicoid-specific matrix allowing provisioning eusociality to evolve directly from non-nesting parasitoidy, nesting parasitoidy, and provisioning-nesting states.

Similar to all other 10S analyses, the Scolioides were maximally to nearly maximally supported as non-nesting parasitoids, and the total clade Formicidae were nearly maximally supported as eusocial. Indeed, these analyses agree with the simultaneous superstate transition (- SS) analyses in providing a (slim) majority of support for eusociality at the Formicoidea node; they are further similar, in that the most complex model (-AS-F) split the remainder of the non-eusocial probability at this node between nesting and non-nesting parasitoidy, albeit with somewhat greater support for the non-nesting condition. The main difference between the -AS and -SS analyses is that the respective models with the highest likelihoods supports ∼50% non-nesting parasitoidy in the former, and ∼50% nesting parasitoidy in the latter. In other words, the state of the non-eusocial remainder at the formicoid node is inverted.

*6. Covarion-like models*. Similar to the -AS models, the covarion-like models implemented two rate matrices, with the distinction being that the second matrix can be switched on or off at any point in the tree given its single hidden character. The second matrix either allowed a total jump to eusociality from non-nesting parasitoidy (**10S-CV2-H**: Fig. 20A.2) or a jump from non-nesting parasitoidy and a skip nesting parasitoidy (**10S-CV2-I**: Fig. 20A.3). The “jump and skip” model (-CV2-I) performed significantly better than the “jump” model (-CV2-H), and in fact better than all other model variations (Table 5).

Both of the covarion-like models approximated the results of the ant-specific model that implemented two base matrices (-AS-G), as well as the stationary/non-stationary rjMCMC averaging on the base matrix analysis (10S-RF-RJ: Fig. 19A.4). Specifically, these models less strongly support eusociality at the Formicoidea node, with the remainder of the probability mass divided between provisioning, nesting-parasitoidy, and “other”. The covarion-like models stand out in that the Formicapoidina node is no longer unambiguously supported as nesting, which is the case for -AS-G and -RF-RJ via the aggregate probabilities of nesting parasitoidy and provisioning.

The sampling behavior of the alternative matrices differed between the two covarion-like models. In the more-restrictive -CV2-H model, the “base” matrix was dominant throughout the phylogeny, without transition to the second matrix, but the posterior probability of sampling the second matrix increased toward the Formicapoidina, with near total split in the Formicoidea. In contrast, stochastic character mapping of matrix usage under the less-restrictive and better-fitting-CV2-I model supports transition to the second matrix along the stem of the Formicoidea. See the supplementary material for these stochastic character maps.

## Discussion

We took multiple approaches to modeling the transformations of morphological and behavioral characters. For the morphological data, we monitored state probabilities across all nodes in our 303-taxon 50CB tip-dating analysis. These estimates were conducted with all of the characters dependent on the same parameter set as they were in a single partition. We relied on the 303-t-50CB analysis for our interpretations as this was the joint estimation which employed the same model as was used to estimate the phylogeny itself (Mk+Γ). In contrast, for our comparative 86-taxon ASE analysis, the node-by-node states for all of the 43 characters were estimated independently. We took a similar approach for our basic diet (D3) and nesting (N2) analyses, effectively assuming that these characters were unlinked via independent analyses on both the 303-t-50CB and ASE trees (Fig. 19A.1–5).

Because all obligately eusocial species are nesting and provisioning (whether continuously or discretely), it is intuitive that these traits have some degree of correlation in their evolution. To treat all three characters as dependent, we combined them into a “supercharacter”, allowing us to model the transition rates among the substates of each subcharacter, similar to the approaches of Pagel & Meade (2006) and Beaulieu *et al*. (2013). In our basic analyses (5S, 10S-NS, 10S-RF-RJ), we forced all substates to transition one-by-one in order by setting the probability of simultaneous substate transitions to 0, as previously done by Beaulieu *et al*. (2013). This set of transition matrices therefore represents our intuitive null hypothesis of stepwise gain of nesting from a non-nesting ancestor followed by provisioning from a parasitoid ancestor, and eventually the origin of obligate eusociality, as evinced by the Vespidae and Apidae. Interestingly, the more-complex models of this set (10S-NS: Fig. 19A.7, 10S-RF-RJ: Fig. 19A.8, Fig. 20A.4) did favor the stepwise pathway for the Formicoidea, while the most restrictive model (5S: Fig. 19A.6) favored dependence on nesting as an ancestral condition with simultaneous transition to provisioning and eusociality.

In order to differentiate the stepwise and “skip” scenarios of eusocial origins in ants, we constructed three alternative model sets. The first set uses the single base matrix but allows for simultaneous substate transitions, with different degrees of stringency (“jump” only: 10S-SS-A; “skip” only: -SS-B; or “jump” and “skip”: -SS-B). The other two sets allow for time heterogeneity through the implementation of two independent rate matrices, one of which was invariably the stepwise base matrix, and the other being one of the -SS models. The most extreme model set (10S-AS) assumes that the evolution of eusociality in ants is distinct relative to other eusocial Aculeata, *i.e.*, an event with singular conditions, by specifying the -SS matrix for the Formicoidea. We relaxed this assumption through the use of a covarion-like model, where a third “hidden” rate matrix controls the switch from one or the other matrices (10S-CV2). Hidden state models (*e.g.*, Beaulieu *et al*. 2013, Beaulieu & O’Meara 2016) have been highlighted as a potential solution for the detection of spurious correlations among characters in phylogenetic comparative methods by Uyeda *et al*. (2018) and have been recently extended to account for “hierarchical” (developmental) characters (Tarasov 2019).

We found that the models which provided ant-specific rate matrices (-AS) had the best fit among all evaluated models (Table 5), followed by the single-matrix simultaneous transition models (-SS). The base model (-RF-RJ), the covarion-like model (-CV2-H), and the stepwise ant-specific model (-AS-G) were multiple log likelihood units worse fitting. The well-fitting -AS and -SS models unanimously supported simultaneous transition to provisioning and eusociality in the Formicoidea, regardless of the estimated condition(s) at the Formicapoidina node (Fig. 20A.5–10), similar to the most restrictive base matrix (5S). In contrast, the base, stepwise ant-specific, and covarion-like models provided some support for the necessity of a non-eusocial provisioning intermediate, as the state probabilities at the formicoid node were split among eusociality, non-eusocial provisioning, nesting parasitoidy, and “other” conditions (Fig. 20A.1– 4). What do these results mean for the evolution of ants; was the evolution of eusociality in Formicoidea mechanistically unique relative to other obligately eusocial insects, or have other factors influence this pattern?

While we expect that the evolution of superorganismality in Formicoidea was contingent on the preadaptations of nesting and provisioning, our modeling analyses indicate simultaneous transition of provisioning and eusociality, a pattern not supported for other obligately eusocial insects. On one hand it is quite possible that this pattern is driven by the loss of information over time (Vermeij 2006); on the other, the ants do appear to be unique among obligately eusocial groups. In terms of species-level diversity, the crown Formicidae are one, two, and three orders of magnitude more diverse than Isoptera, obligately eusocial Apidae (Apini, Meliponini), and obligately eusocial Vespidae (Vespinae), respectively (Table 6). The sense that ants are unique also strengthens when accounting for time, as the crown clades of Isoptera and Formicidae both survived the End Cretaceous crisis and may have been coeval in their transition to superorganismality (Evangelista *et al*. 2019), and even more so given other estimates of a Jurassic crown age for Isoptera (*e.g.*, Legendre *et al*. 2015, Bourguignon *et al*. 2015, Jouault *et al*. 2021c).

It is possible that the evolution of obligate eusociality in the ants was relatively switch-like, correlated with the stabilization of within-sex winged-wingless polyphenism. While winglessness has evolved numerous times across the Hymenoptera (Hanna & Abouheif 2021), no other superorganismal hymenopteran has wingless females and no other aculeate lineage has intragenerational wingless and winged female forms. This polyphenism has been suggested to potentiate the evolution of worker variation as wing buds are known to regulate the allometric development of discrete worker castes in *Pheidole* (Rajakumar *et al*. 2018), for example. Accounting for the uncertainty of wingless females †@@@idae **fam. nov.**, our ancestral state estimations favored the inheritance of both eusociality and winged-wingless polyphenism in the most recent common ancestor (MRCA) of the total clade Formicidae, which would seem to buttress the simultaneous transition scenario. However, we recommend caution regarding this interpretation, as lineage turnover is clearly important in the evolution of the Formicoidea (Fig. 17), pointing to the possible instability of intermediate lineages and their certain loss. Notably, the derivation of adaptations for terrestrial locomotion prior to the fixation of eusociality in the Formicoidea does suggest, that the ancestor of the ants went to the ground prior to deriving wing polyphenism (Fig. 21), at least given current knowledge of absence of wingless forms in †@@@idae **fam. nov.**

**Fig. 21.**
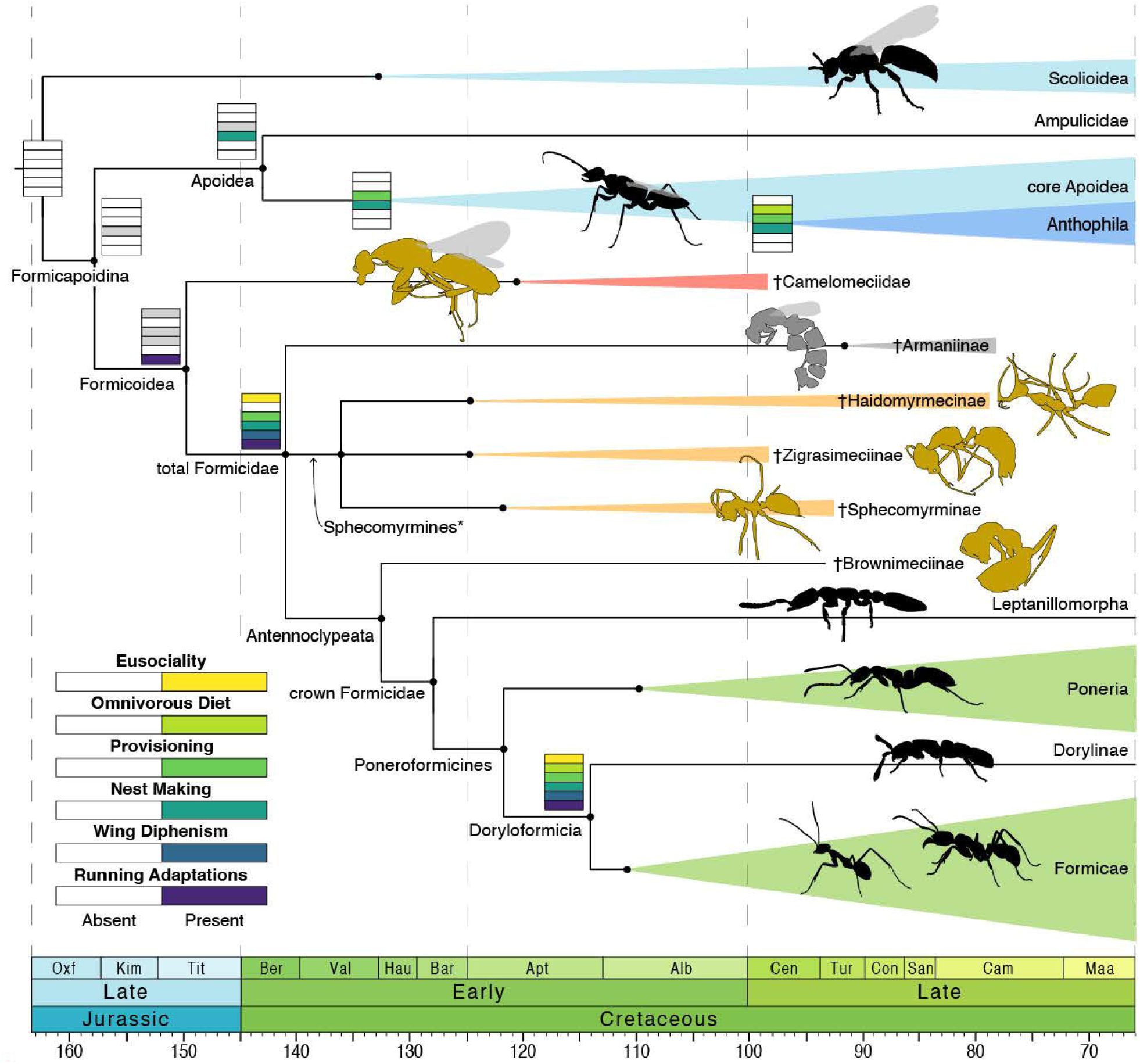
Time-scaled evolution of key traits in the ants and their closest relatives. The origin of obligate eusociality in total Formicidae was preceded by derived locomotory mechanisms of the mesosoma and legs. The best-fitting models for our behavioral data favor simultaneous transition of provisioning and eusociality, with considerable uncertainty about nesting. Differential survival among crown Formicidae based on diet is suggested by the massive diversity of omnivorous Formicae relative to the predatory Poneria and specialist Leptanillomorpha and Dorylinae. Rectangles show reconstructed states; where these are not indicated, no changes are estimated. Black silhouettes mark crown clades, yellow mark amber-preserved stem clades, and grey marks a compression-fossil stem clade. Timescale in millions of years. Sphecomyrmines may be non-monophyletic.

Our results regarding the pathway to eusociality in the Formicoidea are ultimately inconclusive, albeit suggestive of the “ant singularity” scenario. Our study has also explored the application of heterogeneous models for trait evolution while including fossil information for both topology, age, state estimation. Our work indicates several lines of future study. As †@@@idae **fam. nov.** and Formicidae were found to be sistergroups here, it may be possible to extract further information from these fossils for analysis. For example, the mandibles of all camelomeciids are highly derived in form; modeling of female mandibles across the Aculeata may provide circumstantial evidence that †@@@idae **fam. nov.** have a provisioning, hence nesting, lifestyle, if there is indeed a correlation among these traits. Additionally, our modeling experiments suggest a potential solution to the problem of evolutionary singularities (*e.g.*, Uyeda *et al*. 2018): rather than treating lineage diversification as a response variable for a given trait (*e.g.*, the SSE model family, Maddison *et al*. 2007), perhaps jointly simulating trees and focal traits under different models for fossils and extant taxa under the fossilized birth-death process. Finally, it is time to reconsider our conception of the ancestor of the Formicoidea. The overall body forms of †@@@idae **fam. nov.**, †Haidomyrmecinae, and Ampulicidae are strikingly similar; it is possible that the ancestor of the Formicapoidina passed through a temporally short-lived “ampulicid”-like condition that was lost in the Formicoidea and barely retained in the Apoidea. Further morphological study and modeling is necessary. See “Survival to the Cenozoic” in Section 5 below for further considerations.

### Section 5: Evolution of ants and the Aculeata

#### Overview

In this section we will discuss the paleoecological implications of our fossil studies and phylogenetic analyses, while considering uncertainty of geochronological timing (Fig. 9). We recognize three main phases in the evolution of ants in the context of Aculeata: **Phase I**, the origin of the Formicoidea as ground-adapted hunters after successive transitions to passive (parasitic) and aggressive entomophagy (Fig. 22A); **Phase II**, the first radiation of the Formicoidea into increasingly angiosperm-filled ecosystems (Fig. 22B); and **Phase III**, the global collapse of forest ecosystems at the K/Pg boundary and the second radiation of Formicoidea, restricted to those crown Formicidae which survived (Fig. 22C). After outlining these phases in more detail, we discuss the potential factors leading to differential survival of ant lineages through the end-Cretaceous crisis to the Cenozoic. We conclude this section by outlining a transformation series from the ancestor of the Formicoidea into the crown Formicidae, with consideration of potential functional consequences of these modifications.

**Fig. 22.**
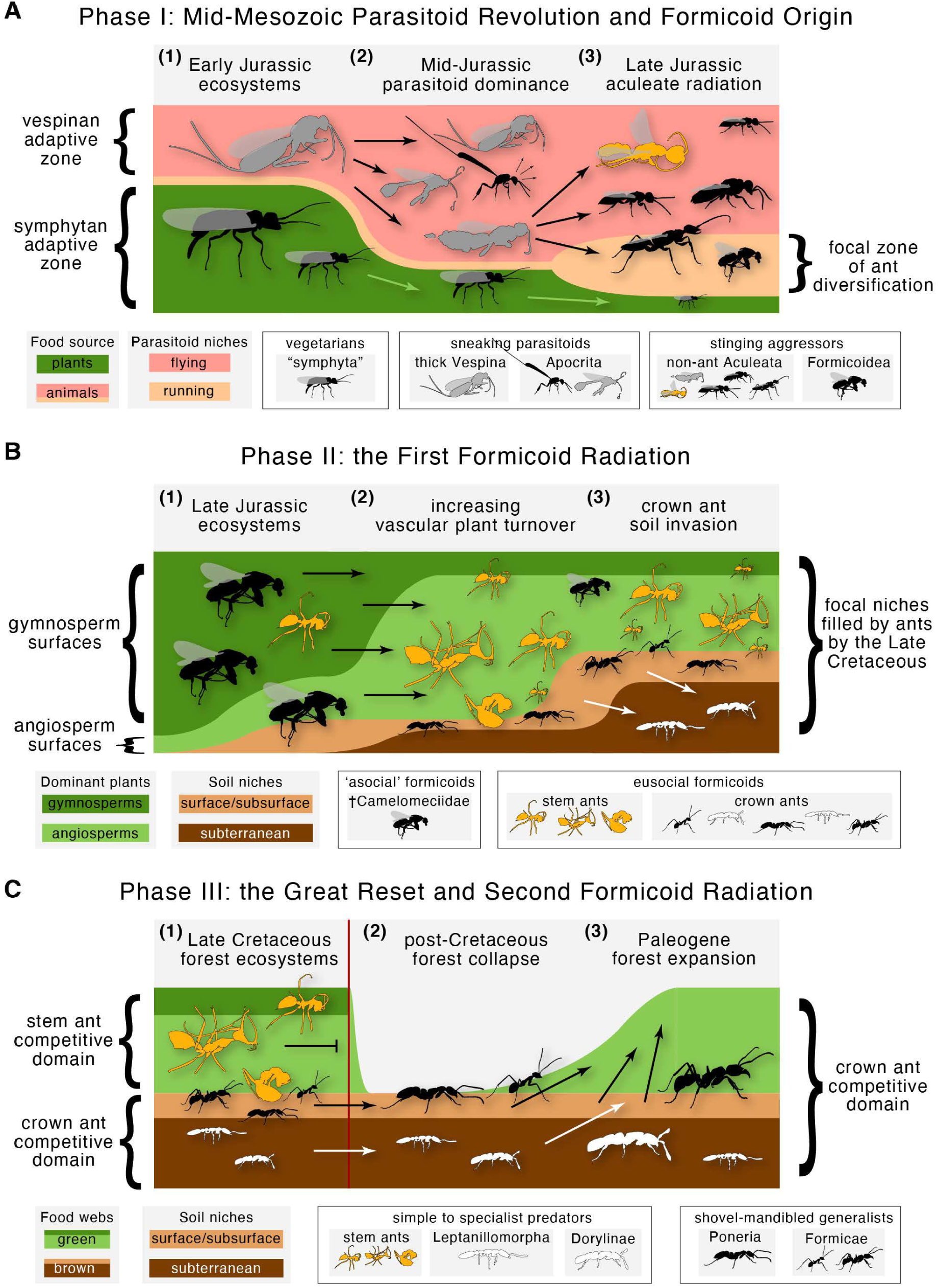
Summary hypothesis for the early aculeate radiation and the origin and evolution of ants. See text for detailed explanation.

#### Outline of ant and aculeate evolution

Based on the fossil record, our topological, dating, and ancestral state estimation results, plus comparison to other studies (Fig. 9), we propose the following narrative hypothesis for the origin and evolution of Aculeata, Formicoidea, and crown Formicidae (Fig. 22). This may be considered an extension to and revision of the hypothesis of Rasnitsyn (1975, 1980, 1988, 2002) for the Hymenoptera and of Wilson & Hölldobler (2005) for the Formicidae.

**Phase I.** The first major event in the post-Permian diversification of the crown Hymenoptera is the dietary transition from phytophagy to entomophagy in the ancestor of the Vespina sometime during the Late Triassic to the Early Jurassic (Figs. 15, 22A.1); by the Late Jurassic, the Vespina had achieved numerical dominance in the fossil record, relative to the “symphytan” grade (Fig. 13). This transition corresponded with wide-scale parallel evolution of parasitism across the Pterygota, potentially resulting in shift from bottom-up to top-down food webs during the first phase of the “mid-Mesozoic parasitoid revolution” (Labandeira & Li 2021). The diversity of life histories among extant Apocrita (Gauld & Bolton 1988) and the Mid-Jurassic preservation of †Bethylonymidae suggests that two guilds had emerged during this revolution: the “sneaking parasitoids” and “stinging aggressors”, with the former encompassing the parasitican grade and the latter the clade Aculeata (Fig. 22A.2).

While there may be no known morphological synapomorphy of the sting apparatus to define Aculeata (see “Notes” for Character 433 in Results Part V), all superfamilies inherited the derived capacity to subdue or pester hosts, prey, or enemies via envenomation; consequently, the “sting” (= terebra, Lieberman *et al*. in press) must be raised for oviposition (Gauld & Bolton 1988), notwithstanding the derived telescoping abdominal tube of Chrysididae. We further hypothesize that this combative approach to host appropriation may have been a preadaptation facilitating the derivation of predatory habits, which have independently evolved multiple times the Aculeata, including Chrysidoidea (Bethylidae), Vespoidea (Vespidae), most Apoidea, and Formicoidea. To our knowledge this specific connection has not been made previously, or at least explicitly. The final step in Phase I was the derivation of thoracicocoxal adaptations for surface locomotion in the predecessor of the Formicoidea from a predominantly flight-dependent ancestor (Figs. 21, 22A.3). Notably, Formicoidea are older than most crown groups of other aculeate lineages which have derived surface-oriented locomotory adaptations (see wingless silhouettes, Figs. 14–16; it is unclear how many wing loss events have occurred in the Bethylidae and when these have arisen).

**Phase II.** After their Late Jurassic origin as terrestrial huntresses among gymnosperm-dominated ecosystems (Figs. 17, 22B.1; Benton *et al*. 2021), the Formicoidea began their morphological and ecological radiation. In the ∼15–20 million year interval from the split of the Formicapoidina into the Apoidea and the Formicoidea, at least one daughter lineage of the latter—the total clade Formicidae—derived obligate eusociality in correlation with, and possibly due to the origin and stabilization of winged-wingless polyphenism in females (Figs. 20A.5–10, 21, 22B.1). Conceivably, long-term maintenance of obligate monogamy in the ancestral formicid, maximizing offspring relatedness, may have been the final selective crux allowing for the evolution of morphologically differentiated worker castes (Boomsma, 2007, 2009; Hughes *et al*., 2008). As evinced by the undescribed ponerine, dolichoderine, and formicine species from Kachin amber included in the present study (Figs. S17–19) and supported by the congruence of divergence dating estimates across studies (Fig. 9), the origins of the major lineages of crown Formicidae had been achieved during the Early Cretaceous.

By the mid Cretaceous, the ant fauna was a mixture of variably specialized stem lineages (Barden *et al*. 2020, Perrichot *et al*. 2020, Boudinot *et al*. 2020), which may have been numerically dominant in or at least experiencing possible preservational bias in the Kachin amber fossil record. It appears that the major cladogenic events of the crown Formicidae closely tracked the transition to obligate eusociality in Dictyoptera (Evangelista *et al*. 2019, although see, *e.g.*, Legendre *et al*. 2015) and the diversification of major extant groups of Angiospermae (Coiro *et al*. 2019) regardless of the age of the crown group (*e.g.*, Budd *et al*. 2021, Silvestro *et al*. 2021). A potential consequence of the Early Cretaceous angiosperm radiation was the increase in palatable leaf litter due to the synchronous evolution of dense leaf veins and other features (Brodribb & Field 2010, Berendse & Scheffer, 2009), which may have facilitated expansion on the brown food web (Kaspari & Yanoviak, 2009) and perhaps increased the concentration of prey. Whatever the sequence, source, or magnitude of food, we hypothesize that by the Late Cretaceous the soil surface and subsurface was populated by crown and stem Formicidae (Fig. 22B.3).

**Phase III.** The final phase of ant evolution considered here is the wide-scale lineage turnover of the Formicoidea presumably caused by the end-Cretaceous extinction event and subsequent second formicoid radiation (Fig. 22C). Although the fossil record of the Late Cretaceous is far too sparse to provide a fine-grained picture of the paleodiversity of Formicoidea in the Ma leading to the crisis (Figs. 9, 11), †Haidomyrmecinae and †Sphecomyrminae were able to persist at least into the Campanian (Barden 2017). The power of the fossil record for distinguishing “turnover” from “mass extinction” and “mass depletion” (*e.g.*, Schachat & Labandeira 2021) is further limited by the scarcity of fossil material from the Paleocene (Barden 2017, Jouault 2021d). However, the Eocene fossil record is densely sampled (LaPolla *et al*. 2013, Barden 2017) and completely lacks stem Formicidae, providing the strongest indication available that †@@@idae **fam. nov.** and the known stem formicid subfamilies were extirpated, including all potential Mesozoic lineages which have not yet been recovered in fossils. To be clear, we recognize that it is possible that the end-Cretaceous extinction crisis may not have been the cause of stem ant extinction, but the pattern of extant and extinct diversity does require consideration. With these caveats in mind, the turnover of formicoid lineages is indeed centered around the K/Pg boundary (Fig. 17), thus we provide evidence-based reasoning for extinction and survival below.

#### Survival to the Cenozoic

To explain the apparent differential survival of lineages across the K/Pg boundary, in addition or as an alternative to the expectation of purely stochastic patterns we propose four hypotheses which are not necessarily mutually exclusive (Fig. 22C): (1) bottom-up perturbation led to extinction of overspecialized lineages, (2) certain specialists survived due to dominance of a stable fundamental niche, (3) collapse of the green food web decimated epigaeic and especially arboreal lineages, and (4) numerical dominance and mid-depth position in ecological networks buttressed other lineages. Hypotheses 1 and 2 may be grouped as “extinction due to specialization”, and 3 and 4 as “survival due to ecological occupation”.

Before continuing, however, it is critical to point out a conundrum: Whereas there are ∼14,000 species of ants alive at the present day (Bolton 2022), there are at least 29,500 species of their sistergroup, the Apoidea (Aguiar *et al*. 2013, Johnson *et al*. 2013). With the evidence of stem ants and the †@@@idae **fam. nov.**, as well as the unequal crown ages of the Formicidae and Apoidea (Fig. 9), it is clear that extinction has played a major role in shaping modern ant biodiversity. Why did ants suffer comparatively more pruning than the Apoidea; what is it about the Apoidea that made them more robust or adaptable over the global biodiversity crisis? This is a question which should be addressed in future study. Note that we employ new taxonomic nomenclature in this section corresponding to our revision of the total clade Aculeata (Results Part III) and defined in the extended transformation series (Results Part IV) (see also Figs. 14– 17).

*1. The saber-tooth hypothesis*. As has long been recognized, overly specialized lineages are prone to extinction (Huxley 1942, Simpson 1944). Based on their highly derived mouthpart and cranial morphologies, it is reasonable to infer that the stem ant subfamilies †Zigrasimeciinae and †Haidomyrmecinae were indeed dietarily specialized, perhaps being apex predators of Cretaceous ecosystems. The postulate of this hypothesis is that the top predator ant fauna was unable to respond by expanding their dietary niche breadth in response to bottom-up trophic network collapse. The hypothesis predicts that: (1) “generalist” lineages of the crown Formicidae should have more surviving nodes in the Cretaceous; (2) “specialist” lineages which survived should have few or no surviving nodes. The pattern of extant lineage diversity across the K/Pg boundary matches these expectations. The Mesozoic diversity of the highly morphologically specialized lineages Leptanillomorpha and Dorylinae is heavily pruned (with the exception of *Martialis* in the former; Borowiec *et al*. 2019), while the more morphologically plesiomorphic or “generalized” Poneria and Formicae retain considerable Cretaceous diversity (Figs. 14, 17; also: Borowiec *et al*. 2019). Further resolution of this question will require discovery of additional fossil Lagerstätte. We do not consider the comparatively less specialized lineages of stem ants such as †*Brownimecia* or †*Gerontoformica* to have suffered extinction to this precise potential cause.

*2. The bunker hypothesis*. We hypothesize that those very highly specialized lineages of crown ants that originated in the Mesozoic were buffered from ecological catastrophe through relative independence from aboveground ecosystems, *i.e.*, they were subterranean. The ecologically relevant concept is that the stark simplification of the global food web divided the surviving lineages of ants into two fundamental nichespaces: the epigaeic (above-ground) and hypogaeic (below-ground). This hypothesis phylogenetically predicts that the specialist lineages which did survive the end-Cretaceous event will have an ancestral state of being hypogaeic. Although we did not specifically evaluate this hypothesis in our study because of uncertainty regarding the fossils, two previous works are available that have tested the pattern using ancestral state reconstruction across extant ants (Lucky *et al*., 2013; Nelson *et al*., 2018), which we will discuss in combination with the next hypotheses due to their continuity of explanation. Notably, Houadria & Menzel (2021) recently found that subterranean (hypogaeic) ants, even those which may forage above ground, were more specialized in their diet relative to epigaeic species.

*3. The burning forest hypothesis*. We hypothesize that ant lineages which were obligately arboreal or dependent on arboreal resources were extinguished due to the End Cretaceous forest-ecosystem collapse (Vajda *et al*., 2014; Field *et al*., 2018). The two pertinent studies, Lucky *et al*., (2013) and Nelson *et al*. (2018) came to different conclusions about the ancestral state of the crown Formicidae but are complementary when framed in the context of the Mesozoic fauna.

Lucky *et al*. (2013) defined seven states for the character of ant ecology (soil only, soil + surface, surface, surface + arboreal, arboreal, arboreal + soil, and all). In their analysis, the ancestor of the crown Formicidae was strictly subterranean, a condition inherited by Leptanillinae, the Poneroformicines, the Poneria, the Doryloformicia, and the Dorylinae; the Formicae, however, were supported as being surface-dwelling. This work provides support for the notion that the subterranean ecology of ants, particularly the various lineages of Poneria, was an important factor in surviving the End Cretaceous catastrophe.

In contrast, Nelson *et al*. (2018) divided the relevant ecological characters into eight states relating to diet (strict predation, herbivory or omnivory), foraging domain (strictly ground-based, arboreal, or ground-arboreal), and nesting domain (strictly in the ground, arboreal or ground-arboreal). They estimated that the most recent common ancestor of the crown Formicidae was strictly predatory, ground-foraging, and ground nesting, with both the Dorylinae and Leptanillinae having these states. Critically, the ancestral poneroformicine in their analysis was capable of foraging above-ground, indicating that the doryline state is derived, and supporting the contention that the obligately subterranean lineage of this group of specialist predators was “bunkered” during the global climate catastrophe of the Cretaceous-Cenozoic transition, while the above-ground relatives were extinguished.

*4. The “bird” and “dinosaur” hypothesis.* Using the Late Cretaceous paleodiversity of Aves—particularly during the Maastrichtian—as a framework (Longrich *et al*. 2011, Field *et al*. 2020), it is possible that the crown lineages of Formicidae attained numerical and ecological dominance relative to stem lineages by the very latest Cretaceous, which would have statistically favored their survival even if extinction were nearly or totally stochastic. This hypothesis predicts that, upon discovery of further Lagerstätte around the K/Pg boundary, crown ants would be either more diverse and/or more abundant than stem ants. Such lineages may have occupied relatively stable mid-depths in the Cretaceous ecosystems, allowing them to adapt to changes in trophic networks thus to expand during the post-Cretaceous recovery (*e.g.*, Field *et al*. 2018, Oliveros *et al*. 2019) and terminal phase of the Angiosperm Terrestrial Revolution (Benton *et al*. 2021). For the sake of speculation, it is possible that crown ants may have gained dominance through derivation of social traits that cannot be evaluated from anatomical data, including more integrated communication, increased foraging efficiency, and improved hygiene. Regardless, a test of this hypothesis cannot be made with the presently available evidence; discovery and analysis of Maastrichtian and Paleocene ant fossil deposits is strictly necessary.

#### Paleomorphological Evolution of the Ants

Because our estimated transformation series (Results Part IV) is the most comprehensive and detailed reconstruction of insect morphological evolution to our knowledge to date, we would like to discuss the potential functional and ecological implications of this series as it pertains to the early evolution of the ants. The restructuring of the body cannot be without some degree of functional consequences, and the stability of these conditions across the Formicoidea requires explanation: Why are the bodies of ants consistently structured the way they are relative to other Hymenoptera? We note that the functional hypotheses provided below are based on mechanical first principles, direct observation of live ants in a lab-based setting, physical manipulation of dead specimens, and experimental and observational evidence from the literature (see the references); they are to be formally tested via comparative kinematic study, simulation, and robotics (*e.g.*, Blanke *et al*. 2017, Nyakatura *et al*. 2019, Büsse *et al*. 2021a, b), which we consider to be a highly desirable goal. For a list of estimated synapomorphies for all sampled formicoid nodes outside of the context of functional hypothesis and interpretation, see Section 1 of Results Part IV. For previous broad-scale hypotheses about the early evolution of the ants synthesizing fossil morphology and phylogeny see Wilson *et al*. (1967a, b), Wilson (1987a, b), Dlussky & Fedoseeva (1988), Hölldobler & Wilson (1990), Appendix 3 of Bolton (2003), Dlussky & Rasnitsyn (2007), Barden & Grimaldi (2016), Perfilieva (2022), and Richter *et al*. (submitted). Note also that we employ taxonomic nomenclature derived from our systematic revision of the ants and the Aculeata (see Results Part III and IV); we disfavor the use of the terms “formicoid”, “poneroid”, or “leptanilloid” for groups within the Formicidae as they imply superfamilial status and can be conflated with “formicoid” in the sense of Formicoidea, as used here.

*1. Formicoidea.* Across all characters and nodes of the Formicoidea, given our present sampling of stem lineages, we detect multiple sets of derived conditions. The first set forms three syndromes likely associated with the Jurassic lifestyle of the formicoid lineage. Based on our estimates, the most recent ancestor of the Formicoidea inherited the following derived syndromes: (1) derivation of the locomotory system, (2) modification of the cranium, mouthparts, and neck, and (3) strengthening of the metasoma. We hypothesize that the modifications of the locomotory system is associated with surface foraging; they are mirrored to various extents by other aculeate lineages with wingless females. With respect to the head and neck modifications, modern ants rely on gripping objects with their mandibles and lifting them for ergonomic functions such as nest building, food carriage, and brood care. Because certain key derivations of the physical structures of the head and neck which define crown ants were inherited by the formicoid ancestor, it is possible that functional changes had already occurred. Finally, the metasomal modifications are likely associated with attacking hosts or prey, as this is a fundamental role of this tagma in living sting-bearing ants.

The morphological derivations likely associated with ground-based locomotion in the ancestor of the Formicoidea are severalfold. With respect to the legs, the fore coxa and trochanter transformed as one unit, and the mid and hind coxae as another. The forecoxa was elongated and the distal foramen was rotated laterally, enclosed ventrally, and completely concealed the proximal portion of the trochanter (Boudinot, 2015; Liu *et al*. 2018) (Fig. 23E, F vs. 23N, O). The trochanter itself was also constricted and bent, potentially maximizing stride length during locomotion (Fig. 23I vs. 23R). In extant ants, one of the two coxotrochanteral condyles is absent, while the remaining condyle is curved ventrally and buttressed dorsally (Fig. 23I). This form may be of value under load carried by the mandibles, with neck strength for the highly mobile head enhanced by the muscular and elongate prothorax (Keller *et al*., 2014; Nguyen *et al*., 2014; Peeters *et al*., 2020). Similarly, the meso- and metathoracic articulations were rotated ventrad (Boudinot, 2015) (Fig. 23E vs. N; Fig. 24), restricting the motion of the legs to the frontal plane while acting as columns supporting the body (Fig. 23F vs. O).

**Fig. 23.**
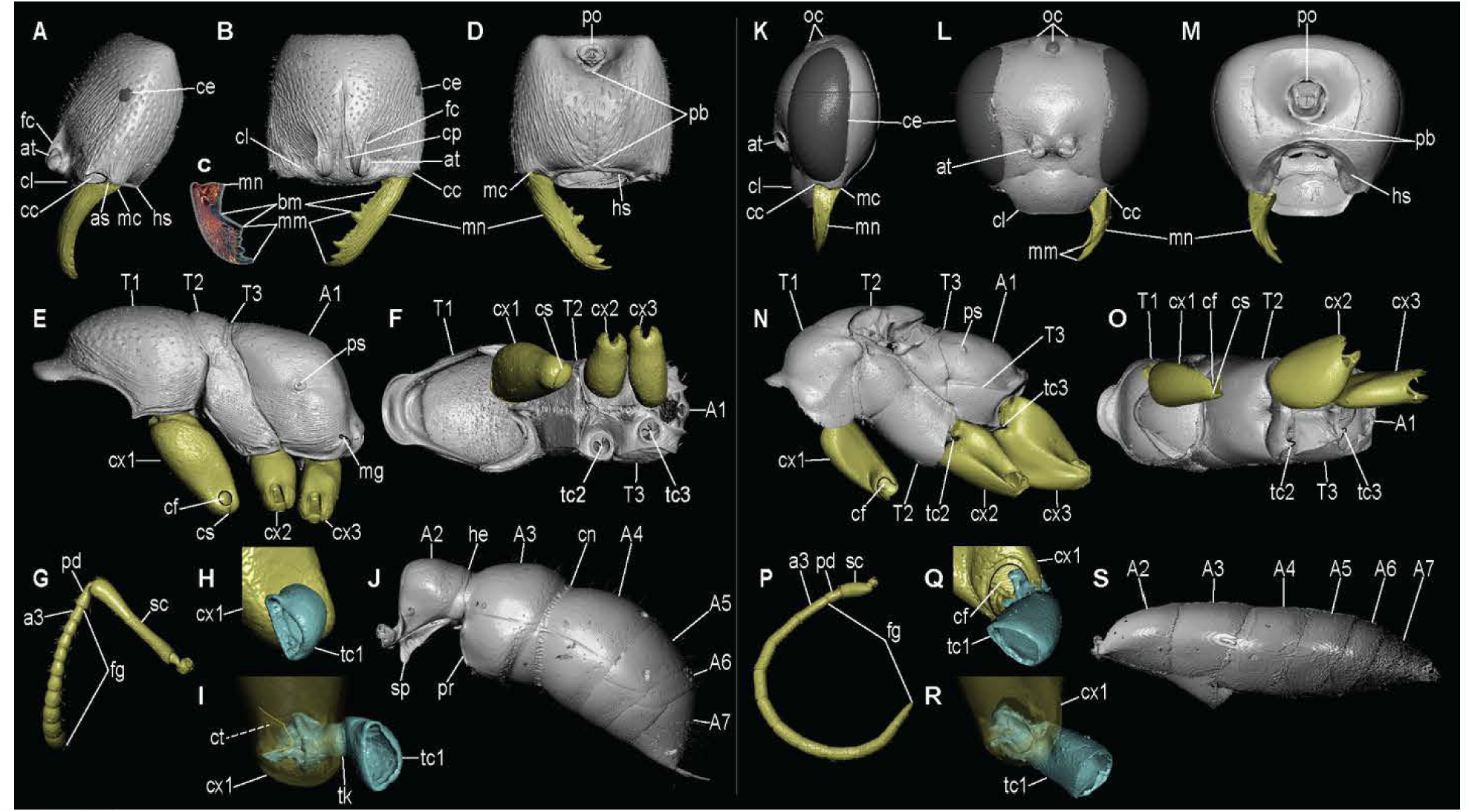
Ant and aculeate morphology, showing formicoid apomorphies. **(A to J)** Formicidae, all of *Amblyopone australis* except **(C)** *Formica integroides.* **(K** to **S)** Pompilidae, *Anoplius* (*Arachnophronctus*) species **(A** to **D, K** to **M)** Crania, **(E, F, N, O)** mesosomata, **(G, P)** antennae, **(H, I, Q, R)** foreleg trochanteral articulations, (J, S) metasomata. Al, propodeum; A2-7, metasomal segments; a3, third antennomere; at, antennal torulus; bm, basal margin; cc, cranial condyle; ce, compound eye; cf, fore coxal foramen; cl, clypeus; cp, posterior extension of clypeus; cs, coxal suture; ct, coxotrochanteral condyle; cx1–3, pro-, meso-, and metacoxae; fc, frontal carina; fg, flagellum; he, helcium; hs, hypostoma; mc, mandibular condyle; mg, metapleural gland; mm, masticatory margin; mn, mandible; oc, ocelli; pb, postgenal bridge; pd, pedicel; po, postocciput; ps, propodeal spiracle; sp, subpetiolar process; pr, prora; T1–3, pro-, meso-, and metathoraxes; tc1, fore trochanter; tc2, 3, meso- and metathoracic condyles; tk, trochanteral constriction.

**Fig. 24.**
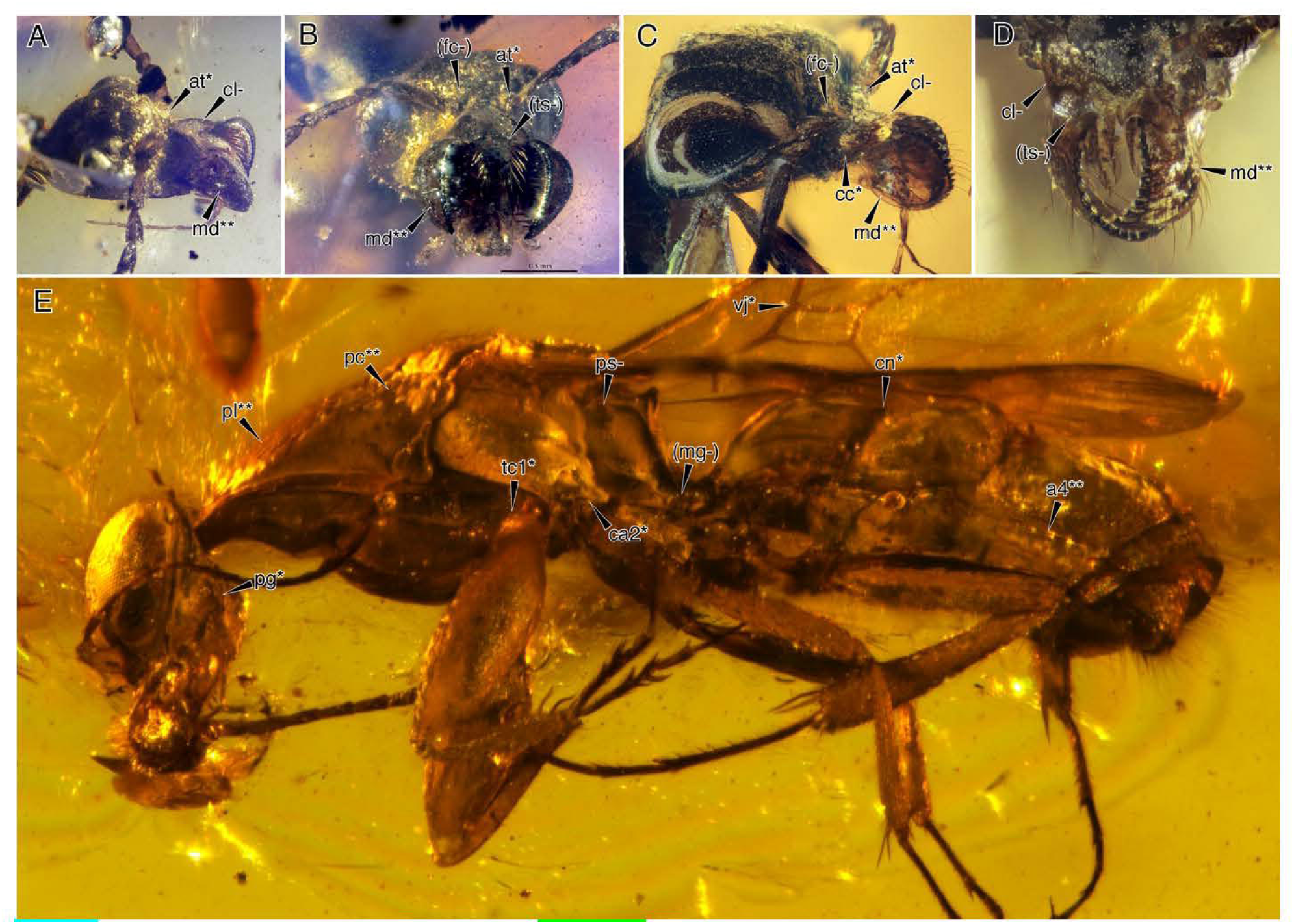
Diagnostic features of †@@@idae fam. nov. No known camelomeciids are wingless, and the social metapleural gland is definitively absent in several species. Not all diagnostic features shown (see also Boudinot *et al.* 2020). **(A, B)** †*Camelomecia janovitzi* holotype in dorsolateral and anterior oblique views. **(C, D)** †*Camelosphecia fossor* holotype (ANTWEB1038930) and **(E)** non-type (BALBuTJ-36) in ventrolateral, anterolateral, and dorsolateral oblique views. *, **, synapomorphies of Formicoidea and †@@@idae, respectively; -, supporting plesiomorphies. **a4\*\***, tubulated fourth abdominal segment; **at\***, partially to completely laterally directed torulus; **ca2\***, closed mesocoxal articulation; **cc\***, enlarged cranial condyle; cl-, strongly projecting clypeus; **cn\***, constricted abdominal segment IV presclerites; **(fc-)**, frontal carinae absent; **(mg-)** metapleural gland absent; **mn\*\***, bowed mandible with elongate masticatory margin; **pc\*\***, posteriorly constricted pronotum; **pl\*\***, elongate pronotum; ps-, high and anterior propodeal spiracle; tc1, torqued protrochanter; **(ts-)**, traction chaetae absent; **vj*,** rsf1 and mf1 meeting at angle. Imagers: Phil Barden (A, B, E), Julio Chaul (C, D) (AntWeb).

Potential foraging-associated adaptations defining the ancestral formicoid involve transformations of the cranium (Fig. 23A, B, D vs. K, L, M; Fig. 24) and metasoma (Fig. 23J vs. 23S; Fig. 24). From a short condition, the cranial postgenal bridge was elongated, thus directing the mouthparts forward (“prognathy”) and allowing the head to lift above the frontal plane; this may have improved the balancing capacity of foragers under load via controlled head movements and the shifting their center of gravity (*e.g.*, Zollikofer 1994; Moll *et al*. 2010). Mandibular gape was potentially modified through enlargement of the dorsal craniomandibular condyle (Richter *et al*., 2019, 2020, 2021, submitted; Boudinot *et al*. 2021; Gronenberg *et al*., 1998; Zhang *et al*., 2020), although the specific biomechanical consequences of this derivation have yet to be explicitly addressed. The range of antennal motion may have also been increased through and rotation of the antennal sockets from a dorsal aspect to a lateral orientation (Richter *et al*. submitted). Likewise, the metasomal range of motion was increased by anterior elongation of the petiole and shrinkage of the anterior articulatory surfaces of the second segment (helcium) into a defined ball-and-socket joint (Dlussky & Fedoseeva, 1988). Three further modifications are reasonably inferred to have maximized the capacity for force transfer during stinging as well as the safety factor, *i.e.*, the failure rate of strenuous actions may have been improved. Development of a petiolar node may have increased metasomal strength (Hashimoto, 1996), while the petiolar sternum gained an anteroventral process which prevents shearing by locking between the hind coxae, and the helcium was buttressed by gain of an anteroventral keel (prora) just posterior to the articulatory sternite.

*2. Formicidae.* Although there is no question that other, closely related extinct lineages are not yet known, we recover several morphological synapomorphies coinciding with the origin of eusociality in the ancestor of the total Formicidae. Most importantly, according to contemporary thinking, are gain of the metapleural gland (Fig. 23E vs. 23N; Fig. 25C; Hölldobler & Wilson 1990) and optimization of the worker’s mesosoma into a flightless power core for surface foraging (see Peeters *et al*., 2020 for discussion of the “power core” notion and hypothesis). The propodeal spiracle to a low and lateral position above the legs, from the high and forward position just posterad the hind wing base (Fig. 23E vs. N), suggesting change in the tracheation of the mesosoma; some stem Formicidae have the spiracle situated near the dorsal margin of the propodeum at about propodeal midlength (*e.g.*, Fig. 25C). Association with soil and soil-surface habitats (*e.g.*, Nelson *et al*., 2018, Lucky *et al*., 2013) is supported by reduction in worker and queen compound eye size to < ½ head length. Metasomal mobility was further increased via pronounced narrowing of the helcium, resulting complete petiolation of the first metasomal segment (Fig. 23J vs. S). Likely associated with protection of the antennal bases, the contact surface of the cranium developed medial longitudinal facial rims (frontal carinae) (Fig. 23A, B vs. K, L), which may also reinforce the cranium against longitudinal compression caused by mandibular closure. Small colony size (Burchill & Moreau, 2016) and solitary foraging (Reeves & Moreau, 2019) are reasonably expected conditions of the total Formicidae ancestor. It is possible that brood transport may be a defining behavioral synapomorphy of the total clade Formicidae (Boudinot *et al*. 2022).

**Fig. 25.**
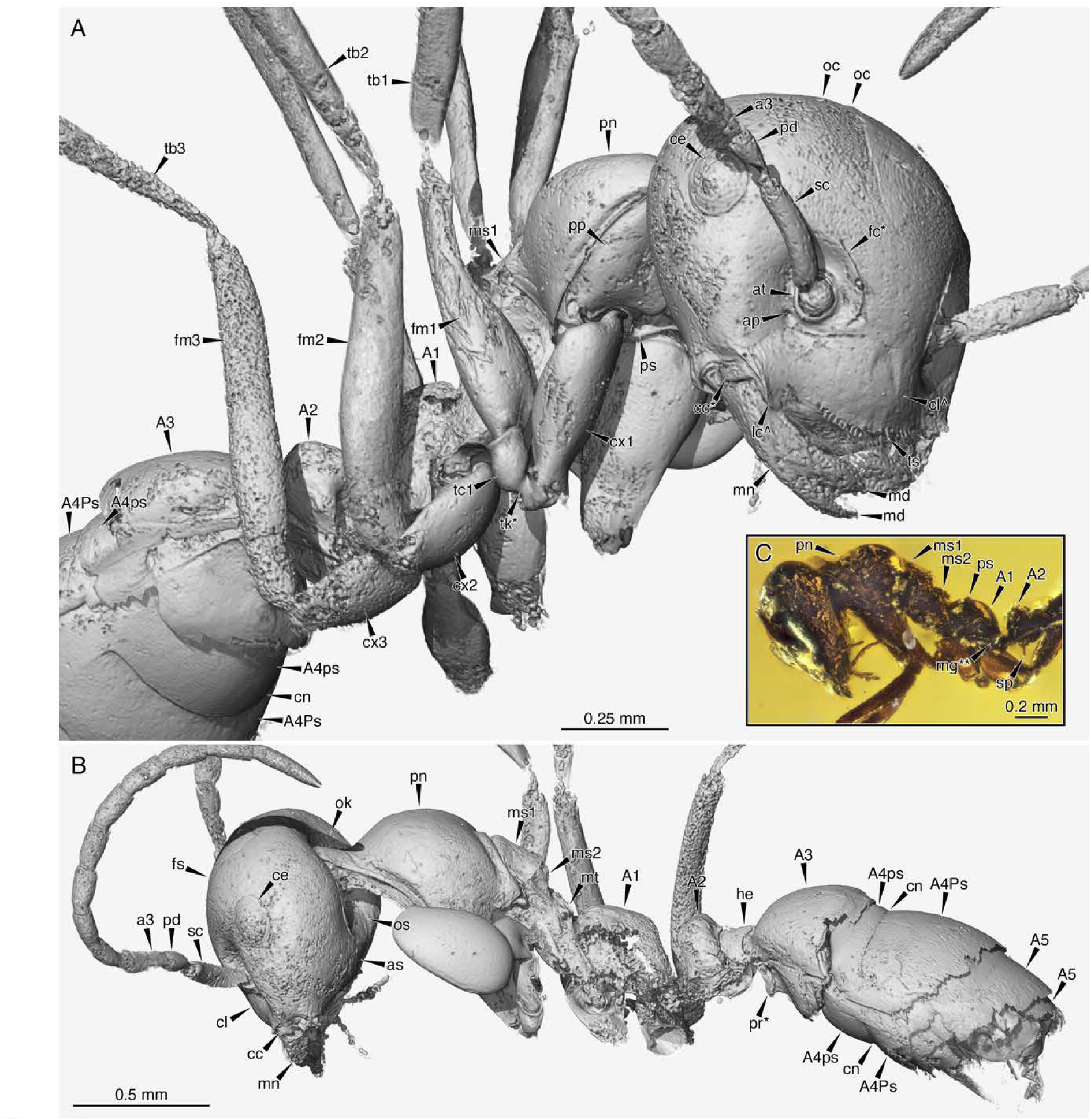
Worker morphology of least-specialized stem Formicidae. †*Gerontoformica* species near *pilosa,* CASENT0811296 **(A, B)** and UFV-LABECOL-009656 **(C). (A)** Body in anteroventral oblique view. **(B)** Body in anterodorsal oblique view. **c,** Cranium, mesosoma, and petiole in profile view. *, **, synapomorphies of Formicoidea and Formicidae, respectively; ^, sphecomyrmine clade apomorphies; **A1,** propodeum; **A2-7**, metasomal segments; A4Ps, abdominal segment IV postsclerite; A4ps, abdominal segment IV presclerite; **a3,** third antennomere; **ap,** anterior tentorial pit; **at,** antennal torulus, **cc,** cranial condyle; **ce,** compound eye; **cl,** clypeus; **cn,** cinctus; **cx1-3,** fore, meso-, and metacoxae; fc, frontal carinae; he, helcium; mg, metapleural gland; **lc,** lateral clypeal lobe; **md,** mandibular teeth; **mn,** mandible; **ms1,** mesoscutum; **ms2,** mesoscutellum; **mt,** metanotum; **oc,** ocelli; **ok,** occipital carina; **os,** occipital contact surface; **pd,** pedicel; **pn,** pronotum; **pr,** prora; **ps,** propodeal spiracle; **sp,** subpetiolar process; **tk,** trochanteral constriction; **ts,** clypeal “traction setae” (chaetae).

3. *Antennoclypeata.* Modifications along the formicid-anntennoclypeatan stem involve simplification of the mesosomal organization in workers (Barden & Grimaldi, 2016), perhaps reflecting skeletomuscular optimization and a decrease in energetic expense of workers (Peeters *et al*., 2020; Peeters & Ito, 2015), plus modifications of the antenna and clypeus possibly resulting in enhanced fine motor control. Specifically, the clypeus is extended posteriorly between the antennal sockets, altering the conformation of extrinsic scapal and pharyngeal muscles (Richter *et al*., 2020), while the antennal scape became elongated and the flagellum shortened. Along with proximal compression and curvature of the pedicel (Richter *et al*. submitted), elongation of the scape produces the typical “geniculate” form of the ant antenna. This was previously hypothesized to improve worker-brood interaction through allowing the focal surfaces of the flagellae to be placed near the mouthparts (Dlussky & Fedoseeva, 1988). We further hypothesize that by separating the flagellar bases widely through scape elongation, worker ants may more easily center themselves along a pheromone trail through controlled sampling of the ground by the odorant sensitive surfaces of the left and right antennae (see, *e.g.*, Draft *et al*. 2018). In other words, scape elongation may indicate a transition from the use of behavioral strategies of aerial odorant reception to surface-based olfaction, and perhaps to physical pheromone trail laying. If this were indeed true, our estimates support multiple transitions to this surface-based trail laying strategy, with scape elongation also occurring in some †Haidomyrmecinae and the †Zigrasimeciinae. Whether short-scaped †*Gerontoformica* were capable of sophisticated trail following is unknown, but it may be possible to evaluate their glandular array and that of other stem ants via µ-CT (*e.g.*, Boudinot *et al*. 2022b; Richter *et al*. submitted). It is also plausible that the expansion of odorant binding receptor genes (McKenzie *et al*., 2016) and diversification of antennal sensilla may have also occurred along this stem, although such an event may have occurred earlier.

4. *Crown Formicidae.* The final derivations inherited by the ancestor of crown Formicidae are repression of ocelli in workers (Fig. 23A, B vs. K, L) (see Character 40 of crown clade Formicidae diagnosis in Results Part IV), reinforcement of the metasoma via petiolar tergosternal fusion, and perhaps most importantly, increased tooth armament of the mandible, enhancing grip (Fig. 23C vs. L; for another discussion, see Perfilieva 2022). Whereas the stem ants plesiomorphically have bidentate and more-or-less strap-shaped mandibles, crown ants display a wide range of mandibular forms and are synapomorphically defined, in part, by additional teeth.

The pattern of tooth distribution within crown ants is distinct between the depauperate leptanillomorphan and massive poneroformicines clades. In the former, representing a heterogeneous assemblage of specialized predators, the additional teeth occur on the basal mandibular margin, improving grip when the mandibles clasp objects to the clypeus. However, several derived mandibular forms have evolved in the leptanillomorphs, mirroring the poneroformicines in small scale. In poneroformicines, the additional teeth occur on the elongate masticatory or tool-edge margin, a conformation which allows objects to be precisely grasped between the mandibles, corresponding to the traditional “elongate mandibular margin” trait previously hypothesized by Wilson (1987a) to define all living ants and to improve foraging and other social tasks.

We further define the “triangular” form of the crown ant mandible as “shovel-shaped” (Richter *et al*. submitted), as the mandibles are not only capable of precise grip, but also to be used as shearing tools for digging. Given the critical role of the nest in regulating social interaction, colonial ergonomics, and development (*e.g.*, Tschinkel 2013, 2021), the effects of mandibular shape variation should be critically explored. Among lineages of the total Formicidae, retention of and reversal to short masticatory margins is associated with development of perioral traction chaetae, as observed in †Sphecomyrmines, Apomyrminae, Amblyoponinae, and some Leptanillinae (Fig. 22C; see Boudinot *et al*. 2020, 2021 for differentiation and discussion of “chaetae” and “setae”). That Cenozoic survival of crown ants was dependent on a combination of traits, not solely dentition, is evinced by †@@@idae **fam. nov.** which convergently gained elongate mandibular grasping margins (Fig. 24E).

### Section 6: Final Considerations and Future Directions

The present study represents a systematic revision of the Aculeata and kin which emphasizes the evolution of the Ants (Formicoidea) and integrates morphological and molecular data following recent advances in and results of phylogenetic modeling (*e.g.*, Pagel & Meade 2006, Ronquist *et al*. 2012, Beaulieu *et al*. 2013, Branstetter *et al*. 2017a, b). Our work provides a review of the Mesozoic fossil record of the Hymenoptera, a new system for aculeate fossils, introduces a new sistergroup of the Formicidae, and uses large-scale ancestral state estimation (ASE) to infer the evolutionary sequence of the ants and other Aculeata. The ASE for the morphological data appears to be a new approach in contrast to our guide studies as we employ the same models for ASE as were used for estimating our tip-dated phylogenies. Further, our ASE for the behavioral data builds on and expands the use of complex transition models, such as our covarion-like model, while also implementing reversible jump MCMC. We would like to conclude Results Part I with some final considerations and a brief outline of future directions for study. See Sections 1–5 of Results Part I for topic-specific conclusions, discussions, and hypotheses; see Part II for our extended topological results, Part III for a taxonomic synopsis of sampled groups, Part IV for the extended transformation series, and Part V for character and state definitions and illustrations.

The original impetus for this work was dual: Intent to update the morphological classification of the total clade Formicidae based on the integration of phenotypic and genotypic data, and the shaking of our assumptions about the groundplan of the ants on one hand by the discovery of the Formicapoidina node (Johnson *et al*. 2013) and, on the other, by a profound sense of consternating enigma arising from the specimens we classify here as †@@@idae **fam. nov.** The strangeness of these fossils continues to halt us in our confidence about the evolution and relationships of Formicoidea and other Aculeata, but we have pushed our data as far as we could in the present work. Surely further Mesozoic Formicoidea will be discovered, which inform and revise our collective hypotheses for the evolution of the ants. It is certain that application of µ- CT technology to study stem ant anatomy will shed light on these questions (*e.g.*, Boudinot *et al*. 2022, Richter *et al*. submitted). We further recognize that the traditional manual scoring of morphological data is a fallible subjective process, for which a key solution will be the development of analytical techniques for the quantitative and three-dimensional data derived from µ-CT scanning.

We hope that future study will make headway with this dataset and its idiosyncrasies, particularly as there are many ways of dissecting the matrix: How should morphological data be partitioned (*e.g.*, Tarasov & Génier 2015, Caldas & Schrago 2018, Rosa *et al*. 2019, Porto *et al*. 2021)? Will hierarchical character scoring and/or hidden state matrix approaches improve the modeling of phenotypic data and provide a pathway for complex trait-based phylogenetic comparative methods (*e.g.*, Tarasov 2019, Hopkins & St. John 2021; also: Uyeda *et al*. 2018)? We specifically wonder if the combined analysis of continuous and discrete phenotypic data will improve phylogenetic estimation, *i.e.*, the treatment of developmental characters (discrete on/off, present/absent variables) as “observed” hidden rate matrices for developmental states (continuous 1–3D variation); continuous data have already been shown to be more information rich, in theory, than discrete characters (Parins-Fukuchi 2018). Additionally, considering the difficulty posed by turnover, it may perhaps be worthwhile to jointly simulate of topology and traits under the fossilized birth-death process.

If parsimony analysis of these data can be performed on the present dataset, how will these results compare to searches under a clock model (Goloboff *et al*. 2019, Mongiardino Koch *et al*. 2021)? Given the limited practical timescale of the actual work presented herein, what will terminal-by-terminal RoguePlot methods reveal about fossil placement for key taxa (Klopfstein & Spasojevic 2019), particularly the ancient and wing-based earliest vespiform Aculeata, and how do they influence date estimates (Luo *et al*. 2020, Spasojevic *et al*. 2021)? How will the dating analyses behave without the age constraints that we implemented? Will refined methods improve estimation of synapomorphies along the backbone of the Aculeata; would “pseudoextinction” or “hypothetical ancestor” approaches provide any insight (*e.g.*, Mongiardino Koch & Parry 2020, Brady & Springer 2021)? These and numerous other methodological and empirical questions remain, not the least of which include resolving conflicts over fossil placement between this study and others. We will be satisfied to see further resolution of the paleomorphological evolution, systematics, and classification of the ants, other Aculeata, and the Hymenoptera, even if some or many of our results are overturned.

#### Part II: Extended Topological Results

### Overview

In order to evaluate placement of the sampled fossils and to reconstruct the transformation leading crownward into the Formicidae, we evaluated a comprehensive sample of Aculeata and three outgroups, Trigonaloidea, Evaniomorpha, and †Ephialtitoidea. Outgroup sampling was intended to be dense in order to improve character polarization near the root of the Aculeata. Because this is the first study to estimate the phylogeny of total clade Aculeata, we describe morphology-only results from our dataset under unconstrained ML analysis (IQ-Tree 2, Minh *et al*. 2020) and soft-constrained (“molecular scaffolded”), clockless Bayesian analysis (MrBayes 3.6.2, Ronquist *et al*. 2012). Both ML and Bayesian analyses were stratified by data completeness. Support values for ML are represented by ultrafast bootstrap (UFB, Hoang *et al*. 2018) and those for Bayesian are posterior probabilities (BPP). Support values are lower in the ML analyses because constraints were not implemented. Therefore, the Bayesian analyses with the “soft” constraint molecular scaffold provide greater sensitivity for fossil placement when extant taxa are part of the group. Treatments in this section begin with outgroups (†Ephialtitoidea, Evanioidea) and proceed through aculeate ingroups into the Formicoidea.

The most important factor we found influencing phylogenetic analysis quality was data completeness, probably due to the fact that less-complete terminals were less likely to have preserved syn- or autapomorphies. Support values decreased for with increasing data incompleteness. The major distinction in our total matrix was between the two primary fossil classes, amber (179 count, 63% average completeness) and compression (252 count, 29% average completeness). The most complete fossil was an undescribed genus of Sierolomorphidae from burmite (90.1% complete); no other fossil was > 90% complete. The most complete compression fossils are between 50–59%. Completeness among amber and compression fossil terminals is reported in Fig. 7. Matrix abbreviations used below are based on completeness strata, for example, “90%C” = 90% complete matrix, *i.e.*, that matrix excluding all terminals which are < 90% complete. See XXX on the online repository Zenodo for tree files (doi: XXX). See also Figs. S1–17 for supplementary figures depicting undescribed taxa from Kachin amber used in the present study.

### Section 1. Outgroup topologies

**† Ephialtitoidea Handlirsch, 1906**. Rooting is a crucial issue for †Ephialtitoidea, which must be addressed with comprehensive sampling of all Apocrita as well as Unicalcarida. Given the current sampling, †*Kuafua* Rasnitsyn & Zhang, 2010 is robustly supported as either sister to a clade comprising most †Ephialtitoidea, or sister to the Evanioidea + Trigonaloidea. Other †Kuafuidae vary in placement. Considering the rooting problem, ephitaltitoid support is here reported with respect to bifurcations. All †Ephialtitoidea are represented by compression fossils.

• **† Ephialtitidae** Handlirsch, 1906. Support for an ephialtitoid bifurcation is strong in analyses with relatively high data completeness (40%C: 1.0 BPP, 97 UFB), but decays rapidly with the addition of less-complete taxa (30%C: 0.42 BPP, 52 UFB). There is limited support for a bifurcation separating the two subfamilies, †Ephialtitinae and †Symphytopterinae. One kuafuid terminal, †*Leptogastrella* Rasnitsyn, 1975 is supported within †Symphytopterinae. The unplaced Early Jurassic species †*Kotaphialtites frankmortoni* Rasnitsyn, 2008 clusters within †Symphytopterinae under the soft-constrained Bayesian conditions but is unplaced in unconstrained ML analysis.
•• **† Ephialtitidae: †Ephialtitinae** Handlirsch, 1906. Most †Ephialtitinae cluster together, often including †Symphytopterinae as an internal bifurcation. Supported bifurcations include: †*Praeproapocritus* Rasnitsyn & Zhang, 2010 as a clade (40%C: 1.0 BPP, 97 UFB), †*Proapocritus* Rasnitsyn, 1975 plus †*Stephanogaster* Rasnitsyn, 1975 (40%C: 0.90 BPP, 62 UFB; 30%C: 0.95 BPP, 54 UFB), †*Proephialtitia* Li *et al*., 2015 as clade (30%C: 1.0 BPP); †*Sessiliventer* Rasnitsyn, 1975 as a clade (40%C: 1.0 BPP, 100 UFB, 30%C: 0.71 BPP, 97 UFB), †*Crephanogaster* Rasnitsyn, 1990 plus †*Acephialtitia* Li *et al*., 2015 (30%C: 0.78 BPP, 94 UFB). Neither †*Asiephialtites* Rasnitsyn, 1975 nor †*Leptephialtites* Rasnitsyn, 1975 are supported as monophyletic. †*Trigonalopterus* Rasnitsyn, 1975 clusters near the ephialtitoid root, near †*Karataus hispanicus* and †*Kuafua*. The oldest fossils, †*Liadobracona* Zessin, 1981, †*Proapocritus praecursor* Rasnitsyn, 1975, and †*Thilopterus* Rasnitsyn *et al*., 2003, are Early Jurassic. †*Liadobracona* is unplaceable, often being recovered within the Aculeata, which is unreasonable. †*Proapocritus praecursor* is recovered in a bifurcation with †*Proapocritus atropus* Rasnitsyn & Zhang, 2010 under the Bayesian conditions (0%C: 0.79 BPP), but is unplaceable without the soft constraints, *i.e.*, in the present ML analysis. Finally, †*Thilopterus* is recovered in a bifurcation with †*Proapocritus elegans* Rasnitsyn & Zhang, 2010 and †*P. longantennatus* Rasnitsyn & Zhang, 2010 (0%C: 0.98 BPP0. Other ephialtitine genera, represented by single terminals in the present sampling, cluster robustly within the subfamily.
•• **† Ephialtitidae: †Symphytopterinae** Rasnitsyn, 1980. Bifurcation supported with the exclusion of †*Brigittepterus* Rasnitsyn *et al*., 2003 and †*Karataus hispanicus* Rasnitsyn & Martínez-Delclòs, 2000, and with inclusion of the kuafuid †*Leptogastrella* (40%C: 1.0 BPP, 97 UFB; 30%C: 0.63 BPP, 50 UFB; 20%C: 91 UFB; 10%C: 0.38 BPP, 75 UFB; 0%C: 0.54 BPP, 67 UFB). †*Brigittepterus* recovered within a bifurcation of the †Ephialtitinae. †*Karataus hispanicus* often spits near the †Ephialtitoidea–all-other-taxa bifurcation, whereas the remaining †*Karataus* receive some support as a clade (*e.g.*, 30%C: 0.74 BPP), but may include †*Karataviola* Rasnitsyn, 1975 (*e.g.*, 30%C: 53 UFB). The ephialtitine †*Tuphephialtites* Zhang *et al*., 2002 clusters with †Symphytopterinae, but is otherwise unstable.
• **† Kuafuidae** Rasnitsyn & Zhang, 2010. Never recovered as monophyletic. †*Kuafua* and †*Arthrogaster* Rasnitsyn, 1975 often cluster close to the †Ephialtitoidea root bifurcation, while †*Leptogastrella* nests within the †Symphytopterinae bifurcation.

**Evanioidea Latreille, 1802**. [Equivalent to Evaniomorpha Rasnitsyn, 1988.] Strongly supported as monophyletic at the 50% and 40% complete levels (50%C: 1.0 BPP, 89 UFB; 40%C: 0.95C, 59 UFB), with the exclusion of †Andreneliidae and †*Archaulacus probus* Li *et al*., 2014, two taxa which cluster with Trigonaloidea (see below). Most †Praeaulacidae are < 40% complete, and completely decay backbone signal. Note that Evanioidea phylogeny has been previously evaluated in a Bayesian framework by Li *et al*. (2018); inclusion of †Othniodellithidae and †*Nevania* (†Nevaniinae, †Praeaulacidae) in crown Evanioidea and exclusion of †Andreneliidae here are key differences from the prior study.

• **Crown Evanioidea**. Comprises Aulacidae, Evaniidae, Gasteruptiidae, †Baissidae, †Othniodellithidae, and possibly †Anomopterellidae and †Nevaniinae (†Praeaulacidae).
•• **Aulacidae** Jurine, 1807. Not monophyletic: **†Hyptiogastritinae** Engel, 2016 sister to gasteruptiid *Gasteruption kirbii* (Westwood, 1851) (50%C: 0.68 BPP; 40%C: 0.75 BPP). **Aulacinae** Jurine, 1807 represented solely by *Pristaulacus stigmaterus* (Cresson, 1864).
•• **Evaniidae** Latreille, 1802. Maximally supported as clade (50%C, 40%C: 1.0 BPP) to the exclusion of †*Habraulacus zhaoi* Li *et al*., 2015 and the clade †*Palaeosyncrasis hongi* Poinar, 2019 and †*Mesevania swinhoei* Basibuyuk & Rasnitsyn, 2000, the three of which strongly cluster with †Othniodellithidae + Evaniidae. The five extant species were chosen to represent a diversified sample based on Sharonowski *et al*. (2019) and are maximally supported as a clade (50%C, 40%C: 1.0 BPP). †*Cretevania* Rasnitsyn, 1975 is monophyletic (50%C, 40%C: 1.0 BPP). †*Lebanevania azari* Basibuyuk & Rasnitsyn, 2002 is in a basal polytomy of Evaniidae with †*Cretevania* and a clade which includes all other Evaniidae (50%C: 1.0 BPP; 40%C: 0.98 BPP; 30%C: 0.76 BPP; 10%C: 0.80 BPP). †*Newjersevania* Basibuyuk *et al*., 2000 is sister to or paraphyletic with respect to crown Evaniidae (Evaniidae + †*Newjersevania*: 40%C: 0.91 BPP; 30%C: 0.85 BPP; 10%C: 0.82 BPP). Other Evaniidae include †*Botsvania* Rasnitsyn & Brothers, 2007, †*Curtevania* Li *et al*., 2018, †*Grimaldivania* Basibuyuk *et al*., 2000, †*Protoparevania* Deans, 2004, †*Sinuevania* Li *et al*., 2018, and †*Sorellevania* Engel, 2006.
•• **Gasteruptiidae** Ashmead, 1900. Monophyletic with inclusion of **†Hyptiogastritinae** (see Aulacidae above for support). Also including: **Gasteruptiinae** Ashmead, 1900, †**Hypselogastriinae** Engel, 2016, and **†Kotujellitinae** Rasnitsyn, 1975. †Hypselogastriinae is polyphyletic: †*Hyptiogastrites* Cockerell, 1917 forming clade with Gasteruptiidae in part (50%C: 0.70 BPP; 40%C: 0.81 BPP), whereas †*Archeofoenus* Engel, 2017 forming clade (40%C: 1.0 BPP, 76 UFB) with two genera unplaced to subfamily, †*Exilaulacus* Li *et al*., 2018 and †*Electrofoenops* Engel, 2017. †Kotujellitinae (comprising †*Kotujellites* Rasnitsyn, 1990 and †*Kotujisca* Rasnitsyn, 1991) is not supported as a clade in any analysis.
•• **† Baissidae** Rasnitsyn, 1975. Paraphyletic with respect to Gasteruptiidae + Aulacidae (30%C supports for paraphyletic splits: 0.81, 0.76, 0.98 BPP). The most complete two †*Manlaya* Rasnitsyn, 1980 species cluster together (30%C: 0.47 BPP; †*M. flexuosa* Ren, 1995, †*M. lacubrua* Rasnitsyn & Ansorge, 2000), but the other species fall out in the †Anomopterellidae + BGA clade variably. †*Humiryssus* Lin, 1980 species form a clade. †*Tillywhimia* Rasnitsyn & Jarzembowski, 1998 are strongly to maximally supported as a clade, and cluster with †Baissidae + Aulacidae + Gasteruptiidae (10%C: 0.59 BPP; 0%C: 0.58 BPP). Other genera in the present matrix are represented by single terminals (†*Baissa* Rasnitsyn, 1975, †*Electrobaissa* Engel, 2013, †*Heterobaissa* Li *et al*., 2018, †*Mesepipolaea* Zhang & Rasnitsyn, 2004). †*Electrobaissa* is strongly supported in a clade with (BPP nearly maximal in all analyses), and as sister to Aulacidae and Gasteruptiidae (40%C: 0.65 BPP; 30%C, 10%C: 1.0 BPP).
•• **† Othniodellithidae** Engel & Huang, 2017. Monophyletic (60%C, 50%C, 40%C, 30%C: 1.0 BPP). In clade with (60%C: 0.97 BPP; 50%C: 0.72 BPP; 40%C: 0.91; 30%C: 0.80) and sister to Evaniidae (60%C, 50%C, 40%C: 1.0 BPP; 30%C: 0.99 BPP).
•• **† Anomopterellidae** Rasnitsyn, 1975 (40%C: 1.0 BPP). Monophyletic (40%C, 30%C, 10%C: 1.0 BPP). Forming clade with †Baissidae + Gasteruptiidae + Aulacidae (“BGA clade”) (40%C: 0.87 BPP; 30%C: 1.0 BPP). Support in the less-complete matrices (*e.g.*, 30%C) drops precipitously because of †Praeaulacidae, most species of which cannot be placed. Sister to †Baissidae + Gasteruptiidae + Aulacidae (40%C for BGA: 1.0 BPP). †*Anomopterella* Rasnitsyn, 1975 strongly supported as paraphyletic with respect to †*Choristopterella* Li *et al*., 2013 (30%C: 0.97 BPP), and probably paraphyletic with respect to †*Synaphopterella* Li *et al*., 2013 (10%C: 0.69 BPP).
•• **† Praeaulacidae: †Nevaniinae** Zhang & Rasnitsyn, 2007. With the exclusion of †*Eonevania robusta* Rasnitsyn & Zhang, 2010 monophyletic (40%C, 30%C, 10%C: 1.0 BPP); †*Eonevania* under ML forms clade with “praeaulacines” †*Sinevania* Rasnitsyn & Zhang, 2010 and †*Sinaulacogastrinus* Zhang & Rasnitsyn, 2008 (20%C: 81 UFB; 10%C: 76 UFB). †Nevaniinae in clade with Evaniidae + †Othniodellithidae + †Anomopterellidae + BGA clade (50%C: 1.0 BPP; 40%C: 0.96 BPP).
• **Stem Evanioidea**. Comprises taxa of †Praeaulacinae and †Cretocleistogastrinae. Likely not including †Nevaniinae *sensu stricto* (†*Nevania robusta* strongly [90 UFB] to maximally [1.0 BPP] supported as member of crown Evanioidea in 50%C analysis).
•• **† Praeaulacidae** Rasnitsyn, 1972. Never monophyletic. †*Archaulacus* Li *et al*., 2014 (†Praeaulacinae) strongly clusters with Trigonaloidea–Aculeata (40%C: 0.94 BPP, 80 UFB); the metasoma of this species is not distinctly dorsally-located on the propodeum, and the second abdominal segment is broad. †*Gulgonga* Oberprieler *et al*., 2012 clusters with †*Archaulacus* outside Evanioidea. †*Eonevania robusta* Rasnitsyn & Zhang, 2010 strongly clusters with †Ephialtitoidea, particularly †*Montsecephialtites zherikhini* Rasnitsyn & Martínez-Delclòs, 2000 (40%C: 0.89 BPP), although placement is more variable in ML analysis. All other species cluster on the stem of the Evanioidea, with limited structuring.
•• **† Praeaulacidae: †Praeaulacinae** Rasnitsyn 1972. Never monophyletic. †*Palaeosyncrasis hongi* Poinar, 2019 and †*Habraulacus zhaoi* Li *et al*., 2015 are strongly supported within crown Evanioidea, the latter unstable in position and the former maximally supported as sister to or conspecific with †*Mesevania swinhoei* Basibuyuk & Rasnitsyn, 2000. †*Sinevania* Rasnitsyn & Zhang, 2010 and †*Sinaulacogastrinus* Zhang & Rasnitsyn, 2008 consistently forming a clade (20%C: 81 UFB; 10%C, 0%C: 0.97 BPP); †*Eonevania* part of this clade in ML analyses, but distant in Bayesian. †*Evanigaster* Rasnitsyn, 1972 and †*Evaniops* Rasnitsyn, 1972 form a highly supported clade (30%C: 0.96 BPP, 81 UFB; 20%C: 0.83; 10%C: 0.97 BPP, 87 UFB; 0%C: 0.96 BPP, 83 UFB). †*Praeaulacon* Rasnitsyn, 1972 supported as clade (10%C: 0.91 BPP; 0%C: 0.92 BPP). †*Aulacogastrinus ater* (Rasnitsyn, 1972) and †*Praeaulacus daohugouensis* Zhang & Rasnitsyn, 2008 highly to maximally supported as clade (10%C: 0.99 BPP). †*Praeaulacinus* Rasnitsyn, 1972 weakly supported as clade (10%C, 0%C: 0.40 BPP). †*Praeaulacus* Rasnitsyn, 1972 never monophyletic.
•• **† Praeaulacidae: †Cretocleistogastrinae** Rasnitsyn, 1990. All sampled species cluster together; †*Westratia* Jell & Duncan, 1986 is not supported as monophyletic. The genera, represented by a single species in this study, form a consistent group (10%C: 0.33 BPP): †*Cretocleistogaster hyperura* Rasnitsyn, 1990, †*Miniwestratia* Rasnitsyn, 1990, and †*Nanowestratia* Rasnitsyn, 1990, with †*Westratia femorata* Rasnitsyn, 1990 well-supported as sister to the former two taxa (10%C: 0.68 BPP). †*Sinowestratia* Zhang & Zheng, 2000 not evaluated.

**Trigonaloidea Cresson, 1887**. Traditionally comprises Trigonalidae and †Maimetshidae, which are here recovered as monophyletic (50%C: 0.71 BPP, 71 UFB; 40%C: 57 BPP, 88 UFB), and probably including the putative evanioid family †Andreneliidae (30%C: 0.65 BPP, 95 UFB; 20%C: 84 UFB; 10%C: 0.48 BPP, 81 UFB; 0%C: 0.61, 0.71 UFB).

• **Trigonalidae** Cresson, 1887. Monophyly of total clade Trigonalidae, *i.e.*, including †*Albiogonalys* Nel *et al*., 2003, uncertain. Crown clade Trigonalidae monophyletic (*e.g.*, 30%C: 0.99 BPP, 99 UFB).
• **† Maimetshidae** Rasnitsyn, 1975. Monophyletic (50%C: 0.99 BPP, 76 UFB; 40%C: 0.99 BPP, 86 UFB; 30%C: 0.99 BPP, 84 UFB; 20%C: 81 UFB; 10%C: 0.70 BPP, 88 UFB; 0%C: 0.92 BPP, 95 UFB). The oldest maimetshid fossils are †*Ahiromaimetsha najlae* Perrichot *et al*., 2011, †*Andyrossia joyceae* (Rasnitsyn & Jarzembowski, 1998), †*Turgonaliscus cooperi* Engel, 2016, and †*Zorophratra corynetes* Engel, 2016, all Barremian).
•• **† Zorophratrinae** Engel, 2016. Monotypic. Nested within †Maimetshinae Rasnitsyn, 1975 (see †Maimetshinae below).
•• **† Maimetshinae** Rasnitsyn, 1975. Divided into two tribes: †Maimetshini Rasnitsyn, 1975 and †Ahiromaimetshini Engel, 2016. **†Ahiromaimetshini** is not supported as monophyletic, as †*Turgonaliscus* is recovered close to †*Guyotemaimetsha enigmatica* Perrichot *et al*., 2004 (10%C: 0.64 BPP; 5%C: 0.67 BPP; 0%C: 0.66 BPP). With that exception, †*Ahiromaimetsha* is recovered outside of the **†Maimetshini**, a well-supported clade with the inclusion of **†*Zorophratra*** (50%C: 0.95 BPP; 40%C: 0.98 BPP, 88 UFB; 30%C: 0.99 BPP, 86 UFB; 20%C: 92 UFB; 10%C: 0.69 BPP, 94 UFB; 0%C: 0.89 BPP, 98 UFB). †*Ahiromaimetsha* is further supported as to †Maimetshini + †*Zorophratra* (50%C: 0.78 BPP, 46 UFB; 40%C: 0.89 BPP, 66 UFB; 30%C: 0.96 BPP, 63 UFB; 20%C: 69 UFB; 10%C: 0.39 BPP, 70 UFB; 0%C: 0.50 BPP, 80 UFB). †*Turgonalus* clusters outside of the †Maimetshini + †*Zorophratra* clade but is not recovered with †*Ahiromaimetsha*. †*Andyrossia* is highly supported as close to †*Iberomaimetsha rasnitsyni* Ortega-Blanco *et al*., 2011 (5%C, 0%C: 0.95 BPP), with †*I. pallida* Perrichot & Perkovsky, 2016 weakly supported as sister (5%C, 0%C: 0.53 BPP).
• **† Andreneliidae** Rasnitsyn & Martínez-Delclòs, 2000. Monotypic. Within Trigonaloidea, supported as sister to †*Albiogonalys* Nel *et al*., 2003 (30%C: 0.63 BPP, 80 UFB; 20%C: 80 UFB; 10%C: 0.68 BPP; 0%C: 0.68 BPP).

### Section 2. Aculeate topologies

**Aculeata Latreille, 1802**. [Equivalent to Vespomorpha Laicharting, 1781, as used by Rasnitsyn (1988).] The backbone topology of the Aculeata has recently been largely resolved via phylogenomic analysis (*e.g.*, Branstetter *et al*. 2017, Peters *et al*. 2017), providing an ideal opportunity to reevaluate fossil placement while also informing ancestral state estimation procedures. No prior study has explicitly attempted to evaluate placement of the †Bethylonymoidea, a putative aculeate superfamily (Rasnitsyn 1975, 1988; Rasnitsyn & Quicke 2002), which would thus include the oldest representatives of the Aculeata during Mid Jurassic times. Other important taxa include the †Angarosphecidae, which have been presumed to represent the oldest crown Apoidea (*e.g.*, Rasnitsyn & Jarzembowski 1998), hence Vespaculeata. These oldest angarosphecids have been used to calibrate the Apoidea in divergence dating analysis (*e.g.*, Branstetter *et al*. 2017). Additionally, many taxa have been placed in the crown groups of other superfamilies without statistical analysis. The Bayesian analyses are emphasized due to the molecular scaffold, and results from the 50% complete + †Bethylonymidae tip-dating analysis (“50CB”, see main text methods) are also reported. For the phylogenetic classification used here, see Section 4 below.

• **Aculeata: *incertae sedis***. One superfamily and one genus are *incertae sedis* in the Aculeata after the present analyses.
•• **† Bethylonymoidea** Rasnitsyn, 1975. The most complete bethylonymoids are between 30– 39%, thus analyses at this level are emphasized. At 30% matrix completeness, four Jurassic †Bethylonymidae Rasnitsyn, 1975 are included, representing one †*Bethylonymus* Rasnitsyn, 1975 and three †*Bethylonymellus* Rasnitsyn, 1975, plus one putative Early Cretaceous bethylonymid, †*Meiagaster cretaceus* Rasnitsyn & Ansorge, 2000. In this analysis, †Bethylonymidae are recovered as a grade on the stem of the Vespaculeata (30%C: 0.45 BPP), with the Jurassic taxa clustering together (30%C: 0.32 BPP). In ML analysis, Chrysidoidea and Dryinoidea are erroneously recovered as a clade, but notably with the Jurassic species a grade at their base (30%C: 97 UFB). †*Meiagaster* is one node stemward from the artificial chrysidoid-dryinoid clade in the ML analysis (30%C: 67 UFB) or one node crownward toward the vespaculeates in Bayesian (30%C: 0.53 BPP). †*Allogaster ovata* Ren, 1995 (the genus an unrecognized junior homonym of *Allogaster* Thomson, 1864 [Cerambycidae]) is unplaceable, albeit the terminal clusters surprisingly with †*Montsecosphex jarzembowskii* Rasnitsyn & Martínez-Delclòs, 2000. In Bayesian analyses with more taxa but a lower level of completeness, bethylonymids cluster in what is effectively a basal polytomy of the Aculeata, with uninformative support values. Contrastingly, ML analysis at the 20% completeness level supports a †Bethylonymidae clade with exclusion of the unplaceable †*Renymus* (= †*Allogaster*) (20%C: 64 UFB) that is sister to the remaining Aculeata (20%C: 81 UFB), as well as a clade of †*Bethylonymus* and †*Bethylonymellus* (20%C: 80 UFB), the latter weakly supported as a clade within a polytomy of the former (20%C: 64 UFB). Support degrades to meaninglessness in less-complete analysis (10%C and 0%C levels). The conclusion thus is that the †Bethylonymidae, as represented by the most complete Jurassic fossils, belong to a basal polytomy of the Aculeata, with some probability that they belong on the stem of the chrysidoid, dryinoid, or vespaculeate crown nodes.
•• **† Falsiformicidae** Rasnitsyn, 1975. Originally considered sister to the Formicoidea. †*Falsiformica* Rasnitsyn, 1975 is maximally supported as a clade with the inclusion of the putative scolebythid †*Siccibythus*; the latter had been transferred to the †Falsiformicidae by Rasnitsyn *et al*. (2020). This clade is robustly supported as sister to the putative apoid genus **†*Burmasphex*** Melo & Rosa, 2018 (50%C: 0.99 BPP; 40%C: 0.94 BPP; 30%C: 0.98 BPP; 50CB: 0.97 BPP). The †*Falsiformica* + †*Burmasphex* clade is weakly supported as sister to Apoidea (50%C: 0.45 BPP, support within the Vespaculeata: 50%C: 1.0 BPP), sister to †Bethylonymidae and the two sister to Vespaculeata (40%C: 0.49 BPP; 50CB: 0.37 BPP) or in a bethylonymid grade on that stem (30%C: 0.31 BPP; 0%C: 0.22 BPP), or as a member of the total Dryinoidea again in a bethylonymid grade (10%C: 0.26 BPP). Two undescribed taxa from burmite (unique specimen identifiers: CASENT0844568, 0844587) also cluster with †*Falsiformica* + †*Burmasphex*. Because of the remaining uncertainty, †Falsiformicidae is here considered unplaced in Aculeata.
• **Chrysidoidea** Latreille, 1802. The three major questions pertaining the chrysidoid fossil record are placement of †Chrysobythidae, †Falsiformicidae (see Aculeata *incertae sedis*, above), and the monotypic family †Plumalexiidae. Other fossil Chrysidoidea are supported as members of their given families with high support (see raw data matrix for taxon sampling).
•• **Bethylidae**: †*Holopsenella primotica* Engel *et al*., 2016, †*Holopsenelliscus pankowskiorum* Engel, 2019, and †*Lancepyris opertus* Azevedo & Azar, 2012 consistently formed the first and second splits of the total Bethylidae clade (50%C: 1.0, 0.99, 0.67 BPP; 40%C: 0.94, 0.94, 0.75 BPP; 30%C: 0.96, 0.96, 0.65 BPP). Two additional, undescribed species from burmite (CASENT0844562, 084474) were nested within the crown (50%C: 0.94 BPP; 40%C: 0.93 BPP; 30%C: 0.95 BPP). Note that several additional Mesozoic Bethylidae are known, but these were not sampled in the present study.
•• **Chrysididae**: Five describe and two undescribed Mesozoic species were sampled. A specimen from burmite (CASENT0844579) is consistently recovered as sister to the remainder of the family (60%C: 0.98 BPP; 50%C: 0.93 BPP; 30%C: 0.70 BPP; 10%C: 0.72 BPP). †*Burmasega ammirabilis* Lucena & Melo, 2018 is consistently recovered with Amiseginae Mocsary, 1889 (40%C: 0.92 BPP; 30%C: 0.3 BPP), the two including Loboscelidiinae Maa & Yoshimoto, 1961 with high support at greater matrix completeness (60%C: 0.87 BPP; 50%C: 0.85 BPP). The relatively incomplete †*Protamisega khetanga* Evans, 1973 clusters with the Amiseginae + Loboscelidiinae clade (30%C: 0.39 BPP; 10%C: 0.46 BPP; 0%C: 0.45 BPP). †*Miracorium tetrafoveolatum* Lucena & Melo, 2018 is weakly supported stemward one node (50%C, 60%C: 0.52 BPP). Besides a undescribed species burmite (CASENT0844585) with maximal support as sister to the crown Chrysidinae in all analyses in which it is included, other described, sampled taxa have limited signal (†*Auricleptes nebulosus* Lucena & Melo, 2018, †*Bohartiura glabrata* Lucena & Melo, 2018, †*Hypocleptes rasnitsyni* Evans, 1973, and †*Procleptes hopejohnsonae* McKellar & Engel, 2014).
•• **Scolebythidae**: All fossils attributed to Scolebythidae except †*Siccibythus musculosus* Cockx & McKellar, 2016 (see †Falsiformicidae below) cluster with the crown clade, including the Barremian fossils †*Libanobythus milkii* Prentice & Poinar, 1996, †*Mirabythus lechrius* Cai *et al*., 2012, and †*Zapenesia libanica* Engel & Grimaldi, 2007. At the 40% completeness level, the total Scolebythidae clade weakly includes †*Plumarius* Philippi, 1873 (Plumariidae), for which sequence data are not available.
•• **Plumariidae**: See Scolebythidae above and †Plumalexiidae below.
•• **† Chrysobythidae** Melo & Lucena, 2019. Robustly recovered as monophyletic (50%C: 0.97 BPP; 40%C: 0.99 BPP; 30%C, 10%C, 0%C, 50CB: 1.0 BPP) and sister to the total clade **†Bethylidae** Forster, 1856 (50%C: 0.99 BPP; 40%C: 0.89 BPP; 30%C: 0.93 BPP; 10%C: 0.76 BPP; 0%C: 0.93 BPP; 50CB: 0.99 BPP).
•• **† Plumalexiidae** Brothers, 2011. Tip-dating analysis places †*Plumalexius* Brothers, 2011 as sister to Plumariidae Bischoff, 1904 with high support (50CB: 0.99 BPP), and the two in Chrysidoidea with similar consistency (50CB: 0.97). In morphology-only analysis, only those runs with highest data completeness resolved meaningful support, where †*Plumalexius* was recovered as member of the total Chrysidoidea (60%C: 0.6) or sister to the Aculeata (40%C: 0.71), with limited support for a sistergroup relationship to Plumariidae (50%C: 0.58 BPP). ML results are not reported due to the lack of soft constraints.
• **Dryinoidea** Haliday, 1833. Represented by three families, Dryinidae Haliday, 1833, Embolemidae Förster, 1856, and Sclerogibbidae Ashmead, 1901, with fossils attributed to each.
•• **Dryinidae**: monophyletic with the sampled fossils in all virtually all analyses (50%C, 40%C, 30%C, 50CB: 1.0 BPP), with support decaying in analyses with < 30% completeness.
•• **Embolemidae**: monophyletic including the Cretaceous compression fossil †*Baissobius minimus* Rasnitsyn, 1996 (30%C: 1.0 BPP). At the 10% completeness level, support diminishes. Other putative Embolemidae tend to cluster with the core clade represented by *Ampulicomorpha* Ashmead, 1893, although with some affinity for Dryinidae.
•• **Sclerogibbidae**: †*Sclerogibba cretacica* Martynova *et al*., 2019 clusters weakly with Dryinoidea in morphology-only analysis (50%C: 0.71 BPP) but more strongly in tip-dating (50CB: 0.98 BPP). the older †*Sclerogibbodes embioleia* Engel & Grimaldi, 2006 is strongly supported as on the stem of Sclerogibbidae (40%C: 0.98 BPP; 30%C: 0.96 BPP; 10%C: 0.79 BPP; 0%C: 0.82 BPP).
• **Vespoidea** Laicharting, 1781. Comprises Rhopalosomatidae Ashmead, 1896 and Vespidae Laicharting, 1781, plus †*Prosphex* Grimaldi & Engel, 2019 and an undescribed taxon from burmite (CASENT0844576) very similar to †*Alivespa* Wu *et al*., 2020 (50CB: 0.92 BPP; 40%: 0.47 BPP). Placement of †*Prosphex* in the superfamily is uncertain. Note that †*Angarosphex bleachi* and †*A. consensus* cluster this superfamily (see †Angarosphecidae below).
•• **Rhopalosomatidae**: †*Eorhopalosoma* Engel, 2008 forms an unequivocal clade with extant Rhopalosomatidae. Despite being transferred to the “apoid” form family †Angarosphecidae, †*Mesorhopalosoma cearae* Darling, 1990 is supported as sister to †*Eorhopalosoma* and the extant rhopalosomatids (10%C: 0.63 BPP; 0%C: 0.80 BPP).
•• **Vespidae**: †*Protovespa haxairei* Perrard & Carpenter, 2017 is recovered as sister to crown Vespidae (50%C: 1.0 BPP; 40%C: 0.90 BPP; 30%C: 0.80 BPP). Within the crown Vespidae are recovered species of †*Priorvespa* Carpenter & Rasnitsyn, 1990, †*Curiosivespa* Rasnitsyn, 1975, and the Raritan ?*Symmorphus senex* Carpenter, 2000. Wu *et al*. (2020) described a new genus of fossil Vespidae, †*Alivespa*, which they also recover within the crown.
• **Pompiloidea** Latreille, 1804 *sensu novum*. Comprises Sierolomorphidae, Tiphiiformes, and core Pompiloidea.
•• **Sierolomorphidae** Krombein, 1951. One fossil species has previously been assigned to the family, †*Loreisomorpha nascimbenei* Rasnitsyn, 2000. This species, along with an undescribed taxon from burmite (CASENT0844589) are recovered as a clade (50%C: 0.84 BPP; 40%C: 0.82 BPP; 30%C: 0.80 BPP; 10%C: 0.65 BPP; 0%C: 0.48 BPP).
•• **Tiphiiformes**. †*Architiphia rasnitsyni* Darling, 1990 and †*Thanatotiphia nyx* Engel *et al*., 2009 have been attributed to Thynnidae and Tiphiidae, respectively. †*Architiphia* is unstable, clustering with *Pepsis* (**Pompilidae**) (40%C: 0.89 BPP) or †*Cretofedtschenkia* Osten, 2007 (30%C: 0.76 BPP; 10%C: 0.73 BPP) in a pompiloid polytomy. †*Thanatotiphia* is unplaceable with the current dataset, despite inclusion of the male unciform process. Possibly this is due to parallel wing venation reduction and gain of a “pseudosting” in Bradynobaenidae.
•• **Core Pompiloidea**. Comprises Pompilidae Latreille, 1804, Sapygidae Latreille, 1810, Mutillidae Latreille 1802, †Bryopompilidae Rodriguez *et al*., 2016, †Burmusculidae Zhang *et al*., 2018. Note that †*Eubaissodes*, †*Ilerdosphex*, †*Pompilopterus ciliatus*, †*P. montsecensis*, and †*P. wimbledoni* cluster with this superfamily (see †Angarosphecidae below).
••• **† Burmusculidae**: strongly supported as a stem lineage of **Pompilidae**, given the present sampling (60%C, 50%C: 0.99 BPP); compression fossils attributed to the “†Angarosphecidae” disrupt support for this clade.
••• **† Bryopompilidae**: supported as sister to Sapygidae + Mutillidae (70%C: 0.82 BPP; 60%C: 0.89 BPP; 50%C: 0.95 BPP; 40%C: 0.91 BPP; 30%C: 0.95BPP); support decays with matrices < 30% complete due to interruption by wing fossils (*e.g.*, †*Pompilopterus keymerensis* Rasnitsyn & Jarzembowski, 1998).
••• **Sapygidae**: †*Cretosapyga* clusters with the Sapygidae + Mutillidae clade, with some probability of being sister to Mutillidae (60%C: 0.82 BPP; 50%C: 0.74 BPP; 40%C: 0.70 BPP; 30%C: 0.65 BPP), deserving renewed focus. The Crato compression fossil †*Cretofedtschenkia*, as noted for Tiphiiformes above, clusters outside of the Pompiloidea.
••• **Mutillidae**: Of the two Mesozoic fossils attributed to the family, †*Cretavus sibiricus* Sharov, 1957 and †*Mesomutilla aptera* Zhang, 1985, only the latter was evaluated, and it was found the be unplaceable in Aculeata, lacking support for any specific relationship (hence *incertae sedis* in Aculeata, **stat. nov.**). An undescribed species from burmite (CASENT0844568) was robustly recovered as sister to the Mutillidae, however (70%C: 1.0 BPP; 60%C, 50%C: 0.97 BPP; 40%C: 0.96 BPP; 30%C: 0.95 BPP).
• **Scolioidea** Latreille, 1802. Comprises the extant families Scoliidae Latreille, 1802 and Bradynobaenidae de Saussure, 1892. Fossils attributed to Scolioidea are all considered to be Scoliidae, and have been placed either in the †Archaeoscoliinae Rasnitsyn, 1993 or Proscoliinae Rasnitsyn, 1977. Because the latter subfamily includes *Proscolia* Rasnitsyn, 1977, placement in Proscoliinae implies that these fossils are members of the crown group of the family; any fossil placement to family also implies crown-group membership in the superfamily.
•• **Crown Scoliidae**: Highly to maximally supported clade, with the exclusion of all Mesozoic fossils, including those attributed to Proscoliinae (50%C: 0.98 BPP; 40%C, 30%C: 0.99 BPP; 10%C, 0%C: 1.0 BPP).
•• **† Archaeoscoliinae**: Comprises three Mesozoic genera, †*Archaeoscolia* Rasnitsyn, 1993, †*Cretaproscolia josai* Rasnitsyn & Martínez-Delclòs, 1999 (note that second species in genus attributed to Proscoliinae), and †*Protoscolia* Zhang *et al*., 2002. Not monophyletic: **†*Protoscolia*** and “proscoliine” **†*Sinoproscolia*** Zhang *et al*., 2015 form polytomy at base of total Scolioidea, with †*Archaeoscolia* and †*Cretaproscolia* a highly supported clade with the remainder of the scoliid taxa (“Proscoliinae” + crown Scoliidae clade, or “total Scoliidae”) (50%C: 0.98 BPP; 40%C: 0.99 BPP; 10%C: 0.98 BPP; 0%C: 0.99 BPP). †*Protoscolia* Zhang *et al*., 2002 maximally supported as monophyletic (40%C, 30%C, 10%C, 0%C: 1.0 BPP).
•• **Proscoliinae**: Comprises two fossil genera, †*Cretoscolia* Rasnitsyn, 1993, †*Sinoproscolia yangshuwanziensis* Zhang *et al*., 2015, and †*Cretaproscolia asiatica* Zhang, 2006. Not monophyletic: †*Sinoproscolia* never forms clade with †*Cretoscolia* and is strongly supported as outside of the total Scoliidae clade (see †Archaeoscoliinae above for support). Additionally, “Proscoliinae” excluding †*Sinoproscolia* forming grade to crown Scoliidae. †*Cretaproscolia* Rasnitsyn & Martínez-Delclòs, 1999 highly supported as a clade (10%C, 0%C: 0.98 BPP). †*Cretoscolia* is never monophyletic, although †*C. formosa*, †*C. laiyangica*, and †*C. rasnitsyni* form a clade with some consistency (10%C: 0.75 BPP; 0%C: 0.91 BPP), clustered with †*C. montsecana* (10%C: 0.29 BPP; 0%C: 0.33 BPP).
• **Apoidea** Latreille, 1802. The vast majority of Mesozoic fossils attributed to the Apoidea are placed in the form taxon †Angarosphecidae. With the exception of †*Burmasphex*, which is never recovered with the Apoidea (see †Falsiformicidae above), all taxa attributed to the †Angarosphecidae are compression fossils. Conversely, most fossils attributed to other apoid families are preserved in amber, except a putative member of Apidae from the Crato formation (Osten 2007, p. 354, Fig. 11.71). These other amber apoid fossils are attributed to Ampulicidae, Apidae, †Melittosphecidae, Pemphredonidae, or Sphecidae as broadly conceived, *i.e.*, prior to molecular insights into apoid phylogeny (Sann *et al*. 2018, Branstetter *et al*. 2017, Peters *et al*. 2007). †Angarosphecidae is treated last because of its complexity.
•• **Ampulicidae** Shuckard, 1840. Six taxa representing putative Mesozoic Ampulicidae were sampled, †*Apodolichurus sphaerocephala* Antropov, 2000, †*Cretampulex gracilis* Antropov, 2000, †*Gallosphex cretaceus* Schlüter, 1978, †*Mendampulex monicularis* Antropov, 2000, an undescribed species from burmite (CASENT0844571), and †*Cariridris bipetiolata* Brandão & Martins-Neto, 1990, a species originally attributed to Formicidae. Note that †*Angarosphex penyaleveri* and †*A. saxosus* cluster with this family (see †Angarosphecidae below). **†*Apodolichurus*** clusters weakly with ***Heterogyna*** *nocticola* Ohl, 2006 (Heterogynaidae Nagy, 1969) (60%C: 0.48 BPP; 30%C: 0.54 BPP; 10%C, 0%C: 0.82 BPP) and more strongly so when †*Mendampulex* is included (50%C: 0.76 BPP), with a sistergroup relationship between the two indicated at the 50% completeness level (50%C: 0.90 BPP). Support for †*Apodolichurus* + *Heterogyna* disappears at the 40% completeness level. The putative sphecid **†*Burmastatus*** *triangularis* Antropov, 2000 clusters with the possible *Heterogyna* + *Apodolichurus* clade as its sistergroup (60%C: 0.46 BPP; 10%C: 0.79 BPP; 0%C: 0.95 BPP; see also “Sphecidae” below for more on placement of †*Burmastatus*). **†*Cretampulex*** is not supported as a member of the Ampulicidae, rather it comes out in a basal polytomy of the Apoidea, weakly clustering with the remainder of the superfamily (50%C: 0.52 BPP; 40%C: 0.58 BPP); at lower levels support for its placement disappears entirely **†*Gallosphex*** and **†*Mendampulex*** are supported as a clade (30%C: 0.86 BPP) clustering very weakly crown Ampulicidae (30%C: 0.24 BPP; 10%C: 0.30 BPP; 0%C: 0.21 BPP). The undescribed species CASENT0844571 is highly to maximally supported as stem Ampulicidae (70%C: 1.0 BPP; 60%C: 0.84 BPP; 50%C: 0.91 BPP; 40%C: 0.54 BPP; 30%C: 0.52 BPP). Finally, **†*Cariridris*** falls within Formicoidea, a clade supported at 0.74 BPP in the 5% complete analysis and at 0.67 BPP in the 0%C analysis, in both cases clustering with Formicae.
•• **Apidae** Latreille, 1802. Two Mesozoic fossils are attributed to Apidae, a Crato fossil (see Osten 2007) and †*Cretotrigona prisca* Michener & Grimaldi, 1988 from Raritan amber. The putative Crato apid clusters in a basal polytomy of the Apoidea near †*Cretobestiola* Pulawski & Rasnitsyn, 2000, †*Archisphex proximus* Rasnitsyn & Jarzembowski, 1998, †*A. boothi* Jarzembowski, 1991, and †*Trichobaissodes antennatus* Rasnitsyn, 1975 (10%C: 0.26 BPP; 5%C: 0.32 BPP; 0%C: 0.20 BPP). ML analysis strongly conflicts, as the Crato apid is recovered as a member of Anthophila (10%C, 0%C: 99 UFB). **†*Cretotrigona***, in contrast, always places as Apidae (70%C, 60%C, 50%C, 40%C, 30%C, 10%C, 0%C: 1.0 BPP).
•• **† Melittosphecidae** Poinar & Danforth, 2006. †*Melittosphex burmensis* Poinar & Danforth, 2006 always places in Apoidea (60–30%C: 1.0 BPP), with support for Apoidea interrupted by †*Cretosphecium lobatum* Pulawski & Rasnitsyn, 2000 below the 30% completeness threshold (see †Angarosphecidae below). Within Apoidea, †*Melittosphex* surprisingly clusters with the Megachilidae + Apidae clade (60%C: 0.87 BPP; 50%C, 40%C: 0.85 BPP; 30%C: 0.68 BPP; 10%C: 0.71 BPP; 0%C: 0.57 BPP).
•• **Pemphredonidae** Dahlbom, 1835. Pemphredonidae is a polyphyletic assemblage (Branstetter *et al*. 2017, Sann *et al*. 2018). Eleven species attributed to Pemphredonidae were included as terminals, and they cluster together as a clade with limited support (10%C: 0.30 BPP; 5%C: 0.36 BPP; 0%C: 0.32 BPP). This Cretaceous pemphredonid clade comes out in a basal polytomy of Apoidea; more extant taxa and reevaluation of the morphological boundaries of the pemphredonid clades are necessary to resolve this problem. Within this weakly-supported Cretaceous clade, †*Cretoecus spinicoxa* Budrys, 1993 is recovered as sister to a well-supported clade comprising †*Cretospilomena familiaris* Antropov, 2000, †*Colmepsiterona cumcarena* Cockx & McKellar, 2018, †*Psolimena electra* Antropov, 2000, †*Menopsila dupeae* Bennett *et al*., 2014, and †*Lisponema singularus* Evans, 1969 (60%C: 0.92 BPP; 50%C: 0.91 BPP; 40%C: 0.95 BPP; 30%C: 0.93 BPP; 10%C: 0.95 BPP; 0%C: 0.96 BPP). †*Psolimena*, †*Menopsila*, and †*Lisponema* are strongly supported as a clade (50%C: 0.97 BPP; 40%C: 1.0 BPP; 30%C: 0.97 BPP; 10%C: 0.99 BPP; 0%C: 0.98 BPP). The oldest putative pemphredonid, **†*Iwestia provecta*** Rasnitsyn & Jarzembowski, 1998, is a wing from the Lulworth formation (Berriasian). †*Iwestia* clusters within the main Cretaceous pemphredonid clade (*i.e.*, that clade excluding †*Cretoecus*) (5%C: 0.51 BPP; 0%C: 0.60 BPP); note that this fossil is unplaced in the ML analysis.
•• **Sphecidae** Latreille, 1802. Two Mesozoic species attributed to Sphecidae were included, †*Burmastatus triangularis* and †*Cirrosphex admirabilis* Antropov, 2000. †*Burmastatus* is unstable and clusters variably with a *Heterogyna* + †*Apodolichurus* clade (see Ampulicidae above), or with fossils attributed to Pemphredonidae (50%C, 40%C: 0.47 BPP). Likewise, †*Cirrosphex* is unplaceable in basal apoid polytomy, but clustering away from Ampulicidae. Neither species is supported as near Sphecidae *sensu stricto* (*i.e.*, the clade recovered by molecular analysis).
•• **† Angarosphecidae** Rasnitsyn, 1975. This is a highly heterogeneous form taxon which, even with exclusion of †*Burmasphex* (see †Falsiformicidae above) and †*Mesorhopalosoma* (see Rhopalosomatidae above), is never recovered as a clade. Likewise, four of the five genera in the group with more than one species are non-monophyletic, with the exception of †*Cretobestiola* Pulawski & Rasnitsyn, 2000. The other four genera are †*Angarosphex* Rasnitsyn, 1975, †*Archisphex* Evans, 1969, †*Baissodes* Rasnitsyn, 1975, and †*Pompilopterus* Rasnitsyn, 1975. The treatment of this family will proceed with monotypic genera, including those for which only a single terminal was sampled, followed by the five polytypic groups.
••• **† Angarosphecidae, monotypic genera**. **†*Cretosphecium lobatum*** Pulawski & Rasnitsyn, 2000 is one of two species in the genus, and clusters with Anthophila (10%C: 0.64 BPP; 5%C: 0.64 BPP; 0%C: 0.48 BPP). **†*Eubaissodes completus*** Zhang, 1992 clusters in the **Pompiloidea** (40%C: 0.50 BPP), support for which is completely decayed below the 40% completeness threshold due to uncertainty of placement for other fossil taxa and is pulled into a basal vespaculeate polytomy below 30% completeness; there is some limited support for a relationship of this species with †*Baissodes* (see below). **†*Ilerdosphex wenzae*** Rasnitsyn, 2000 clusters with Pompilidae *sensu novum* within **core clade Pompiloidea** with the same support as †*Eubaissodes* at the 40% completeness level, but clusters with Apoidea at lower completeness, albeit with meaningless support (< 0.10 BPP) due to uncertainty. If the core pompiloid placement is further supported, †*Ilerdosphex* would represent the oldest member of the superfamily, being Barremian in age. The best supported position for **†*Montsecosphex*** Jarzembowski, Rasnitsyn & Martínez-Delclòs, 2000 is at the 30% level, where it is close to †*Eubaissodes* (30%C: 0.53 BPP), clustering in the core Pompiloidea. In less-complete analyses, †*Montsecosphex* remains clustered with the Pompiloidea. **†*Oryctobaissodes armatus*** Rasnitsyn, 1975 weakly clusters with †*Ilerdosphex* (30%C: 0.33 BPP; 10%C: 0.64 BPP; 5%C: 0.65 BPP; 0%C: 0.55 BPP). **†*Trichobaissodes antennatus*** Rasnitsyn, 1975 is unplaceable in the Vespaculeata, although it does cluster with †*Archisphex boothi*, †*A. proximus*, the putative Crato apid, and †*Cretobestiola* (10%C: 0.26 BPP; 5%C: 0.32 BPP; 0%: 0.20 BPP). **†*Vitimosphex vividus*** Rasnitsyn, 1975 is one of two species in the genus, and it is strongly supported as sister to †*Angarosphex magnus* (Darling, 1990) (30%C: 0.90 BPP) or both †*A. magnus* and †*A. parvus* (Darling, 1990) (10%C: 0.95 BPP; 5%C: 0.93 BPP; 0%C: 0.92 BPP), the latter two being Crato fossils. The †*Vitimosphex* + Crato †*Angarosphex* species clade is unplaceable in Vespaculeata.
••• **† Angarosphecidae: †*Angarosphex***. This form genus comprises 15 species at the time of writing, all of but one of which were sampled, and are treated stratified by age. **(1)** †*Angarosphex **bleachi*** Rasnitsyn & Jarzembowski, 1998 and †*Angarosphex **consensus*** Rasnitsyn & Jarzembowski, 1998 are the oldest species attributed to this form genus, being wing fossils from the Hauterivian Weald clay formation; both species cluster with **Vespoidea** with limited support (10%C: 0.46 BPP; 5%C: 0.34 BPP; 0%C: 0.36 BPP). **(2)** The next oldest species are †*A. baektoensus* Jon *et al*., 2019, †*A. goldringi* Jarzembowski, 1991 (a wing), †*A. lithographicus* Rasnitsyn & Ansorge, 2000, and †***Apisphex penyaleveri*** (Rasnitsyn & Martínez-Delclòs, 2000) **gen. nov. comb. nov.** †*Angarosphex **baektoensis*** weakly clusters with Apoidea in a basal polytomy (30%C: 0.36 BPP; less-complete analyses ∼0.1 BPP), or more meaningfully in a vespaculeate polytomy. †*Angarosphex **goldringi*** clusters with †*A. myrmicopterus* Rasnitsyn, 1975 with limited support (10%C, 5%C: 0.38 BPP; 0%C: 0.33 BPP) in a vespaculeate polytomy. †*Angarosphex **lithographicus*** clusters with †*A. **venulosus*** (Zhang, 1985) (30%C: 0.50 BPP; less complete: ∼0.2 BPP) on the stem of Vespoidea, along with †*A. strigosus* (30%C: 0.43 BPP; 10%C: 0.22 BPP; 5%C: 0.26 BPP; 0%C: 0.30 BPP). †*Apisphex **penyaleveri*** clusters with **Ampulicidae** (10%C: 0.45 BPP; 5%C: 0.55 BPP; 0%C: 0.57 BPP). **(3)** The remaining nine species are Aptian to Albian in age, and include †*A. beiboziensis* (Hong, 1984), †*A. lithodes* (Zhang, 1985), †*A. magnus*, †*A. myrmicopterus*, †*A. niger* Rasnitsyn, 1990 (not sampled), †*A. pallidus* Rasnitsyn, 1986, †*A. parvus*, †*A. saxosus* Zhang *et al*., 2018, and †*A. strigosus* (Zhang, 1992). †*Angarosphex **beiboziensis*** clusters with Vespidae with higher support than †*A. consensus* and †*A. bleachi* (10%C: 0.66 BPP; 5%C: 0.56 BPP; 0%C: 0.56 BPP). †*Angarosphex **lithodes*** is unplaceable in Vespaculeata, although it does cluster with Apoidea with meaningless support (10–0%C). †*Angarosphex **magnus*** and †*A. **parvus*** cluster with †*Vitimosphex* (see above). †*Angarosphex **myrmicopterus*** clusters with †*A. goldringi* (see above). †*Angarosphex **pallidus*** is unplaceable in Vespaculeata (30–0%C). †*Angarosphex **saxosus*** clusters with **Ampulicidae** (40%C: 0.47 BPP; 30%C: 0.42 BPP; 10%C: 0.30 BPP; 5%C: 0.31 BPP; 0%C: 0.46 BPP). Finally, †*A. **strigosus*** clusters with †*A. lithographicus* and †*A. venulosus* (see above).
••• **† Angarosphecidae: †*Archisphex***. Not monophyletic. Comprises six species, all of which were sampled: †*Archisphex boothi* Jarzembowski, 1991, †*A. catalunicus* (Ansorge, 1993) (wing), †*A. crowsoni* Evans, 1969 (wing), †*A. curvus* Rasnitsyn & Jarzembowski, 1998 (wing), †*A. incertus* (Rasnitsyn, 1975), and †*A. proximus* Rasnitsyn & Jarzembowski, 1998 (wing). †*Archisphex **crowsoni*** Evans, 1969 is both the type and oldest species of the genus; this wing from the Wadhurst clay formation (Valanginian) clusters with the Apoidea but has no meaningful support. †*Archisphex **boothi*** and †*A. **proximus*** strongly cluster together (10%C, 5%C: 0.96 BPP; 0%C: 0.93 BPP), and together cluster in with the Apoidea, albeit with very limited to no support. †*Archisphex **catalunicus*** and †*A. **curvus*** are very weakly drawn together (10–0%C: ∼0.18 BPP), the two in a basal polytomy of the Vespaculeata. Finally, †*A. **incertus*** is unplaceable in the vespaculeate polytomy (10–0%C).
••• **† Angarosphecidae: †*Baissodes***. Not monophyletic. Comprises four species, all but one of which were sampled: †*B. grabaui* Ren, 1995 (not sampled), †*B. longus* Rasnitsyn, 1986, †*B. magnus* Rasnitsyn, 1975, and †*B. robustus* Rasnitsyn, 1975. †*Baissodes **longus*** does not meaningfully cluster with the other species and is unplaceable in the Vespaculeata. †*Baissodes **magnus*** and the type species †*B. **robustus*** are, however, weakly supported as a clade with †*Eubaissodes* (*†B. robustus* + †*Eubaissodes*: 10%C: 0 BPP; 5%C: 0.45 BPP; 0%C: 0.47 BPP; †*B. magnus* + †*B. robustus* + †*Eubaissodes*: 10%C: 0.27 BPP; 5%C: 0.19 BPP; 0%C: 0.24 BPP).
••• **† Angarosphecidae: †*Cretobestiola***. Monophyletic (10%C: 0.72 BPP; 5%C: 0.65 BPP; 0%C: 0.55 BPP). Unplaceable in vespaculeate polytomy (10–0%C).
••• **† Angarosphecidae: †*Pompilopterus***. Not monophyletic. Comprises nine species, all of which are sampled: †*Pompilopterus ciliatus* Rasnitsyn, 1975 (wing), †*P. corpus* Rasnitsyn & Jarzembowski, 1998, †*P. difficilis* Rasnitsyn & Jarzembowski, 1998 (wing), †*P. keymerensis* Rasnitsyn & Jarzembowski, 1998 (wing), †*P. leei* Rasnitsyn & Jarzembowski, 1998 (wing), †*P. montsecensis* Rasnitsyn, 2000, †*P. noguerensis* Rasnitsyn & Martínez-Delclòs, 2000, †*P. wimbledoni* Rasnitsyn & Jarzembowski, 1998 (wing), and †*P. worssami* Rasnitsyn & Jarzembowski, 1998 (wing). The oldest two fossils are †*P. **difficilis*** and †*P. **wimbledoni***, both from Berriasian Lulworth formation; the former clusters with Apoidea without meaningful support, whereas the latter clusters with **Pompiloidea** (5%C: 0.39 BPP; 0%C: 0.41 BPP) with some affinity to †*Bryopompilus* (5%C: 0.30 BPP; 0%C: 0.35 BPP). The next oldest fossils are †*P. **keymerensis***, †*P. **leei***, and †*P. **worssami***, all from the Hauterivian Weald clay; †*P. keymerensis* weakly clusters with Pompiloidea (10%C: 0.20 BPP; 5%C: 0.19 BPP 0%C: 0.22 BPP); †*P. leei* very weakly clusters with Sierolomorphidae (5–0%C, < 0.15 BPP), so is unplaced in Vespaculeata; and †*P. worssami* clusters with Vespoidea without meaningful support. †*Pompilopterus **corpus***, †*P. **montsecensis***, and †*P. **noguerensis*** are Barremian; †*P. corpus* weakly clusters with †*Ilerdosphex* and †*Oryctobaissodes* (10%C: 0.21 BPP; 5%C: 0.22 BPP; 0%C: 0.16 BPP) (see those genera above); †*P. montsecensis* clusters with *Pepsis* (**Pompilidae**) at the 30% complete level (30%C: 0.59 BPP) but is otherwise unplaceable in Vespaculeata; †*P. noguerensis* is unplaceable in Vespaculeata. Finally, †*P. ciliatus* clusters with *Pepsis* (**Pompilidae**) (0%C: 0.77 BPP).

### Section 3. Formicoid topologies

Support for the most important splits in this study are reported in Fig. 17, namely Formicoidea– Apoidea, †@@@idae–Formicidae, †@@@idae, †Sphecomyrmines–Antennoclypeata, and †Brownimeciinae–crown-Formicidae. Results are reported clade-by-clade, based on the present results (see Section 4.5 for systematic outline). A minimum of taxonomic actions is taken to ease cataloging.

**• †@@@idae fam. nov.** Two genera. See Fig. 17f or support at this node. Topology within †@@@idae is unresolved, except that the TONG-112 female (figured in Barden & Grimaldi 2016) receives some support as sister to the remainder of the family (60–0%C BPPs: 0.77– 0.93; 50–0%C UFBs: 23–67); †*Camelosphecia* (ANTWEB1038930, NIGP163574) is maximally supported as a clade.

• **Total clade Formicidae**. See Fig. 17 for support at this node.
•• **Total clade Formicidae, *incertae sedis* to subfamily**. **†*Archaeopone*** Dlussky, 1975, a form genus comprising male compression fossils, is not recovered as monophyletic. The two species, †*A. **kzylzharica*** Dlussky, 1975 and †*A. **taylori*** Dlussky, 1983, are unplaceable in the total Formicidae. **†*Baikuris*** Dlussky, 1987. Highly supported as Formicidae. Non-monophyletic based on the present analysis, thus representing a form taxon for unplaceable male stem Formicidae. Four species are attributed to this genus, all of which were sampled. †*Baikuris **casei*** Grimaldi *et al*., 1997, †*B. **mandibularis*** Dlussky, 1987, and †*B. **mirabilis*** Dlussky, 1987 are all unplaceable on the stem of Formicidae. In contrast, †*B. **maximus*** Perrichot, 2015 clusters with †Armaniinae (50%C: 0.54 BPP; 40%C: 0.48 BPP; 30%C: 0.72 BPP; 10%C: 0.39 BPP; 5%C: 0.31; 0%C: 0.35); despite this, †*B. maximus* bears little resemblance to the armaniines, thus should also be considered unplaced among stem Formicidae. **†*Burmomyrma rossi*** Dlussky, 1996 is the least-complete Mesozoic amber taxon attributed to Formicidae, as it is missing the anterior half of its soma. It has been attributed to Aneuretinae (Dlussky 1996) and †Falsiformicidae (Lucena & Melo 2018). In present analyses, †*Burmomyrma* clusters with Leptanillomorpha (10%C: 0.88 BPP; 5%C: 0.78 BPP; 0%C: 0.78) almost certainly due to the reduced wing venation, a “negative” set of synapomorphies. This position is considered implausible based on the unreduced (*i.e.*, large) third abdominal segment and because the key cranial features of the leptanillomorphs cannot be evaluated. For these reasons, †*Burmomyrma* is here treated as *incertae sedis* in total Formicidae. †***Petropone petiolata*** Dlussky, 1975, a worker-based fossil originally attributed to Ponerinae and preserved at < 5% completeness, is unplaceable in total Formicidae.
•• **†Armaniinae** Dlussky, 1983. Seven genera have attributed to this subfamily, †*Archaeopone* Dlussky, 1975, †*Armania* Dlussky, 1983, †*Khetania*, Dlussky, 1999, †*Orapia* Dlussky *et al*., 2004, †*Poneropterus* Dlussky, 1983, †*Pseudarmania* Dlussky, 1983, and the worker-based taxon †*Dolichomyrma* Dlussky, 1975. Most of these taxa are very incompletely preserved and receive diminishing to no support for any specific placement among the total Formicidae. Three of these genera are excluded: †*Archaeopone* is unplaceable in total Formicidae and lacks even those features indicative of relationship with †Armaniinae; †*Poneropterus* is supported as part of †Sphecomyrmines; and †*Dolichomyrma* clusters within the crown Formicidae. Because the subfamily is definable based on three well-supported synapomorphies, a shortened pronotum, presence of 1r-rs, and large body size, we retain the placement of most of †*Khetania* and †*Orapia* in the subfamily but recognize †*Armania* and †*Pseudarmania* as tribe †**Armaniini** Dlussky, 1983 **stat rev.** due to the comparatively high support for this clade. Considerable debate has been had about the placement of this group (*e.g.*, Dlussky 1983, Wilson 1987, Bolton 2003, LaPolla *et al*. 2013, Barden & Engel 2019).
••• **†Armaniinae: Armaniini stat. rev**. Comprises two genera and four species. **†*Armania robusta*** Dlussky, 1983, †*A. **curiosa*** (Dlussky, 1983), and **†*Pseudarmania rasnitsyni*** Dlussky, 1983, cluster consistently (30%C: 0.56 BPP; 10%C: 0.47 BPP; 5%C: 0.43 BPP; 0%C: 0.41 BPP). †*Pseudarmania **aberrans*** fails to cluster with the †Armaniini but can be attributed to the tribe based on the small, squamiform, and longitudinally sulcate petiole which is shared with †*P. rasnitsyni*.
••• **†Armaniinae: *incertae sedis* to tribe**. **†*Orapia rayneri*** Dlussky *et al*., 2004 clusters with †*Armania* and †*Pseudarmania* with limited support (30%C: 0.42 BPP; 10%C: 0.14 BPP; 5%C: 0.11 BPP; 0%C: 0.13 BPP), whereas †*O. **minor*** Dlussky *et al*., 2004 never places close to †Armaniinae. **†*Khetania*** Dlussky, 1999 is a form genus comprising three very poorly preserved compression fossils from the Emanra formation of the Khetana River. Two species formerly attributed to †*Armania* are here placed, forming †*K. **capitata*** (Dlussky, 1999) **n. comb.** and †*K. **pristina*** (Dlussky, 1999) **n. comb.** †*Khetania pristina* is similar to †*Armania* in having a short pronotum and robust body, while †*A. capitata* is barely recognizable (< 5% complete). A relationship of †*K. **mandibulata*** Dlussky, 1999 with †*Archaeopone taylori* is diminishingly supported (10%C: 0.34 BPP; 5%C 0.26 BPP; 0%C:
0.16 BPP), thus ignored.
•• **†Sphecomyrmines**. Comprises two subfamilies, one unplaced tribe, and three unplaced genera. See Fig. 17 for support at this node.
••• **†Sphecomyrmines: *incertae sedis* to subfamily**. **†*Cretomyrma*** Dlussky, 1975 comprises one species, †*C. unicornis* Dlussky, 1975 (= †*C. arnoldii* Dlussky, 1975 **j. syn.**), which was recovered in the †Sphecomyrmines (10%C: 0.59 BPP; 5%C: 0.38 BPP; 0%C: 0.16), often clustering with a Raritan male attributed to †*Sphecomyrma* from the AMNH (AMNH-NJ-242; 10%C: 0.71 BPP; 5%C: 0.70 BPP; 0%C: 0.64 BPP). Specific placement in the †Sphecomyrmines is not possible due to poor preservation. **†*Dlusskyidris zherichini*** (Dlussky, 1975), a male-based taxon, is unplaced in the Sphecomyrmines (50%C: 0.75 BPP; 40%C: 0.49 BPP; 30%C: 0.66 BPP; 10%C: 0.35 BPP). The 40% complete support is interrupted by †Armaniinae, which traversed into the Sphecomyrmines, whereas the 10% complete support is degraded by the highly incomplete †*Poneropterus*, and support below the 10% threshold is obliterated. **†*Poneropterus sphecoides*** Dlussky, 1983 clusters strongly with †Haidomyrmecinae based on wing venation (10%C: 0.76 BPP; 5%C: 0.55 BPP; 0%C: 0.54 BPP). Because of poor preservation preventing evaluation of key synapomorphies, †*Poneropterus* is considered *incertae sedis* in †Sphecomyrmines.
••• **†Sphecomyrmines: †Zigrasimeciinae** Borysenko, 2017. Comprises, according to Borysenko (2017), †*Boltonimecia canadensis* (Wilson, 1985) and species of †*Zigrasimecia* Barden & Grimaldi, 2013 (see also Cao *et al*. 2020b). **†*Zigrasimecia*** is maximally supported as clade in all analyses, with an undescribed species (CNU009193) representing the definitive sister group to the remaining species, also with maximal support. **†*Boltonimecia*** Borysenko, 2017 is only included at the 40% completeness level, and despite being poorly preserved is recovered as sister to †*Zigrasimecia* with maximal support in all but the least complete analyses (5–0%C). When †*Boltonimecia* is excluded, *i.e.*, at high matrix completeness, †*Zigrasimecia* strongly clusters with †Haidomyrmecines (70%C, 60%C: 1.0 BPP; 50%C: 0.92 BPP). At less-complete levels, †Zigrasimeciinae is highly supported as part of a clade including the †*Gerontoformica pilosa* species group, †*Myanmyrma* including †*M. mauradera* (40%C: 0.94 BPP; 30%C: 0.91 BPP; 10%C: 0.95 BPP; 5%C: 0.79 BPP; 0%C: 0.74 BPP). Within the *pilosa*–*mauradera*–†*Zigrasimecia* clade, †*Myanmyrma* and †*M. mauradera* are supported as the sistergroup to †Zigrasimeciini (40%C, 30%C: 0.85 BPP; 10%C: 0.86 BPP; 5%C: 0.78 BPP; 0%C: 0.72 BPP). Because of the conflict between the high- and low-completeness matrices, †Zigrasimeciini is here treated as unplaced to subfamily in Sphecomyrmines.
••• **†Sphecomyrmines: †Haidomyrmecinae** Bolton, 2003. An extremely well-supported subfamily defined by numerous cranial autapomorphies. Sister to †*Zigrasimecia* at high matrix completeness (see †Zigrasimeciinae above). Support for this clade evaporates at the 40% completeness level, where several male-based terminals, numerous †*Gerontoformica* species, and the poorly preserved †*Boltonimecia* are added. At completeness levels ≤ 40%, there is weak and diminishing support for †Haidomyrmecinae as sister to the remaining Sphecomyrmines (30%C: 0.66 BPP; 10%C: 0.59 BPP; 5%C: 0.38 BPP; 0%C: 0.10 BPP), with the placement of †*Dlusskyidris* considered unresolved. Among sampled species, within †Haidomyrmecinae, †*Ceratomyrmex* Perrichot *et al*., 2016 and †*Linguamyrmex* Barden & Grimaldi, 2017 form a maximally supported clade within a polytomy of †*Haidomyrmodes* Perrichot *et al*., 2008, †*Haidoterminus* McKellar *et al*., 2013, and †*Haidomyrmex* Dlussky, 1996.
••• **†Sphecomyrmines: †Sphecomyrminae** Wilson & Brown, 1967. Three genera are here attributed to †Sphecomyrminae: †*Gerontoformica* Nel & Perrault, 2004, †*Myanmyrma* Engel & Grimaldi, 2005, and †*Sphecomyrma* Wilson & Brown, 1967, as well as several males attributed to either †*Gerontoformica* or †*Sphecomyrma*. **†*Gerontoformica*** divides into three clades, the †*G. **orientalis*** species group (50%C: 0.10 BPP; 40%C: 0.33 BPP; 30%C: 0.31 BPP; 10%C: 0.29 BPP; 5%C: 0.25 BPP; 0%C: 0.26) which is often paraphyletic with respect to †*Sphecomyrma freyi*, the †*G. **pilosa*** species group (50%C: 0.94 BPP; 40%C: 0.70 BPP; 30%C: 0.75 BPP; 10%C: 0.79 BPP; 5%C, 0%C: 0.75 BPP), and †*M. mauradera*, which is maximally supported as sister to the poorly-preserved species †*Myanmyrma gracilis* Engel & Grimaldi, 2005 (40–0%C). **†*Myanmyrma*** and †*M. mauradera* share unique, elongate sickle-shaped mandibles, and presence of tractions setae on both the clypeus and labrum. **†*Sphecomyrma*** is here defined by absence of chaetae (“traction setae”) on the clypeus, in distinction to †*Gerontoformica*, plus apomorphic presence of an anteromedian clypeal lobe. Because †*S. **mesaki*** Engel & Grimaldi, 2005 is poorly preserved, it does not place consistently among stem Formicidae. †*S. **freyi*** Wilson *et al*., 1967, on the other hand, clusters with the †*G. orientalis* species group with diminishing support (50%C: 0.39 BPP; 30%C: 0.32 BPP; 10%C: 0.31 BPP; 5%C: 0.26 BPP; 0%C: 0.13 BPP).
•• **†Brownimeciinae** Bolton, 2003. Monotypic. Sister to crown Formicidae, forming Antennoclypeata. For support, see Fig. 17. Note that support is interrupted by the unplaceable male-based species of †*Baikuris*.
•• **Crown Formicidae.** See Fig. 17 for support.
••• **Crown Formicidae, *incertae sedis* to subfamily.** Five genera are considered *incertae sedis* among subfamilies and higher clades of crown Formicidae: †*Afromyrma* Dlussky *et al*., 2004, †*Afropone* Dlussky *et al*., 2004, †*Canapone* Dlussky, 1999, †*Cariridris* Brandão & Martins-Neto, 1990, and †*Cretopone* Dlussky, 1975. **†*Afromyrma petrosa*** Dlussky *et al*., 2004 clusters with Myrmeciinae (10%C: 0.77 BPP; 5%C: 0.46 BPP; 0%C: 0.49 BPP). **†*Afropone oculata*** Dlussky *et al*., 2004 clusters within crown Formicidae with meaningful support, but otherwise is unplaceable. †*Afropone **orapa*** Dlussky *et al*., 2004 is unplaceable among total Formicidae due to its incompleteness but is retained in the genus until further evidence is adduced. **†*Canapone dentata*** Dlussky, 1999 clusters with Poneria (40%C: 0.58 BPP; 30%C: 0.69 BPP; 10%C: 0.61 BPP; 5%C: 0.38 BPP; 0%C: 0.39 BPP), despite posterolateral cranial lobes which are suggestive of Ectatomminae. **†*Cariridris bipetiolata*** Brandão & Martins-Neto, 1990 clusters very weakly within crown Formicidae, but is well-supported within Formicoidea and total Formicidae; see Ampulicidae in Section 3.3 above for support values. **†*Cretopone magna*** Dlussky, 1975
••• **Crown Formicidae: Ponerinae** Lepeletier de Saint-Fargeau, 1835. One species is recovered in this subfamily, an undescribed species from burmite.
••• **Crown Formicidae: Dolichoderinae** Forel, 1878. Two species are maximally recovered with Dolichoderinae, †*Chronomyrmex medicinehatensis* McKellar *et al*., 2013 and an undescribed species from burmite. **†*Chronomyrmex*** McKellar *et al*., 2013 is supported as sister to Dolichoderinae (50%C: 0.88 BPP; 40%C: 0.91 BPP; 30%C, 10%C: 0.92 BPP; 5%C: 0.90 BPP; 0%C: 0.93 BPP).
••• **Crown Formicidae: Formicinae** Latreille, 1809. Four unexpected species are associated with Formicinae based on present analyses, and two are maximally recovered with the subfamily, †*Kyromyrma neffi* Grimaldi & Agosti, 2000 and an undescribed species from burmite. **†*Kyromyrma*** Grimaldi & Agosti, 2000 is supported as a crown member of Formicinae (60%C, 50%C, 40%C: 1.0 BPP; 30%C: 0.90 BPP; 10%C: 0.83 BPP). At the highest data completeness, †*Kyromyrma* is recovered as a member of Lasiini (50%C: 0.74 BPP), at lower levels it is sister to the remainder of the subfamily to the exclusion of *Myrmelachista* Roger, 1863. **†*Cananeuretus occidentalis*** Engel & Grimaldi, 2005 are recovered as sister to the crown Formicinae (40%C: 0.99 BPP; 30%C: 0.98 BPP; 10%C: 0.83 BPP; 5%C: 0.80 BPP; 0%C: 0.66 BPP), probably due to the lack of biseriate dentition which otherwise characterizes most Dolichoderomorpha. **†*Dolichomyrma longiceps*** Dlussky, 1975, originally attributed to †Armaniinae then excluded from the Formicidae by Borysenko (2017), is supported at a member of the total Formicidae to the exclusion of its apparent sistergroup, †*Cananeuretus* (10%C: 0.83 BPP; 5%C: 0.80 BPP; 0%C: 0.66 BPP). †*Dolichomyrma **latipes*** Dlussky, 1975 is fragmentary, represented by the gaster, propleurae, ventral head, and leg remnants, and is unplaced in the formicid polytomy at the < 5% completeness level; it is retained in †*Dolichomyrma* however, as it was recovered from the same formation as †*D. longiceps*. Finally, the poorly preserved putative dolichoderine †*Eotapinoma macalpini* Dlussky, 1999 clusters with Formicinae, again to the exclusion of †*Cananeuretus* (30%C: 0.90 BPP; 10%C: 0.83 BPP; 5%C: 0.80 BPP; 0%C: 0.66 BPP).

## Part III: Synopsis of Family-group Classification

### Overview

In this section, the family-group classification used here for the Aculeata, Formicoidea, Evanioidea, and Trigonaloidea, is presented. Because no catalog is available which includes authority and dates for all family-group names of the Aculeata, each suprageneric name listed in the synopsis includes a reference. This effort benefitted from the systematic literature and particularly the catalogs of Azevedo *et al*. (2018), Pulawski (2020), Engel (2005a), Bolton (2021), Ward *et al*. (2021), and personal communications. Mistakes should be attributed to the lead author of the present work. Because the ranking espoused here combines formal names governed by the ICZN and informal names, taxa with regulated rank are provided with taxon authorities to convey priority; all regulated names are set in bold-face font, with superfamilies underlined. Overall, the hierarchy is asymmetrical, which represents the recovered diversification pattern. Notes clarifying taxon delimitation are provided after the synopsis. For clades which end in “-ines”, we suggest the pronunciation /aCniCs/ (“eye-knees” [letter symbols in forward slashes from Oxford English Dictionary]). Taxa which are diagnosed with estimated synapomorphies or with special remarks in Results Part IV are indicated with an asterisk (*).

### Section 1: Synopsis

**<u>X.</u>** Superfamily **<u>Evanioidea</u>**\* Latreille, 1802. [***Note 1.***]

**X.1.** Stem, paraphyletic, or *incertae sedis* groups:

**X.A.** Family **†Anomopterellidae** Rasnitsyn, 1975.

**X.B.** Family **†Praeaulacidae** Rasnitsyn, 1972. [*Paraphyletic!*]

**X.C.** Family **†Vectevaniidae** Li *et al*., 2018.

**X.#.#.** Subfamily **†Nevaniinae** Zhang & Rasnitsyn, 2007.

X.2. Crown clade Evanioidea*.

X.2.1. Clade Aulaciformes* Grimaldi & Engel, 2005. [***Note 2.***]

X.2.#. *Incertae sedis* in Aulaciformes:

**X.#.#.** Subfamily †**Hyptiogastritinae** Engel, 2006.

**X.#.#.** Subfamily †**Kotujellitinae** Rasnitsyn, 1975.

**X.#.#.#.** Tribe †**Archeofoenini** Engel, 2017.

**X.#.#.#.** Tribe †**Electrofoenini** Cockerell, 1917.

X.2.1. †Baissidae–Gasteruptiidae clade.

**X.D.** Family †**Baissidae** Rasnitsyn, 1975. [*Paraphyletic! **Note 3.***]

X.2.1.1. Aulacidae–Gasteruptiidae Clade *. [***Note 4.***]

**X.E.** Family **Aulacidae** Shuckard, 1841.

**X.F.** Family **Gasteruptiidae** Ashmead, 1900.

**X.F.A.** Subfamily **Hyptiogastrinae** Crosskey, 1953.

**X.F.B.** Subfamily **Gasteruptiinae** Ashmead, 1900.

X.2.2. Clade Evaniiformes* Grimaldi & Engel, 2005. [***Note 5.***]

**X.D.** Family †**Othniodellithidae\*** Engel & Huang, 2016b.

**X.E.** Family **Evaniidae\*** Latreille, 1802. [***Note 6.***]

X’. Clade Trigaculeata*. [***Note 7*.**]

**<u>Y.</u>** Superfamily **<u>Trigonaloidea</u>\*** Cresson, 1887. [***Note 8*.**]

Y.#. *Incertae sedis* in superfamily:

**Y.A.** Family †**Andreneliidae** Rasnitsyn & Martinez-Delclòs, 2000. **Superfam. trans.** [***Note 9.***]

Y.1. †Maimetshidae–Trigonalidae clade*.

**Y.B.** Family **Trigonalidae\*** Cresson, 1887. [***Note 10.***]

**Y.B.A.** Subfamily **Orthogonalinae** Carmean & Kimsey, 1998.

**Y.B.A.** Subfamily **Trigonalinae** Cresson, 1887.

**Y.B.A.A.** Tribe **Nomadinini** Cameron, 1899.

**Y.B.A.B.** Tribe **Trigonalini** Cresson, 1887.

**Y.C.** Family †**Maimetshidae\*** Rasnitsyn, 1975.

**Y.C.A.** Subfamily †**Maimetshinae** Rasnitsyn, 1975.

**Y.C.A.A.** Tribe †**Ahiromaimetshini** Engel, 2016.

**Y.C.A.B.** Tribe †**Maimetshini** Rasnitsyn, 1975.

**Y.C.A.** Subfamily †**Zorophratrinae** Engel, 2016.

**<u>#.</u>** Superfamily **<u>†Panguoidea</u>** Li *et al*., 2020a. [***Note 11.***]

**#.#.** Family **†Panguidae** Li *et al*., 2020a.

Y’. Total clade Aculeata Latreille, 1802.

Y’.1. *Incertae sedis* to superfamily or higher clade:

**<u>Z.</u>** Superfamily **<u>†Bethylonymoidea</u>\*** Rasnitsyn, 1975. [***Note 12.***]

**Z.#.** Family **†Bethylonymidae\*** Rasnitsyn, 1975.

**Z.#.#.** Subfamily †**Bethylonyminae** Rasnitsyn, 1975 **stat. nov.** [***Notes 13, 14.***]

#. Unplaced to superfamily:

**#.#.** Family **†Falsiformicidae\*** Rasnitsyn, 1975. **Superfam. trans.** [***Note 15.***]

**<u>A.</u>** Superfamily **<u>Chrysidoidea</u>\*** Latreille, 1802.

A.1. †Plumalexiidae–Plumariidae clade*. [***Note 16.***]

**A.A.** Family **Plumariidae** Bischoff, 1914.

**A.B.** Family **†Plumalexiidae** Brothers, 2011. [***Note 17.***]

A.2. Bethylidae–Chrysididae clade*.

A.2.1. †Chrysobythidae–Bethylidae clade*.

**A.C.** Family **†Chrysobythidae\*** Melo & Lucena, 2019.

**A.D.** Family †**Holopsenellidae** Engel *et al*., 2016a. [***Note 18.***]

**A.E.** Family **Bethylidae\*** Haliday, 1839. [***Note 19.***]

A.E.#. Extant subfamilies:

**A.E.A.** Subfamily **Bethylinae** Haliday, 1839.

**A.E.B.** Subfamily **Epyrinae** Kieffer, 1914.

**A.E.C.** Subfamily **Lancepyrinae** Azevedo & Azar, 2012.

**A.E.D.** Subfamily **Mesitiinae** Kieffer, 1914.

**A.E.E.** Subfamily **Pristocerinae** Mocsáry, 1881.

**A.E.F.** Subfamily **Scleroderminae** Kieffer, 1914.

A.E.#. Extinct subfamily:

**A.E.G.** Subfamily †**Protopristocerinae** Nagy, 1974.

A.2.2. Chrysididae–Scolebythidae clade*.

**A.E.** Family **Chrysididae\*** Latreille, 1802. [***Note 20*.**]

A.E.#. *Incertae sedis* in family:

**A.E.D.** Subfamily **Loboscelidiinae** Ashmead, 1903b.

**A.E.A.** Subfamily **Chrysidinae** Latreille, 1802.

A.E.1. Amiseginae–Cleptinae clade.

**A.E.B.** Subfamily **Amiseginae** Krombein, 1957.

**A.E.C.** Subfamily **Cleptinae** Latreille, 1802.

**A.F.** Family **Scolebythidae\*** Evans, 1963.

**A.F.A.** Subfamily **Scolebythinae** Evans, 1963.

**A.F.B.** Subfamily **Pristapenesiinae** Engel *et al*., 2013.

A’. Clade Dryinaculeata.

**<u>B.</u>** Superfamily **<u>Dryinoidea</u>\*** Haliday, 1833. [***Note 21.***]

**B.A.** Family **Sclerogibbidae\*** Ashmead, 1902c. [***Note 22.***]

**B.A.A.** Subfamily †**Sclerogibbodinae** Engel & Grimaldi, 2006a.

**B.A.B.** Subfamily **Sclerogibbinae** Ashmead, 1902c.

B.A’. Clade Dryinomorpha*. [***Note 23.***]

**B.B.** Family **Embolemidae\*** Förster, 1856.

**B.C.** Family **Dryinidae\*** Haliday, 1833. [***Note 24.***]

B.C.1. Extant subfamilies:

**B.C.A.** Subfamily **Anteoninae** Perkins, 1912.

**B.C.B.** Subfamily **Aphelopinae** Perkins, 1912.

**B.C.C.** Subfamily **Bocchinae** Richards, 1939.

**B.C.D.** Subfamily **Dryininae** Haliday, 1833.

**B.C.E.** Subfamily **Gonatopodinae** Kieffer, 1906 in Kieffer & Marshall, 1906.

**B.C.F.** Subfamily **Thaumatodryininae** Perkins, 1905.

B.C.2. Extinct subfamilies:

**B.C.G.** Subfamily †**Archaeodryininae** Olmi *et al*., 2020.

**B.C.H.** Subfamily †**Burmadryininae** Olmi *et al*., 2014.

**B.C.I.** Subfamily †**Palaeoanteoninae** Olmi & Bechly, 2001.

**B.C.J.** Subfamily †**Ponomarenkoinae** Olmi, 2010.

**B.C.K.** Subfamily †**Protodryininae** Olmi & Guglielmino, 2012.

**B.C.L.** Subfamily †**Raptodryininae** Olmi *et al*., 2020.

B’. Clade Vespaculeata* / Vespiformes*.

B’.1. Clade Vespoides*.

**<u>C.</u>** Superfamily **<u>Vespoidea</u>\*** Laicharting, 1781.

C.#. *Incertae sedis* in superfamily:

C.#. †*Prosphex* genus group*. [*Warning! **Note 25.***]

**C.A.** Family **Rhopalosomatidae\*** Ashmead, 1896b. [***Note 26.***]

**C.A.A.** Subfamily **Rhopalosomatinae**\* Ashmead, 1896b.

**C.A.A.** Subfamily **Olixoninae** Engel, 2008.

**C.B.** Family **Vespidae**\* Laicharting, 1781.

C.B.#. *Incertae sedis*, likely stem-vespid subfamilies:

**C.B.#.** Subfamily †**Priorvespinae** Carpenter & Rasnitsyn, 1990.

**C.B.#.** Subfamily †**Protovespinae** Perrard & Carpenter, 2017.

C.B. Crown clade Vespidae*. [***Notes 27, 28.***]

**C.B.A.** Subfamily **Stenogastrinae** Bequaert, 1918. [***Note 29*.**]

C.B.1. Core clade Vespidae*.

C.B.1.1. Masarinae–Euparagiinae clade.

**C.B.B.** Subfamily **Euparagiinae** Ashmead, 1902b.

**C.B.C.** Subfamily **Gayellinae** Bradley, 1922.

**C.B.D.** Subfamily **Masarinae** Latreille, 1802.

C.B.1.2. Eumeninae–Vespinae clade*.

**C.B.E.** Subfamily **Eumeninae** Leach, 1815.

C.B.1.2.1. Zethinae–Vespinae clade.

C.B.1.2.1.#. Relationships within clade uncertain:

**C.B.F.** Subfamily **Raphiglossinae** Ashmead, 1902.

**C.B.G.** Subfamily **Zethinae** de Saussure, 1855.

C.B.1.2.1.1. Polistinae–Vespinae clade.

**C.B.H.** Subfamily **Polistinae** Lepeletier de Saint-Fargeau, 1835.

**C.B.I.** Subfamily **Vespinae** Laicharting, 1781.

**<u>C’.</u>** Superfamily **<u>Pompiloidea</u>**\* Latreille, 1804. [***Note 30.***]

**D.A.** Family **Sierolomorphidae\*** Krombein, 1951.

D.A’. Clade Tiphiopompiloides*.

D.A.1. Clade Tiphiiformes*.

**D.B.** Family **Tiphiidae\*** Leach, 1815. [***Note 31.***]

D.B.#. *Incertae sedis* in family:

**D.B.#.** Subfamily †**Thanatotiphiinae** Engel *et al*., 2009. [***Note 32.***]

D.B.1. Crown Tiphiidae clade.

**D.B.A.** Subfamily **Brachycistidinae** Malloch, 1926.

**D.B.B.** Subfamily **Tiphiinae** Leach, 1815.

D.B’. Clade Thynnimorpha*.

**D.C.** Family **Chyphotidae** Ashmead, 1896a*.

**D.C.A.** Subfamily **Chyphotinae** Ashmead, 1896a.

**D.C.B.** Subfamily **Typhoctinae** Schuster, 1949.

**D.D.** Family **Thynnidae** Erichson, 1842.

**D.D.A.** Subfamily **Anthoboscinae** Turner, 1912.

**D.D.B.** Subfamily **Diamminae** Turner, 1907.

**D.D.C.** Subfamily **Methochinae** André, 1899.

**D.D.D.** Subfamily **Myzininae** Ashmead, 1903a.

**D.D.E.** Subfamily **Thynninae** Erichson, 1842.

D.A.2. Core Pompiloidea clade.

D.A.2.1. Clade Pompiliformes*.

**D.E.** Family †**Burmusculidae** Zhang *et al*., 2018.

**D.F.** Family **Pompilidae** Latreille, 1804.

**D.F.A.** Subfamily **Ceropalinae** Radoszkowski, 1888.

**D.F.B.** Subfamily **Ctenocerinae** Arnold, 1934.

**D.F.C.** Subfamily **Notocyphinae** Fox, 1895.

**D.F.D.** Subfamily **Pepsinae** Lepeletier de Saint-Fargeau, 1845.

**D.F.E.** Subfamily **Pompilinae** Latreille, 1804.

D.A.2.2. Clade Mutilliformes*.

**D.G.** Family †**Bryopompilidae** Rodriguez *et al*., 2016.

D.G.1. Crown clade Mutilliformes*.

**D.G.** Family **Mutillidae\*** Latreille, 1802. [***Note 33.***]

**D.G.A.** Subfamily **Myrmosinae** Fox, 1895.

D.G.A.1. Core clade Mutillidae*.

**D.G.B.** Subfamily **Dasylabrinae** Brothers & Lelej, 2017.

**D.G.C.** Subfamily **Mutillinae** Latreille, 1802.

**D.G.D.** Subfamily **Myrmillinae** Bischoff, 1920.

**D.G.E.** Subfamily **Rhopalomutillinae** Schuster, 1949.

**D.G.F.** Subfamily **Pseudophotopsinae** Bischoff, 1920.

**D.G.G.** Subfamily **Sphaeropthalminae** Schuster, 1949.

**D.G.H.** Subfamily **Ticoplinae** Nagy, 1970.

**D.H.** Family **Sapygidae\*** Latreille, 1810.

**D.H.A.** Subfamily †**Cretosapyginae** Bennett & Engel, 2005.

D.H.1. Crown clade Sapygidae*.

**D.H.B.** Subfamily **Fedtschenkiinae** André, 1899.

**D.H.C.** Subfamily **Sapyginae** Latreille, 1810.

B’.2. Clade Scolioides*.

**<u>E.</u>** Superfamily **<u>Scolioidea</u>\*** Latreille, 1802.

**E.A.** Family **Bradynobaenidae\*** de Saussure, 1892. [***Note 34.***]

**E.A.A.** Subfamily **Apterogyninae** André, 1899.

**E.A.B.** Subfamily **Bradynobaeninae** de Saussure, 1892.

**E.B.** Family **Scoliidae\*** Latreille, 1802. [***Note 35.***]

E.B.#. *Incertae sedis* in family, monophyly uncertain:

**E.B.#.** Subfamily †**Archaeoscoliinae** Rasnitsyn, 1993. [*Warning!* ***Note 36.***]

E.B.1. Crown clade Scoliidae*.

**E.B.A.** Subfamily **Proscoliinae** Rasnitsyn, 1977.

**E.B.B.** Subfamily **Scoliinae\*** Latreille, 1802.

**E.B.B.A.** Tribe **Campsomerini**\* Betrem in Betrem & Bradley, 1972. [***Note 37*.**]

**E.B.B.B.** Tribe **Scoliini\*** Latreille, 1802.

E’. Clade Formicapoidina*.

**<u>F.</u>** Superfamily **<u>Formicoidea</u>\*** Latreille, 1809. [*See below*.]

**<u>G.</u>** Superfamily **<u>Apoidea</u>\*** Latreille, 1802. [***Notes 38, 39*.**]

G.#. Stem groups or otherwise *incertae sedis* in superfamily:

**G.#.** Family †**Allommationidae** Rosa & Melo, 2021. [*Monogeneric.*]

**G.#.** Family †**Angarosphecidae\*** Rasnitsyn, 1975. [*Paraphyletic! **Note 40.***]

**G.#.** Family †**Spheciellidae** Rosa & Melo, 2021. [*Monogeneric.*]

**G.#.#.** Subfamily †**Burmastatinae** Antropov, 2000a. [*Monogeneric.*]

**G.#.#.** Subfamily †**Apodolichurinae** Antropov, 2000a. [*Monogeneric.*] [***Note 41.***]

**G.#.#.** Subfamily †**Mendampulicinae** Antropov, 2000a. [*Monogeneric.*]

**G.A.** Family **Ampulicidae\*** Shuckard, 1840.

G.A.#. *Incertae sedis* in family.

**G.A.#.#.** Tribe †**Cretampulicini** Antropov, 2000. [*Monogeneric.*]

G.A.1. Crown clade Ampulicidae*:

**G.A.A.** Subfamily **Ampulicidae** Shuckard, 1840. [***Note 42.***]

**G.A.A.A.** Tribe **Ampulicini** Shuckard, 1840.

**G.A.B.** Subfamily **Dolichurinae** Dahlbom, 1842.

**G.A.B.A.** Tribe **Aphelotomini** Ohl & Spahn, 2009. [***Note 43.***]

**G.A.B.B.** Tribe **Dolichurini** Shuckard, 1840.

G.1. Core Apoidea clade*. [***Notes 44.***]

**G.B.** Family †**Cirrosphecidae** Antropov, 2000a. [***Note 45.***]

G.B.A.1. Placement tested in the present study:

**G.B.A.** Subfamily †**Cirrosphecinae** Antropov, 2000a.

G.B.A.#. Placement untested in the present study:

**G.B.A.** Subfamily †**Glenocephalinae** Rosa & Melo, 2021.

G.2. *Incertae sedis* within core Apoidea clade:

**G.C.** Family **Heterogynaidae** Nagy, 1969. [***Note 46.***]

G.C.#. Placement in family untested:

**G.C.#.** Subfamily **†Ptilocosminae** Rosa & Melo, 2021.

**G.C.A.** Subfamily **Heterogynaidae** Nagy, 1969.

G.2’. Crabronidae–Anthophila clade*.

G.3. Crabronidae–Sphecidae clade*.

**G.D.** Family **Crabronidae\*** Latreille, 1802.

G.D.#. *Incertae sedis* in family:

**G.D.#.#.** Tribe †**Discoscapini** Poinar, 2020. [*Monogeneric*.] [***Note 47.***]

**G.D.#.#.** Tribe †**Protomicroidini** Antropov, 2010. [*Monogeneric*.]

**G.D.A.** Subfamily **Crabroninae** Latreille, 1802.

G.D.A.1. Crabronine clade.

**G.D.A.A.** Tribe **Crabronini** Latreille, 1802.

**G.D.A.A.** Subtribe **Anacrabronina** Ashmead, 1899.

**G.D.A.B.** Subtribe **Crabronina** Latreille, 1802.

**G.D.A.B.** Tribe **Oxybelini** Leach, 1815.

G.D.A.2. Larrine clade.

**G.D.A.C.** Tribe **Bothynostethini** Fox, 1895.

**G.D.A.C.A.** Subtribe **Bothynostethina** Fox, 1895.

**G.D.A.C.B.** Subtribe **Scapheutina** Menke, 1968.

**G.D.A.D.** Tribe **Gastrosericini** André, 1886.

**G.D.A.E.** Tribe **Larrini** Latreille, 1810.

**G.D.A.F.** Tribe **Miscophini** Fox, 1895. [*Paraphyletic*.]

**G.D.A.G.** Tribe **Palarini** Schrottky, 1909.

**G.D.A.H.** Tribe **Trypoxylonini** Lepeletier de Saint-Fargeau, 1845.

**G.D.B.** Subfamily **Dinetinae** Fox, 1895. [*Monogeneric*.]

**G.E.** Family **Mellinidae** Latreille, 1802.

**G.E.A.** Subfamily **Mellininae** Latreille, 1802.

**G.E.A.A.** Tribe **Mellinini** Latreille, 1802. [*Monogeneric*.]

**G.E.A.B.** Tribe **Xenosphecini** Parker, 1966. [*Monogeneric*.]

**G.F.** Family **Sphecidae\*** Latreille, 1802.

G.F.1. Sphecine clade.

G.F.#. *Incertae sedis* within sphecine clade:

**G.F.#.#.** Tribe **Stangeellini** Bohart & Menke, 1976.

**G.F.A.** Subfamily **Ammophilinae** André, 1886. [*No tribal classification*.]

**G.F.B.** Subfamily **Sphecinae** Latreille, 1802.

**G.F.B.A.** Tribe **Prionychini** Bohart & Menke, 1963.

**G.F.B.B.** Tribe **Sphecini** Latreille, 1802.

G.F.1’. Sceliphrine clade.

**G.F.D.** Subfamily **Chloriontinae** Fernald, 1905. [*Monogeneric*. *No tribes.*]

**G.F.E.** Subfamily **Sceliphrinae** Ashmead, 1899b.

**G.F.E.A.** Tribe **Podiini** de Saussure, 1892.

**G.F.E.B.** Tribe **†Protosceliphrini** Antropov, 2014. [*Monogeneric*.]

**G.F.E.C.** Tribe **Sceliphrinae** Ashmead, 1899b.

G.3’. Bembicidae–Anthophila clade*.

**G.G.** Family **Astatidae** Lepel. de Saint-Fargeau, 1845. [***Note 48.***]

G.G.#. *Incertae sedis* in family:

**G.G.#.#.** Tribe **†Cretastatini** Rosa & Melo, 2021. [*Monogeneric*.]

**G.G.A.** Subfamily **Astatinae** Lepeletier de Saint-Fargeau, 1845.

**G.H.** Family **Entomosericidae** Dalla Torre, 1897. [***Note 49.***]

**G.H.A.** Subfamily **Entomosericinae** Dalla Torre, 1897.

**G.H.A.A.** Tribe **Entomosericini** Dalla Torre, 1897. [*Monogeneric*.]

**G.I.** Family **Eremiaspheciidae** Menke, 1967.

**G.I.A.** Subfamily **Eremiaspheciinae** Menke, 1967.

**G.I.A.A.** Tribe **Eremiaspheciini** Menke, 1967. [*Monogeneric*.]

**G.I.A.B.** Tribe **Laphyragogini** Bohart & Menke, 1976. [*Monogeneric*.]

**G.J.** Family **Bembicidae\*** Latreille, 1802. [***Note 50.***]

G.J.#. *Incertae sedis* in family:

**G.J.#.#.** Tribe †**Megaroliini** Rosa & Melo, 2021. [*Monogeneric*.]

**G.J.#.#.** Tribe †**Pristinopterini** Rosa & Melo, 2021. [*Monogeneric*.]

**G.J.A.** Subfamily **Bembicinae** Latreille, 1802.

**G.J.A.A.** Tribe **Bembicini** Latreille, 1802.

**G.J.A.B.** Tribe **Helocausini** Handlirsch, 1925.

G.J.A.#. *Incertae sedis* in subfamily:

**G.J.A.#.#.** Subtribe **Exeirina** Dalla Torre, 1897.

**G.J.A.#.#.** Subtribe **Gorytina** Lepeletier de Saint Fargeau, 1845.

**G.J.A.#.#.** Subtribe **Handlirschiina** Nemkov & Lelej, 1996.

**G.J.A.#.#.** Subtribe **Spheciina** Nemkov & Ohl, 2011. [*Paraphyletic*.]

**G.J.A.#.#.** Subtribe **Stictiellina** Bohart & Horning, 1971.

**G.J.A.#.#.** Subtribe **Stizina** Costa, 1859.

**G.J.B.** Subfamily **Nyssoninae** Latreille, 1804.

**G.J.B.B.** Tribe **Alyssontini** Dalla Torre, 1897.

**G.J.B.B.** Tribe **Nyssonini** Latreille, 1804.

**G.J.B.B.A.** Subtribe **Nurseina** Nemkov & Lelej, 2013.

**G.J.B.B.B.** Subtribe **Nyssonina** Latreille, 1804.

G.4. Pemphredonidae–Anthophila clade*.

G.#. *Incertae sedis* in Pemphredonidae–Anthophila clade:

**G.K.** Family **Psenidae** Costa, 1858.

**G.K.A.** Subfamily **Psenidae** Costa, 1858. [*Clade weakly supported.*]

**G.K.A.A.** Tribe **Odontosphecini** Menke, 1967. [*Monogeneric*.]

**G.K.A.B.** Tribe **Psenini** Costa, 1858.

G.K.#. *Incertae sedis* in family, placements untested:

**G.K.#.#.** Tribe †**Rasnitsynapini** Antropov, 2011. [*Monogeneric*.]

**G.K.#.#.#.** Subtribe †**Helosericina** Rosa & Melo, 2021. [*Monogeneric*.]

**G.K.#.#.#.** Subtribe †**Odontosericina** Rosa & Melo, 2021. [*Monogeneric*.]

G.5. Pemphredonidae–Philanthidae clade*.

**G.L.** Family **Pemphredonidae** Dahlbom, 1835.

**G.L.A.** Subfamily **Pemphredoninae** Dahlbom, 1835.

**G.L.A.A.** Tribe **Pemphredonini** Dahlbom, 1835.

**G.L.A.B.** Tribe **Spilomenini** Menke, 1989.

**G.L.A.C.** Tribe **Stigmini** Bohart & Menke, 1976.

G.L.A.#. *Incertae sedis* in Pemphredoninae, may belong in other family:

**G.L.A.#.** Tribe †**Palangini** Antropov, 2011. [*Monogeneric*.]

**G.M.** Family **Philanthidae\*** Latreille, 1802.

**G.M.A.** Subfamily **Philanthinae** Latreille, 1802.

**G.M.A.A.** Tribe **Aphilanthopini** Bohart, 1966.

**G.M.A.B.** Tribe **Cercerini** Lepeletier de Saint-Fargeau, 1845.

**G.M.A.C.** Tribe **Philanthini** Latreille, 1802.

**G.M.A.C.A.** Subtribe **Philanthina** Latreille, 1802.

**G.M.A.C.B.** Subtribe **Philanthinina** Menke, 1967.

**G.M.A.D.** Tribe **Pseudoscoliini** Menke, 1967.

G.5’. Ammoplanidae–Anthophila clade:

**G.N.** Family **Ammoplanidae** Evans, 1959. [***Note 51.***]

G.6. Clade Anthophila* Latreille, 1804. [***Note 52.***]

G.7. Extinct families:

**G.O.** Family †**Melittosphecidae** Poinar & Danforth, 2006. [*Monogeneric*. ***Note 53.***]

**G.P.** Family †**Paleomelittidae** Engel, 2001. [*Monogeneric*.]

G.7’. Extant families:

**G.Q.** Family **Melittidae** Schenck, 1860.

G.8. Clade 1.

**G.R.** Family **Andrenidae** Latreille, 1802.

G.9. Clade 2.

**G.S.** Family **Halictidae** Thomson, 1869.

G.10. Clade 3.

G.11. Clade 4.

**G.T.** Family **Stenotritidae** Cockerell, 1934.

**G.U.** Family **Colletidae** Lepeletier de Saint-Fargeau, 1841.

G.11’. Clade 5.

**G.V.** Family **Apidae** Latreille, 1802.

**G.W.** Family **Megachilidae** Latreille, 1802.

**<u>F.</u>** Superfamily **<u>Formicoidea</u>**\* Latreille, 1809. [***Note 54.***]

**F.A.** Family **†@@@idae**\* **fam. nov.** [***Note 55.***]

**F.B.** Family **Formicidae**\* Latreille, 1809. [“Total clade Formicidae”. ***Note 56.***]

F.B.#. *Incertae sedis* in family:

**F.B.A.** Subfamily **†Armaniinae\*** Dlussky, 1893.

**F.B.A.A.** Tribe **†Armaniini\*** Dlussky, 1893 **stat. nov.**

F.B.1. Clade †Sphecomyrmines*.

**F.B.B.** Subfamily **†Haidomyrmecinae\*** Bolton, 2003.

**F.B.B.A.** Tribe **†Aquilomyrmecini trib. nov.**

**F.B.B.B.** Tribe **†Haidomyrmecini stat. nov.**

**F.B.C.** Subfamily **†Sphecomyrminae\*** Wilson & Brown, 1967.

**F.B.D.** Subfamily **†Zigrasimeciinae\*** Borysenko, 2017.

**F.B.D.A.** Tribe **†Zigrasimeciini\*** Borysenko, 2017.

F.B.2. Clade Antennoclypeata*.

**F.B.E.** Subfamily **†Brownimeciinae** Bolton, 2003.

F.B.3. Crown clade Formicidae*.

F.B.4. Clade Leptanillomorpha*.

**F.B.F.** Subfamily **Leptanillinae** Emery, 1910. [***Note 57.***]

**F.B.F.A.** Tribe **Opamyrmini** Boudinot & Griebenow **trib. nov.**

**F.B.F.B.** Tribe **Leptanillini** Emery, 1910.

**=** Tribe **Anomalomyrmini** Taylor in Bolton, 1990. **Syn. nov.**

**F.B.G.** Subfamily **Martialinae** Rabeling & Verhaagh, 2008.

F.B.4’. Clade Poneroformicines*.

F.B.5. Clade Poneria*. [***Note 58.***]

F.B.6. Clade Amblyoponomorpha*.

**F.B.H.** Subfamily **Amblyoponinae\*** Forel, 1893.

**F.B.I.** Subfamily **Apomyrminae** Dlussky & Fedoseeva, 1988.

**F.B.J.** Subfamily **Agroecomyrmecinae** Carpenter, 1930.

**F.B.J.A.** Tribe **Agroecomyrmecini** Carpenter, 1930.

**F.B.J.B.** Tribe **Ankylomyrmini** Bolton, 2003.

**F.B.K.** Subfamily **Paraponerinae** Emery, 1901.

**F.B.L.** Subfamily **Ponerinae\*** Lepeletier de Saint-Fargeau, 1835.

**F.B.L.A.** Tribe **Platythyreini** Emery, 1901.

**F.B.L.B.** Tribe **Ponerinae\*** Lepeletier de Saint-Fargeau, 1835.

**F.B.M.** Subfamily **Proceratiinae** Emery, 1895. [***Note 59*.**]

**F.B.M.A.** Tribe **Probolomyrmecini** Perrault, 2000.

**F.B.M.B.** Tribe **Proceratiini** Emery, 1895.

F.B.5’. Clade Doryloformicia*.

**F.B.N.** Subfamily **Dorylinae** Leach, 1815.

F.B.7. Clade Formicae*.

F.B.8. Clade Myrmechoderines*.

F.B.9. Clade Myrmeciomorpha*.

**F.B.O.** Subfamily **Myrmeciinae** Emery, 1877.

**F.B.O.A.** Tribe **Myrmeciini** Emery, 1877.

**F.B.O.B.** Tribe **Prionomyrmecini** Wheeler, 1915.

**F.B.P.** Subfamily **Pseudomyrmecinae\*** Smith, M. R., 1952.

F.B.9’. Clade Dolichoderomorpha*.

**F.B.Q.** Subfamily **Aneuretinae** Emery, 1913.

**F.B.Q.A.** Tribe **Aneuretinae** Emery, 1913.

**F.B.Q.B.** Tribe **†Pityomyrmecini** Wheeler, 1915.

**F.B.R.** Subfamily **Dolichoderinae\*** Forel, 1878.

F.B.R.1. Extant tribes:

**F.B.R.A.** Tribe **Bothriomyrmecini** Dubovikoff, 2005.

**F.B.R.B.** Tribe **Dolichoderini** Forel, 1878.

**F.B.R.C.** Tribe **Leptomyrmecini** Emery, 1913.

**F.B.R.E.** Tribe **Tapinomini** Emery, 1913.

F.B.R.#. Extinct tribes, *incertae sedis* in subfamily.

**F.B.R.D.** Tribe **†Miomyrmecini** Carpenter, 1930.

**F.B.R.F.** Tribe **†Zherichiniini** Dlussky, 1988.

F.B.8’. Clade Formicomyrmines*.

F.B.10. Clade Formicomorpha*.

**F.B.S.** Subfamily **†Formiciinae** Lutz, 1896.

**F.B.T.** Subfamily **Formicinae\*** Latreille, 1809. [***Note 60.***]

F.B.T.1. Lasiomorph grade, split 1:

**F.B.T.A.** Tribe **Myrmelachistini** Forel, 1912.

F.B.T.2. Lasiomorph grade, split 2 / lasioformicine clade:

**F.B.T.B.** Tribe **Lasiini** Ashmead, 1905.

F.B.T.3. Formicoform radiation, split 1:

**F.B.T.C.** Tribe **Melophorini** Forel, 1912.

F.B.T.4. Formicoform radiation, core clade:

F.B.T.5. Arboreal clade:

**F.B.T.F.** Tribe **Gesomyrmecini** Ashmead, 1905.

**F.B.T.I.** Tribe **Oecophyllini** Emery, 1895.

F.B.T.6. Unusual clade:

**F.B.T.D.** Tribe **Camponotini** Forel, 1878.

**F.B.T.H.** Tribe **Myrmoteratini** Emery, 1895.

F.B.T.#. Tribes with unclear relationships:

**F.B.T.E.** Tribe **Formicini** Latreille, 1809.

**F.B.T.G.** Tribe **Gigantiopini** Ashmead, 1905.

**F.B.T.J.** Tribe **Plagiolepidini** Forel, 1886.

**F.B.T.K.** Tribe **Santschiellini** Forel, 1917.

F.B.10’. Clade Myrmicomorpha*.

**F.B.U.** Subfamily **Ectatomminae\*** Emery, 1895. [***Note 61.***]

**F.B.U.A.** Tribe **Heteroponerini** Bolton, 2003.

**F.B.U.B.** Tribe **Ectatommini** Emery, 1895.

**F.B.V.** Subfamily **Myrmicinae\*** Lepeletier de Saint-Fargeau, 1835.

F.B.V.1. Clade 1 from Branstetter *et al*. (2017a).

**F.B.V.A.** Tribe **Myrmicini** Lepeletier de Saint-Fargeau, 1835.

**F.B.V.B.** Tribe **Pogonomyrmecini** Ward *et al*., 2015.

F.B.V.2. Clade 2 from Branstetter *et al*. (2017a).

**F.B.V.C.** Tribe **Stenammini** Ashmead, 1905.

F.B.V.3. Clade 3 from Branstetter *et al*. (2017a).

**F.B.V.D.** Tribe **Attini** Smith, F., 1858. [***Note 62.***]

F.B.V.4. Clade 4 from Branstetter *et al*. (2017a).

**F.B.V.E.** Tribe **Crematogastrini** Forel, 1893.

**F.B.V.F.** Tribe **Solenopsidini** Forel, 1893.

### Section 2: Notes

#### Evanioidea

***Note 1.*** The circumscription of the Evanioidea crown clade employed here follows the extended phylogenetic results (Results Part II). It approximates the Neoevanioides of Engel (2006), but with the exclusion of †Anomopterellidae and consideration of †Baissidae as *incertae sedis* in the group, rather than as part of the Aulaciformes. Here, the Aulaciformes includes Gasteruptiidae and Aulacidae, while the Evaniiformes is extended to include †Othniodellithidae in addition to Evaniidae. †Andreneliidae is here placed with Trigonaloidea based on our phylogenetic estimates. For the comparative purposes and to guide future research, the topological results of Jouault *et al*. (2021) are here summarized: †Praeaulacidae paraphyletic with respect to †Othniodellithidae; †Praeaulacidae + †Othniodellithidae (†Anomopterellidae ((†Andreneliidae, Evaniidae), (†Baissidae paraphyletic with respect to (Gasteruptiidae, Aulacidae)))).

***Note 2.*** †Hyptiogastritinae and †Kotujellitinae are considered to be subfamilies of Gasteruptiidae (*e.g.*, Engel 2017), but were not recovered as a clade with *Gasteruption* in the present study, thus are treated as *incertae sedis* to family. Likewise, †Archeofoenini is attributed to †Hyptiogastritinae and †Electrofoenini to the Aulacinae. The results of our study are not the final word on the matter, and we expect refinement of the higher classification of the group via further critical analysis, particularly with a broader taxon sampling. See Li *et al*. (2021) for an alternative classification.

***Note 3*.** In the present study, †Baissidae was supported as paraphyletic with respect to the Aulacidae–Gasteruptiidae clade, congruent with the results of Jouault *et al*. (2021).

***Note 4.*** Rasnitsyn & Öhm-Kuhnle (2021) synonymized Aulacidae with Gasteruptiidae, following Townes (1950). We prefer to retain the distinction between the two families pending concentrated revision of the Aulaciformes.

***Note 5.*** The Evaniiformes have been expanded to include †Othniodellithidae based on results from the present study. Jouault *et al*. (2021) recovered †Othniodellithidae within an otherwise monophyletic †Praeaulacidae.

***Note 6.*** Although the name †Cretevaniidae Rasnitsyn, 1975 is available for †*Cretevania*, no formal taxonomic rearrangement of the Evaniidae is undertaken here, as further analyses are desirable.

#### Trigaculeata

***Note 7.*** Recircumscription of Trigaculeata is probably necessary as there is some genomic signal for the relatedness of Ceraphronoidea with the Aculeata (Branstetter *et al*. 2017). Furthermore, we did not sample Stephanoidea, which are plausibly related to the Trigaculeata (Peters *et al*. 2017).

***Note 8.*** Recently Rasnitsyn & Öhm-Kuhnle (2021) placed †*Albiogonalys* in †Praeaulacidae, arguing that the venation is distinctly different from Trigonalidae, and remarking that the metasoma is attached high on the propodeum. Examining the available figures, we were unable to come to a conclusion about the metasomal attachment. We tentatively retain †*Albiogonalys* as *incertae sedis* in the Trigonaloidea based on our present analyses. We hope that the fossil can be µ-CT scanned to resolve the uncertainty clouding this fossil.

***Note 9.*** We tentatively place †Andreneliidae in the Trigonaloidea based on our present analyses. Detailed restudy of †*Andrenelia* is necessary.

***Note 10.*** See Carmean & Kimsey (1998) for taxonomic and morphological synopsis of extant members of Trigonalidae.

***Note 11*.** †Panguoidea was recently defined by Li *et al*. (2020a) for a singular new species, †*P. yuangu*. We place the superfamily in Trigaculeata based on its overall phenotypic similarity and hypothesize that it is sister to the Aculeata including †Bethylonymoidea but consider it unplaced pending phylogenetic hypothesis testing. See Note 2 of the Trigaculeata diagnosis below for further information.

#### Total clade Aculeata

***Note 12.*** †Bethylonymidae is *incertae sedis* in the total clade Aculeata based on the present analyses.

***Note 13.*** †Bethylonyminae is recognized here to distinguish taxa with well-supported placement in the group.

***Note 14.*** We provide †*Renymus* **nom. nov.** for †*Allogaster* Ren, a junior homonym of *Allogaster* Thomson, 1864 of the Cerambycidae. †*Renymus* clustered with †Bethylonymidae in our analyses.

***Note 15.*** †Falsiformicidae was originally considered to be the sister of the Formicidae (Rasnitsyn 1975) but was suggested to be closer to Bethylidae in a recent published abstract (Perrichot *et al*. 2014). The family is here treated as *incertae sedis* in the total clade Aculeata based on our analyses. Our results suggest close relationship of †Falsiformicidae and †*Burmasphex*, a putative apoid (Melo & Lucena 2018). We tentatively retain †*Burmasphex* and other (unnamed) fossil taxa as close to †Falsiformicidae based on our results and pending further study and hypothesis testing. See Notes 3–5 of the familial diagnosis below for further detail.

***Note 16.*** The placements of Plumariidae and †Plumalexiidae are uncertain, but they are grouped together here based on the present analyses. Brothers & Melo (2021) recovered this grouping in their recent parsimony analysis of the Chrysidoidea with select outgroups.

***Note 17.*** See also Rasnitsyn & Brothers (2020) for further information about the †Plumalexiidae. ***Note 18.*** †Holopsenellidae, formerly a subfamily of Bethylidae, was recently elevated to family status by Lepeco & Melo (2022), who considered the group *incertae sedis* in the Vespaculeata (“Aculeata *s. str.*”) based on internalization of the seventh metasomal segment in females and the proximal constriction of the third metasomal segment, features which were not previously appreciated. Because the family’s constituent genus, †*Holopsenella*, consistently clustered with Bethylidae in the present study, we tentatively retain the family in Chrysidoidea as presently defined. Revision of its scoring and reanalysis are recommended.

***Note 19*.** The Bethylidae were insufficiently sampled in the present study to provide internal structure.

***Note 20.*** The sistergroup relationship of Amiseginae + Cleptinae, recovered by Pauli *et al*. (2021), is recognized here, although this clade was not detectable with the data used in the present study. Loboscelidiinae remains *incertae sedis* in the family, but it was suggested that the subfamily may be sister to Amiseginae. Further study is necessary and will, according to Pauli *et al*. (2021), require considerable revision at the generic level.

***Note 21.*** Dryinoidea = Dryiniformes Bennett & Engel, 2005.

***Note 22.*** Argaman (1988) proposed a suprageneric classification of Sclerogibbidae, recognizing three subfamilies (Caenosclerogibbinae, Probethylinae) and two tribes within Sclerogibbinae (Parasclerogibbini, Tanynotini). All of these taxa were synonymized with the nominotypical subfamily by Engel & Grimaldi (2006a), who also recognized a new extinct subfamily,

†Sclerogibbodinae. See Engel & Grimaldi (2006a) and Perkovsky *et al*. (2020) for the most recent generic synopsis of the family, including a key to all extant and extinct genera in the latter.

***Note 23.*** The sistergroup relationship of Dryinidae and Embolemidae was previously postulated by Rasnitsyn in Rasnitsyn & Quicke (2002). See the note on the diagnosis of this clade below for more detail.

***Note 24.*** The Dryinidae were insufficiently sampled in the present study to provide internal structure.

***Note 25.*** The monophyly of the “†*Prosphex* genus group” is quite uncertain and deserves renewed attention. Li *et al*. (2020b) considered †*Prosphex* to be a member of †Panguidae stating that the eighth abdominal (seventh metasomal) segment is exposed, sharing a series of wing venation plesiomorphies plus an apparent derived condition of the hind wing, “r-m aligned with 1-M in a smooth vein, with free M end lost and with free end of A lost as well, resulting in a trigonalyd-like pattern”. Our analyses did not recover †*Prosphex* outside of the Aculeata, thus we treat the genus as a member of the Aculeata, rather than †Panguoidea.

***Note 26.*** See the notes on the diagnosis of Rhopalosomatidae below for discussion of generic relationships.

***Note 27.*** To the best of our knowledge there does not exist a catalog of the family-group names, their attributions, and synonymies for the Vespidae. For this reason, we are uncertain about the attributions of some names provided above, specifically Polistinae, Gayellinae, and Stenogastrinae. However, we thank J. Carpenter for sharing his records of taxon attribution.

***Note 28.*** The internal classification of the crown Vespidae shown here follows the phylogenomic results of Bank *et al*. (2017) and Piekarski *et al*. (2018). See the Vespidae diagnosis below for notes on fossil placement and topology.

***Note 29*.** Stenogastrinae Bequaert, 1918 was originally recognized at family rank as Ischnogastrinae Ashmead, 1902a, but was replaced after *Ischnogaster* Guérin-Méneville, 1838 in Duperrey (1838) was found to be a junior synonym of *Stenogaster* Guérin-Méneville, 1831 in Duperrey (1831). See p. 32 of Carpenter & Kojima (1996) and article 40.2 of the ICZN (2021).

***Note 30.*** We choose here to simplify the classification of this clade by recognizing it as a single superfamily, the Pompiloidea, with these constituent families: Sierolomorphidae, Tiphiidae, Chyphotidae, Thynnidae, †Burmusculidae, Pompilidae, †Bryopompilidae, and Mutillidae. These groups were previously classified in a multi-superfamily scheme based on molecular phylogenetic results. Specifically, Pilgrim *et al*. (2008) split Tiphiidae and the polyphyletic Bradynobaenidae and recognized Tiphioidea, Thynnoidea, Pompiloidea, while Branstetter *et al*. (2017b) cleaved Sierolomorphidae from Tiphioidea and gave this small group a new status as Sierolomorphoidea. For our single superfamily system, we therefore consider Pompiloidea to be the senior synonym of Sierolomorphoidea Krombein, 1951 **syn. nov.**, Tiphioidea Leach, 1815 **syn. nov.**, and Thynnoidea Erichson 1842 **syn. nov.** With the proposal of the present superfamilial consolidation, our hope is to contribute to the stabilization of pompiloid taxonomy and to encourage explicit comparative morphological studies at higher and lower levels, *i.e.*, of the families contained in this group relative to one another, and of this group relative to other superfamily-ranked clades. Because the names recognizing the interrelationships of the family-group taxa of Pompiloidea are informal—Tiphiopompiloides, Tiphiiformes, Pompiliformes, Mutilliformes—they will be robust to subsequent topological findings, such as removal of Myrmosinae from Mutillidae (*e.g.*, Waldren 2021), or the description or reshuffling of Cretaceous stem families. We expect that new, meaningful morphological synapomorphies across the superfamily will be discovered.

***Note 31.*** The clade recognized here as Tiphiiformes has had a long and complicated history, with the root-name group “Tiphiidae” having been used for a variety of circumscriptions. The tiphiiform lineages have been traditionally associated with Mutillidae, Scoliidae, and Bradynobaenidae *sensu lato*, at some point with these taxa considered altogether to form “Scolioidea”, a superfamily now known to represent a polyphyletic assemblage of fossorial aculeates. We retain the Tiphiidae, Chyphotidae, and Thynnidae as families given their ready diagnosability and to ensure that a rank change cascade is not initiated within these respective families. For pre-molecular taxonomic synopses of these almost terminally afflicted or otherwise astonishingly confused groups, see Rasnitsyn (1986), Kimsey (1991, 2006, 2009), Boni Bartalucci (2004).

***Note 32.*** Although †Thanatotiphiinae is unplaced in the Tiphiiformes based on our present analyses, we conservatively retain it in Tiphiidae and consider it *incertae sedis* therein.

***Note 33.*** See Brothers & Lelej (2017) for a morphological phylogeny and family-group synopsis of Mutillidae. Waldren (2021), using phylogenomic data in his dissertation work, has moved to raise Myrmosinae to family level based on its unstable position; no pointed dissent is offered here. We conservatively retain Myrmosinae within the Mutillidae until an incisive test is published.

***Note 34.*** Argaman (1994) substantially revised the classification of Apterogyninae, adding four tribes and seven genera, for a total of five and 10, respectively. The classification was simplified by Pagliano (2002), leaving only four genera in the group. See Pagliano & Romano (2018) for the most recent catalog of bradynobaenid genus and species names, albeit including Chyphotidae.

***Note 35.*** The higher classification of crown Scoliidae is under revision by Khouri on phylogenomic data (Khouri *et al*. 2022), as the taxonomy of the family is in serious need of attention. The high-water mark of splitting in the family is that of Argaman (1996), in which he recognized three subfamilies (including one new, Colpinae), 28 tribes (21 new), and 143 genera of which 62 were proposed as new. In his morphological cladistic study of *Scolia*, Golfetti (2019) estimated that about 560 extant species can be attributed to the family; this stands in contrast to Formicidae, for example, where 337 genera encompass >14,000 species, and the ant genera *Pheidole* and *Camponotus* alone comprise >1,100 and >1,000 species, respectively. Therefore, toward simplifying the extremely fragmented and unstable internal classification of Scoliidae, we have provided estimated synapomorphies for the total clade of the family, the crown clade, traits to be evaluated in *Proscolia* (Proscoliinae), the Scoliinae, the Campsomerini and Scoliini, and the core clades within these two tribes. Importantly, Khouri *et al*. (2022) found compelling support for the non-monophyly of Campsomerini, as *Colpa* was robustly recovered as sister to the Scoliini. No further revisionary efforts are taken in the present work, recognizing the importance of expanding the taxon sampling of the family (Khouri *et al*. 2022). The subfamily and tribal systems employed here follow Osten (2005); see also Kimsey & Brothers (2016). See Note 1 of the Scoliidae diagnosis below for discussion of fossil placement.

***Note 36.*** Monophyly of the †Archaeoscoliinae is uncertain, and several Mesozoic genera remain unplaced.

***Note 37.*** Some publications recognize Bradley (1957) as the authority for the name Campsomerini. This appears to be in error, as the cited work is a treatment of the genus *Campsomeris*, albeit this being the concept of Campsomerini retained in the literature. According to Kimsey & Brothers (2016), the appropriate attribution is Campsomerini (erstwhile Campsomerinae) Betrem in Betrem & Bradley, 1972.

***Note 38.*** The internal classification of the Apoidea is still in flux (*e.g.*, Branstetter *et al*. 2017, Sann *et al*. 2018, 2021), particularly with respect to fossil taxa. For this reason, and to hopefully benefit future study, we provide a classification of the Apoidea to tribal level. The classification espoused below follows the three phylogenomic studies cited in this note, with authority citations for spheciform apoids following Pulawski (2020). Names of Anthophila follow the suggestions of Engel (2005a). A number of taxon names were added or moved after establishment of the present system. We refrain from posing new formal names or creating rank changes beyond what has already been done as these efforts should be the subject of focused phylomic study, *i.e.*, synthesized based on analysis of phenomic and genomic data. Finally, based on our present analyses, we recognize †*Apisphex penyaleveri* **gen. nov. comb. nov.** as the oldest definitive apoid wasp.

***Note 39.*** Rosa & Melo (2021) took a series of taxonomic actions which affect the classification here reported: (a) †*Discoscapa* Poinar, 2020 placed in Crabronidae and †Discoscapidae Poinar, 2020 considered to be a tribe within Crabroninae; (b) two new families erected, †Allommationidae and †Spheciellidae; (c) †Cirrosphecinae raised to family with one new subfamily erected therein, †Glenocephalinae; (d) †Apodolichurini elevated to subfamily rank and transferred to Heterogynaidae from Ampulicidae; (e) †Mendampulicini elevated to subfamily rank and retained in Ampulicidae; (f) subfamily †Ptilocosminae erected in Heterogynaidae; (g) †Burmastatinae lowered to tribal rank and placed in Astatinae. The authors also support the argumentation of Melo & Rosa (2018) and retain †*Burmasphex* in the Apoidea. Other actions are not here listed. Because these actions were taken after the extended phylogenetic results of the present study were written, we were unable to address the topological hypothesis of Rosa & Melo (2021). We tentatively list †Allommationidae and †Spheciellidae as *incertae sedis* in the superfamily here, although Rosa & Melo (2021) consider these taxa to be nested within the core Apoidea based on presence of the mesepisternal ridge and form of the mesocoxa. Likewise, we indicate †Burmastatinae, †Cirrosphecinae, †Apodolichurinae, and †Mendampulicinae as *incertae sedis* based on our results; we make no comment here on the actions of Rosa & Melo (2021) except to note the incongruence between our systems at present. Critically, we did not recover †*Burmasphex* in the Apoidea, so we list this genus twice, once with †Falsiformicidae and once here.

***Note 40.*** See the notes on the diagnosis of Apoidea below for discussion of the †Angarosphecidae problem.

***Note 41.*** Rosa & Melo (2021) placed †Apodolichurinae in Heterogynaidae based on a suite of morphological characters. We tentatively leave this subfamily as *incertae sedis* in the Apoidea, as its placement is uncertain based on our analyses.

***Note 42.*** Based on our topological results, we provisionally removed †Apodolichurini, †Mendampulicini, †*Gallosphex*, and †*Cariridris* from Ampulicidae, and we set †Cretampulicini as *incertae sedis* to subfamily.

***Note 43.*** Aphelotomini is attributed to Ohl & Spahn, 2009 in the literature, including Pulawski (2020). An online early version of the work did appear in 2009, but the immutable online version has a publication date of 5 January 2010. Because the taxon name is not registered in ZooBank, it would not be available until the final online version is released. To resolve the date of publication for Aphelotomini, it will be necessary to determine when *Cladistics* mailed the physical copies of volume 26, issue 1. We conservatively retain 2009 as the year of publication but refer to Ohl & Spahn (2010) in our literature cited section.

***Note 44.*** The fossil placement within the core Apoidea clade was highly uncertain.

***Note 45.*** Of the †Cirrosphecidae, †*Cirrosphex* was recovered as sister to the remainder of the core Apoidea in our analyses. The other genus, †*Haptodioctes*, as well as the subfamily †Glenocephalinae were established by Rosa & Melo (2021) after we had completed the analytical phase of our study.

***Note 46.*** Heterogynaidae is *incertae sedis* in the superfamily Apoidea. Despite having substantially more sequence data than Sann *et al*. (2018), Sann *et al*. (2021) were unable to resolve the placement of this family. It is possible that Heterogynaidae are highly derived bembicids in or near Nyssonini (Sann *et al*. 2021). Rosa & Melo (2021) added both †Apodolichurinae and a new subfamily, †Ptilocosminae, to the Heterogynaidae. The †Apodolichurinae are of uncertain placement in our analyses.

***Note 47.*** †Discoscapini was transferred to Crabroninae by Rosa & Melo (2021) based on the form of the mesepisternal sulcus and an unpublished topological analysis. We were unable to include †*Discoscapa* in our study and tentatively treat the fossil and its tribe as *incertae sedis* in the Crabronidae due to its uncertain relationship with Crabroninae and Dinetinae.

***Note 48.*** Astatidae is here treated as two parts: Astatidae (Astatinae) and Dinetinae. Between the penultimate manuscript submission and the ultimate, the formerly *incertae sedis* Astatidae (Astatinae) was recovered as sister to Bembicidae + Pemphredonidae–Anthophila by Sann *et al*. (2021). Dinetinae was placed in the Crabronidae–Sphecidae clade based on the results of Sann *et al*. (2018). Additionally, †Cretastatini was recently erected for a species from burmite by Rosa & Melo (2021), who also considered †Burmastatini and Eremiaspheciidae (Eremiaspheciini) as members of the Astatidae (Astatinae).

***Note 49.*** Entomosericini and Eremiaspheciini were elevated to family rank and considered of uncertain placement in the Bembicidae–Anthophila clade by Sann *et al*. (2021). The former was formerly a member of Pemphredonidae (Pulawski 2020), and the latter a member of Philanthidae (Bohart & Menke 1976). Laphyragogini is retained in the Eremiaspheciinae following Pulawski (2020) as this tribe was not sampled in Branstetter *et al*. (2017) or Sann *et al*. (2018, 2021). Rosa & Melo (2021) placed Eremiaspheciini in Astatinae.

***Note 50.*** We were unable to evaluate †Megaroliini or †Pristinopterini as these fossil taxa had been described after our analyses were complete.

***Note 51.*** Ammoplanidae was recovered as sister to Anthophila in Sann *et al*. (2018, 2021) and as sister to Pemphredonidae in Branstetter *et al*. (2017b). In the extended transformation series of the present study, the group is treated as close to Pemphredonidae due to our source data and guiding tree; future study should evaluate the transformational consequences of the Sann *et al*. topology. Our present sampling only includes Ammoplanina Evans, 1959.

***Note 52.*** The internal structure of Anthophila here represents the congruent results of Sann *et al*. (2018) and Branstetter (2017b). For internal classification, see Michener (2007).

***Note 53.*** Rosa & Melo (2021) considered †*Melittosphex* as *incertae sedis* in the Aculeata, noting that the fossil had the appearance of the Mutilliformes (Mutillidae + Sapygidae). The present study found maximal support for †*Melittosphex* with Anthophila. Ideally, †*Melittosphex* should be re-evaluated using µ-CT technology, although the setae will be difficult to evaluate

#### Formicoidea

***Note 54.*** The formicoid classification overview provided here extends to the level of subtribe in order to reflect the validity and prevailing usages of family-group names, although most of the tribes and the one subtribe are not diagnosed here.

***Note 55.*** No subfamily-ranked classification is proposed for the †@@@idae, although there is meaningful structural variation between †*Camelomecia*, †*Camelosphecia*, and the various undescribed forms.

***Note 56.*** Several genera are left *incertae sedis* in the family; these are listed in Results Part IV below.

***Note 57.*** *Opamyrma* Yamane *et al*., 2008 is a highly distinct lineage of Leptanillinae that has been diagnosed in the context of Formicidae as a whole, Amblyoponinae, and Leptanillomorpha, based on workers, queens, and males (Yamane *et al*. 2008, Boudinot 2015, Yamada *et al*. 2020, Griebenow 2020). As the genus would be *incertae sedis* in the Leptanillinae without a tribe, despite its known phylogenetic relationships, it is here recognized as a new tribe and is diagnosed below. The genus has only very recently been found to be a leptanillomorph (Borowiec *et al*. 2019).

***Note 58.*** With the exception of the Amblyoponomorpha, the internal phylogeny of the Poneria is as yet unresolved; see Branstetter *et al*. (2017a), Borowiec *et al*. (2019).

***Note 59.*** Although Probolomyrmecini is known to render Proceratiini paraphyletic (*e.g.*, Brady *et al*. 2006), these tribes are not synonymized here as global phylogenomic and morphological revision of the Proceratiinae is required.

***Note 60.*** Following the phylogenomic results of Blaimer *et al*. (2015), Ward *et al*. (2016) revised the tribal system of Formicinae and Boudinot *et al*. (2022) estimated transformation series across the subfamily. Based on the results of the latter study, two morphological series are discernible: The lasiomorph grade, which is paraphyletic with respect to the formicoform radiation clade. These grades and clades roughly correspond to Bolton’s (2003) lasiine and formicine tribe groups, respectively.

***Note 61.*** Based on phylogenomic analysis of a broad sample of Ectatomminae and Heteroponerinae, Camacho *et al*. (2022) synonymized the latter under the former as well as the Typhlomyrmecini Emery, 1911 under Ectatommini. As previously recognized (Keller 2011, Feitosa & Prada-Achiardi 2019, Camacho *et al*. 2020), the morphological differences between Ectatommini and Heteroponerini are limited, compared to Martialinae versus Leptanillinae, and Apomyrminae versus Amblyoponinae.

***Note 62*.** Since the molecular phylogenetic revision of the Myrmicinae by Ward *et al*. (2015), wherein numerous tribes were sunk and genera were redistributed, there has been use of a single subtribal name: Attina (*e.g.*, Calheiros *et al*. 2019, Sosa-Calvo *et al*. 2019, Cardenas *et al*. 2021, Teixeira *et al*. 2021, among many others). This name has not been used in a manner conferring availability, and we recommend that its usage be recognized as informal to avoid the formalization of subtribes throughout the Myrmicinae or other clades. With that noted, it will be possible to provide genus group diagnoses and keys in the Myrmicinae and Dorylinae, among other subfamilies. A key will be to conduct these studies in a synoptic manner so that the system can be comprehended from a single source rather than iteratively synthesized by each worker or researcher.

### Section 3: Summary of taxonomic actions

Although numerous actionable taxonomic actions are possible from the results of the present work (see Results Part II above and Part IV below), only a subset of these changes are formally recognized here. A summary of these actions in the order of their appearance in the extended topological results section is as follows. – **(1)** †Andreneliidae Rasnitsyn & Martínez-Delclòs, 2000 transferred from Evanioidea Latreille, 1802, placed in Trigonaloidea Cresson, 1887 **new grouping**. – **(2)** †*Renymus* **nom. nov.** replaces †*Allogaster* Ren, 1995 (†Bethylonymidae), a junior homonym of *Allogaster* Thomson, 1864 (Cerambycidae). – **(3)** †Bethylonyminae Rasnitsyn, 1975 **stat. nov.**recognized for †*Bethylonymus* and †*Bethylonymellus* to the exclusion of †*Renymus* and †*Meiagaster*. – **(4)** †Falsiformicidae Rasnitsyn, 1975 excluded from Vespaculeata and Chrysidoidea Latreille, 1802, *incertae sedis* in Aculeata; **superfam. trans.** –**(5)** †*Burmasphex* Melo & Rosa, 2018 excluded from Apoidea Latreille, 1802, *incertae sedis* in Aculeata **new grouping**. – **(6)** †*Cursoribythus* Cockx & McKellar *incertae sedis* in Scolebythidae **subfam. transfer**. – **(7)** †*Mesomutilla aptera* Zhang, 1985 *incertae sedis* in Aculeata **fam. transfer**. – **(8)** Pompiloidea as senior synonym of Sierolomorphoidea Krombein, 1951 **syn. nov.**, Tiphioidea Leach, 1815 **syn. nov.**, and Thynnoidea Erichson 1842 **syn. nov.** –**(9)** †Allommationidae, †Spheciellidae, †Burmastatinae, †Cirrosphecinae, †Apodolichurinae, and †Mendampulicinae *incertae sedis* in Apoidea **new grouping** / **fam. transfer**. – **(10)** †*Angarosphex penyaleveri* Rasnitsyn & Martínez-Delclòs, 2000 is transferred to †*Apisphex* **gen. nov.**, forming †*Ap. penyaleveri* **comb. nov.**– **(11)** †*Cariridris* Brandão & Martins-Neto, 1990 transferred from Ampulicidae Shuckard, 1840, placed in Formicidae Latreille, 1809 *incertae sedis* **fam. transfer**. – **(12)** †@@@idae **fam. nov.** described and placed in Formicoidea Latreille, 1809. – **(13)** †*Burmomyrma* Dlussky, 1996 transferred from †Falsiformicidae, placed in total Formicidae *incertae sedis* **fam. transfer**. – **(14)** †*Petropone* Dlussky, 1975 transferred from Aculeata *incertae sedis*, placed in total Formicidae *incertae sedis* **fam. transfer**. – **(15)** †*Poneropterus* Dlussky, 1983 transferred from Formicidae *incertae sedis*, placed in clade Sphecomyrmines (stem Formicidae) *incertae sedis* **new grouping**. – **(16)** †Armaniini Dlussky, 1983 **stat. rev.** – **(17)** †*Armania capitata* Dlussky, 1999 and †*A. pristina* Dlussky, 1999 transferred to form-genus †*Khetania* Dlussky, 1999, creating †*K. capitata* (Dlussky, 1999) **comb. n.** and †*K. pristina* (Dlussky, 1999) **comb. n.** – **(18)** †*Cretomyrma unicornis* Dlussky, 1975 is treated as the senior synonym of †*C. arnoldii* Dlussky, 1975 **syn. n.** – **(19)** †*Dlusskyidris* Bolton, 1994 transferred from Formicidae *incertae sedis*, placed in clade Sphecomyrmines (stem Formicidae) *incertae sedis* **new grouping**. – **(20)** †*Afromyrma* Dlussky *et al*., 2004 excluded from Myrmicinae Lepeletier de Saint-Fargeau, 1835, placed in crown Formicidae *incertae sedis* **new grouping**. – **(21)** †*Afropone* Dlussky *et al*., 2004 excluded from Ponerinae Lepeletier de Saint-Fargeau, 1835, placed in crown Formicidae *incertae sedis* **new grouping**. – **(22)** †*Canapone* Dlussky, 1999 excluded from Ponerinae, placed in crown Formicidae *incertae sedis* **new grouping**. – **(23)** †*Cananeuretus* Engel & Grimaldi, 1975 excluded from Aneuretinae Emery, 1913, placed in total clade Formicinae Latreille, 1809 **new grouping**. – **(24)** †*Dolichomyrma* Dlussky, 1975 transferred from Aculeata *incertae sedis*, placed in total Formicinae **new grouping**. – **(25)** †Aquilomyrmecini **trib. nov.** and †Haidomyrmecini **stat. nov.** established in †Haidomyrmecinae; – **(26)** Opamyrmini Boudinot & Griebenow **trib. nov.** established for *Opamyrma* (Leptanillinae). – **(27)** Anomalomyrmini Taylor, 1990 synonymized under Leptanillini Emery, 1910 **syn. nov.**

## Part IV: Extended Transformation Series

### Overview

To trace structural and functional transformations through the Formicoidea and into the crown of the Formicidae, we evaluated morphology for a diversified sample of extant Aculeata and comprehensive sample of Mesozoic aculeates, plus outgroups. Below we outline an inferred groundplan for the Aculeata, and list crownward synapomorphies in the Formicoidea and other clades through phylogenetically defined nodes, with analysis of function as guided by the *Principles of Insect Morphology* (Snodgrass 1935) and *Comparative Biomechanics* (Vogel 2013). For this reason, the extended transformation series here is the first functional morphological characterization of the ants and is the first to trace structural and functional evolution from the Aculeata into the formicid crown. Not all transformations below are inferred to be specific adaptations, but rather that most have functional consequences, providing a testable framework for future study. It is plausible that some derived features may be the result of pleiotropic modifications to alternative developmental pathways. Three major research directions arising are the need to conduct molecular experimentation to identify morphogenic causes of hymenopteran phenotype, analysis of modularity and codependency of phenotypic features, and the creation of a digital atlas of hymenopteran three-dimensional structure from micro-CT scan data and function based on biomechanical quantification. The transformation series proposed here may be considered a revision of or alternative hypothesis to Rasnitsyn (1988, 2002).

Below we list each of the clade-defining states which are either transitions (*apomorphies*) or key plesiomorphies. These states are accompanied by a statistical polarity estimate in the harvest worksheet, for which we use the following abbreviations from the 576 character reconstruction: **CB50** = BPP from total-evidence dating ASE analysis using the 50% complete plus †Bethylonymidae matrix (303 taxa total); **SFA** = BPP for ASE from the Scoliida (Scolioidea + Formicoidea + Apoidea) matrix. Abbreviations for the nesting analyses are as follows: **303t-JC** = equal-rates transition model; **303t-F81** = variable-rates transition model. Abbreviations for the diet analyses are as follows: **303t-U-JC** = unordered, equal-rates; **303t-U-F81** = unordered, variable rates; **303t-O-JC** = ordered, equal-rates; and **303t-O-N0** = ordered, prior for rate of 1➔0 set to be minimal. Finally, the abbreviation **5S** = supercharacter analysis.

For character state distributions among the 575 scored terminals, see the character definitions plus the raw data matrix and results deposited on Zenodo (doi: XXX). Topological support for this phylogenetic classification is given in Section 3 above. For certain terms, Hymenoptera Anatomy Ontology (HAO) permanent URLs (purls) (Yoder *et al*. 2010) are provided.

## SECTION 1. Transformation series from Aculeata into Formicoidea

### Y’. Total clade Aculeata Latreille, 1802

***Clade comprising*:** • **Chrysidoidea** Latreille, 1802, • clade **Dryinaculeata**. • ***Incertae sedis*** in Aculeata: †**Bethylonymidae**, Rasnitsyn, 1975, †**Falsiformicidae** Rasnitsyn, 1975 **superfam. transfer**, †*Burmasphex* Melo & Rosa, 2018 **superfam. transfer**, †*Mesomutilla aptera* Zhang, 1985 **superfam. transfer**.

#### Synapomorphies

**22.** Occipital carina not extending to hypostoma (50CB: 0.98). ***Function*:** It is possible that this condition reflects modification of the postocciput such that the cranium is raised anterodorsad from the frontal plane (strict hypognathy). Detailed study of occipital form and cranial comportment is necessary to determine the function of this probable synapomorphy. ***Polarity*:** The state for the Trigonaloidea–Aculeata node is equivocal.

**\*.** Scape shape and length are scored as a complex: scape not globular (char. **152**, 50CB: 0.88); scape slightly elongate, length > 2 x width (char. **153**, 50CB: 0.96). ***Function*:** The scape is the proximalmost segment of the antenna and bears on the internal surface of its articulatory bulb (“bulbus”) the insertions of tentorioscapal and scapopedicellar muscles (*e.g.*, Richter *et al*. 2019, 2020) which control much of the antennal range of motion. Variation in the length of the scape directly influences the mechanical advantage of the scapopedicellar articulation. Features of the proximalmost region of the scape, the bulbus and bulbus neck, are also meaningfully variable (Keller 2011).

**390.** Abdominal segment II with laterotergite (CB50: 0.90). ***Function*:** The function of the laterotergite is ambiguous based on present analysis but is retained in some Formicoidea (see Perrault 2004 for further information). ***Polarity*:** Expression of the laterotergite in more stemward nodes with equivocal CB50; root node CB50: 0.81.

***Note*:** Although modification of the ovipositor into a “sting” is the canonical synapomorphy of the Aculeata, there is conflicting evidence and insufficient anatomical documentation to define this condition anatomically. See notes for character 433 for further detail.

#### Select non-wing plesiomorphies

**15.** Cranium including eyes lateromedially wider than long, whether anteroposteriorly as in prognathy, or dorsoventrally as in hypognathy (CB50: 0.93). ***Function*:** Head proportion variation has no single functional consequence, as transformation of the cranial length and width by necessity changes developmental and mechanical factors of internal tissue and architecture. In general, broader heads may be optimized for powerful mandibular grip, whereas long heads may be optimized for fast mandibular closure (Paul 2001). Extremes in cranial proportionality can serve very specific ecological roles, however, as in the tubular crania of gynes (“queens”) in the bamboo-dwelling formicine species *Camponotus* (*Myrmostenus*) *mirabilis* Emery, 1903.

**21.** Occipital carina present (CB50: 1.0). ***Function*:** The occipital carina is a rim which limits the thoracic contact surface of the cranium. It has previously been inferred to be derived at least twice among the broad-waisted or “symphytan” Hymenoptera (Vilhelmsen 2011). Form of the occipital carina varies considerably within Formicoidea and among Hymenoptera more generally, where it occurs. These forms are correlated with a wide range of functions, often acting as a stop which limits motion of the cranium in one or more directions relative to the pronotum or propleurae, but also serving as a protective scrobe when the head is flexed to the prothorax, among other mechanical and behavioral roles. When the occipital carina is absent, the contact surface which it delimits is still usually distinguished by surface sculpture and other cuticular features. See note at end of Formicoidea synapomorphies for list of contact surface rims.

**\*.** Compound eyes and ocelli expressed in all sexes and forms (char. **37**, CB50: 1.0; char. **40**, CB50: 1.0). ***Function*:** Ancestral aculeates were sighted.

**45.** Cranial condyle, articulating with mandibular acetabulum, inconspicuous (CB50: 1.0). ***Function*:** The cranial condyle is the second articulation of the mandible, derived in Ectognatha (Hexapoda) Blanke *et al*. (2015a), which fixes the motion of the mandibles around a proximal dicondylic hinge. In hypognathous insects, the cranial condyle is located anterior to the mandibular condyle; in prognathous insects, the condyle is dorsal. See also Snodgrass (1935) and Blanke *et al*. (2015b) for further information.

**\*.** Masticatory margin of mandible tridentate (char. **65**, CB50: 0.82) and short (char. **48**, CB50: 1.0). ***Function*:** The presence of teeth enhances traction of the tool edge of the mandibular margins and may be used for a variety of functions. In Hymenoptera, these teeth are usually triangular and used for gripping objects.

**78.** Buccal cavity at least as broad as long (CB50: 1.0). ***Function*:** Transformation to a long buccal cavity is associated with elongation of the mouthparts, often for nectar consumption as in Anthophila (Apoidea).

**147.** Antennomere count: Either female > 12 or male > 13, or both ≥ 13 (50CB: 0.92 for Aculeata, 0.99 for deeper nodes). ***Function*:** No clear mechanical or chemosensory function can be attributed to many-annulate flagellae, or to stabilization of antennomere count at 13 between sexes.

**175.** Pronotum with anterior lobe or “neck” (CB50: 0.96). ***Function*:** The anteromedian lobe of the pronotum serves as a contact surface, both projecting the craniothoracic articulation (“neck”) anteriorly and preventing overextension of the cranium during extreme dorsal abduction. This lobe was observed to be absent in various non-aculeate outgroups.

**213.** Parascutal carinae present, margining the mesoscutum dorsad the fore wing insertion (CB50: 0.87). ***Function*:** Similar to the occipital carina, the parascutal carina delimits a contact surface, which in this case is termed the preaxilla, a cuticular area which also bears processes that are important for fore wing stroke mechanics (Snodgrass 1935, HAO purl: 0000800).

**\*.** Female notauli present (char. **217**, CB50: 0.99) and extending completely across mesoscutum (char. **218**, CB50: 0.96). ***Function*:** Notauli, by definition, correspond to the median border of certain indirect or dorsoventral flight muscles (HAO purl: 0000647). The mechanical consequence of notaular development is unclear but may relate to structural reinforcement.

**223.** Prepectus present and freely mobile (CB50: 0.94). ***Function*:** The prepectus is an intersegmental sclerite situated anterior to the mesopectus which bears the origin of spiracular occlusor muscles (Gibson 1985, HAO purl: 0000811). Functional consequences of variation in prepectal development have yet to be studied in detail, although studies of form are available (*e.g.*, Brothers 1975, Gibson 1985).

**\*.** Distal procoxal cavity “open”, exposing membrane and base of trochanter (chars. **269–271**, CB50: 1.0) and meso-and metathoracic coxal cavities “open”, revealing thoracicocoxal articulations laterally (chars. **272**, **273**, CB50: 1.0). ***Function*:** The distal procoxal cavity surrounds the dicondylic coxotrochanteral articulation, while the meso- and metathoracic coxal cavities engage with the proximal articulatory surfaces of their respective coxae. The open condition of these three articulations appears to be a groundplan feature of the Hexapoda. Variation in the conformation and development of these articular regions influences mechanical force transfer of the procoxotrochanteral and meso- and metathoracicocoxal muscles. In the case of the distal procoxal cavity among Hymenoptera, there is a transformational trend toward reinforcement, rotation, and closure via ventral suturing (Boudinot 2015).

**292.** Hind coxa linear, *i.e.*, with a single longitudinal axis (CB50: 1.0). ***Function*:** The metacoxa has three primary functions: columnar support of the body, swinging the telopod of the hind leg around the coxa’s proximal dicondylic hinge for locomotion via action of the thoracicocoxal muscles, and housing the levator and depressors of the telopod, *i.e.*, the coxotrochanteral muscles.

**\*.** Paired apicoventral mid and hind tibial spurs present (chars. **309–312**, CB50: > 0.95). ***Function*:** Meso- and metatibial spurs function variably in grooming and traction. Bolton (2003) inferred that both spurs of both legs were pectinate in the groundplan of the total Formicidae; this may also be true for the Aculeata.

**324.** Pretarsal claws dentate (CB50: 0.89). ***Function*:** Dentition of the pretarsal claws increases tarsal traction but is difficult to associate with specific functional or ecological dynamics.

**\*.** Propodeal spiracle slit-shaped (char. **350**, CB50: 1.0) and situated high and anteriorly, close in proximity to hind wing base (char. **349**, CB50: 1.0). ***Function*:** Slit-shaped propodeal spiracles are often larger than circular spiracles and are associated with demanding muscular movement, as for flight in most Hymenoptera or in running in various ants, such as Formicini (Formicinae).

**\*.** Abdominal segment III (metasomal II) without transverse groove dividing tergum and sternum into pre- and postsclerites (chars. **400**, **401**, CB50: 1.0). ***Function*:** Without the transverse groove, abdominal tergum and sternum III slide freely beneath those sclerites of segment II.

**A.** Nesting behaviors absent (303t-JC: 1.0; 303t-F81: 1.0). ***Note*:** Nesting behavior is here considered to be any form of developmental chamber preparation, including transport and placement of hosts or prey in preformed cavities. Parasitoids attacking hosts *in situ* or kleptoparasites taking advantage of brood or food-stocks which are parceled by nest-making hosts are scored as non-nesters. Fossils were categorically scored as unknown (“?”), yet absence of nesting behavior is supported at all of the deepest nodes, including the outermost outgroup, †Ephialtitoidea. Among sampled taxa, transition to nesting is recovered three times, once in the ancestor of the Vespidae, of *Pepsis* (Pompilidae), and of the Formicapoidina.

**B.** Diet carnivorous, lifestyle parasitoidal (*i.e.*, employing a single host) (303t-U-JC, 303t-U-F81, 303t-O-JC, 303t-O-N0: 1.0). ***Note*:** For this character, we distinguish parasitoidy, predation, and all other diets, the latter category including omnivory, pollinivory, passive scavenging, and kleptoparasitism. Transitions to predation from parasitoidy are supported once in the ancestor of the Vespidae and at least twice in the Formicapoidina (see Formicoidea below).

**5S.** Non-nesting, non-eusocial parasitoid (303t-TD: 1.0; 303t-ST: 1.0).

***Note 1*:** Plesiomorphies of the body were selected based on whether or not they varied along the stem or into the crown of the Formicidae, as well as whether they have been previously inferred to be ancestral to extant ants (*e.g.*, Bolton 2003).

***Note 2*:** After completion of all analyses, we observed that grooming brushes on the posterior surface of the metatibia were widespread among the aculeatan superfamilies, spanning the crown node. These brushes are present in various other Apocrita, including Evanioidea. Although it is too late to formally include expression of these brushes in the formal analyses, we do hypothesize that the brush is a plesiomorphy of the Aculeata. See also definition of Formicidae below.

#### Venational plesiomorphies

*Fore wing venation*: C present (char. **461**, CB50: 1.0); R present and tubular distal to pterostigma (char. **463**, CB50: 1.0); crossvein 1r-rs absent (char. **464**, CB50: 1.0); crossvein 2r-rs anterior junction at about pterostigma midlength (char. **469**, CB50: 1.0); “marginal cell 1” (= 2rrs) closed (char. 471), with apex pointed to narrowly rounded (char. **475**) (CB50s: 1.0); Rsf1 (first free abscissa of Rs distad R+Rs) present (char. **481**) and anterior juncture distinctly separated from Pterostigma (char. **482**) (CB50s: 1.0); M+Cu present as tubular vein (char. **490**, CB50: 1.0); Mf1 (first free abscissa of M distad M+Cu) present (char. **487**) and without distinct curvature (char. 488) (CB50s: 1.0); Rs+M present, tubular (char. **492**); Rsf2 (first Rs abscissa distad Rs+M) present (char. **493**, CB50: 1.0); Mf2 or Mf3 (first and second abscissae of M distad Rs+M, divided by 1m-cu) present, Mf2 shorter than Rs+M (char. **497**, CB50: 1.0), and Mf3 possibly shorter Rs+M (char. **498**, CB50: 0.68); M distal to Mf3 tubular (char. **501**, CB50: 0.99); adventitious veins between Rsf and M absent (char. **502**, CB50: 1.0); crossvein 2rs-m present (char. **511**) and joining Rs distal to 2r-rs by less than one of its lengths (char. **515**, CB50: 1.0); crossvein 3rs-m present (char. **517**, CB50 for Trigonaloidea + Aculeata: 0.77, prone to loss) and separated from distal wing margin by at least one of its lengths (char. **519**, CB50: 1.0); crossvein 1m-cu present (char. **530**), without distinct curve (char. **531**) and joining Rs+M at or distal to split of Rsf and M (char. **532**) (CB50s: 1.0); crossvein 2m-cu possibly present (char. **538**, support equivocal); crossvein 1cu-a present, situated at about the split of M+Cu (char **549**, support equivocal); Cu2 reaching 1A (char. **558**, CB50: 0.91); 2A and crossvein 1a-a absent (chars. **559**, **560**, CB50: 1.0).

*Hind wing venation*: Basal hamuli absent (char. **561**, CB50: 1.0); C present (char. **562**, CB50: 0.92; Rsf (first free abscissa of Rs) present (char. **563**, CB50: 0.99); crossvein r-m present (char. **565**, CB50: 0.99) and joining Rs distal to the split of R+Rs (char. **566**, CB50: 1.0); Mf2 (free abscissa of M distad Mf, 1rs-m juncture) present (char **568**, CB50: 0.90); Cuf present (char. **569**, CB50: 0.85); crossvein cu-a present (char. **570**, CB50: 0.99); basal cell not distally elongate (char. **572**, CB50: 0.99); jugal lobe well-developed (char. **574**, CB50: 0.85).

***Note*:** Due to the importance of wing venation for the fossil record of the Aculeata, groundplan plesiomorphies of this clade are here listed. See the character definition section below for extended description of these states, and Fig. 4 for a demonstration of the venation system employed in the present work. Functional comments are not provided, suffice it to note that the major transformational trends crownward in the Formicoidea is venational reduction and increase of proximal wing membrane support by modification of abscissal location; trends in other superfamilies differ.

### A’. Clade Dryinaculeata

*Clade comprising*: • Clade **Vespaculeata**, • **Dryinoidea** Haliday, 1833.

#### Synapomorphies

**147.** Antennomere count: Female ≤ 12 and/or male ≤ 13 (50CB: 0.70). ***Function*:** The evolutionary cause for this stabilization is unclear. ***Polarity*:** Support for this estimate is affected by the incompleteness of many fossil antennae. 50CB for Vespaculeata and all internal nodes: 1.00. Reversal to the ≥ 13 state is implied for Sclerogibbidae.

### B’. Clade Vespaculeata

*Clade comprising*: • Clade **Scolioides**, • clade **Vespoides**.

#### Synapomorphies

**87.** Clypeus loosely integrated with cranium, projecting from head some distance (CB50: 0.77). ***Function*:** Detailed study of clypeofrontal anatomy is necessary. The clypeus bears labral attachments and the origin of prepharyngeal muscles (Richter *et al*. 2019, 2020), hence influences suctorial dynamics. Variation in clypeal form is associated with structural changes of the tentorium, antennal torulus, and frontal region. ***Note*:** The muscles originating on the clypeus are cibarial muscles in various other insect orders, inserting on the epipharynx. In extant Formicidae, this part of the epipharynx is fused with the hypopharynx, thus forming a prepharynx (Richter *et al*. 2019, 2020). ***Polarity*:** The state of clypeal integration is equivocal for Dryinaculeata, Scolioides, Formicapoidina, and Formicoidea. Resolution of the condition at these nodes should be improved through future study to refine the characterization of clypeal integration.

**146.** Antennomere count sexually dimorphic (CB50: 0.60). ***Function*:** The evolutionary cause for this stabilization is unclear. ***Polarity*:** This estimate is affected by the difficulty of evaluating intersexual dimorphism in fossils. 50CB for Vespoides and all internal nodes: 1.00.

**223.** Prepectus immobilized, reduced (CB50: 0.64). ***Function*:** Gibson (1985) notes that the prepectus may be fused to the pronotum in Vespidae, or to the mesopectus in other Aculeata, indicating that this ancestral condition may represent a tendency. Topology of the spiracular occlusor muscle varies with prepectal conformation. Further anatomical study in Aculeata is necessary to resolve functional consequences of prepectal transformation. ***Polarity*:** Support for this estimate is affected by fossil incompleteness. Vespoides CB50: 0.64, Scoliida: 0.96. Conditions within the Scoliida reflect complete fusion of the prepectus with the mesopectus.

**401.** Abdominal segment III (metasomal II) presternite present (CB50: 0.80). ***Function*:** The presternite of abdominal segment III is delimited by a sulcus which serves as a catch or stop for the preceding sternum during ventral flexion of the abdomen. Because of the catching or locking of abdominal sterna II and III which increases rigidity of the metasomal venter, overextension is mechanically limited or prevented, and stinging may be enhanced. Specifically, force may be more effectively transferred from the abdominal depressors originating within the propodeum and inserting on abdominal segment II, while also providing resistance to telescoping. ***Polarity*:** CB50 is maximal within Vespaculeata. CB50 for character 400, segment III pretergite present, is equivocal at the vespaculeate node.

**432.** Abdominal tergum VIII (metasomal VII) of female completely internalized (CB50: 0.97). ***Note*:** See notes for this condition and character 433 in the formal character description of Section 5 below for more detail.

***Notes*:** Distal elongation of the hind wing radial (“basal”) cell (char. **572**) is equivocal for this node but is strongly supported as the condition for the Vespoidea (CB50: 0.90), Pompiloidea *sensu novum* (CB50: 0.93), and Apoidea (CB50: 0.90), albeit equivocally so for Scoliida (CB50: 0.65).

### B’.2. Clade Scolioides

*Clade comprising*: • Clade **Formicapoidina**, • **Scolioidea** Latreille, 1802.

#### Synapomorphies

**63.** Mandible at least unidentate, *i.e.*, unidentate or bidentate from three- or more-dentate condition (50CB: 0.86). ***Function*:** No obvious function is correlated with this inferred transformation. ***Polarity*:** This condition equivocal for Vespaculeata.

**227.** Longitudinal (craniocaudally oriented) mesopectal sulcus present (CB50: 0.60). ***Function*:** Insufficient anatomical information is available to predict or define the function of this groove. Based on direct examination of Formicidae, the sulcus separates the pleurocoxal and indirect flight muscles. ***Polarity*:** Although support is equivocal for this condition, a craniocaudally oriented sulcus on the mesopectus is observed in Scoliidae, Apoidea, and Formicoidea. Among Vespoides, a morphologically and probably functionally similar sulcus is observed in some Vespidae, and very few Pompiloidea *sensu novum*, including Pompiliformes, †*Bryopompilus*, and †*Fedtschenkia*. Support was probably diminished to some degree by the fact that no observed Bradynobaenidae have this sulcus, and that the trait could not be evaluated for many fossils. For these reasons, we interpret presence of this sulcus as a synapomorphy of clade Scolioides.

**523.** Fore wing “submarginal cell 1” (= 1rrs) at least as long as “submarginal cells 2” and “3” (= 3rsm, 4rsm) when these are present (CB50: 0.64). ***Polarity*:** CB50 for Formicapoidina 0.76. The Scoliida node is possibly influenced by venational reduction of Bradynobaenidae (Scolioidea), correlated with an independent origin of female aptery.

**S.** Nesting non-eusocial parasitoid (303t-TD: 0.88; 303t-ST: 0.94). ***Note*:** Nesting is equivocally supported at the Vespaculeata node when estimated across a distribution of trees (303t-TD: 0.60 vs. 303t-TD: 0.40 for non-nesting) and is well-but misleadingly supported from single tree analysis (303t-SD: 0.80 vs. 303t-SD: 0.20 for non-nesting). Both analytical regimes, however, support nesting behavior at the Scoliida node. The state coding for the individual analysis of nesting (character **A**) was more restricted, with various tiphioids and Scolioidea scored as non-nesters; in this case, nesting was definitively supported for the Formicapoidina (303t-JC: 0.97; 303t-F81: 0.95). Reversal from nesting non-eusocial parasitoid to non-nesting non-eusocial parasitoid is weakly to equivocally supported for Rhopalosomatidae, and strongly supported for Mutillidae. Core Apoidea, *i.e.*, excluding Ampulicidae, are reconstructed as continuous provisioners in analysis of character **S**.

### E’. Clade Formicapoidina

*Clade comprising*: • **Apoidea** Latreille, 1802, • **Formicoidea** Latreille, 1809.

#### Synapomorphies

**227.** Longitudinal mesopectal sulcus present (CB50: 0.85). ***Function*:** The longitudinal mesopectal sulcus divides the lateral mesopectus into dorsal and ventral portions, which in alates bear the origins of coxal and indirect flight muscles, respectively. Functional variation will need to be ascertained by detailed skeletomuscular analysis. Deeper stemward patterns of “pleural” evolution in Hymenoptera remain unresolved (Gibson 1993). ***Polarity*:** Support for this condition in Scoliida and more stemward nodes up to Dryinaculeata is equivocal. Formicapoidina CB50: 0.85, Formicoidea: 0.98. Convergently derived in the Pompilidae + †Burmusculidae clade.

**550.** Fore wing crossvein 1cu-a anterior junction “prefurcal”, *i.e.*, proximal to the branching point of M+Cu (50CB: 0.76). ***Polarity*:** CB50 for Apoidea: 0.92; Formicoidea: 0.84.

***Note 1*:** There is equivocal support for antennomere III being longer than antennomere IV at this node (CB50: 0.61). Support increases for the total Ampulicidae and core Apoidea (CB50: 0.88, 0.75, respectively). Although support is equivocal for the Formicapoidina, we provisionally recognize that this condition as a candidate synapomorphy, especially given that the majority of sampled Apoidea and fossil Formicoidea had a proportionally elongate third antennomere, not to mention Leptanillomorpha and some Poneria. All sampled Scolioidea, in contrast, had a shorter third antennomere with the exception of *Trisciloa* and two species of †*Cretoscolia*. See the raw data matrix for further information on character state distribution.

***Note 2*:** The overall similarity of alate †Haidomyrmecinae, †@@@idae **fam. nov.**, Ampulicidae, and Sphecidae suggests that the Formicapoidina may have passed through an ancestor roughly similar to these groups, although it is also possible that this rough body form was derived in parallel. Future ancestral state modeling should provide special consideration for the formicapoidinan node.

### **F.** Superfamily Formicoidea Latreille, 1809

*Clade comprising*: • **†@@@idae fam. nov.**, • **Formicidae** Latreille, 1809.

#### Synapomorphies

**10.** Postgenal bridge elongate (50CB: 0.86; SFA for crown Formicidae: 0.97). ***Function*:** Elongation of the postgenal bridge is associated with prognathy (char. 11 below), but crucially in ants involves derivation of the internal postgenal ridge which bears muscular origins, including that of the mandibular abductor (“opener”) (Richter *et al*. 2020), representing an optimization of the internal organization of musculature. Moreover, postgenal bridge elongation is also associated with increased cranial mobility via transformation of the craniothoracic articulation, wherein the postocciput is modified into a ball-like articulation fitting into the socket-like prothoracic cervix, which is set in the countersunk occiput (Vilhelmsen 1999, 2011; Nguyen *et al*. 2014; Keller *et al*. 2014; Liu *et al*. 2019; Richter *et al*. 2019, 2020).

**11.** Cranium prognathous (50CB: 0.70; SFA for crown Formicidae: 0.93). ***Function*:** Prognathy is determined by angle of the postocciput relative to the length of the cranium and represents a spectrum of cranial orientation out of the transverse plane to hyperprognathy, in which the mouthparts are held in the frontal plane. In combination with postgenal elongation, the prognathous condition both increases the distance mechanical advantage of cranial motion, as well as raises the mouthparts anterodorsally out of the transverse plane. With mouthparts closer to the frontal plane, contact with anteriorly located objects is facilitated, as well as the action of raising objects held in the mandibles above the frontal plane. ***Note*:** Cranial orientation in Aculeata is complex and difficult to discretize in a simple manner. See Boudinot *et al*. (2021) and Richter *et al*. (in prep./2022) for further discussion.

**44.** Cranial/clypeal condyle enlarged (50CB: 0.98). ***Function*:** Enlargement of the cranial condyle, the mandibular acetabulum (its opposing articulation), and by consequence their contact surfaces, allows the mandible to have a wider gape (Richter *et al*. 2020). This has been described as a “cam” or slide-lock mechanism previously in the literature (Gronenberg *et al*. 1998; Zhang *et al*. 2020), where it has also been demonstrated that the enlarged condyle allows for biaxial mandibular potion, *i.e.*, the mandibles open laterally and rotate along their longitudinal axes near the extremes. Further study is necessary.

**142.** Antennal toruli directed laterally (50CB: 0.90; SFA for crown Formicidae: 0.86). ***Function*:** Given the prognathous condition of the cranium, the lateral orientation of the antennal socket is caused by the dorsad elevation of the medial rim of the torulus above the lateral rim. Ancestrally, the toruli were directed anteriorly (dorsally). Consequently, the antennae are directed at a dorsolateral oblique angle relative to the dorsal cranial surface, allowing for further lateral to ventrolateral motion, such that the scapes may be approximately aligned in the frontal plane. Medial overextension is mechanically prevented by the medial rim of the torulus in Formicoidea and by the frontal carinae in Formicidae. ***Note*:** Richter *et al*. (in prep./2022) redefined this condition as their character 25: Face between toruli bulging, thus toruli oriented more laterally. They referred to this synapomorphy as the “frontal bulge”.

**177.** Pronotum muscular, elongate (50CB: 0.75; SFA for crown Formicidae: 0.83). ***Function*:** Elongation of the pronotum is associated with elongation and increase in volume of craniothoracic musculature, increasing neck strength under load (Nguyen *et al*. 2014; Keller *et al*. 2014). Elongate pronota have evolved in other groups of insects, *e.g.*, Raphidioptera, Mantispidae, but these conformations are starkly different and result in different muscular topologies and volumes as compared to Formicidae. ***Polarity*:** The condition of the pronotum is equivocal at all nodes from Aculeata crownward to the Formicapoidina, but is well-supported for Formicoidea, Antennoclypeata, and internal nodes.

**269.** Procoxal distal foramen “closed” (50CB: 0.97; SFA for crown Formicidae: 0.98). ***Function*:** The distal procoxal foramen of Formicoidea is completely “closed”, *i.e.*, the rim of the foramen is ventrally sutured and completely encircles the trochanter, which itself is characteristically constricted and kinked. Based on prior (Boudinot 2015) and present observations, this condition in extant ants is associated with presence of only a single coxal condyle, representing a key exception to generalization that all coxotrochanteral hinges are dicondylic with an anteroposterior axis (Snodgrass 1935). With a single coxal condyle which embraces the trochanteral acetabulum, itself extended on an arm-like process, the formicid coxotrochanteral articulation rotates around a lateromedial axis. Because of the trochanteral kink, the telopod (those segments of the leg distal to the coxa) swings in or near the sagittal plane, maximizing stride length. Some degree of lateromedial freedom in the coxotrochanteral articulation is observable in live ants, enhanced by the slightly flexible trochanterofemoral articulation, allowing the legs to sway, perhaps increasing balance on variably even surfaces. A similar condition of distal procoxal foramen closure is observed in *Myrmosa* (Pompiloidea: Mutillidae), a taxon with apterous females, but anatomical details need to be evaluated. Note that the procoxa of the Formicoidea is further distinguished by its proportions, being in general about twice as long dorsoventrally as wide lateromedially (Liu *et al*. 2019); unfortunately, this condition was not scored for the present study.

**\*.** Modifications of the meso- and metacoxae and their thoracic articulations for walking represent a complex set of characters: mesothoracic coxal cavity “closed” (char. **272**, 50CB: 0.94); metathoracic coxal cavity “closed” (char. **273**, 50CB: 0.92; SFA for crown Formicidae 0.99); proximal articulations of mid- and hind coxae constricted and internalized (char. **288**, CB50: 0.85); metacoxa length with two axes, rather than one (char. **292**, 50CB: 0.93; SFA for crown Formicidae: 0.97). ***Function*:** “Closure” of the meso- and metathoracic coxal cavities in ants is associated with modification of the proximal surfaces of the coxae into ball-like articulatory structures, fitting within the thoracic coxal sockets. This conformation is also associated with curvature of the coxal axis, such that both the meso- and metacoxae have two axes (although the former was not scored). The first axis is dorsoventral, representing the proximal axis of coxal rotation, oriented dorsoventrally, rather than at an oblique ventrolateral angle, as observed in the ancestral condition as retained in such taxa as Pompilidae (illustrated in Fig. 3 of the main text). This first axis also represents, to a large degree, the columnar support of the mid and hind legs. With the legs directed laterally, the second axis is lateromedial relative to the body and describes the longitudinal orientation of the coxotrochanteral muscles. When the mid and hind legs are rotated on their first axis, the second axis swings in the frontal plane, parallel to the idealized surface of the ground. In this way, excursion of the coxal apices is maximally distant from the midline of the body as represented by the first or dorsoventral coxal axis, increasing the effective leg length and consequently, stride length. Similar modifications are also observed in taxa with apterous females or which seek prey or hosts within the topmost soil layers, including various Chrysidoidea (Plumariidae, Bethylidae), some Dryinoidea (at least one species of Sclerogibbidae), *Olixon* (Vespoidea: Rhopalosomatidae), Tiphiiformes and Mutillidae (Pompiloidea), and Bradynobaenidae (Scoliidae). Note that not all of these groups have the exact pattern of apomorphy as indicated above. The occurrence of closed thoracicocoxal articulations in *Loboscelidia* (Chrysididae) is very likely an adaptation for self-defense given their probable myrmecophily. In contrast, the occurrence of closed cavities in Scoliidae are probably due to the need for reinforcing limbs, given that they must dig up through soil after pupation before they can feed as adults or mate. These parallelisms should be subjected to comparative anatomical and mechanical study. ***Polarity*:** 50CBs of char. 272 (mesothoracic closure) for Scolioides, Scolioidea, and Formicapoidina are equivocal. Taxon sampling is apparently important here, as the SFA results strongly conflict (> 0.99 for “closure” as ancestral to the Scoliida and Formicapoidina). See Gibson 1985 for a treatment of mesothoracicocoxal articulation variation, emphasizing non-aculeate Hymenoptera.

**370.** Petiole (abdominal segment II, AII) anterior foramen at least subsessile, *i.e.*, with some length of anterior neck immediately posterad the propodeal articulation and associated sclerotic structures (50CB: 0.80). ***Function*:** The anterior neck, or “peduncle”, of ants is an elongation of the petiole between the anterior articulatory surfaces and the node. This elongation is associated with increase in the length of the levator muscle of abdominal segment III (AIII, metasomal II, or “postpetiolar”), except in those highly derived instances where the peduncle is emusculated and the anterodorsal petiolar node surface is restricted to a posterior location as observed in Myrmicinae (Hashimoto 1996). Peduncular elongation may also increase the range of excursion for the AII–AIII articulation, depending on the proportions of abdominal segment II.

**375.** Petiolar tergum with distinct posterior face (50CB: 0.97; SFA for crown Formicidae: 0.99). ***Function*:** Presence of a distinct posterior face on the petiolar tergum is here taken as the key criterion for the development of a node. Internally, the petiolar node bears the origins of the abdominal segment III depressors, for origin surface area increases and muscle fiber length variation is increased when the posterodorsal face is present (Dlussky & Fedoseeva 1988; Hashimoto 1996).

**377.** Node of petiole lateromedially broader than anteroposteriorly long (50CB: 0.97). ***Function*:** The physiological significance of this condition is uncertain, although it does represent the groundplan conformation as here estimated.

**382.** Subpetiolar process of abdominal segment II present (50CB: 0.88). ***Function*:** The “subpetiolar” or anteroventral process of abdominal sternum II (metasomal I, “petiole”) is situated in a location which approximates the dorsoventral midlength of the hind coxae in all instances where it occurs. Thus, at full ventral metasomal flexion, the subpetiolar process would be inserted between the metacoxae. In this position, lateromedial shearing of the propodeal–petiolar articulation as during the action of stinging live prey would be resisted, because the process, hence petiole, is physically sandwiched by the hind coxae, thus physically limited in motion. In brief, the subpetiolar process increases the safety factor of stinging. ***Polarity*:** This is another character for which taxon sampling is important, as demonstrated by the differences between the SFA and 50CB analyses, wherein the latter includes fossil terminals, but the former does not. SFA support for process presence in crown Formicidae is equivocal (0.68) but is strongly supported in all internal backbone nodes of Formicoidea in 50CB (≥ 0.95). Bolton (2003) inferred that subpetiolar process absence was ancestral for the total Formicidae.

**385.** Petiolar tergum with dorsal, anterolateral paired carinae (CB: 0.81). ***Function*:** Anterolateral tergal carinae often end anteriorly with triangular processes, which limit motion of the petiole during extreme dorsolateral flexion. Presence of these processes were not recorded, however. The carinae themselves may stiffen the petiolar tergum against anteroposterior compression. ***Polarity*:** This condition was difficult to observe for fossils and the carinae are also observable in Scolioidea and Apoidea. Focused examination may demonstrate that these carinae may be more widespread, although they are at least inferred to be a groundplan feature of the formicoids.

**403.** Anterior articulatory surfaces of abdominal segment III (helcium) narrowed (50CB: 0.91). ***Function*:** Narrowing of the helcium is associated with formation of notches on abdominal segment II (petiole) for reception of segment III (Hashimoto 1996), increasing the tightness or rigidity of the intersegmental articulation, hence improving resistance to anteroposterior compression during stinging and force transfer from petiolar depression. This rigidity also transforms the tergal and sternal protractors of segments II and III into depressors and levators, respectively (Hashimoto 1996), thus increasing the muscular force transmitted in these directions. Additionally, the sulci delimiting the pre- and post-sclerites of abdominal segment III (the “helcial tergites and sternites”) act as stop mechanisms reducing or preventing dorsal and ventral overextension. Extreme narrowing, thus forming a ball-and- socket joint is reconstructed for Formicidae (char. 404), below.

**411.** Abdominal sternum III with prora (50CB: 0.90). ***Function*:** The “prora” is a sclerotic thickening of the contact surface of abdominal sternum III, where the sternum meets that of abdominal sternum II (the petiole) at full ventral flexion. As for the occipital carina, the prora has considerable variation in form, with differing functional consequences. For further information, see char. 412 at this node as well as char. 413 for the crown Formicidae below.

**412.** Prora in form of longitudinal keel (50CB: 0.93). ***Function*:** The prora in this condition extends anteriorly away from the sternum, thus decreasing the distance between the petiolar sternum, consequently limiting the range of ventral excursion for the third abdominal segment. At full ventral flexion, with the prora locked in contact with the petiolar sternum, the helcial articulation (that of the joint between segments II and III) becomes rigid, improving force transfer during stinging action. The prora, moreover, bears proprioceptor setae, thus transmitting information about the degree of metasomal flexion. Among †@@@idae and stem Formicidae, the proral keel varies from lateromedially narrow to broad.

**418.** Abdominal segment IV transversely constricted by transverse sulcus, “cinctus present” (50CB: 0.75; SFA for crown Formicidae: 0.96). ***Function*:** The constriction of abdominal segment IV is physically represented by a transverse sulcus, which limits the overlap of abdominal tergum and sternum III during dorsal and ventral flexion. This sulcus is often associated with “tubulation” (tergosternal alignment of abdominal segment IV) and the “postpetiolation” syndrome (most prominently represented by reduction in size of abdominal segment III) (*e.g.*, Taylor 1978; Perrault 2004). ***Polarity*:** 50CB for total Formicidae: 0.84; 1.0 for Antennoclypeata and crown Formicidae. This is a key departure from the groundplan inferred for the Formicidae by Bolton (2003), wherein the fourth abdominal segment was characterized as lacking the transverse sulcus.

**483.** Fore wing Rsf1 and Mf1 meeting at distinct, oblique angle (CB50: 0.78). ***Polarity*:** All internal formicoid nodes with CB50 > 0.95. This condition paralleled in Apoidea, Chrysidoidea, and various non-aculeate outgroups.

**5S.** Continuous provisioning present (303t-TD, 303t-SD: 0.95). ***Note*:** The stated posterior probability is an aggregate of the two highest-scoring states, which are nesting eusocial continuous-provisioner (303t-TD, 303t-ST: 0.48) and nesting non-eusocial continuous-provisioner (303t-TD, 303t-ST: 0.47), meaning that continuous provisioning is strongly supported but eusociality is strictly equivocal. It is likely that this continuously-provisioning formicoid ancestor was a predator, given that no direct transitions from parasitoidy to omnivory are observed in our dataset. However, the individual analysis of diet (**B**) reconstructs the most recent common ancestor of the Formicoidea as equivocal (**303t-O-N0: 0.51**; 303t-O-JC: 0.70; 303t-U-F81: 0.58; 303t-U-JC: 0.71). This is attributable to the scoring of diet among the Formicoidea. †@@@idae were categorically scored as unknown because it is not possible to count host or prey from the fossil record but are weakly supported as predators rather than parasitoids (**303t-O-N0: 0.50**; 3303t-O-JC: 0.69; 303t-U-F81: 0.58; 03t-U-JC: 0.70). Most stem Formicidae were scored as unknown because differentiating predators from omnivores in species with “generalized” morphology is impossible. In contrast, because stem Formicidae are unambiguously supported as workers, which by necessity forage, in the supercharacter analysis all terminals represented by apterous specimens were scored as continuous provisioners.

In the diet-only analysis (**B**), a selection of stem ants was scored as predators based on obvious morphological specializations: †*Myanmyrma mauradera*, †Haidomyrmecinae, and †Zigrasimeciini. Morphological evidence for predation includes the traction-setae-bearing cranial horns and increased dentition of the haidomyrmecines, the aggressive chaetae (“traction setae”) on the clypeus, labrum, and inner mandibular faces of the zigrasimeciines, and the labral chaetae and elongate, curved mandibles of †*M. mauradera*. That these adaptations would be for herbivory or plant-based liquid diets is unlikely, given the dietary requirements of plant-based diets and the known range of modifications for plant-based liquid intake. Suggestions that †*Zigrasimecia* may be pollinivores based on the row-like arrangement of perioral setae, reminiscent of apid leg combs, are unfounded, particularly as pollen has never been observed on these ants unlike †*Prosphex* and various coeval Coleoptera (*e.g.*, Peris *et al*. 2020). Additionally, elongate, sickle-like mandibles have evolved multiple times in members of the crown Formicidae which are specialist-predators (*e.g.*, Ponerinae: *Leptogenys*, *Myopias*, *Plectroctena*, *Promyopias*), but never in the omnivorous Formicae clade; the sickle-shaped mandibles of the dulotic formicine *Polyergus* are short, in contrast to †*M. mauradera*. Predation was highly supported for total Formicidae (**303t-O-N0: 0.96**; 303t-O-JC, 303t-U-F81: 0.97; 303t-U-JC: 0.98), including †Armaniinae. Crown ants were maximally supported as predators (1.0 BPP).

#### Key plesiomorphies

**\*.** Maxillary palp 6-merous, labial palp 4-merous. ***Note*:** Palp formulae were not recorded in our matrix because of the difficulty of evaluating palpomere counts in amber fossils, and the absence of these structures in impression fossils. Bolton (2003) inferred that this Hymenopteran plesiomorphy is retained in the ancestor of the total Formicidae, which is also reasonable for the formicoid node. A number of other groundplan features of the mouthparts established by Bolton could not be evaluated.

**\*.** Metasternal process absent. Metacoxal cavities of thorax open to propodeal foramen. Metacoxae closely approximated. ***Note*:** These is groundplan features, established by Bolton (2003), was not evaluated in the present study due to difficulty or impossibility of scoring for fossil terminals. Importantly, †*Kyromyrma*, a Cretaceous formicine, has wide-set metacoxae, a derived feature shared with Lasiini.

**217.** Female (gyne/queen) mesonotum with paired notauli. ***Polarity*:** Support for the retention of notauli at the formicoid, camelomeciid, and formicapoidan nodes is equivocal. Presence of notauli is well supported at the Scoliida and Vespaculeata nodes (CB50: 0.89, 0.93, respectively). Bolton (2003) inferred presence of female notauli to be a groundplan feature of the total Formicidae. See char. 217 under the total Formicidae node below for further discussion.

**324.** Pretarsal claws dentate. ***Note*:** Pretarsal claw dentition is supported as retained from the aculeate ancestor through the Formicoidea and to the anntennoclypeatan node, where support diminishes from CB50 > 0.97 to 0.80. This condition was inferred as a groundplan feature of total Formicidae by Bolton (2003).

**\*.** Helcium axial, *i.e.*, situated at about midheight of abdominal segment III. ***Polarity*:** Axiality describes the dorsoventral location of the helcium relative to the isosegmental postsclerites. When the helcium is above segment midheight, it is termed “supraaxial” (char. 410), when it is at midheight, it is termed “axial”, and below midheight “infraaxial” (char. 409) (Perrault 2004; Keller 2011). Among Formicoidea, the infraaxial condition has evolved independently in poneriines *Apomyrma* (Amblyoponinae) and Ponerini (Ponerinae), and in the formican Dolichoderomorpha and Formicomyrmines, with axiality retained in Myrmeciomorpha and regained in *Oecophylla* (Formicinae). Support for infraaxiality at the Formicae node is equivocal (CB50: 0.58). Supraaxiality has been derived in *Anomalomyrma* (Leptanillinae) and Amblyoponinae. Refined anatomical characterization is needed to improve our understanding of petiolar and AIII evolution.

***Note*:** A number of additional head characters were developed by Richter *et al*. (in prep./2022) based on a comparative µ-CT analysis of a stem ant, †*Gerontoformica gracilis*, and a series of crown ants and outgroups. Although †@@@idae could not be evaluated in that study, we list the new characters here in the hopes that they can be examined for specimens of the new family. These characters include a short dorsal tentorial arm (their char. 57), presence of a transverse line of setae on the labrum (their char. 73), and development of a pharyngeal gland (their char. 145). Other characters from Richter *et al*. (in prep./2022) are addressed below where appropriate.

### F.A. Family †@@@idae fam. nov

***LSID*:** XXX. [NOTE: Will be provided after manuscript accepted.]

***Type genus*:** †*Camelomecia* Barden & Grimaldi, 2016.

***Clade comprising*:** • †*Camelomecia* Barden & Grimaldi, 2016, • †*Camelosphecia* Boudinot *et al*., 2020.

***Stratigraphic horizon*.** Kachin burmite, Myanmar; 98.8 Ma based on U–Pb zircon dating (Shi *et al*. 2012).

***Etymology*.** Root †*Camelomec-* from the type genus. *Camelo*- was originally chosen in reference to the artiodactyl mammal, judged to be less beautiful than horses by some, and *mecia*- in reference to ants. Gender feminine.

#### Synapomorphies

**48.** Mandibular masticatory margin elongate (50CB: 0.94). ***Function*:** Elongation of the masticatory margin, the primary tool edge of the mandible, increases friction during contact of the appendage with an object. In camelomeciids, friction is enhanced by an array of very small to large teeth. *Note*: Convergent with Poneroformicines.

**51.** Mandible strongly bowed, thus bowl-like, having a strongly convex dorsal surface (50CB: 0.99). ***Function*:** A parallel condition is observed convergently in *Tatuidris* (Agroecomyrmecinae, Poneria). Both taxa have a dense vestiture of chaetae (“traction setae”) on the ventral mandibular surface (see, *e.g.*, Fig. 12B of Keller 2011), suggesting both a specialized diet (Jacquemin *et al*. 2014) and complex mechanism for object grasping. The range of objects grasped between the mandibles beyond prey is unclear, *i.e.*, there is no available evidence that the mandibles were used for nest construction.

**68.** Chaetae on inner (ventral) mandibular surface robustly developed (50CB: 0.94). ***Function*:** These setae increase the contact surface area, hence friction, of the mandibles surrounding the tool edge represented by the masticatory margin. ***Polarity*:** We recommend detailed re-examination of this condition, particularly among crown Formicidae using scanning electron microscopy.

**176.** Anterior pronotal rim upturned, often bearing seta fringe (50CB: 0.94). ***Function*:** In general, setae are innervated, so would provide tactile signal, but as the setae are not always present, the rim has no unambiguous function. ***Polarity*:** Autapomorphy.

**178.** Female pronotum elongate, length ≥ width (CB50: 0.94). ***Function*:** As noted above (chars. 10, 177 of Formicoidea), pronotal elongation is associated with increased neck strength in the context of object manipulation in the Formicoidea. The condition reconstructed here is of comparatively extreme pronotal elongation. Similar elongation has been derived independently in various other Aculeata, including some Bethylidae plus *Loboscelidia* (Chrysidoidea), Sclerogibbidae (Dryinoidea), *Sierolomorpha* (Sierolomorphidae), and *Aelurus* (Tiphiiformes). Functional analysis for these additional taxa will depend on their life histories and new anatomical study.

**191.** Pronotum broadly constricted posteriorly, just anterad mesoscutum (50CB: 0.97). ***Function*:** Narrow pronotal constriction is observable in some Apoidea. In these taxa, the pronotum bulges anterad the constriction, and the bulge corresponds to the origins of craniothoracic musculature, indicating functional association with neck strength.

**419.** Abdominal segment IV tubulated, *i.e.*, tergosternal margins aligned (50CB: 0.96). ***Function*:** Alignment of the lateral tergosternal margins, associated with partial to complete tergosternal fusion among extant Formicidae (Taylor 1978; Bolton 2003; Perrault 2004; Fisher & Bolton 2017), limits or prevents the tergum from overlapping the sternum, thus increasing the rigidity of the fourth abdominal segment, and improving force transfer during the action of stinging. ***Polarity*:** Tergosternal alignment of abdominal segment IV has arisen independently as a synapomorphy of the Poneria (50CB: 0.86), and apparently of the Ectatomminae (including Heteroponerini Bolton 2003). “Tubulation”, as this condition is known (Taylor 1978), also occurs in Dorylinae, although this was not evaluated in detail here.

**538.** Fore wing crossvein 2m-cu, when present, joining M at or proximal to 2rs-m crossvein (50CB: 0.62). ***Polarity*:** Support is low as 2m-cu is absent in three of the six sampled †@@@idae. Crossvein 2m-cu never occurs in Formicidae, thus loss is inferred to be parallel between the families.

**543.** Fore wing medial cell 2 (“discal cell 2”) broader anteroposteriorly than long proximodistally when present (*i.e.*, distally enclosed by 2m-cu) (CB50: 0.99).

**Extended diagnosis.** *Both sexes recognizable as Formicoidea by the following synapomorphic conditions* (Fig. 22, compare to Figs. 21A–J, 23): **(a)** cranium prognathous and postgenal bridge elongated; **(b)** cranial condyle (articulating with mandible) enlarged; **(c)** antennal toruli directed laterally, away from one another (although toruli directed dorsally in male); **(d)** protrochanter constricted and twisted; meso- and metathoracic coxal foramina “closed” (*i.e.*, these concealed in lateral view due to internalization of proximal coxal articulations); **(e)** abdominal segment II (metasomal I) petiolated; **(f)** abdominal sternum II with anteroventral process fitting between metacoxae at full anterior metasomal flexion; **(g)** abdominal sternum III (metasomal II) with anterior keel (prora); first free abscissae of Rs and M meeting at an angled juncture.

*Distinguished from the Formicidae total clade by the following synapomorphies* (Fig. 22): **(h)** masticatory mandibular margins elongated (short in stem Formicidae); **(i)** mandible strongly bowed, almost cup-shaped (nearly unique, occurs rarely among crown ants); **(j)** anterior pronotal margin upturned; **(k)** female pronotum anteroposterior length at least as long as lateromedial width (versus length less than width); **(l)** pronotum posteriorly constricted; abdominal segment IV tubulated, *i.e.*, margins of tergum and sternum meeting along their whole length (variable in Formicidae); **(m)** fore wing crossvein 2m-cu, when present, joining M at or proximal to 2rs-m crossvein (in contrast to Apoidea, Scolioidea; 2m-cu lost in some camelomeciids and in all Formicidae); **(n)** fore wing medial cell 2, when 2m-cu present, anteroposteriorly wider than proximodistally long (in contrast to Apoidea, Scolioidea).

*Identification supported by the following plesiomorphies*: **(o)** cranioclypeal integration weak, *i.e.*, clypeus projecting anteriorly, away from head (versus strongly integrated); **(p)** clypeus lacking chaetae (“traction setae”; apomorphically present in the sphecomyrmine clade and Amblyoponinae); **(q)** frontal carinae absent between antennal toruli (presence synapomorphic for total Formicidae); **(r)** head width, including eyes, usually wider than anteroposteriorly long (long head synapomorphic for total Formicidae); **(s)** propodeal spiracle located high and anteriorly on the segment (versus low and posterior position, synapomorphic for Formicidae); **(t)** fore wing Cu2 reaching 1A (occurs sporadically among stem and crown Formicidae).

### **F.B.** Family Formicidae Latreille, 1809 total clade

***Clade comprising*:** • **†Armaniinae** Dlussky, 1983, • clade **†Sphecomyrmines**, • clade **Antennoclypeata**. • ***Incertae sedis*** to higher clade or subfamily: †*Archaeopone* Dlussky, 1975; †*Baikuris* Dlussky, 1987; †*Burmomyrma* Dlussky, 1999; †*Petropone* Dlussky, 1975.

***Note*:** Total clade Formicidae is equivalent to the Formicidae of Bolton (2003) and pan Formicidae of Borysenko (2017).

#### Synapomorphies

**2.** Apterous individuals present (50CB: 1.00; SFA: 0.96). ***Note*:** Apterous or flightless females have been derived independently in each of the major clades of Aculeata, as evaluated at the superfamilial rank, including Chrysidoidea (Plumariidae, some Bethylidae), Dryinoidea (Sclerogibbidae), Vespoidea (Rhopalosomatidae: *Olixon*), various Tiphiiformes, Pompiloidea (Mutillidae), Scolioidea (Bradynobaenidae), Apoidea (Heterogynaidae), and of course, Formicoidea (Formicidae). The only exception is the small (< 20 extant spp., 2 recorded extinct spp.) and isolated family Sierolomorphidae. The evolution of flightlessness is generally considered to be related to energy conservation and habitat or resource stability (Roff 1990; McNab 1994) and is observed here to be correlated with modifications of the coxal articulations (among sampled taxa in the present study, 100% of those terminals expressing aptery also have at least one of the modifications associated with terrestrial locomotion, namely characters 269, 272, 273, 288, and 292). Based on our tip-dating analysis (50CB), a Cretaceous origin of aptery among sampled Aculeata is only supported for Tiphiiformes (although possibly in parallel within this clade) and Formicoidea. The enigmatic †Aptenoperissidae (Proctotrupomorpha?) represents another origin of Cretaceous flightlessness. Other instances of aculeate flightlessness plausibly evolved in the Cenozoic, but denser taxon sampling in this Era is necessary. In Aculeata, females are usually the apterous sex, although in rare cases, some male ants may be wingless (*e.g.*, some species of *Hypoponera*, *Cardiocondyla*, *Technomyrmex*, and various socially-parasites). The polarity of aptery at the Formicoidea and †@@@idae nodes are equivocal (50CB: 0.51), but strongly supported as absent at higher nodes (50CB > 0.90). For this reason, we state that aptery is “definitively present” in Formicidae, but equivocal for †@@@idae. It is possible but unknown whether some species of †*Camelomecia* had apterous females; if such flightless females were present in those species presently known, then this would be evidence for ancestral eusociality in the Formicoidea.

**4.** Female winged/wingless dimorphism present (50CB: 0.94; SFA: 0.97). ***Note*:** Expression of females with and without wings in a single species is strong indication for reproductive division of labor, hence eusociality. Trible & Kronauer (2017) argue that size-based polyphenic expression in female ants, particularly of the flight apparatus, is the primary factor determining reproductiveness or caste identity of an individual. Because evidence of dimorphism is absent in †@@@idae and other Formicoidea which are known only from alates, these taxa were scored as uncertain (“?”) in contrast to those formicoids which have apterous females (*e.g.*, various †Sphecomyrmines, †*Brownimecia*, and of course, crown Formicidae). 50CB for Formicoidea and †@@@idae is equivocal, supporting absence at 0.51 for both nodes. For this reason, we interpret reproductive division of labor, a key criterion of eusociality, as “definitively present” in Formicidae, and equivocal or uncertain for †@@@idae or the most recent common ancestor of the Formicoidea.

**31.** Malar space longer than broad (50CB: 0.93). ***Function*:** Malar space proportion is not a functional character but does belie space use within the cranium. This condition is not strictly dependent on compound eye reduction, as other aculeate lineages express small eyes which are situated close to the mandibular bases (*e.g.*, various Tiphiiformes, various Mutillidae, and all Bradynobaenidae).

**36.** Female compound eyes reduced (50CB: 0.96; SFA: 0.96). ***Function*:** Reduced eye size is associated with epigaeic to hypogeic lifestyles, with complete repression occurring as an extreme in the latter. Other aculeate taxa with reduced eyes include various Chrysidoidea (Plumariidae, Scolebythidae, various Bethylidae), various Tiphiiformes with apterous females, several Pompiloidea (*Sapyga* and most Mutillidae), and Bradynobaenidae. All females are apterous in Mutillidae and Bradynobaenidae, while some tiphioid females are not.

**40.** Ocelli repressed in workers (50CB: 0.85; SFA: 0.91). ***Note*:** Because the expressed (or “present”) condition has been assumed to be ancestral to the crown Formicidae (*e.g.*, Bolton 2003), we estimated the marginal likelihoods of the SFA matrix for character polarity under JC (equal state frequencies in matrix, equal transformation frequencies from 0 to 1, 1 to 0; with or without root conditioned on ocellar expression), F81 (equal frequencies, unequal transformation frequencies), and a “Dollo” model in which the root node of the SFA matrix was conditioned as having ocelli, and the stationary frequency of ocellar repression to expression was set to 10^10^. JC with root expression performed the best, with LnL -42.76, followed by JC without prior state conditioning (LnL -43.31) then F81 (- 44.05). The “Dollo” model does not adequately describe the data (LnL -57.80). This reconstruction is unlikely to be due to biased taxonomic sampling given our diversified sampling scheme. To make clear the pattern of ocellar expression and repression, we here outline the distribution of the two character states. Among outgroups, ocelli are sporadically repressed, a state observed in the following terminals: few Chrysidoidea (*Plumarius*, Plumariidae; *Pristocera*, Bethylidae), *Olixon* (Rhopalosomatidae, Vespoidea), various Tiphiiformes, Mutillidae (Pompiloidea), and Bradynobaenidae (Scolioidea). Among Formicidae, ocelli are repressed in workers of the following clades and sampled terminals: Leptanillomorpha, most Poneria (Apomyrminae, Amblyoponinae, Agroecomyrmecinae, Proceratiinae, all Ponerinae except *Harpegnathos*, clearly representing a re-expression), *Lioponera* (the sampled doryline), and most Formicae. Among sampled Formicae, ocelli are expressed in some Myrmeciomorpha and various Formicinae, but not in Dolichoderomorpha or Myrmicomorpha. Among sampled formicines, ocelli are absent in Myrmelachistini (the sistergroup to all other Formicinae), *Cladomyrma* (the sistergroup to Lasiini), various Lasiini, and some members of the major radiation spanned by the Melophorini and Camponotini. In virtually all species, with extremely rare exception, male Formicidae express ocelli, thus the morphogenic machinery for ocellar development is retained in most species of ants, even when they are not observed in workers. If repression of ocelli in workers biases the survival capacity of various clades, then this should also be considered an adaptation, likely conserving resources which would be invested in cuticle and neural mass during development.

**87.** Clypeal disc completely integrated with cranium (50CB: 0.86). ***Function*:** Clypeal variation influences pharyngeal function due to the clypeopharyngeal musculature. Specific functional consequences of the integrated form of the clypeus as observed here are unknown and necessitate further anatomical study. See also char. 139 under Antennoclypeata.

**116.** Frontal carinae present medial to the antennal toruli (50CB: 0.83; SFA for crown Formicidae: 0.92). ***Function*:** The frontal carinae are a pair of anteroposteriorly-oriented to posterolaterally-directed cuticular ridges which form a dorsomedial rim margining the cranial or frontal contact surface which opposes the antennal scapes immediately posterad the toruli. The carinae limit or prevent overextension of the antennae when they are flush with the cranium and rotated mediad. In addition to increasing the safety factor of scapal rotation, when the scapes are abutting the frontal contact surface, they are often provided some degree of protection from damage which may be caused by other insects. That danger from ants and perhaps other insects is a persistent selective factor is evinced by the frequent development of the frontal contact surface into scrobes, observable even in stem Formicidae, such as †*Zigrasimecia*. Additionally, due to their anteroposterior orientation, they may contribute to cranial support against longitudinal compression, as caused by contraction of the mandibular adductor (closer) muscles. Form of the frontal carinae varies considerably across the Formicidae, and they may be lost entirely. ***Polarity*:** 50CB increases to 1.0 one node within Formicidae and remains > 0.95 into the crown. Absence of frontal carinae is a key plesiomorphy of †@@@idae. Note that longitudinal ridges corresponding to the “frontal carinae” of ants are sporadically observed in other groups, such as Ampulicidae, *Chalybion* (Sphecidae), some Scoliidae, and a few Tiphiopompiloides.

**167.** Mesosomal profiles of apterous individuals diagonal, rather than parallel to the ground (CB50: 0.99). ***Function*:** No specific function can be ascribed to the parallel form of the mesosoma, due to the complexity of this tagma. However, this condition is important because it is the first form of mesosomal simplification from the alate condition for which there is evidence. With the pronotum elevated above the propodeum and with the procoxa elongated, the cranium is lifted away from the ground, perhaps enabling ants to engage with a broader range of surface-based objects. Transitions to the parallel condition have occurred frequently within the crown and represent further modifications of internal mesosomal architecture which have yet to be documented in detail. The groundplan condition proposed by Bolton (2003) implies the diagonal condition but is not explicit.

**217.** Female (gyne/queen) mesonotum without paired notauli (CB50: 0.92). ***Function*:** Loss of the notauli has no clear functional consequences, as the basic function of notauli is uncertain. ***Polarity*:** Notauli are strongly supported as absent from the total Formicidae node crownward to each of the major clades (Leptanillomorpha, Poneroformicines [Poneria, Doryloformicia]). Consequently, notaular presence in gynes represents regain, and should be investigated for evidence of function. Male notauli were not evaluated but are inferred intuitively here to be plesiomorphically retained. Loss of female notauli contrasts with Bolton’s (2003) hypothesis that they are retained in total Formicidae.

**249.** Metapleural gland present (50CB: 0.97; SFA for crown Formicidae: 0.98). ***Function*:** The metapleural gland is structurally represented by gland cells which open externally to a secretory surface which is surrounded by cuticular outgrowth which forms a reservoir (“bulla”, “atrium”) with internal hairs directed toward a small orifice located dorsad the metacoxal articulation on the lateral surface of the mesosoma. Form of the bulla and its orifice are meaningfully variable across ant phylogeny, are lost completely in some lineages (*e.g.*, various Camponotini), and are often absent altogether in males. The function or functions of the secretions of this gland have been controversial (*e.g.*, Brown 1968; Hölldobler & Engel-Siegel 1984; reviewed by Yek *et al*. 2011), but because the structure is also observable in workers of stem Formicidae and is otherwise a consistent feature of females among crown group clades, one or more social functions are assumed. In general, the consensus is that the gland has antiseptic thus social hygiene function (Maschwitz *et al*. 1970; Maschwitz 1974; Yek *et al*. 2011). For this reason, myrmecologists have considered the metapleural gland to be an important element during the early evolution of ants (*e.g.*, Wilson 1987), and remains so in most lineages today. ***Note*:** Wilson (1987) stated that presence of the metapleural gland is “the closest we have to a single diagnostic character separating Formicidae from all other [Aculeata]”. This position is supplanted by the modifications of the procoxa (char. 269 of Formicoidea), as the structural form of the procoxa is retained in all workers, queens, and all males—with no known losses—and is present even when the metapleural gland is absent (as in, *e.g.*, various Camponotini or many males), and is also retained when petiolation is completely lost (Boudinot 2015). Similarly, modifications of the meso- and metathoracicocoxal articulations are more consistent in presence of the metapleural gland. A similar procoxal conformation is observed in Myrmosinae (Pompiloidea: Mutillidae), sporadic convergences of meso- and metathoracicocoxal articulations in other apterous Aculeata, but in all cases a suite of features are distinguishing in this case, not the least of which is “geniculate” antennae, as defined for Antennoclypeata below.

**349.** Propodeal spiracle low and lateral (50CB: 0.94; SFA for crown Formicidae: 0.92). ***Function*:** Insufficient anatomical work has been done on the tracheal system of Hymenoptera to induce a well-grounded hypothesis for the locational transformation of the propodeal spiracle reconstructed here. The principal works are Snodgrass (1910, 1935, 1956) on *Apis mellifera* and insects generally, and Keister (1963) on the worker of *Camponotus pennsylvanicus*, following Janet’s serials. Conceivably, the shift of the propodeal spiracle from a position just posterad the hind wing to a more posterior, ventral, and lateral location improves aeration of the meso- and metathoracic thoracicocoxal musculature. That the low and lateral condition is also observed in alates suggests that respiration, importantly exhalation (Hetz & Bradley 2005), during high intensity running may be a selection factor. In various crown and stem taxa, the propodeal spiracle has shifted to a position at or near the dorsolateral propodeal margin (*e.g.*, some †*Gerontoformica*, Pseudomyrmecinae), but in these cases the spiracle remains distant from the metanotal sulcus. Anatomical comparison of Aculeata other than *Apis mellifera* and specific sampling of conspecific workers, queens, and males for a diversified set of ants is a necessity moving forward.

**404.** Helcium strongly narrowed (50CB: 1.0; SFA for crown Formicidae 0.96). ***Function*:** This transformation is from the broadly constricted condition, which is related to rigidity of the helcial articulation during stinging (see char. 403 for Formicoidea above). The extreme condition of “strong narrowing” renders the helcial articulation into a ball-and-socket joint (Bolton 1990; Hashimoto 1996; Perrault 2004), increasing the extent of rotational freedom (Dlussky & Fedoseeva 1988). In various crown ant taxa (*e.g.*, Proceratiinae, Ponerinae, Ectatomminae *s. l.*) a tongue-and-groove system is present, wherein the anterolateral margins of the helcial tergum fit into notches of the petiolar sternum or of the tergosternum in cases of fusion (Perrault 2004; Keller 2011), limiting lateromedial shear. Reversals to the broad and unnarrowed conditions have arisen twice in Leptanillomorpha (Leptanillinae: *Opamyrma*, *Anomalomyrma*) and once in the Poneria (Amblyoponinae).

**498.** Fore wing Mf3 (that abscissa of M immediately distal to the juncture of 1m-cu) at least as long as Rs+M (CB50: 0.93).

**517.** Fore wing crossvein 3rs-m absent (CB50: 0.96). ***Polarity*:** 50CB for loss at the Formicoidea and †@@@idae nodes are 0.63 and 0.61, respectively. We interpret loss of this crossvein as a synapomorphy of Formicidae, with parallel loss or losses in camelomeciids, for which two of the six scored species retain this crossvein.

**558.** Fore wing Cu2 not reaching 1A vein (CB50: 0.66). ***Polarity*:** Support increases crownward into the Formicidae.

**572.** Hind wing basal cell without distal elongation (50CB: 0.98). ***Polarity*:** This is an unstable character at the Formicoidea node, recovered as equivocally absent due to intermediate presence in two of the six camelomeciids sampled. Additionally, SFA for the crown node is affected by the inapplicability of this condition for the Leptanillinae.

**5S.** Eusocial (303t-TD: 0.95; 303t-ST: 0.99). ***Note*:** Eusociality was scored as true for all stem Formicidae with apterous individuals following analysis of the main dataset of 576 characters (see character 4 above). Support for eusociality in the most recent common ancestor of the †Sphecomyrmines, Antennoclypeata, and crown Formicidae are maximal under both analytical regimes.

#### Key plesiomorphy

**\*.** Mandibular dentition was scored as a complex of characters. Retention of unidentate mandibles from the ancestral scoliidan is strongly supported (char. **63**, CB50: 0.96). Absence of the bidentate and tridentate conditions are well supported (chars. **64**, **65**, CB50s: 0.80, 0.88), while absence of the tetra- to many-dentate mandibles are strongly supported (chars. **66**, **67**, CB50s: 0.99, 1.0). This result is contrary to the intuitive hypothesis of Bolton (2003) that the bidentate condition was a groundplan feature of the total Formicidae. There remains some probability that the mandibles may have been tri- or more-dentate, as the CB50 for at least three teeth in the ancestor of the Scoliida is equivocal (0.58), however support for absence in the ancestor of the Formicapoidina is somewhat high (0.83).

***Note*:** An elongate antennomere 3 (char. **158**) has been adduced as an ancestral feature of the Formicidae (Bolton, 2003). CB50 for Formicapoidina: 0.61; Apoidea: 0.74; Formicoidea: 0.73; Formicidae total clade: 0.73. Additionally, due to saturation (parallel loss), the presence of the second tibial spur on the mid- and hind legs is equivocal for the Antennoclypeata, Formicidae crown, and Poneroformicines. Neither labral or clypeal chaetae (“traction setae”; chars. **76**, **97**) are supported as ancestral to the Formicoidea or the Formicidae, implying parallel gains. However, Richter *et al*. (in prep./2022) recently redefined labral setation characters based on the developmental specification of a proximal, transverse line of setae in ants, considering seta size and shape to be an additional variable. With this characterization, a proximal hair line on the labrum is tentatively supported as a synapomorphy of the total clade Formicidae. Further study of antennal sensilla diversity is necessary to resolve the claims of Hashimoto (1990). Finally, metatibial grooming brushes are absent in crown Formicidae, †Brownimeciinae, †Zigrasimeciinae, and †Sphecomyrminae; they are present in at least some †@@@idae and are uncertain in †Haidomyrmecinae. Loss of the metatibial grooming brush may be a synapomorphy of the total clade Formicidae, but direct re-evaluation of the Mesozoic formicoid material is necessary to evaluate this hypothesis (see also Aculeata plesiomorphies above). Key studies on the groundplan of the Formicidae include Wilson *et al*. (1967a,b), Taylor (1978), Bolton (2003), Keller (2011), Boudinot (2015), Liu *et al*. (2019), Richter *et al*. (2019, 2020), Peeters *et al*. (2020), Richter *et al*. (in prep./2022).

### **F.B.A.** Subfamily †Armaniinae Dlussky, 1983

***Clade comprising*: • †Armaniini** Dlussky, 1983 **stat. rev.** (†*Armania* Dlussky, 1983; †*Pseudarmania* Dlussky, 1983). • ***Incertae sedis*** in subfamily: †*Khetania* Dlussky, 1999 **subfam. transfer**; †*Orapia* Dlussky *et al*., 2004.

***Note*:** Bayesian posterior probabilities for synapomorphies listed here are reconstructed for †Armaniini; conformation of †*Orapia* is indicated in the polarity notes. Function is only noted for char. 177 below, as three-dimensional structural form of †Armaniinae is difficult or impossible to ascertain due to the fact that all constituent members of the subfamily are compression fossils. The best support for †Armaniinae is derived from large body size (char. 14), short pronota (char. 177), and presence of 1r-rs (char. 464). The dorsomedial groove of the petiolar node, originally used as a diagnostic feature for the group (Dlussky 1983), was unfortunately not included in the present study.

#### Synapomorphies

**\*.** Postgenal bridge relatively short, being < 0.5 x head length (char. 10: CB50: 1.0), and directed more-or-less ventrally, *i.e.*, comparatively hypognathous (char. 11: CB50: 1.0). ***Polarity*:** This reversal complex is strongly supported from the synapomorphic condition of the Formicoidea. Stemward, support for long and prognathous heads at CB50 0.94 at Formicidae total clade node. †*Orapia* have longer, prognathous heads, conforming to the formicid groundplan.

**14.** Head width ≥ 1 mm, as proxy for large body size (CB50: 0.99). ***Polarity*:** †Armaniini are large compared to other stem Formicidae, a condition strongly supported (CB50 0.91 at total Formicidae node). The cranium of †*Orapia minor* may be narrower than this threshold but is not measurable due to preservation.

**64.** Mandible bidentate (CB50: 1.0). ***Polarity*:** Homoplastic with respect to †Sphecomyrmines; see char. 64 at that node for further information. †*Orapia rayneri* is apparently unidentate.

**177.** Pronotum short, without distinct anterior and dorsal faces (CB50: 0.99). ***Function*:** Shortened pronota are associated with claustral founding in crown Formicidae (Keller *et al*., 2014). Because very little is known or discernable about the biology of †Armaniinae, assuming claustral gynes is tenuous.

**404.** Helcium distinct but broadened relative to total Formicidae (CB50: 0.99). ***Polarity*:** This condition may apply specifically to †*Armania*, as the plesiomorphic narrow condition is observable in †*Pseudarmania* and †*Orapia*.

**464.** Fore wing first radial-radial sector crossvein (1r-rs) present (CB50: 0.94). ***Polarity*:** This transition is strongly supported, as absence is at CB50 1.0 for total Formicidae. Both †*Orapia* species display 1r-rs.

### **F.B.1.** Clade †Sphecomyrmines

***Clade comprising*:** • **†Haidomyrmecinae** Bolton, 2003, • **†Sphecomyrminae** Wilson & Brown, 1967, • **†Zigrasimeciinae** Borysenko, 2017. • ***Incertae sedis*** to subfamily and tribe:

†*Cretomyrma* Dlussky, 1975; †*Dlusskyidris* Bolton, 1994 **fam. transfer**; †*Poneropterus* Dlussky, 1983 **subfam. transfer**.

#### Synapomorphies

**64.** Mandible at most bidentate (CB50: 0.89). ***Polarity*:** In contrast to the total Formicidae (CB50: 0.80) and Antennoclypeata (CB50: 0.98), the bidentate condition is supported as present in the ancestor of the †Sphecomyrmines. Among these stem ants, the second mandibular tooth has been lost in most †*Zigrasimecia*. Condition among known †Haidomyrmecinae is variable (see Barden *et al*. 2020, Perrichot *et al*. 2020).

**76.** Labrum armed with chaetae (“traction setae”; CB50: 0.87). ***Function*:** Coarse, often peg-like labral and clypeal chaetae are situated on the contact surfaces of the respective sclerites, in opposition to the mandibles. With the mandibles at full closure, objects— particularly soft-bodied prey—would be pressed against these setae, which either act as additional teeth in the case of the peg form, or as friction surfaces in the parallel case of †*Camelosphecia* sp. (Fig. 1). Thus, perioral chaetae increase effectiveness of mandibular clasping. ***Polarity*:** Presence of these setae is uncertain for most †*Gerontoformica*. Loss of these setae are implied for †*Sphecomyrma*.

**97.** Clypeus armed with chaetae (CB50: 0.98). ***Function*:** See char. 76 above. ***Note*:** Lost in †*Sphecomyrma*.

***Note 1*:** The lateral portions of the clypeus are extended as broad flat lobes in many †Sphecomyrmines (char. 90, CB50: 0.62 for absence), particularly †*Gerontoformica*. Homology of this condition with †Zigrasimeciini and †Haidomyrmecinae is uncertain. It is possible that these lobes conceal the mandibular mandalus, which is associated with the mandibular gland orifice, and was inferred by Bolton (2003) to be absent in the groundplan of the total Formicidae. The “trulleum” described by Gotwald (1969) is probably restricted to Myrmicinae; application of this term to other subfamilies represents misidentification of the mandibular acetabulum.

***Note 2:*** In the CB50 ancestral state reconstruction analysis, †Haidomyrmecinae and †Zigrasimeciini were recovered as sister taxa, supported by three putative synapomorphies: (1) mandibles rotated in their sockets (see definition of char. **46** for more detail, CB50: 0.95); (2) perceived anterior clypeal margin broadly and deeply concave (char. **80**, CB50: 0.96; char. **90**, CB50: 0.79).

### **F.B.B.** Subfamily †Haidomyrmecinae Bolton, 2003

***Clade comprising*:** • **†Aquilomyrmecini trib. nov.** [type genus: †*Aquilomyrmex* Perrichot *et al*., 2020] (†*Aquilomyrmex*, †*Chonidris* Perrichot *et al*., 2020, †*Dhagnathos* Perrichot *et al*., 2020), • **†Haidomyrmecini stat. nov.** [type genus: †*Haidomyrmex* Dlussky, 1996] (†*Dilobops* Lattke & Melo, 2020, •• **†*Haidomyrmex* clade** [••• **†*Ceratomyrmex* subclade** {†*Ceratomyrmex* Perrichot *et al*., 2016, †*Linguamyrmex* Barden & Grimaldi, 2017, †*Proceratomyrmex* Perrichot *et al*., 2020}, ••• **†*Haidomyrmex* subclade** {†*Haidomyrmex*, †*Haidomyrmodes* Perrichot *et al*., 2008, †*Haidoterminus* McKellar *et al*., 2013}]).

***Note*:** Species of †Aquilomyrmecini and †*Dilobops* were not available for study at the time of character development, thus the synapomorphies listed below apply to the †*Haidomyrmex* clade. See the diagnostic key in Boudinot *et al*. (2020) for defining features of the tribes and suprageneric groups of the †Haidomyrmecinae.

#### Synapomorphies

**12.** Anterior portion of head, bearing mouthparts and clypeus, narrowed and elongated (CB50: 0.99). ***Function*:** This forms a complex with chars. **60**, **81**, **85**, and **138**, associated with the putative trap-jaw mechanism of these ants (*e.g.*, Perrichot *et al*. 2016, 2020, Barden *et al*. 2020), which also bear elongate setae analogous to the “trigger hairs” of extant trap-jaw species. ***Polarity*:** Unique among Aculeata.

**60.** Basal mandibular tooth produced as sabre-like tong (CB50: 0.99). ***Polarity*:** Unique among Aculeata.

**81.** Anterior clypeal margin migrated posteriorly, up to or past antennal toruli, resulting in longitudinal paired lines on the clypeus (CB50: 0.99). ***Polarity*:** Unique among Aculeata.

**85.** Median portion of clypeus (that region between the anterior tentorial pits) longer than wide (CB50: 0.95). ***Polarity*:** Autapomorphic among †Sphecomyrmines; these proportions also observed in various crown Formicidae.

**138.** Antennal toruli located at or posterior to head midlength (CB50: 0.96). ***Polarity*:** Convergently derived in the formicines *Gigantiops* and *Gesomyrmex*, as well as various non-formicoid taxa sampled in the present study (see character definition below for further detail).

**154.** Scape elongate, length ≥ 4 x width (CB50: 0.80). ***Function*:** Possibly related to the elongation of the cranium observed in char. **12**. ***Polarity*:** The shorter scape of †*Haidomyrmodes* is supported as a reversal. Scape elongation is moderately supported as a synapomorphy of the †Sphecomyrminae (CB50: 0.77), and is convergently derived with respect to Antennoclypeata, and a number of non-formicoid taxa sampled in the present study (see character definition below for further detail).

**171.** Mesonotum of worker with two bulges in profile view (CB50: 0.97). ***Polarity*:** Supported as a homoplastic derivation with respect to †Sphecomyrminae (CB50: 0.94) in the CB50 ancestral state reconstruction analysis; condition equivocal for the †Sphecomyrmines node (CB50: 0.55).

**282.** Protibial calcar apically unifid (versus bifid or with vellum) (CB50: 0.93). ***Polarity*:** In the CB50 ancestral state reconstruction analysis, equivocal for the †Haidomyrmecinae + †Zigrasimeciinae and †Sphecomyrmines nodes (CB50: < 0.68).

### **F.B.C.** Subfamily †Sphecomyrminae Wilson & Brown, 1967

***Clade comprising*:** • †*Gerontoformica* Nel & Perrault, 2004, • †*Myanmyrma* Engel & Grimaldi, 2005, • †*Sphecomyrma* Wilson & Brown, 1967.

***Note*:** See Boudinot *et al*. (2020) for diagnostic key to genus.

#### Synapomorphies

**90.** Anterolateral portions of clypeus extended anteriorly as lobate processes (CB50: 0.98). ***Polarity*:** In the CB50 analysis, recovered as a homoplastic transformation with respect to †Zigrasimeciinae (CB50: 0.99).

**120.** Medial frontal carinae evenly curving laterally, forming semicircle around antennal toruli (CB50: 0.94). ***Function*:** The shape of these rims may reflect a distinct range of antennal motion. ***Polarity*:** Autapomorphic among sampled taxa, although *Lioponera longitarsus* (Dorylinae) homoplastically approaches this form. The only exception to this trait in this subfamily is †*Myanmyrma*.

**154.** Scape elongate, length ≥ 4 x width (CB50: 0.77). ***Polarity*:** Scape elongation is moderately supported as a synapomorphy of the †Haidomyrmecinae (CB50: 0.80), and is convergently derived with respect to Antennoclypeata, and a number of non-formicoid taxa sampled in the present study (see character definition below for further detail).

**171.** Mesonotum of worker with two bulges in profile view (CB50: 0.94). ***Polarity*:** Supported as a homoplastic derivation with respect to †Haidomyrmecinae (CB50: 0.97) in the CB50 ancestral state reconstruction analysis; condition equivocal for the †Sphecomyrmines node (CB50: 0.55).

**409.** Helcium infraaxial (*i.e.*, articulatory surfaces of abdominal segment III below midheight of segment) (CB50: 0.94). ***Polarity*:** Among Formicoidea, convergent with *Apomyrma*, Ponerini, Dolichoderomorpha, and Formicomyrmines. Also present in †*Cretomyrma*, which is otherwise unplaceable in the †Sphecomyrmines do the poor preservation.

### **F.B.D.** Subfamily †Zigrasimeciinae Borysenko, 2017

***Clade comprising*:** • **†Zigrasimeciini** Borysenko, 2017 (†*Protozigrasimecia* Cao *et al*., 2020b, †*Zigrasimecia* Barden & Grimaldi, 2013). • ***Incertae sedis*** in subfamily: †*Boltonimecia* Borysenko, 2017.

***Note*:** Because †*Boltonimecia* was not included in the ancestral state estimation analyses, the synapomorphies listed below apply to †Zigrasimeciini only. See also Cao *et al*. (2020b) for further diagnostic traits of the subfamily and tribe.

#### Synapomorphies

**47.** Mandible with extreme torsion such that lateral mandibular margin obscured by basal margin (CB50: 0.98). ***Function*:** The mandibular torsion may allow the mandibles to open and close in such a manner that prey objects are put in maximal contact among the mandibular, clypeal, and labral chaetae (“traction setae”, Cao *et al*. 2020b; Boudinot *et al*. 2020). ***Polarity*:** Autapomorphic.

**68.** Chaetae present on inner mandibular surface (CB50: 0.99). ***Polarity*:** Supported as absent at the †Sphecomyrmines node; uncertain for †Haidomyrmecinae.

**75.** Labrum massively enlarged and apically bilobate (CB50: 0.99). ***Polarity*:** Autapomorphy among sampled taxa (some *Rhopalothrix* Mayr, 1870, *e.g.*, have massive labra; †*Myanmyrma mauradera* (Barden & Grimaldi, 2016) has a massive labrum which is arcuate apically*, i.e.*, without a distal median notch.

**90.** Anterolateral portions of clypeus extended anteriorly as lobate processes (CB50: 0.99). ***Polarity*:** In the CB50 analysis, recovered as a homoplastic transformation with respect to †Sphecomyrminae (CB50: 0.98).

**131.** Antennal scrobe present on face (CB50: 0.98). ***Polarity*:** Autapomorphic among †Sphecomyrmines; also present in †*Boltonimecia* (see Cao *et al*. 2020b). Homoplastic with respect to several clades of crown clade Formicidae.

***Note*:** Tubulation of abdominal segment IV (char. **419**) and a shortened poststernite IV (char. **420**) are equivocally supported as derivations of the †Zigrasimeciini (CB50: < 0.59); the former character is variable among sampled species, and the latter is uncertain for most sampled species.

### **F.B.2.** Clade Antennoclypeata

***Clade comprising*:** • **†Brownimeciinae** Bolton, 2003 (†*Brownimecia* Grimaldi *et al*., 1997), •

**crown clade Formicidae** Latreille, 1809.

#### Synapomorphies

**139.** Clypeus extending posteriorly between antennal toruli (50CB: 0.91; 50CB for Formicidae: 0.99; SFA for Formicidae: 0.91). ***Function*:** Extension of the clypeus between the antennal toruli represents a pronounced reorganization of the internal structures of the dorsal cranium, affecting the clypeus, the toruli, and the supraclypeal area. Because the clypeus bears the origins of the clypeoprepharyngeal muscles (Richter *et al*. 2019, 2020), this may alter suctorial capacity, while the toruli structure the basic rotational mechanism of the antenna via physical interaction with the scapal bulbus. Additionally, the supraclypeal area corresponds spatially to the medial ampulla of the antennal circulatory organ (Matus & Pass 1999, Pass 2000), variation of which may influence the physiology of hemolymph distribution in the antennae. Specific functional consequences of this transformation are uncertain and should be subject to further anatomical study.

**\*.** Scape length was scored as a complex of characters, for which the following three transitions are recorded: scape length ≥ 4 x width (char. **154**, 50CB: 0.88); scape length ≥ 1/2 x head length (char. **155**, 50CB: 0.90); scape length ≥ 1/2 x pedicel + flagellum length (char. **156**, 50CB: 0.90). ***Function*:** Elongation of the scape and compaction of the flagellum result in the traditional “geniculate” conformation of ant antennae. Because the out-lever of the scape increased while the in-lever of the bulbus remains more-or-less constant, elongation of the scape results in greater distance mechanical advantage. For any unit of input force, the excursion distance and speed of the scape apex is relatively greater than that of the ancestrally short scape. Moreover, small contractions of the tentorioscapal muscles result in finer motor control of the scape apex and flagellum. With reduction of flagellar length, the focal point between the antennal apices is zoned just beyond the reach of the mandibles, which was previously proposed as a modification improving social communication (Dlussky & Fedoseeva 1988, Dlussky & Rasnitsyn 2007), whether via worker-queen, worker-worker, or worker-larva antennation.

### **F.B.3.** Family Formicidae Latreille, 1809 crown clade

***Clade comprising*:** • Clade **Leptanillomorpha**, • clade **Poneroformicines**. • ***Incertae sedis*** in crown clade of family: †*Afromyrma* Dlussky *et al*., 2004 **subfam. transfer**, †*Afropone* Dlussky *et al*., 2004 **subfam. transfer**, †*Canapone* Dlussky, 1990 **subfam. transfer**, †*Cariridris* Brandão & Martins-Neto, 1990 **fam. transfer**, †*Cretopone* Dlussky, 1975 **fam. transfer**.

#### Synapomorphies

**\*.** Masticatory mandibular margin at least 3-dentate (char. **65**, CB50: 0.79), but probably multi-dentate, with somewhere between 4–9 teeth (char. **66**, CB50: 0.85). ***Function*:** Increase in mandibular tooth count enhances grip when the mandibles are pulled closed by the adductor muscles, whether the manipulated object is clasped to the clypeus or grasped between the masticatory margins. In Leptanillomorpha, the additional teeth occur on the basal mandibular margin (char. 61), whereas in the Poneroformicines, those teeth are on the masticatory margin (char. 48). ***Polarity*:** Support for additional teeth is strong for the leptanillomorphan and poneroformicine nodes (CB50: > 0.97) and is maximal for the next few internal nodes.

**229.** Epicnemial carina on anterolateral margin of mesopectus present (CB50: 0.87). ***Function*:** The epicnemium is the anterior contact surface of the mesopectus which opposes the procoxae (HAO: 0000294). The epicnemial carinae form the lateral rim of the epicnemial area, thus also forming the margin between the anterior and lateral surfaces of the mesopectus, similar to the occipital and frontal carinae. Often, the epicnemial carinae form the lateral margin of a longitudinal impression of the mesopectus, thus completing a scrobe for the reception of the procoxae. It is possible that the epicnemial carinae prevents slippage of the procoxae posteriorly or posterolaterally, thus increasing the safety factor of procoxal remotion. As for other scrobes, the epicnemial carina in conjunction with the longitudinal epicnemial impression may protect the coxae from damage during predation or combat. The epicnemial carinae are lost in Dolichoderinae (with possible regain in *Dolichoderus*), and most Formicinae (with possible regains). ***Polarity*:** Equivocal for Antennoclypeata.

**324.** Pretarsal claw edentate (50CB: 0.73; SFA: 1.0). ***Function*:** Loss of the tooth of the pretarsal claw may reflect adaptation to epigaeic or hypogaeic habitats during the evolution of crown Formicidae, as previously hypothesized in studies of ecological ancestral state estimation (Lucky *et al*. 2013; Nelson *et al*. 2018). ***Polarity*:** Loss of claw tooth is more strongly supported in Poneroformicines (50CB: 0.92), while presence has limited support Antennoclypeata (50CB: 0.80). Absence is strongly supported further stemward nodes (50CB: > 0.95). SFA analysis, lacking fossils, supports absence of the pretarsal claw tooth for crown Formicidae stemward to the root node (Scolioides), further supporting the important role of fossil material for evolutionary and phylogenetic morphology.

**350.** Propodeal spiracle circular or elliptical, from slit-shaped (50CB: 0.87; SFA: 0.96). ***Function*:** Reversal to slit-shaped from the circular or elliptical shape has occurred multiple times in crown ants including the monotypic Paraponerinae, some Ponerinae, some Dorylinae, Myrmeciinae, various Formicinae (*Melophorus*, Formicini, various Camponotini), some Dolichoderinae (*e.g.*, some Neotropical *Forelius* species), some Ectatomminae *sensu lato*, and some Myrmicinae (*e.g.*, *Ocymyrmex*). These transitions are often associated with very active lifestyles, diurnal surface foraging in exposed habitats (Formicini), and xerophily (*e.g.*, *Melophorus*, *Ocymyrmex*). It is plausible, therefore, that the elliptical-to-circular propodeal spiracle shape is associated with less metabolically demanding lifestyles or adaptation to moist habitats. Further insight may be provided by study of spiracular anatomy and musculature. ***Polarity*:** Retention of slit-shaped spiracle equivocal for Antennoclypeata (50CB: 0.74).

**369.** Petiole with tergosternal fusion (SFA: 0.79). ***Function*:** Tergosternal fusion increases rigidity of the petiole, thus increasing the safety factor during high stress action and improving force transfer from the petiolar depressor muscles. ***Polarity*:** Support for this transition is equivocal for 50CB.

**413.** Prora in form of transverse lip-like projection (50CB: 0.95). ***Function*:** The basic function of the prora is retained (see char. 411 under Formicoidea), but the zone of contact is modified, being lateromedially broader. In addition to increasing the contact surface area, this lateromedial extension may also accommodate for rotation of the helcial joint around the longitudinal axis.

**#.** Based on their micron-scale comparative anatomy of †*Gerontoformica* and various extant Aculeata, Richter *et al*. (in prep./2022) proposed seven of novel groundplan hypotheses for the crown Formicidae (their character indices italicized in parentheses): (*42*) hypostomal corners developed as teeth; (*46*) paramandibular process aligned with hypostomal corner; (*86*) atala / abductor swelling of mandible in the form of a distinct, rounded process; (*97*) mandible “triangular” or shovel-shaped (note that this is supported as a synapomorphy of the Poneroformicines here, char. **48**); (*108*) 0md11 muscle fibers attaching via cuticular filaments; (*128*) 0la11 muscle originating proximally on prementum; and (*143*) *Musculus pharyngoepipharyngalis* comprising two, rather than one bundle.

***Note 1*:** Absence of apicoventral “plantar lobes” of the tarsi (char. **329**) is definitely true for the crown clade Formicidae (CB50: 1.0), but presence of these lobes is uncertain for most stem ants, including †*Brownimecia*.

***Note 2*:** Cranial shape and tentorial conformation are promising character systems for refining the synapomorphy-based definition of crown Formicidae, as †*Brownimecia* has the dome-shaped cranium observed in generalized †Sphecomyrmines. The orientation of the tentoria of crown ants parallel to the longitudinal axis of the mandibular adductors (closers) may be a synapomorphy to the exclusion of †*Brownimecia*, but refined observation is necessary.

***Note 3*:** Location of the compound eyes distinctly in the posterior half of the head (char. **39**) is moderately supported as a plesiomorphy of Antennoclypeata (CB50: 0.82), equivocally so for the crown Formicidae and Leptanillomorpha (CB50: 0.57, 0.53, respectively). Eyes set at or anterior to head midlength (char. **38**) is equivocal for the Poneroformicines, Poneria, Doryloformicia, and Formicidae (CB50: 0.65, 0.74, 0.74, 0.75, respectively). Integration or characterization of additional anatomical data is necessary to inform the reconstruction of this trait.

### **F.B.4.** Clade Leptanillomorpha

***Clade comprising*:** • **Leptanillinae** Emery, 1910 (Opamyrmini Boudinot & Griebenow **trib. nov.**, Leptanillini Emery, 1910 = Anomalomyrmini Taylor, 1990 **syn. nov.**), • **Martialinae** Rabeling & Verhaagh, 2008 (*Martialis* Rabeling & Verhaagh, 2008).

***Note*:** Although not subject to synapomorphy estimation, we provide a formal diagnosis of Opamyrmini **trib. nov.** below following the results of our present analyses plus additional comparative observations. Further, we synonymize Anomalomyrmini Taylor, 1990 with Leptanillini, 1910 **syn. nov.** based on consideration of worker and male morphology contextualized by phylogenomic inference (Griebenow 2020, 2021). To maintain reciprocally monophyletic genera that are diagnosable based on all adult castes *Anomalomyrma* and *Protanilla* will soon be synonymized, while all genera within the tribe Leptanillini will be subsumed into one (Griebenow, in prep.).

#### Synapomorphies

**24.** Occipital carina visible in full-face view (CB50: 0.94). ***Note*:** Visibility of the occipital carina in Leptanillomorpha is a consequence of hyperprognathy (char. 25 below). The carina is visible sporadically amongst Poneroformicines (including many Ponerinae), Myrmeciomorpha, and *Myrmoteras* (Formicinae).

**25.** Cranium hyperprognathous, with postocciput migrated to extreme posterior portion of head, albeit still concealed (CB50: 0.99). ***Function*:** Hyperprognathy directs the mandibles forward such that the long axis of the cranium is approximately in line with the longitudinal axis of the mesosoma. Because the life histories of leptanillomorphans remain largely unknown except for specialization as hypogaeic predators, the ecological consequences of hyperprognathy are uncertain. *Note*: This condition was not observed to be true among other sampled Formicidae. Further anatomical study is recommended.

**37.** Compound eyes repressed (“absent”) in worker caste (CB50: 0.99). ***Function*:** Reduction to complete repression of compound eyes is associated with subterranean ecology. Complete repressing strongly suggests a permanent hypogaeic lifestyle. Compound eye repression also occurs in, for example, *Apomyrma* (Apomyrminae), some Amblyoponinae, and some Ponerinae.

**61.** Teeth present on basal mandibular margin (CB50: 0.95). ***Function*:** As with addition of teeth to the masticatory tool edge of the mandible, increase of teeth on the basal tool edge increases friction. In the cases *Opamyrma* (Leptanillinae), these basal teeth function during the act of clasping objects to the clypeus or labrum which is also armed with chaetae (“traction setae”). The basal tooth of *Leptanilla* (Leptanillinae), in contrast, is oriented such that grasp between the mandibles may be achieved. In *Protanilla* and kin (Leptanillinae), the mandibles function appear to function as trap jaws, albeit without an obvious lock mechanism (Richter *et al*. 2021), whereas the mandibles of *Martialis* (Martialinae) remain a mystery. Teeth on the basal mandibular margin have been independently gained in Poneroformicines (Apomyrminae, Ponerinae: *Harpegnathos*) and various Myrmechoderines. ***Notes*:** The basal mandibular teeth of *Opamyrma* are qualitatively distinct from those of *Leptanilla* and *Yavnella*, being in the form of small serrations that are proximomediad the longitudinal mandibular sulcus and dissimilar in shape from the apical two teeth, which are themselves distinct in shape. In contrast, the teeth of *Leptanilla* and *Yavnella* are distad the sulcus and in the form of the apical tooth pair.

**93.** Anterior clypeal margin evenly linear to concave between mandibular insertions (CB50: 0.95). ***Function*:** No obvious function can be ascribed to these clypeal shapes. Some degree of convexity (char. 92) is the general condition among other Formicidae. ***Notes*:** This estimate implies that the highly variable anteromedian clypeal process of *Leptanilla* is derived.

**116.** Frontal carinae absent (CB50: 0.95). ***Function*:** Loss of frontal carinae in Leptanillomorpha is associated with the far anterior migration of the antennal toruli and hyperprognathy. Parallel losses have occurred under different circumstances in various poneroformicine taxa.

**142.** Antennal toruli directed dorsally (CB50: 0.95). ***Function*:** Reversal of torular orientation from the lateral to dorsal aspect is associated in Leptanillomorpha with hyperprognathy, and functionally aligns the antennae with the longitudinal axis of the body. This modification is again probably due to hypogaeism. This is a reversal from the synapomorphic condition of the Formicoidea, and also occurs in *Apomyrma*, *Discothyrea* (Proceratiinae), and Dorylinae.

**158.** Antennomere III longer than IV (CB50: 0.88). ***Function*:** No discrete function can be ascribed to lengthening of antennomere III. Dedicated, comparative study of antennal anatomy may reveal functional correlations. Importantly, this character is prone to homoplasy among Formicidae.

**\*.** A series of wing venation reduction transformations define Leptanillomorpha based on our ancestral state estimates. These include the following eight characters: **(1)** fore wing “marginal cell 1” (= 2rrs) not pointed or narrowly rounded, possibly being broadly rounded (char. **475**, CB50: 0.89), ***note*** that the pointed or narrowly rounded condition is a reconstructed symplesiomorphy of all sampled taxa, including Evanioidea; **(2)** fore wing free M after Rs+M nebulous, spectral, or absent (char. **501**, CB50: 0.95), ***note*** that this contrasts strongly to the symplesiomorphic condition of all sampled taxa; **(3)** fore wing crossvein 2r-sm (enclosing “submarginal cell 2”) absent (char. **511**, CB50: 0.91), ***note*** that loss of this crossvein has occurred in Poneria (*e.g.*, *Apomyrma*, *Tatuidris* [Agroecomyrmecinae], Proceratiinae) and Doryloformicia (Dorylinae, some Dolichoderinae, some Myrmicinae, and all Formicinae); **(4)** fore wing crossvein 1m-cu (enclosing “discal cell 1”) absent (char. **530**, CB50: 0.90), ***note*** that 1m-cu loss also occurs in Proceratiinae, some Ponerinae, some Dorylinae, some Dolichoderinae, some Myrmicinae, and numerous Formicinae; **(5)** hind wing free Rs (Rsf) spectral or absent (char. **563**, CB50: 0.95), ***note*** that hind wing Rsf is lost sporadically, and among sampled taxa includes *Apomyrma*, some Amblyoponinae, some Proceratiinae, Dorylinae, all Dolichoderinae, *Cladomyrma* (Formicinae), and Crematogastrini (Myrmicinae); **(6)** hind wing r-m crossvein nebulous, spectral, or absent (char. **565**, CB50: 0.95), ***note*** that this is also lost in *Apomyrma*, some Proceratiinae, some Ponerinae, Dorylinae, some Pseudomyrmecinae, Dolichoderinae, and *Cladomyrma* (Formicinae); **(7)** hind wing free Cu (Cuf) nebulous, spectral, or absent (char. **569**, CB50: 0.89), ***note*** that this is also lost in *Apomyrma*, some Proceratiinae, some Ponerinae, Dorylinae, some Dolichoderinae, and Crematogastrini (Myrmicinae); and **(8)** hind wing cu-a absent (char. **570**, CB50: 0.95), ***note*** that among sampled taxa, also lost in *Apomyrma* and *Leptomyrmex* (Dolichoderinae).

***Notes*:** SFA results are not reported for Leptanillomorpha as UCE sequence data were not available for *Martialis* (Martialinae) or *Opamyrma* (Opamyrmini, Leptanillinae). Surprisingly, despite evidence for the monophyly of Leptanillomorpha being only recently uncovered (Borowiec *et al*. 2019), the clade is supported by 16 synapomorphies, plus a series of marginal apomorphies which are not here reported. The degree of convergence between Leptanillomorpha and *Apomyrma* (Apomyrminae, Poneria) is remarkable. Among sampled taxa, the better synapomorphies are those of the cranium, all associated with hyperprognathy and hypogaeism. Because the wing venational characters are largely losses, they should be interpreted as weaker. A key plesiomorphy for Leptanillomorpha is retention of short masticatory mandibular margins (char. 48, CB50: 0.99) as this condition contrasts to the synapomorphic condition of the Poneroformicines. Consequently, the elongate mandibles of *Martialis* and *Protanilla* (Leptanillini, Leptanillinae) are interpreted as independent gains within the leptanillomorph clade.

### F.B.F.A. Tribe Opamyrmini Boudinot & Griebenow trib. nov

**Clade comprising:** Opamyrma hungvuong Yamane *et al*., 2008.

***Note*:** For brevity, the following single letter abbreviations are used: L = Leptanillini, M = Martialinae, O = Opamyrmini.

***Diagnosis***. *Sharing with Leptanillini the following likely synapomorphies (i.e., **Leptanillinae** are diagnosed relative to Martialinae by the following)*: **(a)** male mandibles edentate, nub-like (vs. mandibles bidentate, linear; *Scyphodon* with derived, large, almost plate-like mandibles, *Noonilla copiosa* Petersen, 1968 with derived, elongate mandibles); **(b)** female labrum with chaetae (vs. chaetae absent); **(c)** forewing Mf1 reaching Sc+R+Rs, Rsf1 very small or not developed at all (vs. Mf1 meeting Rsf distant from Sc+R+Rs); **(d)** forewing free R distad pterostigma absent (vs. present; pterostigma not developed in *Leptanilla*, *Noonilla*, *Scyphodon*, *Yavnella*); **(e)** distalmost abscissa of Rsf linear to posteriorly curved, not meeting anterior wing margin (vs. anteriorly curved, meeting Rf); **(f)** when developed, forewing crossvein cu-a interstitial, *i.e.*, anterior junction of cu-a situated at about the split of Mf1 and Cuf from M+Cu (vs. cu-a prefurcal, *i.e.*, proximad; not developed in *Yavnella*, *Scyphodon*, and most *Leptanilla*); **(g)** male genital cupula not ring-like, being either “absent”/indiscernible (some L) or only developed ventrally (O, some L) (vs. cupula ring-like, complete; note that a male *Protanilla* [morphospecies zhg-vn01] has a cupula that is developed ventrally but not dorsally, which implies that this condition is a symplesiomorphy of *Opamyrma* and *Protanilla*). **Note:** The larvae of *Opamyrma*, *Protanilla*, and *Leptanilla* are stenocephalous; whereas the larvae of *Opamyrma* and *Protanilla* are thick (pogonomyrmecoid), those of *Leptanilla* are very thin (leptanilloid). Additionally, there is a longitudinal flange which overhangs the metapleural gland in Leptanillinae but not Martialinae.

*Distinguished from other Leptanillomorpha by the following likely autapomorphies (see Boudinot 2015 for male Martialis, Griebenow 2020, 2021 for Leptanillini, and Yamada et al. 2020 for precise morphological depictions of Opamyrma)*: **(h)** female mandibular teeth distinct in form, with a pointed apical tooth and a subrhomboidal and proximally directed subapical tooth, with these subtended by fine serration (vs. teeth spiniform and widely spaced [M] and teeth subtriangular and roughly equal in size [L], with [some L] or without [other L] teeth on the basal margin); **(i)** female mandible with a single median traction chaeta (vs. chaetae absent [M] or multiple [L]); **(j)** male antennal toruli situated distant from posterior clypeal margin (vs. toruli contacting or very nearly clypeus); **(k)** female occiput grossly enlarged such that it is distinctly visible behind the occipital carina in full-face view; **(l)** female petiole elongate, sessile and subrectangular, length ∼ 2 x width, excluding anterior and posterior constrictions (vs. petiole shorter [L, except for the bizarrely elongated, tubular condition in *Yavnella laventa* Griebenow *et al*., submitted] or pedunculate [M]); **(m)** female petiolar sternum extremely long and narrow, vase-like in ventral view (vs. sternum shorter, wider, or with tergosternal fusion rendering abdominal sternum II undelimited laterally); **(n)** female abdominal segment III de-petiolated, *i.e.*, posterior margins not constricted, postsclerites tube-like (vs. AIII partially [M] to completely petiolated [L]; AIII is completely petiolate in workers and some queens of Leptanillini, with exceptional occurrence among certain *Protanilla* males, *e.g.*, *Protanilla* morphospecies TH03); (**o**) female postsclerites of abdominal segment IV shorter than postsclerites of abdominal segment III and subequal in length to V, VI (vs. longer than IV and slightly [M] to much longer than V, VI [L]; note that postsclerites IV are subequal in length to V and VI in some *Leptanilla*); (**p**) female abdominal tergum IV without distinct transverse sulcus (vs. sulcus distinct); **(q)** female abdominal segment VII elongate, tergum massive, dome-line in lateral view, and incapable of complete retraction (vs. segment IV not distinctly elongate, tergum smaller, capable of complete or nearly complete retraction); **(r)** male gonostyli (= telomeres) extremely long and narrow, length > 10 x width (vs. gonostyli absent or present, but not of this specific conformation); **(s)** male lateropenites (= digiti) malleate, *i.e.*, stalked with a distinct apical ellipsoid process (vs. lateropenites absent or present, but not of this specific conformation; volsellae completely lost in *Noonilla* and *Scyphodon*).

*Further distinguished from Leptanillini by the following conditions*: **(t)** compound eyes of the queen caste, when developed, situated slightly anterad head midlength (possible autapomorphy; vs. slightly posterad; note that the queens of various *Leptanilla* completely lack eyes); **(u)** male mandalus of mandible ≤ 0.5 x mandible length (possible plesiomorphy; vs. mandalus enlarged, > 0.5 x mandible length [autapomorphy]; *Scyphodon*, *Noonilla* with proportionally smaller mandali due to mandibular enlargement or elongation); **(v)** forewing discal cell closed (vs. open [synapomorphy], although note that males of *Anomalomyrma* are unknown); **(w)** petiole tergosternal fusion of all castes incomplete, restricted to presclerites (vs. this fusion complete in all castes [synapomorphy]); **(x)** female abdominal segment III postsclerites unfused (vs. fused [L, M]; polarity uncertain based on present analyses; condition in *Anomalomyrma* not confirmed based on dissection, but apparent from external examination).

### F.B.4’. Clade Poneroformicines

*Clade comprising*: • Clade **Poneria**, • clade **Doryloformicia**.

#### Synapomorphies

**48.** Masticatory margin elongate (CB50: 0.82; SFA: 0.98). ***Function*:** This is the key functional derivation of the Poneroformicines. Elongation of the masticatory tool edge allows for object grasping between the mandibles, improving precision and efficacy of manipulating objects of variable proportions, topographies, and other physical properties. Previously, elongation of the mandibular tool edge had been attributed to the groundplan of the crown Formicidae (Wilson 1987) but is here well- to strongly supported as a synapomorphy of the Poneroformicines. Richter *et al*. (in prep./2022) provide clarification of mandibular morphology, particularly the “triangular” or shovel-shaped form, which should be evaluated in future study. See char. **61** under Leptanillomorpha for further discussion.

**#.** Torular apodeme present (*not sampled in the present study*). **Note:** Richter *et al*. (2020) discovered that a large and variable apodeme develops from the internal medial to anteromedial rim of the torulus, on or near the antennal acetabulum. One muscle, *Musculus frontobuccalis lateralis / M. tentoriooralis* (0hy2), originates on the torular apodeme and inserts on the “oral arm” of the sitophore plate, which is a prepharyngeal sclerite that stabilizes and presumably controls suctorial dynamics. Published observations of apodeme presence include *Brachyponera* (Ponerinae), *Dorylus* (Dorylinae), *Formica* (Formicinae), and *Wasmannia* (Myrmicinae) (Richter *et al*. 2019, 2020; Boudinot *et al*. 2021); published observations of absence among crown ants are confined to the Leptanillomorpha, including *Opamyrma* (Yamada *et al*. 2020), *Protanilla* (Leptanillinae) (Richter *et al*. 2021). Absence of the apodeme in †*Gerontoformica gracilis* and various non-ant Aculeata was recently confirmed by Richter *et al*. (in prep./2022). In the present study, we have also observed presence in *Amblyopone* (Amblyoponinae) and absence in *Leptanilla* (Leptanillinae).

**#.** Three additional, tentative synapomorphies for Poneroformicines were discovered and described in Richter *et al*. (in prep./2022) (their character numbering italicized in parentheses: (*39*) oral margin of hypostoma “shouldered” rather than straight or evenly curved; (*82*) 0an3 muscle fibers originating on dorsal tentorial arms; and (*134*) buccal tube abruptly widened distally.

***Note*:** Because of parallel variation in abscissa presence and cell shape, transformations of wing venation are challenging to reconstruct with confidence within the poneroformicines.

### **F.B.5.** Clade Poneria

***Clade comprising*:** • **Agroecomyrmecinae** Carpenter, 1930 (•• **Agroecomyrmecini** Carpenter, 1930: †*Agroecomyrmex* Wheeler, 1910, †*Eulithomyrmex* Carpenter, 1935, *Tatuidris* Brown & Kempf, 1968; •• **Ankylomyrmini** Bolton 2003: *Ankylomyrma* Bolton, 1973,), • clade **Amblyoponomorpha**, • **Paraponerinae** Emery, 1901 (*Paraponera* Smith, F., 1858), • **Ponerinae** Lepeletier de Saint-Fargeau, 1835, • **Proceratiinae** Emery, 1895 (•• **Probolomyrmecini** Perrault, 2000, **Proceratiini** Emery, 1895).

***Note*:** Because the internal topology of the Poneria is unresolved, synapomorphies are only provided for Amblyoponomorpha and Ponerinae. As Agroecomyrmecinae + Paraponerinae is one of the better supported clades joining subfamilies (Brady *et al*. 2006, Moreau *et al*. 2006, Branstetter *et al*. 2017), a list of possible synapomorphies are included after Note 1 below. See Bolton (2003) and Keller (2011) for synopses of poneriine morphology, including definitions of the Proceratiinae and Paraponerinae. Additionally, Proceratiini is known to be paraphyletic with respect to Probolomyrmecini, but further anatomical and molecular analyses are necessary before tribal recircumscription.

#### Synapomorphies

**145.** Antennal sockets concealed by laterally expanded “frontal lobes” (frontal carinae or medial torular arches) (CB50: 0.85). ***Function*:** The “concealed” state represents an extreme of laterad orientation of the antennal sockets, wherein the medial torular arches are produced laterally over and nearly or completely concealing the lateral torular arch. This development is associated with a proximal bend between the bulbus neck and scape such that the scape is oriented mediad or posteromediad to some degree relative to the main axis of the bulbus neck. No single functional distinction can be drawn at the present time without quantitative mechanical experimentation. This condition is approximated in other taxa, including Myrmicomorpha (Formicae).

**419.** Abdominal segment IV tubulated (CB50: 0.86). ***Function*:** See 419 for †@@@idae above.

**420:** Abdominal poststernite IV shorter than posttergite IV (CB50: 0.81). ***Function*:** Proportional reduction in the length of poststernite IV directs the abdominal apex and sting below the frontal plane, reducing the flexional distance require for the sting to be directed between the mouthparts for prey subduction.

***Note 1*:** Meso- and metatibiae retain both spurs in Poneria; ASE reconstructs the two-spur condition in Formicae as secondary gains. Among poneriines, *Paraponera* represents the only known transition to omnivory.

***Note 2*:** Derived states shared between Agroecomyrmecinae (*Tatuidris*, *Ankylomyrma*) and Paraponerinae include the following: **(1)** mandible at least 10-dentate (although *Ankylomyrma* 5- dentate; char. **67**; other sampled Poneria with this state: *Proceratium*, most Ponerinae); **(2)** antennal scrobe present, receiving entire antenna in repose, with a longitudinal carina or wedge-shaped ridge which separates the scape and flagellum (char. **129**; scrobes with this conformation are unique); **(3)** long axes of radicle and scape offset at distinct angle (char. **149**; among sampled Poneria, also occurring in *Platythyrea*); **(4)** promesonotal articulation immobile due to fusion (char. **165**; among Poneria, also occurring in Proceratiinae, further anatomical study necessary); **(5)** abdominal segment III strongly petiolated (char. **405**; unique among sampled Poneria, exact conformation differs between the two subfamilies; and **(6)** stridulitrum of abdominal pretergite IV present (char. **423**; among Poneria, also occurs in Ponerinae). The extant genera of Agroecomyrmecinae further share: compound eyes situated at posterior apex of scrobe; pronotum with anterior and lateral faces offset by distinct carina; and abdominal tergum IV strongly vaulted.

### **F.B.6.** Clade Amblyoponomorpha

***Clade comprising*:** • **Amblyoponinae** Forel, 1893, • **Apomyrminae** Dlussky & Fedoseeva, 1988 (*Apomyrma* Brown *et al*., 1972).

***Note:*** Synapomorphies are provided below for both the Amblyoponomorpha and the Amblyoponinae.

#### Amblyoponomorpha synapomorphies

**39.** Compound eyes set distinctly in the posterior half of the head (CB50: 0.70). ***Polarity*:** Signal for this synapomorphy increases at the Amblyoponinae node (CB50: 0.97).

**48.** Mandible not triangle shaped, being lateromedially narrow (CB50: 0.72). ***Polarity*:** This is a reversal from the triangular form with an elongate masticatory margin recovered for the Poneroformicines, and likely reflects the specialized diet of the constituent clades, although little is known about the life history *Apomyrma*.

**61.** Mandibular teeth occurring on basal mandibular margin (CB50: 0.81). ***Polarity*:** Also observed in Leptanillomorpha, *Paraponera*, *Harpegnathos*, *Myrmecia*, *Pseudomyrmex*, and Dolichoderinae.

**76.** Labrum bearing chaetae (“traction setae”, “spicules”, or “peg-like setae) (CB50: 0.88). ***Polarity*:** While widespread in stem Formicidae, only observed in Leptanillinae and Amblyoponinae among crown Formicidae.

#### Amblyoponinae synapomorphies

**20.** Posterior head margin broadly concave in full-face view (CB50: 0.96). ***Polarity*:** Among sampled Formicidae, also observed in *Odontomachus*, Dolichoderomorpha, *Myrmelachista*, *Aphomomyrmex*, *Myrmoteras*, *Camponotus*, *Solenopsis*, and *Wasmannia*.

**49.** Mandible elongate bar-shaped (CB50: 0.97). ***Polarity*:** Among sampled Formicidae, long narrow mandibles are also observed in *Odontomachus*, *Harpegnathos*, *Myrmecia*, and *Myrmoteras*, although they are known to occur in other groups (*e.g.*, Strumigenina, Dacetina).

**56.** Mandibular teeth in the form of fangs (CB50: 1.0). ***Polarity*:** Unique among sampled Formicidae, although the teeth of *Myrmecia* are similar.

**97.** Anterior clypeal margin with chaetae (“traction setae”, “clypeal pegs”) (CB50: 0.99).

***Polarity*:** Unique among crown Formicidae, including *Apomyrma*.

**103.** Genae with anterolateral spiniform processes (CB50: 0.99). ***Polarity*:** Unique among crown Formicidae. Also present in †*Brownimecia*.

**375.** Petiolar tergum without distinct posterior face (CB50: 0.99). ***Polarity*:** This reversal is nearly unique among crown Formicidae but is also observed in Leptanillini.

**390.** Petiolar tergum with distinct laterotergites (CB50: 0.96). ***Polarity*:** Among Poneria, also present in some Ponerinae (*Platythyrea*, *Neoponera*).

**404.** Helcium broad, although still somewhat constricted (CB50: 0.95). ***Polarity*:** Considered traditionally to be an ancestral feature, a broad helcium is here strongly inferred to be a synapomorphy of the Amblyoponinae and may involve concomitant muscular derivations.

**410.** Helcium supraaxial, *i.e.*, set above segment midheight (CB50: 0.96). ***Polarity*:** Among crown Formicidae, also occurring in Leptanillini.

**458.** Pterostigma enlarged (CB50: 0.96). ***Polarity*:** Also occurring in *Apomyrma*, Leptanillini, *Proceratium*, *Platythyrea*, *Lioponera*, some Dolichoderinae, and *Stegomyrmex*.

### F.B.L. Subfamily Ponerinae Lepeletier de Saint-Fargeau, 1835

***Clade comprising*:** • **Platythyreini** Emery, 1901, • **Ponerini** Lepeletier de Saint-Fargeau, 1835.

***Note*:** Synapomorphies are provided below for both the Ponerinae and the Ponerini. The internal topologies of both tribes are unresolved. For mutual diagnoses of the tribes, see also Bolton (2003).

#### Ponerinae synapomorphies

**35.** Males with compound eyes notched medially (CB50: 0.60). ***Polarity*:** Support stabilizes for the Ponerini (CB50: 0.94). Medially emarginate male compound eyes were also observed in *Stigmatomma*, *Paraponera*, Myrmeciomorpha, *Dolichoderus*, some Formicinae, Ectatomminae, and some Myrmicinae.

**38.** Compound eyes situated in anterior half of head (CB50: 0.96). ***Polarity*:** Unique among Poneria; among sampled Ponerinae, reversed to a more posterior position in *Diacamma*.

**67.** Mandible at least 10-dentate (CB50: 0.89). ***Polarity*:** Support stabilizes for the Ponerini (CB50: 0.95). Many-dentate mandibles among Poneria are also observed for *Paraponera* and *Tatuidris*. See Dolichoderinae below for distribution among Doryloformicia.

**117.** Medial frontal carinae fused with medial torular arches (CB50: 0.97). ***Polarity*:** This condition is approximated in various Amblyoponinae Bolton (2003), but was does not occur in the sampled terminals

**369.** Petiole without tergosternal fusion (CB50: 0.77). ***Polarity*:** Except for *Adetomyrma* (Amblyoponinae, not sampled), unique among Poneria. Independently derived in Ectatomminae.

#### Ponerini synapomorphies

**59.** Male mandible reduced to nonfunctional spiniform or spatulate processes (CB50: 0.99). ***Polarity*:** This is one of the best defining feature of Ponerini, as observed by Bolton (2003). Derived independently several times among crown Formicidae, including *Opamyrma*, *Apomyrma*, and various Myrmicinae (although none sampled in the present study).

**123.** Frontal carinae close-set and curving medially posterad antennal toruli, thus having a “pinched” (CB50: 0.98). ***Polarity*:** Also recognized as a strong feature for the tribe by Bolton (2003). Not observed among other sampled crown Formicidae.

**409.** Helcium infraaxial (CB50: 0.94). ***Polarity*:** Except for *Apomyrma*, unique among Poneria. Reversed to the axial state in some Ponerinae, but these taxa not sampled in the present study.

### F.B.5’. Clade Doryloformicia

*Clade comprising*: • Dorylinae Leach, 1815, • clade **Formicae**.

***Note*:** No unequivocal synapomorphies were recovered for the Doryloformicia, which remains one of the most morphologically insoluble groups known since the first major molecular phylogenetic studies of the Formicidae (Moreau *et al*. 2006, Brady *et al*. 2006). We strongly suspect that this loss of signal is due to pruning of the intermediate lineages of Dorylinae at the End Cretaceous.

### **F.B.7.** Clade Formicae

*Clade comprising*: • Clade **Myrmechoderines**, • clade **Formicomyrmines**.

#### Synapomorphies

**399.** Abdominal segment III without tergosternal fusion, except for helcium (CB50: 0.97). ***Function*:** Loss of fusion allows dorsoventral flexion of the tergum and sternum relative to one another and is interpreted to be regained in Tapinomini (Dolichoderinae), the *Prenolepis* genus group (Formicinae: Lasiini), Plagiolepidini exclusive of *Anoplolepis* (Formicinae), and Ectatomminae *sensu lato*. ***Polarity*:** Tergosternal fusion is supported for Poneria (CB50: 0.98), but weakly so for crown Formicidae, Poneroformicines, and Doryloformicia (CB50: 0.72, 0.74, 0.67, respectively).

**409.** Helcium infraaxial, *i.e.*, set below segment midheight (CB50: 0.58). ***Polarity*:** Although equivocal, the majority of Formicae have an infraaxial helcium. Exceptions include Myrmeciomorpha, *Oecophylla*, *Wasmannia*, and *Crematogaster*.

**551.** Fore wing crossvein 1cu-a anterior junction considerably proximal to branching of M+Cu (CB50: 0.76). ***Polarity*:** This condition occurs sporadically in Poneria.

**B.** Diet omnivorous (303t-U-JC, 303t-U-F81, 303t-O-JC: 0.99). ***Polarity*:** From the morphological perspective, most recently codified by Bolton (2003), this is a novel and perhaps surprising synapomorphy. Among Formicae, *Myrmecia* (Myrmeciinae) was scored as purely predaceous, while *Acanthoponera* (Ectatomminae *sensu lato*) was scored as unknown due to the paucity of information for this genus. *Nothomyrmecia* (Myrmeciinae) is known to supplement its diet with foraged liquid, while Dolichoderomorpha, Formicinae, and Myrmicomorpha are virtually all omnivores, acting as scavengers, predators, and angiosperm-derived liquids, whether directly from extrafloral nectaries or hemipteran anal waste. It is plausible that this dietary transition is accompanied by dietary pathway derivations.

***Note*:** Development of a petiolar peduncle is equivocal at this node (char. **370**, CB50: 0.56 for sessile condition). A peduncle is present in Myrmeciomorpha (scored as TRUE for all sampled terminals, CB50: 0.74), Aneuretinae, and is moderately supported as ancestral for the Myrmicomorpha (CB50: 0.78). If the petiole were pedunculate in the ancestral formican, then the sessile condition would be derived in parallel between Dolichoderinae and Formicinae.

### **F.B.8.** Clade Myrmechoderines

*Clade comprising*: • Clade **Dolichoderomorpha**, • clade **Myrmeciomorpha**.

***Note*:** No unequivocal synapomorphies of the myrmechoderine clade were recovered. However, multiseriate mandibular dentition (*i.e.*, teeth with alternating sizes, char. **55**) is a possible shared derived trait of the Dolichoderomorpha and Myrmeciomorpha.

### **F.B.9.** Clade Myrmeciomorpha

***Clade comprising*:** • **Myrmeciinae** Emery, 1877 (Myrmeciini Emery, 1877, Prionomyrmecini Wheeler, 1915), • **Pseudomyrmecinae** Smith, M. R., 1952.

***Note*:** Because only one myrmeciine terminal was sampled in the present study, synapomorphies within this clade are only presented for Pseudomyrmecinae. See Ward & Brady (2003) for inferred morphological synapomorphies of Myrmeciinae.

#### Synapomorphies

**24.** Occipital carina visible in full-face view (CB50: 0.82). ***Polarity*:** The occipital carina is strongly supported as concealed for the Myrmechoderines (CB50: 0.99). Infrequently derived among other Formicae (*e.g.*, *Myrmoteras*).

**35.** Medial margins of male compound eyes emarginate (CB50: 0.66). ***Polarity*:** Despite the equivocal support, true for all sampled Myrmeciomorpha and independently gained among some Dolichoderinae. Support is moderate for an “entire” or not-notched medial eye margin at the myrmechoderine node (CB50: 0.81).

**158.** Antennomere III longer than IV (CB50: 0.63). ***Polarity*:** True for all sampled Myrmeciomorpha, as well as for the sampled dolichoderomorphs *Aneuretus*, *Bothriomyrmex*, *Dolichoderus*, and *Linepithema*.

**\*.** Two mesotibial (char. **310**, CB50: 0.72) and two metatibial spurs (char. **312**, CB50: 0.74) present. ***Polarity*:** Loss of one of the spurs of each of the meso- and metatibiae are supported as a synapomorphy of the Doryloformicia (CB50: 0.92), countering the Dollo-like expectation that once lost, spurs will not be regained. Notably, some male Ectatomminae express a second tibial spur, suggesting that the genetic capacity for developing the second spur is retained in at least some Formicae. Fossils may be crucial for testing the reconstruction hypothesis from our estimates; for example, in its original description, the unsampled Cenozoic fossil †*Formicium* (†Formiciinae Lutz, 1986) is stated to have “two short, strong [metatibial] spurs”. Likewise, reevaluation of the putative Cretaceous ectatommine, †*Canapone*, is necessary as the original description did not include observations of the spurs. Future discoveries will, of course, provide critical information.

**324.** Pretarsal claws bidentate (CB50: 0.81). ***Polarity*:** Clearly an independent re-gain among crown Formicidae, which are reconstructed as lacking additional dentition (CB50: 0.73; Leptanillomorpha CB50: 0.96; Poneroformicines CB50: 0.92). Additional teeth on the pretarsal claws are also expressed in the poneriines *Paraponera*, *Platythyrea*, *Harpegnathos*, and *Leptogenys*, and in the formican clade Ectatomminae. Bolton (2003, pp. 28, 29) notes that some small *Tetraponera* species have secondarily reduced or lost pretarsal claw teeth.

**369.** Abdominal segment II tergosternal fusion absent (CB50: 0.81). ***Polarity*:** Fusion is equivocally supported as ancestral for the Myrmechoderines (CB50: 0.69), but strongly to moderately supported as retained in the Dolichoderomorpha (CB50: 0.93) and Formicomyrmines (CB50: 0.84). Absence of fusion is universal among Myrmeciomorpha (see, *e.g.*, Bolton 2003).

**376.** Node of petiolar tergum anteroposteriorly longer than dorsoventrally tall (CB50: 0.59). ***Polarity*:** Although this synapomorphy is equivocally supported, the petiolar node is longer than tall in all sampled Myrmeciomorpha and is not so for most sampled Formicae.

**390.** Petiole with well-defined laterotergites (CB50: 0.70). ***Polarity*:** As for char. 376, equivocally supported but true for Myrmicomorpha and false for all Formicae, with the exception of Ectatomminae in this case.

**405.** Abdominal segment III (metasomal II) strongly petiolated (CB50: 0.87). ***Polarity*:** This implies reversal in *Nothomyrmecia*, contrary to the hypothesis of Taylor (1976). Convergently derived to a stronger degree in Myrmicinae. An “undeveloped” postpetiole is maximally supported as the condition for the Myrmechoderines and Formicae.

**409.** Helcium of abdominal segment III axial, *i.e.*, set at segment midheight (CB50: 0.84). ***Polarity*:** The infraaxial conditions is equivocally supported for Myrmechoderines (CB50: 0.59) and Formicae (CB50: 0.58).

***Note 1*:** Support values for conditions shared among all sampled Myrmeciomorpha are surprisingly low. This may be due to influence from the fossil dolichoderines (†*Chronomyrmex*, undescribed Burmite sp.) which were unconstrained in the ancestral state reconstruction analyses. A second factor contributing to the low support may be rate variability of the traits among crown Formicidae, particularly as Myrmeciomorpha display a number of features which are variable among Poneria.

***Note 2*:** Large body size, as measured by head width (≥ 1.0 mm) as a proxy (char. **14**), may be ancestral for the Myrmeciomorpha. All scored terminals have relatively large heads, but the support for one or the other condition was estimated to be perfectly equivocal at this node (CB50: 0.50).

### F.B.P. Subfamily Pseudomyrmecinae Smith, M. R., 1952

***Clade comprising*:** • **Pseudomyrmecini** Smith, M. R., 1952 (*Myrcidris* Ward, 1990, *Pseudomyrmex* Lund, 1831, *Tetraponera* Smith, F., 1852).

***Note*:** For additional defining characters, see also Ward & Downie (2005).

#### Synapomorphies

**139.** Clypeus not extending posteriorly between antennal toruli (CB50: 0.90). ***Polarity*:** This reversal is strongly supported for the Pseudomyrmecinae and crown Formicinae (CB50: 0.98; 0.85 for total Formicinae). The plesiomorphic condition of extending between the toruli is maximally supported for the Dolichoderomorpha, Myrmechoderines, and Formicae, and strongly so for the Myrmeciomorpha (CB50: 0.94) although all Myrmeciinae display this condition.

**179.** Pronotum with dorsolateral longitudinal margination between lateral and dorsal surfaces (CB50: 0.91). ***Polarity*:** This putative synapomorphy should be tested with a much broader sample of Pseudomyrmecinae, due to the known variability of pronotal margination throughout the subfamily. Maximally supported as absent for the Dolichoderomorpha, Myrmechoderines, and Formicae, and strongly supported as absent for the Myrmeciomorpha (CB50: 0.94).

**295.** Female meso- and/or metafemora swollen in appearance (CB50: 0.90). ***Polarity*:** This mechanical adaptation probably associated with arboreality is recovered as absent in the dolichoderomorphan (CB50: 0.99), myrmeciomorphan (CB50: 0.94), myrmechoderine (CB50: 1.0), and formican (CB50: 1.0) nodes. Expanded taxon sampling and detailed anatomical study within the subfamily with improve our understanding of this evolution of this trait.

**376.** Node of petiolar tergum anteroposteriorly longer than dorsoventrally tall (CB50: 0.93). ***Polarity*:** Equivocal for Myrmeciomorpha (CB50: 0.58), although this clade more strongly supported as having a wide node (char. **377**, CB50: 0.89). A node which is wider than long is maximally supported for the Formicae, Dolichoderomorpha, and Formicomyrmines, and strongly so for the Myrmechoderines (CB50: 0.99).

**423.** Stridulitrum on abdominal pretergite IV (“gastral” pretergite I) present (CB50: 0.91). ***Polarity*:** Among Formicae, also present in some Ectatomminae, and various Myrmicinae. Strongly supported as absent for the Myrmeciomorpha as a whole, and maximally so for the Dolichoderomorpha, Myrmechoderines, and Formicae

### F.B.9’. Clade Dolichoderomorpha

***Clade comprising*:** • **Aneuretinae** Emery, 1913 (Aneuretini Emery, 1913), • **Dolichoderinae** Forel, 1878.

***Note*:** See Bolton (2003) for definition of Aneuretinae.

#### Synapomorphies

**21.** Occipital carina absent (CB50: 0.79). ***Polarity*:** The carina was observed to be absent in all sampled dolichoderomorph species and is equivocally supported as present in Formicae probably due to the influence of the carina-less Formicinae.

**38.** Compound eyes located in anterior half of head (CB50: 0.55). ***Polarity*:** More-posteriorly set compound eyes were observed in *Dolichoderus* and *Leptomyrmex* among sampled dolichoderomorphs. It may be the case that Aneuretinae and Dolichoderinae independently derived anteriorly set eyes, as an alternative to the hypothesis of reversal in the two genera noted here. Anatomical study of the ocular nerves may provide further insight into this question.

**55.** Masticatory mandibular margin with serrated teeth of alternating size (CB50: 1.0). ***Polarity*:** Equivocal for Myrmeciomorpha and Myrmechoderines nodes (CB50: < 0.64). We hypothesize that the future inclusion of *Nothomyrmecia* and †*Prionomyrmex* will result in recovery of this condition as a synapomorphy of the Myrmechoderines.

**95.** Anterior clypeal margin with a median emargination or notch which is shouldered laterally by convex laminar processes (CB50: 0.99). ***Function*:** Seifert (2016) hypothesized that the medial notch may allow the mouthparts greater capacity for protrusion for the purposes of consuming liquids, including food and water, particularly in tight spaces. The development of the notch is particularly refined in *Tapinoma*, the focal group of Seifert’s study. ***Polarity*:** Autapomorphy among Formicidae and strongly reduced in *Bothriomyrmex*.

**177.** Pronotum very short, without strong anterior and dorsal faces in profile view (CB50: 0.83). ***Polarity*:** Convergently derived with respect to various Poneria (*Paraponera*, *Tatuidris*, Proceratiinae), as well as Formicinae and Myrmicinae; see note for Formicomyrmines below for further detail.

**213.** Parascutal carina of alates absent (CB50: 0.67). ***Polarity*:** Regained in *Dolichoderus*, and convergently lost in Formicinae.

**294.** Mesotrochantellus indistinct or completely absent (CB50: 0.94). ***Polarity*:** The presence of trochantella is more variable among crown ants than has been assumed (*e.g.*, Grimaldi *et al*. 1997). Here, the mesotrochantellus was recovered as an ancestral state of the crown clade Formicidae (CB50: 0.68), Leptanillomorpha (CB50: 0.83), Poneroformicines (CB50: 0.70), Doryloformicia (CB50: 0.70), Formicae (CB50: 0.61), Myrmechoderines (CB50: 0.60), and the Formicomyrmines (CB50: 0.51). Although these estimations largely fall in the “equivocal” category, they reflect the widespread presence of mesotrochantella among extant species, as documented by Keller (2011) in his SEM Atlas of Formicidae, available on AntWeb.

**356.** Propodeal lobes reduced or absent (CB50: 088). ***Polarity*:** Propodeal lobes are retained in Myrmeciomorpha (CB50: 0.81) and Myrmicomorpha (CB50: 0.97). They are reconstructed as present in the ancestor of the Formicomyrmines (CB50: 0.79), with loss in the Formicinae (absent in all sampled species except *Oecophylla*).

**382.** Subpetiolar process absent (CB50: 0.84). ***Polarity*:** Present universally in sampled Leptanillomorpha, Poneria, and Dorylinae. Among Formicae, convergently lost in Pseudomyrmecinae and Formicinae, although with regains in that subfamily.

**385.** Anterolateral carinae of petiolar tergum absent (CB50: 0.80). ***Polarity*:** Present in most sampled ant taxa with the exception of *Opamyrma*, *Apomyrma*, and Formicinae, in addition to the Formicomorpha.

**411.** Prora of abdominal sternum III absent (CB50: 0.83). ***Polarity*:** Similar to char. 385, present in most taxa except *Opamyrma*, *Apomyrma*, Dolichoderomorpha, and Formicinae, although also lost in *Crematogaster*, among sampled Myrmicinae.

**418.** Cinctus (constriction) of abdominal segment IV absent (CB50: 1.0). ***Polarity*:** Among sampled crown Formicidae, convergently lost in *Apomyrma*, *Simopelta*, and Formicinae.

***Note*:** Reduction of the sting was recovered as a moderately-supported synapomorphy of Dolichoderomorpha (char. **435**, CB50: 0.86). We strongly suspect this is a spurious result caused by the variable placement of stem Dolichoderinae, particularly the undescribed species from Burmite. Data from stem Aneuretinae are highly desirable.

### F.B.R. Subfamily Dolichoderinae Forel, 1878 total clade

***Clade comprising*:** • **†Miomyrmecini** Carpenter, 1930, • **†Zherechinini** Dlussky, 1988, • **Bothriomyrmecini** Dubovikoff, 2005, • **Dolichoderini** Forel, 1878, • **Leptomyrmecini** Emery, 1913, • **Tapinomini** Emery, 1913. • Mesozoic taxa ***incertae sedis*** in subfamily:

†*Chronomyrmex* McKellar *et al*., 2013, †*Eotapinoma* Dlussky, 1988.

***Note*:** Placement of the Burmite dolichoderine was uncertain in the ancestral state reconstruction topology searches, thus the CB50s here are reported for both the total and crown nodes excluding that fossil, where total includes †*Chronomyrmex* plus the crown clade.

#### Total clade synapomorphy

**61.** Mandibular teeth present on basal margin (CB50: 0.92). ***Polarity*:** Strongly supported as true for crown Formicidae (CB50: 0.92). Uncertainty about the condition of basal teeth due to the Burmite dolichoderine was propagated among the dolichoderomorphan and myrmeciomorphan nodes, where such teeth were equivocally reconstructed as absent (CB50: 0.71 for both). Basal teeth are strongly to maximally supported as absent for the Myrmechoderines (CB50: 0.98) and Formicae, respectively. Also true for the Burmite dolichoderine, despite its variable placement in topology searches.

**435.** Sting reduced to non-functionality and terminal opening of metasoma without coronula of setae (CB50: 1.0). ***Polarity*:** Sting reduction is convergent with Formicomorpha, with the primary external difference being the lack of setae surrounding the terminal abdominal opening.

#### Crown clade synapomorphies

**67.** Mandible at least 10-dentate (CB50: 0.94). ***Polarity*:** Mandibles were strongly supported as fewer dentate for the total Dolichoderinae (CB50: 0.99) and maximally so for higher nodes to the Formicae.

**87.** Disc of clypeus, including medial and lateral portions, produced anteriorly from cranium (CB50: 0.89). ***Polarity*:** The clypeal disc is strongly supported as less produced or not produced at all for total Dolichoderinae (CB50: 0.95), Myrmechoderines (CB50: 1.0), and Formicae (CB50: 0.99). The produced condition is supported as independently derived in the Myrmicomorpha (CB50 for Ectatomminae: 0.97; for Myrmicinae: 0.99), and is observed in Formicinae and Poneria.

**563.** Hindwing Rsf2 spectral or absent (CB50: 0.91). ***Polarity*:** Strongly to maximally supported as tubular higher nodes to the Formicae.

***Note*:** Support for eye placement anterior to head midlength (char. **38**) is equivocal for the Dolichoderinae but would be supported as a derived condition given that the Formicae node is moderately supported as having eyes at or somewhat posterior to head midlength (CB50: 0.79).

### F.B.8’. Clade Formicomyrmines

*Clade comprising*: • Clade **Formicomorpha**, • clade **Myrmicomorpha**.

***Note 1*:** No unequivocal synapomorphies were recovered for the Formicomyrmines in our dataset. However, a head which is broader than long (char. **15**) is surprisingly observed in most members of the Formicomyrmines. The lack of signal at this node in our analysis is unaccounted for.

***Note 2*:** The development of the alate pronotum (char. **177**), associated with colony foundation strategy (Keller *et al*. 2014), is equivocal at the formicomyrmine and myrmicomorphan nodes (CB50 for both: 0.57), very weakly supporting the derived short and weak condition. This tentative result mirrors the parsimony-based reconstruction for this trait in Keller *et al*. (2014). However, it should be noted that their starting phylogeny treated the long-pronotal subfamily Ectatomminae as sister to Formicinae + Myrmicinae based on the multigene phylogeny of Rabeling *et al*. (2008) (similar to Moreau *et al*. 2006, Brady *et al*. 2006), rather than as sister to the Myrmicinae as recovered from the refined sampling and modeling of Kück *et al*. (2011), Moreau *et al*. (2013), Economo *et al*. (2018), and Borowiec *et al*. (2019), and with phylogenomic data by Branstetter *et al*. (2017a). Future study incorporating more anatomical information will be necessary to resolve the transformation of this important trait.

***Note 3*:** Richter *et al*. (in prep./2022) proposed five groundplan hypotheses for the Formicomyrmines (their character indices italicized in parentheses): (*46*) paramandibular process shifted mediad relative to hypostomal tooth, with concave connection; (*47*) secondary hypostomal knob/bulge present; (*50*) medial tentorial lamella forming medium-sized lobe; (*113*) stipital flange not raised; and (*147*) maxillary gland present. The taxon sampling of this study was very limited, so these should be considered preliminary hypotheses.

### **F.B.10.** Clade Formicomorpha

***Clade comprising*:** • Clade **Formicomorpha** (**†**Formiciinae Lutz, 1896 [**†**Formiciini Lutz, 1896], Formicinae Latreille, 1809).

***Note 1*:** See Lutz (1986) for rationale of †Formiciinae placement in Formicomorpha.

### **F.B.T.** Subfamily Formicinae Latreille, 1809

*Clade comprising*: • **Myrmelachistini** Forel, 1912, • **lasioformicine** clade (•• **Lasiini**, Ashmead 1905, **core Formicinae** [••• **Melophorini** Forel, 1912, **formicoform** radiation {•••• **Camponotini** Forel, 1878, **Formicini** Latreille, 1809, **Gesomyrmecini** Ashmead, 1905, **Gigantiopini** Ashmead, 1905, **Myrmoteratini** Emery, 1895, **Oecophyllini** Emery, 1895, **Plagiolepidini** Forel, 1886, **Santschiellini**, Forel, 1917). • Mesozoic taxa *incertae sedis* in subfamily: †*Kyromyrma* Grimaldi & Agosti, 2000.

#### Synapomorphies

**39.** Eyes set distinctly in posterior half of head (CB50: 0.92). ***Polarity*:** The markedly posterior position of the compound eyes of Formicinae is a condition shared with some Leptanillinae, Amblyoponinae, Agroecomyrmecinae, *Leptomyrmex*, and various Myrmicinae. Eyes are set less posteriorly in Formicinae, however, in *Myrmelachista*, *Cladomyrma*, *Oecophylla*, *Aphomomyrmex*, and *Myrmoteras*, suggesting reversals.

**139.** Clypeus not extending posteriorly between antennal toruli (CB50: 0.85). ***Polarity*:** Support for this condition increases at the crown formicine node (CB50: 0.98). Among sampled crown formicines, the posterior extension of the clypeus was observed in *Myrmelachista*, *Lasiophanes*, *Santschiella*, *Gesomyrmex*, and *Aphomomyrmex*. The condition of extension between the toruli is strongly to maximally supported as a plesiomorphy (CB50 for Formicomyrmines: 0.94; CB50 for Formicae: 1.0). This condition is homoplastic with respect to Pseudomyrmecinae (see above).

**213.** Parascutal carina of mesoscutum, above forewing base, absent (CB50: 0.63). ***Polarity*:** The reported support value applies to the crown node, as alates for stem Formicinae are unknown. Although the value is equivocal, none of the examined crown Formicinae express the parascutal carina, supporting the interpretation of this condition as a synapomorphy of the group. The carina is strongly supported as a plesiomorphy of the Formicae (CB50: 0.94), moderately so for the Formicomyrmines (CB50: 0.89), and maximally so for the Ectatomminae and Myrmicinae.

**264.** Metacoxae wideset (CB50: 1.0). ***Polarity*:** This is a condition attributed to the Lasiini tribe group by Bolton (2003) which is here maximally supported as a synapomorphy of the total Formicinae, indicating that the close-set condition of various tribes is a reversal. This condition is not observed among Dolichoderinae, despite the strong convergence.

**294.** Mesotrochantellus indistinct to completely absent (CB50: 0.98). ***Polarity*:** Among sampled Formicinae, the mesotrochantellus is redeveloped in *Myrmoteras*. Lack of a distinct mesotrochantellus is supported as convergent with some Poneria, the Dolichoderinae, and various Myrmicinae.

**356.** Propodeal lobes indistinct or absent (CB50: 0.84). ***Polarity*:** Support for indistinct to absent propodeal lobes increases at the crown node (CB50: 0.99), and is convergent with respect to the Dolichoderomorpha, and various Leptanillomorpha (with the exception of *Opamyrma*). Among sampled Formicinae, propodeal lobes were redeveloped in *Oecophylla*.

**378.** Petiolar node squamiform, *i.e.*, much taller than wide (CB50: 0.98). ***Polarity*:** Convergent with respect to various Dolichoderinae and *Hypoponera* (Ponerinae). The node transformed to other conformations among sampled Formicinae in *Santschiella* and *Oecophylla*.

**382.** Subpetiolar process absent (CB50: 0.98). ***Polarity*:** Among sampled crown Formicinae, re-expressed in *Santschiella* and *Myrmoteras*. Loss of the subpetiolar process is convergent with respect to *Pseudomyrmex* and the Dolichoderinae. This condition is probably correlated with the derivation of wide-set metacoxae, which allows the petiole to fit broadly between the legs, rather than relying on the process to prevent torsion.

**385.** Petiolar tergum without anterolateral carinae (CB50: 0.98). ***Polarity*:** These carinae are maximally supported as absent for the crown Formicidae. Loss is convergent with *Opamyrma*, *Apomyrma*, and the Dolichoderomorpha.

**407.** Abdominal poststernite III (“gastral I”) low shouldered posterior to the narrow helcium *i.e.*, with the tergosternal line strongly curving anteriorly before narrowly curving posterolaterally (CB50: 0.99). ***Polarity*:** This is another feature attributed by Bolton (2003) to his broadly conceived (*i.e.*, artificial) Lasiini, further supporting the lasiine syndrome as an approximation of the ancestral form of the Formicinae. The low shouldered condition has been independently derived in Tapinomini and Bothriomyrmecini (Dolichoderinae) and has been reversed in parallel among most but not all of the tribes in the major formicine radiation (Gigantiopini, Santschiellini, Gesomyrmecini, Oecophyllini, Formicini. Myrmoteratini, Camponotini). Note that the topology of this radiation is still uncertain (Blaimer *et al*. 2015).

**411.** Prora absent (CB50: 0.98). ***Polarity*:** Loss of the prora is convergent with respect to *Opamyrma*, *Apomyrma*, Dolichoderomorpha, and *Crematogaster*, among sampled taxa.

**418.** Cinctus of abdominal segment IV (“gastral II”) absent (CB50: 0.98). ***Polarity*:** Loss of the cinctus dividing pre- and postsclerites of abdominal segment IV is supported as convergent with respect to *Apomyrma*, *Simopelta*, and Dolichoderomorpha among sampled taxa.

**434.** Sting reduced to nonfunctionality and terminal metasomal opening fringed by a ring of hairs (“coronula of acidopore”) (CB50: 0.98). ***Polarity*:** Reduction of the sting has been derived in parallel with the Dolichoderinae, with derivation of the coronula being the major external distinguishing feature of Formicinae.

**511.** Forewing crossvein 2rs-m absent (CB50: 0.69). ***Polarity*:** The support value reported here is for the crown Formicinae, as alates are not known for Mesozoic members of the subfamily. Crossvein 2rs-m is never observed among Formicinae. Notably, †Formiciinae retain 2rs-m, despite the autapomorphic derivation of their venational conformation. From a broader phylogenetic perspective, with the loss of 2rs-m, the proximal juncture of free M is often distal to 2r-rs, resulting in extension of Rs+M distad that crossvein, a condition suggested by Rasnitsyn (2021) to be unique to †Peleserphidae among Proctotrupomorpha and perhaps to Hymenoptera as a whole. Free M also diverges distad 2rs-m in some *Ravavy* (Dolichoderinae).

***Note*:** There is equivocal support for absence of the occipital carina (char. **21**) as a synapomorphy of the total Formicinae (CB50: 0.57), and slightly higher support for the crown Formicinae (CB50: 0.68). Among sampled crown formicines, the carina was only observed in *Gigantiops*, *Santschiella*, *Oecophylla*, *Anoplolepis*, *Myrmoteras*, and *Camponotus*. Given that the Myrmelachistini and Lasiini comprise the sistergroups of the first two splits in the subfamily, it is expected that occipital carina absence will be more-strongly supported if subjected to further analysis.

### F.B.10’. Clade Myrmicomorpha

***Clade comprising*:** • **Ectatomminae** Emery, 1895, *•* **Myrmicinae** Lepeletier de Saint-Fargeau, 1835.

#### Synapomorphies

**87.** Clypeus projecting anteriorly from cranium (CB50: 0.71). ***Polarity*:** Although support for clypeal production at the myrmicomorphan node is equivocal, the condition is strongly supported as ancestral for Ectatomminae (CB50: 0.97) and Myrmicinae (CB50: 0.99). Among sampled ants, convergent with respect to Dolichoderomorpha, many Formicinae, and Poneria.

**250.** Metapleural gland orifice dorsoventrally oriented and slit-shaped (curved or not), not overhung by dorsal carina (CB50: 0.93). ***Polarity*:** The gland orifice is maximally supported as having a different, although unspecified, conformation for the Formicomyrmines and Formicae nodes. The original characterization is based on Bolton’s (2003) observations, which he suggested may be a synapomorphy of Ectatomminae and Myrmicinae, although he had not classified them as a group together.

***Note*:** Bolton (2003, p. 45) pointed out that the helcial pretergite of *Rhytidoponera* (Ectatomminae) and various Myrmicinae is produced anteriorly as a median lobe (Hashimoto 1996), and may constitute a synapomorphy of the Myrmicomorpha, as here constituted. It was not possible to evaluate this condition in the present work.

### **F.B.U.** Subfamily Ectatomminae Emery, 1895

*Clade comprising*: • **Ectatommini** Emery, 1895, • **Heteroponerini** Bolton, 2003.

***Note*:** Taxon sampling of Ectatomminae was limited, and thus supported synapomorphies may be overturned or refined with evaluation of more terminals.

#### Synapomorphies

**86.** Clypeus with median longitudinal carina (CB50: 0.92). ***Polarity*:** Sporadically derived among sampled Formicinae, including *Gigantiops*, *Anoplolepis*, *Lepisiota*, *Formica*, and *Camponotus*.

**135.** Posterolateral portions of head produced as distinct lobes (CB50: 0.98). ***Polarity*:** Among sampled crown Formicidae, only observed in Ectatomminae.

**183.** Anteroventral pronotal corner with spiniform process (CB50: 0.98). ***Polarity*:** Among sampled crown Formicidae, convergently derived in *Amblyopone*.

**324.** Pretarsal claw bifurcate (CB50: 0.86). ***Polarity*:** Multidentate pretarsal claws have been derived in Ponerinae perhaps once with reversal, and in *Paraponera*.

**369.** Petiolar tergosternal fusion absent (CB50: 0.94). ***Polarity*:** This condition represents a reversal which is independently derived in Ponerinae.

**390.** Abdominal segment II with defined laterotergite (CB50: 0.91). ***Polarity*:** Although this is expected to be a Dollo-like character, regain of the laterotergite is strongly supported for the Ectatomminae; the laterotergite is observed in all sampled Myrmeciomorpha. Absence of the laterotergite in Leptanillomorpha probably has a strong influence on the reconstruction of this feature.

**399.** Abdominal segment III (“postpetiole”) with tergosternal fusion (CB50: 0.96). ***Polarity*:** Formicae are strongly supported as lacking tergosternal fusion (CB50: 0.97), which is estimated to be regained in *Tapinoma*, *Prenolepis*, Plagiolepidini, and Ectatomminae.

**419.** Abdominal segment IV tubulated, *i.e.*, lateral margins of tergum and sternum meeting and aligned for their length (CB50: 0.94). ***Polarity*:** Unique among Formicae, and present in most Poneria with the exception of *Apomyrma* and *Adetomyrma* (Amblyoponinae, unsampled).

**420.** Abdominal poststernite IV shorter than posttergite IV (CB50: 0.94). ***Polarity*:** Unique among sampled Formicae. Also observed in *Opamyrma* and most Poneria (except *Apomyrma* and *Feroponera*, among sampled taxa).

**488.** Forewing Mf1 distinctly curved (CB50: 0.82). ***Polarity*:** Among sampled crown Formicidae, also observed in various Poneria (*Fulakora*, *Stigmatomma*, *Paraponera*, *Diacamma*, *Simopelta*, *Odontomachus*) and in some Formicae (*Myrmecia*, *Tetraponera*, *Gesomyrmex*, *Formica*, *Manica*).

**521.** Forewing 1rrs cell (“submarginal cell 1”) prestigmal length less than half total cell length (CB50: 0.93). ***Polarity*:** Variable among Formicidae; among Formicae, also observed in *Myrmecia*, *Pseudomyrmex*, *Formica*, *Camponotus*, *Manica*, *Stegomyrmex*, and *Crematogaster*.

**551.** Forewing crossvein 1cu-a prefurcal but separated proximad from Mf1 by less than its own length (CB50: 0.94). ***Polarity*:** Among Formicae, this somewhat prefurcal condition was also observed for *Myrmecia* and *Linepithema*; all other sampled terminals have a strongly prefurcal condition (1cu-a separated by more than one of its lengths proximad Mf1). The somewhat prefurcal condition is strongly supported as an ancestral condition of crown Formicidae (CB50: 0.96), and maximally so for the Poneroformicines, Poneria, Doryloformicia, and Formicae.

**568.** Hindwing Mf2 tubular (CB50: 0.86). ***Polarity*:** This condition is strongly supported as an expansion of the tubular state for Mf2 among Formicae (CB50 for non-tubular Mf2 for Formicae: 0.92; for Myrmechoderines: 0.94; for Formicomyrmines: 0.96).

***Note*:** See Formicomyrmines above for discussion of alate pronotal development (char. **177**).

### **F.B.V.** Subfamily Myrmicinae Lepeletier de Saint-Fargeau, 1835

***Clade comprising*:** • **Myrmicini** Lepeletier de Saint-Fargeau, 1835, • **Pogonomyrmecini** Ward *et al*., 2015, **core Myrmicinae** clade (•• **Stenammini** Ashmead, 1905, **ACS clade** {••• **Attini** Smith, F., 1858, **Crematogastrini** Forel, 1893, **Solenopsidini** Forel, 1893}).

#### Synapomorphies

**39.** Compound eyes situated at about head midlength (CB50: 0.86). ***Polarity*:** Despite uncertainty of eye position for the Formicae, eyes situated at head midlength is strongly supported as a condition of the ancestral myrmicine. Within Myrmicinae, eyes situated posterior to head midlength were observed in *Stegomyrmex* and *Crematogaster*.

**149.** Radicle and scape offset at a distinct angle (CB50: 0.89). ***Polarity*:** Also observed in the ectatommine *Acanthoponera*, and the poneriines *Paraponera*, *Tatuidris*, and *Platythyrea*.

**165.** Promesonotal articulation immobile due to fusion between pronotum and mesonotum (CB50: 0.96). ***Polarity*:** Among Doryloformicia, also observed in *Lioponera*, *Rhytidoponera* and *Lepisiota*; among Poneria, observed for *Paraponera*, *Tatuidris*, and Proceratiinae.

**216.** Alate mesoscutum with anteromedian line (CB50: 0.92). ***Polarity*:** Among sampled Formicidae, also observed for *Dolichoderus*, and the major formicine radiation (*i.e.*, all tribes with the exclusion of Melophorini, Lasiini, Myrmelachistini); observed to be absent in *Solenopsis*.

**348.** Propodeum armed (CB50: 0.95). ***Polarity*:** Among sampled Myrmicinae, lost in *Solenopsis*. Also observed in *Acanthoponera*, *Lepisiota*, *Santschiella*, *Dolichoderus*, *Aneuretus*, Proceratiinae, and various Ponerinae.

**\*.** Posterior foramen of petiole with tergosternal fusion and in the form of a perfect circle in posterior view (char. **395**, CB50: 0.99) and postpetiole with pretergites (helcium) fused and forming a perfect circle in anterior view (char. **396**, CB50: 0.99), consequently helcial sternite exposed in lateral view (char. **402**, CB50: 0.96). ***Polarity*:** The first of these two correlated characters were recognized as an autapomorphy of Myrmicinae by Bolton (2003) and were used to remove the poneriines *Tatuidris* and *Ankylomyrma* from the subfamily.

**405.** Abdominal segment III strongly petiolated, *i.e.*, considerably reduced in size and with distinct posterior face (CB50: 0.99). ***Polarity*:** Formation of a postpetiole is supported as independently derived in the ancestor of the Myrmeciomorpha, and is also observed in *Martialis*, Leptanillini, *Paraponera*, and *Tatuidris*.

**421.** Abdominal tergum IV enlarged, being larger than the petiole, postpetiole, and abdominal segments V+ (CB50: 0.99). ***Polarity*:** Independently derived in Leptanillini, *Paraponera*, and *Tatuidris*.

## SECTION 2: Transformation series from Apocrita into Aculeata

In this section, we provide an extended transformation series for non-focal clades included in our ancestral state analyses, formatted as for the main taxa. As for the main transformation series, we recommend that these results be considered working hypotheses to be tested with expanded taxon sampling and additional anatomical data. †Ephialtitoidea Handlirsch, 1906 is not treated as sampling of Unicalcarida and non-aculeate Apocrita was too limited to meaningfully evaluate character polarities. Please refer to the raw morphological data matrix for character distributions.

### **X.** Superfamily Evanioidea Latreille, 1802

***Clade comprising*: • Evanioidea crown** clade (Aulaciformes Grimaldi & Engel, 2005, Evaniiformes Grimaldi & Engel, 2005). • **Stem Evanioidea** or ***incertae sedis*** in superfamily: †**Anomopterellidae** Rasnitsyn, 1975, †**Nevaniinae** Zhang & Rasnitsyn, 2007, †**Praeaulacidae** Rasnitsyn, 1972 (*note: non-monophyletic; see extended phylogenetic results above*; †Praeaulacinae Rasnitsyn, 1972, †Cretocleistogastrinae Rasnitsyn, 1990), †**Vectevaniidae** Li *et al*., 2018 (†*Vectevania* Cockerell, 1922 [*not included in analyses*]).

***Note 1*:** Because the ancestral state reconstruction analyses employed the 50CB matrix, those non-bethylonymid fossils with < 50 % completeness were excluded, including †Anomopterellidae, most †Baissidae, and all †Praeaulacidae with the exception of †Nevaniinae. For this reason, the synapomorphies listed below apply to the crown clade of the Evanioidea, tempered by variable placement of †Nevaniinae. As only one terminal of extant Aulacidae and Gasteruptiidae were included, family-level synapomorphies are not provided. Synapomorphies are listed, however, for the Aulacidae + Gasteruptiidae total clade, the †Othniodellithidae + Evaniidae clade, the total Evaniidae, †*Cretevania*, crown Evaniidae + †*Grimaldivania* + †*Iberoevania* + †*Protoparevania*, and crown Evaniidae.

***Note 2*:** †*Rasnitsevania* Jouault *et al*., 2020 from burmite was attributed to the †Nevaniinae, but the description appeared too late to include in the present study.

#### Crown Evanioidea synapomorphies

**279.** Protrochanter elongate, length > 3 x width (CB50: 0.92). ***Polarity*:** Estimate influenced by extant taxa, stem Aulacidae, stem Gasteruptiidae, and †Othniodellithidae. Other evanioid fossils with protrochanter length uncertain. *Gasteruption* with a shorter protrochanter.

**293.** Mesotrochanter elongate, length > 3 x width (CB50: 0.96). ***Polarity*:** As for char. 279 above. *Gasteruption* with a shorter mesotrochanter.

**\*** Propodeometasomal articulation separated dorsad from metacoxae (char. **359**, CB50: 0.99) above propodeal midheight (char. **361**, CB50: 0.99), with the propodeum forming a completely fused sclerotic bridge ventrad propodeal foramen (char. **360**, CB50: 0.99). ***Polarity*:** †Nevaniinae and a number of †Praeaulacidae show intermediate conditions. Focused ancestral state reconstruction may recover a supported pathway (see Li *et al*. 2015 for a current transformational hypothesis).

**494.** Forewing Rsf between Rs+M and 2r-rs (= “2rs” of Engel 2017) linear to very weakly curved, without distinct kink or strong curve (CB50: 0.89). ***Polarity*:** Among sampled extant taxa, not applicable for *Hyptia*; distinct curvature or kinking was observed for †Nevaniinae, numerous †Praeaulacidae, some †Baissidae, and †*Hypselogastrion*.

### **X.2.1.** Clade Aulaciformes Grimaldi & Engel, 2005

***Clade comprising*:** • **Aulacidae** Shuckard, 1841 (Aulacinae Shuckard, 1841), • **Gasteruptiidae** Ashmead, 1900 (Hyptiogastrinae Crosskey, 1953, Gasteruptiinae Ashmead, 1900). • Paraphyletic with respect to Aulacidae, Gasteruptiidae: **†Baissidae** Rasnitsyn, 1975. **• *Incertae sedis*** to family: *•* **†Hyptiogastritinae** Engel, 2006, • †**Kotujellitinae** Rasnitsyn, 1975. • ***Incertae sedis*** to family and subfamily: **†Archeofoenini** Engel, 2017 (†*Archeofoenus* Engel, 2017, †*Protofoenus* Cockerell, 1917), • **†Electrofoenini** Cockerell, 1917 (†*Electrofoenia* Jouault *et al*., 2020, †*Electrofoenops* Engel, 2017, †*Electrofoenus* Cockerell, 1917, †*Exilaulacus* Li *et al*., 2018).

***Note 1*:** Synapomorphies are listed for the Aulaciformes, the recovered crown Aulacidae + Gasteruptiidae clade (“crown AG clade”, *i.e.*, excluding the stem grade family †Baissidae), and the “†ArElEx” clade (†*Archeofoenus*, †*Electrofoenops*, †*Exilaulacus*).

***Note 2*:** The archeofoenine genus †*Protofoenus* and the electrofoenine genera †*Electrofoenia* and †*Electrofoenus* were not included in the present analyses.

#### Aulaciformes synapomorphies

**\*** Mandible < 3 dentate (char. **65**, CB50: 0.86), possibly unidentate (char. **63**, CB50: 0.69). ***Polarity*:** Sampled extant AG clade species 3-dentate, as well as †*Electrofoenops*; other †ArElEx clade members < 3-dentate, as well as †*Hypselogastrion*, †*Kotujellites*, and †*Kotujisca*.

**201.** Propleurae forming tube-like collar supporting head (CB50: 0.87). ***Polarity*:** Stabilizes within the Aulaciformes. Also observed in †Othniodellithidae and *Evania*. Could not be evaluated for the most fossil Evanioidea.

**209.** Median portion of mesoscutellum, between parapsidal lines, bulging dorsally (CB50: 0.97). ***Polarity*:** Unique among all sampled taxa. The median portion of the mesoscutum is not bulging in †*Hypselogastrion* and could not be evaluated in either †*Kotujellites* or †*Kotujisca*.

**220.** Alate female with notauli that meet posteromedially, forming Y-shaped groove (CB50: 0.79). ***Polarity*:** Support stabilizes within the Aulaciformes. The notauli do not form a Y-shaped groove in crown and stem Evaniidae, where this trait could be evaluated.

**388.** Abdominal segments II and III fused tergally, forming cone-to funnel-shaped “petiole” (CB50: 0.98). ***Polarity*:** Unique among sampled taxa.

**511.** Fore wing 2rs-m nebulous to absent (CB50: 0.81). ***Polarity*:** This reconstruction is more ambiguous than the support values indicate. Crossvein 2rs-m is tubular in *Pristaulacus*, †*Exilaulacus loculatus* Li *et al*., 2018, †*Hypselogastrion*, †*Kotujellites*, and †*Kotujisca*, suggesting either ancient retention in these taxa with parallel losses, or re-expression. Extended taxon sampling and application of Bayesian mixture models are suggested to resolve this question.

**517.** Fore wing crossvein 3rs-m nebulous to absent (CB50: 0.95). ***Polarity*:** Among Evanioidea, a tubular 3rs-m is only retained in †Othniodellithidae, †*Palaeosyncrasis*, and †*Mesevania*.

**524.** Fore wing “submarginal cell 2” length and width subequal (CB50: 0.84). ***Polarity*:** Among sampled Evanioidea, also observed in †*Grimaldivania* and †*Iberoevania*; cell with differing proportions in †*Archaeofoenus engeli*, †*Electrofoenops rasnitsyni*, †*Hypselogastrion*, †*Kotujellites*, and †*Kotujisca*.

**532.** Fore wing crossvein 1m-cu antefurcal, *i.e.*, joining Rs+M proximal to the distal split of Rsf2 and Mf2 (CB50: 0.91). ***Polarity*:** Among sampled Evanioidea, also observed in various †*Cretevania*, and †*Grimaldivania*.

**534.** Fore wing first medial cell (“discal cell 1”) small to very small and rectangular, with proximodistal length ∼ 3 x anteroposterior width (CB50: 0.97). ***Polarity*:** Very consistent among sampled Evanioidea with the exception of *Pristaulacus*, which has a first medial cell with a different conformation.

**542.** When defined, fore wing medial cell 2 (“discal cell 2”) proportionally small, with surface area less than that of “submarginal cells 2, 3” (CB50: 0.72). ***Polarity*:** Support increases among internal nodes, but reconstruction is difficult due to inapplicability for several terminals. The ancestral condition estimated for Evanioidea is to have a proportionally large second medial cell.

**544.** Fore wing medial cell 2 proximodistal length ≥ 2 x anteroposterior length (CB50: 0.96). ***Polarity*:** Support diminished due to inapplicability among many terminals. However, consistent among taxa of the Aulaciformes, albeit with differing proportions observed in *Pristaulacus*.

**552.** Fore wing second cubital cell (“subdiscal 1”) anteroposteriorly wider than proximodistally long (CB50: 0.94). ***Polarity*:** *Pristaulacus*, *Gasteruption*, and †*Archeofoenus* with different proportions.

* Hind wing venation reduced: Rsf spectral or absent (char. **563**, CB50: 0.95); crossvein 1-m nebulous to absent (char. **565**, CB50: 0.95); Mf2 nebulous to absent (char. **568**, CB50: 0.92); Cuf nebulous to absent (char. **569**, CB50: 0.97); cu-a absent (char. **570**, CB50: 0.96). ***Polarity*:** Paralleled in Evaniidae, for which venational reduction appears to have occurred in multiple lineages (see raw data matrix for character distributions).

#### †ArElEx clade synapomorphies

**163.** Mesosoma compact, being dorsoventrally tall, anteroposteriorly short, and heavily sclerotized (CB50: 0.99). ***Polarity*:** The mesosoma is not compact in †*Hyptiogastrites electrinus* Cockerell, 1917, and could not be evaluated for †*Electrofoenops rasnitsyni* Turrisi & Ellenberger, 2019. Also observed in total Evaniidae.

**362.** Propodeal foramen situated at the apex of a conical projection (CB50: 0.99). ***Polarity*:** Also observed in *Pristaulacus*. Metasomal articulation is not situated on a cone in †*Hyptiogastrites*, nor the putative gasteruptiids †*Hypselogastrion*, †*Kotujellites*, and †*Kotujisca*. The articulation in †*Cretevania rubusensis* Peñalver *et al*., 2020 approaches this condition, and was scored as TRUE.

**523.** Fore wing 1rrs cell (“submarginal 1”) as long or longer than 2rsm, 3rsm (“submarginal 2, 3”) (CB50: 0.95). ***Polarity*:** Among sampled Evanioidea, also observed in *Pristaulacus*, †*Palaeosyncrasis*, †*Kotujellites*, †*Kotujisca*, and †*Cretevania*.

**558.** Fore wing Cu2 (posterior branch of free Cu) not reaching 1A (CB50: 0.97). ***Polarity*:** Among sampled Evanioidea, also observed in *Gasteruption*, *Hyptia*, and †*Sorellevania*.

#### Crown AG clade synapomorphies

**206.** Mesoscutum longer than broad in dorsal view (CB50: 0.77). ***Polarity*:** The mesoscutum is shorter in †*Electrofoenops*; otherwise, an elongate pronotum is unique among Evanioidea.

**365.** Metasomal articulation contacting metanotum (CB50: 0.85). ***Polarity*:** The articulation is distant in *Pristaulacus* for which the articulation is situated on a cone-like projection. The articulation contacts the metanotum in †*Hypselogastrion* and †*Kotujellites* and is uncertain in †*Kotujisca*. Almost completely unique except for the praeaulacines †*Sinaulacogastrinus* and †*Sinevania*.

**366.** Metasoma long, thin, and tubular, with segments only gradually increasing in width posteriorly (CB50: 0.85). ***Polarity*:** As for 365, except that the praeaulacines do not display this condition.

### **X.2.2.** Evaniiformes Grimaldi & Engel, 2005

*Clade comprising*: • †**Othniodellithidae** Engel & Huang, 2016b, • **Evaniidae** Latreille, 1802.

#### Synapomorphies

**138.** Antennal toruli situated at or posterior to head midlength (CB50: 0.98). ***Polarity*:** Moderately supported as a plesiomorphy of the Evanioidea (CB50: 0.82), but uncertain or absent for all sampled terminals which are outside of the Evaniiformes. This reconstruction should be considered provisional until a greater sample of Apocrita can be evaluated and analyzed.

**368.** Metasoma after abdominal segment II (metasomal I) very short and lateromedially compressed, forming oval or teardrop shape in lateral view (CB50: 0.96). ***Polarity*:** Observed in †*Palaeosyncrasis*. Condition uncertain in †*Xenodellitha*. Otherwise, unique among all sampled taxa.

**369.** Abdominal segment II (metasomal I) with tergosternal fusion (CB50: 0.86). ***Polarity*:** As for char. 368, but much more difficult to evaluate for fossils, and not as unique due to variation among Aculeata.

**433.** Ovipositor longer than metasoma (CB50: 0.87). ***Polarity*:** Although other Evanioidea have elongate ovipositors, this condition is attributable in part to the compact form of the “gaster” of †Othniodellithidae and Evaniidae. However, the ovipositor is markedly short in crown Evaniidae.

**453.** Fore wing Rsf clearly not reaching distalmost wing margin (CB50: 0.58). ***Polarity*:** Support for this condition is maximal for †Othniodellithidae and is strong for total Evaniidae (CB50: 0.92). Rsf does reach the distalmost margin in †*Palaeosyncrasis* and †*Mesevania*.

**538.** Fore wing crossvein 2m-cu nebulous to absent (CB50: *). ***Polarity*:** Support for this condition is equivocal at the †Othniodellithidae–Evaniidae node but is maximally supported for †Othniodellithidae and strongly so for the total Evaniidae (CB50: 0.98). The equivocal reconstruction is probably due to †*Palaeosyncrasis*, which has a well-defined second sectoriomedial crossvein. 2rs-m is also lost in *Gasteruption*, †*Archeofoenus*, †*Hyptiogastrites*, †*Electrofoenops diminuta*, and †*Kotujisca*.

### X.D. †Othniodellithidae Engel & Huang, 2016

**Clade comprising: •** †*Othniodellitha* Engel & Huang, 2016, • †*Xenodellitha* Engel, 2017.

#### Synapomorphies

**19.** Cranium elongated posterad compound eyes (CB50: 0.99). ***Polarity*:** Among sampled Evanioidea, also observed in extant Aulacidae and Gasteruptiidae, and †*Archeofoenus engeli* Turrisi & Ellenberger, 2019.

**30.** Malar space (area between mandibular base and compound eye) virtually absent (CB50: 0.99). ***Polarity*:** Uncertain for most fossil terminals. Also observed in †*Palaeosyncrasis* and †*Cretevania tenuis* Li *et al*., 2018.

**48.** Mandibular masticatory margin elongate (CB50: 0.99). ***Polarity*:** Almost unique among Evanioidea. Among sampled taxa, also occurring in and defining a clade of stem Aulacidae (†*Exilaulacus*, †*Archeofoenus*, and †*Electrofoenops*).

**66.** Mandible at least 4-dentate (CB50: 0.99). ***Polarity*:** Unique among sampled Evanioidea for which tooth count could be evaluated.

**86.** Clypeal disc with median longitudinal carina (CB50: 1.0). ***Polarity*:** Unique among sampled Evanioidea for which the clypeus could be evaluated.

**93.** Anterior clypeal margin linear (CB50: 1.0). ***Polarity*:** Also observed in *Gasteruption*. Otherwise, unique among those Evanioidea for which the clypeus could be evaluated. Other Evanioidea reconstructed as having a convex anterior clypeal margin.

**111.** Face produced anteriorly, forming massive process which bears the antennal toruli (CB50: 1.0). ***Polarity*:** Unique among all sampled taxa.

**140.** Antennal toruli separated from clypeus by more than one toruli diameter (CB50: 1.0). ***Polarity*:** Among Evanioidea for which this condition could be evaluated, also observed for *Gasteruption*.

**142.** Antennal toruli directed more-or-less laterally (CB50: 1.0). ***Polarity*:** Unique among all Evanioidea for which this condition could be evaluated.

**\*.** Scape not subglobular, not bulbous or swollen medially (char. **152**, CB50: *), being rather somewhat elongate, with a length > 2 x but < 4 x width (char. **153**, CB50: *). ***Polarity*:** Strictly reporting at the †Othniodellithidae + Evaniidae node, these two characters are equivocally reconstructed. However, they are maximally supported as the condition for †Othniodellithidae and strongly so for the total Evaniidae (CB50: 0.99). The scape was observed to be bulbous in *Evania* and †*Mesevania*, and ≥ 4 x width in †*Botsvania*, most †*Cretevania*, and †*Iberoevania*. The scape is < 2 x width in †*Mesevania* and †*Newjersevania nascimbenei* Basibuyuk *et al*., 2000.

**187.** Pronotum laterally scrobiculate, *i.e.*, with a broad furrow capable of receiving the fore leg when it is retracted dorsally and medially (CB50: 1.0). ***Polarity*:** Could not be evaluated for other fossil Evanioidea. Crown Evaniidae without such a scrobe.

**451.** Fore wing Rf distinctly not reaching wing apex (CB50: 1.0). ***Polarity*:** Highly variable among sampled Hymenoptera. However, strongly supported as ancestral for the Evanioidea and the †Othniodellithidae-Evaniidae clade (CB50 for both: 0.96). Convergent with respect to crown Evaniidae (CB50: 0.96).

**498.** Fore wing Mf3 shorter than Rs+M (CB50: 1.0). ***Polarity*:** Among Evanioidea, a short third Medial abscissa is also observed in †*Palaeosyncrasis*, †*Exilaulacus latus* Li *et al*., 2018, †*Kotujisca*, †*Grimaldivania*, and †*Mesevania*.

**514.** Fore wing crossvein 2rs-m joining Rs at or very near to 2r-rs resulting in anterior “petiolation” of cell 2rm (CB50: 0.99). ***Polarity*:** Among sampled Evanioidea for which cell 2rm is enclosed, also observed in †*Cretevania*.

**525.** Fore wing “submarginal cell 2” quadrangular to rectangular, proximodistal length ≥ 3 x anteroposterior width (CB50: 0.99). ***Polarity*:** Among sampled Evanioidea, also observed in †*Cretevania*.

**540.** Fore wing cross vein 2m-cu joining M at or proximal to 2rs-m crossvein (CB50: 0.82). ***Polarity*:** Not applicable for †*Othniodellitha*.

**\*.** When defined, fore wing medial cell 2 (“discal cell 2”) proportionally small, with surface area less than that of “submarginal cells 2, 3” (char. **542**, CB50: 0.83), with cell being proximodistally longer than anteroposteriorly broad (char. **543**, CB50: 0.83). ***Polarity*:** Not applicable for †*Othniodellitha*.

### X.E. Evaniidae Latreille, 1802 total clade

***Clade comprising*: •** †***Cretevania*** Rasnitsyn, 1975, • **EGIP clade** (•• †*Grimaldivania* Basibuyuk *et al*., 2020, †*Iberoevania* Peñalver *et al*., 2010, †*Protoparevania* Deans, 2004, crown Evaniidae). • ***Incertae sedis*** in family: †*Burmaevania* Shih *et al*. (2020), †*Lebanevania* Basibuyuk & Rasnitsyn, 2002, †*Mesevania* Basibuyuk & Rasnitsyn, 2002, †*Newjersevania* Basibuyuk *et al*., 2002, †*Palaeosyncrasis* Poinar, 2019, †*Sinuevania* Li *et al*., 2018, †*Sorellevania* Engel, 2016.

***Note*:** Listed synapomorphies are divided into those applying to the total clade Evaniidae, †*Cretevania*, the “EGIP” clade (crown Evaniidae + †*Grimaldivania* + †*Iberoevania* + †*Protoparevania*), and the crown Evaniidae itself. For the total clade Evaniidae, evaluations were made including and excluding †*Palaeosyncrasis* (itself probably a junior synonym of †*Mesevania*); these are noted in the polarity comments.

#### Total clade Evaniidae synapomorphies

**17.** Head ventrally elongate and strongly flattened posteriorly, fitting against the vertically truncate mesosoma (CB50: 0.99). ***Polarity*:** Unique among all sampled taxa and shared among all sampled extant species. Uncertain for several fossils, including †*Palaeosyncrasis*.

**114.** Face with medially situated and longitudinal frontal carina (CB50: 0.75). ***Polarity*:** Among crown Evaniidae, the medial frontal carina was observed to be absent for *Hyptia*; also observed for †*Lebanevania*; otherwise, absent for all Evanioidea for which the face could be evaluated. Support becomes near maximal at the crown evaniid node (CB50: 0.98).

**144.** Antennal toruli directed dorsally (assuming hypognathy) (CB50: 0.85). ***Polarity*:** Unique among those Evanioidea for which this condition could be evaluated, including †*Palaeosyncrasis*.

**163.** Mesosoma very compact, being dorsoventrally tall, anteroposteriorly short, and heavily sclerotized (CB50: 0.95). ***Polarity*:** The mesosoma is compact in †*Palaeosyncrasis*, and support becomes maximal for the remainder of the total Evaniidae. Also observed for the †ArElEx clade.

**483.** Fore wing Rsf1 and Mf1 meeting at a distinct angle, with the vertex directed distally (CB50: 0.56). ***Polarity*:** Although equivocal at the total Evaniidae node, strongly supported for †*Cretevania*, the EGIP clade, and crown Evaniidae (CB50: ≥ 0.96). The first free abscissae of Rs and M are more linear in †*Xenodellitha*, †*Lebanevania*, †*Mesevania*, and †*Palaeosyncrasis*.

**511.** Fore wing crossvein 2rs-m nebulous to absent (CB50: 0.94). ***Polarity*:** Crossvein 2rs-m is tubular in †*Palaeosyncrasis*, †*Mesevania*, and †*Cretevania minor* **517.** Fore wing crossvein 3rs-m nebulous to absent (CB50: 0.96). ***Polarity*:** Among Evanioidea, a tubular 3rs-m is only retained in †Othniodellithidae, †*Palaeosyncrasis*, and †*Mesevania*.

#### †Cretevania synapomorphies

**154.** Scape relatively elongate, with its length ≥ 4 x width (CB50: 0.82). ***Polarity*:** The scape length is < 4 x width in some †*Cretevania*.

**514.** Fore wing crossvein 2rs-m joining Rs at or very near to 2r-rs resulting in anterior “petiolation” of cell 2rm (CB50: 0.95). ***Polarity*:** Among sampled Evanioidea for which cell 2rm is enclosed, also observed in †Othniodellithidae.

**521.** Fore wing 1rrs cell (“submarginal 1”) prestigmal length greater than half total cell length (CB50: 0.89). ***Polarity*:** Among sampled Evanioidea, also observed in Aulacidae, Gasteruptiidae, †*Exilaulacus latus*, †*Archeofoenus engeli*, †*Kotujellites*, †*Kotujisca*, †*Lebanevania*, and †*Mesevania*. First radial cell shorter in †*Cretevania alcalai*, †*C. montoyai*, and †*C. vesca*.

**522.** Fore wing 1rrs cell extremely long and narrow, with anteroposterior length being only slightly wider than that of costal cell, and with proximodistal length ≥ 3 x anteroposterior width (CB50: 0.99). ***Polarity*:** Also observed in crown Evaniidae.

**523.** Fore wing 1rrs cell (“submarginal 1”) as long or longer than 2rsm, 3rsm (“submarginal 2, 3”) (CB50: 0.94). ***Polarity*:** Among sampled Evanioidea, also observed in the †ArElEx clade, *Pristaulacus*, †*Palaeosyncrasis*, †*Kotujellites*, †*Kotujisca*, and †*Cretevania*.

**525.** Fore wing “submarginal cell 2” quadrangular to rectangular, proximodistal length ≥ 3 x anteroposterior width (CB50: 0.99). ***Polarity*:** Among sampled Evanioidea, also observed in †Othniodellithidae.

#### EGIP clade synapomorphies

**372.** Abdominal segment II in form of perfectly straight, narrow tube for its entire length (CB50: 0.98). ***Polarity*:** Also observed in †*Palaeosyncrasis*, and the praeaulacines †*Evanigaster*, †*Evaniops*. The petiole of †*Cretevania* is clubbed posteriorly.

**488.** Fore wing Mf1 distinctly curved (CB50: 0.89). ***Polarity*:** Strongly supported for crown Evaniidae (CB50: 0.98) and observed sporadically among stem taxa. Not applicable for *Hyptia*.

**501.** Fore wing Mf2+ or Mf3+ nebulous to absent (CB50: 0.95). ***Polarity*:** Among sampled Evanioidea, also observed in †*Curtevania*, †*Newjersevania*, and †*Sinuevania*.

#### Crown clade Evaniidae synapomorphies

**31.** Malar space longer dorsoventrally than broad anteroposteriorly (assuming hypognathy) (CB50: 0.96). ***Polarity*:** Also observed in crown Aulacidae, †*Electrobaissa*, †*Hypselogastrion*, †*Kotujellites*, †*Kotujisca*, and †*Cretevania minor* Rasnitsyn, 1975.

**85.** Median portion of clypeus, between anterior tentorial pits, dorsoventrally longer than lateromedially wide (assuming hypognathy) (CB50: 0.97). ***Polarity*:** Could not be evaluated for most fossil Evanioidea, including all stem Evaniidae, so this synapomorphy should be considered provisional at best.

**94.** Anterior clypeal margin produced as a distinct median lobe (CB50: 0.87). ***Polarity*:** Among sampled crown Evaniidae, the clypeus was not medially produced in *Evaniella*. Otherwise, unique among those Evanioidea for which the clypeus could be evaluated.

**115.** Longitudinal carinae present laterad the antennal toruli (“lateral frontal carinae”) (CB50: 1.0). ***Polarity*:** Present in all sampled crown Evaniidae, and absent in all Evanioidea for which the lateral region of the face could be critically evaluated.

**243.** Alate with upper metapleural area (that area dorsad the endophragmal pit) reduced, being present as a small, longitudinally aligned region, much smaller than the proportionally massive lower metapleural area (CB50: 0.89). ***Polarity*:** The upper metapleural area does not have this conformation in *Evaniella*. It was not possible to evaluate this condition for stem Evaniidae. For this reason, support at higher nodes is weak.

**256.** Mesocoxae set in ventral thoracic sockets, with sternal area nearly reaching ventralmost surface of coxae (CB50: 0.83). ***Polarity*:** The moderate support is misleading, as this condition was observed in only three of the five sampled crown Evaniidae and could not be evaluated for stem Evaniidae. The mesocoxae were observed to be not set in sockets in stem AG clade members.

**364.** Dorsal surface of propodeum between the metanotum and the dorsally migrated metasomal articulation convex (CB50: 0.99). ***Polarity*:** Also observed in †*Sinuevania*, otherwise unique among Evanioidea.

**433.** Ovipositor short, not longer than metasoma (CB50: 1.0). ***Polarity*:** The ovipositor is also relatively short in some species of the †ArElEx clade, †*Botsvania*, †*Mesevania*, †*Protoparevania*, †*Sinuevania*, and †*Sorellevania*.

**451.** Fore wing Rf distinctly not reaching wing apex (CB50: 1.0). ***Polarity*:** See polarity comments for †Othniodellithidae above.

**485.** Fore wing Mf1 long to extremely long, Mf1 length > 4 x Rsf1 length (CB50: 0.95). ***Polarity*:** Also observed in †*Newjersevania*, various Chrysididae, *Rhopalosoma*, Vespidae, *Methocha*, various Apoidea, and *Stigmatomma*.

**489.** Fore wing Mf1 strongly curved and closely approximating Sc+R+Rs such that first radial cell is long and very narrow distally (CB50: 0.99). ***Polarity*:** Unique among sampled taxa.

**516.** Fore wing crossvein 2rs-m joining Rs distal to 2r-rs by greater than one of its lengths (CB50: 0.99). ***Polarity*:** Among sampled Evanioidea, also observed in Aulacidae, Gasteruptiidae, †*Kotujellites*, †*Mesevania*, †*Newjersevania*, and †*Protoparevania*. Analysis including both of the latter taxa may meaningfully inform the age and composition of the crown Evaniidae.

**522.** Fore wing 1rrs cell extremely long and narrow, with anteroposterior length being only slightly wider than that of costal cell, and with proximodistal length ≥ 3 x anteroposterior width (CB50: 0.99). ***Polarity*:** Also observed in †*Cretevania*.

***Note 1*:** There is a distinct trend among total Evaniidae toward having fore wing crossvein 1cu-a becoming distinctly prefurcal (char. **548**) but there is variability which indicates parallelism.

***Note 2*:** The hind wing venation of Evaniidae displays a similar trend toward reduction as observed for the Aulacidae + Gasteruptiidae clade; see that node for further details.

### X’. Clade Trigaculeata

***Clade comprising*: • Trigonaloidea** Cresson, 1887, • †**Panguoidea** Li *et al*., 2020 (••†**Panguidae** Li *et al*., 2020), • **Aculeata** Latreille, 1802.

***Note 1*:** No unequivocal synapomorphies for the Trigaculeata as sampled here were detected, largely due to the uncertainty at deeper nodes. We enthusiastically anticipate phenome-scale anatomical data for the Apocrita.

***Note 2*:** †Panguoidea was defined after the results of the present phylogenetic analyses were written in detail. The single species attributed to this superfamily, †*P. yuangu* Li *et al*., 2020, agrees with the overall morphology of the Trigaculeata, having a median pronotal lobe (“neck”), trigonaloid-like wing venation, a very short, aculeate-like ovipositor, and an aculeate-proportioned scapes. Contradicting the estimated synapomorphy of the Aculeata, the occipital carina appears to extend to the hypostoma. Similar to †Falsiformicidae *sensu stricto*, the first metasomal segment is petiolate, but otherwise the *P. yuangu* fails the diagnosis of the family with the exception of having an axial helcium. Furthermore, the pronotum of †*P. yuangu* is very short and weak, unlike †Bethylonymoidea, Chrysidoidea, and Dryinoidea, suggesting that the elongate pronotum is a synapomorphy of the Aculeata. Overall, we agree with Li *et al*. (2020) and consider the superfamily close to the Aculeata and Trigonaloidea. Due to the derived condition of having a short ovipositor, we hypothesize that †Panguoidea are sister to the Aculeata, including †Bethylonymoidea; this should be tested in future study.

### **Y.** Superfamily Trigonaloidea Cresson, 1887

***Clade comprising*: • †Maimetshidae** Rasnitsyn, 1975, • **Trigonalidae** Cresson, 1887. • ***Incertae sedis*** in superfamily: †**Andreneliidae** Rasnitsyn & Martinez-Delclòs, 2000 **superfam. trans.**, †*Albiogonalys* Nel *et al*., 2003, †*Archaulacus* Li *et al*., 2014.

***Note*:** Synapomorphies for the Trigonaloidea are provided for two estimated nodes: †*Albiogonalys* + †Maimetshidae + Trigonalidae (“total clade Trigonaloidea”), and †Maimetshidae + Trigonalidae (“MT” clade). Because quite a few features were difficult to evaluate for fossils especially of the mesosoma, and as the extant taxa clustered together, there is limited support for certain reconstructions, compounded if they are variable in the crown clade. For this reason, we limit our reporting to those characters which could be evaluated with consistency; we direct the reader to the raw data matrix for character distributions.

#### Total clade Trigonaloidea synapomorphies

**48.** Mandibular masticatory margin elongate, thus mandible more-or-less triangular in form (CB50: 0.86). ***Polarity*:** Reversed in †*Iberomaimetsha rasnitsyni*, and uncertain for a number of trigonaloids which were not included in the ancestral state analyses.

**62.** Mandibular dentition asymmetrical, with one mandible having more teeth than the other (CB50: 0.97). ***Polarity*:** Occasionally observed in Formicidae, for example, but otherwise diagnostic of the Trigonaloidea.

**66.** Mandible at least 4-dentate (CB50: 0.87). ***Polarity*:** Apparently without reversals in the superfamily.

**329.** Tarsomeres I–IV with apicomedian ventral lobate setae (CB50: 0.88). ***Polarity*:** Although supported as a synapomorphy based on the present analyses, we suspect that these setae are a retained groundplan feature of the Apocrita, particularly as these structures correspond to the plantulae (*e.g.*, Beutel *et al*. 2020, references therein).

**523.** Fore wing cell 1rrs (“submarginal cell 1”) as long or longer than 2rsm, 3rsm (“submarginal 2, 3) (CB50: 0.82). ***Polarity*:** This is the condition of all sampled Trigonaloidea, although it could not be evaluated for †*Cretogonalys*.

#### MT clade synapomorphies

**19.** Cranium elongated posterad compound eyes (CB50: 0.73). ***Polarity*:** Stabilizes for the total Trigonalidae and †Maimetshidae, with reversal stabilizing in one of the maimetshid subclades.

**158.** Antennomere III longer than IV (CB50: 0.84). ***Polarity*:** Among sampled Trigonaloidea, the third antennomere is subequal or longer than the fourth in †*Iberomaimetsha pallida*, †*Maimetshasia*, †*Zorophratra*, and †*Albiogonalys*.

**279.** Protrochanter elongate, length > 3 x width (CB50: 0.97). ***Polarity*:** †*Albiogonalys* with a relatively shorter protrochanter.

**483.** Fore wing Rsf1 and Mf1 meeting at a distinct oblique angle (CB50: 0.93). ***Polarity*:** Consistent for most sampled trigonaloids, with the exception of †*Guyotemaimetsha*, †*Turgonaliscus*, and †*Albiogonalys*.

**488.** Fore wing Mf1 distinctly curved (CB50: 0.82). ***Polarity*:** The only exceptions among sampled Trigonaloidea are †*Iberomaimetsha pallida*, †*Maimetsha arctica*, †*Albiogonalys*, and *Bareogonalos*.

**491.** Fore wing M+Cu angling strongly from juncture with crossvein cu-a (CB50: 0.89). ***Polarity*:** Crown Trigonalidae are supported as having reversed to the less- or non-angled condition (CB50: 0.96). All sampled extinct Trigonaloidea, however, display this condition. It is possible that the angled conformation may be a synapomorphy of the extinct species to the exclusion of crown Trigonalidae, but further analyses are necessary to evaluate this alternative hypothesis.

***Note*:** Fore wing Rsf extending to the distalmost wing margin was supported as a synapomorphy of the crown Trigonalidae (CB50: 0.96), but we suspect this may be an artefact due to the small body size of most trigonaloid fossils.

### Y.B. Family Trigonalidae Cresson, 1887

***Clade comprising*: • Orthogonalinae** Carmean & Kimsey, 1998 (*Orthogonalys* Schultz, 1905), • **Trigonalinae** Cresson, 1887 (Nomadinini Cameron, 1899, Trigonalini Cresson, 1887). • ***Incertae sedis*** in family: †*Cretogonalys* Rasnitsyn, 1977.

***Note*:** Synapomorphies are provided for the total clade Trigonalidae as estimated here (*i.e.*, including †*Cretogonalys*) and for the crown clade.

#### Total clade Trigonalidae synapomorphies

**548.** Fore wing crossvein 1cu-a anterior juncture postfurcal, *i.e.*, distal to the split of M+Cu (CB50: 0.86). ***Polarity*:** Observed in †*Cretogonalys*, extant Trigonalidae, and †*Zorophrata*.

#### Crown clade Trigonalidae synapomorphies

**87.** Clypeal disc broadly projecting ventrally over mandibles (CB50: 0.93). ***Polarity*:** Difficult to evaluate for fossil taxa, but of those which could be seen well enough, the disc was not projecting.

**113.** Face with transverse process dorsad antennal toruli (CB50: 0.96). ***Polarity*:** †*Cretogonalys* without this process. It was not possible to evaluate this trait for †*Albiogonalys*. Otherwise, absent among other sampled Trigonaloidea.

**142.** Antennal toruli directed more-or-less laterally (CB50: 0.96). ***Polarity*:** Unique among sampled Trigonaloidea, including †*Cretogonalys* and †*Albiogonalys*.

**514.** Fore wing crossvein 2rs-m joining Rs at or very near to 2r-rs (CB50: 0.98). ***Polarity*:** Among sampled fossil Trigonaloidea, only occurring in †*Turgonalus*.

### Y.C. Family †Maimetshidae Rasnitsyn, 1975

***Clade comprising*: • †Maimetshinae** Rasnitsyn, 1975 (•• †**Ahiromaimetshini** Engel, 2016 [†*Ahiromaimetsha* Perrichot *et al*., 2011, †*Turgonaliscus* Engel, 2016, †*Turgonalus* Rasnitsyn, 1990], •• †**Maimetshini** Rasnitsyn, 1975 [†*Afrapia* Rasnitsyn & Brothers, 2009, †*Afromaimetsha* Rasnitsyn & Brothers, 2009, †*Ahstemiam* McKellar & Engel, 2011, †*Andyrossia* Rasnitsyn & Jarzembowski, 2000, †*Burmaimetsha* Perrichot, 2013, †*Guyotemaimetsha* Perrichot *et al*., 2004, †*Iberomaimetsha* Ortega-Blanco *et al*., 2011, †*Maimetsha* Rasnitsyn, 1975, †*Maimetshasia* Perrichot, 2013, †*Maimetshorapia* Rasnitsyn & Brothers, 2009]), • †**Zorophratrinae** Engel, 2016 (†*Zorophratra* Engel, 2016).

***Note*:** Most of the fossils attributed to the †Maimetshidae were not included in the ancestral state reconstruction analyses due to the fossils’ incompleteness. However, among those that were included, two main clades were supported in the ancestral state reconstruction topology searches, for which polarity estimates are provided: †*Ahstemiam*, †*Iberomaimetsha rasnitsyni*, and †*Maimetshasia* (“AIM clade”), and †*Burmaimetsha*, †*Iberomaimetsha nihtmara*, †*Guyotemaimetsha*, †*Maimetsha*, and †*Maimetshorapia* (“BIGMM clade”). †*Cretogonalys*, previously attributed to †Maimetshidae, was supported as outside the family.

#### †Maimetshidae synapomorphies

**494.** Fore wing Rsf between Rs+M and 2r-rs linear, neither kinked or distinctly curved (CB50: 0.88). ***Polarity*:** This is supported as a synapomorphy of the AIM and BIGMM clades to the exclusion of †*Ahiromaimetsha* (CB50 for inclusion of †*Ahiromaimetsha* = 0.75 for kinked or curved state). Within the AIM clade, †*Iberomaimetsha pallida* has a kinked or curved Rsf in this position, as does †*Cretogonalys* (supported as sister to crown Trigonalidae).

**513.** Fore wing 2rs-m joining Rs distinctly proximal to 2r-rs crossvein, resulting in anterior “petiolation” of cell 2rsm (“second submarginal”) (CB50: 0.92). ***Polarity*:** Observed in all †Maimetshidae except for †*Maimetsha*, †*Turgonalus*, and †*Zorophratra*; 2rsm is not petiolated in †*Albiogonalys*.

**520.** Fore wing crossvein 3rs-m situated close to 2rs-m, such that cell 3rm (“submarginal 3”) anteroposteriorly wider than proximodistally long (CB50: 0.95). ***Polarity*:** Apparently reverse in †*Ahstemiam*. Not applicable for a number of maimetshids and could not be evaluated for †*Cretogonalys*. †*Albiogonalys* with a longer cell.

**524.** Fore wing “submarginal cell 2” length and width subequal (CB50: 0.93). ***Polarity*:** The recovered sister to the †Maimetshidae in the ancestral state reconstruction topology search,

†*Ahiromaimetsha*, has a cell with unequal length and width. Additionally, this condition was not observed in †*Turgonaliscus*, †*Turgonalus*, †*Albiogonalys*, and was observed for *Orthogonalys*, indicating some degree of homoplasy.

**532.** Fore wing 1m-cu antefurcal, *i.e.*, joining Rs+M proximal to split of Rsf2 and Mf2 (CB50: 0.95). ***Polarity*:** Also observed in *Orthogonalys*, otherwise largely consistent among sampled Trigonaloidea with the exception of †*Ahiromaimetsha*, †*Turgonalus*, and †*Albiogonalys*.

**550.** Fore wing crossvein 1cu-a anterior junction prefurcal, *i.e.*, proximal to the splitting of M+Cu (CB50: 0.94). ***Polarity*:** Observed in all †Maimetshidae, with the exception of †*Zorophratra*. Crossvein cu-a is not prefurcal in †*Cretogonalys*, †*Albiogonalys*, and crown Trigonalidae.

**565.** Hind wing r-m nebulous to spectral (CB50: 0.51). ***Polarity*:** Although equivocal, r-m was only observed to be tubular in †*Ahiromaimetsha* and †*Iberomaimetsha*, among sampled †Maimetshidae. Crossvein r-m is also tubular in †*Cretogonalys*, †*Albiogonalys*, and crown Trigonalidae.

#### AIM clade synapomorphies

**19.** Cranium not extended posterior to compound eyes (CB50: 0.82). ***Polarity*:** This represents a reversal from the elongated condition of the Trigonaloidea, at least as supported by our analyses.

**555.** Fore wing Cu1 and Cu2 (CuA and CuP) diverging at about a right angle (CB50: 0.84). ***Polarity*:** Uncertain for several †Maimetshidae.

**558.** Fore wing Cu2 reaching 1A (CB50: 0.96). ***Polarity*:** Among sampled †Maimetshidae, also observed in †Maimetshorapia. See raw data for character distribution.

#### BIGMM clade synapomorphies

**496.** Second abscissa of Rs+M present and longer than first abscissa (CB50: 0.93). ***Polarity*:** Among sampled crown Trigonalidae, only applicable to *Orthogonalys*. This appears to be the condition which places †*Iberomaimetsha nihtmara* in the BIGMM clade. See raw data matrix for further details of character distribution.

***Notes*:** Compound eyes which converge medially toward the mandibles (char. **34**) was equivocal for the †Maimetshidae but was moderately supported as a synapomorphy of the AIM clade (CB50: 0.80) and equivocally so for the BIGMM clade (CB50: 0.59). In some †Maimetshidae, it appears that the cranium is supported to a greater extent by the propleurae (char. **201**), however this was difficult to evaluate for most taxa and was provisionally scored as true only for †*Ahiromaimetsha*, †*Guyotemaimetsha*, †*Maimetsha arctica*, and †*Maimetshasia*.

### Y’. Aculeata Latreille, 1802

See the main transformation series (*i.e.*, leading to Formicoidea) above for definition based on the present analyses.

### **Z.** #. Superfamily †Bethylonymoidea Rasnitsyn, 1975

***Clade comprising*:** • **†Bethylonymidae** (•• **†Bethylonyminae** Rasnitsyn, 1975 **stat. nov.** (••• †*Bethylonymus* Rasnitsyn, 1975, †*Bethylonymellus* Rasnitsyn, 1975); •• ***Incertae sedis*** in family: †*Meiagaster* Rasnitsyn & Ansorge, 2000, †*Renymus ovata* (Ren, 1995) **nom. nov.**) ***Note*:** Due to their incompleteness, the compression fossil taxa †*Meiagaster* and †*Renymus* were not included in the ancestral state estimation analysis. Therefore, the results below only apply to †*Bethylonymus* and †*Bethylonymellus*, the two of which probably render one another paraphyletic.

#### Synapomorphies

**11.** Cranium prognathous (CB50: 1.0). ***Polarity*:** Convergent with various Dryinoidea and some Vespaculeata (some Vespidae, Sierolomorphidae, Formicoidea, Ampulicidae).

**15.** Cranium narrower than long (CB50: 0.95). ***Polarity*:** Convergent with some Chrysidoidea, Embolemidae, †Falsiformicidae, and Formicidae.

**136.** Antennal toruli extremely anteriorly situated, such that they are at or overhanging the anterior clypeal margin (CB50: 0.99). ***Polarity*:** Convergent with various Chrysidoidea, Formicoidea.

**453.** Forewing Rsf extending to or nearly to distalmost wing margin (CB50: 0.63). ***Polarity*:** Support increases at the first internal node of the family (CB50: 0.79). Variable among Aculeata.

**482.** Forewing Rsf1 anterior juncture with Rf contacting or nearly contacting pterostigma (CB50: 0.97). ***Polarity*:** Variable among Aculeata.

### Z’.#. Family †Falsiformicidae Rasnitsyn, 1975

***Clade comprising*: • Strictly:** †*Falsiformica* Rasnitsyn, 1975, †*Siccibythus* Cockx & McKellar, 2016. • **Uncertain, associated forms:** †*Burmasphex* Melo & Rosa, 2018, two undescribed forms (CASENT0844587 Fig. S1, CASENT0844568 Fig. S2); †*Taimyrisphex* Evans, 1973 (see Rasnitsyn 1980).

***Note 1*:** The †Falsiformicidae and associated forms were recovered within the Pompiloidea *sensu novum* clade in the ancestral state analysis topology searches, a result which we remain skeptical about. However, because this was the case, we note the posterior probabilities here reported relate directly to the probability distribution of the Vespoides and internal nodes. To provide greater value, we also report in the polarity notes the conditions of †Bethylonymidae, Dryinaculeata, Vespaculeata, Vespoides, Pompiloidea *sensu novum*, and Tiphiopompiloides. Additionally, as †*Burmasphex* and two undescribed forms (CASENT0844587, CASENT0844568) cluster with †Falsiformicidae, we report transformations to †*Falsiformica* itself. The new genus represented by CASENT0844568 was recovered as sister to †*Burmasphex* + †*Falsiformica*.

***Note 2*:** Synapomorphies are traced for: (1) the †Falsiformicidae group including †*Burmasphex* and the two undescribed forms, †*Burmasphex* and form CASENT0844568 (“node 1”); (2) †*Burmasphex* + †*Falsiformica* + CASENT0844568 (“node 2”); (3) †*Burmasphex* + †*Falsiformica* (“node 3”); and †*Falsiformica* including †*F. musculosa* (Cockx & McKellar, 2016) **n. comb.** (“†Falsiformicidae *sensu stricto*”).

***Note 3*:** †Falsiformicidae were recently transferred from the Scolioidea *sensu* Rasnitsyn to the Vespoidea *sensu lato* by Rasnitsyn *et al*. (2020); this superfamilial composition is the paraphyletic set of all non-chrysidoid, non-dryinoid, and non-apoid Aculeata. This argument was based on the sexually dimorphic antennomere count and the internalization of female metasomal tergum VII. These characters were scored as uncertain in the present study (chars. **146**, **432**), with the exception of the male CASENT0844575 for antennomere count, which we now question. Recovery of the †Falsiformicidae within the Vespaculeata does predict that these two conditions would be observed. We retain our conservative placement until future study can confidently place the family.

***Note 4*:** Here we affirm the transfer of †*Siccibythus* to †Falsiformicidae by Rasnitsyn *et al*. (2020) based on the numerous derived features they share (see †Falsiformicidae *sensu stricto* synapomorphies below). We caution that the distinction between †*Falsiformica* and †*Siccibythus* recognized by Rasnitsyn *et al*. (2020) is tenuous. Specifically, †*Falsiformica* was redefined as those specimens with a tubular 1m-cu fore wing crossvein and

†*Siccibythus* as those specimens where this crossvein is at most nebulous. Given the high frequency of 1m-cu tubularity loss, as observed across the sampled taxa and as reconstructed, we consider this as weak evidence for reciprocal monophyly. Although we had considered synonymizing the two genera, we retain them here to encourage taxonomic stability.

***Note 5*:** †*Burmasphex*, a putative apoid (Melo & Rosa 2018), clustered with †Falsiformicidae in our analyses. Rosa & Melo (2021) explicitly treated the genus as a member of the †Angarosphecidae. Further scrutiny of fossils attributed to this genus is necessary. See also comments for Apoidea, below.

#### †Falsiformicidae group node 1 synapomorphies

**11.** Head prognathous, with an elongate postgenal bridge (CB50: 0.94). ***Polarity*:** Convergent with †Bethylonymidae, Bethylidae, various Dryinoidea, and Formicoidea.

**18.** Head wedge-shaped, elongate posterior to compound eyes (CB50: 0.92). ***Polarity*:** Also observed in Dryinidae, various Apoidea, and various Evaniomorpha.

**30.** Malar space well-defined, but not longer than broad (CB50: 0.77). ***Polarity*:** Only a synapomorphy in the context of Pompiloidea *sensu novum*, where the †Falsiformicidae group was recovered in ancestral state reconstruction topology searches.

**136.** Antennal toruli extremely-anteriorly situated, abutting or overhanging anterior clypeal margin (CB50: 0.93). ***Polarity*:** Among sampled taxa, also observed in many Chrysidoidea, Sclerogibbidae, Bradynobaenidae, *Martialis heureka*, *Lioponera longitarsus*, some †Ephialtitoidea, and †Bethylonymidae.

**143.** Antennal toruli directed toward the mandibles (CB50: 0.95). ***Polarity*:** Also observed in Bethylidae, Scolebythidae, Sclerogibbidae, *Olixon* (Rhopalosomatidae), Sierolomorphidae, some Tiphiiformes *sensu lato*, Mutillidae, *Apterogyna* (Bradynobaenidae), and some Apoidea.

**187.** Pronotum laterally scrobiculate, capable of receiving fore leg when leg raised and tucked inwards (CB50: 0.96). ***Polarity*:** Also observed in Bethylidae, Chrysididae, Dryinidae, Sierolomorphidae, *Tiphia* (Tiphiidae), Bradynobaenidae, Ampulicidae and many other Apoidea.

**223.** Prepectus indistinct, either fused or absent (exact state uncertain) (CB50: 0.71). ***Polarity*:** Support for this condition increases among internal nodes but is dominated by the uncertainty of this state among most included species.

**324.** Pretarsal claws edentate (CB50: 0.64). ***Polarity*:** Subapical teeth are present in †*Burmasphex* but absent in other species recovered in the †Falsiformicidae group. Such teeth are reconstructed as appearing and disappearing among various sampled clades.

**488.** Forewing Mf1 distinctly curved (CB50: 0.87). ***Polarity*:** Mf1 is curved in all sampled members of the †Falsiformicidae group except for one species of †*Falsiformica* (CNU-HYM-MA-2017089). This state varies homoplastically among sampled Aculeata.

**573.** Hindwing jugal lobe absent (CB50: 0.68). ***Polarity*:** Unobserved for most species sampled in the group. Variable across the Aculeata.

#### †Falsiformicidae node 2 synapomorphies

**86.** Clypeal disc with a median longitudinal carina (CB50: 0.88). ***Polarity*:** Also observed in Bethylidae, some Chrysididae, some Dryinidae, some Apoidea, and some Formicidae.

**207.** Mesoscutum width nearly twice length (CB50: 0.88). ***Polarity*:** Most frequently observed in Chrysidoidea: Bethylidae, Chrysididae, and some fossil Scolebythidae. Also observed in Dryinidae, *Olixon*, and Leptanillinae (Formicidae).

**212.** Lateral portions of Mesoscutum massively enlarged, such that each lateral portion is wider than the medial portion, and the two together with conspicuously greater surface area overall (CB50: 0.96). ***Polarity*:** Observed only in †*Burmasphex*, CASENT0844568, and some fossil Dryinidae.

#### †Falsiformicidae node 3 synapomorphies

**348.** Propodeum armed with posterior dorsolateral projections (CB50: 0.73). ***Polarity*:** Among species recovered in the †Falsiformicidae group, observed for †*Burmasphex* and most but not all †*Falsiformica*.

**375.** Abdominal tergum II (metasomal I) in lateral view with distinct posterior face offset from a dorsal face (CB50: 0.94). ***Polarity*:** Virtually unique, with the exception of Formicoidea and *Apterogyna* (Bradynobaenidae).

**403.** Articulatory surfaces of abdominal segment III (metasomal II) narrowed relative to remainder of segment (CB50: 0.85). ***Polarity*:** Formation of a “helcium” was also observed in Rhopalosomatinae, †*Eorhopalosoma*, some Vespidae, some Tiphiiformes, some Mutillidae, *Apterogyna*, *Cerceris* (Philanthidae), and Formicoidea. Comparison of internal anatomy will be useful.

#### †Falsiformicidae sensu stricto synapomorphies

**10.** Postgenal relatively long, with length ≥ 0.5 x width (CB50: 0.90). ***Polarity*:** Also observed in Bethylidae, †Chrysobythidae, *Amisega* (Chrysididae), some Dryinidae, *Ampulex* (Apoidea), some fossil Chrysidoidea, and Formicoidea. Among sampled †Falsiformicidae, could only be evaluated for two of the four terminals.

**178.** Pronotum in dorsal view nearly square, with anteroposterior length of disc ≥ lateromedial width (CB50: 0.97). ***Polarity*:** Observed for Chrysidoidea (infrequent), Sclerogibbidae, *Olixon* (Rhopalosomatidae), total clade Sierolomorphidae, and *Aelurus* (Tiphiiformes).

**179.** Pronotum with dorsolateral longitudinal margination between lateral and dorsal surfaces (CB50: 0.97). ***Polarity*:** Observed even less frequently than the prior condition, being confirmed present for very few Chrysidoidea, the Sclerogibbidae, *Olixon*, Sierolomorphidae, and various Formicidae (although these not scored).

**320.** Tarsomeres II–V of any leg swollen lateromedially or broadly laminate (CB50: 0.99). ***Polarity*:** A very strong synapomorphy of †*Falsiformica* which also observed in Rhopalosomatidae, although the form is distinct.

**326.** Pretarsus of fore leg with arolium grossly enlarged, spread by conspicuous, wide, paired, ellipsoid sclerites (CB50: 0.99). ***Polarity*:** Another strong synapomorphy of †*Falsiformica*. Also observed in Dryinidae, Sclerogibbidae, and *Olixon*.

**327.** Pretarsi of meso- and metalegs also enlarged as for the fore leg (CB50: 0.99). ***Polarity*:** As for char. 326, but not observed in Sclerogibbidae.

**378.** Node of petiole (metasomal segment I) dorsoventrally tall and very narrow anteroposteriorly, thus squamiform in shape (CB50: 0.98). ***Polarity*:** Among sampled taxa, a squamiform petiole was only observed in Formicidae: *Hypoponera* (Ponerinae), various Dolichoderinae, and Formicinae. ***Note*:** Inferred to be a synapomorphy of †Falsiformicidae by Rasnitsyn *et al*. (2020).

**409.** Articulatory surfaces of abdominal segment III (metasomal II) set below midheight of segment (CB50: 0.98). ***Polarity*:** An “infraaxial helcium” was also observed in *Chyphotes* (Chyphotidae) and Formicidae, including *Apomyrma* (Amblyoponomorpha), Ponerini (Ponerinae), Dolichoderomorpha, and Formyrmines.

**463.** Fore wing R not extending beyond pterostigma (CB50: 0.97). ***Polarity*:** This reduction was also observed in some Chrysidoidea, at least one dryinid fossil, Tiphiinae, Bradynobaenidae, and various Formicidae.

**\*.** Fore wing 2rrs cell (marginal 1) distally open (char. **471**, CB50: 0.98), consequently cell not pointed or narrowly rounded (char. **475**, CB50: 0.98). ***Polarity*:** An open 2rrs cell was observed in some Evaniidae, various Chrysidoidea, Embolemidae, Sclerogibbidae, Tiphiinae, and various Formicidae.

**501.** Fore wing Media nebulous to absent distad Rs+M (CB50: 0.98). ***Polarity*:** Observed in various Evanioidea, some Trigonaloidea, most Chrysidoidea, most Dryinoidea, *Olixon*, Bradynobaenidae, some Ammoplanidae, and various Formicidae. Also absent in CASENT0844567, but not CASENT0844568.

**511.** Fore wing crossvein 2rs-m nebulous to absent (CB50: 0.96). ***Polarity*:** Observed in various Evanioidea, most Chrysidoidea, Dryinoidea, *Olixon*, some Scoliidae, Bradynobaenidae, various Apoidea, and various Formicidae.

**517.** Fore wing crossvein 3rs-m nebulous to absent (CB50: 0.97). ***Polarity*:** More wide-spread than loss of 2rs-m; see character definition and raw data for state distribution.

**530.** Fore wing crossvein 1m-cu nebulous to absent (CB50: 0.97). ***Polarity*:** Observed to be absent in a few Evanioidea, many Chrysidoidea, Dryinidae, Sclerogibbidae, *Olixon*, Bradynobaenidae, and various Formicidae.

**538.** Fore wing crossvein 2m-cu nebulous to absent (CB50: 0.98). ***Polarity*:** As for loss of 3rs-m, more wide-spread than loss of 1m-cu; see character definition and raw data for state distribution. Also observed in CASENT 0.844567.

**558.** Fore wing Cu2 not reaching 1A vein (CB50: 0.95). ***Polarity*:** Observed in various Evanioidea, most Chrysidoidea, most Dryinoidea, Bradynobaenidae, and most Formicidae.

**\*.** Hind wing venation reduced: Rsf absent (char. **563**, CB50: 0.92); r-m crossvein nebulous to absent (char. **565**, CB50: 0.98); Mf2 nebulous to absent (char. **568**, CB50: 0.97); Cuf absent (char. **569**, CB50: 0.96); crossvein cu-a absent (char. **570**, CB50: 0.97). ***Note*:** Absence of closed hind wing cells was recognized as a synapomorphy of †Falsiformicidae by Rasnitsyn *et al*. (2020).

***Other states*:** Char. **15** (head broader lateromedially than long) shared with all constituent clades of Dryinaculeata and all constituent clades, to the exclusion of †Bethylonymidae. Char. **65** (mandible at least tridentate) shared with Dryinaculeata and †Bethylidae, but not other taxa recovered in the Vespoides in the ancestral state reconstruction analysis. Char. **87** (clypeal disc projecting from cranium) shared with Dryinaculeata and many of the constituent clades, to the exclusion of †Bethylonymidae. Closure of the meso- and metathoracic coxal cavities (chars. **272**, **273**) was observed for some †*Falsiformica*, but could not be confidently evaluated for most taxa in the tentative †Falsiformicidae group. Finally, Rasnitsyn *et al*. (2020) recognize presence of two narrow, upcurved processes on the male abdominal sternum IX as a synapomorphy for the family (unscored in our study but confirmed to be present).

### **A.** Superfamily Chrysidoidea Latreille, 1802

*Clade comprising*: • †**Plumalexiidae–Plumariidae** clade, • **†Chrysobythidae–Bethylidae** clade, • **Chrysididae–Scolebythidae** clade.

***Note***: Topological support for relationships among these clades is uncertain, see Results Part II above; recovered synapomorphies for the Bethylidae–Chrysididae clade are presented below as falsifiable hypotheses.

#### Synapomorphies

**136.** Antennal toruli situated at or extending anteriorly beyond anterior clypeal margin (50CB: 0.73). ***Polarity*:** Support for this condition is equivocal but included here to encourage for further hypothesis testing.

**139.** Clypeus extending posteriorly between antennal toruli (50CB: 0.62). ***Polarity*:** Support for this condition is equivocal but included here to encourage for further hypothesis testing.

**384.** Abdominal tergum II very broadly overlying sternum posteriorly, but narrowly so anteriorly (50CB: 0.93). ***Note*:** This condition was recovered as a synapomorphy of this clade by Brothers & Carpenter (1993).

**517.** Forewing crossvein 3rs-m not expressed (50CB: 0.85).

***Note*:** Support for either state of character 213 (parascutal carina of mesoscutum dorsad forewing articulation present or absent) is equivocal (50CB: 0.55). However, all internal clades of Chrysidoidea with the exception of Chrysididae lack this structure, implying either loss in the chrysidoid ancestor with “re-gain” of the carina in chrysidids, or parallel loss among the major groups.

### **A.1.** †Plumalexiidae–Plumariidae clade

***Clade comprising*:** • **†Plumalexiidae** Brothers, 2011 (†*Plumalexius* Brothers, 2011), • **Plumariidae** Bischoff, 1914 (*Maplurius* Roig-Alsina, 1994, *Mapluroides* Diez, 1994, *Myrmecopterina* Bischoff, 1914, *Myrmecopterinella* Day, 1977, *Plumarius* Philippi, 1873, *Plumaroides* Brothers, 1974a).

#### Synapomorphies

**201.** Propleurae forming tube supporting head (CB50: 0.88).

**548.** Pterostigma grossly enlarged (CB50: 0.86). ***Function*: *Polarity*:** Present analyses support parallel enlargement of the pterostigma between the †Plumalexiidae–Plumariidae clade, the †Chrysobythidae, Bethylidae, and the crown clade of the Scolebythidae.

**483.** Forewing Rsf1 and Mf1 meeting at a distinct angle, with the vertex of the angle directed vertically (CB50: 0.787). ***Polarity*:** Among Chrysidoidea, this condition is estimated to be gained in parallel between the †Plumalexiidae–Plumariidae clade and the Scolebythidae– Chrysididae clade.

**523.** Forewing “submarginal cell 1” (= 1rrs) as long or longer than both “submarginal cells 2” and “3” (= 3rsm, 4rsm) when these are present (CB50: 0.84). ***Polarity*:** A shorter submarginal cell is equivocally supported as ancestral for the Chrysidoidea and is estimated to be independently gained in the *Amisega*–*Loboscelidia* clade of the Chrysididae.

**548.** Forewing crossvein 1cu-a anterior junction postfurcal (CB50: 0.81). ***Polarity*:** Supported as gained in parallel between this clade and the Chrysididae (CB50: 0.80).

**558.** Forewing Cu2 not reaching 1A (CB50: 0.88). ***Polarity*:** Equivocally supported as parallel apomorphies of the Bethylidae–Chrysididae clade and the Scolebythidae (CB50: ≤ 0.55).

**561.** Hindwing basal hamuli present (CB50: 0.87). ***Polarity*:** Gained in parallel in some crown clade Bethylidae.

***Note* 1:** Support for character 15 (head wider than long) as a synapomorphy of this clade is equivocal (CB50: 0.57).

***Note* 2:** Because only two species of this tentative clade were sampled (†*Plumalexius rasnitsyni* and *Plumarius* sp.), no further synapomorphies can be provided. See the morphological data matrix for comparisons between these two taxa. Brothers & Melo (2021) recovered one unambiguous synapomorphy for this grouping (their character 42, state 4): endophragmal pit situated anteriorly, with the sulcus extending acutely ventrad pit. See Brothers & Melo (2021) and Brothers (2011) for parsimony optimizations of further characters.

### **A.2.** Bethylidae–Chrysididae clade

*Clade comprising*: • **†Chrysobythidae–Bethylidae** clade, • **Chrysididae–Scolebythidae** clade.

#### Synapomorphies

**219.** Alate female with notauli that are both parallel and extend from the anterior mesoscutal margin to the posterior (CB50: 0.77). ***Function*:** No precise prediction of function can be deduced at present. However, this form implies a distinct conformation of the mesothoracic indirect flight muscles.

**501.** Forewing free M distal to 1m-cu (Mf2+ or Mf3+) nebulous, spectral, or absent (CB50: 0.88). ***Polarity*:** Reversal to tubularity is supported for the crown clade of the Scolebythidae (CB50: 0.99).

**502.** Forewing with 3–4 “adventitious veins” which are present as creases or tubular to spectral abscissae. ***Polarity*:** Difficult to observe for most fossils, thus support for absence is marginally moderate for the Chrysidoidea and †Plumalexiidae–Plumariidae clade (CB50: 0.75, 0.76, respectively).

**563.** Hindwing Rsf spectral or absent (CB50: 0.87). ***Polarity*:** This character is supported as unreversed at the family level within the Bethylidae–Chrysididae clade. Rsf of the hindwing is moderately supported as ancestral for the Chrysidoidea (CB50: 0.88), and strongly supported as retained in the tentative †Plumalexiidae–Plumariidae clade (CB50: 0.99).

**565.** Hindwing r-m crossvein nebulous, spectral, or absent (CB50: 0.87). ***Polarity*:** The pattern and support for this condition is identical to character 563 above.

**568.** Hindwing Mf2 nebulous, spectral, or absent (CB50: 0.77). ***Polarity*:** Supported as unreversed at the family level within the Bethylidae–Chrysididae clade.

**569.** Hindwing Cuf nebulous, spectral, or absent (CB50: 0.79). ***Polarity*:** Supported as unreversed at the family level within the Bethylidae–Chrysididae clade.

**570.** Hindwing cu-a absent (CB50: 0.87). ***Polarity*:** Supported as unreversed at the family level within the Bethylidae–Chrysididae clade.

***Note* 1:** Support for character 216 (loss of anteromedian line of mesoscutum) as a synapomorphy of this node is equivocal (CB50: 0.56), although all clades within this group are reconstructed as lacking this feature, albeit with some uncertainty due to the included fossils.

***Note* 2:** Support for character 549 (forewing crossvein 1-cua interstitial) is equivocally reconstructed as ancestral to this clade (CB50: 0.55).

***Note* 3:** See also character 558 for the tentative †Plumalexiidae–Plumariidae clade.

### **A.2.1.** †Chrysobythidae–Bethylidae clade

*Clade comprising*: • **†Chrysobythidae** Melo & Lucena, 2019, • **Bethylidae** Haliday, 1839.

#### Synapomorphies

**30.** Malar space between compound eye and mandibular articulation virtually absent (CB50: 0.75). ***Polarity*:** †Chrysobythidae is variable for this trait (CB50: 0.75).

**86.** Clypeal disc with median longitudinal carina (CB50: 0.87). ***Polarity*:** Support for this condition in the ancestors of the †Chrysobythidae and Bethylidae is maximal or nearly so (CB50: 1.0, 0.99, respectively).

**94.** Clypeus produced anteromedially as a lobe (CB50: 0.88). ***Polarity*:** Support for this condition is maximal in the two constituent clades.

**143.** Antennal toruli rotated such that they are directed anteriorly, over the mandibles, rather than dorsally (CB50: 0.90). ***Polarity*:** As for the above character, this condition stabilizes in the ancestors of the constituent clades.

**187.** Pronotum laterally scrobiculate, capable of receiving the fore leg when this limb is tucked upwards and inwards (CB50: 0.68). ***Polarity*:** This condition is uncertain for some of the †Chrysobythidae, thus is equivocally reconstructed for that family as well as the †Chrysobythidae–Bethylidae clade.

**241.** Metapleural area scrobiculate, capable of receiving the mid leg when this limb is tucked upwards and inwards (CB50: 0.84). ***Polarity*:** As for the above characters, this condition stabilizes within the clade.

**273.** Mesocoxal cavity closed in lateral view (CB50: 0.55). ***Polarity*:** The uncertainty of character polarity at this node reflects the uncertainty for most †Chrysobythidae but is strongly supported as ancestral for the Bethylidae (CB50: 0.98).

**493.** Forewing Rsf2 absent such that 1m-cu apparently joining Rs+M and 2m-cu apparently joining “second submarginal cell (= 3rm) (CB50: 0.65). ***Polarity*:** This condition is strongly reconstructed as ancestral in the †Chrysobythidae (CB50: 1.0), but equivocally so in the Bethylidae (0.71).

**558.** Forewing Cu2 not reaching 1A (CB50: 0.63). ***Polarity*:** Although equivocal at this node, the support increased for the †Chrysobythidae (CB50: 0.85) (support for Bethylidae: CB50: 0.63). As noted for the †Plumalexiidae–Plumariidae clade above, this apomorphic condition is homoplastic among various Chrysidoidea.

### A.C. Family †Chrysobythidae Melo & Lucena, 2019

***Clade comprising*:** • †*Aureobythus* Melo & Lucena, 2019, • †*Bethylochrysis* Melo & Lucena, 2019, • †*Chrysobythus* Melo & Lucena, 2019.

#### Synapomorphies

**34.** Compound eyes converging medially toward the mandibles (CB50: 0.95). ***Polarity*:** This is an autapomorphy for the †Chrysobythidae in the context of the sampled Chrysidoidea.

**48.** Masticatory margin of mandible elongate, *i.e.*, mandible triangular in form (CB50: 0.99). ***Polarity*:** Autapomorphic among the sampled Chrysidoidea.

**66.** Mandible 4–9-dentate (CB50: 0.99). ***Polarity*:** Also true for †*Plumalexius* and crown Bethylidae.

**93.** Anterior clypeal margin linear, neither convex nor concave (CB50: 0.96). ***Polarity*:** Also observed in crown Scolebythidae and the Chrysididae, for which this condition is synapomorphic (50CB: 1.0, 0.78, respectively).

**177.** Pronotum elongate (CB50: 0.99). ***Polarity*:** Convergently derived in Scolebythidae.

**229.** Mesopectus with dorsoventrally oriented carina at about the anterolateral margin (CB50: 0.72). ***Polarity*:** Among Chrysidoidea, this condition is only duplicated in Chrysididae. The equivocal support reflects the observational uncertainty of this state among sampled †Chrysobythidae.

**219.** Notauli of mesoscutum either not completely transversing mesoscutum or not parallel (CB50: 0.99). ***Polarity*:** This apomorphic condition within the Chrysidoidea is supported as lost in parallel between the †Chrysobythidae and Bethylidae.

**279.** Protrochanter elongate, length > 3 x width (CB50: 0.99). ***Polarity*:** Also observed in various Bethylidae (CB50 for short protrochanters in the †chrysobythid–bethylid ancestor is equivocal, 0.59).

**345.** Propodeum with dorso- and posterolateral carinae, *i.e.*, propodeum marginate (CB50: 1.0). ***Polarity*:** Presence is equivocal for Bethylidae (CB50: 0.56).

**348.** Propodeum with angle at dorsal posterolateral corners, *i.e.*, propodeum armed (CB50: 0.99). ***Polarity*:** Presence is equivocal for Bethylidae (SB50: 0.56).

**382.** Abdominal sternum II with anteroventral process which fits between metacoxae at full metasomal flexion (CB50: 0.75). ***Polarity*:** Observed only for †*Aureobythus*, but nearly unique among sampled Chrysidoidea with the sole exception of the chrysidid *Amisega*.

**392.** Abdominal segment III with distinct laterotergite (CB50: 0.74). ***Polarity*:** Convergently derived in Chrysididae (CB50: 0.63, with condition stabilizing among internal nodes). The marginally equivocal support is likely due to uncertainty among †Chrysobythidae; this condition was only observable for †*Chrysobythus*.

**502.** Forewing without adventitious creases or veins (see also 502 for Bethylidae–Chrysididae clade above) (CB50: 0.81). ***Polarity*:** As for other traits, observable only for one terminal (†*Aureobythus*). This crease or fold system is generally observable for other Chrysididae.

### A.D+E. Families Bethylidae Haliday, 1839 + †Holopsenellidae Engel *et al*., 2016a

***Clade comprising*:** • **†Holopsenellidae** Engel *et al*., 2016a • **Bethylidae** Haliday, 1839 with seven subfamilies (see Azevedo *et al*. 2018 for synopsis and key): **†Protopristocerinae** Nagy, 1974, **Bethylinae** Haliday, 1839, **Epyrinae** Kieffer, 1914, **Lancepyrinae** Azevedo & Azar, 2012, **Mesitiinae** Kieffer, 1914, **Pristocerinae** Mocsáry, 1881, **Scleroderminae** Kieffer, 1914.

#### Synapomorphies

**5.** Body dorsoventrally flattened (CB50: 0.57). ***Polarity*:** Maximally supported as a synapomorphy of crown Bethylidae (CB50: 1.0).

**11.** Head prognathous (CB50: 0.97). ***Polarity*:** Maximally supported for crown group.

**19.** Head cube-shaped, with cranium elongated posterad eyes (CB50: 0.94). ***Polarity*:** Maximally supported for crown group.

**207.** Mesoscutum nearly twice as wide as long (CB50: 0.96). ***Polarity*:** Maximally supported for crown group.

**272.** Mesocoxal cavity closed in lateral view (CB50: 0.96). ***Polarity*:** Unreversed among total clade Bethylidae.

**280.** Fore femur grossly swollen along its entire length (CB50: 0.79). ***Polarity*:** Uncertain for some stem fossils, otherwise strongly supported for crown Bethylidae (CB50: 0.96). .

**295.** Meso- and/or metafemora of female swollen (CB50: 0.77). ***Polarity*:** Nearly maximal for crown Bethylidae (0.99).

***Note*:** The crown group of the sampled Bethylidae, excluding †*Holopsenella*, †*Holopsenelliscus*, and †*Lancepyris*, is also supported by the following synapomorphies: char. **10**, postgenal bridge elongate (CB50: 1.0); char. **15**, head longer than broad (CB50: 1.0; equivocal for total clade: CB50: 0.70); char. **149**, radicle offset from scape by distinct angle (CB50: 0.95); char. **256**, mesocoxae set in ventral thoracic sockets (CB50: 1.0, support for total Bethylidae node equivocal due to uncertainty: CB50: 0.57); char. **263**, mesocoxae wide-set (CB50: 1.0, support for close-set mesocoxae at CB50: 0.91); char. **288**, proximal articulations of meso- and metacoxae constricted and internalized (CB50: 1.0, uncertain for stem fossils); char. **342**, propodeum in form of elongate rectangular box (CB50: 1.0, supported as short in stem Bethylidae, CB50: 0.91); char. **390**, abdominal segment II laterotergite lost (CB50: 0.98, uncertain for stem Bethylidae); char. **463**, forewing R nebulous to spectral or absent distal to pterostigma (CB50: 0.78, support for tubular R in the ancestor of the total Bethylidae is maximal); char. **471**, forewing “marginal cell 1” (= 2rrs) open distally (CB50: 0.79, maximally supported as closed in total Bethylidae); char. **493**, forewing Rsf2 absent (CB50: 0.99, equivocal for total clade due to uncertainty, CB50: 0.71); char. **505**, forewing with “crease crossvein” between two of the longitudinal folds of the adventitious vein system (CB50: 0.95, could not be evaluated for all fossils); char. **511**, forewing crossvein 2rs-m absent or not tubular (CB50: 0.99, support moderate for presence in stem Bethylidae, CB50: 0.78); char. **530**, forewing crossvein 1m-cu absent or not tubular (CB50: 0.97, support is strong for retention of this vein at the total bethylid node, CB50: 0.92); char. **553**, forewing Cuf absent or not tubular (CB50: 0.77, strongly supported as retained at the total bethylid node, CB50: 0.99).

### **A.2.2.** Chrysididae–Scolebythidae clade

*Clade comprising*: • **Chrysididae** Latreille, 1802, • **Scolebythidae** Evans, 1963.

#### Synapomorphies

**92.** Anterior clypeal linear or concave (CB50: 0.73). ***Polarity*:** Although support for this condition at this node is equivocal, the linear to concave anterior margin stabilizes at the ancestral nodes of the Scolebythidae (CB50: 0.90) and Chrysididae (CB50: 0.90).

**386.** Abdominal tergum II (metasomal I) with distinct anterior concavity which fits against the propodeum when the metasoma is dorsally flexed, this concavity also laterally margined by dorsoventrally oriented carinae (CB50: 0.92). ***Polarity*:** Support increases crownward to the Scolebythidae (CB50: 0.98) and Chrysididae (CB50: 0.96), with uncertainty reflecting unobserved conditions in fossil taxa.

**471.** Forewing “marginal cell 1” (= 2rrs) distally open (CB50: 0.85). ***Polarity*:** As above, support increases among internal nodes (Scolebythidae CB50: 0.97, Chrysididae CB50: 0.98).

**511.** Forewing crossvein 2rs-m nebulous, spectral, or absent (CB50: 0.84). ***Polarity*:** Support increases crownward to the Scolebythidae (CB50: 0.95) and Chrysididae (CB50: 0.99).

### A.E. Family Chrysididae Latreille, 1802

***Clade comprising*:** • Four subfamilies as of Kimsey & Bohart (1990): **Amiseginae** Krombein, 1957, **Cleptinae** Latreille, 1802, **Loboscelidiinae** Ashmead, 1903, **Chrysidinae** Latreille, 1802. • ***Incertae sedis*** to subfamily: †*Auricleptes* Lucena & Melo, 2018, †*Burmasega* Lucena & Melo, 2018, †*Dahurochrysis* Rasnitsyn, 1990, †*Eochrysis* Doweld, 2015, †*Miracorium* Lucena & Melo, 2018.

***Note*:** Because sampling within the Chrysididae was dense relative to other Chrysidoidea, synapomorphies are provided at the family level as well as for the total Chrysidinae (†*Azanichrum* + †*Bohartiura* + an undescribed burmite fossil CASENT0844585), total Chrysidinae node 2 (including the undescribed fossil), and the crown Chrysidinae. The most recent study on chrysidid morphology was that of Lucena & Almeida (2022), who did morphology-only topology and chronogram estimation in a Bayesian framework using 300 characters and used parsimony to reconstruct transformation series across their sampled taxa.

#### Synapomorphies of Chrysididae

**21.** Occipital carina absent (CB50: 0.64). ***Polarity*:** Variable and uncertain among stem Chrysididae, thus support remains low to the crown Chrysidinae (CB50: 0.96).

**63.** Mandible at most unidentate (CB50: 0.57). ***Polarity*:** Support increases crownward, although trait variable.

**93.** Anterior clypeal margin linear (CB50: 0.78). ***Polarity*:** Support increases and remains stable among internal nodes of Chrysididae.

**154.** Scape length ≥ 4 x width (CB50: 0.67). ***Polarity*:** As above, support stabilizes among internal nodes.

**158.** Antennomere III (flagellomere I) longer than antennomere IV (flagellomere II) (CB50: 0.79). ***Polarity*:** Stabilizes among internal nodes.

**184.** Transverse line or groove present on pronotum (CB50: 0.79). ***Polarity*:** Increases and stabilizes among internal nodes. A line which completely traverses the pronotum is a supposed synapomorphy of the Cleptinae and is supported as lost in the ancestor of the Chrysidinae crown (CB50: 0.93).

**187.** Pronotum laterally scrobiculate, capable of receiving leg when folded up and inwards (CB50: 0.89). ***Polarity*:** Stabilizes as present among crown nodes.

**252.** Medial lobes of ventral mesopectal surface expanded, forming large plates (CB50: 0.52). ***Polarity*:** Uncertain for most fossils but stabilizes as present in Cleptinae and Chrysidinae (CB50: ≥ 0.90).

**277.** Procoxa with more-or-less dorsoventrally oriented carina on posterolateral surface (CB50: 0.59). ***Polarity*:** Uncertain for most fossils but supported as present in Cleptinae and Chrysidinae (CB50: ≥ 0.84).

**348.** Propodeum armed at dorsal posterolateral corners (CB50: 0.76). ***Polarity*:** Stabilizes within crown.

**392.** Abdominal segment III (metasomal II) with defined laterotergite (CB50: 0.63). ***Polarity*:** Stabilizes as present among crown nodes, although uncertain or variable for some fossils.

**432.** Abdominal tergum VIII of female fully internalized (CB50: 0.85). ***Polarity*:** Convergent with respect to Vespaculeata (see below).

**439.** Abdomen of female with ≤ 5 exposed segments, ≤ 6 in males (≤ 4 and ≤ 5 metasomal) (CB50: 0.88). ***Polarity*:** Stabilizing among crown nodes.

**483.** Forewing Rsf1 and Mf1 meeting at distinct oblique angle (CB50: 0.60). ***Polarity*:** Stabilizing among crown nodes, but variable.

**492.** Forewing Rs+M nebulous, spectral, or absent (CB50: 0.80). ***Polarity*:** Equivocal at the Chrysididae–Scolebythidae node (CB50: 0.51) and at that of the total Scolebythidae (CB50: 0.59), stabilizing as present in the crown Scolebythidae (CB50: 0.92) and absent within Chrysididae (CB50: 1.0).

**\*.** Forewing pterostigmal break crease extends posteroapically to Mf (char. **510**, CB50: 0.87), where these two meet, crease dividing into two distinct grooves which surround expected location of Mf (char. **508**, CB50: 0.87).

**531.** Forewing crossvein 1m-cu distinctly curved or sinuate (CB50: 0.63). ***Polarity*:** Stabilizing among crown nodes, although variable or uncertain.

**\*.** Forewing crossvein 1cu-a not interstitial (char. **549**, CB50: 0.82), rather being postfurcal (char. **548**, CB50: 0.80).

**558.** Forewing Cu2 not reaching 1A (CB50: 0.67). ***Polarity*:** Support increases within Chrysididae and is supported as a synapomorphy of the crown Scolebythidae (CB50: 0.99). State equivocal at the Chrysididae–Scolebythidae node (CB50: 0.53).

#### Synapomorphies of the total Chrysidinae

**136.** Antennal toruli extremely anteriorly situated, at or overhanging anterior clypeal margin (CB50: 1.0). ***Polarity*:** Maximally supported for total Chrysidinae.

**\*** Propodeum with short dorsal face (char. **343**, CB50: 0.80), consequently wedge-shaped in profile view (char. **344**, CB50: 0.82). ***Polarity*:** Maximally supported at the total and crown Chrysidinae nodes, and as FALSE at more-stemward nodes. State uncertain for †*Bohartiura*.

#### Synapomorphies of the total Chrysidinae node 2

**188.** Pronotum with laterally situated carina which is dorsoventrally oriented (CB50: 0.91). ***Polarity*:** Strongly supported as plesiomorphic in crown Chrysidinae (CB50: 1.0) and as absent at the total Chrysidinae node (CB50: 0.91).

**204.** Prosternum massive and broadly exposed (CB50: 1.0). ***Polarity*:** This condition is equivocal at the total Chrysidinae node (CB50: 0.57).

**\*** Mesopectus with epicnemial carina (char. **229**, CB50: 0.90) which margins a distinct anterior mesopectal scrobe (char. **330**, CB50: 0.99). ***Polarity*:** Strongly supported as TRUE for the crown Chrysidinae (CB50: 0.99, 1.0, respectively) and as FALSE for the total Chrysidinae node (CB50: 0.91, 0.99, respectively).

**\*.** Mesopectus with a scrobe posterior to the epicnemial carina which receives the mesoleg when the leg is flexed upwards and inwards (char. **234**, CB50: 0.98). ***Polarity*:** Equivocal at more-stemward nodes, and maximal at crown Chrysidinae node.

**239.** Metanotum lateral region in form of an angular process which is triangular in cross-section (CB50: 0.98). ***Polarity*:** Maximally supported for crown Chrysidinae, and strongly supported as FALSE for the total Chrysidinae node (CB50: 0.95).

**440.** Abdomen of female with ≤ 4 exposed segments, male with ≤ 5 (CB50: 0.97). ***Polarity*:** Maximally supported as TRUE for crown Chrysidinae, and as FALSE for the total Chrysidinae clade.

**442.** Abdominal segment II anteriorly truncate in dorsal view, with paired lobate process lateral to the propodeal articulation (CB50: 0.98). ***Polarity*:** Maximally supported as TRUE for crown Chrysidinae and as FALSE for the total Chrysidinae clade.

**443.** Abdomen ventral surface concave after sternum II (CB50: 0.98). ***Polarity*:** Maximally supported as true for the crown Chrysidinae, and strongly supported as FALSE for the total Chrysidinae clade (CB50: 0.99).

**444.** Abdominal terga II and III longer than IV+ (CB50: 0.99). ***Polarity*:** Maximally supported as TRUE for crown Chrysidinae, and strongly supported as FALSE for the total Chrysidinae clade (CB50: 0.93).

#### Synapomorphies of the crown Chrysidinae

**184.** Transverse line or groove on pronotum absent (CB50: 0.93). ***Polarity*:** Equivocal at the total Chrysidinae and total Chrysidinae nodes (CB50 < 0.64).

**199.** Pronotum distinctly separated from tegula (CB50: 0.98). ***Polarity*:** Support is maximal for the pronotum touching or nearly touching the tegula at more-stemward nodes (CB50: 1.0).

**207.** Mesoscutum width nearly twice its length (CB50: 0.94). ***Polarity*:** Equivocal at the total Chrysidinae node 2 (CB50: 0.53), and moderately supported as narrower at the total Chrysidinae node (CB50: 0.77).

**234.** Metapleural area included in posterior mesopectal scrobe (CB50: 0.94). ***Polarity*:** Metapleural area strongly supported as not included in posterior mesopectal scrobe at the total Chrysidinae node 2 (CB50: 0.90).

**486.** Forewing Mf1 long to extremely long, being ≥ 4 x length of Rsf1 (CB50: 0.93). ***Polarity*:** Strongly supported as FALSE for the total Chrysidinae node 2 (CB50: 0.91) and the total Chrysidinae (CB50: 0.93). Convergently derived in †*Burmasega* and †*Miracorium*.

**573.** Hindwing jugal lobe present (CB50: 0.92). ***Polarity*:** Uncertain for total Chrysidinae node 2 (CB50 for absence of jugal lobe: 0.64), but strongly supported as absent for the total Chrysidinae clade (CB50: 0.97).

***Notes*:** Presence of an antennal scrobe on the face (char. **126**) is homoplastic between the Amiseginae–Loboscelidiinae clade and the Chrysidinae when Cleptinae are not sister to the remainder of the family. Only one character is strongly supported as uniting Amiseginae and Loboscelidiinae (char. **553**, forewing Cuf not tubular).

### A.F. Family Scolebythidae Evans, 1963

***Clade comprising*:** • **Scolebythinae** Evans, 1963 (*Clystopsenella* Kieffer, 1911, *Scolebythus* Evans, 1963), • **Pristapenesiinae** Engel *et al*., 2013 (†*Boreobythus* Engel & Grimaldi, 2007, †*Ectenobythus* Engel *et al*., 2013, †*Eobythus* Lacau *et al*., 2000, †*Libanobythus* Prentice & Poinar, 1996, †*Mirabythus* Ren *et al*., 2019, †*Nadezhdabythus* Zhang *et al*., 2020, †*Necrobythus* Engel *et al*., 2013, †*Sphakelobythus* Engel *et al*., 2013, †*Uliobythus* Engel & Grimaldi, 2007, †*Zapenesia* Engel & Grimaldi, 2007, *Pristapenesia* Brues, 1933, *Ycaploca* Nagy, 1975). • ***Incertae sedis*** to subfamily: †*Cursoribythus* Cockx & McKellar, 2016 **subfam. transfer**.

***Note 1*:** †*Siccibythus* Cockx & McKellar, 2016 was transferred to the †Falsiformicidae by Rasnitsyn *et al*. (2020) while the present work was in revision. Rasnitsyn *et al*. also considered transferring †*Cursoribythus*, but noted high uncertainty, so we retain the original placement here until further resolution can be attained.

***Note 2*:** Among the genera listed above, one species each was sampled from †*Necrobythus*, †*Zapenesia*, *Clystopsenella*, and *Ycaploca*, plus one undescribed species from Burmite (CASENT0844570). Because the two described fossil species were recovered as a clade sister to the remaining three sampled taxa, a distinction is here drawn among “total node 1” encompassing all sampled taxa, “total node 2” which excludes †*Necrobythus* and †*Zapenesia*, and the crown node which includes the two extant species.

#### Synapomorphies, total node 1

**19.** Head elongate posterad compound eyes (CB50: 0.60). ***Polarity*:** Increasing to crown node (CB50: ≥ 0.92).

**201.** Propleurae forming narrow tube-like collar supporting head (CB50: 0.84). ***Polarity*:** Convergent with Plumariidae.

**204.** Prosternum massive and broadly exposed (CB50: 0.78). ***Polarity*:** Also occurring in Chrysidinae.

#### Synapomorphies, total node 2

**143.** Antennal toruli directed toward mandibles (CB50: 0.94). ***Polarity*:** Uncertain at total node 1 (CB50 for toruli having another conformation: 0.65).

**224.** Prepectus fused with mesopectus but remaining visible (CB50: 0.56). ***Polarity*:** Increasing at the crown node (CB50: 0.93), and equivocal for total node 1 (CB50: 0.59 for prepectus being unfused).

**280.** Female fore femur grossly swollen (CB50: 0.85). ***Polarity*:** Support maximal for swollen femora in the crown Scolebythidae.

**295.** Female meso- and/or metafemora swollen (CB50: 0.84). ***Polarity*:** Support becomes maximal for the crown Scolebythidae.

**391.** Abdominal segment II laterotergite dorsoventrally broad (CB50: 0.92). ***Polarity*:** Support is maximal for this condition at the Scolebythidae crown node.

**492.** Forewing Rs+M tubular (CB50: 0.99). ***Polarity*:** This reconstruction represents regain of the tubular expression of the Rs+M abscissa for total node 2 and the crown node of the Scolebythidae.

#### Synapomorphies, crown node

**274.** Procoxa extended posterior to trochanteral articulation as distinct lobe (CB50: 1.0). ***Polarity*:** Procoxa unmodified for total nodes 1 and 2 (CB50: 1.0 for both).

**349.** Propodeal spiracle situated on lateral propodeal surface (CB50: 1.0). ***Polarity*:** Equivocal for total node 1.

**350.** Propodeal spiracle circular (CB50: 0.91). ***Polarity*:** Support for a slit-shaped propodeal spiracle is equivocal for total node 2 (CB50: 0.64) but is moderate for total node 1 (CB50: 0.86).

**427.** Female abdominal sternum VII (metasomal VI) with posteromedian triangular structure (CB50: 0.88). ***Polarity*:** Observed only for *Clystopsenella*, so this hypothesis should be subjected to further testing with expanded taxon sampling.

**454.** Most or all of distal fourth of fore wing without tubular or nebulous venation (CB50: 0.99).

**458.** Pterostigma grossly enlarged (CB50: 0.99).

**\*.** Forewing “marginal cell 1” (= 2rrs) closed (char. **471**, CB50: 0.99), apex of cell pointed to narrowly rounded (char. **475**, CB50: 0.98), with apex somewhat “downturned” toward posterior margin (char. **477**, CB50: 0.99). ***Polarity*:** Closure of marginal cell 1 (char. 471) represents a reversal from the condition at the Chrysididae–Scolebythidae node, as well as total nodes 1 and 2 (CB50: ≥ 0.96).

**483.** Forewing Rsf1 and Mf1 meeting at distinct oblique angle (CB50: 0.98). ***Polarity*:** Convergent with Chrysididae.

**493.** Forewing Rsf2 absent (CB50: 0.99). ***Polarity*:** Well-supported as a synapomorphy.

**501.** Forewing free M tubular distal to Rs+M (CB50: 0.99). ***Polarity*:** The non-tubular condition is strongly supported at more-stemward nodes (CB50: ≥ 0.97).

4. **530.** Forewing 1m-cu tubular (CB50: 0.99). ***Polarity*:** Moderately supported at the two more-stemward nodes (CB50 = 0.89).

### A’. Clade Dryinaculeata

See the main transformation series (*i.e.*, leading to Formicoidea) above for definition based on the present analyses.

### **B.** Superfamily Dryinoidea Haliday, 1833

*Clade comprising*: • **Sclerogibbidae** Ashmead, 1901, clade **Dryinomorpha**.

#### Synapomorphies

**553.** Fore wing CuF nebulous to absent (CB50: 0.53). ***Polarity*:** Although equivocal, only observed in the extant sampled sclerogibbid; support for this condition is strong to maximal among other internal dryinoid nodes.

**558.** Fore wing CuF not reaching 1A (CB50: 0.54). ***Polarity*:** Similar to char. 554, although support is equivocal at this node, CuF meeting 1A was not observed for extant Dryinoidea and the fossil which could be evaluated for this condition.

**563.** Hind wing Rsf2 spectral to absent (CB50: 0.55). ***Polarity*:** Could not be evaluated for fossils. Rsf2 is not developed in the sampled crown dryinoids.

**565.** Hind wing r-m crossvein nebulous to absent (CB50: 0.55). ***Polarity*:** This crossvein was observed to be absent in crown dryinoids as well as for †*Sclerogibbodes embioleia* Engel & Grimaldi, 2006a.

**570.** Hind wing cu-a absent (CB50: 0.58). ***Polarity*:** Exactly as for char. 565 above.

***Note*:** Support for the venational reduction characters provisionally recorded as dryinoid synapomorphies above is limited largely due the difficulty of evaluating fossil taxa.

### **B.A.** Family Sclerogibbidae Ashmead, 1902

***Clade comprising*:** • †**Sclerogibbodinae** Engel & Grimaldi, 2006a (†*Sclerogibbodes* Engel & Grimaldi, 2006a), • **Sclerogibbinae** Ashmead, 1902.

#### Synapomorphies

**11.** Cranium prognathous (CB50: 0.71). ***Polarity*:** The low support reflects uncertainty for †*Sclerogibba cretacica* Martynova *et al*., 2019.

**147.** Antennomere count ≥ 13 for both sexes (CB50: 0.99). ***Polarity*:** As noted for the Dryinaculeata above, this is a strongly supported reversal to the many-annulate condition.

**295.** Female meso- and/or metafemora muscularly swollen (CB50: 0.54). ***Polarity*:** True for those sampled taxa which could be evaluated; could not be observed for †*Sclerogibba cretacica*.

**501.** Fore wing Mf nebulous distal to Rs+M (CB50: 0.98). ***Polarity*:** This was observed to be true for all sampled Sclerogibbidae for which this condition could be evaluated, likewise for Dryinidae. Mf distal to Rs+M was observed to be tubular for most Embolemidae, with the exception of †*Cretembolemus*.

***Notes*:** Occipital carina presence (char. **21**) was variable among sampled taxa. Reduction and loss of 2rs-m (char. **511**, CB50: 0.79) and of 1m-cu (char. **530**, CB50: 0.79) is ambiguously reconstructed as a synapomorphy of the Dryinidae–Embolemidae clade. Regarding the former, 2rs-m was observed to be tubular in †*Sclerogibba cretacica*, although absent in the extant species, suggesting parallel loss. Although only one rs-m crossvein was observed in †*S. cretacica*, the following the suggested interpretation of Rasnitsyn (1988) and Rasnitsyn in Rasnitsyn & Quicke (2002) for other species. Regarding 1m-cu, this crossvein was observed to be tubular in the extant embolemid and nebulous to absent in the extant sclerogibbid, again suggesting parallel reductions.

### B.A’. Clade Dryinomorpha

*Clade comprising*: • **Dryinidae** Haliday, 1833, • **Embolemidae** Förster, 1856.

***Note*:** Rasnitsyn & Quicke (2002) hypothesized that Dryinidae and Embolemidae were their closest relatives based on these two conditions: “antenna attached well above clypeus” and “pronotum circular, being fused with symprepectus”. The former character could not be evaluated for most of the dryinid fossils; thus the “distant” conformation was recovered as a synapomorphy of the Embolemidae. Future study should clarify the diagnosis of this clade.

#### Synapomorphies

**148.** Antenna 10-merous in females (CB50: 0.96). ***Polarity*:** Male Embolemidae have 11-merous antennae, whereas those of females and both sexes of Dryinidae are 10-merous, at least among sampled taxa. It is plausible that the antennomere count of male dryinids stabilized at 10 independently, as such intersex variation is known to occur in Formicidae. That both sexes in Dryinidae are 10-merous is likely a synapomorphy of the family, but this was not evaluated as a separate trait in the present study.

**324.** Pretarsal claw unidentate (CB50: 0.74). ***Polarity*:** The uncertainty of this estimate is associated with those fossils which could not be evaluated.

**511.** Fore wing crossvein 2rs-m nebulous to absent (CB50: 0.79). ***Polarity*:** Support for this condition stabilizes as strong among internal nodes. Crossvein 2rs-m was observed to be tubular in †*Sclerogibba cretacica*, although absent in the extant species, suggesting parallel loss. Although only one rs-m crossvein was observed in †*S. cretacica*, the following the suggested interpretation of Rasnitsyn (1988) and Rasnitsyn in Rasnitsyn & Quicke (2002) for other species.

### **B.B.** Family Embolemidae Förster, 1856

***Clade comprising*:** • **Core Embolemidae** clade (†*Baissobius* Rasnitsyn, 1975, crown Embolemidae clade [•• *Ampulicomorpha* Ashmead, 1893, *Embolemus* Westwood, 1833]). • ***Incertae sedis*** in family: †*Cretembolemus* Olmi *et al*., 2014, †*Embolemopsis* Olmi *et al*., 2010.

***Note*:** Ancestral state reconstruction analysis only included fossil *Ampulicomorpha* and an extant species; other taxa were too incomplete to include. See Rasnitsyn (1996) for morphological information, and the polarity comments below for information on character state distributions within the family.

#### Synapomorphies

**15.** Cranium anteroposteriorly longer than dorsoventrally tall (CB50: 0.98). ***Polarity*:** Associated with the conification of the face. Although head proportions does vary considerably among Aculeata, the combination of this and the following trait are highly unusual.

**16.** Cranium cone-shaped, with antennal toruli situated on an anterior process which broadens toward the mandibles and posterior head margin (CB50: 1.0). ***Polarity*:** Unique among sampled Hymenoptera.

**21.** Occipital carina absent (CB50: 0.92). ***Polarity*:** Unique among sampled Dryinoidea for which this condition could be evaluated.

**36.** Female compound eyes reduced in size, being < ½ head length in full-face view (CB50: 0.96). ***Polarity*:** Variable among sampled Embolemidae: reduced in †*Ampulicomorpha*, †*Baissobius*, but not the extant species, †*Cretembolemus*, or †*Embolemopsis*. Reanalysis of this condition including a broader taxon sample may overturn this provisional synapomorphy.

**138.** Antennal toruli located at about head midlength when viewed anteriorly (CB50: 0.99). ***Polarity*:** Could not be evaluated for †*Baissobius*, †*Cretembolemus*, or †*Embolemopsis*.

**140.** Antennal toruli distinctly separated from clypeus (CB50: 0.78). ***Polarity*:** Among the sampled fossil taxa, this could only be evaluated for †*Ampulicomorpha janzeni*.

**\*** Scape elongated, with length ≥ 4 x width (char. **154**, CB50: 0.97), and length > ½ head length (char. **155**, CB50: 0.99). ***Polarity*:** Scape length could not be evaluated for †*Baissobius*, †*Cretembolemus*, or †*Embolemopsis*.

**158.** Antennomere III longer than IV (CB50: 0.97). ***Polarity*:** Could be evaluated for †*Baissobius minimus*, and true for various Dryinidae.

**345.** Propodeum with dorsolateral and posterolateral margins carinate (CB50: 0.82). ***Polarity*:** Also observed for the sampled extant species of Sclerogibbidae; could not be evaluated for most fossil Embolemidae.

***Note*:** Forewing crossvein 1cu-a was strongly supported as prefurcal for the Embolemidae (char. **550**, CB50: 0.93). However, 1cu-a does not have this condition in †*Baissobius* and †*Embolemopsis*, indicating that may not be synapomorphy of the family, particularly with inclusion of these taxa in the ancestral state estimation analyses.

### **B.C.** Family Dryinidae Haliday, 1833

***Clade comprising*:** • 12 subfamilies (see Olmi 1984, 1990, Tribull 2015, Olmi *et al*. 2020): •• Extinct lineages: †**Archaeodryininae** Olmi *et al*., 2020, †**Burmadryininae** Olmi *et al*., 2014, †**Palaeoanteoninae** Olmi & Bechly, 2001, †**Ponomarenkoinae** Olmi, 2010, †**Protodryininae** Olmi & Guglielmino, 2012, †**Raptodryininae** Olmi *et al*., 2020, •• Extant lineages: **Anteoninae** Perkins, 1912, **Aphelopinae** Perkins, 1912, **Bocchinae** Richards, 1939, **Dryininae** Haliday, 1833, **Gonatopodinae** Kieffer in Kieffer & Marshall, 1906, **Thaumatodryininae** Perkins, 1905.

#### Synapomorphies

**11.** Cranium prognathous (CB50: 0.99). ***Polarity*:** Although not all extant Dryinidae have prognathous crania (some males), this condition was observed to be generally true for all sampled taxa, with some uncertainty where the cranium could not be evaluated in detail.

**18.** Cranium more-or-less wedge-shaped (CB50: 1.0). ***Polarity*:** Unique among Dryinoidea and pronounced in males. Also observed among the taxa of the recovered †*Falsiformica* group.

**19.** Cranium elongate posterad compound eyes (CB50: 0.77). ***Polarity*:** Variable among Aculeata; not true for all crown Dryinidae.

**182.** Female pronotum with grossly expanded flange, forming a broad collar around the propleurae, and partially encircling the posteromedian muscular portion of the notum (CB50: 0.71). ***Polarity*:** This condition was observed for all Dryinidae for which females could be evaluated.

**187.** Pronotum laterally furrowed such that it can receive the fore leg when this appendage is folded dorsomedially (CB50: 0.76). ***Polarity*:** Observed for all sampled Dryinidae for which this condition could be evaluated.

**191.** Pronotum broadly constricted posteriorly, just anterior to mesoscutum (CB50: 0.67). ***Polarity*:** Not observed to be the condition in *Deinodryinus atriventris* but was observed to be true for *Dryinus gulfensis* and those fossils for which this trait could be evaluated.

**286.** Propretarsus with one of the claws reduced or absent (CB50: 0.99). ***Polarity*:** Observed for all sampled Dryinidae for which this condition could be evaluated.

**295.** Female meso- and/or metafemora muscularly swollen (CB50: 0.76). ***Polarity*:** Observed for all sampled Dryinidae for which the female could be evaluated.

**310.** Metatibia with a single apical spur (CB50: 0.97). ***Polarity*:** True for those Dryinidae which could be evaluated. Also recorded as the condition for †*Sclerogibbodes*.

**471.** Fore wing 2rrs cell (“marginal 1”) distally open (CB50: 0.98). ***Polarity*:** This cell was also observed to be open in some but not all Embolemidae and was open in all sampled Dryinidae.

**492.** Fore wing Rs+M nebulous to absent (CB50: 0.97). ***Polarity*:** Rs+M was observed to be tubular in all sampled Embolemidae and Sclerogibbidae, and nebulous to absent for all sampled Dryinidae for which this condition could be evaluated.

**501.** Fore wing Mf nebulous distal to Rs+M (CB50: 0.98). ***Polarity*:** This was observed to be true for all sampled Dryinidae for which this condition could be evaluated, likewise for Sclerogibbidae. Mf distal to Rs+M was observed to be tubular for most Embolemidae, with the exception of †*Cretembolemus*.

### B’. Clade Vespaculeata

See the main transformation series (*i.e.*, leading to Formicoidea) above for definition based on the present analyses.

### B’.1. Clade Vespoides

*Clade comprising*: • **Vespoidea** Laicharting, 1781, • clade **Pompiloidea** Latreille, 1804 *sensu novum*.

#### Synapomorphies

**30.** Malar space virtually absent (CB50: 0.69). ***Polarity*:** We recognize that this condition is equivocal in our analyses. Support for this condition stabilizes for the Vespoidea (CB50: 0.91) and Pompiloidea *sensu novum* (CB50: 0.78). Scolioides and Formicapoidina were equivocally reconstructed as having a broader malar space (CB50: 0.58, 0.62, respectively). For this reason, we consider the present synapomorphy as provisional, and definitely in need of further scrutiny. Malar space development is variable within the major vespaculeate clades.

### **C.** Superfamily Vespoidea Laicharting, 1781

*Clade comprising*: • †*Prosphex* genus group, • **Rhopalosomatidae** Ashmead, 1896, • **Vespidae** Laicharting, 1781.

***Note*:** We are dissatisfied with certain characterizations of vespoid morphology after *post hoc* reassessment of variability, for which reason we provide explicit notes for features that we recommend be reevaluated in future study.

#### Synapomorphies

**35.** Medial margins of compound eyes emarginate (CB50: 0.79). ***Polarity*:** The condition of eyes without medial notches was strongly supported for the Pompiloidea *sensu novum* (CB50: 0.97) and maximally so for the Scolioides (CB50: 1.0), although we recognize that emarginate eyes do occur in those clades.

**251.** Medial lobes of ventral mesopectal surface expanded ventrally, rather than medially, forming distinct plates which at least partially overlap the mesocoxae (CB50: 0.75). ***Polarity*:** This condition is estimated to be reversed in *Parischnogaster* and *Euparagia*, and it could not be evaluated for the majority of fossils with the exception of CASENT0844576 and †*Eorhopalosoma lohrmanni*. Various Tiphiiformes display a similar condition. “Mesosternum with posterad paired lobes covering the bases of the mesocoxae”, corresponding to the definition here, was adduced as a synapomorphy of the total clade Rhopalosomatidae by Lohrmann *et al*. (2020).

**353.** Propodeal ventrolateral margin extended posteriorly to posterodorsally, thus distinctly separating the propodeal foramen from the metacoxae (CB50: 0.97). ***Polarity*:** This is recovered as a strong synapomorphy of the superfamily, which was not observed in other sampled Aculeata.

**453.** Fore wing Rsf extending to or almost to distalmost wing margin (CB50: 0.75). ***Polarity*:** Although equivocal at the Vespoidea node, this is strongly supported for the Rhopalosomatidae (CB50: 0.95), moderately for the †*Prosphex* group + Vespidae node (CB50: 0.83), and strongly for the total Vespidae node (CB50: 0.97). Additionally, the Vespaculeata are moderately supported as not having Rsf reach the wing apex (CB50: 0.77). Support for the Vespoides and Pompiloidea *sensu novum* lacking this condition is equivocal (CB50: 0.70 for both). The reconstruction supports optimization for strong flight in Vespoidea, in comparison to Pompiloidea *sensu novum*, Dryinoidea, and Chrysidoidea.

***Note*:** Rhopalosomatidae and Vespidae are both supported as having a distinct, broad, translucent to opaque laminar process extending from the ventrolateral propodeal margin above the metacoxae, shielding the metasomal insertion in lateral view (char. **354**). This lamina was scored as absent in CASAENT0844576; consequently, our ancestral state estimation analysis supports independent origin between the two families. Without this fossil, however, the lamina is an apparent synapomorphy of the superfamily.

### **C.A.** Family Rhopalosomatidae Ashmead, 1896

***Clade comprising*:** • **Crown Rhopalosomatidae: •• Rhopalosomatinae** Ashmead, 1896 (*Liosphex* Townes, 1977, *Paniscomima* Enderlein, 1904, *Rhopalosoma* Cresson, 1865), •• **Olixoninae** Engel, 2008 (*Olixon* Cameron, 1887), •• ***Incertae sedis*** in crown: †*Cretolixon* Lohrmann *et al*., 2020. • ***Incertae sedis*** in family: †*Eorhopalosoma* Engel, 2008, †*Mesorhopalosoma* Darling, 1990.

***Note 1*:** †*Mesorhopalosoma* was moved out of the Rhopalosomatidae by Osten (2007) but was recovered with the family in our topology searches; it is thus provisionally treated as a taxon that is *incertae sedis* in the group. See also Boudinot & Dungey (2020). †*Cretolixon* was recently described by Lohrmann *et al*. (2020) and placed by them in the crown clade, but with uncertain placement therein. We were unable, unfortunately, to include this taxon in our present analyses. See that study for morphological features which will be important to consider for future phylogenetic study.

***Note 2*:** We recovered †*Eorhopalosoma* as sister to the crown Rhopalosomatidae, which comprised Olixoninae and Rhopalosomatinae. For this reason, we list synapomorphies at the levels of total and crown Rhopalosomatidae, and for the Rhopalosomatinae. “Mesosternal lobes” directed posterad over mesocoxal bases is a putative synapomorphy of the total clade Rhopalosomatidae (Lohrmann *et al*. 2020) but was here recovered as a synapomorphy of the Vespoidea. Further comparative definition of the states of the mesosternal lobes are necessary.

#### Total Rhopalosomatidae synapomorphies

**30.** Malar space distinctly developed (CB50: 0.80). ***Polarity*:** Although some crown Vespidae have well-developed malar areas, our analysis recovers the “virtually absent” condition as a retained plesiomorphy of the family.

**\*** Anterior clypeal margin not convex or angular (char. **92**, CB50: 0.98), being rather linear (char. **93**, CB50: 0.86). ***Polarity*:** The clypeus of all sampled extant Vespidae was observed to be convex, angular, or with angular projections.

**184.** Pronotum with transverse line or groove on lateral to lateromedial surfaces (CB50: 0.83). ***Polarity*:** This may be a plesiomorphy of the Vespoidea, but support is equivocal at this node. Extant Vespidae were observed to lack this sulcus.

**314.** Meso- and metatibial spurs extremely elongate, with the posterior spur ≥ ½ basitarsal length (CB50: 0.97). ***Polarity*:** Also observed in some Tiphiiformes, Scolioides, and spheciform Apoidea.

**370.** Abdominal segment II (petiole) not sessile, being rather subsessile to pedunculate (CB50: 0.91). ***Polarity*:** Supported as independently derived in various Vespidae, including Stenogastrinae, Zethinae, and Polistinae.

**392.** Abdominal segment III (metasomal II) with distinct laterotergite (CB50: 0.93). ***Polarity*:** Among sampled Vespoidea, also observed in *Euparagia*; equivocally reconstructed as an ancestral feature of Pompiloidea *sensu novum* (CB50: 0.57) and as absent in the ancestor of the Vespoidea (CB50: 0.75).

**403.** Abdominal segment III anterior articulatory surfaces narrowed relative to remainder of segment (CB50: 0.78). ***Polarity*:** Supported as reversed to a broad condition in *Olixon*, and as independently derived in Vespidae at least once, being observed in *Zethus* and *Mischocyttarus*.

**459.** Pterostigma strongly reduced (CB50: 0.78). ***Polarity*:** Support increases for the crown Rhopalosomatidae (CB50: 0.90). A reduced pterostigma is also observed in *Euparagia* among sampled crown Vespidae, and in †*Curiosivespa* and †*Priorparagia*. The pterostigma is unreduced in †*Priorvespa* and †*Protovespa*.

**462.** Fore wing C contacting Sc+R+Rs for most or all of its length (CB50: 0.95). ***Polarity*:** Unique among all sampled Aculeata.

**482.** Fore wing Rsf1 anterior juncture with Rf contacting or nearly contacting pterostigma (CB50: 0.97). ***Polarity*:** Also strongly supported for the Vespidae, but not the condition in the prospective †*Prosphex* group. Given the lingering uncertainty about †*Prosphex*, it is possible that Rsf1 contacting or nearly contacting the pterostigma may be a synapomorphy of the Vespoidea with some degree of reversal.

**517.** Fore wing crossvein 3rs-m nebulous to absent (CB50: 0.93). ***Polarity*:** Crossvein 3rs-m was observed to be absent in all sampled macropterous Rhopalosomatidae, and to be retained in all Vespidae except for *Pseudomasaris*.

**533.** Fore wing crossvein 1m-cu and Mf3 forming a straight line (CB50: 0.95). ***Polarity*:** This is the condition for all sampled macropterous Rhopalosomatidae and was not observed in any vespid.

**538.** Fore wing crossvein 2m-cu nebulous to absent (CB50: 0.95). ***Polarity*:** Crossvein 2m-cu was observed to be absent in all sampled macropterous Rhopalosomatidae and was retained in all Vespidae for which this condition could be evaluated.

**548.** Fore wing crossvein 1cu-a anterior junction distinctly “postfurcal”, *i.e.*, distal to split of Mf and Cuf (CB50: 0.88). ***Polarity*:** Crossvein 1cu-a was observed to be postfurcal for all sampled macropterous Rhopalosomatidae and was reconstructed as independently postfurcal in some Vespidae.

**561.** Hind wing basal hamuli present (CB50: 0.83). ***Polarity*:** Among Vespoidea, only observed for Rhopalosomatidae. Could not be evaluated for fossil Vespidae.

**571.** Hind wing cu-a crossvein interstitial to prefurcal, *i.e.*, at or proximal to the split of Mf and Cuf (CB50: 0.85). ***Polarity*:** The postfurcal condition is supported as ancestral for Vespoides and Vespoidea (CB50: 0.89, 0.75, respectively). The interstitial to prefurcal condition is observed in *Euparagia* and *Pseudomasaris* among sampled crown Vespidae.

**572.** Hind wing sectoral cell (“basal cell”) without distal elongation (CB50: 0.85). ***Polarity*:** The distal elongation is observed in all sampled Vespidae for which this condition could be evaluated and is also supported as ancestral for the Vespoides and Vespoidea (CB50: 0.86, 0.90, respectively).

#### Crown Rhopalosomatidae synapomorphies

**138.** Antennal toruli situated at or posterior to head midlength (CB50: 0.84). ***Polarity*:** Difficult to examine for †*Eorhopalosoma*, so should be subject to reevaluation.

**175.** Pronotum without distinct anteromedian lobe forming “neck” (CB50: 0.85). ***Polarity*:** Estimated to be independently derived in crown Vespidae.

#### Rhopalosomatinae synapomorphies

**206.** Mesoscutum anteroposteriorly longer than lateromedially broad in dorsal view (CB50: 0.97). ***Polarity*:** The mesoscutum was observed to be broader in both sampled †*Eorhopalosoma* species and in *Olixon*.

***Note 1*:** The traditional synapomorphy adduced for the family (char. **160**, paired chaetae at apices of flagellomeres) are present on extant Rhopalosomatinae, very small (reduced) to absent in *Olixon*, and have been previously recorded as absent in †*Eorhopalosoma gorgyra* Engel, 2008, although they are present in †*E. lohrmanni*. Apparently, for these reasons, such setae were supported as a synapomorphy of the crown group, for which we retain skepticism. The apical chaetae were adduced as a synapomorphy of the total clade Rhopalosomatidae by Lohrmann *et al*. (2020).

***Note 2*:** The mesepimeron of †*Eorhopalosoma lohrmanni* Boudinot & Dungey, 2020 and sampled Rhopalosomatinae is present as a distinct narrow sclerite posterior to the pleuropleural suture of the meso- and metapectus (char. **237**). However, †*E. lohrmanni* was incorrectly scored as “absent”, thus this feature is strongly supported in our dataset as a synapomorphy of the crown Rhopalosomatidae.

***Note 3*:** A distinct mesotrochantellus (char. **294**) is sporadically observed among extant Vespoidea and was almost impossible to score for fossils. Expression of a mesotrochantellus was consequently estimated to be a synapomorphy of the Rhopalosomatinae (CB50: 0.94), which we are uncertain about.

***Note 4*:** Expanded or lateromedially swollen or laminate female tarsomeres II–V (char. **320**) is considered to be a strong synapomorphy of the Rhopalosomatidae (Lohrmann *et al*. 2020). Unfortunately, both described species of †*Eorhopalosoma* are male, thus cannot be properly evaluated for this condition, perhaps leading to the lack of support for this condition in our ancestral state estimation analyses.

***Note 5*:** Presence of apicomedian lobate setae on the tarsi (char. **326**) was recovered as a synapomorphy of †*Eorhopalosoma* (CB50: 0.98).

***Note 6*:** Hind wing r-m is V- rather than U-shaped in †*Eorhopalosoma lohrmanni* and *Rhopalosoma* (char. **567**); see the formal character definition for more information.

***Note 7*:** Upcurved male gonopods (“parameres”) is a strong synapomorphy of the Rhopalosomatidae (Lohrmann *et al*. 2020) but was not scored here because of the emphasis on females. An upcurved female sting, and female pretarsus with “arolium and cuticular plate greatly enlarged and claw sensor elongated”, were also recorded as synapomorphies of the total Rhopalosomatidae by Lohrmann *et al*. (2020). These conditions were not operationalized here, and we cannot comment on the plausibility of the proposed polarity.

### ##. †*Prosphex*–Vespidae clade

*Clade comprising*: • †*Prosphex* genus group, • **Vespidae** Laicharting, 1781.

***Note*:** As this clade was reasonably supported in topology searches, we include those traits which were recovered as synapomorphies for the †*Prosphex* genus group plus Vespidae. However, we recognize that the support for the †*Prosphex* group is weak. See notes for the two constituent lineages for further details about certain reservations.

#### Synapomorphies

**14.** Head width ≥ 1 mm (CB50: 0.80). ***Polarity*:** Obviously, this is a variable across the Aculeata. Among extant Rhopalosomatidae, *Liosphex varius* and *Olixon* observed to have narrower crania.

**524.** Fore wing “submarginal cell 2” length and width subequal (CB50: 0.75). ***Polarity*:** Observed to be true for both †*Prosphex* group terminals, and all sampled Vespidae except for *Parischnogaster* and †*Curiosivespa zigrasi*.

### C.#. †*Prosphex* genus group

***Clade comprising*:** • †***Prosphex*** Grimaldi & Engel, 2019, Burmite fossil CASENT0844576 (Fig. S8).

***Note*:** The two species treated as the informal “†*Prosphex* genus group” here group together in 50CB tip-dating and ancestral state estimation analyses. The synapomorphies listed below simply reflect those features which draw the terminals together; the grouping should be considered highly provisional. Specimen CASENT0844576 is very similar morphologically to †*Alivespa* Wu *et al*., 2020 which was placed in †Priorvespinae. See also Vespidae below. Based on a number of features, we are concerned that †*Prosphex* may really be a species of Chrysidoidea, including: (1) antennal toruli close to anterior portion of head; (2) clypeus broadly projecting from cranium; and (3) upper metapleural areas and dorsolateral margins of propodeum with strong triangular processes. Further examination is necessary. Moreover, the two conditions supported as synapomorphies below are prone to homoplasy and should be considered weak evidence for the existence of this group.

#### Synapomorphies

**48.** Mandibular masticatory margin elongate, thus mandible triangular in form (CB50: 0.92). ***Polarity*:** The mandibles of various Vespidae are elongate, but the blade does not attain a triangular shape. Triangular mandibles were observed, however, in *Zethus* and *Mischocyttarus*.

**550.** Fore wing crossvein 1cu-a anterior junction “prefurcal”, *i.e.*, proximal to the split of Mf and Cuf (CB50: 0.90). ***Polarity*:** Almost unique among sampled Vespoidea, with the exception of Euparagiinae + Masarinae, †*Priorparagia*, and †*Symmorphus senex*.

### **C.B.** Family Vespidae Laicharting, 1781

***Clade comprising*:** • 9 extant subfamilies (see Carpenter 1981, 1988, 1989, Bank *et al*. 2017, Perrard *et al*. 2017, Piekarski *et al*. 2018, also Griffin 1939): **Stenogastrinae** Ashmead, 1902, •• **core Vespidae** clade: ••• <u>Masarinae–Euparagiinae clade</u> (**Euparagiinae** Ashmead, 1902, **Gayellinae** Bradley, 1922, **Masarinae** Latreille, 1802), <u>Eumeninae–Vespinae clade</u> (**Eumeninae** Leach, 1815, •••• **Zethinae** de Saussure, 1855, **Rhaphiglossinae** Ashmead, 1902, ••••• <u>Polistinae–Vespinae clade</u> (**Polistinae** Lepeletier de Saint-Fargeau, 1836, **Vespinae** Laicharting, 1781). • 2 extinct subfamilies: †**Priorvespinae** Carpenter & Rasnitsyn, 1990 (†*Alivespa* Wu *et al*., 2020, †*Priorvespa* Carpenter & Rasnitsyn, 1990), †**Protovespinae** Perrard & Carpenter in Perrard *et al*., 2017 (†*Protovespa* Perrard & Carpenter in Perrard *et al*., 2017). • Genus *incertae sedis* in family: †*Archaeovespa* Wu *et al*. 2021.

***Note 1*:** Wu *et al*. (2020) described †*Alivespa* from burmite and evaluated its phylogenetic position using a sample of 55 characters for five fossil Vespidae, seven extant Vespidae, and one terminal for Rhopalosomatidae and Tiphiidae. Given the very strong morphological similarity of the burmite specimen CASENT0844576 (“†*Prosphex* genus group” above and Fig. Fig. S8), to †*Alivespa*, the placement of the described fossil in Vespidae is uncertain and requires further study. †*Alivespa* was treated as a member of †Priorvespinae, thus we conservatively retain this placement.

***Note 2*:** Wu *et al*. (2021) also described the distinct genus †*Archaeovespa* from burmite, evaluating its phylogenetic position using an expanded dataset from Perrard *et al*. (2017), to which they added 15 characters, one extant species, and their three new species. Based on their analysis, the authors found that †*Archaeovespa* is close to Masarinae, adducing a single unreversed autapomorphy: hind wing cu-a “angled with apex of A” (their char. 82, from Perrard *et al*. 2017). Oddly, the extant species they figure does not appear to have this condition (their Fig. 6). The phylogenetic placement of this fossil remains to be shown convincingly, and is treated as *incertae sedis* here, as done by the authors. Note further uncertainty: The article was released online early, with ZooBank credentials, which should make 2020 the valid year of publication; however, the article and taxonomic action have not appeared in the ZooBank register as of 28 Dec 2020.

***Note 3*:** We list synapomorphies for the total Vespidae (*i.e.*, including †*Protovespa*), the crown Vespidae (Stenogastrinae + core Vespidae), core Vespidae (Euparagiinae–Polistinae), and the Eumeninae–Vespinae clade (represented in our phylogeny by Polistinae, Eumeninae, and Zethinae). The core Vespidae is of particular interest as recent phylogenomic works have revealed that Stenogastrinae is sister to the remainder of the crown group (Bank *et al*. 2017, Piekarski *et al*. 2018), overturning the morphology-based Euparagiinae-sistergroup hypothesis (Carpenter 1981, 1988). Recently, Jouault *et al*. (2021a) described a species attributed to †*Curiosivespa* from the Crato formation and discussed the placement of this genus in Euparagiinae.

#### Total Vespidae synapomorphies

**194.** In dorsal view, posterior margin of pronotum narrowly arcuate (CB50: 0.98). ***Polarity*:** Observed for all crown Vespidae plus †*Protovespa*, †*Curiosivespa striata*, and †*Symmorphus senex*; the pronotal arc was determined to be broader in †*Priorparagia*. The form of the pronotum in dorsal view could not be evaluated for †*Priorvespa*, †*Protovespa*, and other species of †*Curiosivespa*.

**451.** Fore wing Rf extending to or almost to wing apex (CB50: 0.86). ***Polarity*:** Among sampled fossils for which this trait could be evaluated, the derived condition was observed in †*Protovespa*, †*Priorvespa*, and at least one species of †*Curiosivespa*, as well as for the crown terminals *Parischnogaster* and *Mischocyttarus*. This implies multiple reversals within the crown Vespidae.

**494.** Fore wing Rsf between Rs+M and 2r-rs linear, not kinked or curved (CB50: 0.83). ***Polarity*:** Among sampled Vespidae, this abscissa of Rsf was only observed to be kinked or curved in *Euparagia*.

**\*.** Fore wing medial cell 1 (“discal cell 1”) proportionally elongate, with proximodistal length ≥ 4 x its anteroposterior width (char. **535**, CB50: 0.89), and also exceeding the maximum anteroposterior width of the wing (char. **536**, CB50: 0.97). ***Polarity*:** An elongate first medial cell observed for all sampled Vespidae. This condition was also approximated in the sampled crown Rhopalosomatidae, but with its length not always exceeding the maximum fore wing width.

**556.** Fore wing Cuf distal to 1m-cu directed proximally (CB50: 0.96). ***Polarity*:** Unique among Vespoidea, and present among all sampled Vespidae, extant and extinct.

#### Crown Vespidae synapomorphies

**94.** Anterior clypeal margin produced as median lobe (CB50: 0.80). ***Polarity*:** Could not be evaluated for †*Curiosivespa*. With reevaluation, may be found to be a synapomorphy of the Vespoidea.

**138.** Antennal toruli situated at or posterior to head midlength (CB50: 0.81). ***Polarity*:** Could not be evaluated for †*Curiosivespa*. With critical reappraisal, toruli at midlength may be found to be a vespoid synapomorphy.

**227.** Mesopectus with a distinctly deep pit subtending a flange at the base of the fore wing attachment (CB50: 0.72). ***Polarity*:** Support for this condition is equivocal at the crown Vespidae node, which probably reflects uncertainty from the fossils, as well as a potential reversal in Masarinae (as represented by *Pseudomasaris* in our dataset).

**339.** Posterior propodeal surface with a median longitudinal sulcus that extends from the metanotum to the propodeal foramen (CB50: 0.99). ***Polarity*:** Could not be evaluated for fossil Vespidae, thus the alternative hypothesis that this condition is a synapomorphy of the total clade Vespidae cannot be ruled out. See the character definition section for notes on the form of the propodeum in Rhopalosomatidae.

**460.** Pterostigma distinctly situated in apical third of fore wing (CB50: 0.96). ***Polarity*:** Observed in all sampled extant Vespidae, as well as for †*Curiosivespa* and †*Priorparagia*; the pterostigma is more proximally situated in †*Protovespa*.

**486.** Fore wing Mf1 long to extremely long, being ≥ 4 x the length of Rsf1 (CB50: 0.86). ***Polarity*:** Observed among all sampled Vespidae except †*Protovespa*, which was recovered as sister to the crown group, and in †*Symmorphus senex*, which does not have sufficient preservational fidelity to score with confidence.

#### Core Vespidae synapomorphies

**195.** Lateral portion of pronotum, *i.e.*, that region which overlaps the mesoscutum laterally, dorsoventrally expanded, visible as lateromedially broad shoulders in dorsal view (CB50: 0.96). ***Polarity*:** Among crown Vespidae, observed for all sampled taxa except for Stenogastrinae, which has a dorsoventrally narrow lateral pronotal region, appearing as a long, narrow digitate process in lateral and dorsal views (autapomorphy, char. **196**).

**307.** Anterior apical areas of the meso- and metatibiae with an even to messy row of short, stout chaetae (“traction setae”) near the anteroapical tibial margins (CB50: 0.85). ***Polarity*:** Observed to be absent in *Parischnogaster* (Stenogastrinae) and present in all other sampled extant Vespidae. Could not be scored for fossil taxa. This condition has not been previously recognized in the literature and has uncertain functional consequences.

**562.** Fore wing C present (CB50: 0.71). ***Polarity*:** The low support for this condition as a synapomorphy of the core Vespidae reflects fossil uncertainty, and possibly homoplastic presence of C in CASENT0844576. Observed for all sampled extant Vespidae except *Parischnogaster*.

#### Eumeninae–Vespinae clade synapomorphies

**214.** Parategula present, *i.e.*, posterior apex of parascutal carina expanded as a lobate, laminate, or subdigitate process (CB50: 0.97). ***Polarity*:** Among sampled fossils, the parategular expansion has been observed to be absent in †*Protovespa*; among sampled extant terminals, this development is absent in *Parischnogaster*, *Euparagia*, and *Pseudomasaris*. Although the Polistinae are usually considered to lack a parategula, the posterior apex of the parascutal carina was observed in our study to be expanded to some degree in *Mischocyttarus*, thus was scored as present. Parategular absence in Polistinae and Vespinae should probably be considered a secondary reduction, or parallel reductions once the form of the parascutal carina is subjected to future scrutiny.

**444.** First two metasomal segments comparatively long, together being distinctly longer than remainder of metasoma (CB50: 0.96). ***Polarity*:** These segments were observed to be shorter in other sampled crown Vespidae, and in †*Priorvespa*, †*Curiosivespa*, and †*Symmorphus senex*.

**548.** Fore wing crossvein 1cu-a distinctly postfurcal (CB50: 0.92). ***Polarity*:** Among sampled Vespidae, observed for Eumeninae, Zethinae, and Polistinae; variable among fossils. Independently derived with respect to Rhopalosomatidae.

**554.** Fore wing Cuf1 distinctly angled posteroapically from M+Cu, and about as long as 1cu-a (CB50: 0.97). ***Polarity*:** Unique among all sampled Vespoidea, and virtually unique among other Aculeata.

***Note 1*:** Character 43 (ocellar deformation) was recovered as a synapomorphy of the crown Vespidae, but after *post hoc* trait evaluation, this should be considered a mischaracterization.

***Note 2*:** Presence of a longitudinal crease or plait (char. **456**) is supported as a synapomorphy of the Zethinae–Vespinae clade (CB50: 0.97).

***Note 3*:** Fore wing with a sinuate crossvein 3rs-m (char. **518**) was reconstructed as a synapomorphy of the core Vespidae (*i.e.*, crown taxa with the exclusion of Stenogastrinae) (CB50: 0.77). However, understanding the ancestral condition is complicated by the general absence of this crossvein in the Rhopalosomatidae, Dryinoidea, and Chrysidoidea.

***Note 4*:** Fore wing crossvein 1m-cu distinctly curved or sinuate (char. **531**) was reconstructed as a synapomorphy for the Zethinae–Vespinae clade (CB50: 0.80), but we note that this condition was also observed in both *Euparagia* and *Pseudomasaris*, as well as most fossils (†*Priorvespa*, †*Priorparagia*, some †*Curiosivespa*).

### C’. Superfamily Pompiloidea Latreille, 1804 *sensu novum*

*Clade comprising*: • **Sierolomorphidae** Krombein, 1951, • clade **Tiphiopompiloides**.

***Note*:** No unequivocal synapomorphies were recovered for the Pompiloidea *sensu novum*, although several conditions are supported as stabilized among the first few internal nodes (see below). Complex patterns of parallelism among the pompiloidine clades were detected. There is equivocal support for a distinct angle between the radicle and scape as a shared apomorphy (char. **149**, CB50: 0.53), with reversal to the linear condition in Tiphiidae (CB50: 0.90) and Pompiliformes (CB50: 0.70). It is possible that a distinct angle may be synapomorphic for the Vespoidea as this condition is supported for Scolioidea (CB50: 0.83) and Apoidea (CB50: 0.90), but considerable uncertainty remains. Refined characterization or quantification will contribute to the resolution of the pattern of radicle and scape transformation.

### **D.A.** Family Sierolomorphidae Krombein, 1951

***Clade comprising*:** • †*Loreisomorpha* Rasnitsyn, 2000, *Proscleroderma* Kieffer, 1905, *Sierolomorpha* Ashmead, 1903, Burmite genus CASENT0844589 (Fig. S9).

***Note 1*:** See Argaman (1990) for inclusion of *Proscleroderma* in Sierolomorphidae, Lelej & Mokrousov (2015) for a catalog of species in the family, and Mokrousov *et al*. (2018) for the most recent addition to the group.

***Note 2*:** The uncertain fossil CASENT0844580 (Fig. S9) clustered with Sierolomorphidae in the ancestral state estimation topology searches. To understand how this occurred, estimated synapomorphies for that node are provided (“clade 1”), in addition to those for the total Sierolomorphidae. We strongly suggest against formal interpretation of the provisional CASENT0844580 + Sierolomorphidae clade given the highly variable placement of this fossil, and the observation that all characters which draw the group together are prone to homoplasy.

#### Clade 1 synapomorphies

**143.** Antennal toruli directed toward the mandibles (CB50: 0.94). ***Polarity*:** Among Pompiloidea *sensu novum*, supported as an independent derivation of Mutilliformes (CB50: 0.90). Also observed in *Chyphotes*, *Aelurus*, *Myzinum,* and †*Bryopompilus*.

**187.** Pronotum with a lateral furrow which can receive the fore leg when this appendage is folded dorsomedially (CB50: 0.83). ***Polarity*:** Among Pompiloidea *sensu novum*, also observed in *Tiphia* and †*Thanatotiphia*.

**488.** Fore wing Mf1 distinctly curved (CB50: 0.87). ***Polarity*:** Distinct curvature of Mf1 was also observed in Tiphiiformes, for which the condition was equivocally reconstructed as a synapomorphy.

**550.** Fore wing crossvein 1cu-a anterior junction prefurcal, *i.e.*, proximal to the split of M+Cu (CB50: 0.88). ***Polarity*:** Also observed in some Tiphiiformes, †*Bryopompilus*, and one species of †*Burmusculus*.

#### Total Sierolomorphidae clade synapomorphies

**14.** Head width ≥ 1 mm (CB50: 0.79). ***Polarity*:** Among Pompiloidea *sensu novum*, large body size—as indicated by the head width proxy—is supported as independently derived for Chyphotidae (CB50: 0.96), Mutilliformes (CB50: 0.82), with reversals.

**21.** Occipital carina absent (CB50: 0.73). ***Polarity*:** The occipital carina is also absent in *Sapyga* and *Dasymutilla* among sampled Pompiloidea *sensu novum*. Uncertain for †*Loreisomorpha*.

**64.** Mandible bidentate (CB50: 0.79). ***Polarity*:** Uncertain for †*Loreisomorpha*. Supported as an independent derivation in Tiphiiformes (CB50: 0.92) and Pompiliformes (CB50: 0.94).

**94.** Anterior clypeal margin with a distinct median lobate process (CB50: 0.78). ***Polarity*:** Among sampled Pompiloidea *sensu novum*, also observed in *Tiphia* and crown Sapygidae.

**178.** Pronotum elongate and muscular, more-or-less box-shaped (CB50: 0.94). ***Polarity*:** Strongly supported as a synapomorphy of total Sierolomorphidae, and convergently derived in *Olixon* and *Aelurus* among Vespoides. Observed in both stem species of the family.

**213.** Parascutal carina absent (CB50: 0.77). ***Polarity*:** Could not be evaluated for †*Loreisomorpha*. Convergently lost for total Tiphiiformes (CB50: 0.81) and crown Sapygidae (CB50: 0.97); reconstructed as trending toward loss independently between the Pompiliformes and Mutilliformes.

***Note*:** Only one condition strongly draws the stem Sierolomorphidae together, that of 2m-cu being close to the distal fore wing apex, separated by less than one of its lengths (char. **541**, CB50: 0.97). This is Also observed in Thynnidae and crown Sapygidae.

### **D.A** ’. Clade Tiphiopompiloides

*Clade comprising*: • **Tiphiiformes**, • **core Pompiloidea clade**.

***Note*:** No unequivocal synapomorphies were detected for this node.

### **D.A.1.** Clade Tiphiiformes

*Clade comprising*: • **Tiphiidae** Leach, 1815, • **clade Thynnimorpha**. • *Incertae sedis* in clade: †*Architiphia* Darling, 1990.

***Note*:** †*Architiphia* is conservatively retained in Tiphiiformes, although its placement is variable.

#### Synapomorphies

**64.** Mandible bidentate (CB50: 0.92). ***Polarity*:** Uncertain for †*Loreisomorpha*. Supported as an independent derivation in total Sierolomorphidae (CB50: 0.79), and Pompiliformes (CB50: 0.94).

**213.** Parascutal carina absent (CB50: 0.81). ***Polarity*:** Convergently lost for total Sierolomorphidae (CB50: 0.77) and crown Sapygidae (CB50: 0.97); reconstructed as trending toward loss independently between the Pompiliformes and Mutilliformes.

**263.** Mesocoxae wide set (CB50: 0.74). ***Polarity*:** Among sampled Tiphiiformes, the mesocoxae were close-set in *Methocha*; support for the wide-set condition stabilizes among the internal tiphioid nodes. Some Mutillidae were also observed to have wide set mesocoxae (*Dasymutilla*, *Rhopalomutilla*).

**284.** Female protarsus with row of psammochaetae, or broad, flat to grooved “fossorial setae” (CB50: 0.73). ***Polarity*:** Observed in all sampled extant Tiphiiformes with the exception of *Methocha*; also observed in *Dasymutilla* and *Rhopalomutilla*.

**288.** Proximal articulatory surfaces of meso- and metacoxae distinctly narrowed relative to the remainder of the segments (CB50: 0.91). ***Polarity*:** Also observed in crown Mutillidae, among Pompiloidea *sensu novum*.

**292.** Metacoxae distinctly bent between the proximal articulation and distal portion (CB50: 0.89). ***Polarity*:** Among Pompiloidea *sensu novum*, also observed in crown Mutillidae.

**488.** Fore wing Mf1 distinctly curved (CB50: 0.70). ***Polarity*:** Observed among all sampled Tiphiiformes with the exception of *Myzinum* and †*Thanatotiphia*.

***Note*:** A distinctly curved first free medial abscissa was equivocally reconstructed as a synapomorphy of Tiphiiformes (CB50: 0.69) but was observed in all sampled species within the group with the exception of *Myzinum*. Expanded taxon sampling may resolve the polarity of this condition at the tiphioid node.

### **D.B.** Family Tiphiidae Leach, 1815

***Clade comprising*:** • **Brachycistidinae** Malloch, 1926, • **Tiphiinae** Leach, 1815. ***Incertae sedis*** in family: †**Thanatotiphiinae** Engel *et al*., 2009 (†*Thanatotiphia* Engel *et al*., 2009).

***Note*:** The placement of †*Thanatotiphia* is unstable, but the “unciform” abdominal sternum VII of the male does strongly suggest relationship with the superfamily.

#### Synapomorphies

**34.** Compound eyes converging toward mandibles (CB50: 0.90). ***Polarity*:** Expanded taxon sampling may refine the estimates for this condition. Moderately supported for the Pompiliformes (see below).

**87.** Clypeus not projecting anteriorly from cranium (CB50: 0.80). ***Polarity*:** Supported as independently derived in Chyphotidae and crown Mutillidae.

**139.** Clypeus extending posteriorly between antennal toruli (CB50: 0.90). ***Polarity*:** Strongly supported as independently derived in Thynnidae.

**234.** Mesopectus posteriorly scrobiculate, *i.e.*, with a broad dorsoventrally oriented groove which can receive the mesoleg when this appendage is flexed dorsomedially (CB50: 0.90). ***Polarity*:** Supported as independently derived in Thynnidae.

**251.** Medial lobes of ventral mesopectal surface expanded posteriorly as large, distinct plates which at least partially overlap the mesocoxae (CB50: 0.50). ***Polarity*:** Among sampled Pompiloidea *sensu novum*, also observed in *Myzinum* and *Aelurus* (core Thynnidae clade).

**\*** Mesopectus ventrally scrobiculate, accommodating mesocoxa when leg flexed dorsoventrally (char. **254**, CB50: 1.0), the scrobe being deep and associated with relatively elongate mesocoxae (char. **255**, CB50: 1.0). ***Polarity*:** A ventrally scrobiculate mesopectus was also observed in Thynnidae but not in Chyphotidae; the depth of the thynnid scrobe is comparatively shallow.

**256.** Mesocoxae set in a ventral thoracic socket, with the sternal area nearly reaching the ventralmost surface of the coxa (CB50: 0.90). ***Polarity*:** Also observed in the core Thynnidae and Mutillidae clades.

**273.** Metathoracic coxal cavities closed in lateral view (see definition for further detail) (CB50: 1.0). ***Polarity*:** Supported as independently derived in Chyphotidae (*Chyphotes*) and Thynnidae (*Myzinum*, *Methocha*). Also observed in crown Mutillidae.

**304.** Meso- and metatibiae clavate, being grossly swollen (CB50: 1.0). ***Polarity*:** Also observed in the core Thynnidae clade (as represented by *Aelurus* and *Myzinum*).

**304.** Meso- and metatibiae clavate, being grossly swollen (CB50: 1.0). ***Polarity*:** Also observed in core Thynnidae.

**310.** Mesotibia with a single ventroapical spur (CB50: 0.90). ***Polarity*:** Among sampled Pompiloidea *sensu novum*, the condition of having only one mesotibial spur was observed in *Methocha*. All other sampled species had a 2,2 tibial spur formula (*i.e.*, two meso- and metatibial spurs, respectively), including fossils.

**314.** Meso- and/or metatibial spurs extremely elongate, with the posterior (or sole) spur length ≥ ½ basitarsal length (CB50: 0.90). ***Polarity*:** Among sampled Pompiloidea *sensu novum*, also observed for core Thynnidae.

**370.** Anterior foramen of metasomal petiole (abdominal segment II) with a distinct anterior neck, whether short (subsessile) or long (pedunculate) (CB50: 0.90). ***Polarity*:** Among Pompiloidea *sensu novum*, the pedunculate condition of the petiole is supported as independently derived in Tiphiidae, Chyphotidae, Methochinae, and core Mutillidae.

**458.** Pterostigma grossly enlarged (CB50: 0.90). ***Polarity*:** Also observed in *Methocha*, *Chyphotes*, and the fossil CASENT0844580.

**\*** Fore wing 2rrs cell (marginal 1) not apically pointed or narrowly rounded (char. **475**, CB50: 0.80), rather with its apex strongly “downcurved” (see definition for further detail, char. **479**, CB50: 0.90). ***Polarity*:** The pointed or narrowly rounded condition was observed for most fossils of the Pompiloidea *sensu novum*, and for *Sierolomorpha*, *Methocha*, *Aelurus*, *Chyphotes*, *Sapyga*, *Myrmosa*, and *Rhopalomutilla*.

**483.** Fore wing Rsf1 and Mf1 meeting at a distinct angle (CB50: 0.80). ***Polarity*:** Also observed in *Methocha*, *Myrmosa*, *Smicromyrmilla*, the stem mutillid CASENT0844578, and CASENT0844580.

**531.** Fore wing crossvein 1m-cu distinctly curved or sinuate (CB50: 0.90). ***Polarity*:** Also observed in *Sierolomorpha*, Chyphotidae, *Aelurus*, †*Architiphia*, one species of †*Burmusculus*, and †*Cretofedtschenkia*.

**535.** Fore wing medial cell 1 (discal 1) proportionally elongate, with proximodistal length ∼ 4 x its anteroposterior width (CB50: 0.90). ***Polarity*:** Also observed in Chyphotidae, *Myzinum*, *Pepsis*, and the Mutilliformes.

**539.** Fore wing crossvein 2m-cu curved or sinuate (CB50: 0.80). ***Polarity*:** Supported as convergently derived in Thynnidae and crown Sapygidae.

### **D.B** ’. Clade Thynnimorpha

*Clade comprising*: • **Chyphotidae** Ashmead, 1896, • **Thynnidae** Erichson, 1841.

#### Synapomorphies

**2.** Apterous females present (CB50: 0.80). ***Polarity*:** Female aptery is supported as having originated at least three times independently in the Pompiloidea *sensu novum*: at least once in Tiphiidae, once in the ancestor of the Thynnimorpha, and in the ancestor of the crown Mutillidae.

**36.** Female compound eyes reduced in size, being < ½ head length (CB50: 0.80). ***Polarity*:** Supported as independently derived in Tiphiidae (Brachycistidinae), with a reversal inferred for *Myzinum* within the Thynnimorpha. Although reduction of the compound eyes is associated with the derivation of apterous females, the sampled Mutillidae were observed to be variable for this condition.

**300.** Meso- and/or metatibiae with strong, scattered chaetae (“spurs” or “traction setae”) (CB50: 0.90). ***Polarity*:** Support for expression of these chaetae is equivocal for the Thynnimorpha and Tiphiidae, as such chaetae were observed to be present in *Tiphia* and absent in *Brachycistis*. These were also observed as present for *Pepsis*, *Fedtschenkia*, and *Myrmosa*, with each occurrence strongly supported as independent.

**431.** Male abdominal sternum VII “unciform”, *i.e.*, in the form of an upturned stinger-like process (CB50: 0.90). ***Polarity*:** Observed in †*Thanatotiphia*, *Brachycistis*, and all sampled Thynnimorpha except *Aelurus*.

### **D.C.** Family Chyphotidae Ashmead, 1896a

***Clade comprising*:** • **Chyphotinae** Ashmead, 1896a (•• *Chyphotes* Blake, 1886), • **Typhoctinae** Schuster, 1949 (•• **Eotillini** Schuster, 1949 [*Eotilla* Schuster, 1949, *Prototilla* Schuster, 1949], •• **Typhoctini** Schuster, 1949 [*Typhoctes* Ashmead, 1899, *Typhoctoides* Brothers, 1974b]).

#### Synapomorphies

**14.** Head width ≥ 1 mm (CB50: 0.96). ***Polarity*:** Supported as independently derived for total Sierolomorphidae (CB50: 0.79), Mutilliformes (CB50: 0.82), with reversals.

**40.** Ocelli repressed in females (CB50: 0.98). ***Polarity*:** Expression of the ocelli is retained in some Tiphiidae and some Thynnidae. Repression is independently derived in Mutillidae.

**87.** Clypeus not projecting anteriorly from cranium (CB50: 0.97). ***Polarity*:** Supported as independently derived in Tiphiidae and crown Mutillidae.

**272.** Mesothoracic coxal cavity closed in lateral view (see definition for further detail) (CB50: 1.0). ***Polarity*:** Supported as independently derived in Thynnidae, where the closed condition was observed in *Methocha* and *Myzinum*, and the open condition in *Aelurus*. Mesocoxal cavity closure was also observed in Mutillidae.

**294.** Mesotrochantellus distinct and well-developed (CB50: 0.96). ***Polarity*:** Also observed in *Brachycistis* and Pompiloidea.

**370.** Anterior foramen of metasomal petiole (abdominal segment II) with a distinct anterior neck, whether short (subsessile) or long (pedunculate) (CB50: 0.97). ***Polarity*:** Among Pompiloidea *sensu novum*, the pedunculate condition of the petiole is supported as independently derived in Tiphiidae, Chyphotidae, Methochinae, and core Mutillidae.

**392.** Abdominal segment III (metasomal II) without defined laterotergite (CB50: 0.98). ***Polarity*:** The laterotergite of abdominal segment III is supported as independently lost in *Tiphia* and some Mutillidae.

**403.** Abdominal segment III anterior articulatory surfaces narrowed relative to remainder of segment (CB50: 0.98). ***Polarity*:** Supported as independently derived in *Tiphia* and *Dasymutilla*, among sampled Pompiloidea *sensu novum*.

**512.** Fore wing crossvein 2rs-m sinuate (CB50: 0.98). ***Polarity*:** Also observed in *Brachycistis*, *Pepsis*, †*Architiphia*, one species of †*Burmusculus*, and †*Cretofedtschenkia*.

**520.** Fore wing crossvein 3rs-m situated close to 2rs-m such that cell 3rm (submarginal 3) anteroposteriorly wider than proximodistally long (CB50: 0.96). ***Polarity*:** Also observed in *Myzinum*, *Pepsis*, and various fossil Pompiloidea *sensu novum*.

**521.** Fore wing cell 1rrs (submarginal 1) prestigmal length > ½ cell length (CB50: 0.95). ***Polarity*:** Also observed in *Myzinum*, *Pepsis*, crown Sapygidae, core Mutillidae, and †*Cretofedtschenkia*.

**531.** Fore wing crossvein 1m-cu distinctly curved or sinuate (CB50: 0.97). ***Polarity*:** Also observed in *Sierolomorpha*, Tiphiidae, *Aelurus*, †*Architiphia*, one species of †*Burmusculus*, and †*Cretofedtschenkia*.

**535.** Fore wing medial cell 1 (discal 1) proportionally elongate, with proximodistal length ∼ 4 x its anteroposterior width (CB50: 0.99). ***Polarity*:** Also observed in Tiphiidae, *Myzinum*, *Pepsis*, and the Mutilliformes.

**561.** Hind wing basal hamuli present (CB50: 0.97). ***Polarity*:** Observed in some Mutillidae and †*Burmusculus*.

**571.** Hind wing crossvein cu-a interstitial or prefurcal, *i.e.*, at or proximal to the split of M+Cu (CB50: 0.96). ***Polarity*:** Also observed in *Sierolomorpha*, *Myzinum*, *Myrmosa*, and *Rhopalomutilla*.

**572.** Hind wing distal elongation of sectoral cell (basal cell) absent (CB50: 0.97). ***Polarity*:** Supported as a synapomorphy of core Mutillidae and also observed in *Sierolomorpha*.

### **D.D.** Family Thynnidae Erichson, 1841

*Clade comprising*: • **Anthoboscinae** Turner, 1912, • **Diamminae** Turner, 1907, • **Methochinae**

André, 1899, • **Myzininae** Ashmead, 1903a, • **Thynninae** Erichson, 1841.

***Note*:** Because taxon sampling within the Thynnidae was limited to Methochinae, Myzininae, and Thynninae in the present study, we recommend the use of caution when interpreting the listed synapomorphies. However, because several conditions were found to be supported as synapomorphies for *Aelurus* and *Myzinum*, these are listed below (“core Thynnidae clade”). This Thynninae–Myzininae clade was supported by Pilgrim *et al*. (2008); character polarities would be positively informed by inclusion of Anthoboscinae and Diamminae, which were found to be outsize of the core Thynnidae, but closer than Methochinae, the sister to the remainder of the family.

#### Thynnidae synapomorphies

**139.** Clypeus extending posteriorly between antennal toruli (CB50: 0.93). ***Polarity*:** Strongly supported as independently derived in Tiphiidae.

**234.** Mesopectus posteriorly scrobiculate, *i.e.*, with a broad dorsoventrally oriented groove which can receive the mesoleg when this appendage is flexed dorsomedially (CB50: 0.93). ***Polarity*:** Supported as independently derived in Tiphiidae. Not observed in Chyphotidae.

**244.** Lower metapleural area, *i.e.*, that region ventrad the endophragmal pit, strongly reduced, being present as a dorsoventrally narrow but anteroposteriorly long triangle ventrad the propodeum (CB50: 0.96). ***Polarity*:** Supported as an autapomorphy of Thynnidae among Pompiloidea *sensu novum*.

**254.** Mesopectus ventrally scrobiculate, accommodating mesocoxa when leg flexed dorsomedially (CB50: 0.99). ***Polarity*:** Also observed in Tiphiidae, but the tiphiid scrobe is comparatively deep.

**410.** Anterior articulatory surfaces of abdominal segment III (metasomal II) supraaxial, *i.e.*, situated above segment midheight (CB50: 0.95). ***Polarity*:** Supported as independently derived in *Sierolomorpha*, *Tiphia*, and *Typhoctes*. Evaluation of the skeletomuscular anatomy of the anterior metasoma of Pompiloidea *sensu novum* will inform the evolutionary patterns of this condition.

**516.** Fore wing crossvein 2rs-m joining Rs distal to 2r-rs by greater than one of its lengths (CB50: 0.81). ***Polarity*:** Also observed in *Pepsis*, crown Sapygidae, †*Cretofedtschenkia*, †*Architiphia*, some †*Burmusculus*, and the stem mutillid fossil (CASENT0844578).

**539.** Fore wing crossvein 2m-cu curved or sinuate (CB50: 0.82). ***Polarity*:** Supported as convergently derived in Tiphiidae and crown Sapygidae.

#### Core Thynnidae synapomorphies

**143.** Antennal toruli which are directed toward the mandibles (CB50: 0.94). ***Polarity*:** Also observed for *Chyphotes*.

**256.** Mesocoxae set in ventral thoracic sockets, with the sternal area nearly reaching the ventralmost surface of the coxae (CB50: 0.95). ***Polarity*:** Also observed in Tiphiidae and core Mutillidae.

**296.** Meso- and metafemora strongly compressed and expanded with scrobes to receive their respective tibiae (see definition for further detail) (CB50: 0.97). ***Polarity*:** Unique among sampled Pompiloidea *sensu novum*.

**304.** Meso- and metatibiae clavate, being grossly swollen (CB50: 0.97). ***Polarity*:** Also observed in Tiphiidae.

**314.** Meso- and/or metatibial spurs extremely elongate, with the posterior (or sole) spur length ≥ ½ basitarsal length (CB50: 0.96). ***Polarity*:** Among sampled Pompiloidea *sensu novum*, also observed in Tiphiidae.

**409.** Abdominal tergum VII of female and VIII of male modified as “pygidial plate” (CB50: 0.91). ***Polarity*:** Among sampled Pompiloidea *sensu novum*, also observed in *Tiphia* and *Rhopalomutilla*.

### **D.A.2.** Core Pompiloidea clade

*Clade comprising*: • Clade **Pompiliformes**, • Clade **Mutilliformes**.

#### Synapomorphies

**227.** Longitudinal mesopectal sulcus present (CB50: 0.81). ***Polarity*:** Among Pompiloidea *sensu novum*, observed in Pompilidae, †*Bryopompilus*, and Mutillidae (CASENT0844578). Supported as reversed to absence in crown Mutillidae. Because the placement of fossils remains uncertain, this should be interpreted with caution. The distinct form observed in Pompilidae was not scored.

**294.** Mesotrochantellus distinct and well-developed (CB50: 0.71). ***Polarity*:** Supported as reduced in *Fedtschenkia* and *Smicromyrmilla*; observed for †*Burmusculus*, *Pepsis*, and most Mutillidae.

**400.** Abdominal tergum III (metasomal II) without transverse sulcus dividing the anterior articulatory surface (presclerite) from the posterior surface (postsclerite) (CB50: 0.86). ***Polarity*:** The ancestral state reconstruction analysis strongly supports reversal to the present condition in crown Mutillidae. Among fossils for which this condition could be evaluated, it was only observed for †*Cretofedtschenkia*.

**453.** Fore wing Rsf extending or nearly extending to distalmost wing margin (CB50: 0.89). ***Polarity*:** Among Pompiloidea *sensu novum*, also observed in *Methocha* and *Aelurus*. Within Pompiloidea, observed in *Sapyga, Myrmosa*, †*Bryopompilus*, †*Burmusculus*, and †*Cretosapyga*. Clearly, the fossils have informed the reconstruction at this node.

### **D.A.2.1.** Clade Pompiliformes

***Clade comprising*:** • †**Burmusculidae** Zhang *et al*., 2018 (†*Burmusculus* Zhang & Rasnitsyn, 2018), **Pompilidae** Latreille, 1804 (**Ceropalinae** Radoszkowski, 1888, **Ctenocerinae** Arnold, 1934, **Notocyphinae** Fox, 1895, **Pepsinae** Lepeletier de Saint-Fargeau, 1845, **Pompilinae** Latreille, 1804).

***Note*:** Because we sampled only one extant terminal in Pompilidae, we cannot provide synapomorphies for the family. For recent phylogenies of Pompilidae, see Pitts *et al*. (2006), Waichert *et al*. (2015), and Rodriguez *et al*. (2017). Also note that †*Burmusculus* has been suggested to be a possible junior synonym of †*Taimyrisphex* Evans, 1973 by Rasnitsyn *et al*. (2020) due to the wing venation and broad metapostnotum.

#### Synapomorphies

**64.** Mandible bidentate (CB50: 0.95). ***Polarity*:** Uncertain for †*Loreisomorpha*. Supported as an independent derivation in total Sierolomorphidae (CB50: 0.79), and Tiphiiformes (CB50: 0.92).

**242.** Upper metapleural area, *i.e.*, that region dorsad the endophragmal pit, expanded posteriorly (CB50: 0.98). ***Polarity*:** Supported as an autapomorphy of Pompiliformes among Pompiloidea *sensu novum*.

***Note*:** There is moderate support for compound eyes converging toward mandibles (char. **34**, CB50: 0.81) at the Pompiliformes node. However, among sampled taxa this was only observed for some †*Burmusculus*, so this should be considered uncertain.

### **D.A.2.2.** Clade Mutilliformes

***Clade comprising*:** • **†Bryopompilidae** Rodriguez *et al*., 2016 (†*Bryopompilus* Engel & Grimaldi, 2006b), • **Mutillidae** Latreille, 1802, • **Sapygidae** Latreille, 1810.

***Note*:** Synapomorphies are provided for the total and crown Mutilliformes, *i.e.*, including or excluding †*Bryopompilus*.

#### Total Mutilliformes Synapomorphies

**14.** Head width ≥ 1 mm (CB50: 0.82). ***Polarity*:** Supported as independently derived for total Sierolomorphidae (CB50: 0.79), Chyphotidae (CB50: 0.96), with reversals.

**65.** Mandible tridentate (CB50: 0.92). ***Polarity*:** Tridentate mandibles represent an autapomorphy among sampled Pompiloidea *sensu novum*. Among fossil pompiloidines which could be sampled, observed for †*Bryopompilus* (mandibles more-dentate), †*Cretosapyga*, and the stem mutillid (CASENT0844578); other fossil taxa for which this could be evaluated had two or fewer teeth. *Dasymutilla* with derived unidentate mandibles.

**143.** Antennal toruli directed toward mandibles (CB50: 0.90). ***Polarity*:** Independently derived in Sierolomorphidae and some Thynnoidea.

**518.** Fore wing crossvein 3rs-m sinuate (CB50: 0.93). ***Polarity*:** Also observed in *Tiphia*, *Myzinum*, *Aelurus*, and most Tiphiopompiloides fossils with the exception of some †*Burmusculus*. 3rs-m is linear in the examined *Myrmosa*.

**535.** Fore wing medial cell 1 (discal 1) proportionally elongate, with proximodistal length ∼ 4 x its anteroposterior width (CB50: 0.83). ***Polarity*:** Also observed in Tiphiidae, Chyphotidae, *Myzinum*, and *Pepsis*.

**555.** Fore wing Cu1 and Cu2 (anterior and posterior branches of Cuf) diverging at approximately a right angle (CB50: 0.83). ***Polarity*:** Also observed in *Brachycistis*, one species of †*Burmusculus*, and *Pepsis*. Supported as modified to another conformation in the core Mutillidae.

#### Crown Mutilliformes Synapomorphies

**150.** Radicle and scape nearly perpendicular (CB50: 0.84). ***Polarity*:** Uncertain for †*Cretosapyga* and also observed for *Aelurus*, but otherwise strongly supported as a synapomorphy of crown Mutilliformes.

**288.** Proximal articulatory surfaces of meso- and metacoxae distinctly narrowed relative to the remainder of the segment (CB50: 0.91). ***Polarity*:** Observed for all sampled extant Mutillidae, uncertain for CASENT0844578; independently derived in Tiphiiformes.

### D.G. Family Mutillidae Latreille, 1802

*Clade comprising*: • **Myrmosinae** Fox, 1895, • **core Mutillidae** clade (•• **Dasylabrinae** Brothers & Lelej, 2017, **Mutillinae** Latreille, 1802, **Myrmillinae** Bischoff, 1920, **Rhopalomutillinae** Schuster, 1949, **Pseudophotopsinae** Bischoff, 1920, **Sphaeropthalminae** Schuster, 1949, **Ticoplinae** Nagy, 1970).

***Note*:** Synapomorphies are listed for the total and crown clade Mutillidae, *i.e.*, including or excluding the burmite fossil CASENT0844578 (Fig. S11), as well as for the core Mutillidae (*i.e.*, excluding Myrmosinae).

#### Total Mutillidae synapomorphies

**151.** Radicle obscured by flangelike expansion of scape (CB50: 0.98). ***Polarity*:** An apparent autapomorphy of Mutillidae among sampled Hymenoptera.

**282.** Protibial calcar unifid (CB50: 0.99). ***Polarity*:** Observed in CASENT0844578. Uncertain for most fossil taxa, but unique among sampled extant Pompiloidea *sensu novum*.

#### Crown Mutillidae synapomorphies

**2.** Apterous females present (CB50: 0.89). ***Polarity*:** The condition of the female represented by CASENT0844578 is unknown. For this reason, female aptery is more strongly supported for the crown rather than the total node (CB50: 0.59).

**40.** Ocelli repressed in females (CB50: 0.97). ***Polarity*:** Ocellar repression is independently derived in some Thynnoidea and is uncertain for CASENT0844578.

**87.** Clypeus not projecting anteriorly from cranium (CB50: 0.95). ***Polarity*:** Supported as independently derived in Tiphiidae and Chyphotidae.

**227.** Mesopectus without longitudinal sulcus (CB50: 0.65). ***Polarity*:** Supported as a reversal from the present state (see Pompiloidea polarity comment above), with support stabilizing for core Mutillidae (CB50: 0.94).

**272.** Mesothoracic coxal cavity closed in lateral view (CB50: 0.73). ***Polarity*:** Although the support for this condition at the crown Mutillidae node is equivocal, all extant species were observed to have closed mesocoxal cavities.

**273.** Metathoracic coxal cavity closed in lateral view (CB50: 0.74). ***Polarity*:** As for the mesocoxal cavities, closure of the metacoxal cavities was observed in all sampled crown Mutillidae.

**292.** Metacoxae distinctly bent between proximal articulation and distal portion (CB50: 0.87). ***Polarity*:** Observed in all sampled Tiphiiformes and crown Mutillidae.

**324.** Pretarsal claws unidentate (CB50: 0.60). ***Polarity*:** Observed in all sampled extant Mutillidae; CASENT0844578 with dentate claws. The unidentate condition is supported as independently derived at least twice in Tiphiiformes (*Brachycistis* and *Aelurus*).

**351.** Propodeal spiracle vestibulate, *i.e.*, with an anterior hood-like projection (CB50: 0.75). ***Polarity*:** Unique among sampled Pompiloidea *sensu novum*; supported as lost in *Rhopalomutilla*.

**400.** Abdominal tergum III (metasomal II) without transverse sulcus dividing the anterior articulatory surface (presclerite) from the posterior surface (postsclerite) (CB50: 0.86). ***Polarity*:** Presence of the “cinctus” or transverse sulcus is strongly supported as a reversal within the Pompiloidea.

**423.** Abdominal tergum IV (metasomal III) with anteromedian stridulitrum (CB50: 0.75). ***Polarity*:** Observed in all sampled extant Mutillidae except for *Rhopalomutilla*. Unique among Pompiloidea *sensu novum*.

#### Core Mutillidae synapomorphies

**154.** Scape elongate, with length ≥ 4 x width (CB50: 0.85). ***Polarity*:** Among Pompiloidea *sensu novum*, also observed in *Myzinum*.

**165.** Promesonotal articulation immobile due to fusion, observed in females (CB50: 0.94). ***Polarity*:** Unique among Pompiloidea *sensu novum*.

**256.** Mesocoxae set in a ventral thoracic socket, with the sternal area nearly reaching the ventralmost surface of the coxa (CB50: 0.95). ***Polarity*:** Also observed in Tiphiiformes.

**370.** Anterior foramen of metasomal petiole (abdominal segment II) with a distinct anterior neck, whether short (subsessile) or long (pedunculate) (CB50: 0.84). ***Polarity*:** Observed for all sampled core Mutillidae. Otherwise, supported as independently derived in Tiphiidae, Chyphotidae, and Methochinae.

**521.** Fore wing cell 1rrs (submarginal 1) prestigmal length > ½ cell length (CB50: 0.83). ***Polarity*:** Also observed in Chyphotidae, *Myzinum*, *Pepsis*, crown Sapygidae, and †*Cretofedtschenkia*.

**523.** Fore wing cell 1rrs (submarginal 1) as long or longer than the two rsm cells (submarginal 2, 3) (CB50: 0.81). ***Polarity*:** Also observed in *Chyphotes*, †*Thanatotiphia*.

**555.** Fore wing Cu1 and Cu2 (anterior and posterior branches of Cuf) not diverging at approximately a right angle (CB50: 0.86). ***Polarity*:** Supported as a modification from the right-angled condition of the total Mutilliformes.

**572.** Hind wing distal elongation of sectoral cell (basal cell) absent (CB50: 0.91). ***Polarity*:** Supported as a synapomorphy of Chyphotidae and also observed in *Sierolomorpha*.

**573** Hind wing jugal lobe absent (CB50: 0.90). ***Polarity*:** Also observed in *Sierolomorpha* and *Typhoctes*.

### D.H. Family Sapygidae Latreille, 1810

***Clade comprising*:** • †**Cretosapyginae** Bennett & Engel, 2005, • **Fedtschenkiinae** André, 1899, **Sapyginae** Latreille, 1810. ***Incertae sedis*** in family: †*Cretofedtschenkia* Osten, 2007.

***Note*:** Synapomorphies are provided for the total Sapygidae and crown Sapygidae, *i.e.*, with and without †*Cretosapyga*. †*Cretofedtschenkia* was not included in the ancestral state reconstruction analysis due to the incompleteness of the fossil.

#### Total Sapygidae synapomorphies

**542.** Fore wing medial cell 2 (discal 2) proportionally small, with area less than that of 2 and 3rm (submarginal 2, 3) (CB50: 0.74). ***Polarity*:** A proportionally small second medial cell was also observed in *Sierolomorpha*, Sierolomorphidae CASENT0844589, one species of †*Burmusculus*, and both †*Cretosapyga* and †*Cretofedtschenkia*.

#### Crown Sapygidae synapomorphies

**94.** Anterior clypeal margin with a distinct median lobate process (CB50: 0.97). ***Polarity*:** Among sampled Pompiloidea *sensu novum*, also observed in Sierolomorphidae and *Tiphia*.

**69.** Prementum elongate, length ≥ 4 x width (CB50: 0.98). ***Polarity*:** This fluid feeding adaptation is supported as an autapomorphy among Pompiloidea *sensu novum*. The condition is unknown for †*Cretosapyga*.

**78.** Buccal cavity (external mouth opening of cranium) longer than broad (CB50: 0.98). ***Polarity*:** See polarity comment for premental elongation.

**85.** Median portion of clypeus craniocaudally longer than lateromedially wide (CB50: 0.95). ***Polarity*:** Unique among Pompiloidea *sensu novum*; †*Cretosapyga* with different proportions.

**94.** Anterior clypeal margin with a median lobate process (CB50: 0.97). ***Polarity*:** Independently derived in Sierolomorphidae; apparently absent in †*Cretosapyga*.

**158.** Antennomere III longer than IV (CB50: 0.91). ***Polarity*:** Also observed for *Sierolomorpha* and †*Burmusculus shennongi* among sampled Pompiloidea *sensu novum*.

**213.** Parascutal carina absent (CB50: 0.97). ***Polarity*:** Could not be evaluated for †*Loreisomorpha*. Convergently lost for total Tiphiiformes (CB50: 0.81) and total Sierolomorphidae (CB50: 0.77); reconstructed as trending toward loss independently between the Pompiliformes and Mutilliformes.

**277.** Procoxal “basicostal suture” or carina distinct and extending most of the distance from the base of the coxa to its apex (see definition for further detail) (CB50: 0.98). ***Polarity*:** Could not be evaluated for fossils. A long “basicostal” carina was also observed in *Pepsis* and *Aelurus*, indicating independent elongation of the carina.

**401.** Abdominal sternum III (metasomal II) without transverse sulcus dividing anterior articulatory surface (presclerite) from the posterior surface (postsclerite) (CB50: 0.98). ***Polarity*:** Unique among all sampled Pompiloidea *sensu novum*, extant and extinct; note that this could not be evaluated for all fossils.

**494.** Fore wing Rsf between Rs+M and 2r-rs linear, not kinked or curved (CB50: 0.94). ***Polarity*:** Also observed in most Pompiliformes and *Rhopalomutilla*, among sampled Tiphiopompiloides.

**516.** Fore wing crossvein 2rs-m joining Rs distal to 2r-rs by greater than one of its lengths (CB50: 0.96). ***Polarity*:** Also observed in Thynnidae, *Pepsis*, †*Cretofedtschenkia*, †*Architiphia*, some †*Burmusculus*, and the stem mutillid fossil (CASENT0844578).

**521.** Fore wing cell 1rrs (submarginal 1) prestigmal length > ½ cell length (CB50: 0.92). ***Polarity*:** Also observed in Chyphotidae, *Myzinum*, *Pepsis*, core Mutillidae, and †*Cretofedtschenkia*.

**539.** Fore wing crossvein 2m-cu curved or sinuate (CB50: 0.93). ***Polarity*:** Supported as convergently derived in Tiphiidae and Thynnidae.

### B’.2. Clade Scolioides

See the main transformation series (*i.e.*, leading to Formicoidea) above for definition based on the present analyses.

### **E.** Superfamily Scolioidea Latreille, 1802

*Clade comprising*: • **Bradynobaenidae** de Saussure, 1892, • **Scoliidae** Latreille, 1802.

#### Synapomorphies

**14.** Body size large, as evaluated by the head width proxy (≥ 1 mm) (CB50: 0.68). ***Polarity*:** All measured Scolioidea had head widths greater than 1 mm. Support stabilizes within the Scolioidea.

**149.** Radicle and scape offset at a distinct angle (CB50: 0.83). ***Polarity*:** Strongly supported as homoplastic among Vespoides; see polarity comment for Sierolomorphidae.

**177.** Alate pronotum anteroposteriorly short (CB50: 0.85). ***Polarity*:** All sampled Scolioidea had short pronota, including those fossils for which this condition could be evaluated. Pronotal reduction is variably reconstructed in Apoidea (see below).

**256.** Mesocoxae set in a ventral thoracic socket, with the sternal area nearly reaching the ventralmost coxal surfaces (CB50: 0.75). ***Polarity*:** Observed for all sampled extant taxa, but uncertain for all fossils. Support for this condition is maximal at the Bradynobaenidae and Scoliinae nodes and is strong for the crown Scoliidae (CB50: 0.98).

**263.** Mesocoxae wide-set (CB50: 0.63). ***Polarity*:** Observed in all sampled extant scolioid taxa, but uncertain for nearly all fossils with the exception of †*Sinoproscolia* which retains close-set coxae. This does suggest that †*Sinoproscolia* may be a stem group to the Scolioidea as a whole, rather than stem to Scoliidae as currently treated. Strongly supported for the Bradynobaenidae and crown Scoliidae (CB50: 0.99, 0.97, respectively).

**272.** Mesothoracic coxal cavity closed in lateral view (CB50: 0.71). ***Polarity*:** Observed in all sampled extant Scolioidea; uncertain for all fossils attributed to the group.

**282.** Protibial calcar unifid (CB50: 0.67). ***Polarity*:** Uncertain for all fossils, but true for all sampled extant Scolioidea. Characterizations of the form of the calcar should be critically refined.

**283.** Protibial calcar extremely strongly curved, almost forming a complete circle (CB50: 0.85). ***Polarity*:** Exactly as for char. 272.

**284.** Female protarsus with a row of psammochaetae (“fossorial setae”) (CB50: 0.63). ***Polarity*:** Exactly as for char. 272.

**299.** Metafemora with posteroapical lamella which is parallel to the longitudinal axis of the segment (CB50: 0.77). ***Polarity*:** Uncertain for *Proscolia* and most fossils. Observed to be present in †*Archaeoscolia* and †*Cretaproscolia*, and absent in †*Cretoscolia*.

**410.** Anterior articulatory surfaces of abdominal segment III (metasomal II) set distinctly above segment midheight (supraaxial) (CB50: 0.65). ***Polarity*:** Observed for all extant species; uncertain for all fossils.

**429.** Abdominal tergum VII of female and VIII of male (metasomal VI, VII) modified as “pygidial plate” (CB50: 0.63). ***Polarity*:** Observed for all extant species and uncertain for all fossils with the exception of †*Cretaproscolia asiatica*, for which the pygidial plate was observed to be present.

**457.** Fore and hind wings with apical longitudinal wrinkles (CB50: 0.84). ***Polarity*:** Observed in all extant Scolioidea, as well as for †*Cretaproscolia josai* and †*Sinoproscolia*; could not be evaluated for other fossils. Also observed in various large-bodied Apoidea.

### **E.A.** Family Bradynobaenidae de Saussure, 1892

***Clade comprising*:** • **Apterogyninae** André, 1899 (*Apterogyna* Latreille, 1809, *Micatalga* Argaman, 1994, *Macroocula* Panfilov, 1954, *Gynecaptera* Skorikov, 1935), • **Bradynobaeninae** de Saussure, 1892 (*Bradynobaenus* Spinola, 1851).

#### Synapomorphies

**2.** Females apterous (CB50: 0.97). ***Polarity*:** Clearly and independent derivation relative to Formicidae, Heterogynaidae, the various Pompiloidea *sensu novum*, and *Olixon*, among many other hymenopteran clades with apterous females.

**87.** Clypeal disc not projecting from cranium (CB50: 0.95). ***Polarity*:** Uniquely derived among Scolioidea; independently derived in Apoidea, for which insufficient signal was detected to identify the stem of transformation.

**104.** Genal secretory organ present (CB50: 0.99). ***Polarity*:** Unique among sampled Hymenoptera.

**138.** Antennal toruli extremely anteriorly-situated such that anterior torular margin is at or overhanging anterior clypeal margin (CB50: 0.96). ***Polarity*:** With the exception of some crown Formicidae, unique among Scolioides.

**150.** Radicle and scape nearly perpendicular (CB50: 0.92). ***Polarity*:** Virtually unique among sampled taxa, with the exception of *Aelurus* (Thynnidae), various Pompiloidea, and *Tatuidris* (Formicidae).

**157.** Male with grossly enlarged flagellum that has a smooth and shining surface (CB50: 0.99). ***Polarity*:** Unique among all sampled Hymenoptera.

**164.** Male mesosoma massively enlarged, appearing swollen or otherwise balloon-like (CB50: 0.97). ***Polarity*:** The condition of Bradynobaenidae was observed to be unique, although some male ants, for example, do have a “steroidal” facies.

**187.** Pronotum laterally scrobiculate, *i.e.*, with broad furrow or groove which can receive the fore leg when this appendage is flexed dorsomedially (CB50: 0.97). ***Polarity*:** The lateral pronotal scrobe of Bradynobaenidae is pronounced. Supported as independently derived in Apoidea, albeit the function of the broad lateral groove is less clear.

**227.** Longitudinal mesopectal sulcus absent (CB50: 0.96). ***Polarity*:** Despite uncertainty at the Scolioidea node for presence (CB50: 0.54), absence of the craniocaudally oriented sulcus on the mesopectus is strongly supported as ancestral for the Bradynobaenidae. Total Scoliidae, crown Scoliidae, and Scoliinae are all supported as retaining the groove (CB50: 0.62, 0.78, 0.98, respectively).

**288.** Proximal articulations of meso- and metacoxae distinctly constricted relative to the remainders of their segments (CB50: 0.97). ***Polarity*:** Unique among Scolioidea, but uncertain for all fossils.

**292.** Longitudinal axes of proximal articulation and main body of metacoxa at a distinct angle (CB50: 0.98). ***Polarity*:** As for char. 288.

**329.** Tarsomeres I–IV with apicomedian ventral lobate setae (CB50: 0.97). ***Polarity*:** Unique among sampled Scolioidea.

**370.** Metasomal petiole (metasomal segment I) with distinct anterior neck (peduncle) (CB50: 0.93). ***Polarity*:** Unique among Scolioidea, including fossil taxa.

**385.** Metasomal tergum I with paired anterodorsal carinae (CB50: 0.97). ***Polarity*:** Unique among the extant Scolioidea; uncertain for fossils.

15. **454.** Most or all of distal fourth to third of fore wing completely lacking venation (CB50: 0.98). ***Polarity*:** Also observed in crown Scoliidae, for which our analysis supported an independent derivation of this condition. All Mesozoic fossils attributed to Scoliidae have venation in their apical wing portion, including †*Araripescolia*, which we were unable to score. It is plausible to us that these fossils may actually be stem to the Bradynobaenidae plus Scoliidae, but to treat them as such here would be subjective, given our topological results.

**455.** Wing venation reduced to the “bradynobaenid pattern”, with the abscissae being particularly heavily sclerotized (see character definition for further detail) (CB50: 0.99). ***Polarity*:** Unique among all sampled Hymenoptera.

**\***Fore wing R not present distal to pterostigma (char. **463**, CB50: 0.96); Rsf1 absent (char. **481**, CB50: 0.99); fore wing Rs+M absent (char. **492**, CB50: 0.97); Rsf between Rs+M and 2r-rs linear (char. **494**, CB50: 0.90); Mf2+ absent (char. **501**, CB50: 0.96); crossvein 2rs-m absent (char. **511**, CB50: 0.92); crossvein 3rs-m absent (char. **517**, CB50: 0.95); crossvein 1m-cu absent (char. **530**, CB50: 0.93); 1cu-a prefurcal (char. **550**, CB50: 0.96), separated from M+Cu split by more than one of its lengths (char. **551**, CB50: 0.97); Cuf absent (char. **553**, CB50: 0.96); Cu2 not reaching 1A (char. **558**, CB50: 0.93). ***Polarity*:** Character states 481, 492, 530, 550, 551, 553, and 558 are unique among Scolioidea; states 463 and 501 are unique among sampled Scolioidea; and 494, 511, and 517 require specific comments. The state for 494 is also observed in Scoliinae and is variable among fossil Scoliidae (see raw data). The state for 511 has been convergently lost in core Campsomerini and *Scolia*; among fossils, 2rs-m is only absent in †*Cretaproscolia*. The state for 517, reduction of 3rs-m, also occurs in *Proscolia*, but is otherwise observed for all Scolioidea, including fossils.

**\*** Hind wing Rsf absent (char. **563**, CB50: 0.96), r-m absent (char. **565**, CB50: 0.96), Mf2 absent (char. **568**, CB50: 0.92), Hind wing Cuf absent (char. **569**, CB50: 0.92), distal elongation of hind wing basal cell absent (char. **572**, CB50: 0.98). ***Polarity*:** Character states 563, 565, 568, and 569 are unique among Scolioidea, while the state for 572 has been derived independently in core Scoliini.

### **E.B.** Family Scoliidae Latreille, 1802

***Clade comprising*:** • **Stem Scoliidae grade** (•• †**Archaeoscoliinae** Rasnitsyn, 1993 [†*Archaeoscolia* Rasnitsyn, 1993, †*Cretaproscolia* Rasnitsyn & Martínez-Delclòs, 1999, †*Floriscolia* Rasnitsyn, 1993], •• *incertae sedis* to subfamily among stem groups (†*Araripescolia* Nel *et al*., 2013, †*Cretoscolia* Rasnitsyn, 1993, †*Protoscolia* Zhang *et al*., 2002, †*Sinoproscolia* Zhang *et al*., 2015), • **crown Scoliidae clade** (•• **Proscoliinae** Rasnitsyn, 1977 [*Proscolia* Rasnitsyn, 1977], •• **Scoliinae** Latreille, 1802 [••• **Campsomerini** Bradley, 1957, ••• **Scoliini** Latreille, 1802]).

***Note 1*:** †*Floriscolia* is Cenozoic and was thus excluded from all analyses in the present work. Exclusion of †*Araripescolia* was an error of omission; we were unable to receive a copy of this work. †*Araripescolia* was described from the Crato formation and attributed by Nel *et al*. (2013) to the crown group near Campsomerini. Based on the results of our ancestral state analyses, we exclude †*Araripescolia* from the crown of the Scoliidae (see polarity notes below, particularly for the crown Scoliidae and Scoliinae nodes). Regarding the Eocene compression fossil †*Floriscolia*, we are less certain; there is insufficient preservation to evaluate the body, and some of the key venational features cannot be confidently interpreted. However, given that the fore wing Rf distal to the pterostigma is not strongly “downturned” (char. 478, crown Scoliidae below), we have sufficient evidence to consider the genus unplaced on the stem of the family. Finally, we recognize that †*Oryctopterus* Rasnitsyn, 1975 should have been included in our analysis given the putative sistergroup relationship with Scoliidae proposed by Rasnitsyn (1975).

***Note 2*:** Toward synthetic revision of the Scoliidae buttressed by phylogenomic analysis (Khouri *et al*. 2022; present data), we provide synapomorphies for seven maximally supported groups: (1) †*Protoscolia* plus the crown clade (“total Scoliidae”), (2) the crown clade, (3) Scoliinae, (4) Campsomerini (excluding *Colpa*), (5) core Campsomerini, (6) Scoliini (including *Colpa*), and (7) core Scoliini. Characters which could not be evaluated for *Proscolia*, but which may be synapomorphic for the crown node, are isolated in an additional subsection. Note that core Campsomerini is sister to *Trisciloa* Gribodo, 1893, while core Scoliini is sister to *Colpa* Dufour, 1841, the latter of which was attributed to its own subfamily by Argaman.

***Note 3*:** Many of the synapomorphies defining the Scoliidae are associated with stark modifications apparently representing skeletomuscular optimizations for digging, which is a crucial and perhaps deadly phase of the scoliid lifecycle. Altogether, they give the Scoliidae an appearance of miniature, spiny, and fossorial gorillas.

#### Total Scoliidae synapomorphies

**61.** Teeth present on basal mandibular margin (CB50: 0.96). ***Polarity*:** Uncertain for Scolioidea, as such teeth were observed to be present in *Bradynobaenus* but absent in *Apterogyna*. Among fossil Scoliidae, could only be evaluated for †*Protoscolia*. Not observed in Apoidea.

**66.** Mandible 4-dentate (CB50: 0.94). ***Polarity*:** Strongly supported as a synapomorphy of total Scoliidae, for which this condition could be evaluated for †*Protoscolia*. Mandibles are multidentate in *Bradynobaenus*.

**264.** Metacoxae wide-set (CB50: 0.97). ***Polarity*:** Observed in all sampled extant Scoliidae, and uncertain for most fossils with the exception of †*Sinoproscolia* and some †*Cretoscolia*, which have close-set metacoxae.

**265.** Metasternal region between metacoxae with a lateromedially broad, flat process (CB50: 0.97). ***Polarity*:** Exactly as for wide-set metacoxae.

**335.** Propodeum with paired longitudinal lines (CB50: 0.98). ***Polarity*:** Observed in all sampled extant Scoliidae, plus †*Archaeoscolia*, †*Cretoscolia*, and †*Protoscolia*. See char. 336 for crown Scoliidae below.

**470.** Fore wing crossvein 2r-rs anterior junction at distal apex of pterostigma (CB50: 0.93). ***Polarity*:** Observed in all Scoliidae, including fossils, and not observed in Bradynobaenidae. Also true for the unscored †*Araripescolia*.

**475.** Fore wing 2rrs cell (marginal 1) not apically pointed or narrowly rounded (CB50: 0.91). ***Polarity*:** Observed for all sampled extant and extinct Scoliidae, with the exception of †*Sinoproscolia*, with has the ancestral pointed to rounded condition.

**488.** Fore wing Mf1 distinctly curved (CB50: *). ***Polarity*:** Observed in Scoliinae and the majority of fossil Scoliidae, including †*Araripescolia*. For this reason, although the linear condition was equivocally reconstructed for the total Scoliidae (CB50: 0.61), the weight of evidence supports the curved condition. The ancestral state estimation only included †*Protoscolia normalis*, for which the curvature of Mf1 is unknown. Fossil Scoliidae which have a linear Mf1 include †*Archaeoscolia hispanica*, †*Cretoscolia Formosa*, †*Protoscolia sinensis*, and †*Sinoproscolia*.

**497.** Fore wing Mf2 as long or longer than Rs+M (CB50: 0.76). ***Polarity*:** Observed in Scoliinae and most fossil Scoliidae, with the exception of †*Cretoscolia laiyangica*, †*C. rasnitsyni*, †*Protoscolia sinensis*, and †*Sinoproscolia*.

**512.** Fore wing crossvein 2rs-m sinuate (CB50: 0.96). ***Polarity*:** Inapplicable for core Campsomerini and *Scolia*. Variable among fossil Scoliidae (see raw data).

**518.** Fore wing crossvein 3rs-m sinuate (CB50: 0.89). ***Polarity*:** Inapplicable to *Proscolia*. Otherwise, observed in Scoliinae and most fossils (see raw data).

**525.** Fore wing “submarginal cell 2” (2rm) quadrangular to triangular, at least three time as long proximodistally as wide anteroposteriorly (CB50: 0.75). ***Polarity*:** Cell in *Proscolia* with other shape. Observed in all fossil Scoliidae with the exception of †*Sinoproscolia*, †*Araripescolia*.

**531.** Fore wing 1m-cu distinctly curved or sinuate (CB50: 0.92). ***Polarity*:** Observed in all sampled Scoliidae, with the exception of †*Archaeoscolia senilis*, †*Cretoscolia brasiliensis*, and †*Araripescolia*.

**535.** Fore wing medial cell 1 (discal cell) proportionally elongate, with proximodistal length ∼ 4 x its anteroposterior length (CB50: 0.67). ***Polarity*:** *Proscolia* with a cell of other shape; observed in most fossils with the exception of †*Cretoscolia laiyangica*, †*C. rasnitsyni*, and †*Araripescolia*.

**539.** Fore wing 2m-cu curved or sinuate (CB50: *). ***Polarity*:** Although not observed in *Proscolia*, this was the condition of all fossil Scoliidae for which this trait could be evaluated.

**540.** Fore wing 2m-cu interstitial or prefurcal, *i.e.*, joining M at or proximal to 2rs-m (CB50: 0.94). ***Polarity*:** Observed in most fossil Scoliidae, with the exception of †*Sinoproscolia*; inapplicable for Bradynobaenidae.

**542.** Fore wing medial cell 2 (discal 2) proportionally small, with a surface area less than that of submarginal 2, 3 (CB50: 0.76). ***Polarity*:** Observed in most fossil Scoliidae, with the exception of †*Cretoscolia laiyangica* and †*Sinoproscolia* (†*Araripescolia* intermediate). *Proscolia* with a larger cell.

**544.** Fore wing medial cell 2 (discal 2) proximodistal length ≥ 2 x anteroposterior width (CB50: 0.89). ***Polarity*:** Observed in all fossil Scoliidae including †*Araripescolia* with the exception of †*Cretaproscolia josai* and some †*Cretoscolia*; reversed to the shorter condition in core Campsomerini; inapplicable in Bradynobaenidae and core Scoliini.

**562.** Hind wing C present (CB50: 0.54). ***Polarity*:** Although the support is equivocal, this reflects the uncertainty of the fossils, none of which could be evaluated for C. Otherwise, observed in all extant Scolioidea.

#### Crown Scoliidae synapomorphies

**19.** Head of both sexes more-or-less cuboidal, with cranium elongate posterad compound eyes (CB50: 0.77). ***Polarity*:** Although female Bradynobaenidae have long heads posterior to the compound eyes, males do not. Independently derived in Ampulicidae and other Apoidea.

**30.** Malar space virtually absent (CB50: 0.89). ***Polarity*:** Could not be evaluated for stem Scoliidae, but true for all sampled extant terminals within the family. Supported as independently derived for Apoidea.

**116.** Longitudinal carinae present mediad antennal toruli (“medial frontal carinae”) (CB50: 0.84). ***Polarity*:** Unique among Scolioidea; reversed in core Scoliini.

**142.** Antennal toruli directed laterally, *i.e.*, away from one another (see definition for more detail) (CB50: 0.92). ***Polarity*:** Observed for all crown Scoliidae, but could not be evaluated for any sampled fossils, thus support at the total Scoliidae node is equivocal.

**258.** Mesocoxae extremely foreshortened, being hardly or not at all produced away from the body (CB50: 0.92). ***Polarity*:** Observed for all sampled extant Scoliidae, but uncertain for all fossils.

**273.** Metathoracic coxal cavity closed in lateral view (CB50: 0.70). ***Polarity*:** As for char. 258; uncertain for *Proscolia*.

**277.** “Basicostal carina” of procoxa elongate (see character definition for further detail, CB50: 0.59). ***Polarity*:** Exactly as for char. 273.

**300.** External surfaces of meso- and metatibiae with strong, scattered spurs (chaetae, “traction setae”) (CB50: 0.93). ***Polarity*:** Observed in all sampled extant Scoliidae plus *Bradynobaenus*; absent in †*Cretoscolia*, †*Protoscolia*; present in †*Archaeoscolia*; uncertain for †*Sinoproscolia*.

**322.** Tarsomeres apically expanded and conical (CB50: 0.99). ***Polarity*:** Also observed in *Bradynobaenus*, but not in *Apterogyna*. The tarsomeres of †*Cretaproscolia* and †*Cretoscolia* were observed to be narrow and linear; the condition for other fossils is uncertain.

**324.** Pretarsal claws unidentate (CB50: 0.87). ***Polarity*:** Observed for all extant Scoliidae plus *Apterogyna*; uncertain for most fossils with the exception of †*Cretoscolia promissiva*, which is also unidentate.

**336.** Propodeal lines extending to propodeal foramen without meeting, thus propodeum divided into three well-defined sections (CB50: 0.95). ***Polarity*:** Observed for all extant Scoliidae plus †*Archaeoscolia* and †*Cretoscolia*; not observed in †*Protoscolia* and †*Sinoproscolia*.

**454.** Most or all of distal fourth to third of fore wing completely lacking venation (CB50: 1.0). ***Polarity*:** See polarity comments for this character at the Bradynobaenidae node.

**478.** Marginal cell with apex strongly “downcurved”, with Rf separating widely from wing margin (CB50: 0.91). ***Polarity*:** Variable among fossils, not applicable to Bradynobaenidae. Observed in †*Archaeoscolia*, some †*Cretoscolia*, and †*Araripescolia*; not observed in †*Cretaproscolia*, some †*Cretoscolia*, †*Protoscolia*, and †*Sinoproscolia*. Note that †*Protoscolia* represent an intermediate state (char. 477).

**571.** Hind wing cu-a interstitial or prefurcal (CB50: 0.78). ***Polarity*:** Inapplicable for Bradynobaenidae; postfurcal for all fossils for which this could be evaluated, including

†*Araripescolia*. Reversed to postfurcal in core Campsomerini, core Scoliini.

#### *Characters predicted to be present in Proscolia

*Note: Characters listed in this subsection are strongly to maximally supported at the Scoliinae node, but are unknown for* Proscolia*, thus may actually constitute synapomorphies of either crown Scoliidae or simply the Scoliinae*.

**45.** Craniomandibular articulation modified: lateral base of mandible in form of cylindrical bar (CB50: 1.0). ***Polarity*:** State uncertain for *Proscolia*, which must be evaluated before this synapomorphy is accepted formally as defining Scoliinae, as it may be unique to the Scoliidae. Not observed in other taxa.

**78.** Buccal cavity longer than broad (CB50: 0.95). ***Polarity*:** As for char. 45 above.

**184.** Pronotum with distinct transverse line or groove on lateral surfaces (CB50: 0.70). ***Polarity*:** Because specimens of *Proscolia* were unavailable for examination, we scored this terminal as uncertain for this condition. Our reconstruction predicts, with some reservation, that *Proscolia* will indeed have the transverse line. Supported as independently derived in Apoidea.

**197.** Lateral portions of pronotum, *i.e.*, those areas lateral to the mesoscutum, dorsoventrally broad and posteriorly notched (CB50: 0.58). ***Polarity*:** This is another condition which is uncertain for but predicted to be true in *Proscolia* based on our present analysis. Support is maximal for this condition among all nodes of the Scoliinae.

**203.** Propleurae deeply concave, receiving cranium when head tucked ventrally (CB50: 0.57). ***Polarity*:** Maximally supported as a synapomorphy for Scoliinae, but condition unknown for and weakly predicted to be true in *Proscolia*.

**205.** Pronotum miniscule when observed externally, forming a longitudinally oriented lamella (CB50: 0.66). ***Polarity*:** Support for this condition at the Scoliinae node is maximal; the state for *Proscolia* is unknown.

**234.** Mesopectus posteriorly scrobiculate*, i.e.*, with a broad dorsoventrally oriented furrow or groove which receives the hind leg when this appendage is flexed dorsomedially (CB50: 0.60). ***Polarity*:** Observed in all Scoliinae, unknown for *Proscolia*.

**245.** Both upper and lower metapleural areas enlarged, being nearly as large together as the propodeum in profile view (CB50: 0.59). ***Polarity*:** Observed in all Scoliinae, unknown for *Proscolia*.

#### Scoliinae synapomorphies

**35.** Medial margins of compound eyes notched (CB50: 0.92). ***Polarity*:** Observed in all Scoliinae; supported as independently derived with respect to Vespoidea, some Formicidae, and some Apoidea.

**69.** Prementum elongate, proximodistal length ≥ 4 x lateromedial width (CB50: 0.96). ***Polarity*:** The prementum is shorter in *Proscolia* and Bradynobaenidae but could not be evaluated for Mesozoic fossils.

**266.** Metasternal process with a linear anterior margin (CB50: 0.99). ***Polarity*:** Observed for all sampled Scoliinae. The anterior margin is angled in *Proscolia*.

**310.** Mesotibia with a single ventroapical spur (CB50: 0.96). ***Polarity*:** Observed for all Scoliinae; the two-spurred condition is strongly supported as retained in Bradynobaenidae and was observed for a †*Cretoscolia* species (see raw data).

**301.** Metatibial traction spurs arranged in at least two more-or-less even, transverse rows which are buttressed by strong, raised cuticular ridges (CB50: 0.73). ***Polarity*:** Uncertain for *Proscolia*. Equivocally supported as a synapomorphy of Scoliinae and as ancestral for Scoliini (CB50: 0.73, 0.72, respectively) due to presence of this conformation in *Colpa*. This can either be interpreted as a synapomorphy of the subfamily or independently derived between *Colpa* and Campsomerini. Without any further anatomical or developmental evidence to weigh, we tentatively interpret this as a synapomorphy of the Scoliinae based on the principle of parsimony. Direct examination of *Proscolia* will be critical for resolving the polarity of this trait.

**428.** Abdominal sternum VII (metasomal VI) of female with paired horn-like processes on the lateral margins (CB05: 0.73). ***Polarity*:** Exactly as for char. 301 above, down to the support values. In this case, we argue that this morphogenetic expression is unique, and thus unlikely to evolve twice independently given the current knowledge. The discovery of intermediate taxa would provide further evidence for the independent hypothesis if these are to be recovered from the fossil record. Furthermore, we suggest that that female expression represents a case of morphogen cooption from the male, for which sternal prongs are strongly supported as synapomorphic for the subfamily (see char. 438 below).

**438.** Abdominal sternum IX (metasomal VIII) of male with three widely spaced, spiniform processes (CB50: 0.99). ***Polarity*:** Unique among sampled Hymenoptera, although we note that spiniform processes or prongs are indeed observed among other groups (*e.g.*, Tiphiiformes, Dorylinae).

**494.** Fore wing Rsf between Rs+M and 2r-rs linear (CB50: 0.89). ***Polarity*:** Also observed in Bradynobaenidae and various fossil Scoliidae (see raw data).

**547.** Fore wing crossvein 1cu-a sinuate (CB50: 0.95). ***Polarity*:** 1cu-a is not sinuate in *Proscolia*, nor for most of the fossil Scoliidae with the exception of †*Cretaproscolia* and †*Araripescolia*.

**548.** Fore wing 1cu-a postfurcal, joining M distal to split of M+Cu (CB50: 0.97). ***Polarity*:** Among fossil Scoliidae, only observed for †*Protoscolia* and †*Cretoscolia*; not observed for †*Araripescolia*, which has an interstitial 1cu-a.

**556.** Fore wing Cuf distal to M+Cu directed proximally (CB50: 0.98). ***Polarity*:** Not observed in *Proscolia* but observed in most fossil Scoliidae with the exception of †*Cretoscolia laiyangica*, †*Protoscolia*, and †*Araripescolia*.

#### Campsomerini synapomorphies

**248.** Upper metapleural area, *i.e.*, that region dorsad the endophragmal pit, with a longitudinal carina in its dorsal fourth (see character definition for further detail, CB50: 0.99). ***Polarity*:** Unique to Campsomerini.

**483.** Fore wing Rsf1 and Mf1 meeting at a distinct oblique angle (CB50: 0.91). ***Polarity*:** Unique among sampled extant Scoliidae; among fossils, only observed in †*Cretoscolia brasiliensis*. †*Araripescolia* with Rsf1 and Mf1 more-or-less aligned.

#### Core Campsomerini synapomorphies

**33.** Compound eyes converging away from the mouth to some degree (CB50: 0.77). ***Polarity*:** Among sampled Scolioidea, also observed in *Proscolia*; among fossils, could only be evaluated for †*Cretoscolia laiyangica*, which did not have converging eyes.

**235.** Mesopectus in cross-section broadly V-shaped (CB50: 0.99). ***Polarity*:** Not observed in *Trisciloa*, nor in any other Scolioidea.

**314.** Meso- and/or metatibial spurs extremely elongate, the posterior or sole spur being > ½ basitarsus length (CB50: 0.98). ***Polarity*:** Strongly supported as independently derived in *Colpa*.

**341.** Dorsal face of propodeum with a distinct, raised dorsomedial triangle (see character definition for further detail, CB50: 0.99). ***Polarity*:** Unique among all sampled Hymenoptera.

**430.** “Pygidial plate” of female covered in stout, appressed setae which taper to a fine point and resemble scales (CB50: 0.99). ***Polarity*:** Unique among all sampled Hymenoptera.

**511.** Fore wing crossvein 2rs-m absent (CB50: 0.93). ***Polarity*:** Convergently lost in Bradynobaenidae and *Scolia*; among fossils, 2rs-m is only absent in †*Cretaproscolia*.

**544.** Fore wing medial cell 2 (discal 2) proximodistal length < 2 x anteroposterior width (CB50: 0.92). ***Polarity*:** Support at the Campsomerini node is decayed, but still indicates the long condition (CB50: 0.73). The condition of Campsomerini is otherwise strongly supported as a reversal.

**571.** Hind wing cu-a postfurcal (CB50: 1.0). ***Polarity*:** Supported as an independent derivation (reversal) from core Scoliini.

#### Scoliini synapomorphies

**472.** Fore wing 2rrs cell (marginal 1) small, approximately equal in size to or smaller than pterostigma (CB50: 0.98). ***Polarity*:** Supported as independently derived in *Proscolia* and the campsomerine *Cathimeris*; not observed in fossil taxa, including †*Araripescolia*.

#### Core Scoliini synapomorphies

**116.** Longitudinal carinae absent mediad antennal toruli (“medial frontal carinae”) (CB50: 0.97). ***Polarity*:** This represents a reversal to absence from the synapomorphic present condition of crown Scoliidae.

**267.** Metasternal process with a transverse sulcus just anterad the posterior margin (CB50: 0.99). ***Polarity*:** Observed to be unique to the core Scoliini, to the exclusion of *Colpa*.

**301.** Metatibial traction spurs messily arranged, *i.e.*, not in transverse rows which are buttressed by strong, raised cuticular ridges (CB50: 1.0). ***Polarity*:** See the polarity comments for this character at the Scoliinae node.

**428.** Abdominal sternum VII (metasomal VI) of female lacking paired horn-like processes on the lateral margins (CB05: 1.0). ***Polarity*:** Exactly as for char. 301 above, down to the support values.

**498.** Fore wing Mf3 as long or longer than Rs+M (CB50: 0.97). ***Polarity*:** Among sampled extant Scoliidae, also observed in *Trisciloa*; variable among fossil Scoliidae (see raw data).

**538.** Fore wing crossvein 2m-cu nebulous to absent (CB50: 0.96). ***Polarity*:** Unique among sampled Scoliidae, including †*Araripescolia* and the sampled fossils.

**571.** Hind wing cu-a postfurcal (CB50: 1.0). ***Polarity*:** Supported as an independent derivation (reversal) from core Campsomerini.

**572.** Distal elongation of hind wing basal cell absent (CB50: 0.98). ***Polarity*:** Derived independently from Bradynobaenidae.

### E’. Clade Formicapoidina

See the main transformation series (*i.e.*, leading to Formicoidea) above for definition based on the present analyses.

### G. Superfamily Apoidea, Latreille, 1802

*Clade comprising*: \***Extant taxa:** • **Ampulicidae** Shuckard, 1840, • **core Apoidea** clade.

\***Extinct taxa:** • ***Incertae sedis*** in superfamily: **†Angarosphecidae** Rasnitsyn, 1975 (†*Angarosphex* Rasnitsyn, 1975, †*Archisphex* Evans, 1969, †*Baissodes* Rasnitsyn, 1975, †*Cretobestiola* Rasnitsyn & Pulawski, 2000, †*Cretosphecium* Pulawski & Rasnitsyn, 1999, †*Cretosphex* Rasnitsyn, 1975, †*Decasphex* Zheng *et al*., 2021, †*Eosphecium* Pulawski & Rasnitsyn, 1999, †*Ilerdosphex* Rasnitsyn, 2000, †*Montsecosphex* Rasnitsyn & Martínez-Delclòs, 2000, †*Oryctobaissodes* Rasnitsyn, 1975, †*Pompilopterus* Rasnitsyn, 1975, †*Trichobaissodes* Rasnitsyn, 1975, †*Vitimosphex* Rasnitsyn, 1975), **†Allommationidae** Rosa & Melo, 2021 (†*Allommation* Rosa & Melo, 2021), **†Spheciellidae** Rosa & Melo, 2021 (†*Spheciellus* Rosa & Melo, 2021). • ***Incertae sedis*** to family: †**Burmastatinae** Antropov, 2000 (†*Burmastatus* Antropov, 2000), †**Apodolichurinae** Antropov, 2000 (†*Apodolichurus* Antropov, 2000), †**Mendampulicinae** Antropov, 2000 (†*Mendampulex* Antropov, 2000), †*Gallosphex* Schlüter, 1978, †*Trigampulex* Antropov, 2000. • ***Incertae sedis*** to superfamily: †*Burmasphex* Melo & Rosa, 2018.

***Note*:** †Angarosphecidae is a critical but confounding grouping for understanding the tempo of apoid evolution. In the present study, taxa attributed to this family were found to have strong to weak affinities with Apoidea, with some terminals clustering outside of the superfamily. See the Extended Topological Results and Apoid Calibration sections for further detail. Note that †Angarosphecidae are hypothesized to be distinguished from the remainder of the Apoidea by plesiomorphic absence of the deep constriction between the pronotum and mesoscutum (synapomorphy of Apoidea with reversal in Anthophila, char. 192 below), rather having a pronotum which closely fits over the mesoscutum. See Zheng *et al*. (2021) for a key to the genera based on fore wing venation and a taxonomic synopsis at the species level.

#### Synapomorphies

**30.** Malar space virtually absent (CB50: 0.69). ***Polarity*:** Although with equivocal support at this node, almost all of the extant terminals sampled had strongly reduced malar spaces. Supported as independently derived from Scoliidae.

**149.** Radicle and scape offset at a distinct angle (CB50: 0.90). ***Polarity*:** Strongly supported as homoplastic among Vespoides; see polarity comment for Sierolomorphidae.

**177.** Pronotum anteroposteriorly short (CB50: 0.60). ***Polarity*:** Although support is equivocal, the character distribution for this state does suggest that an elongate pronotum may have been a secondary gain in Ampulicidae among Apoidea.

**184.** Pronotum with transverse line or groove on lateral surfaces (CB50: 0.88). ***Polarity*:** Supported as an independent derivation relative to Scolioidea. Observed in the majority of sampled Apoidea, extant and extinct. Alate Formicoidea lack this line altogether.

**187.** Pronotum with a broad lateral furrow or groove (CB50: 0.92). ***Polarity*:** The lateral pronotal line of Apoidea is often set in a broad furrow which resembles a scrobe for the fore leg, but for which the function is uncertain.

**\*** Pronotum broadly constricted just anterior to mesoscutum (char. **191**, CB50: 0.95), with the muscular portion posterior to the constriction also separated from the mesoscutum by a deep and narrow impression which resembles an incision in profile view (char. **192**, CB50: 0.97). ***Polarity*:** The conformation described by char. 192 is unique, but the constriction is also observed in †@@@idae and some aquilomyrmecine †Haidomyrmecinae (*e.g.*, †*Chonidris*).

**198.** Pronotal lobe, *i.e.*, that posterolateral portion of the pronotum which conceals the first mesosomal spiracle, grossly enlarged and dorsoventrally compressed, thus appearing as a semicircular extension of the pronotum (CB50: 0.98). ***Polarity*:** One of the canonical synapomorphies of the Apoidea. Observed in all sampled extant species and for a number of fossils. See the character definition and raw data for more information.

**200.** Pronotum forming a complete ring posterior to the procoxae (CB50: 0.97). ***Polarity*:** Another strong synapomorphy of the Apoidea as for the pronotal lobe; see the character definition and raw data for more information.

**226.** Mesopectus with a deep pit subtending a flange at the base of the fore wing articulation (CB50: 0.83). ***Polarity*:** Uncertain for the majority of fossils. Observed for most sampled extant Apoidea with the exception of *Heterogyna*, *Ammophila*, *Sphex*, *Prionyx*, and *Sphecius*. Probably associated with internal modification(s).

**228.** Mesopectus with a dorsoventrally oriented sulcus present in the anterior half of the sclerite (CB50: 0.73). ***Polarity*:** Observed in most sampled Apoidea, including fossils; the uncertainty probably reflects that of several of the fossil terminals. See the raw data for further character distribution information.

**241.** Metapleural area scrobiform, *i.e.*, surface of metapleural area directed posteriorly due to bulging mesopectus, and with capacity for receiving a leg if upfolded (CB50: 0.87). ***Polarity*:** Support for this condition stabilizes among the first several internal nodes and indicates reversal in some clades.

**261.** Ventral surface of the metapectus, bearing the medial metacoxal articulations, strongly produced ventrally such that the metacoxal foramen is directed laterally and the hinge of motion is nearly vertical (CB50: 0.95). ***Polarity*:** Autapomorphic for Apoidea.

**\*.** Propodeum with paired longitudinal lines (char. **335**, CB50: 0.95), these lines meeting posteriorly, thus delimiting a dorsal triangular region with its vertex directed posteriorly (char. **338**, CB50: 0.95). ***Polarity*:** The “propodeal triangle” was not observed in †*Cretobestiola* but was observed in various amber fossils. Other fossil taxa were uncertain. See raw data.

***Note*:** The interpretation for curvature of 2m-cu (char. **539**) is uncertain; the crossvein is curved in many extant Apoidea but is variable for fossils. Elongation of the fore wing medial cell 2 (discal 2) such that its proximodistal length is ≥ 2 x anteroposterior width (char. **544**) is supported as independently derived between Ampulicidae and the Crabronidae–Anthophila clade but could be a synapomorphy of the Apoidea as a whole. The ancestral dental count for the mandibles was supported as at least unidentate (char. 63, CB50: 0.99); the bidentate condition (char. 64) was unstable among most of the backbone node but stabilizes at the family level in our analyses. For this reason, we do not report dental count reconstructions below, suffice it to note that Crabronidae are supported as unidentate (CB50: 0.96), Sphecidae variable, but stabilizing at bidentate (CB50: 0.54, 0.92), Bembicidae bidentate (CB50: 0.96), and Pemphredonidae– Anthophila clade unidentate with variation among shallower nodes (CB50: 0.86).

### G.A. Family Ampulicidae Shuckard, 1840

***Clade comprising*:** • **Ampulicinae** Shuckard, 1840 (Ampulicini Shuckard, 1840), • **Dolichurinae** Lepeletier de Saint-Fargeau, 1845 (•• Aphelotomini Ohl & Spahn, 2009, Dolichurini Dahlbom, 1842). • ***Incertae sedis*** in family: †**Cretampulicini** Antropov, 2000 (†*Cretampulex* Antropov, 2000).

***Note*:** The undescribed burmite genus CASENT0844594 (Fig. S12) clustered on the stem of the total Ampulicidae in the ancestral state estimation analysis, which we remain skeptical about. Additionally, the undescribed burmite taxon CASENT0844571 was recovered as sister to the crown Ampulicidae. For these reasons, synapomorphies are provided for the first, second, and third total nodes (*i.e.*, including CASENT0844594, †*Cretampulex* Antropov, and CASENT0844571, respectively), as well as the crown node.

#### Total Ampulicidae node 1 synapomorphies

**143.** Antennal toruli directed toward the mandibles to some degree (CB50: 0.90). ***Polarity*:** Observed in Ampulicidae, including fossils, plus *Heterogyna* and *Chalybion*, which are both supported as independent origins of this condition.

#### Total Ampulicidae node 2 synapomorphies

**342.** Propodeum elongate, in the form of a rectangular box (CB50: 0.98). ***Polarity*:** Supported as independently derived in core Apoidea.

**488.** Fore wing Mf1 distinctly curved (CB50: 0.80). ***Polarity*:** Observed in both sampled extant species, CASENT0884571, †*Cretampulex*, and †*Gallosphex*, but not †*Apodolichurus*; uncertain for †*Mendampulex*. Mf1 is linear in most core Apoidea but curved in various fossils (see raw data).

#### Total Ampulicidae node 3 synapomorphies

**14.** Body size large, as evaluated by the head width proxy (≥ 1 mm) (CB50: 0.94). ***Polarity*:** Large body size relative to this proxy is supported as independently derived among other Apoidea.

**33.** Compound eyes converging away from the mouth to some degree (CB50: 0.77). ***Polarity*:** Among sampled Apoidea, supported as independently derived within Crabronidae and within Philanthidae.

**142.** Antennal toruli directed laterally, *i.e.*, away from one another (CB50: 0.97). ***Polarity*:** Due to the combination of orally directed mandibles, the toruli of these Ampulicidae are directed anterolaterally, assuming the prognathy. Unique among sampled Apoidea with the exception of *Chalybion*. Among fossils, observed for CASENT0844571, but not for †*Cretampulex* or †*Mendampulex*.

**177.** Pronotum anteroposteriorly elongate (CB50: 0.98). ***Polarity*:** Support for the short condition is moderate at the two preceding nodes (CB50: 0.88 for total 1, 0.80 for total 2). Among extant Apoidea, comparatively elongate pronotum was observed in Ampulicidae, while among fossils, this state was scored as true for †*Apodolichurus*, CASENT0844571, †*Mendampulex*, †*Gallosphex*, and some impression fossils (see raw matrix).

**179.** Pronotum with longitudinal margination between lateral and dorsal surfaces (CB50: 0.97). ***Polarity*:** Unique to Ampulicidae among Apoidea; not observed in Scolioidea, and independently derived in some Formicidae. Among fossils, marginate pronota were observed in †*Apodolichurus*, CASENT0844571, †*Mendampulex*, †*Gallosphex*, and was apparent in some impression fossil taxa; the pronotum was observed to be unmargined in †*Cretampulex*.

**229.** Mesopectus with dorsoventrally oriented carina situated approximately at the anterolateral margin of the sclerite (CB50: 0.95). ***Polarity*:** Condition uncertain for †*Cretampulex*. Also observed in *Oxybelus*, *Sphecius*, and *Pluto*, among sampled extant Apoidea. See raw data for fossil state distribution.

**348.** Propodeum armed (CB50: 0.93). ***Polarity*:** Observed in CASENT0844571, but not in †*Apodolichurus*; uncertain for †*Cretampulex*, †*Mendampulex*, and †*Gallosphex*.

**370.** Abdominal segment II (metasomal I) with an anterior neck (peduncle) (CB50: 0.94). ***Polarity*:** As for char. 348; present in various core Apoidea.

**382.** Metasomal sternum I with an anteroventral process that fits between the metacoxae at full metasomal flexion (CB50: 0.77). ***Polarity*:** Observed in *Dolichurus* and CASENT0844571, but not in *Ampulex*.

**385.** Metasomal tergum I with paired anterodorsal carinae (CB50: 0.79). ***Polarity*:** Exactly as for char. 382 above.

**402.** Helcial sternite exposed in lateral view (see character definition for more detail, CB50: 0.99). ***Polarity*:** The helcial sternite is concealed in the majority of sampled Apoidea.

**518.** Fore wing crossvein 3rs-m sinuate (CB50: 0.95). ***Polarity*:** Observed in most sampled Apoidea, including compression fossils; variable among amber fossils (see raw data).

**544.** Fore wing medial cell 2 (discal 2) with proximodistal length ≥ 2 x anteroposterior width (CB50: 0.95). ***Polarity*:** Supported in our analyses as an independent derivation with respect to the Crabronidae–Anthophila clade. We treat this with heightened skepticism, although we do also recognize that this reconstruction is informed by our sampling of fossil taxa.

#### Crown Ampulicidae synapomorphies

**116.** Longitudinal carinae present mediad antennal toruli (“medial frontal carinae”) (CB50: 0.93). ***Polarity*:** Among sampled Apoidea, independently derived in *Chalybion*. Absent in †*Cretampulex* and †*Mendampulex*.

**295.** Female meso- and/or metafemora muscularly swollen (CB50: 0.83). ***Polarity*:** Not observed in any fossils attributable to the family.

**410.** Anterior articulatory surfaces of abdominal segment III (metasomal II) set distinctly above segment midheight (supraaxial) (CB50: 0.83). ***Polarity*:** Although the support value is moderate, this appears to be a strong synapomorphy of crown Ampulicidae.

**523.** Fore wing 1rrs cell (submarginal 1) shorter than submarginal cells 2, 3 (CB50: 0.89). ***Polarity*:** Uncertain for CASENT0844571 and †*Mendampulex*. Otherwise, observed in †*Cretampulex*; inapplicable for †*Apodolichurus*.

**535.** Fore wing medial cell 1 (discal cell) proportionally elongate, with proximodistal length ∼ 4 x its anteroposterior length (CB50: 0.79). ***Polarity*:** Not observed in fossils attributable to the family. Variable among Apoidea (see raw data).

***Note*:** Female notauli present in Ampulicidae (char. **217**), but polarity highly uncertain. See comments for Crabronidae–Anthophila clade below.

#### **G.1.** Core Apoidea clade

***Clade comprising*:** • **Heterogynaidae** Nagy, 1969 (*Heterogyna* Nagy, 1969; †Ptilocosminae Rosa & Melo, 2021: †*Ptilocosmus* Rosa & Melo, 2021, †*Thigmocosmus* Rosa & Melo, 2021, †*Aulacicocosmus* Rosa & Melo, 2021), †**Cirrosphecidae** Antropov, 2000 (†Cirrosphecinae Antropov, 2000: †*Cirrosphex* Antropov, 2000, †*Haptodioctes* Rosa & Melo, 2021; †Glenocephalinae Rosa & Melo, 2021: †*Glenocephalus* Rosa & Melo, 2021), • Crabronidae–Anthophila clade.

***Note*:** †*Cirrosphex* Antropov was recovered as sister to the core Apoidea, so both the stem and crown nodes of core Apoidea are addressed below. The placement of other genera of the †Cirrosphecidae have not been tested. Within the core Apoidea, the fossils have very uncertain placement, so subsequent clades are not provided with synapomorphies, with the exception of the total Anthophila.

#### Total core Apoidea Synapomorphies

**400.** Abdominal tergum III (metasomal II) without a transverse sulcus dividing the anterior articulatory surface (pretergite) from the posterior region (posttergite) (CB50: 0.81). ***Polarity*:** Presclerites are observed in Ampulicidae and some Apoidea, but not in †*Cirrosphex* and others (see raw data).

**571.** Hind wing cu-a interstitial or prefurcal (CB50: 0.90). ***Polarity*:** Observed in most core Apoidea (see raw data).

### Crown core Apoidea Synapomorphies

**324.** Pretarsal claws unidentate (CB50: 0.81). ***Polarity*:** Among sampled extant Apoidea, bidentate claws were observed in Ampulicidae, Sphecidae, and Anthophila. Among fossils for which this condition could be evaluated, the bidentate claws were observed in †*Cretampulex*, CASENT0844571, †*Mendampulex*, †*Gallosphex*, †*Cirrosphex*, †*Pittoecus*, †*Melittosphex*, and †*Cretotrigona*; unidentate claws were observed for †*Apodolichurus*, and the majority of other taxa (see raw data).

### G.2’. Crabronidae–Anthophila clade

*Clade comprising*: • **Crabronidae–Sphecidae clade**, • **Bembicidae–Anthophila clade**.

***Note*:** All Mesozoic fossils attributed to the Pemphredonidae clustered together on the stem of the Crabronidae–Anthophila clade in our ancestral state estimation analyses, with the addition of the otherwise sphecid-like CASENT0844566. We suspect this is a pattern of “stemward slippage” which is due to missing data, either absent sequences or uncertain states (“?”).

#### Crabronidae–Anthophila clade synapomorphies

**78.** Buccal cavity longer than broad (CB50: 0.76). ***Polarity*:** All rootward nodes from the Crabronidae–Anthophila clade to the Apoidea maximally support a cavity which is as shorter.

**217.** Female lacking notauli on mesoscutum (CB50: 0.95). ***Polarity*:** Female notauli were observed in crown and fossil Ampulicidae, *Pluto*, †*Burmastatus*, and †*Psolimena*. Variable among other Vespida, being absent in most Vespoidea, various Pompiloidea *sensu novum*, Scolioidea, and most ants. Because presence of female notauli in Vespida is the minority condition, it is plausible that these represent independent gains. It is just as likely, however, that the notauli have been lost in parallel. Further taxonomic sampling and comparative anatomy will contribute to the resolution of this conundrum.

**259.** Ventral surface of mesopectus, bearing medial mesocoxal articulation, strongly produced ventrally such that the mesocoxal foramina are directed laterally with a nearly vertical hinge (CB50: 1.0). ***Polarity*:** This is a very strong synapomorphy of the Crabronidae– Anthophila clade, only reversed in *Cerceris* (Philanthidae) among sampled Apoidea. *Heterogyna* does not display this condition, but its polarity is uncertain due to the lack of topological resolution for this taxon. The majority of spheciform fossils for which this could be evaluated had the ventral production, with the exception of those fossils attributable to Ampulicidae and CASENT0844594.

**260.** Ventrally expanded ventral surface of mesopectus lateromedially expanded, forming lobe-like plates which *do not* overlap the mesocoxae (CB50: 1.0). ***Polarity*:** This additive condition represents the ancestral form of the mesopectal lobes in the Crabronidae– Anthophila clade; the lobes are reduced or otherwise narrowed synapomorphically for the Anthophila.

**263.** Mesocoxae wide-set (CB50: 0.76). ***Polarity*:** Wide-set mesocoxae are observed in the majority of sampled taxa within the Crabronidae–Anthophila clade, with the exception of *Sphecius*, Ammoplanidae, Pemphredonidae, and *Apis*. Uncertain for most fossils (see raw data for state distribution).

**310.** Mesotibia with a single ventroapical spur (CB50: 0.92). ***Polarity*:** Presence of the spur in Sphecidae is strongly supported as a reversal in our analyses. However, we recognize that this is contrary to expectations as the pattern is not Dollo-like, which is an intuitively valid criticism. We expect that expanded taxon sampling may contribute significant information for this reconstruction, particularly as Astatidae, Mellinidae, Nyssonini, and Stizini retain two spurs (Bohart & Menke, 1976). Given this pattern, we list the single-spur condition as a synapomorphy as this was the result supported by our data, and to highly the need for sampling. We also note that a similar “reversal” two the two-spur condition was supported for the Myrmeciinae within the clade Formicae of the Formicidae (see above).

**494** Fore wing Rsf between Rs+M and 2r-rs linear, not kinked or distinctly curved (CB50: 0.61). ***Polarity*:** Reversal to the kinked or curved condition is supported for *Bembix*, Philanthidae, and various Anthophila, among our sampled Apoidea. The linear condition is supported as independently derived in *Ampulex*. See raw data for fossil state distribution.

**535.** Fore wing medial cell 1 (discal 1) proportionally elongate, with proximodistal length ∼ 3 x anteroposterior width (CB50: 0.61). ***Polarity*:** Although equivocally supported for this node, observed in most sampled terminals of the Crabronidae–Anthophila clade; exceptions include *Plenoculus*, Ammoplanidae, and some Anthophila. Supported as independently derived in Vespida and Scolioidea.

**544.** Fore wing medial cell 2 (discal 2) with proximodistal length ≥ 2 x anteroposterior width (CB50: 0.86). ***Polarity*:** See comment for Total Ampulicidae node 3.

### **G.3.** Crabronidae–Sphecidae clade

***Clade comprising*:** • **Crabronidae** Latreille, 1802, • **Mellinidae** Latreille, 1802 (•• Mellini Latreille, 1802 [*Mellinus* Fabricius, 1790], •• Xenosphecini Parker, 1966 [*Xenosphex* Williams, 1954]), • **Sphecidae** Latreille, 1802.

***Note*:** Mellinidae was recovered in the Crabronidae–Sphecidae clade by Sann *et al*. (2018, 2021) but was not sampled in our study.

#### Synapomorphies

**241.** Metapleural area not scrobiform, *i.e.*, surface of metapleural area forming more-or-less flat, undisrupted surface with mesopectus (CB50: *). ***Polarity*:** Support for this condition is almost exactly equivocal. The “scrobiform” condition was observed in *Chalybion* and *Trypoxylon* of the Sphecidae and Crabronidae, respectively, suggesting a complicated pattern of transformation and indicating need for anatomical study. The non-scrobiform condition is supported as independently derived in *Sphecius* (Bembicidae) and Anthophila, with some reversals in the latter.

**284.** Female protarsus with a row of psammochaetae (CB50: 0.72). ***Polarity*:** Among sampled Crabronidae–Sphecidae, absent in *Chalybion* and *Trypoxylon*. Supported as independently derived in Bembicidae and the Pemphredonidae–Philanthidae clade.

**300.** Meso- and metatibiae with strong, scattered traction chaetae (CB50: 0.60). ***Polarity*:** Variable among sampled Crabronidae–Sphecidae: present in *Sphex*, *Prionyx*, *Oxybelus*, *Plenoculus*, and *Tachysphex*, absent in *Chalybion* and *Trypoxylon*.

**328.** Arolia enlarged, with an expanded unguitractor plate (CB50: 0.93). ***Polarity*:** Among sampled Crabronidae–Sphecidae, absent only in *Plenoculus*. Among other sampled Apoidea, observed to be present only in *Sphecius*.

**483.** Fore wing Rsf1 and Mf1 meeting at a distinct angle (CB50: 0.57). ***Polarity*:** Supported as an independent derivation of crown Ampulicidae and the Pemphredonidae–Anthophila clade. Observed in all sampled Crabronidae–Sphecidae with the exception of *Ammophila* and *Oxybelus*.

**531.** Fore wing crossvein 1m-cu distinctly curved or sinuate (CB50: 0.74). ***Polarity*:** Variable among Ampulicidae (see above); also observed in *Bembix*, *Cerceris*, and most sampled Anthophila.

**539.** Fore wing crossvein 2m-cu curved or sinuate (CB50: 0.73. ***Polarity*:** Also observed in some Ampulicidae, Bembicidae, *Philanthus*, and most Anthophila. Could very well be ancestral to the superfamily; should be tested with extended taxon sampling.

**568.** Hind wing Mf2 nebulous to absent (CB50: 0.61). ***Polarity*:** Among sampled Crabronidae– Sphecidae, only observed in *Chalybion*. Also absent in Pemphredonidae, Ammoplanidae, and some Anthophila.

***Note:*** Orientation of the compound eyes—whether orally-converging (char. 33), parallel, or posteriorly-converging (char. 34)—is variable among groups of Crabronidae and Sphecidae, and probably preserves important phylogenetic signal. Our sampling was too limited to confidently assert what the ancestral eye forms of the two subfamilies were, or to what degree they have converged.

### G.D.A. Family Crabronidae Latreille, 1802

***Clade comprising*:** • **Crabroninae** Latreille, 1802 (•• Crabronine clade [Crabronini Latreille, 1802, Oxybelini Leach, 1815], •• Larrine clade [Bothynostethini Fox, 1895, Gastrocericini André, 1886, Larrini Latreille, 1810, Miscophini* Fox, 1895 {*paraphyletic}, Palarini Schrottky, 1909, Trypoxylonini Lepeletier de Saint-Fargeau, 1845]), • **Dinetinae** Fox, 1895 (*Dinetus* Panzer, 1806). • ***Incertae sedis*** in family: †**Discoscapini** Poinar, 2020, **†Protomicroidini** Antropov, 2010.

***Note*:** The synapomorphies below should be considered with the reservation as Dinetinae were not sampled in the present study; this subfamily is supported as sister to Crabroninae *sensu stricto* (Sann *et al*. 2018, 2021). Focused effort to define the Crabronidae is necessary. In our study, the Crabronidae are supported as relatively plesiomorphic in comparison to Sphecidae. Most of the characters below are simple and some are coarse, thus should be considered “weak”. The strongest characters are the foreshortening of the mesocoxae and modification of the pretarsi, although these features should be evaluated with a broader taxon sampling.

#### Crabronidae synapomorphies

**14.** Head width ≤ 1 mm (CB50: 0.74). ***Polarity*:** This is a surprising result. All sampled Crabronidae with the exception of *Tachysphex* had small heads, hence small body size. We treat this reconstruction with reservation, as expanded taxon sampling is necessary.

**227.** Longitudinal mesopectal sulcus absent (CB50: 0.89). ***Polarity*:** Strongly supported as independently lost in *Heterogyna*, some Sphecidae, Crabronidae, Bembicidae, and Anthophila. Absent in all sampled Crabronidae.

**258.** Mesocoxae extremely foreshortened, thus hardly nor not at all produced away from the body (CB50: 0.96). ***Polarity*:** Supported as independently derived in *Sphecius*, Pemphredonidae + Ammoplanidae, and Anthophila. This feature deserves dedicated anatomical study.

**277.** “Basicostal carina” dorsoventrally elongate, approaching coxal apex (CB50: 0.90). ***Polarity*:** Among sampled Apoidea, supported as independently derived in Bembicidae, within Philanthidae, and within Anthophila.

**325.** Pretarsal claws very strongly reflexed, being in exact line with the apical tarsomere which itself is ventrally softened and covered with a plush layer of pubescence (CB50: 0.90). ***Polarity*:** Among sampled Crabronidae, not observed for *Plenoculus*; supported as a loss in this case. The form of the crabronid pretarsus is unique among all sampled Hymenoptera.

**569.** Hind wing Cuf absent (CB50: 0.71). ***Polarity*:** Absent in all sampled Crabronidae with the exception of *Tachysphex*, for which presence is supported as a reversal to the elongate condition. Hind wing C has also been lost in *Heterogyna* and *Ceratina*, among sampled extant Apoidea.

***Note*:** Four characters were recovered as synapomorphies of the third internal node of the Crabronidae in our analyses, *i.e.*, *Plenoculus* (Miscophini) plus *Tachysphex* (Larrini: Gastrosericina): (1) posterolateral mandibular margin with notch and tooth (char. **54**, CB50: 0.95); (2) female forefemur grossly swollen along its entire length (char. **280**, CB50: 0.94); (3) pterostigma strongly reduced (char. **459**, CB50: 0.92); and (4) fore wing cross vein 2r-rs anterior junction at distal apex of pterostigma (char. **470**, CB50: 0.87). See character definitions and raw data for state distributions among sampled taxa.

### G.F. Family Sphecidae Latreille, 1802

***Clade comprising*:** • **Sphecidae** Latreille, 1802 (•• Sphecine clade [Ammophilinae André, 1886, Sphecinae Latreille, 1802, Stangeellini Bohart & Menke, 1976 {*incertae sedis* to subfamily}], •• Sceliphrine clade [Chloriontinae Fernald, 1905, Sceliphrinae Ashmead, 1899]).

***Note*:** The synapomorphies below should be considered with the reservation that Mellinidae, which was recovered as sister to the Sphecidae by Sann *et al*. (2018). Note that while the crabronine *Ectemnius* was sister to *Ammophila* in Branstetter *et al*. (2017), Sann *et al*. (2018) found the genus to be nested within Crabronina, a result which we agree with based on morphological similarity. Overall, the Sphecidae are supported as relatively derived among Apoidea, particularly in comparison to Crabronidae.

#### Sphecidae synapomorphies

**138.** Antennal toruli located at approximate midlength of head (CB50: 0.98). ***Polarity*:** Supported as independently derived in Sphecidae, Bembicidae, Psenidae (at least *Pluto*), Philanthidae, and Anthophila. Cranial skeletomuscular anatomical data will contribute to resolving the polarity of this condition.

**152.** Scape subglobular in shape, being bulbous or swollen medially (CB50: 0.85). ***Polarity*:** Observed in *Chalybion*, *Sphex*, and *Prionyx*, but not *Ammophila*. As with other characters, expanded taxon sampling will be crucial.

**186.** Pronotum completely divided by a shallow, transverse sulcus, with the anterior pronotal portion narrowed anteromedially (CB50: 0.96). ***Polarity*:** Observed in all sampled sphecid species with the exception of *Prionyx*. Among sampled Apoidea, also observed in Ampulicidae and *Plenoculus*.

**187.** Pronotum without lateral scrobiform furrow (CB50: 0.95). ***Polarity*:** Supported as an independent origin with respect to *Sphecius* and the Pemphredonidae–Anthophila clade.

**246.** Metapleural area elongated such that upper metapleural area present as dorsoventrally aligned subtriangular area situated very distant dorsally from mesocoxa, with apex directed ventrad (CB50: 0.99). ***Polarity*:** Unique among sampled Apoidea and probably underlying meaningful skeletomuscular modifications.

**268.** Metasternal area forming dorsoventrally oriented rectangular sclerite which separates the metacoxal cavities from the propodeal foramen (CB50: 0.99). ***Polarity*:** As for char. 246, unique among sampled Apoidea.

**[310].** Mesotibia with two spurs (CB50: 0.95). ***Polarity*:** See comments for Crabronidae– Anthophila clade above. We are skeptical about this result based on our limited taxon sampling.

**355.** Propodeum with distinct but short tubular extension surrounding metasomal articulation (CB50: 0.96). ***Polarity*:** Another feature observed in all sampled Sphecidae with the exception of *Prionyx*.

**\*** Metasomal petiole (metasomal segment I) distinctly necked (“pedunculate”) (char. **370**, CB50: 0.89), this peduncle comprising the sternum only, with the tergum restricted to posterior portion of segment (char. **371**, CB50: 0.99). ***Polarity*:** A necked petiole was also observed in Ampulicidae and *Pluto* (Psenidae) but is extreme in form in Sphecidae (char. 371).

**390.** Metasomal petiole without a distinct laterotergite (CB50: 0.96). ***Polarity*:** Loss of the petiolar laterotergite is supported as independently derived in Sphecidae, Pemphredonidae– Ammoplanidae, and Anthophila, among sampled Apoidea.

**521.** Fore wing 1rrs cell (submarginal 1) prestigmal length greater than half total cell length (CB50: 0.92). ***Polarity*:** Among sampled Apoidea, also observed in various Crabronidae (*Oxybelus*, *Tachysphex*), Bembicidae, and some Anthophila.

**536.** Fore wing medial cell 1 (discal 1) relatively long, with anteroposterior length exceeding the maximum anteroposterior width of the wing (CB50: 0.99). ***Polarity*:** Among sampled Apoidea, also observed in Bembicidae and *Clypeadon* (Philanthidae).

### G.3’. Bembicidae–Anthophila clade

***Clade comprising*:** • **Astatidae** Lepeletier de Saint-Fargeau, 1845 (Astatinae: *Astata* Latreille, 1796, *Dryudella* Spinola, 1843, *Uniplectron* Parker, 1966; †Cretastatini Rosa & Melo, 2021: †*Cretastatus* Rosa & Melo, 2021), • **Entomosericidae** Dalla Torre, 1897 (Entomosericinae Dalla Torre, 1897: Entomosericini Dalla Torre, 1897) • **Eremiaspheciidae** Menke, 1967 (Eremiaspheciinae Menke, 1967: Eremiaspheciini Menke, 1967; Laphyragogini Bohart & Menke, 1976), • **Bembicidae** Latreille, 1802, • **Pemphredonidae–Anthophila** clade.

***Note*:** Bembicidae was recovered as sister to the Pemphredonidae–Anthophila clade with maximum support by Branstetter *et al*. (2017) but was weakly supported as sister to the Crabronidae–Sphecidae clade by Sann *et al*. (2018). Our results match the topology of the prior study, which is congruent with that of Sann *et al*. (2021). Astatidae (Astatinae) was supported as sister to Bembicidae + Pemphredonidae–Anthophila clade in the majority of analyses of Sann *et al*. (2021), so we place the family there, although we have not presently sampled any of its constituent taxa.

#### Synapomorphies

**138.** Antennal toruli situated at about head midlength (CB50: 0.81). ***Polarity*:** Supported as independently derived with respect to Sphecidae. Reversed in the sampled Ammoplanidae and Andrenidae, among sampled Anthophila.

**300.** Meso- and/or metatibiae with traction chaetae (CB50: *). ***Polarity*:** Support at the Bembicidae–Anthophila and Pemphredonidae–Anthophila nodes is equivocal (CB50: 0.53, 0.54), as well as for the Crabronidae–Anthophila node (CB50: 0.63). However, tibial traction chaetae are present in all extant spheciform members of the group, with loss strongly supported in Anthophila regardless of the condition at the two deeper nodes.

**310.** Mesotibia with a single ventroapical spur (CB50: 0.98). ***Polarity*:** As noted for more rootward nodes within the Apoidea, this is a complicated character to interpret due to limited taxon sampling. Whatever the ancestral condition of the second mesotibial spur, its absence is strongly supported at this node, and is important in the comprehensive treatment of spheciform Apoidea by Bohart & Menke (1976).

### G.J. Bembicidae Latreille, 1802

***Clade comprising*:** • **Bembicinae** Latreille, 1802 (Bembicini Latreille, 1802, Helocausini Handlirsch, 1925), • **Nyssoninae** Latreille, 1804 (Alyssontini Dalla Torre, 1897, Nyssonini Latreille, 1804). • ***Incertae sedis*** in family: †Megaroliini Rosa & Melo, 2021, †Pristinopterini Rosa & Melo, 2021.

#### Bembicidae synapomorphies

**79.** Clypeal disc bulbous and swollen, projecting anteriorly (assuming hypognathy) (CB50: 0.99). ***Polarity*:** This clypeal form was unique among examined taxa.

**88.** Anterolateral portions of clypeus broadly curved posteromedially around labral base (assuming hypognathy) (CB50: 0.99). ***Polarity*:** Observed in various Anthophila (see character definition and raw data for more information).

**92.** Anterior clypeal margin not convex or angular, rather being broadly concave (CB50: 0.92). ***Polarity*:** The broadly concave condition was not explicitly scored; the transformation from the ancestral convex condition is strongly supported, however.

**191.** Pronotum flush with the mesoscutum, without a posterior constriction just anterad the intersegmental articulation (CB50: 0.95). ***Polarity*:** Convergently derived in Anthophila. This condition effectively represents a reversal from the constricted form which defines, in part, the Apoidea.

**215.** “Oblique scutal carina” present, *i.e.*, a transverse to diagonal carina present on the posterolateral lobe of mesoscutum which overhangs the fore wing articulation (CB50: 0.98). ***Polarity*:** Unique among Apoidea (Bohart & Menke 1976). Among other Bembicidae, the oblique scutal carina is also known to be present in Gorytini (variable), Nyssonini, Stictiellini, and Stizini, while it is absent in Alyssonini, Heliocausini, †Megaroliini, and †Pristinopterini (Bohart & Menke 1976, Sann *et al*. 2018, Rosa & Melo 2021).

**227.** Longitudinal mesopectal sulcus absent (CB50: 0.93). ***Polarity*:** Supported as independently lost in *Heterogyna*, some Sphecidae, Crabronidae, Bembicidae, and Anthophila.

**247.** Upper and lower metapleural areas with more-or-less parallel anterior and posterior margins, such that the entire metapleural area is in the form of a dorsoventrally long strip (CB50: 0.98). ***Polarity*:** Among sampled Apoidea, also observed in *Heterogyna*, *Trypoxylon*, and †*Psolimena electra*.

**277.** “Basicostal carina” of fore coxa conspicuously elongate (CB50: 0.94). ***Polarity*:** Supported as independently derived within Apoidea for Crabronidae, some Philanthidae, and some Anthophila.

**459.** Pterostigma strongly reduced, being absent or virtually so (CB50: 0.89). ***Polarity*:** The pterostigma is also reduced in Crabronidae, Sphecidae, and various Anthophila.

**470.** Fore wing crossvein 2r-rs anterior junction situated at distal apex of pterostigma (CB50: 0.94). ***Polarity*:** As for char. 459 above: 2r-rs is also at the distal pterostigmal apex in some Crabronidae, some Sphecidae, and some Anthophila.

**483.** Fore wing Rsf1 and Mf1 more-or-less linear, not meeting at a distinct oblique angle (CB50: 0.93). ***Polarity*:** Among sampled Apoidea, also observed in *Ammophila*, *Oxybelus*, some Anthophila, and many fossils.

**486.** Fore wing Mf1 long to extremely long, length ≥ 4 x that of Rsf1 (CB50: 0.93). ***Polarity*:** Among sampled Apoidea, also observed in most Sphecidae, *Trypoxylon*, a few Anthophila, and †*Cretampulex*.

**495.** Fore wing Rsf2 perpendicular to the longitudinal axis of the fore wing (CB50: 0.97). ***Polarity*:** Among sampled Apoidea, also observed in *Trypoxylon*, a few Anthophila, and various fossils.

**519.** Fore wing crossvein 3rs-m situated close to apical wing margin, separated by less than one of its lengths (CB50: 0.92). ***Polarity*:** Among sampled Apoidea for which this condition is applicable, also observed in *Ampulex*, *Tachysphex*, Philanthidae, *Apis*, and various fossils.

**521.** Fore wing 1rrs cell (submarginal 1) prestigmal length greater than half total cell length (CB50: 0.87). ***Polarity*:** Among sampled Apoidea, also observed in Sphecidae, some Crabronidae, some Anthophila, and various fossils.

**\*** Fore wing medial cell 1 (discal 1) extremely long, with a proximodistal length exceeding the maximum anteroposterior width of the wing (char. **535**, CB50: 0.99) and which is > 2/5 the maximum proximodistal length of the wing (char. **536**, CB50: 0.99). ***Polarity*:** A first medial cell which is longer than the fore wing width is supported as a synapomorphy of Sphecidae, Bembicidae, and is also observed in *Clypeadon*. The extreme length relative to the fore wing length is unique among sampled Apoidea and is otherwise observed in Vespoidea.

**541.** Fore wing crossvein 2m-cu close to distal apex of wing, within one of its own lengths (CB50: 0.87). ***Polarity*:** Among sampled Apoidea, also observed in *Ampulex*, *Chalybion*, *Tachysphex*, some Philanthidae, *Apis*, and various compression fossils.

**548.** Fore wing crossvein 1cu-a anterior junction distinctly postfurcal, *i.e.*, distal to the split of M+Cu (CB50: 0.88). ***Polarity*:** Nearly unique among sampled Apoidea, being sporadically observed in other major groups (*Thyreus*, †*Cretotrigona*, two †*Pompilopterus* wings, and †*Trichobaissodes*).

**555.** Fore wing Cu1 and Cu2 diverging at approximate right angle (CB50: 0.90). ***Polarity*:** Among sampled Apoidea, also observed in *Ampulex*, Pemphredonidae, Ammoplanidae, some Philanthidae, some Anthophila, some amber fossils, and many compression fossils.

**556.** Fore wing Cuf2 and Cuf3 directed proximally from anterior juncture to posterior juncture (CB50: 0.98). ***Polarity*:** Nearly unique among sampled Apoidea, with the exception of *Clypeadon* and various fossils.

### **G.4.** Pemphredonidae–Anthophila clade

*Clade comprising*: • **Pemphredonidae–Philanthidae** clade, • clade **Anthophila** Latreille, 1804

#### Synapomorphies

**106.** Face with one or two longitudinally oriented “subantennal sutures” between posterior clypeal margin and torular rim (CB50: 0.82). ***Polarity*:** Not observed in *Ammoplanops* or *Pulverro*; otherwise apparently unique, given the present sample of Apoidea. Some ants (*e.g.*, Ectatomminae) also have clypeotorular lines. This condition is correlated with having the toruli separated from the posterior clypeal margin, although the clypeotorular lines are not always observed even when the toruli are distant, *e.g.*, in Camponotini. See char. 140 for Pemphredonidae–Philanthidae clade below for more details.

**149.** Radicle and scape linear, not offset at a distinct angle (CB50: 0.93). ***Polarity*:** Equivocal at the Bembicidae–Anthophila node as *Bembix* with linear scape and radicle, while *Sphecius* with these portions at distinct angle. Extended taxon sampling in Bembicidae will contribute to resolution of the polarity of this character.

**429.** Abdominal tergum VII of female and VIII of male (metasomal VI, VII) modified as “pygidial plate”, *i.e.*, produced posteriorly and/or marginate laterally, often with medial sculpturation (CB50: 0.81). ***Polarity*:** Supported as independently derived in Crabronidae (*Oxybelus*, *Tachysphex*), Bembicidae (*Sphecius*), and the tentative Pemphredonidae– Anthophila clade.

### **G.5.** Pemphredonidae–Philanthidae clade

***Clade comprising*:** • **Ammoplanidae** Evans, 1959, • **Pemphredonidae** Dahlbom, 1835 (Pemphredonini Dahlbom, 1835, Spilomenini Menke, 1989, Stigmina Bohart & Menke, 1976), • **Philanthidae** Latreille, 1802, • **Psenidae** Costa, 1858.

***Note 1*:** The placement of Ammoplanidae was uncertain at the time of our analyses; it was imperfectly supported as close to Pemphredonidae by Branstetter *et al*. (2017) and was well-supported as sister to Psenidae with the two forming the sister group of the Anthophila in Sann *et al*. (2018). We provisionally report morphological results from the Branstetter *et al*. (2017) topology. Additionally, we note that Psenidae was not included in our dataset.

***Note 2*:** We report results for the following nodes from our tree, recognizing the tentative relationship of Ammoplanidae as of 2020: Pemphredonidae–Philanthidae clade (“PePh clade”), and the putative Pemphredonidae–Ammoplanidae clade (“PeAm clade”). Sann *et al*. (2021) recently recovered Ammoplanidae as sister to the Anthophila.

#### Pemphredonidae–Philanthidae clade synapomorphies

**140.** Antennal toruli separated from clypeus by more than one torular diameter (CB50: 0.63). ***Polarity*:** Among sampled Apoidea, this condition was observed in *Ammophila*, *Pluto*, Philanthidae, and several Anthophila. Support stabilizes for the Philanthidae.

**279.** Protrochanter elongate, proximodistal length > 3 x lateromedial width (CB50: 0.86). ***Polarity*:** Observed in all sampled species of the tentative Pemphredonidae–Philanthidae clade, including Ammoplanidae and two of the fossil species, †*Prolemistus apiformis* and †*Psolimena dupeae*; other fossils without an elongate protrochanter, or condition uncertain.

**280.** Female fore femur grossly swollen (CB50: 0.88). ***Polarity*:** Also observed among various Crabronidae and Anthophila.

**284.** Female protarsus with a row of psammochaetae (fossorial chaetae) (CB50: 0.84). ***Polarity*:** Among sampled Apoidea, also observed in most Sphecidae, most Crabronidae, and Bembicidae; supported as independent derivations in each group.

#### Pemphredonidae–Ammoplanidae clade synapomorphies

**78.** Buccal cavity length ≤ width (CB50: 0.73). ***Polarity*:** The support for this condition is equivocal, and probably rightly so given our limited taxon sampling. Among sampled Apoidea, a short cavity is observed in Ampulicidae, *Heterogyna*, *Chalybion*, *Trypoxylon*, and the PeAm clade.

**263.** Mesocoxae wide-set (CB50: 0.74). ***Polarity*:** Among sampled Apoidea, also observed in *Sphecius* and *Apis*; uncertain for most fossils.

**390.** Abdominal segment II (metasomal petiole) without defined laterotergites (CB50: 0.84). ***Polarity*:** Also absent in most sampled Anthophila, with the exception of *Dieunomia*, and absent in *Heterogyna* and *Sphecidae*.

**488.** Fore wing Mf1 distinctly curved (CB50: 0.63). ***Polarity*:** Among sampled Apoidea, also observed in Ampulicidae, *Dieunomia*, *Megachile*, *Ceratina*, and a number of fossils attributed to the Pemphredonidae (see raw data for state distribution).

**568.** Hind wing Mf2 nebulous to absent (CB50: 0.61). ***Polarity*:** Among terminals of the Bembicidae–Anthophila clade, only observed in the tentative Pemphredonidae– Ammoplanidae clade and some Anthophila.

### G.M. Philanthidae Latreille, 1802

***Clade comprising*:** • **Aphilanthopini** Bohart, 1966, • **Cercerini** Lepeletier de Saint-Fargeau, 1845, • **Philanthini** Latreille, 1802, • **Pseudoscoliini** Menke, 1967.

***Note*:** We report results for the following nodes from our tree: Philanthidae node 1 (*Cerceris*, *Clypeadon*, *Philanthus*) and Philanthidae node 2 (*Clypeadon*, *Philanthus*). Eremiaspheciini was recently recognized as a family unplaced in the Bembicidae–Anthophila clade by Sann *et al*. (2021).

#### Philanthidae node 1 synapomorphies

**11.** Head relatively prognathous (CB50: 0.91). ***Polarity*:** Among sampled Apoidea, this condition was also observed in *Ampulex* and *Ceratina*. Support stabilizes as strong for Philanthidae node 2.

**22.** Occipital carina extending to hypostoma (CB50: 0.93). ***Polarity*:** Equivocal for the Pemphredonidae–Philanthidae node. Among sampled Apoidea, also observed in *Tachysphex* and *Pluto*; could not be evaluated for most fossils.

**69.** Prementum elongate, at proximodistal length ≥ 4 x lateromedial width (CB50: 0.98). ***Polarity*:** Supported as independently derived in Sphecidae, Crabronidae, Bembicidae, and Anthophila.

**102.** Lateral regions of male clypeus with dense brush of glandular setae (CB50: 0.98). ***Polarity*:** Nearly unique among all sampled taxa, with the exception of the sampled *Melitta*, although the forms are not identical.

**112.** Face broad and weakly convex across its entire surface (CB50: 0.98). ***Polarity*:** This strange facial form was only observed in Philanthidae.

**258.** Mesocoxae comparatively elongate, being projecting distinctly away from the body (CB50: 0.94). ***Polarity*:** This condition represents an elongation of the coxae; the foreshortened condition is strongly supported for the Anthophila (CB50: 0.95), the Pemphredonidae– Ammoplanidae node (Cb50: 0.93), and moderately so at the Pemphredonidae–Philanthidae and Pemphredonidae–Anthophila nodes (CB50: 0.80, 0.81, respectively). Foreshortened coxae are also observed in Crabronidae (CB50: 0.96), suggesting independent derivation.

**351.** Propodeal spiracle with anterior hood-like projection (CB50: 0.92). ***Polarity*:** Among sampled Apoidea, only observed in Philanthidae and *Thyreus*.

**355.** Propodeum with distinct but short tubular extension surrounding metasomal articulation (CB50: 0.98). ***Polarity*:** Supported as independently derived in Sphecidae, at least once in Crabronidae (*Trypoxylon*), and Philanthidae.

**385.** Abdominal tergum II (metasomal petiole) with one or two anterodorsal carinae (CB50: 0.89). ***Polarity*:** Among sampled Apoidea, also observed in *Dolichurus*, *Oxybelus*, and *Plenoculus*; supported as independent derivations for each.

**400.** Abdominal segment III (metasomal II) with sulcus dividing anterior contact surface (“presclerite”) and posterior non-contact surface (“postsclerite”) (CB50: 0.98). ***Polarity*:** Among sampled Apoidea, also observed in Ampulicidae, *Tachysphex*, *Sphecius*, and most Anthophila; uncertain for anthophilan and most other fossils.

**418.** Abdominal segment IV (metasomal III) with sulcus dividing anterior contact surface (“presclerite”) from posterior non-contact surface (“postsclerite”) (CB50: 0.95). ***Polarity*:** Supported as independently derived in Ampulicidae (*Dolichurus*), Crabronidae (*Tachysphex*), Philanthidae, and Anthophila. Sulci of abdominal segments V (char. 425) and VI (char. 426) are also observed in Philanthidae, and rarely in some ants (*e.g.*, *Sphinctomyrmex*).

**494.** Fore wing Rsf between Rs+M and 2r-rs kinked or distinctly curved (CB50: 0.85). ***Polarity*:** Among sampled Apoidea, also observed in *Dolichurus*, *Bembix*, and various Anthophila. See raw data for fossil distribution.

**519.** Fore wing crossvein 3rs-m situated close to apical wing margin, separated from the margin by less than one of its lengths (CB50: 0.91). ***Polarity*:** Among sampled Apoidea for which 3rs-m is present observed in *Ampulex*, *Tachysphex*, Bembicidae, *Apis*, and various fossils.

#### Philanthidae node 2 synapomorphies

**33.** Compound eyes converging posteriorly to some degree (CB50: 0.90). ***Polarity*:** Observed sporadically among sampled Apoidea, including Ampulicidae, various Crabronidae, *Eufriesea*, *Apis*, and †*Cretotrigona*; condition among fossils variable.

**108.** Face with a single or medial “subantennal suture” which contacts the torulus medially (CB50: 0.97). ***Polarity*:** Also observed in the sampled Andrenidae, Colletidae, Halictidae, Euglossini, and Apini.

**193.** Pronotum posterior margin notched medially (CB50: 0.97). ***Polarity*:** Among sampled Apoidea, also observed in *Dieunomia*.

**277.** “Basicostal carina” of fore coxa elongate (CB50: 0.91). ***Polarity*:** Elongation of the posterolateral carina of the fore coxa is supported as independently derived among Apoidea in Crabronidae, Bembicidae, Philanthidae node 2, and is also observed in various Anthophila.

**294.** Mesotrochantellus distinct and well-developed (CB50: 0.83). ***Polarity*:** Among sampled Apoidea, also observed in *Chalybion* and *Bembix*.

**329.** Tarsomeres I–IV with apicoventral lobate seta (CB50: 0.89). ***Polarity*:** Observed sporadically among other Apoidea, including *Dolichurus*, *Chalybion*, and *Ammophila*; strongly supported as independently derived in these groups.

### **G.6.** Clade Anthophila Latreille, 1804

***Clade comprising*: †Melittosphecidae** Poinar & Danforth, 2006, †**Paleomelittidae** Engel, 2001, **Andrenidae** Latreille, 1802, **Apidae** Latreille, 1802, **Colletidae** Lepeletier de Saint-Fargeau, 1841, **Halictidae** Thomson, 1869, **Megachilidae** Latreille, 1802, **Melittidae** Schenck, 1860, **Stenotritidae** Cockerell, 1934.

***Note*:** The two extinct Anthophila included in the ancestral state estimation analyses were recovered on the stem of the group. For this reason, results are reported for Anthophila node 1 (including fossils), and for Anthophila node 2 (excluding fossils). Our intent is to understand how the tested characters behaved, as well as to capture information about reconstruction certainty. An interpretation of the ancestral form of the total Anthophila should probably integrate features at both nodes. Reference is made to Michener (2007) for those characters which he adduced as support for the monophyly of Anthophila.

#### Anthophila node 1 synapomorphies

**7.** Plumose setae present somewhere on body (CB50: 1.0). ***Polarity*:** Observed in very few species of Formicidae (not sampled) and Mutillidae. The plumose setae of †*Melittosphex* are apparently present at least on the hind leg (Danforth & Poinar 2011). Plumose setae on abdominal tergum II (char. **387**) is also supported as a synapomorphy of the Anthophila (CB50: 0.99). Plumose setae are “character a” of Michener (2007) for the monophyly of Anthophila.

**154.** Scape relatively elongate, with proximodistal length ≥ 4 x width (CB50: 0.4). ***Polarity*:** Only some sampled Anthophila had shorter scapes; these include *Dieunomia*, *Megachile*, and *Centris*. Both †*Melittosphex* and †*Cretotrigona* were observed to have relatively long scapes. Outside of the Anthophila, long scapes were also observed in *Ammophila* and *Bembix*.

**228.** Dorsoventral mesopectal sulcus in anterior half of sclerite absent (CB50: 0.80). ***Polarity*:** Presence is strongly supported at the Bembicidae–Anthophila node (CB50: 0.92) and moderately so at the Pemphredonidae–Anthophila node (CB50: 0.88). Absence of the longitudinal mesopectal sulcus is strongly supported for the Anthophila (char. **227**), but the polarity is uncertain as the reconstruction is equivocal for both of the higher nodes. See definitions and the raw data for character state distributions.

**300.** Meso- and/or metatibiae without traction chaetae (CB50: 0.96). ***Polarity*:** Strongly supported as a synapomorphic loss within the Pemphredonidae–Anthophila clade. Adduced as “character j” of Michener (2007) for the monophyly of Anthophila.

**318.** Metabasitarsus anteroposteriorly flattened to some degree (CB50: 0.91). ***Polarity*:** Flattening of the metabasitarsus is not observed in all Anthophila but is nevertheless supported as a synapomorphy for the clade. This condition was also observed in †*Melittosphex* and †*Cretotrigona*. Expanded metabasitarsomeres are “character c” of Michener (2007) for the monophyly of Anthophila.

**324.** Pretarsal claw with at least one median or subapical tooth present (CB50: 0.89). ***Polarity*:** Among sampled Apoidea, also observed in Ampulicidae and some Sphecidae, strongly indicating independent origin. See definition and raw data for further information on state distribution outside of the superfamily. Bidentate pretarsal claws are “character m” of Michener (2007) for the monophyly of Anthophila.

**390.** Abdominal segment II (metasomal petiole) without defined laterotergites (CB50: 0.84). ***Polarity*:** Also absent in Ammoplanidae, extant Pemphredonidae (uncertain for fossils), *Heterogyna*, and *Sphecidae*. Among sampled Anthophila, laterotergites were observed in *Dieunomia*.

**400.** Abdominal segment III (metasomal II) with sulcus dividing contact surface (“presclerite”) and non-contact surface (“postsclerite”) (CB50: 0.69). ***Polarity*:** Among sampled Apoidea, also observed in Ampulicidae, *Tachysphex*, *Sphecius*, and Philanthidae; uncertain for anthophilan and most other fossils.

#### Anthophila node 2 synapomorphies

**21.** Occipital carina absent (CB50: 0.92). ***Polarity*:** Among sampled Anthophila, an occipital carina was only observed for *Thyreus*; the condition in the fossils is uncertain. Among the spheciform Apoidea, the carina was uniquely observed to be absent in Ammoplanidae.

**93.** Anterior clypeal margin more-or-less linear (CB50: 0.97). ***Polarity*:** Observed for all sampled Anthophila; uncertain for the fossil terminals.

**191.** Pronotum not constricted posteriorly (CB50: 0.98). ***Polarity*:** This simplification of the pronotum is supported as independently derived in Bembicidae. Uncertain for the fossils, for which reason this condition is equivocally supported at node 1.

**192.** Pronotum flush with mesoscutum, not separated by a transverse notch (CB50: 0.99). ***Polarity*:** Also observed as an independent derivation in *Cerceris* (Philanthidae). Uncertain for the fossils, thus equivocal at node 1.

**241.** Metapleural area not scrobiform, *i.e.*, with metapleural area not directed posteriorly (CB50: 0.78). ***Polarity*:** A scrobiform metapleural area is supported as regained within the Anthophila, being observed in *Thyreus*, *Centris*, and *Eufriesea*. The non-scrobiform condition was observed in most Sphecidae, most Crabronidae, and *Sphecius*.

**260.** Ventral lobes of mesopectus reduced (CB50: 0.93). ***Polarity*:** Could not be observed in fossil taxa, and for *Dasypoda* and *Melitta*. The ventral mesopectal lobes are supported as lateromedially expanded for the Crabronidae–Anthophila clade (see above).

**295.** Female meso- and/or metafemora swollen in appearance (CB50: 0.93). ***Polarity*:** Among sampled Anthophila, the femora are thick in all species but for *Apis* and the two sampled fossils.

**343.** Propodeum without a dorsal face extending posterad beyond the metanotum in profile view (CB50: 0.97). ***Polarity*:** Observed uniquely in Anthophila among sampled Apoidea. The propodeum of *Dieunomia* is strongly supported as secondarily elongate.

**386.** Abdominal tergum II (metasomal I) with distinct anterolaterally-situated and dorsoventrally oriented margins which border an anteromedian cavity that fits against the propodeum when the metasoma is flexed dorsad (CB50: 0.99). ***Polarity*:** Autapomorphic among sampled Apoidea. Could not be confirmed for fossil terminals.

**390.** Abdominal segment II (metasomal petiole) without defined laterotergites (CB50: 0.98). ***Polarity*:** Uncertain for the fossil terminals. Supported as independently derived for Pemphredonidae + Ammoplanidae (should this be confirmed as a clade), Sphecidae, and Heterogynaidae.

**401.** Abdominal segment III (metasomal II) without transverse sulcus (“gradulus”) separating anterior contact surface (“presternite”) from posterior non-contact surface (“poststernite”) (CB50: 0.95). ***Polarity*:** Sporadically derived among various Vespida and reversed to the “present” condition in *Thyreus* and *Eufriesea*, among sampled Anthophila.

**418.** Abdominal segment IV (metasomal III) with transverse sulcus (“gradulus”) dividing anterior contact surface (“presclerite”) from posterior non-contact surface (“postsclerite”) (CB50: 0.95). ***Polarity*:** Supported as independently derived in Ampulicidae (*Dolichurus*), Crabronidae (*Tachysphex*), Philanthidae, and Anthophila.

**425.** Abdominal segment V (metasomal IV) with transverse sulcus (“gradulus”) dividing anterior contact surface (“presclerite”) from posterior non-contact surface (“postsclerite”) (CB50: 0.80). ***Polarity*:** Independently derived in Philanthidae and *Tachysphex*, among sampled Apoidea. Uncertain for the fossils and *Dasypoda*, *Thyreus*, and *Eufriesea*; absent in *Megachile* and *Apis*. There is equivocal support for a transverse sulcus on segment VI, the difference in scoring being that *Centris* is uncertain for this line.

**512.** Fore wing crossvein 2rs-m sinuate (CB50: 0.93). ***Polarity*:** Observed in most Anthophila, with the exceptions of *Dieunomia* and *Apis*. Among sampled spheciform Apoidea for which 2rs-m is present, also observed for *Tachysphex* and *Sphecius*. Not observed for †*Melittosphex*; not applicable for †*Cretotrigona*; observed for “Crato Apidae” (see raw data).

**562.** Hind wing C absent (CB50: 0.96). ***Polarity*:** Hind wing C was observed for *Caupolicana*, among sampled Anthophila. Present for most spheciform Apoidea with the exceptions of *Dolichurus*, *Oxybelus*, *Plenoculus*, and *Pluto*.

***Note*:** Curvature of the anterolateral clypeal portions around the labral base was equivocally supported as a synapomorphy of the Anthophila (char. **88**, CB50: 0.55); this condition is variable among the bees and may have arisen more than once.

## Part V: Character and State Definitions

### Overview

Part V of this article is divided into four sections. The first two adduce the behavioral characters we developed for the modeling of eusocial evolution, the third outlines the transition models, and the fourth includes all morphological characters and their corresponding illustrations. To ease reference to the character state sets analyzed in the present study, we have organized these as tables. Note that the tables of behavioral state definitions include multistate (nonbinary) variables, in distinction to the morphological data used for topology searches. These multistate (nonbinary) variables can be considered aggregates of several observed states. See these sections for further information.

### Section 1: Independent behavioral characters

The two variables listed here, **A** and **B**, are behavioral traits which were evaluated independently from one another and the 576 morphological character matrix above. In other words, these characters were used for ancestral state reconstruction, but not for topological searches.

**Table 7.**
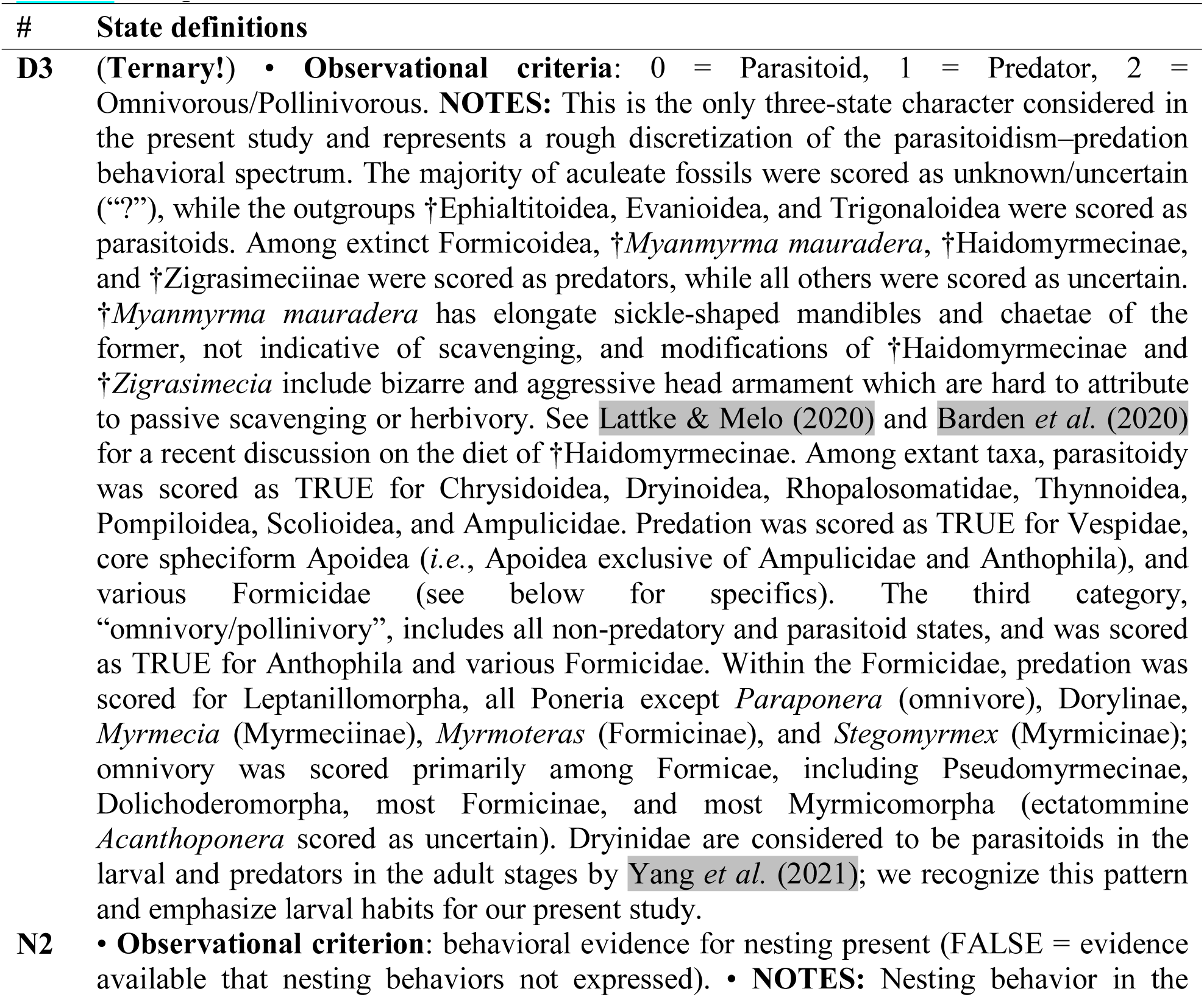
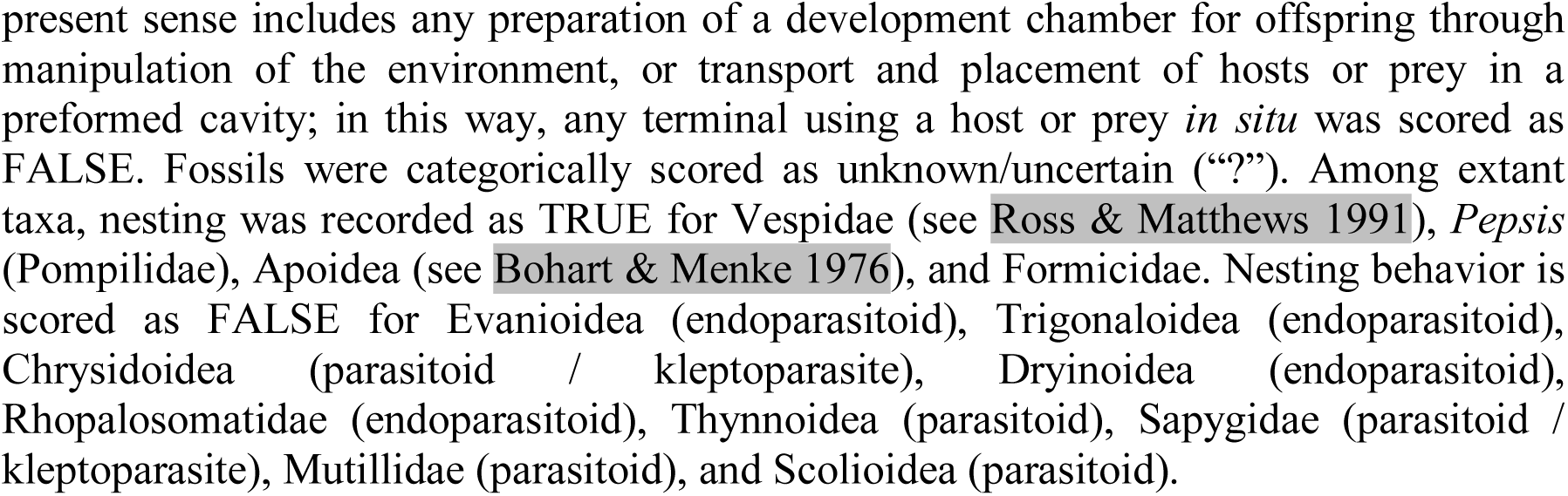
Independent behavioral characters.

### Section 2: Social behavior supercharacter

The social behavior “supercharacter” (**S**) comprises twelve “superstates”, themselves composed of alternative combinations of three “substates” (###). The substates describe dietary (X##), nesting (#X#), and social behaviors (##X); the superstates simplify meaningful combinations of substates. Five superstates are considered in our model; excluded substate combinations are listed after the superstate definitions. Scoring rationales for substates and evidence sources are listed after state definitions. A pair of alternative instantaneous (Q) rate matrices were averaged via reversible jump MCMC and are listed at the end of this section. We follow Boomsma & Gawne (2018) in defining eusociality as a comparatively discrete variable. Specifically, we recognize the distinction between *facultative* and *obligate eusociality*, with the latter category applicable to species with irreversible worker- and reproductive-caste fates.

**Table 8.**
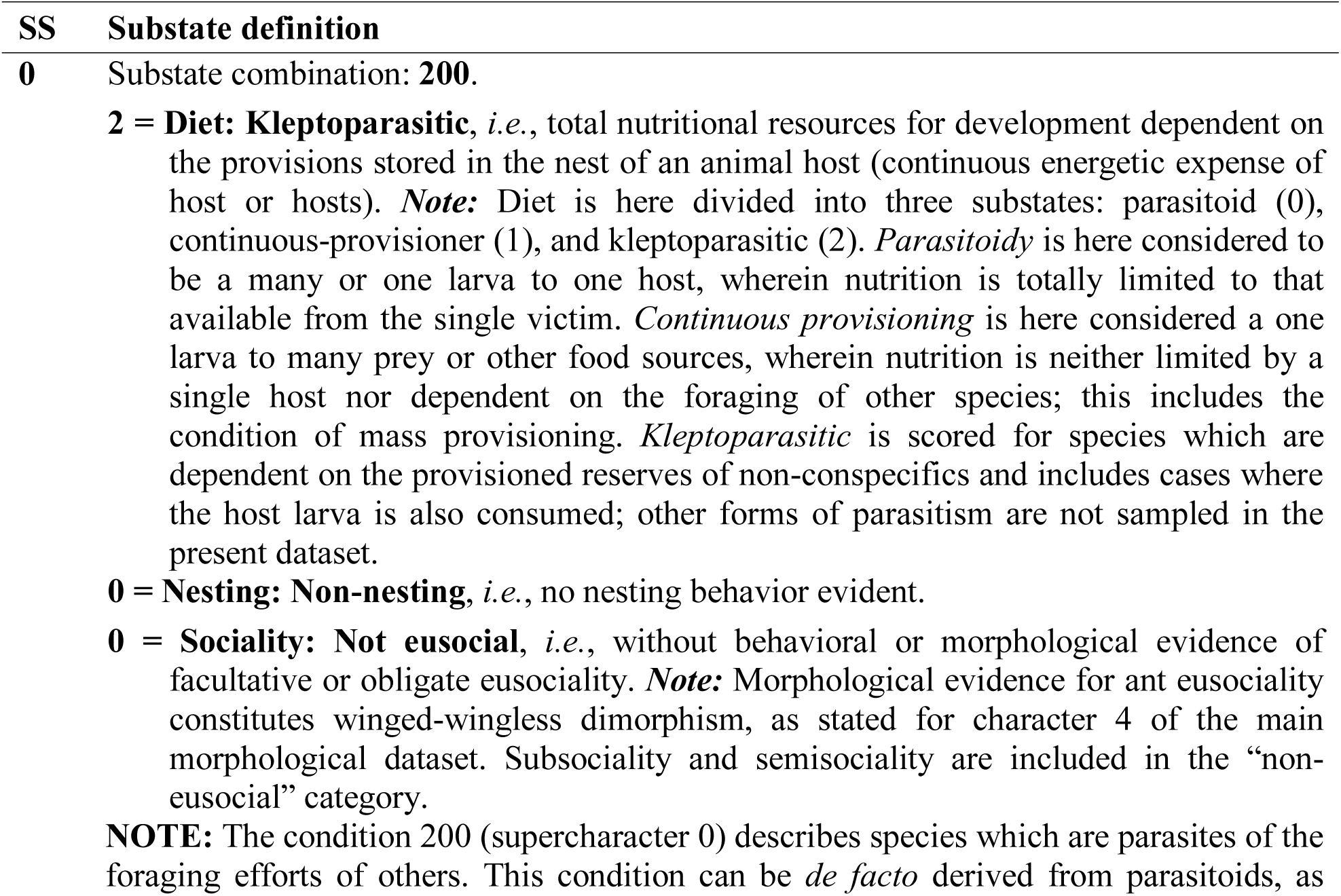
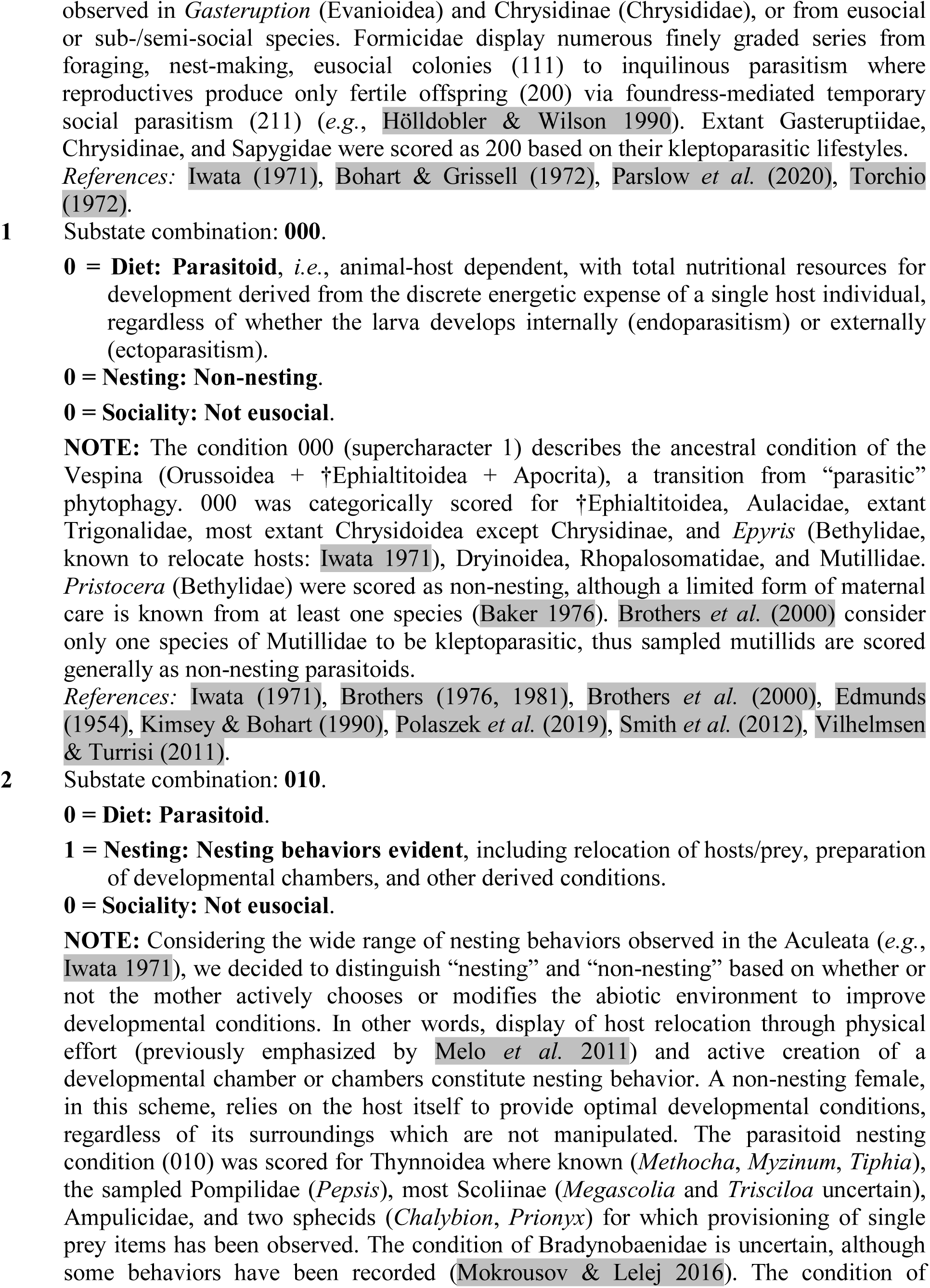
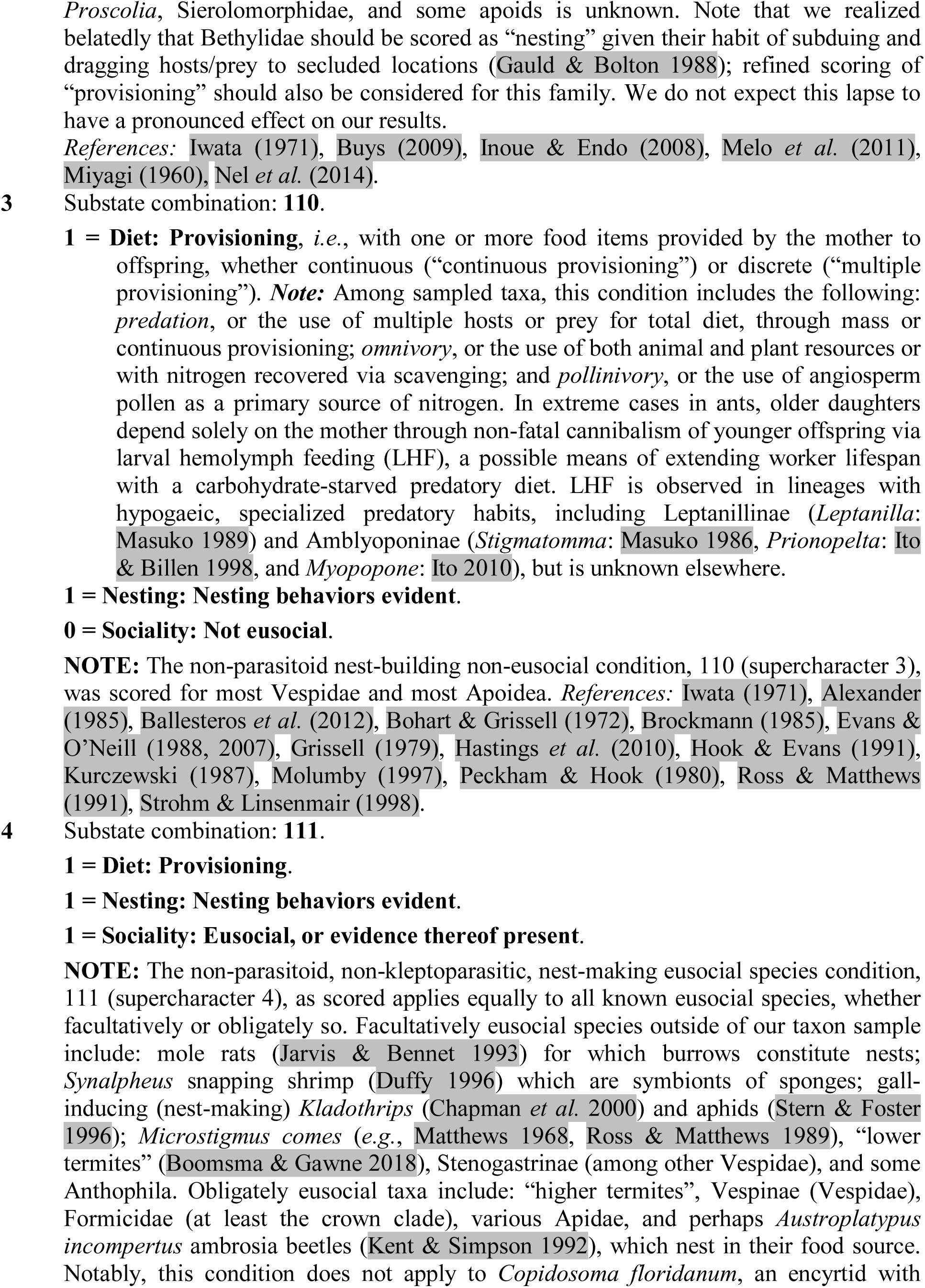
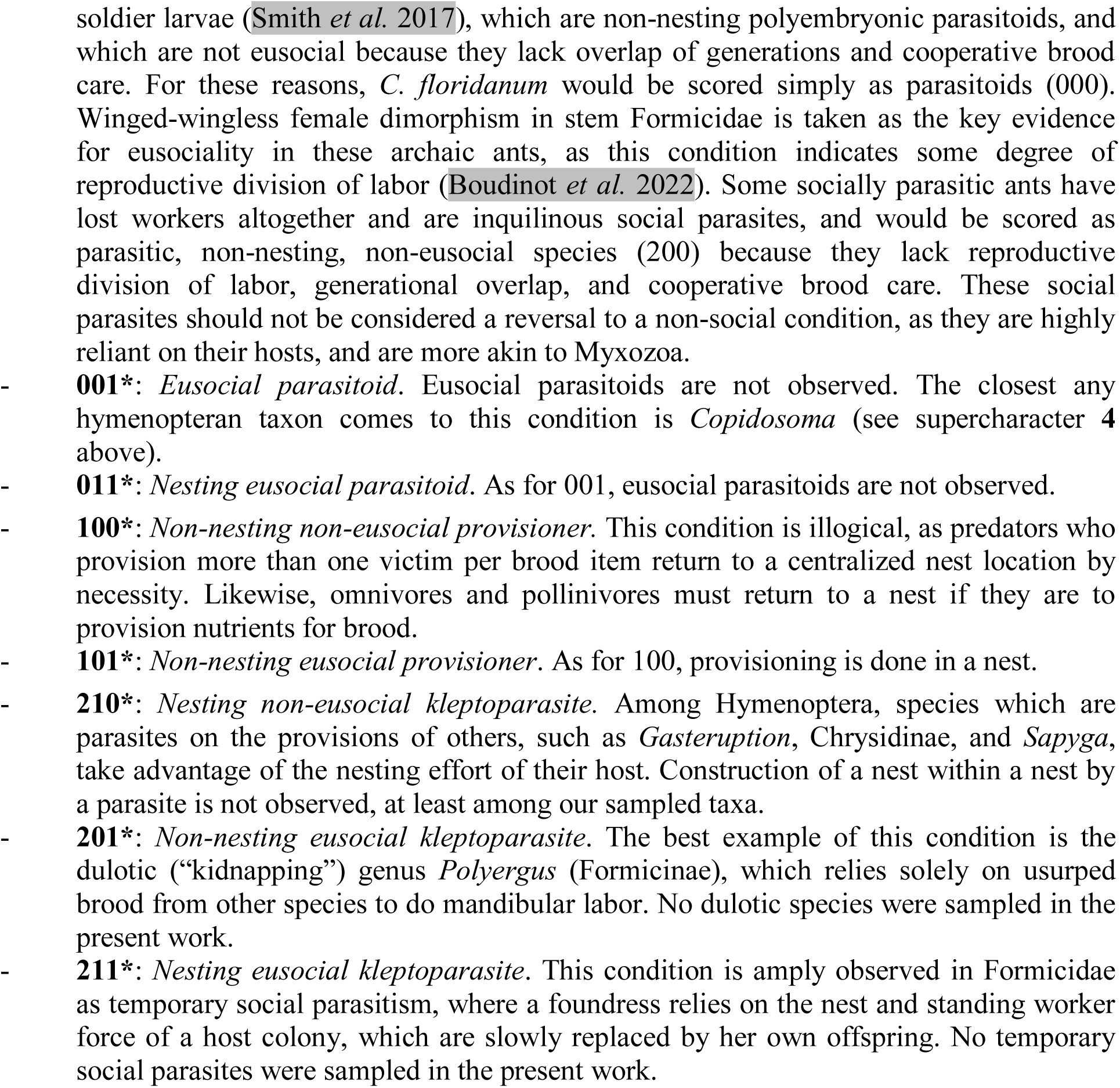
The social behavior supercharacter. SS = “superstate”; * = condition not observed.

### Section 3: Transition models

#### Complex transition model

We derived our instantaneous rate matrix for a five-superstate dataset (**5S**) from a base matrix including all twelve superstates (**12S**). In our models, each superstate transition proceeds stepwise by transition of one of the substates, analogous to the codon model for nucleotides. Rate variable codes for the two models are as follows: r = rate; odd numbers = forward transitions; even numbers = reverse transitions; suffix letters = linear transition sequences to eusociality. If rates are equal for any numbered rate series, then transitions among these are independent on the node-specific state; when such rates are unequal, transitions are state-dependent. Below is the complete 12-superstate matrix (**12S**) with rate variables indicated for each possible single-substate-step transition.

**Table 9.** Total Q matrix, showing all possible transition combinations (**12S**).

| - | 000 | 001 | 010 | 011 | 100 | 101 | 110 | 111 | 200 | 201 | 210 | 211 |
| --- | --- | --- | --- | --- | --- | --- | --- | --- | --- | --- | --- | --- |
| 000 | - | **r9a** | **r7a** | 0.0 | **r1a** | 0.0 | 0.0 | 0.0 | **r3a** | 0.0 | 0.0 | 0.0 |
| 001 | **r10a** | - | 0.0 | **r7b** | 0.0 | **r1b** | 0.0 | 0.0 | 0.0 | **r3b** | 0.0 | 0.0 |
| 010 | **r8a** | 0.0 | - | **r9b** | 0.0 | 0.0 | **r1c** | 0.0 | 0.0 | 0.0 | **r3c** | 0.0 |
| 011 | 0.0 | **r8b** | **r10b** | - | 0.0 | 0.0 | 0.0 | **r1d** | 0.0 | 0.0 | 0.0 | **r3d** |
| 100 | **r2a** | 0.0 | 0.0 | 0.0 | - | **r9c** | **r7c** | 0.0 | **r5a** | 0.0 | 0.0 | 0.0 |
| 101 | 0.0 | **r2b** | 0.0 | 0.0 | **r10c** | - | 0.0 | **r7d** | 0.0 | **r5b** | 0.0 | 0.0 |
| 110 | 0.0 | 0.0 | **r2c** | 0.0 | **r8c** | 0.0 | - | **r9d** | 0.0 | 0.0 | **r5c** | 0.0 |
| 111 | 0.0 | 0.0 | 0.0 | **r2d** | 0.0 | **r8d** | **r10d** | - | 0.0 | 0.0 | 0.0 | **r5d** |
| 200 | **r4a** | 0.0 | 0.0 | 0.0 | **r6a** | 0.0 | 0.0 | 0.0 | - | **r9e** | **r7e** | 0.0 |
| 201 | 0.0 | **r4b** | 0.0 | 0.0 | 0.0 | **r6b** | 0.0 | 0.0 | **r10e** | - | 0.0 | **r7f** |
| 210 | 0.0 | 0.0 | **r4c** | 0.0 | 0.0 | 0.0 | **r6c** | 0.0 | **r8e** | 0.0 | - | **r9f** |
| 211 | 0.0 | 0.0 | 0.0 | **r4d** | 0.0 | 0.0 | 0.0 | **r6d** | 0.0 | **r8f** | **r10f** | - |

#### Simplified transition model

In order to obtain an empirical set of superstates, seven impossible or unobserved superstates were removed (see above). Therefore, the following ordered substate transitions could generally occur in our **5S** empirical dataset: 200 (*kleptoparasitism*) ←→ 000 (*parasitoidy*) ←→ 010 (*nesting parasitoidy*) ←→ 110 (*progressive provisioning*) ←→ 111 (*eusociality*). To allow for transitions outside of this ordered series, we added “jump” rates which allow transitions from parasitoidy directly to eusociality (**rS1**: 000 → 111) and from nesting parasitoidy directly to eusociality (**rS2**: 010 → 111). The reverse rates (rn1, rn2) were disallowed (*i.e.*, set to 0.0) as no apparent reversals (losses of eusociality) were sampled in our matrix. We used reversible jump MCMC to average over the possible rate matrices, either disallowing a given transition (rate set to a constant of 0.0) or allowing a given transition with a rate drawn from a (1,1) gamma distribution. These alternatively modeled transitions include reversal from *parasitoidy* to *progressive provisioning* (**r2c**: 110 → 010), reversal from *nesting* to *non-nesting in parasitoids* (**r8a**: 010 → 000), and the two “jump” rates (**rS1**, **rS2**). To each of these four transitions, we applied a 50:50 reversible jump mixture prior on the sampling of the disallowed or gamma rates. All transitions are state-dependent in this matrix, *i.e.*, it is fully ordered, although independence can be achieved if two sampled rates are equal. Rate codes are implemented as for the base **12S** matrix.

**Table 10.** Reduced matrix (**5S**), showing only those transitions which are empirically observed in our dataset. This matrix was used for the “extreme stepwise” analysis.

| - | 200 | 000 | 010 | 110 | 111 |
| --- | --- | --- | --- | --- | --- |
| 200 | - | **r3a** | 0 | 0 | 0 |
| 000 | **r4a** | - | **r7a** | 0 | **rS1** |
| 010 | 0 | **r8a** | - | **r1c** | **rS2** |
| 110 | 0 | 0 | **r2c** | - | **r9d** |
| 111 | 0 | **rn1** | **rn2** | **r10d** | - |

**Table 11.**
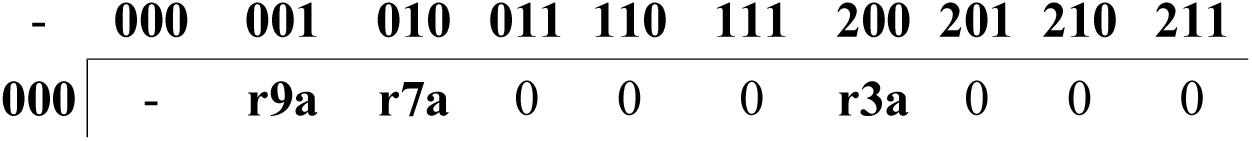
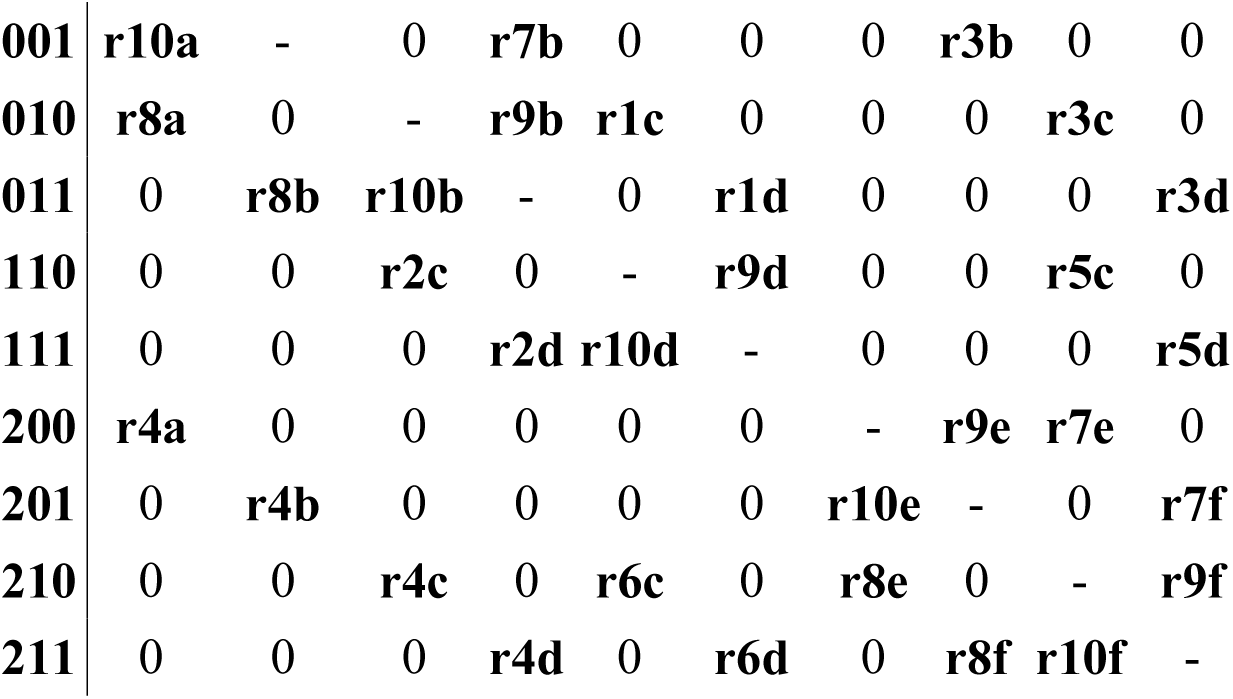
Reduced matrix (**10S**), showing all allowed transitions when simultaneous substate transitions are disallowed. States 100 and 101 are excluded.

| - | 000 | 001 | 010 | 011 | 110 | 111 | 200 | 201 | 210 | 211 |
| --- | --- | --- | --- | --- | --- | --- | --- | --- | --- | --- |
| 000 | - | **r9a** | **r7a** | 0 | 0 | 0 | **r3a** | 0 | 0 | 0 |
| **001** | **r10a** | - | 0 | **r7b** | 0 | 0 | 0 | **r3b** | 0 | 0 |
| **010** | **r8a** | 0 | - | **r9b** | **r1c** | 0 | 0 | 0 | **r3c** | 0 |
| **011** | 0 | **r8b** | **r10b** | - | 0 | **r1d** | 0 | 0 | 0 | **r3d** |
| **110** | 0 | 0 | **r2c** | 0 | - | **r9d** | 0 | 0 | **r5c** | 0 |
| **111** | 0 | 0 | 0 | **r2d** | **r10d** | - | 0 | 0 | 0 | **r5d** |
| **200** | **r4a** | 0 | 0 | 0 | 0 | 0 | - | **r9e** | **r7e** | 0 |
| **201** | 0 | **r4b** | 0 | 0 | 0 | 0 | **r10e** | - | 0 | **r7f** |
| **210** | 0 | 0 | **r4c** | 0 | **r6c** | 0 | **r8e** | 0 | - | **r9f** |
| **211** | 0 | 0 | 0 | **r4d** | 0 | **r6d** | 0 | **r8f** | **r10f** | - |

### Section 4: Morphological characters

**Table 12.** Character and state definitions used to score observational data of comparative morphology. Abbreviations: DC = developmental character; DS = developmental state. Variations of the conceptual type of definitions are indicated parenthetically (*e.g.*, “additive”, “reductive”, *etc.*). Certain body regions specified along the body axes are treated as DCs because body axes are specified during development (*e.g.*, “aboral” and “oral” regions of head). Note that the organization of the definitions is imperfect, but rigid because of the need to interpret the ancestral state estimation results. Of the 576 character-states, all but 15 are illustrated sequentially in Figs. C1–66, with each panel indexed by its character number below. # Definition - General

1 *DC: Sociality. DS: Superorganismality*. **• Observational criterion:** Behavioral evidence for (a) reproductive division of labor, (b) overlap of generations, and (c) cooperative brood care present (*FALSE* = evidence available that one or more of these conditions are not derived). **• NOTES:** The Major Transitions framework (Bourke 2011; Boomsma & Gawne, 2018) poses (b) and (c) as prerequisite conditions for the evolution of obligate eusociality, or “superorganismality”, *i.e.*, differentiation of reproductive (germline) and non-reproductive (soma) individuals. The present state is scored as *TRUE* when evidence for all three conditions is available. Among sampled taxa, obligate eusociality is evinced for various Vespidae, all Formicidae, and various Anthophila. Arguably, socially parasitic inquiline ants which have lost workers are no longer superorganismal, but these have neither reverted to an ancestral condition, nor were such species sampled in the present study. All fossils were scored as unknown (“?”) except for definitive representatives of crown Formicidae. See character 4 below for within-sex morphological differentiation.
2 *DC: Wing development. DS: Wing expression*. **• Observational criterion:** Apterous individuals of any sex present (*FALSE* = apterous individuals not observed). **• NOTES:** Observed in Chrysidoidea (Plumariidae, Bethylidae), Dryinoidea (Dryinidae, Sclerogibbidae), Tiphiiformes, Pompiloidea *sensu stricto* (Myrmosidae–Mutillidae), Scolioidea (Bradynobaenidae *sensu stricto)*, Formicoidea (apterous females unknown in †*Camelomecia*, “†Armaniidae”), Apoidea (Heterogynaidae); scored as “?” for fossils where apterous individual are unknown; *Olixon* (Rhopalosomatidae) scored for apterous and alate characters due to its intermediate, brachypterous, morphology. Apterous males are known in various crown Formicidae, including *Hypoponera* (Ponerinae), *Technomyrmex* (Dolichoderinae), and *Cardiocondyla* (Myrmicinae), among other genera.
3 *DC: Female polyphenism*. *DS: Extreme size polyphenism*. **• Observational criterion:** Apterous females with pronounced size polyphenism, resulting in presence of a “soldier” or major worker caste (*FALSE* = apterous females observed to be monomorphic). **• NOTES:** Observed sporadically among Formicidae, including *Pheidole* and *Cephalotes*; in the present sampling, observed in *Gesomyrmex*, *Oecophylla*, and *Camponotus. Pheidole* and *Cephalotes* have discrete “soldiers” (Rajakumar *et al*. 2018), whereas *Oecophylla* have an extreme size variation.
4 *DC: Female polyphenism*. *DS: Female winged-wingless diphenism*. **• Observational criterion:** Winged and wingless phenotypes of closely related individuals present (*FALSE* = such diphenism not observed). **• NOTES:** Because behavior cannot be scored based on direct observation in fossils, this state defines the morphological proxy for reproductive division of labor and brood care (at least via foraging) as observed in Formicidae. However, we recognize that wingless reproductive females can coexist with winged females within the same colony among most lineages of ants (Peeters 2012). Winged-wingless dimorphism is not observed in other Aculeata. For fossils, scored as *TRUE* for †Haidomyrmecini, †Zigrasimeciini, †Sphecomymini, †Brownimeciinae. Loss of winged queens has occurred sporadically among the crown Formicidae.
5 *DC: Soma. DS: Specific development of dorsoventral body compression*. **• Observational criterion:** Body dorsoventrally flattened to extremely dorsoventrally flattened (*FALSE* = body not observed to be compressed). **• NOTES:** Observed in Bethylidae. Flattening does occur in other taxa, such as certain *Cephalotes* ants and the Miocene †*Palaeobethylus* chrysidids.
6 *DC: Glands and glandular armament. DS: Glands with specific armament expressed.* **• Observational criterion:** White, silky looking glandular tufts of setae (“appeasement glands”) present anywhere on body (*FALSE* = such glands not observed). **• NOTES:** Such glands are observed in symphilic myrmecophiles. Among sampled species, *Loboscelidia fulgens* (Chrysididae: Loboscelidiinae) is the only taxon wherein these glandular structures are present; in this species, appeasement glands occur on the head posterolaterally, anterolaterally on the pronotum, the posterior faces of the coxae, the propodeum just above and lateral to the metasomal articulation, as well as the metasomal peduncle.
7 *DC: “Setoid” sensilla. DS: Specific sensilla form.* **• Observational criterion:** Plumose setae present on body (*FALSE* = such setal form not observed). **• NOTES:** Sensilla nomenclature is imperfect, with conflict occurring at the level of ontologies (see, *e.g.*, definitions for different sensilla in the Hymenoptera and *Drosophila* Anatomy Ontologies). To partially remedy this situation, here we refer to “setoid” sensilla for thin hair-like sensilla. In the original analyses of the present work, including ancestral state estimation, we hypothesized primary homology for the plumose setae of †*Melittosphex* Danforth & Poinar (2011), Poinar & Danforth (2006). We differentiated this condition from that of the “brachyplumose” state of sphaerophthalmine Mutillidae (Brothers & Lelej 2017), which we scored as char. 387. Our reasoning for doing so was based on the observation that †*Melittosphex* shares more specific features in common than with Sphaerophthalminae, contrary to the contention of Rosa & Melo (2021), who removed †*Melittosphex* from Apoidea and considered the taxon *incertae sedis* as close to the sapygid-mutillid clade.
8 *DC: Sensilla. DS: Specific sensilla structure, distribution.* **• Observational criterion:** Body covered in fine dense iridescent pubescence (*FALSE* = such setae not observed). **• NOTES:** Observed in Pompilidae; scored as single-terminal autapomorphy due to the limited taxon sampling of the family.
9 (**Additive on 7**) *DC: Plumose “setoid” sensilla. DS: Specific sensilla structure, orientation, Body setation*. **• Observational criterion:** Body with brilliantly iridescent, appressed, squamiform, plumose seta patches (*FALSE* = such seta patches not observed). **NOTES:** Observed uniquely in *Thyreus*.

**-** Head - *Cranium*

10 *DC: Aboral region of cranium. DS: Extent of sclerotization between oral and occipital foramina.* **• Observational criterion:** Postgenal bridge relatively long, with length almost 0.5 x or clearly > 0.5 x head length (*FALSE* = postgenal bridge not observed to be long; short, more-or-less forming a transverse bar between the postocciput and hypostoma). **• NOTES:** Among sampled extant terminals, the postgenal bridge was observed to be elongated in Bethylidae, Dryinidae, Sclerogibbidae, *Ampulex* (Ampulicidae), and Formicidae; among extinct taxa, observed in a new genus of Scolebythidae, †Plumalexiidae, most Bethylidae, †*Burmasega* (Chrysididae), †*Falsiformica* (†Falsiformicidae), †Aculeata sp. (CASENT0844587, burmite), and Formicoidea; state in †Bethylonymidae uncertain, albeit they appear to be prognathous.
11 *DC: Head. DS: Development of female head orientation.* **• Observational criterion:** Head orientation prognathous, *i.e.*, mouthparts directed more anteriorly than ventrally (*FALSE* = prognathy not observed; head hypognathous, *i.e.*, mouthparts directed ventrally). **• NOTES:** “Prognathy” in Hymenoptera is difficult to objectively evaluate because the geometrical transformations of the head are complex and differ across the various sampled groups. The prognathous condition can occur with or without elongation of the postgenal bridge, as the condition is dependent on the angle of the postocciput relative to the length of the head. Among extant taxa, scored as *TRUE* for Bethylidae, Sclerogibbidae, Dryinidae, some Mutillidae, *Ampulex* (Ampulicidae), Philanthidae, *Ceratina* (Apidae), and Formicidae; among extinct taxa, scored as *TRUE* for †*Sessiliventer* (†Ephialtitidae), †Bethylonymidae, a new genus of Scolebythidae, †Plumalexiidae, all Bethylidae, Dryinidae, †Falsiformicidae, †*Burmasphex* ("†Angarosphecidae"), Sclerogibbidae, and all Formicoidea except for the †Haidomyrmecinae which are scored as "?" due to disagreement about the state of these ants. †*Plumalexius* was described as "hypognathous" by Brothers (2011). The male of *Aelurus* (Thynnidae) is hypognathous while the female prognathous; the female was scored in order to represent the additional transition to prognathy among the sampled taxa. Precise anatomical differentiation and definition of the variation classified as “prognathous” and “hypognathous” is desirable.
12 *DC: Oral region of the head. DS: Specific development of form.* **• Observational criterion:** Oral portion of head, bearing mouthparts and clypeus, narrowed and elongated (*FALSE* = such narrowing and elongation not observed). **• NOTES:** Unique to †Haidomyrmecinae.
13 *DC: Frontal region of the head. DS: Specific form of the oral region.* **Observational criterion:** Fronto-oral portion of head in lateral view transversely concave, with front of head at a distinct oblique angle relative to clypeus and antennal toruli (*FALSE* = such concavity not observed). • **NOTES:** Observed in †*Camelomecia* and †*Camelosphecia.*
14 (**Metric**) *DC: Head. DS: Head proportions*. **• Observational criterion:** Head width (proxy for body size) ≥ 1 mm (*FALSE* = head not observed to be broad; width < 1 mm). **• NOTES:** Scored because of the “miniaturization-for-success” hypothesis of Peeters & Ito (2015) for the Formicidae. This character is quite variable across the sampled taxa, and may be biased among fossils, with larger specimens preserved as compressions and smaller specimens in amber.
15 (**Metric**) *DC: Head. DS: Head proportions*. **• Observational criterion:** Head, including eyes, wider lateromedially than long oral-aborally (hypognathous: dorsoventrally; prognathous: anteroposteriorly) (*FALSE* = head not observed to have this proportion; longer fronto-abfrontally than wide lateromedially). **• NOTES:** Variable across the sampled taxa. In general, most sampled species have broad heads as here defined.
16 *DC: Head. DS: Specific development of head form*. **• Observational criterion:** Head more-or-less cone-shaped, with face—bearing antennal insertions—produced anteriorly, resulting in an elongate dorsal surface and with the antennal toruli being the most anteriorly-apical cranial sclerotic structures (*FALSE* = head not observed to have this shape). **• NOTES:** Unique to Embolemidae. State for associated compression fossils (†*Baissobius*, †*Cretembolemus*, and †*Embolemopsis*) uncertain.
17 *DC: Head. DS: Specific development of head form.* **• Observational criterion:** Head ventrally elongate and strongly flattened posteriorly, fitting against vertically truncate mesosoma (*FALSE* = head not observed to be flattened in this way). **• NOTES:** Synapomorphy of Evaniidae: Ronquist *et al*. (1999): state 1 of character 1; Li *et al*. (2018): state 1 of character 3. Interpretation of this state for fossil Evaniidae was difficult, thus many taxa were assigned "?".
18 *DC: Head. DS: Specific development of head form.* **• Observational criterion:** Head more-or-less wedge shaped, being lateromedially wider than anteroposteriorly long, elongate posterior to the compound eyes, and fronto-abfrontally compressed (*FALSE* = such head shape not observed). **• NOTES:** This specific state is observed in Dryinidae, †Falsiformicidae, †*Burmasphex* ("†Angarosphecidae"), and †Aculeata sp. (CASENT0844587, burmite).
19 *DC: Head. DS: Specific development of head form.* **• Observational criterion:** Head cuboidal, with cranium elongated posterad compound eyes (does not apply when eyes reduced) (*FALSE* = head not observed to have this shape). **• NOTES:** Observed variably across the Aculeata. Originally included due to the use of this state to define the Pemphredonidae (*e.g.*, Bohart & Menke 1976, Antropov 2000a), but expanded to include more-or-less any taxon with heads which have a distinct temple posterad the compound eye; this character, thus, is conceptually an aggregate of various forms. Capturing shape variation of the cranium is difficult in a qualitative framework, so would be valuable to examine via µ-CT scans.
20 *DC: Head. DS: Specific form of aboral surface*. **• Observational criterion:** Aboral head margin broadly concave in full-face or posterior view (*FALSE* = such concavity of aboral head margin not observed; margin narrowly concave to linear). **• NOTES:** Observed in *Dryinus*, and to be variable in Formicidae.
21 *DC: Craniothoracic contact surface of head. DS: Expression of contact surface rim.* **• Observational criterion:** Occipital carina present (*FALSE* = occipital carina not observed). **• NOTES:** Variable across the sampled taxa.
22 (**Additive on 21**) *DC: Contact surface rim. DS: Oral extent of rim*. **• Observational criterion:** Occipital carina extending to hypostoma (*FALSE* = such extension not observed; carina present OR absent). **• NOTES:** Observed in a subset of taxa, e.g., most non-aculeate outgroups, most Vespidae, *Aelurus* (Thynnidae), *Pluto* (Pemphredonidae), Philanthidae, and among Formicidae, observed in the formicines *Gigantiops* and *Santschiella*. Although additive, this character and chars. 23 and 24 are not expected to have strong effect due to wide phylogenetic pattern of occurrence.
23 (**Additive on 21**) *DC: Contact surface rim. DS: Specific form of rim*. **• Observational criterion:** Occipital carina conspicuously flared lateral to the occiput (*FALSE* = such flaring not observed; carina present OR absent). **• NOTES:** Flared occipital carinae occur sporadically across the present sample of taxa, including *Hyptia* (Evaniidae), *Deinodryinus* (Dryinidae), *Ampulex* (Ampulicidae), and *Santschiella* and *Gigantiops* (Formicidae).
24 (**Additive on 21**) *DC: Contact surface rim. DS: Development of rim and cranial surface immediately fronto-orad the rim*. **• Observational criterion:** Occipital carina visible in frontal view (*FALSE* = carina not observed in frontal view; carina present OR absent). **• NOTES:** Frontal or full-face view is defined in the myrmecological literature as orientation of the head wherein the apparent anterior and posterior margins of the cranium are in the same plane of focus. The *TRUE* state is observed sporadically among Formicidae.
25 *DC: Aboral region of head. DS: Development of cranial orientation relative to postocciput*. **• Observational criterion:** Postocciput apparently migrated to or near to the extreme aboralmost portion of head (*FALSE* = postocciput not observed to be in such a position; located more orally). **• NOTES:** This condition may be conceived as “hyperprognathy”. Unique to Leptanillinae *sensu lato*, described as "occiput enlarged" in Boudinot (2015). See Richter *et al*. (2021) for head anatomy of a representative of Leptanillinae.
26 *DC: Craniothoracic contact surface of head. DS: Fronto-oral extent of contact surface.* **• Observational criterion:** Craniothoracic contact surface of head, delimited by “occipital carina”, extending onto fronto-oral head surface, such that the surface is visible in frontal (full-face) view. **• NOTES:** Uniquely observed in *Opamyrma* (Leptanillinae).
27 *DC: Aboral region of head. DS: Specific functional form.* **• Observational criterion:** Aboral region of head (“occipital region”) differentiated, forming transverse lobe at aboral portion of head which has power amplification function for trap-jaw mechanism (*FALSE* = such occipital lobe not observed). **• NOTES:** Uniquely observed in *Myrmoteras.* Functional morphology of this cranial lump is described by Larabee *et al*. (2017).
28 *DC: Aboral region of head. DS: Specific development of head form*. **• Observational criterion:** Cranium elongated and narrowed aborally, such that postocciput apparently situated on neck-like aboro-medial process (*FALSE* = postocciput not observed to be situated on neck). **• NOTES:** Neck-like extensions of the cranium observed in various tropical Formicidae presumably under selection from army ant Dorylinae, but here scored for *Loboscelidia* wherein this appears to be an adaptation for myrmecophily.
29 *DC: Setoid sensilla. DS: Specified arrangement.* **• Observational criterion:** Abfrontal head surface margined by a single row of very long, curved setae, *i.e.*, psammophore present (*FALSE* = such setae not observed). **• NOTES:** Observed sporadically in Formicidae, including *Pogonomyrmex* (Myrmicinae) and *Melophorus* (Formicinae).

*- Eyes*

30 *DC: Compound eyes. DS: Position relative to epistomal line.* • **Observational criterion:** Oral margin of compound eye nearly abutting epistomal line near lateral mandibular base, such that the malar space (lateral area of head between epistomal line of clypeus and compound eye) virtually absent (*FALSE* = such eye position not observed; malar space longer). • **NOTES:** Among sampled taxa, not observed in non-aculeate outgroups, and very frequent in occurrence among Aculeata. Exceptions include some Chrysidoidea (Plumariidae, Scolebythidae, some Chrysididae), Dryinoidea, some Vespoidea, some Pompiloidea *sensu lato*, Bradynobaenidae, a few Apoidea, and most Formicidae. Male *Aelurus* with malar space virtually absent, female with distinct malar space.
31 *DC: Compound eyes. DS: Position relative to epistomal line*. • **Observational criterion:** Oral margin of compound eye relatively distant from epistomal line near lateral mandibular base, such that the malar space is oral-aborally longer than fronto-abfrontally broad (prognathous: anteroposteriorly vs. dorsoventrally; hypognathous: dorsoventrally vs. anteroposteriorly) (*FALSE* = such length of the malar space not observed, space broader than long OR absent). • **NOTES:** This relatively extreme conformation is observed in various Evanioidea, some Chrysidoidea, *Olixon* (Rhopalosomatidae), and most Formicidae.
32 *DC: Compound eyes*. *DS: Specific form.* • **Observational criterion:** Eyes bulging from head very strongly, appearing stalked, such that the face, in dorsal view, nearly trough shaped (*FALSE* = such bulging of eyes not observed). • **NOTES:** Among sampled taxa, observed in the dryinids †*Hybristodryinus magnificus* and *Dryinus gulfensis.*
33 *DC: Compound eyes. DS: Form of medial margins.* • **Observational criterion:** Eyes converging aborally (prognathous: posteriorly; hypognathous: dorsally), *i.e.*, tangent lines drawn on the medialmost margin or margins of the compound eyes not parallel and converge toward the occiput (*FALSE* = such aboral convergence not observed; medial margins parallel OR orally converging). • **NOTES:** Observed sporadically. Scored as *TRUE* for *Olixon* (Rhopalosomatidae), *Aelurus* (Thynnidae), *Sapyga* (Sapygidae), *Myrmosa* (Mutillidae), some Scoliidae, Ampulicidae, Crabronidae, some Philanthidae, and some Apidae. Evaluating this state for extinct taxa is challenging as compression fossils rarely preserve this detail, and specimens entrapped in amber are difficult to observe from the correct angle. Future studies employing µ-CT will provide much needed information about eye conformation.
34 *DC: Compound eyes. DS: Form of medial margins.* • **Observational criterion:** Eyes converging orally (prognathous: anteriorly; hypognathous: ventrally), *i.e.*, tangent lines drawn on the medialmost margin or margins of the compound eyes not parallel and converging toward the mandibles (*FALSE* = such oral convergence not observed; medial margins parallel OR aborally converging). • **NOTES:** Observed sporadically and difficult to score for the reasons outlined in the prior character.
35 *DC: Compound eyes. DS: Form of medial margins.* • **Observational criterion:** Medial eye margins distinctly emarginate or notched (*FALSE* = emargination OR notching not observed; linear to weakly OR nearly indiscernibly concave). • **NOTES:** Notched compound eyes were observed in Vespoidea *sensu stricto* with the exception of *Olixon*, some Pompiloidea *sensu lato*, Scoliinae (Scoliidae), a species of Philanthidae and Pemphredonidae, plus various Formicidae, for which only males are scored. Among the Formicidae, there is considerable variation of the medial eye margins, including obvious vespid-like notches (*e.g*., *Paraponera*, *Platythyrea*) and many intermediates where the line between "emarginate" and "linear" are blurred, and very slight concavity was scored as *FALSE*. Overall, it is best to score this character with the head in full-face view as defined above, and care should be taken with amber fossils due to the difficulty of achieving the sufficiently dorsal (prognathous) or anterior (hypognathous) perspective.
36 *DC: Compound eyes of females. DS: Eye expression*. *•* **Observational criterion:** Compound eye of female reduced in size, being < 1/2 head length in full-face view or completely repressed (absent) (*FALSE* = female eye not observed to have such relative size; larger, often taking up most of head). • **NOTES:** Reduced eyes in females are observed in some Chrysidoidea, some Pompiloidea *sensu lato*, the Bradynobaenidae, and the Formicidae. This state is associated with aptery, and among aculeate males was only observed for ergatoid (wingless, worker-like) male Formicidae, *e.g.*, *Cardiocondyla*, *Hypoponera*. For the putative zigrasimeciine genus †*Boltonimecia*, the posterior lobate processes of the head were interpreted as eyes which are grossly deformed due to preservation.
37 (**Additive on 36**) *DC: Compound eyes. DS: Eye expression*. • **Observational criterion:** Compound eyes expressed in all sexes and forms (*FALSE* = compound eyes not observed in all sexes and forms; eyes completely repressed in at least one morph). • **NOTES:** Compound eyes are not expressed (= “lost”) in workers of various Formicidae.
38 (**Taxic-reductive: Formicidae**) *DC: Apterous female compound eye. DS: Relative position.* **• Observational criterion:** Compound eyes of apterous female located at or aboral (posterior) to head midlength (*FALSE* = eye not observed to be aborally located; eye situated in oral (anterior) head half OR eye large and occupying most of both the oral (anterior) and aboral (posterior) head halves). • **NOTES:** Among the sampled taxa which have apterous individuals, *Plumarius*, *Pristocera*, Thynnoidea, Mutillidae, Bradynobaenidae, and *Heterogyna* have eyes which are invariably set anterior to head midlength, while *Sclerogibba* and *Olixon* have eyes which are invariably posteriorly set. For this reason, the characters employed here which describe location of the compound eyes in apterous individuals are taxically restricted to the Formicidae, wherein there is considerable and informative variation. In cases where compound eyes are completely absent, *e.g.*, *Apomyrma*, reproductive females scored. As noted at the beginning of the character descriptions, fossils of uncertain affinity are scored as "?" when the state is unknown, rather than "-" for inapplicable.
39 (**Taxic-reductive: Formicidae, additive on 38**) *DC: Apterous female compound eye. DS: Relative position.* **• Observational criterion:** Compound eye of apterous female located distinctly in aboral (posterior) half of head (*FALSE* = eye not observed to have this extreme location; eye situated at or oral (anterior) to head midlength). • **NOTES:** Additive with respect to prior character. Among extant Formicidae, observed in *Anomalomyrma* (Leptanillinae), Amblyoponinae *sensu lato*, *Tatuidris* (Agroecomyrmecinae), *Leptomyrmex* (Dolichoderinae), various Formicinae, Ectatomminae *sensu lato*, and some Myrmicinae; observed among most stem Formicidae, with the exception of one species of †*Zigrasimecia.*
40 *DC: Ocelli. DS: Ocelli expression*. • **Observational criterion:** All three ocelli expressed (present) (*FALSE* = ocelli not observed; ocelli incompletely or not at all developed). • **NOTES:** Among worker Formicidae, various degrees of ocellar reduction occur, including complete absence. Among other aculeates, ocelli were observed as absent in female *Plumarius* (Plumariidae), *Pristocera* (Bethylidae), various apterous Pompiloidea *sensu lato*, Bradynobaenidae, and *Heterogyna* (Heterogynaidae).
41 *DC: Ocelli. DS: Ocellar triangle location*. • **Observational criterion:** Ocelli situated aborally (prognathous: posteriorly; hypognathous: dorsally), being at or very near aboral head margin in full face view (*FALSE* = ocelli not observed to be so situated; relatively distant from aboral head margin). • **NOTES:** Observed infrequently, including some Bethylidae (*Epyris*, *Goniozus*), *Bembix* (Bembicidae), *Apis* (Apidae), and †*Allommation* (†Allommationidae). As this is an exceptional state, it is not reductively scored, although it does depend on presence of ocelli.
42 *DC: Ocelli. DS: Ocellar triangle location*. • **Observational criterion:** Situated in about the middle of the face, with the median ocellus within two of its diameters from the antennal toruli (*FALSE* = ocelli not observed to be so situated; relatively distant from toruli). • **NOTES:** Observed uniquely in *Caupolicana* (Colletidae), thus not scored reductively, despite dependence on presence of ocelli, as noted for the prior character.
43 *DC: Ocelli. DS: Ocellar form*. • **Observational criterion:** Any one of the three ocelli deformed (*FALSE* = ocelli not observed to be deformed; more-or-less circular to oval). • **NOTES:** Observed in various Apoidea and in Vespidae. Because this state is exceptional, not scored reductively.

- *Mouth appendages*

44 *DC: Craniomandibular condyle. DS: Condyle size*. • **Observational criterion:** Cranial condyle (that condyle located frontally) enlarged, restricting mandibular condyle (abfrontal condyle) and atala (abductor swelling) to aboral half of mandible (*FALSE* = condyle not observed to be enlarged; smaller, often not easy to distinguish). • **NOTES:** Observed in †*Camelomecia* and Formicidae. The cranial condyle in †*Burmasphex* ("Angarosphecidae") is enlarged, as is that of *Ampulex* (Ampulicidae); other Ampulicidae do not have an enlarged cranial condyle.
45 *DC: Craniomandibular articulation*. *DS: Shape.* • **Observational criterion:** Lateral base of mandible in form of tube-like bar (*FALSE* = condyle not observed to have such a form; smaller, often not easy to distinguish, OR other). **NOTES:** Observed uniquely in Scoliidae, although state unfortunately uncertain in *Proscolia* and the compression fossils assigned to the family.
46 *DC: Mandible. DS: Orientation of mandibular base.* • **Observational criterion:** Mandibles rotated in their sockets: Ventral mandibular articulation (VMA) and abductor swelling (ABS) NOT in sagittal plane of head (*i.e.*, not aligned on the lateral surface of the head), but rather VMA and ABS in diagonal plane, both being on the ventral head surface (assuming prognathy); in this position, the mandibles do not open in the transverse plane (*i.e.*, lateromedially), but rather mandibles open and close on a diagonal plane, swinging open posterolaterally and closing anteromedially (*FALSE* = VMA and ABS on lateral surface of head, with mandibular action restricted to lateromedial adduction and abduction). • **WARNING:** This primary homology hypothesis, proposed by Cai *et al*. (2020), has been falsified through µ-CT scanning of a worker of †*Zigrasimecia* (Richter & Boudinot, unpubl. data.). The topology searches and ancestral state estimation included this definition and the observations, as described in the NOTES subsection, which is retained here for clarity. • **NOTES:** The *TRUE* state is observed in †*Zigrasimecia* and †Haidomyrmecini; we recognize that the precise anatomical mechanism of this motion is as-yet undetermined in either taxon. However, specimens of †*Zigrasimecia* are known wherein the mandibles are opened. See Fig. 7 in Barden *et al*. (2017), for view of ventrally situated VMA and ABS of †*Linguamyrmex*, and Richter *et al*. (2019, 2020) for craniomandibular terminology.
47 *DC: Mandible. DS: Degree of torsion*. • **Observational criterion:** Mandible with extreme torsion such that, in frontal (full-face) view, usual lateral margin obscured by body of mandible and crossed over by basal mandibular margin (*FALSE* = mandible without such torsion; basal margin not crossing lateral margin). • **NOTES:** This degree of torsion is a unique synapomorphy of †*Zigrasimecia.* Rasnitsyn (2002, p. 247, character 104) proposed another form of mandibular torsion as a synapomorphy of the Aculeata exclusive of Chrysidoidea *sensu lato*; his characterization was "mandible with cutting edge twisted into moving plane of mandible". Unfortunately, this described observation was encountered too late to evaluate in the present study.
48 *DC: Mandible. DS: Development of blade*. • **Observational criterion:** Mandible more-or-less triangular, with an elongate masticatory margin, having a lateromedially broad blade and with the lateral and both medial margins oblique (*FALSE* = triangular form not observed; masticatory margin short, mandible more-or-less curved, lateral and medial margins divergent OR subparallel, *i.e.*, falcate OR linear). • **NOTES:** Triangular mandibles were observed in Trigonaloidea (Trigonalidae, †Maimetshidae), *Pristocera* (Bethylidae), Dryinidae, some Vespidae, *Sapyga* (Sapygidae), *Megachile* (Megachilidae), some Apidae, and the “poneroformicoids” (*i.e.*, the clade within the Formicidae comprising the “poneroids” and “formicoids”). Among extinct taxa, triangular mandibles were observed for †*Andrenelia* (†Andreneliidae), various Evanioidea, †Chrysobythidae, †*Camelomecia.*
49 *DC: Mandible. DS: Length and lateromedial curvature.* • **Observational criterion:** Mandible elongate and linear or bar-shaped, *i.e.*, very long (including masticatory margin), lateromedially narrow, lateral and medial margins more-or-less parallel for most of their length (*i.e*., crowbar or forcep like) (*FALSE* = mandible not observed to have this form; form variable). • **NOTES:** Crowbar-like or mandibles have evolved in various Aculeata, and particularly in the Formicidae where they function as trapjaws, *e.g.*, Bolton (2003), Larabee & Suarez (2014), Perrichot *et al*. (2016); note that this character purely is a descriptor of form, not of function, thus non-trapjaw species are scored as *TRUE*.
50 *DC: Mandible. DS: Length and lateromedial curvature.* • **Observational criterion:** Mandible elongate and narrow but curved (*FALSE* = mandible not observed to be elongate and curved; form variable). • **NOTES:** Among all taxa, observed in †*Myanmyrma mauradera* and †*M. gracilis* (stem Formicidae), Crabronidae, and Sphecidae.
51 *DC: Mandible. DS: Length and frontal-abfrontal curvature.* • **Observational criterion:** Mandible strongly bowed, i.e., bowl-like, with strongly convex frontal surface (prognathous: dorsal; hypognathous: anterior) (*FALSE* = mandible not observed to be bowed; form variable). • **NOTES:** Bowed mandibles were observed in the fossils †*Camelomecia* and †*Camelosphecia* in the extant ant *Tatuidris*; a similar but distinct conformation was observed in the leptanilline *Anomalomyrma taylori*. If analogy is accepted with *Tatuidris*, then †*Camelomecia* species may have been top predators (*e.g.*, Jacquemin *et al*. 2014).
52 *DC: Mandible. DS: Size.* • **Observational criterion:** Mandible hypertrophied, with outline nearly the same size in lateral view as cranium (*FALSE* = mandible not observed to be hypertrophied; form variable). • **NOTES:** Uniquely observed in the Mesozoic "camelo genus indet.", BALBuKL-41.
53 *DC: Mandible. DS: Oral-aboral width along proximodistal axis.* • **Observational criterion:** Mandible, in lateral view, with broad base which very strongly narrows to thin, nearly digitate distal portion (*FALSE* = mandible not observed to have such form; form variable). • **NOTES:** Observed in Apidae, specifically *Thyreus, Ceratina, Centris*, *Eufriesa*, and *Apis*; in the latter, the mandibular blade is secondarily expanded while retaining the narrow profile. Scored as true for †*Cretotrigona* given the very narrow mandibular apex of the holotype as preserved.
54 *DC: Mandible. DS: Notch specification.* • **Observational criterion:** Mandibular abfrontolateral margin (prognathous: ventrolateral; hypognathous: posterolateral) with distinct notch and tooth (*FALSE* = such a notch not observed; margin uninterrupted). **• NOTES:** A character of some non-anthophilan Apoidea (Bohart & Menke 1976) which is also observed among some Mutillidae (*e.g.*, some Myrmosinae). Among sampled taxa, observed in Crabroninae (*Plenoculus*, *Tachysphex*).
55 *DC: Mandibular tooth line. DS: Tooth patterning.* • **Observational criterion:** Masticatory margin with serrate teeth of alternating size (*FALSE* = such a pattern not observed; teeth uniform, irregular, OR absent). **• NOTES:** Small denticles or serrations separated by larger, distinct teeth were only observed in Formicidae. Within this family, they were observed in *Paraponera* (Paraponerinae), *Myrmecia pyriformis* (Myrmeciinae), *Aneuretus* (Aneuretinae), most Dolichoderinae (including †*Chronomyrmex* and the burmite species), the trapjaw formicine *Myrmoteras*, and *Rhytidoponera* (Ectatomminae). The form of these teeth differs among all major groups except *Aneuretus* and the Dolichoderinae, but all are here scored together agnostically as the ecological function and morphogenic mechanisms are unknown.
56 *DC: Mandibular tooth line. DS: Tooth size, proximodistal curvature.* • **Observational criterion:** Teeth in the form of violent, fang-like serrations (*FALSE* = teeth not in this form; form variable). **• NOTES:** Fang-like dentition was only observed in Amblyoponinae sensu stricto, *i.e*., with the exception of *Apomyrma*. Whereas the ecology of *Apomyrma* is unknown, other amblyoponines are generally accepted to be specialist predators of geophilomorph centipedes.
57 *DC: Mandibular tooth line. DS: Specification of position along proximodistal axis.* • **Observational criterion:** Teeth in double row, with one row directly above the other in frontal (full-face) view (*FALSE* = teeth not observed to be in double row; single row OR absent). **• NOTES:** Observed in *Stigmatomma*, *Harpegnathos*, and various †Haidomyrmecinae, a state which is certainly convergent. See also chars. 58, 60.
58 (**See 60**) *DC: Mandible elongation axis. DS: Axis specification point*. *•* **Observational criterion:** Basal angle of mandible extremely elongate and directed anteriorly away from head, with the blade of the mandible retained proximoventrally (*FALSE* = basal angle not observed to have this form; angle not developed OR angle not elongated OR angle elongated but directed anterodorsally, in close association with elements of the cranium OR angle of other form). • **NOTES:** This state is a unique to the ponerine *Harpegnathos* among crown ants but represents a stunning convergence with the †Haidomyrmecinae (see char. 60 for further information). The original formulation of the character, prior to Perrichot *et al*. (2020), was as follows: “Elongate mandible with proximoventral process which bears teeth from the ventral tooth row”.
59 *DC: Mandible. DS: Decoupling of male-female mandible development.* • **Observational criterion:** Male mandible reduced to nonfunctional spatulate to spiniform processes (*FALSE* = male mandible not observed to be reduced; well-developed, form variable). **• NOTES:** Observed sporadically among subfamilies of the crown Formicidae but is certainly a defining synapomorphy of the Ponerini in distinction to the Platythyreini (both Ponerinae) and remaining poneriines. Among sampled taxa, also scored as *TRUE* for *Opamyrma* (Leptanillinae) and *Apomyrma* (Amblyoponinae *sensu lato*); the condition for *Anomalomyrma*, specifically, remains uncertain.
60 (**See 58**) *DC: Mandible elongation axis. DS: Axis specification point*. • **Observational criterion:** Basal angle elongated and directed aboro-frontally (prognathous: anterodorsally), away from the mouth, but in close association with elements of the cranium (*FALSE* = mandible not observed to have this form; mandible of any other form OR angle elongate but not in association with the cranium). • **NOTES:** This character was reformulated after the publication of the †*Aquilomyrmex* genus group taxa in Perrichot *et al*. (2020). The original definition was as follows: “Basal mandibular tooth produced as sabre-like tong directed along the midline of the cranium”. This condition is a unique feature of †Haidomyrmecinae and *Harpegnathos* but is scored “taxically” given Remane’s fourth and fifth criteria of homology. For additional literature, see also Lattke *et al*. (2018), Perrichot *et al*. (2020), and Barden *et al*. (2020). The basal tooth of *Gasteruption* was observed to be elongate and dagger-like but was further maintained in the same plane of the remaining teeth.
61 *DC: Mandibular tooth line. DS: Tooth patterning.* • **Observational criterion:** Teeth occurring on basal mandibular margin (*FALSE* = teeth not observed on basal margin; teeth absent OR restricted to masticatory/incisor margin). **• NOTES:** Observed in Scolioidea, Leptanillinae, Amblyoponinae, *Harpegnathos* (Ponerinae), Myrmeciinae, and the Dolichoderinae. Note that the elongate "bar-like" or "strap-like" mandibles of *Anomalomyrma* are here interpreted to include teeth on the basal margin, which is, based on comparison with *Opamyrma*, the region of the mandibular margin medial to the dorsal longitudinal carina. The proximomedial tooth on the basal margin of the mandible of *Paraponera* (Paraponerinae) is also scored as a "basal tooth"; the "offset basal" tooth of *Lasius* and of *Anoplolepis* are here considered to be on teeth of the basal mandibular margin, and thus scored as *FALSE*. It appears that basal margin teeth are present in *Myrmoteras*, but this was not scored as *TRUE*, given the lack of intermediates.
62 *DC: Mandibular tooth line. DS: Bilateral tooth patterning.* • **Observational criterion:** Tooth count consistently asymmetrical, with one mandible having more teeth than the other (*FALSE* = tooth count not observed to be consistently asymmetrical; inconsistently asymmetrical OR symmetrical). **• NOTES:** Scored as an independent state in order to differentiate asymmetry from the additive condition of three to many teeth. Based on the present sample, it appears that developmentally canalized asymmetry of mandibular dentition is a unique synapomorphy of the Trigonaloidea, *i.e.*, including †Maitmetshidae, but has been recorded by Carmean & Kimsey (1998) as lost in the trigonalid genera *Nomadina*, *Bakeronymus*, and *Pseudonomadina*.
63 *DC: Mandibular tooth line. DS: Tooth patterning.* • **Observational criterion:** Mandible uni- or edentate (*FALSE* = tooth count not observed to be edentate or unidentate OR three or more dentate). **• NOTES:** (1) Mandibular dentition is scored as five separate characters, and these are split into two separate additive sets. The first set is "edentate to bidentate", while the second is "tridentate to many-dentate". This is done as many clades across the sampled taxa either have one or the other set of conditions, with limited exceptions. In effect, this scheme distinguishes between "many teeth" (≥ 3) and few teeth (< 3). In cases where dentition is present on the basal (non-incisor/masticatory) margin, both sets are scored; these cases include *Brachycistis* (Tiphiidae), Scoliidae, and some Formicidae. (2) The uni- and edentate states were lumped as it can be difficult to discern the two conditions in some taxa, as intermediate "blunt" or "angularly rounded" forms can be observed. (3) Reduced dentition, *i.e.*, few teeth, is observed in some Chrysididae, the Sclerogibbidae, Embolemidae, some Vespidae (*Euparagia*, Pseudomasaris), various Pompiloidea *sensu lato*, Scolioidea, and Apoidea. Although known among extant Formicidae, such as *Polyergus*, the occurrence of strict reduced dentition, *i.e.*, without teeth on the basal margin, is sporadic and no such species were sampled here. Among extinct Formicidae, all †*Zigrasimecia*, except for CNU009193, have a single large apical tooth subtended by fine, almost indistinct serrulation, and some †Haidomyrmecini appear to be uni- to bidentate. The unidentate condition was here confirmed by direct holotype observation for †*Brownimecia*.
64 (**Additive on 63**) *DC: Mandibular tooth line. DS: Tooth patterning.* • **Observational criterion:** Mandible at most bidentate (*FALSE* = mandible not observed to be bidentate; edentate, unidentate, tridentate, or more dentate). **• NOTES:** Additive with respect to prior character.
65 *DC: Mandibular tooth line. DS: Tooth patterning.* • **Observational criterion:** Mandible at least tridentate (*FALSE* = mandible not observed to be three or more dentate; edentate, unidentate, bidentate, or multidentate). **• NOTES:** This is the basic *TRUE* category for "many teeth", as defined above. Characters 66 and 67 are additive on the *TRUE* state. All sampled crown Formicidae have at least three teeth, although *Opamyrma* (Leptanillinae) and *Apomyrma* (Amblyoponinae *sensu lato*) serve as exceptions, as their masticatory margins are bidentate, but their basal margins are multidentate.
66 (**Additive on 65**) *DC: Mandibular tooth line. DS: Tooth patterning.* • **Observational criterion:** Mandible at least tetra- to nine-dentate (*FALSE* = mandible not observed to be 4–9-dentate; bidentate, tridentate, or uni- or edentate). **• NOTES:** Multidentate mandibles have evolved a number of times in the Aculeata. Among crown Formicidae, most of our sampled taxa have at least four mandibular teeth in whatever form. Exceptions include the *Discothyrea* species (Proceratiinae), *Odontomachus meinerti* (Ponerinae; other species in this genus with additional teeth). In keeping with the general condition of the Amblyoponinae, the mandibles of *Stigmatomma oregonense* were scored as multidentate; this species effectively has > 10 teeth, but these teeth are arranged are in a double row, with one row above the other.
67 (**Additive on 66**) *DC: Mandibular tooth line. DS: Tooth patterning.* • **Observational criterion:** Mandible ten- to many-dentate (*FALSE* = mandible not observed to be 10+ dentate; fewer-dentate). **• NOTES:** Among all sampled taxa, many-dentate mandibles only occur in the Formicoidea, including the crown clade and †*Camelomecia*. The many- dentate mandibles of †*Camelomecia janovitzi* differ from those of †*Camelomecia* sp. (ANTWEB1038930) in terms of denticle form; whether the male scored from the AMNH corresponds to a female with multidentate mandibles is unknown.
68 *DC: Mandibular “chaetoid” sensilla. DS: Expression.* • **Observational criterion:** Chaetae (“traction setae”, “peg-like setae”) present on abfrontal (prognathous: ventral; hypognathous: posterior) mandibular surface (*FALSE* = such chaetae not observed). **• NOTES:** Similar to the situation for “setoid” sensilla, there is conflict in the use of the term “chaeta” between the Hymenopteran and *Drosophila* research communities. For this reason, we distinguish “setoid” and “chaetoid” sensilla, the being distinctly thickened sensilla which have the function of increasing friction against surfaces, be they animate or inanimate (Boudinot *et al*. 2020). We also refer to them as “chaetae”. Often but not always, these chaetae are short, and under very high magnification (SEM), they have striate surfaces. Here, mandibular chaetae are observed in †*Camelomecia*, †*Zigrasimecia*, and *Tatuidris.* Some *Protanilla* do have this state, and the conformation of other extinct taxa is difficult to ascertain.
69 (**Metric**) *DC: Prementum of labium. DS: Proximodistal length.* • **Observational criterion:** Prementum elongate, at least four times longer than wide (*FALSE* = such elongation not observed; prementum shorter). **• NOTES:** Very difficult to observe in extinct taxa. Among extant taxa, observed in Chrysidinae (Chrysididae), Sapygidae, Scoliinae (Scoliidae), most Crabronidae and Sphecidae, Pemphredonidae, and in all Anthophila. This state may also be true for various Vespidae.
70 *DC: Labial palps. DS: Palp length, shape.* • **Observational criterion:** Labial palps modified for carrying, *i.e.*, palps long, palpomeres flattened and margined with long, curved hairs (*FALSE* = such palpal form not observed). **• NOTES:** Observed in various unscored Vespidae but, in particular, *Zethus*, which was scored.
71 *DC: Labial palps. DS: Relative palpomere length.* • **Observational criterion:** Labial palpomeres 1 and 2 greatly elongate relative to miniscule palpomeres 3 and 4. (*FALSE* = such palpomere proportions not observed) **• NOTES:** Among sampled taxa, observed as a unique synapomorphy of long-tongued bees, *i.e.*, Megachilidae + Apidae.
72 *DC: Labial galeae. DS: Galea length, shape.* • **Observational criterion:** Galea, distal to palp, greatly elongate, forming flat scoop-like process which is longer than stipes (*FALSE* = such galeal form not observed; galea shorter). **• NOTES:** Among sampled taxa, observed as a synapomorphy of the long-tongued bees, and in a few other taxa, *e.g.*, *Parnopes* (Chrysididae) and *Zethus* (Vespidae).
73 *DC: Labial glossa. DS: Glossa length, shape.* • **Observational criterion:** Glossa greatly elongate and pointed, longer than prementum (*FALSE* = such elongation not observed; glossa shorter). **• NOTES:** Another synapomorphy of the long-tongued bees which also occurs in *Parnopes* (Chrysididae) and *Zethus (Vespidae).* In Kimsey & Bohart (1990), listed as characters 8 and 9.
74 *DC: Labrum. DS: Labrum size and distal margin shape.* • **Observational criterion:** Labrum massively enlarged and arc shaped (*FALSE* = such labral form not observed; smaller OR with different shape). **• NOTES:** Massively enlarged labra were observed among sampled taxa in the stem Formicidae (†*Myanmyrma mauradera*, †*Zigrasimecia*), as well as *Bembix* (Bembicidae), *Megachile* (Megachilidae), and *Thyreus* (Apidae). Because the form of the enlarged labra of the apoids differs from that described here (elongate, subtriangular in *Bembix*, more-or-less rectangular in the bees), these taxa are scored as *FALSE*. Further, because the form between the †*Gerontoformica* and †Zigrasimeciinae differs, they are scored as separate characters, leaving the present state as a unique state of the former. Among extant Formicidae, an enlarged labrum is observed in species of *Rhopalothrix* in the Basicerotina [Myrmicinae: Attini]. Note also that the labrum of †Zigrasimeciinae completely conceals the maxillolabial complex, as observed in and scored as a unique state of the Dorylinae.
75 *DC: Labrum. DS: Labrum size and distal margin shape.* • **Observational criterion:** Massively enlarged and distally bilobate (*FALSE* = such labral form not observed; smaller OR not distally bilobate). **• NOTES:** Observed uniquely among sampled taxa in †Zigrasimeciinae (Formicidae).
76 *DC: Chaetoid sensilla. DS: Expression location.* • **Observational criterion:** Chaetae (“traction setae”, “peg-like setae”) expressed on aboral surface of labrum (*FALSE* = such chaetae not observed). **• NOTES:** Labral chaetae were uniquely observed in the Formicoidea, *e.g.*, †*Camelomecia* sp. (ANTWEB1038930), †Zigrasimeciinae, at least some †Haidomyrmecinae, †*Myanmyrma*, the Leptanillinae (*Anomalomyrma*, *Protanilla*, *Opamyrma*), and Amblyoponinae *sensu lato* (*Apomyrma*, *Amblyopone*). In the myrmecological literature, these chaetae are also known as “spicules” or “dentiform setae”. Some other aculeates have thickened setae on the labrum, but these are situated on the distal labral margin, not on the disc of the sclerite.
77 (**Additive-reductive; additive on 76, reductive if 76 *FALSE***) *DC: Chaetoid sensilla. DS: Expression location.* • **Observational criterion:** Labral chaetae restricted to anterior half of sclerite (*FALSE* = chaetae scattered over most of labral surface; "-" = inapplicable due to absence). **• NOTES:** The *TRUE* state was observed in †*Gerontoformica spiralis* and crown ants, where such chaetae are present. The condition could not be evaluated in the haidomyrmecines and was uncertain ("?") in †*Myanmyrma*.
78 (**Metric**) *DC: Oral foramen. DS: Length, width.* • **Observational criterion:** Oral-aborally longer than lateromedially broad (*FALSE* = such oral foramen proportion not observed; as broad or broader than long). **• NOTES:** Oral foramen = "buccal cavity" as used in the myrmecological literature. Observed sporadically among sampled taxa, including Chrysidinae (Chrysididae), most Vespidae, the Sapygidae, Scoliinae (Scoliidae), Crabronidae, *Ammophila* (Sphecidae), Philanthidae, and Anthophila.

- *Clypeus*

79 *DC: Clypeus. DS: Specific shape.* • **Observational criterion:** Clypeal disc bulbous and swollen, projecting orally away from head (anteriorly assuming hypognathy) (*FALSE* = clypeus not observed to have this form). **• NOTES:** Observed uniquely in *Bembix* (Bembicidae).
80 *DC: Clypeus. DS: Specific oral margin shape.* • **Observational criterion:** Perceived oral margin in frontal (full-face) view broadly and deeply concave (*FALSE* = such concavity not observed; convex to shallowly concave or notched). **• NOTES:** Uniquely observed in the hell ants, †Haidomyrmecinae and †Zigrasimeciinae. Notably, †*Boltonimecia*, placed in the †Zigrasimeciini by Borysenko (2017) does not appear to have a deeply concave anterior margin of the head.
81 *DC: Clypeus. DS: Oral margin position.* • **Observational criterion:** Oral clypeal margin apparently migrated aborally, up to or past antennal toruli, resulting in longitudinal paired lines on the clypeus (*FALSE* = such modification not observed). **• NOTES:** Uniquely observed in †Haidomyrmecinae; first described by Perrichot *et al*. (2016).
82 (**Additive on 81**) *DC: Clypeus. DS: Oral margin position.* • **Observational criterion:** Aborally migrated oral clypeal margin produced away from head as horn or lobe, the base of which subtends the antennal toruli (*FALSE* = such form not observed; not of this form OR clypeus without aborally-migrated oral margin). **• NOTES:** Uniquely observed in the haidomyrmecines †*Ceratomyrmex* and †*Linguamyrmex*.
83 *DC: Clypeus. DS: Specific medial clypeus shape.* • **Observational criterion:** Median portion of clypeus, between anterior tentorial pits, raised away from surface of head and margined by distinct arcuate carina (*FALSE* = median clypeal portion not observed to have this form). **• NOTES:** Observed in the ant genera *Opamyrma, Anomalomyrma*, and *Protanilla* (Leptanillinae). Absence in *Leptanilla* presumably explained by loss.
84 *DC: Clypeus. DS: Specific medial clypeus shape.* • **Observational criterion:** Median portion of clypeus raised away from surface of head and in form of nearly pyramidal knob (*FALSE* = median clypeal clypeus not observed to have this form). **• NOTES:** Observed in *Apomyrma* (Amblyoponinae); previously hypothesized to be homologous with the posteriorly carinate clypeus of the leptanillines (Boudinot 2015).
85 (**Metric**) *DC: Clypeus. DS: Proportions.* • **Observational criterion:** Median portion of clypeus, oral-aborally longer (prognathous: anteroposteriorly; hypognathous: dorsoventrally) than lateromedially broad (*FALSE* = median clypeal portion not observed to have these proportions; broader than long). **• NOTES:** Observed in Evaniidae, Gasteruptiidae, *Plumarius* (Plumariidae), some Dryinidae, some Vespidae, Sapygidae, Sphecidae, *Apis* (Apidae), and some Formicidae.
86 *DC: Line of bilateral symmetry. DS: Bilateral slope of the frontal surface of the clypeus.* **• Observational criterion:** Bilateral midline of clypeal disc formed as a ridge between the left and right halves (*FALSE* = such ridging not observed). **• NOTES:** Diagnostic of the Bethylidae, observed in all †Chrysobythidae, and occurring sporadically elsewhere (*Argochrysis*, *Deinodryinus*, some apoids, and some Formicidae, *e.g.*, †Haidomyrmecinae, some Formicinae).
87 *DC: Clypeus oral region. DS: Orad elongation.* • **Observational criterion:** Oral region of clypeus, not just oral margin, broadly projecting orally away from cranium (*FALSE* = disc not observed to project in such a way). **• NOTES:** The clypeus projects anteriorly from the head in many sampled taxa, *e.g.*, Trigonalidae, Plumariidae, Chrysidinae (Chrysididae), Sclerogibbidae, Embolemidae, Vespoidea *sensu stricto*, Pompilidae, Sapygidae, Scoliidae, Ampulicidae, Heterogynaidae, various Anthophila, and various Formicoidea. Future studies will benefit from anatomical analysis of clypeal conformation, as this character is here broadly conceived and surely includes an aggregate of differentiable forms.
88 *DC: Clypeolabral articulation region. DS: Specific form.* • **Observational criterion:** Oral-lateral regions of clypeus broadly curved abfronto-medially (hypognathous: posteromedially) around labral base (*FALSE* = this region not observed to have such a shape). **• NOTES:** Observed in *Bembix* (Bembicidae) and various Anthophila.
89 *DC: Oral-lateral regions of head. DS: Orad elongation.* • **Observational criterion:** Oral-lateral regions of clypeus AND face produced orally past base of mandibular insertion (*FALSE* = such elongation not observed). **• NOTES:** Observed in *Perdita* (Andrenidae), but also occurring in various Halictidae, and extremely derived in some Xeromelissinae.
90 *DC: Oral-lateral regions of clypeus. DS: Orad elongation.* • **Observational criterion:** Oral-lateral regions of clypeus extended orally as lobate processes of varying form, *i.e.*, lateroclypeal lobes present (*FALSE* = such elongation not observed). **• NOTES:** Observed uniquely in stem Formicidae (†*Gerontoformica*, †*Myanmyrma*, †*Boltonimecia*, and †Zigrasimeciini).
91 *DC: Oral-lateral regions of clypeus. DS: Shape of oral margin.* • **Observational criterion:** Lateroclypeal lobes in the form of enlarged flat, rounded laminar lobes (*FALSE* = lobes more angular OR not laminar OR not developed). **• NOTES:** Observed in †*Zigrasimecia*, †*Protozigrasimecia*, and †*Boltonimecia.* Where these lobes are present in other Formicoidea, the lobes are angular, suggesting that this state is a synapomorphy for the two genera where these occur.
92 *DC: Oral-median clypeal region. DS: Shape of oral margin.* • **Observational criterion:** Oral-median clypeal margin angular, regardless of notching (*FALSE* = margin rounded convex or linear). **• NOTES:** This is a broad character which should be refined in future studies by distinguishing angularity versus convexity, and the various forms of notching, as well as dimensions.
93 *DC: Oral-median clypeal region. DS: Shape of oral margin.* • **Observational criterion:** Oral-median margin of clypeus linear (*FALSE* = margin markedly concave or convex). **• NOTES:** Observed sporadically among sampled taxa.
94 *DC: Protuberance on oral-median clypeal region. DS: Protuberance expression.* • **Observational criterion:** Distinct protuberance (“lobe” or “process”) developed medially on oral margin of clypeus, excluding "clypeal shelves" and other non-medially located projections (*FALSE* = such a protuberance not developed). **• NOTES:** Scored as *TRUE* when a distinct lobe is located medially on the anterior clypeal margin of the clypeus; does not include the state of medial production of the body of the clypeus itself. “Clypeal shelves” are paramedian processes of the clypeus orad the toruli that are observed in various Myrmicinae, *e.g.*, *Tetramorium* (Bolton 1994).
95 *DC: Oralmost portion of clypeus. DS: Shape and transparency.* • **Observational criterion:** Oralmost portion of clypeus medially notched; this notch surrounded by convex laminar (thin, variably transparent) processes (*FALSE* = this region not notched AND/OR convex lamina not developed). **• NOTES:** Scored as a unique state of the dolichoderomorphs (Aneuretinae + Dolichoderinae), with the exception of *Bothriomyrmex* (Bothriomyrmecini). Unfortunately, this condition could not be evaluated for putative Mesozoic aneuretines.
96 *DC: Clypeus and labrum. DS: Shape.* • **Observational criterion:** Clypeus and labrum both deeply medially notched; this notch surrounded by narrow and long lobes which project orad from the cranium (*FALSE* = either clypeus or labrum or both not notched). **• NOTES:** Observed uniquely in †*Myanmyrma gracilis* (Formicidae).
97 *DC: “Chaetae” (thick “setoid” sensilla). DS: Chaeta expression.* • **Observational criterion:** Chaetae developed along oral-median margin of clypeus (*FALSE* = such sensilla not developed). **• NOTES:** These stout, short chaetae are also known as “spicules”, “dentiform setae”, “peg-like setae”, or “traction setae” in the myrmecological literature, and were observed in †Sphecomyrmini, †Haidomyrmecini, and Amblyoponinae. Among the stem Formicidae, absent in †*Sphecomyrma*.
98 *DC: Clypeal protuberances proximad “chaetae”. DS: Protuberance expression.* • **Observational criterion:** Cuticular protuberances developed on clypeus, raising chaetae away from body (*FALSE* = such protuberances not developed). **• NOTES:** Observed uniquely in the Amblyoponinae, *e.g.*, *Fulakora*, *Stigmatomma.*
99 *DC: Paramedian clypeal protuberances. DS: Protuberance expression.* • **Observational criterion:** Paramedian, paired dentiform cuticular protuberances developed, *i.e.*, paired denticles present (*FALSE* = such protuberances not developed). **• NOTES:** Observed sporadically in the Myrmicinae (Formicidae), but among sampled taxa, only developed in *Solenopsis* (Solenopsidini).
100 *DC: Setiform sensilla. DS: Sensilla specification on clypeus.* • **Observational criterion:** Unpaired median seta developed at about the medialmost point of oral-median clypeal margin (*FALSE* = such a seta in such a location not developed). **• NOTES:** Observed sporadically in the Formicidae, where this seta often has diagnostic value at the genus level. Morphologically, the unpaired anteromedian seta is part of the "basket seta" series, *i.e.*, the series of setae on the clypeus which overhang the mandible for object manipulation purposes. The basket seta series is also observable on *Myrmosa* (Mutillidae), *Apterogyna* (Bradynobaenidae), and other aculeates with apterous individuals, but this pattern was realized too late to include as a separate character.
101 *DC: Brush of setiform sensilla on medioclypeus. DS: Brush expression.* • **Observational criterion:** Dense brush of long, straight, and orally directed setae developed on the oral- median surface of the clypeus (*FALSE* = such a brush not developed). **• NOTES:** This mysterious toothbrush moustache is uniquely observed in *Martialis* (Leptanillinae *sensu lato*). A less dense brush of setae were also observed on the underside of the clypeal horn of †*Ceratomyrmex* (†Haidomyrmecini), but this conformation considerably different, and was not scored.
102 *DC: Lateroclypeal glandular seta brush. DS: Male brush expression.* • **Observational criterion:** Dense brush of glandular setae developed on the oral margins of the lateroclypeal regions (*FALSE* = such a brush not developed). **• NOTES:** Observed in Philanthidae and Melittidae.

- *Gena and face*

103 *DC: Pleurostomal protuberance. DS: Protuberance expression.* • **Observational criterion:** Spiniform protuberances developed on oral-lateral-most corners of pleurostoma, *i.e.*, “genal armature” present, these in close proximity to the cranial condyle and frontal (dorsal) to the posterolateral clypeal margin (*FALSE* = such protuberances not developed). **• NOTES:** Observed uniquely in †*Brownimecia clavata* and Amblyoponinae *sensu stricto*.
104 *DC: Glandular pits of lateral head cuticle. DS: Pit expression.* • **Observational criterion:** Pits developed on lateral surface of head, *i.e.*, “genal secretory organ” present (*FALSE* = such pits not developed). **• NOTES:** Unique to Bradynobaenidae, described by Brothers (1975; char. 11, Fig. 7), and scored as character 17 of Appendix VI in Brothers & Carpenter (1993).
105 *DC: Oral extension of ocular sulcus. DS: Expression.* • **Observational criterion:** Ocular sulcus extending from compound eye to pleurostoma at the base of the cranial condyle, *i.e.*, “malar line” present (*FALSE* = ocular sulcus not extending to pleurostoma). **• NOTES:** The ocular sulcus corresponds to the ocular ridge internally. It is developed as a line between the compound eyes and cranial condyle infrequently among Aculeata. Here such an extension was observed in Plumariidae, *Amisega* (Chrysididae), Dryinidae, *Olixon* (Rhopalosomatidae), and *Neoponera unidentata* (Formicidae, Ponerinae). In *Plumarius*, this sulcus is almost located on the postgenal bridge, whereas in Chrysididae and Dryinidae, the sulcus is almost on the face; scored as *TRUE* for the sampled ponerine, although the sulcus is carinate and certainly homoplastic.
106 *DC: Clypeotorular sulci. DS: Expression.* • **Observational criterion:** One or two oral-aborally oriented sulci developed between the clypeus and each torulus, *i.e.*, “subantennal sutures” present (*FALSE* = such sulci not developed). **• NOTES:** "Subantennal sutures" are singly or paired per torulus are observable sporadically in Aculeata (*e.g*., Philanthidae, some male Ectatomminae of the Formicidae), but are of widespread diagnostic value in Anthophila (*e.g.*, Michener 2007). Whether these sulci correspond to internal ridges has yet to be confirmed; their morphological function or functions are unknown.
107 (**Additive**) *DC: Clypeotorular sulci. DS: Pairing.* • **Observational criterion:** Two clypeotorular sulci developed per torulus (*FALSE* = either one OR no such sulci developed). **• NOTES:** Additive with respect to the presence of these sulci. Among sampled taxa, this state restricted to *Caupolicana* (Colletidae) and *Eufriesa* (Apidae).
108 (**Additive**) *DC: Clypeotorular sulci. DS: Contact point on torulus.* • **Observational criterion:** Medialmost clypeotorular sulcus contacting its torulus medial to torulus midlength (*FALSE* = sulcus contacts torulus at or lateral to torulus midlength). **• NOTES:** Scored for the medialmost sulcus or the sole sulcus, if only one is present. Additive with respect to "suture" presence. This state observed in Philanthidae (Aphilanthini and Philanthini, but not Cercerini), Andrenidae, Colletidae, Halictidae, and some Apidae.
109 (**Additive**) *DC: Clypeotorular sulci. DS: Contact point on torulus.* • **Observational criterion:** Medialmost clypeotorular sulcus contacting its torulus at torulus midlength (*FALSE* = sulcus contacts torulus more medially or more laterally). **• NOTES:** Scored for the medialmost sulcus or the sole sulcus, if only one is present. Additive with respect to "suture" presence. This state observed in *Ammophila* (Sphecidae), *Pluto* (Pemphredonidae), *Cerceris* (Philanthidae), †*Melittosphex*, Melittidae, and most sampled Apidae.
110 (**Additive**) *DC: Clypeotorular sulci. DS: Contact point on torulus.* • **Observational criterion:** Medialmost clypeotorular sulcus contacting its torulus lateral to torulus midlength (*FALSE* = sulcus contacts torulus at or medial to torulus midlength). **• NOTES:** Scored for the medialmost sulcus or the sole sulcus, if only one is present. Additive with respect to "suture" presence. Among sampled taxa, this state only observed in *Megachile* (Megachilidae).
111 *DC: Frontomedian protuberance. DS: Protuberance expression.* • **Observational criterion:** Massive protuberance developed medially on face; this protuberance bearing the antennal toruli (*FALSE* = such a protuberance not developed). **• NOTES:** Observed uniquely in the extinct evanioid family †Othniodellithidae.
112 *DC: Frontal surface of head. DS: Curvature.* • **Observational criterion:** Face broadly convex across its entire surface (*FALSE* = face flat OR of other form, not matching the form shown in Fig. C112-1). **• NOTES:** To capture the unique form of the face in Philanthidae, initially scored as follows: "Clypeofrontal line at anterior tentorial pit broadly V-shaped, with posterior margin of lateroclypeus subequal in length to posterolateral margin of medioclypeus". However, this description seems inadequate for replication as is, thus it has been substituted by another salient feature of these wasps, the broadly convex face.
113 *DC: Frontomedial protuberance. DS: Protuberance form.* • **Observational criterion:** Face frontomedially produced; regardless of carination, this protuberance transversely oriented between or at the aboral margins of the antennal toruli, whether linear, convex, or concave in form (*FALSE* = protuberance not developed OR protuberance longitudinally oriented). **• NOTES:** Initially scored as the "supraantennal elevation" of Carmean & Kimsey (1998), defining Trigonalidae. However, frontomedial protuberances were observed to be frequent and of variable form across the sampled taxa. Transverse protuberances similar to those of Trigonalidae were observed in various Aculeata, including Sclerogibbidae, some Vespidae, *Sapyga* (Sapygidae), *Myrmosa* (Mutillidae), and various extinct Ampulicidae, plus the extant *Dolichurus*. With respect to *Sapyga*, this protuberance is notably absent in *Fedtschenkia* and †*Cretosapyga* but was unobservable in †*Cretofedtschenkia*. Because the transverse orientation of these protuberances is almost certainly homoplastic across the sampled taxa, future studies may refine the characterization to capture variation specific to the various origins of this structure.
114 *DC: Carination. DS: Carina specification.* • **Observational criterion:** Longitudinally oriented carina present medially on frontal surface of head; this carina not formed from fusion of paired “frontal carinae” (*FALSE* = such a carina not developed OR longitudinal ridge formed by fusion of a pair of carinae). **• NOTES:** Observed sporadically across the sampled taxa, including most Evaniidae, †*Maimetsha arctica* (†Maimetshidae), some apoids, and the heteroponerine *Acanthoponera* (Ectatomminae, Formicidae). In some taxa, such as the ant and the evaniids, the carina extends onto the clypeus; this variation was not scored here, however.
115 *DC: Carination. DS: Carina specification.* • **Observational criterion:** Paired carinae present lateral to the antennal toruli, even if just their anterior margins (*FALSE* = frontal carinae absent OR carinae mediad toruli). **• NOTES:** Observed uniquely in extant Evaniidae; unfortunately, difficult to evaluate for extinct taxa.
116 *DC: Carination. DS: Carina specification.* • **Observational criterion:** Paired carinae present mediad the toruli (“medial frontal carinae”)), even if just their anterior margins; carinae may be nearly completely fused medially (*FALSE* = frontal carinae absent OR carinae laterad toruli). **• NOTES:** In general, when carinae are present on the face, they are parallel to subparallel, being oriented oral-aborally, thus scoring the parallel state independently is redundant if frontal carina presence is also scored. For this reason, the exceptional state of semicircular carinae is scored below.
117 *DC: Medial frontal carinae. DS: Specification relative to toruli.* • **Observational criterion:** Medial frontal carinae anteriorly continuous (“fused”) with medial torular arches (*FALSE* = frontal carinae absent OR unfused, separate). **• NOTES:** This is a canonical synapomorphy of Ponerinae (Formicidae); this state is also known to occur in some Amblyoponinae (see Bolton 2003).
118 *DC: Medial frontal carinae. DS: Specific form.* • **Observational criterion:** Medial frontal carinae oriented more-or-less diagonally or lateromedially AND carinae produced frontally away from the surface of the head as lobes (*FALSE* = frontal carinae absent OR carinae not produced away from head OR carinae oriented more-or-less oral-aborally, at least anteriorly OR carinae medially fused). **• NOTES:** Among sampled taxa, observed uniquely in the male of *Aelurus* (Thynnidae).
119 *DC: Medial frontal carinae. DS: Specific form.* • **Observational criterion:** Medial frontal carinae nearly parallel to long axis of head AND carinae produced frontally away from the surface of the head as lobes (*FALSE* = frontal carinae absent OR carinae not produced away from head OR carinae oriented mor-or-less lateromedially OR carinae medially fused). **• NOTES:** Observed in *Lioponera,* among other Dorylinae.
120 *DC: Medial frontal carinae. DS: Specific form.* • **Observational criterion:** Medial frontal carinae forming semicircle around antennal toruli; carinae evenly, sinuously, or angularly curving laterally (*FALSE* = frontal carinae absent OR carinae not curving laterally aborad toruli, being truncate *or* without curvature, even if transversely oriented).
**• NOTES:** This exceptional state is observed in *Dasymutilla* (Mutillidae) and is diagnostic of †*Gerontoformica* and †*Sphecomyrma* (Boudinot *et al*. 2020).
121 *DC: Medial frontal carinae. DS: Specific form.* • **Observational criterion:** Medial frontal carinae meeting medially on pyramidal-shaped frontal protuberance, *i.e.*, medial pyramid (lobe) present (*FALSE* = frontal carinae absent OR not meeting medially OR not situated on pyramidal protuberance). **• NOTES:** Unique to †Haidomyrmecinae (Perrichot *et al*. 2016).
122 *DC: Medial frontal carinae. DS: Specific form.* • **Observational criterion:** Medial frontal carinae undifferentiated medially (“fused”) forming median longitudinal shelf; carinae may differentiated aborad, *i.e.*, “fusion” may be present for at least half length of frontal carinae (*FALSE* = frontal carinae absent OR carinae distinct medially). **• NOTES:** Fusion may be present posteriorly, as in various Dorylinae. Among scored taxa, however, present only in *Discothyrea*.
123 *DC: Medial frontal carinae. DS: Specific form.* • **Observational criterion:** Medial frontal carinae “pinched” and apparently lobate, *i.e.*, carinae narrowly separated orally and widely diverging aborally, and oral termini of carinae close in proximity to aboral termini of medial torular arches, thus appearing to form large “frontal lobes” (*FALSE* = frontal carinae absent OR carinae not narrowly approximated orally and widely diverging aborally AND/OR carinae not appearing to form lobes). **• NOTES:** Only observed in Formicidae, where this state occurs in some Amblyoponinae and most Ponerini (Ponerinae).
124 *DC: Medial frontal carinae. DS: Specific form.* • **Observational criterion:** Medial frontal carinae extending orally onto clypeus (*FALSE* = frontal carinae absent OR carinae not extending onto clypeal surface). **• NOTES:** This condition occurs uniquely in Formicidae, including †*Myanmyrma mauradera*, *Feroponera*, and *Lioponera*, among other taxa.
125 *DC: Antennocranial contact surfaces. DS: Specific form.* • **Observational criterion:** Scapal contact surfaces of face both (a) longitudinally oriented and (b) situated between lateral frontal carinae. (*FALSE* = these “scrobes” not oriented along the longitudinal head axis OR not situated between lateral frontal carinae). **• NOTES:** Distinct, deep, and protective “antennal scrobes” have evolved independently many times across the Apocrita. Here, each clear example of an independent origin is scored separately as the form of these scrobes is differentiable given the conformation of the head, and their specific orientations and conformations. The particular form described in this character was observed in *Evania* (Evaniidae) and is scored as uncertain ("?") for †*Mesevania*, which appears to have scrobes, albeit paired.
126 *DC: Antennocranial contact surfaces. DS: Specific form.* • **Observational criterion:** “Scapal basin present”, *i.e.*, scapal contact surfaces of face with the following conditions: (1) surfaces medially contiguous, forming a single median depression; (2) this depression longitudinally oriented and longer oral-aborally than lateromedially; and (3) medial depression extending from the clypeus nearly to the ocellar triangle, well past compound eye midlength (*FALSE* = “scrobes” not medially contiguous OR not longitudinally oriented OR not exceeding compound eye midlength). **• NOTES:** This is the "scapal basin" of the Chrysididae (*e.g.*, Kimsey & Bohart 1990). Generally, the scapal basin is present in Amiseginae and Chrysidinae, being absent in Cleptinae and *Loboscelidia*.
127 *DC: Antennocranial contact surfaces. DS: Specific form.* • **Observational criterion:** Scapal contact surfaces of face with the following conditions: (1) surfaces medially contiguous; (2) this depression longitudinally oriented; and (3) depression shorter than one half compound eye length (*FALSE* = “scrobes” not medially contiguous OR not longitudinally oriented OR exceeding one half compound eye length). **• NOTES:** This state is observed in the sampled Crabronidae.
128 *DC: Antennocranial contact surfaces. DS: Specific form.* • **Observational criterion:** Scapal contact surfaces of face in the form of long, paired, longitudinal depressions that are both completely medial to the compound eyes *and* extending past compound eye midlength (not illustrated) (*FALSE* = “scrobes” not paired OR extending laterad compound eyes OR not extending past compound eye midlength). **• NOTES:** As here defined, observed uniquely in †*Colmepsiterona* (Pemphredonidae); see Cockx & McKellar (2018).
129 *DC: Antennocranial contact surfaces. DS: Specific form.* • **Observational criterion:** Scapal contact surfaces of face receiving the entire antenna in repose, with a longitudinal carina or wedge-shaped ridge which separates the flagellum from the scape (*FALSE* = “scrobes” not receiving entire antenna in repose OR without a median, longitudinal carina or wedge). **• NOTES:** Among the Aculeata, unique to the Formicidae, wherein among sampled taxa, this state was observed in *Paraponera* (Paraponerinae) and *Tatuidris* (Agroecomyrmecinae). This state is also present in *Ankylomyrma*, which was recently shown to be sister to *Tatuidris* (Ward *et al*. 2015), as well as the Eocene agroecomyrmecines †*Eulithomyrmex* and †*Agroecomyrmex*.
130 *DC: Antennocranial contact surfaces. DS: Specific form.* • **Observational criterion:** Scapal contact surfaces of face receiving the entire antenna in repose, but: (1) without a longitudinal carina separating the flagellum and scape; and (2) with their medial margins produced as “frontal lobes” that obscure the mandibular insertions in full-face view (*FALSE* = “scrobes” not receiving entire antenna OR with a longitudinal carina OR without such lobes). **• NOTES:** Among sampled taxa, observed uniquely in *Stegomyrmex* (Solenopsidini, Myrmicinae, Formicidae).
131 *DC: Antennocranial contact surfaces. DS: Specific form.* • **Observational criterion:** Scapal contact surfaces of face in the form of a pair of depressions which extend posterolaterally from the toruli to the compound eyes, with their medial margins raised high above the main depression surface (*FALSE* = “scrobes” not paired OR not extending to compound eyes OR are shallow). **• NOTES:** Among sampled taxa, observed uniquely in the †Zigrasimeciinae (†*Boltonimecia*, †*Zigrasimecia*; Formicidae). The species †*Gerontoformica contega* is putatively defined by similar scapal contact surfaces, but these scrobes are shallower
132 *DC: Foveae. DS: Expression.* • **Observational criterion:** “Facial foveae present”, *i.e.*, face, between compound eyes, with paired, deep, distinct depressions which are not contact surfaces (*FALSE* = such foveae absent). **• NOTES:** Observed in many Andrenidae, for which these pits are diagnostic; also occurring in some Colletidae (Michener 2007).
133 *DC: Carination. DS: Specification.* • **Observational criterion:** Face with transverse carina oral to the ocellar triangle, distant from the antennal toruli (*FALSE* = such a carina absent OR carina situated close to toruli, distant from ocelli). **• NOTES:** Synapomorphy of Chrysidini within Chrysidinae, character 5 of Kimsey & Bohart (1990). Among sampled taxa, observed uniquely in *Argochrysis*, although state is uncertain in some Mesozoic Chrysidinae.
134 *DC: Rugose cuticular sculpture. DS: Specification.* • **Observational criterion:** Face with dense transverse rugosity (*FALSE* = face sculpture not rugose OR rugose sculpture irregular, not transverse). **• NOTES:** Unique to the Early Cretaceous evanioid †*Andrenelia* (†Andreneliidae). Also known to occur among some Formicidae, such as the hyperdiverse myrmicine genus *Pheidole* (*e.g.*, Wilson 2003).
135 *DC: Posterolateral cranial regions. DS: Form.* • **Observational criterion:** Cranial sclerite “eared”, *i.e.*, with posterolateral portions produced as distinct lobes (*FALSE* = head not posterolaterally produced). **• NOTES:** Observed sporadically in crown Formicidae, and probably synapomorphic of the Ectatomminae *sensu lato*, including †*Canapone dentata* from Grassy Lake amber from the Late Cretaceous.

- *Periantennal sclerites*

136 *DC: Toruli. DS: Relative positions.* • **Observational criterion:** Antennal toruli extremely anteriorly situated such that: (a) their oral margins are nearly at or are overhanging the oral clypeal margin or the mouth; OR (b) the toruli are set clearly at or oral to the oral margins of the malar areas (*FALSE* = toruli not as orally situated). **• NOTES:** The state of extremely orally set antennal toruli has evolved independently a number of times among the Apocrita. These independent origins, however, are not coded separately as the conformation of the Jurassic family †Bethylonymidae cannot be evaluated, given that these taxa are compression fossils and that the group has been postulated to be sister to the Aculeata by Rasnitsyn (*e.g.*, Rasnitsyn 2002), forming the clade "Vespomorpha". Among sampled taxa, far anterior toruli were observed in most Chrysidoidea *sensu stricto*, the Sclerogibbidae, Bradynobaenidae, and various Formicidae, including *Martialis* (Leptanillinae *sensu lato*), Proceratiinae, and *Lioponera* (Dorylinae). In future studies, it may be possible to differentiate among the independent origins using anatomical criteria; at present, this character is left as aggregate, particularly until bethylonymids can be better placed among the Apocrita. Males of taxa with pronounced sexual dimorphism often have antennae set further away from the oral clypeal margin than their conspecific females.
137 *DC: Cranium. DS: Shape.* • **Observational criterion:** Face produced as a broad and convex process over the mandibles; this process bearing the antennal toruli ventrally, with the toruli directed aborad (*FALSE* = face not produced OR facial process without toruli ventrally OR toruli on underside of process not directed aborad). **• NOTES:** This conformation is observed uniquely in Sclerogibbidae and has been scored as *TRUE* for †*Sclerogibbodes embioleia*. The putative bethylonymid †*Meiagaster* may have this state, but it is challenging to confirm this as the taxon is described from a compression fossil. A similar process is also observed in *Discothyrea* (Proceratiinae, Formicidae), although in this case the toruli are directed dorsally.
138 *DC: Toruli. DS: Relative positions.* • **Observational criterion:** Antennal toruli located at about head midlength or aboral to head midlength (*FALSE* = toruli clearly set in oral half of head). **• NOTES:** This character is broadly scored as *TRUE* when the antennae are at or near head midlength. Scoring may be refined in future studies, particularly via explicit quantification; indeed, this should be done as the current coding does not capture informative variation. In cases of pronounced sexual dimorphism, males will often have antennae at about facial midlength or height, as in *Plumarius* and various Formicidae. As scored, this state is *TRUE* for Evaniidae, Embolemidae, Vespoidea *sensu stricto*, some Scoliidae, the Sphecidae, Bembicinae, some Pemphredonidae, Philanthidae, most Anthophila., and some Formicidae (*Gigantiops*, *Oecophylla*; Formicinae). This state is also *TRUE* for many evaniomorph fossils and most †Ephialtitidae, the latter with the exception of †*Sessiliventer*.
139 *DC: Clypeotorular structures. DS: Spatial relationships.* • **Observational criterion:** Antennal toruli deeply indenting aboral clypeal margin such that the clypeus extends aborally between the toruli (*FALSE* = toruli distant from clypeus OR toruli merely abutting toruli, with clypeus not extending between the torulus). **• NOTES:** This state was observed in most Chrysidoidea *sensu stricto*, Dryinidae, Tiphiidae, some Mutillidae, Ammoplanidae, *Pulverro* (Pemphredonidae), and most crown Formicidae, including †*Brownimecia*. Evaniidae are scored as uncertain ("?") because the posterior extent of the clypeus is unclear based on external examination. As for other torulus-location characters, sexual dimorphism may be observed, with males of some taxa having toruli distant from the clypeus.
140 *DC: Clypeotorular structures. DS: Spatial relationships.* • **Observational criterion:** Antennal toruli distant from clypeus by more than one torular diameter (*FALSE* = toruli closer to, abutting, or indenting aboral clypeal margin). **• NOTES:** Observed in a subset of all taxa which do not have the clypeus extending between the toruli. An intermediate state was scored in the present study, but it was poorly defined and was therefore removed before final analysis. However, there is further information which could be captured by providing well-defined states, such as distinguishing torular abutment of the clypeus and torular indentation, as well as relative metrics of torulus distance once removed from the posterior margin. Only the extreme state is retained with this character. With this scoring, extremely distantly set toruli were observed in *Gasteruptiidae*, †Othniodellithidae, Embolemidae, some Vespoidea *sensu stricto*, *Ammophila* (Sphecidae), *Pluto* (Pemphredonidae), Anthophila, and *Myrmoteras* and *Camponotus* (Formicinae, Formicidae). This latter pair is important, as distant toruli are a canonical diagnostic trait of the Camponotini, and the placement of *Myrmoteras* as sister to this tribe (*e.g*., Blaimer *et al*. 2015) has yet to be understood morphologically.
141 *DC: Toruli. DS: Relative positions.* • **Observational criterion:** Antennal toruli extremely wideset and situated just oral to the compound eyes, thus being directly in line between the mandibles and eyes (*FALSE* = toruli not extremely wideset, not between eyes and mandibles). **• NOTES:** Further information could be captured via characterization of relative intertorular spacing. The present state is only observed in the formicines *Santschiella* and *Gesomyrmex* (Formicidae).
142 *DC: Toruli. DS: Orientation.* • **Observational criterion:** Antennal toruli directed laterally, *i.e.*, away from one another; in this conformation, the medial torular arch is distinctly raised above lateral torular arch (*FALSE* = both toruli directed in parallel, whether anteriorly, ventrally, or dorsally so). **• NOTES:** Laterally directed toruli were observed in †Othniodellithidae, Trigonalidae, Sapygidae, some Mutillidae, Scoliidae, Ampulicidae, *Chalybion* (Sphecidae), and most Formicoidea. Among sampled Formicidae, reversed to dorsally directed (assuming prognathy) in †Haidomyrmecini, Leptanillinae, *Apomyrma* (Amblyoponinae), *Discothyrea* (Proceratiinae), and Dorylinae.
143 *DC: Toruli. DS: Orientation.* • **Observational criterion:** Antennal toruli directed toward the mandibles (*FALSE* = antennal toruli not directed orally). **• NOTES:** Observed in Scolebythidae, Bethylidae, †Chrysobythidae, †Falsiformicidae (including †*Burmasphex*), Sclerogibbidae, *Olixon* (Rhopalosomatidae), Sierolomorphidae, †*Bryopompilus* ("Pompilidae), Thynnidae, Chyphotidae, Mutillidae *sensu lato*, *Apterogyna* (Bradynobaenidae), Ampulicidae, Heterogynaidae, and *Chalybion* (Sphecidae).
144 *DC: Toruli. DS: Orientation.* • **Observational criterion:** Antennal toruli situated on a protuberance; in this location, their orifices are directed aborally (*FALSE* = toruli not on protuberance, not directed aborally). **• NOTES:** Observed in Evaniidae and *Loboscelidia* (Chrysididae); in both cases the toruli are situated on a diapriid-like prominence.
145 *DC: Toruli. DS: Orientation, form.* • **Observational criterion:** Antennal sockets concealed in full-face view by laterally expanded frontal carinae or medial torular arches (*FALSE* = sockets mostly or completely exposed in facial view). **• NOTES:** Socket concealment in this form was observed uniquely in the Formicidae, and within the ants in the poneroid clade (with the exception of *Apomyrma* and *Discothyrea*), a few Formicinae (*Santschiella*, *Gesomyrmex*, and *Camponotus*), and the sphecomyrmines †*Myanmyrma*, including †*M. mauradera*. In retrospect, this state should have been scored as *TRUE* for Ectatomminae *sensu lato*.

- *Antenna*

146 *DC: Antennomeres. DS: Count diphenism.* • **Observational criterion:** Antennomere count sexually diphenic (“dimorphic”) (*FALSE* = antennomere count equivalent between the sexes). **• NOTES:** Observed in Aulacidae, Gasteruptiidae, and Aculeata with the exception of the Chrysidoidea *sensu stricto* and Dryinoidea. Because of the scarcity of associated males and females for fossil taxa, most Mesozoic terminals are scored as uncertain ("?").
147 *DC: Antennomeres. DS: Count.* • **Observational criterion:** Female with > 12 antennomeres OR male with > 13 OR both ≥ 13 (*FALSE* = female with ≤ 12 OR male ≤ 13 OR both < 13). **• NOTES:** Chrysidoidea, Sclerogibbidae, and most sampled non-aculeate taxa have more than twelve antennomeres in the female, or more than thirteen in the male, or both. Exceptions include *Evaniella* (Evaniidae), †*Trigonalopterus* (†Ephialtitidae), all †Bethylonymidae (except the dubiously placed †*Allogaster*), and a subset of extinct chrysidoids.
148 *DC: Antennomeres. DS: Count.* • **Observational criterion:** Antenna 10-merous (*FALSE* = antenna > or < 10-merous). **• NOTES:** Ten-merous antennae are a canonical synapomorphy of the Dryinidae. Numerous taxa in the Formicidae have evolved reduced antennomere counts, among which are sampled *Myrmelachista joycei* and *Solenopsis invicta* with 10-merous antennae. As illustrated, it appears that †*Bethylonymellus bethyloides* has 10-merous antennae, but due to insufficient preservational detail, this taxon was scored as "?" for this state.
149 *DC: Radicle and scape. DS: Relative orientation.* • **Observational criterion:** Radicle ("bulbus neck", “neck of antennal articular process”) and scape offset at a distinct angle (*FALSE* = these parts at a slight angle or not angled). **• NOTES:** The term radicle is used here for the “bulbus neck” of the Formicidae (Keller 2011). Notably, a molecular developmental study of *Tribolium* demonstrated homology of the bulbus (“basal knob”) and radicle with the coxotrochanter, the scape with the femur, the pedicel with the tibia (hence Johnston’s organ with the tibial chordotonal organ), and the multi-annulate flagellum with the tarsus (Toegel *et al*. 2009). (Although be aware that Toegel *et al*. [2009] mislabeled the radicle and bulbus as the “antennifer”, which is part of the cranium by definition [Snodgrass 1935, 1956]). The angle of radicle and scape is offset in some Trigonalidae, Plumariidae (female only), the Bethylidae, Dryinidae, Sclerogibbidae, most Pompiloidea, most Scolioidea, Ampulicidae, Heterogynaidae, Sphecidae, Crabronidae, †*Camelomecia*, and a subset of Formicidae (*Paraponera*, *Tatuidris*, *Platythyrea*, *Acanthoponera*, and the examined Myrmicinae with the exception of *Stegomyrmex* which had a bent scape body and *Crematogaster*).
150 (**Additive**) *DC: Radicle and scape. DS: Relative orientation.* • **Observational criterion:** Radicle and scape perpendicular or nearly so (*FALSE* = these parts at a smaller angle). **• NOTES:** This extreme state was observed in Sapygidae, Mutillidae *sensu lato* (including the two new Cretaceous genera here described), Bradynobaenidae, and *Tatuidris* (Formicidae).
151 *DC: Proximomedial scape region. DS: Form.* • **Observational criterion:** Scape proximomedially produced as a distinct flange which can obscure the radicle from various perspectives (*FALSE* = scape not produced in this way). **• NOTES:** A flange obscuring the radicle is a canonical defining feature of Mutillidae (*e.g.*, Brothers & Lelej 2017). Similar but certainly convergent flanges occur in various Formicidae, such as the genera *Myrmica* and *Pogonomyrmex*.
152 *DC: Scape. DS: Shape.* • **Observational criterion:** Scape ovoid or subglobular in shape, being bulbous and swollen medially (*FALSE* = scape not bulbous medially). **• NOTES:** A subglobular scape was observed in various †Ephialtitoidea, †Baissidae, †Anomopterellidae, *Pristaulacus* (Aulacidae), †Maimetshidae, Trigonalidae, Plumariidae, *Clystopsenella* (Scolebythidae), some Vespidae (*Euparagia*, *Pseudomasaris*), *Pepsis* (Pompilidae), Bradynobaenidae, and *Chalybion* (Sphecidae). Generally, this state occurred in taxa with very short scapes, but not always.
153 *DC: Scape. DS: Proportion.* • **Observational criterion:** Scape “somewhat short”, *i.e.*, with length > 2 x width (*FALSE* = scape length ≤ 2 x width). **• NOTES:** Few sampled Aculeata have scapes which are shorter than 2 x their width.
154 (**Additive**) *DC: Scape. DS: Proportions.* • **Observational criterion:** Scape “intermediate”, *i.e.*, with length ≥ 4 x width (*FALSE* = scape length < 4 x width). **• NOTES:** Additive with respect to the prior character.
155 (**Additive**) *DC: Scape. DS: Relative proportions.* • **Observational criterion:** Scape “elongate”, *i.e.*, length greater than half head length (*FALSE* = scape length ≤ 0.5 x head length). **• NOTES:** Although not compared in the same manner as the prior two characters, this state is effectively additive as no sampled taxon had a scape which was greater than half head length but less than four times as long as wide. For †*Gerontoformica* (stem Formicidae), scoring was based on reported measurements in Barden & Grimaldi (2014).
156 *DC: Scape. DS: Relative proportions.* • **Observational criterion:** Scape length greater than half the length of the funiculus (= pedicel + flagellum) (*FALSE* = scape length ≤ 0.5 x funiculus length). **• NOTES:** This state is not additive with respect to the prior three because flagellar length varies independently. A long scape relative to flagellum is a putative synapomorphy of crown Formicidae (*e.g.*, Dlussky 1983, Borysenko 2017).
157 *DC: Flagellum. DS: Size, cuticle surface.* • **Observational criterion:** Flagellum grossly enlarged *and* with smooth surface (*FALSE* = scape not enlarged OR flagellar surface rough). **• NOTES:** Observed in male Bradynobaenidae. A similar state is observable in the male of the ponerine ant genus *Diacamma* which has rough flagellar cuticle.
158 *DC: Antennomeres. DS: Proportions.* • **Observational criterion:** Antennomere III longer than antennomere IV (*FALSE* = antennomere III shorter than IV). **• NOTES:** The length of antennomere III varies considerably across the sampled taxa. In some cases, antennomere III is extremely long, but further variation will need to be encoded in a future study. Scored for †*Gerontoformica* (stem Formicidae) based on metrics reported in Barden & Grimaldi (2014).
159 *DC: Flagellum. DS: Proportions, collective antennomere shape.* • **Observational criterion:** Apical five antennomeres forming massive, distinct, compact, and ovate club (*FALSE* = these five antennomeres not forming such a club). **• NOTES:** Although clubs of various forms are observed, particularly in the Formicidae (*e.g.*, Bolton 1994), this particular state was uniquely observed in Masarinae (Vespidae).
160 *DC: Chaetiform “setoid” sensilla. DS: Specification.* • **Observational criterion:** Paired chaetae (“bristle-like setae”) present at apices of flagellomeres (= "spines" *sensu* Guidotti 1999) (*FALSE* = such chaetae not developed). **• NOTES:** Unique to Rhopalosomatidae; reduced in *Olixon* (Krogmann *et al*. 2009) and *Eorhopalosoma gorgyra*, Engel 2008.
161 *DC: Villous “setoid” sensilla. DS: Specification.* • **Observational criterion:** Villous patches of setae present on flagellomeres; each seta is as long as or longer than each flagellomere (*FALSE* = such patches absent). **• NOTES:** Uniquely observed in *Plumarius* (Plumariidae). Not illustrated; see, *e.g.*, Brothers (1974a).
162 *DC: Slit-shaped sensilla zones. DS: Specification.* • **Observational criterion:** "Antennal dorsal organs" present. *i.e.*, multiple slit-like pores on dorsal surface of distal antennomeres (*FALSE* = these zones absent). **• NOTES:** According to Olmi in Perkovsky *et al*. (2019), these are observed in dryinid females which hunt Fulgoromorpha.

**-** *Mesosoma*

163 *DC: Winged mesosoma. DS: Overall shape.* • **Observational criterion:** Mesosoma very compact, *i.e.*, anteroposteriorly short and dorsoventrally tall, and heavily sclerotized (*FALSE* = mesosoma neither so compact nor so sclerotized). **• NOTES:** Observed in Evaniidae and related, extinct taxa. State 1 of character 25 in Ronquist *et al*. (1999) and character 68 of Li *et al*. (2018).
164 *DC: Winged mesosoma. DS: Proportions, shape.* • **Observational criterion:** Mesosoma proportionally massive, being greatly enlarged relative to the head and metasoma thus appearing “steroidal”, swollen, or ballooned (*FALSE* = mesosoma smaller; compare Figs. 164-0 and C164-1). **• NOTES:** Humongous mesosomata were observed sporadically among sampled Aculeata, including *Pseudomasaris* (Vespidae), male Bradynobaenidae, and various male Formicidae. The specific morphological pattern of enlargement, however, differs among each of these groups.
165 *DC: Wingless mesosoma, promesonotal articulation. DS: Mobility.* • **Observational criterion:** Promesonotal articulation immobile, with the two segments fused, whether or not a suture is apparent (*FALSE* = this articulation mobile in apterous individuals OR aptery absent). **• NOTES:** (1) "Suture" is used here in the strict sense, meaning a line between two sclerites which were ancestrally unfused. (2) Among clades with aptery, promesonotal fusion occurs in Mutillidae, Heterogynaidae, and Formicidae; such fusion is absent in *Plumarius*, *Pristocera*, *Olixon*, and the Thynnoidea (the state of *Sclerogibba* is uncertain). In crown Formicidae promesonotal fusion was observed in Paraponerinae, Agroecomyrmecinae, Proceratiinae, *Lioponera* (Dorylinae), *Lepisiota* (Formicinae), *Rhytidoponera* (Ectatomminae), and Myrmicinae. Notably, †Zigrasimeciinae includes species with and without such fusion (Cao *et al*. 2020b). Although macropterous, many sphecids have "rigidly" or "solidly" articulated pro- and mesonota (Bohart & Menke 1976). (3) Several scoring scenarios were considered for this character because it is clear that promesonotal fusion has evolved independently in *Heterogyna*, Mutillidae, and Formicidae. Because there is strong *a priori* evidence that promesonotal fusion is the derived state, all taxa without apterous individuals were scored as *FALSE*, thus allowing the three convergent origins to be scored as *TRUE* in the same column, without having to split the character into more than one reductive taxic characters.
166 (**Taxic-reductive**: Thynnoidea) *DC: Wingless mesosoma. DS: Overall shape.* • **Observational criterion:** Mesosomal profile diagonal relative to the long axis of the body, with promesonotum elevated above propodeum and often convex or strongly humped (*FALSE* = mesosoma linear or compact, dorsal margin linear). **• NOTES:** A diagonal mesosomal profile of apterous individuals was observed in Thynnoidea (*Methoca*) and Formicidae. Because the polarity of this state is not clear *a priori*, it was not scored similarly as the prior character (*i.e.*, as *FALSE* for inapplicables). Rather, the character is here taxically split between the Thynnoidea (this character) and the Formicoidea (next character). This was done in order to obviate "missing data attraction" (the draw of terminals together with filled cells relative to terminals with "-" or "?"), as well as to be neutral about the ancestral state of this condition. Among other clades with scored apterous species, a linear (rather than diagonal) profile was observed (*Plumarius*, *Pristocera*, *Sclerogibba*, *Olixon*, Mutillidae, Bradynobaenidae, *Heterogyna*).
167 (**Taxic-reductive**: Formicidae) *DC: Wingless mesosoma. DS: Overall shape.* • **Observational criterion:** Mesosomal profile diagonal relative to the long axis of the body, with promesonotum elevated above propodeum and often convex or strongly humped (*FALSE* = mesosoma linear or compact, dorsal margin linear). **• NOTES:** Among the Formicidae, this was the most frequently observed state, including for stem taxa. During the quality-control phase of scoring an additional character, "promesonotum distinctly domed or humped", was thrown out as it was additive with respect to the present character and provided almost no grouping information.
168 *DC: Carination. DS: Specification.* • **Observational criterion:** Pronotum of wingless individuals “keeled”, *i.e.*, with a medially situated longitudinal carina in the posterior pronotal half (*FALSE* = aptery absent OR pronotum without keel). **• NOTES:** Observed uniquely in *Heterogyna* (Heterogynaidae).
169 *DC: Wingless mesothorax. DS: Circumference.* • **Observational criterion:** Mesothorax strongly constricted ("strangled"), thus giving the mesosoma an "hourglass" shape in profile or dorsal view (*FALSE* = aptery absent OR mesothoracic constriction absent). **• NOTES:** This character is scored in a manner similar to that of promesonotal fusion. Specifically, as a diminishing minority of apterous taxa are *TRUE* for this state there is evidence that it is a derived state, thus terminals without apterous individuals are scored as *FALSE*. Observed in *Methoca* (Thynnidae), and sporadically in crown Formicidae, particularly in the Formicinae (*e.g.*, *Prenolepis*), and among a few species of †*Gerontoformica* (†*G. gracilis*, †*G. pilosa*) plus †*Sphecomyrma freyi*. This mesosomal conformation may represent some skeletomuscular optimization for head movement and locomotion, but functional implications have yet to be tested.
170 *DC: Wingless mesonotum. DS: Differentiation.* • **Observational criterion:** Mesoscutum differentiated into the anterior mesoscutum and posterior mesoscutellum, with their boundaries marked by a transverse keel, hence the mesoscutum is angled in profile view (*FALSE* = aptery absent OR mesonotum without scutal-scutellar differentiation; mesonotum rounded or linear in profile view). **• NOTES:** Because this state is observed uniquely in the Formicidae, and with a restricted taxonomic distribution therein, it is scored as *FALSE* for groups without apterous individuals. Within the Formicidae, this state is only observed in some †*Gerontoformica* species.
171 *DC: Bulge between mesonotum, propodeum in wingless individuals. DS: Expression.* • **Observational criterion:** Distinct bulge present between dorsal plate of mesothorax and propodeum (*FALSE* = aptery absent OR this plate not developed). **• NOTES:** The description of this character is homology neutral. In the total clade of the Formicidae, a distinct plate between the mesonotum and propodeum is sometimes visible, but whether this plate represents the mesoscutellum or metanotum is challenging enough to determine that such a decision is not made here. In stem ants, it is reasonably determinable that this bulge corresponds to the metanotum, as the mesonotum can be differentiated into the mesoscutum and scutellum (see character 170). Because this plate can be present above the mesothoracic or the metathoracic segment or both, referring to it with reference to segment is potentially misleading. Among sampled taxa, the *TRUE* state was applied to *Sclerogibba*, *Dasymutilla*, *Leptomyrmex*, and some Formicinae. Although *Olixon* and *Heterogyna* are generally treated as "apterous" here, they do have wings and functional mesoscutella plus distinct metanota, thus these terminals are scored as *FALSE*.
172 (**Taxic**: Plumariidae) *DC: Wingless mesometanotal articulation. DS: Mobility.* • **Observational criterion:** This articulation freely mobile (*FALSE* = this articulation immobile). **• NOTES:** This unusual condition was observed in *Plumarius*, *Pristocera*, and *Methoca*. To avoid drawing these taxa together based on an obvious convergence, this character is split into three taxic characters, each providing a unique state for each terminal. Moreover, it is expected that detailed anatomical study would demonstrate that the condition among these three taxa is distinct.
173 (**Taxic**: Bethylidae) *DC: Wingless mesometanotal articulation. DS: Mobility.* • **Observational criterion:** This articulation freely mobile (*FALSE* = this articulation immobile). **• NOTES:** See NOTES for prior character.
174 (**Taxic**: Thynnoidea) *DC: Wingless mesometanotal articulation. DS: Mobility.* • **Observational criterion:** This articulation freely mobile (*FALSE* = this articulation immobile). **• NOTES:** See NOTES for prior character.

- *Prothorax*

175 *DC: Anteromedian pronotal lobe. DS: Expression.* • **Observational criterion:** Pronotum with anteromedian lobe forming a "neck" which receives the head when the head is flexed upwards (*FALSE* = such a lobe absent). **• NOTES:** In evaniomorph outgroups the anterior pronotal face does not curve anteriorly, thus there is no neck. Among the Aculeata, a neck is absent in those taxa which support the cranium with the propleurae. Otherwise, the neck is sporadically absent. When the neck is present, it can be present as a rim of more-or-less uniform width as seen in dorsal or dorsolateral oblique view, or it can be present as a distinct medially situated lobe of varying proportions. This rim to lobe variation is true within the Formicidae, and such variation may be informative particularly if Cenozoic fossils are included; indeed, from the perspective of the phylogenetic modeling of continuous variation, it would be ideal to put this variation to use.
176 *DC: Pronotum of alate individuals. DS: Form of margin.* • **Observational criterion:** Alate: pronotum with anterior rim upturned, often bearing fringe of setae (*FALSE* = such a rim absent). **• NOTES:** This state was observed in †*Camelomecia*. See also Barden & Grimaldi (2016). Anatomically, this state can be differentiated from that of having a narrow "neck rim" or complete transverse pronotal sulcus as observed in Cleptinae (Chrysididae), as the "upturned rim" represents an even, very narrow margin of the pronotum which is delimited by a complete groove. This state is analogous to that of having an "epipleurite" on the elytron, as used by coleopterists.
177 *DC: Pronotum of alate individuals. DS: Shape.* • **Observational criterion:** Alate: pronotum muscular and elongate, *i.e.*, with distinct dorsal surface face offset from anterior or declivitous face (*FALSE* = pronotum short, without distinct dorsal face). **• NOTES:** An elongate pronotum is associated with head mobility and strength, thus has implications for object/prey handling and sociality (see, *e.g.*, Keller *et al*. 2014). Here observed in various Aculeata and the enigmatic †Bethylonymidae.
178 (**Additive**) *DC: Pronotum of alate individuals. DS: Shape.* • **Observational criterion:** Alate: pronotum in dorsal view nearly square, with anteroposterior length of disc ≥ lateromedial width of disc, being rectangular to broadly rhomboidal or anteriorly rounded (*FALSE* = pronotum shorter, disc length < width). **• NOTES:** Scored as *TRUE* for some Bethylidae, *Loboscelidia* (Chrysididae), some extinct Scolebythidae, Sclerogibbidae, *Olixon* (Rhopalosomatidae), †*Falsiformica* (†Falsiformicidae), Sierolomorphidae, some fossil Ampulicidae, †*Camelomecia*. Females were scored in cases where sexual dimorphism affects this trait (*e.g.*, Plumariidae).
179 *DC: Pronotum. DS: Shape.* • **Observational criterion:** Pronotum with dorsolaterally situated longitudinal margination between lateral and dorsal surfaces, *i.e.*, dorsal and lateral surfaces offset at distinct angle (*FALSE* = dorsal and lateral surfaces not offset at angle). **• NOTES:** Observed infrequently among sampled aculeates, including some extinct Chrysididae, †*Falsiformica* (†Falsiformicidae), some Dryinidae, Sclerogibbidae, Ampulicidae, and some Formicidae (*e.g.*, *Neoponera*, *Pseudomyrmex*, *Tetraponera*, *Wasmannia*, *Crematogaster*, and the new burmite ponerine).
180 *DC: Female pronotum. DS: Shape.* • **Observational criterion:** Female: disc pronotum distinctly and acutely triangular, distinctly longer than broad in dorsal view and lateromedially narrow (*FALSE* = disc broader, rhomboidal to subrectangular to dome shaped). **• NOTES:** Observed uniquely in some †*Hybristodryinus*; see Perkovsky *et al*. (2019).
181 *DC: Female pronotum. DS: Shape.* • **Observational criterion:** Female: disc pronotum with distinct round process situated dorsomedially, thus giving the pronotum a humped appearance (*FALSE* = disc without median hump). **• NOTES:** Among sampled taxa, observed uniquely in *Dryinus gulfensis* (Dryinidae). Some *Ectatomma* (Formicidae) also have dorsomedian pronotal humps.
182 *DC: Female pronotum. DS: Shape.* • **Observational criterion:** Female: pronotum with grossly expanded flange, forming broad collar around the propleurae and partially encircling the posteromedian muscular portion of the notum (*FALSE* = pronotum without such a flange). **• NOTES:** This state was observed uniquely in female Dryinidae.
183 *DC: Pronotal corner. DS: Shape.* • **Observational criterion:** Anteroventral pronotal corner with distinct spiniform projection (*FALSE* = corner rounded). **• NOTES:** Such pronotal spines occur sporadically among the Formicidae, including some Amblyoponinae, leaf cutter ants (*Atta*, *Acromyrmex*), and various Ectatomminae *sensu lato*.
184 *DC: Pronotal crease of alate individuals. DS: Shape.* • **Observational criterion:** Alate: transverse line or groove on lateral or lateral and medial surfaces of pronotum (*FALSE* = pronotum without such a crease). **• NOTES:** This line or groove probably corresponds to the "anteromedian pronotal ridge" which dorsally delimits the muscular origins of the pronotopropleural arm muscles (*e.g.*, HAP URI: purl.obolibrary.org/obo/HAO_0000131; Vilhelmsen *et al*. 2010), and that has been previously termed the “transnotal sulcus” (Vilhelmsen 2000). It was not possible, however, to confirm this identity, however, as dissections were not conducted. The line observed here is expressed sporadically among the Evanioidea and Aculeata but was consistent in Trigonalidae. It was not observed in any Formicoidea. In extant dryinids, the line was easily observed in males; in females, it appeared that the notal portion which would be anterior to the sulcus or line is grossly expanded, thus forming the "collar" of described above.
185 *DC: Pronotal crease of alate individuals. DS: Extent, shape.* • **Observational criterion:** Alate: pronotum completely divided by a deep, transverse, arc-shaped sulcus; anterior portion of pronotum broad and more-or-less even in width across its length (*FALSE* = crease absent, incomplete, or complete and narrow). **• NOTES:** This state is a putative synapomorphy of the Cleptinae (Chrysididae; character 10 of Kimsey & Bohart 1990). Among fossil taxa, †*Hypocleptes* and a new burmite cleptine were scored as *TRUE*, whereas the state in †*Procleptes hopejohnsonae* is uncertain. also scored as *TRUE* for *Ampulex*, *Chalybion* as the sulcus is almost complete).
186 *DC: Pronotal crease of alate individuals. DS: Extent, shape.* • **Observational criterion:** Alate: pronotum completely divided by a shallow, transverse, arc-shaped sulcus; anterior portion of pronotum narrowed anteromedially (*FALSE* = pronotal crease absent, incomplete, or complete and broad). **• NOTES:** This state was observed in Ampulicidae, Sphecidae, and *Plenoculus* (Crabronidae).
187 *DC: Pronotal sides. DS: Shape.* • **Observational criterion:** Pronotum scrobiculate, *i.e.*, with a more-or-less dorsoventrally oriented groove or concavity which receives the forelegs when these are upfolded forelegs, often associated with lateral bulge of mesopectus (*FALSE* = pronotal sides flat or convex, not shaped to receive legs). **• NOTES:** Deep and broad lateral impressions were observed in †Othniodellithidae, Bethylidae, most Chrysididae, Dryinidae, †Falsiformicidae, *Sierolomorpha* (Sierolomorphidae), *Tiphia* and †*Thanatotiphia* (Tiphiidae), Bradynobaenidae, and most Apoidea.
188 *DC: Pronotal carina of alate individuals. DS: Location.* • **Observational criterion:** Alate: pronotum with laterally situated dorsoventrally oriented carina (*FALSE* = such a carina absent, or carina only present medially). **• NOTES:** A lateral, dorsoventral carina was observed sporadically among sampled taxa, including Evaniidae, Chrysidinae (Chrysididae), some Vespidae, some Pompiloidea *sensu lato*, some Apoidea, and a very few Formicidae (*Tatuidris*, *Wasmannia*). Such carination was often associated with presence of a dorsal transverse carina.
189 *DC: Pronotal carina of alate individuals. DS: Location.* • **Observational criterion:** Alate: pronotum with dorsally situated carina which is continuous across dorsomedian surface of sclerite (*FALSE* = such a carina absent, or carina only present laterally). **• NOTES:** As noted in the prior character, this state is usually associated with presence of a dorsoventral lateral carina, but not always (*e.g.*, extant Scolebythidae).
190 *DC: Posterolateral pronotal carina of alate individuals. DS: Expression.* • **Observational criterion:** Alate: pronotum with sharply margined dorsoventral carina extending from ventral margin of pronotal lobe to dorsolateral corner of pronotum, just above tegula (*FALSE* = such a carina absent). **• NOTES:** This specific carina was observed in *Zethus*, *Pachodynerus,* and *Mischocyttarus* (Vespidae), as well as *Campsomeris pilipes* (Scoliidae). A sulcus which is posteriorly margined by a rounded carina was observed in *Euparagia*, but this was not scored as *TRUE* for the present character.
191 *DC: Pronotum of alate individuals. DS: Shape.* • **Observational criterion:** Alate: pronotum broadly constricted posteriorly, just anterior to mesoscutum (*FALSE* = pronotum not constricted posteriorly). **• NOTES:** This state observed in *Pristocera* (Bethylidae), *Pepsis* (Pompilidae), *Dryinus* (Dryinidae), spheciform apoids, †*Camelomecia*.
192 *DC: Pronotum of alate individuals. DS: Shape.* • **Observational criterion:** Alate: pronotum with muscular posterior portion separated from mesoscutum by deep and narrow incision (*FALSE* = pronotum not constricted, or constriction broad). **• NOTES:** Observed uniquely in non-anthophilan Apoidea.
193 *DC: Posterior pronotal collar of alate individuals. DS: Shape.* • **Observational criterion:** Alate: pronotum posterior collar notched medially (*FALSE* = collar convex or flat, without median concavity). **• NOTES:** Observed in *Clypeadon* and *Philanthus* (Philanthidae), as well as *Dieunomia* (Halictidae).
194 *DC: Posterior pronotal margin of alate individuals. DS: Shape.* • **Observational criterion:** Alate: pronotum posterior margin in dorsal view narrowly arcuate (*FALSE* = this margin broadly arcuate or more-or-less straight). **• NOTES:** The posterior pronotal margin shows considerable variation across the Aculeata, ranging from linear to narrowly arcuate, as scored here. However, it was difficult to discretely categorize variation other than this particular extreme state, although Brothers (1975) did provide a series of alternate states. Narrowly arcuate posterior pronotal margins were uniquely observed in Vespidae, wherein this state was associated with anteroposterior narrowing of medial portion of pronotum in dorsal view and retention of truncate posterolateral region. Among the sampled fossils, observed in †*Protovespa haxairei* and †*Symmorphus senex*.
195 *DC: Pronotum of alate individuals. DS: Shape.* • **Observational criterion:** Alate: lateral portions of pronotum ventral/lateral to mesoscutum dorsoventrally expanded, visible as lateromedially broad shoulders in dorsal view (*FALSE* = these portions narrow OR not “clasping” mesoscutum laterally). **• NOTES:** Observed in all extant Vespidae except for *Parischnogaster* (Stenogastrinae), which has a narrow nearly digitate lateral pronotal portion.
196 *DC: Pronotum of alate individuals. DS: Shape.* • **Observational criterion:** Alate: lateral portions of pronotum ventral/lateral to mesoscutum in lateral view in form of very narrow, acute triangles with their apices at the tegulum (*FALSE* = these portions dorsoventrally broader). **• NOTES:** Uniquely observed *Parischnogaster* (Stenogastrinae, Vespidae).
197 *DC: Pronotum of alate individuals. DS: Shape.* • **Observational criterion:** Alate: lateral portions of pronotum ventral/lateral to mesoscutum dorsoventrally expanded and posteriorly notched (*FALSE* = these portions not expanded, OR if broad then without notch). **• NOTES:** This character captures the unique state observed in Scoliidae, which is reminiscent of both Vespidae and Apoidea.
198 *DC: Pronotal lobe. DS: Shape.* • **Observational criterion:** Pronotal lobe, concealing first mesosomal spiracle, grossly enlarged and dorsoventrally compressed, thus appearing as distinct semicircular extension of pronotum (*FALSE* = pronotal lobe not enlarged, OR only indistinctly bulging from pronotum). **• NOTES:** (1) This is a canonical synapomorphy of the Apoidea. A similar conformation was observed in *Orthogonalys* (Trigonalidae) but was less pronounced thus scored as *FALSE*. Among putative apoid compression fossils, the only taxa for which this state could be confirmed were †*Baissodes robustus*, †*Oryctobaissodes armatus*, and †*Cretosphecium lobatum*. The former two are Aptian "angarosphecid" fossils described by Rasnitsyn (1975) from the Zaza formation in Baissa (Russia), while †*C. lobatum* is an Aptian compression from Mongolia described by Pulawski and Rasnitsyn (Pulawski *et al*. 2000). Among the amber taxa attributed to Apoidea, this state could not be confirmed for but two species (†*Gallosphex cretaceus* and †*Psolimena singularis*). (2) During the proofing process of this matrix "posterior truncation" of the posterolateral pronotal margin was deleted as a character due to arbitrariness. There is considerable variation of the lateral region of the pronotum and of the posterolateral pronotal margin; this variation is almost certainly informative but needs further—perhaps quantitative—study to employ meaningfully.
199 *DC: Pronotum of alate individuals. DS: Shape.* • **Observational criterion:** Alate: pronotum separated from tegula, *i.e.*, mesopectus and mesoscutum contacting one another between pronotal lobe and tegula (*FALSE* = pronotum extending to tegula, separating mesoscutum and mesopectus). **• NOTES:** This is another canonical feature of the Apoidea, although it is not *TRUE* for Ampulicidae and Heterogynaidae, including the new genus CASENT0844594. As for the prior apoid character, this state was impossible to evaluate for the vast majority of compression fossils attributed to the superfamily. Pronotal separation from the tegula also occurs in Chrysidinae (Chrysididae), for which this has been scored as character 11 of Kimsey & Bohart (1990), and the female of *Dryinus gulfensis.* It was confirmed that the pronotum is indeed in contact with the tegulum in †Chrysidinae sp. (CASENT0844585, burmite). Finally, given Krombein’s (1983) reinterpretation of †*Protamisega* relative to the original description (Evans 1973), the separation of the pronotum from the tegula in †*Hypocleptes rasnitsyni* Evans, 1973 is here treated as uncertain ("?").
200 *DC: Posteroventral pronotal corners. DS: Shape.* • **Observational criterion:** Posterolateral corners produced medially posterior to procoxae, thus pronotum forming or almost forming complete ring (*FALSE* = these corners not extending behind coxae). **• NOTES:** This is another canonical synapomorphy of the Apoidea and was described by Brothers (1975) as his character 23 state 2. †*Camelomecia* sp. (ANTWEB1038930) and †*Camelomecia* (male, NIGPAS) clearly have pronota which do not extend medially posterior to the procoxae. It can be seen that the ventrolateral margin of the putative apoid †*Burmasphex* ends dorsal to the procoxa in Fig. 12 of Melo & Rosa (2018).
201 *DC: Propleurae. DS: Shape.* • **Observational criterion:** Propleurae produced anteriorly and forming narrow tube-like collar supporting head (*FALSE* = propleurae not extending beyond pronotum nor supporting head). **• NOTES:** Observed in most Evanioidea, *Orthogonalys* (Trigonalidae), †Maimetshidae, Plumariidae, Scolebythidae, and the female of *Dryinus gulfensis* (Dryinidae). Scores as *TRUE* for †Plumalexiidae based on the original description in Brothers (2011), wherein the propleurae are described as being "anteriorly produced as short neck" (p. 525). Given the distribution of the *TRUE* state of this character among sampled taxa, as well as presence in the "symphyta", this is probably an ancestral state of the Aculeata.
202 (**Additive**) *DC: Propleurae. DS: Fusion.* • **Observational criterion:** Propleurae fused dorsally and ventrally, forming solid tube-like neck supporting head (*FALSE* = propleurae not fused). **• NOTES:** This well-known state (*e.g.*, Goulet & Huber 1993) was observed uniquely in the female of *Plumarius* (Plumariidae).
203 *DC: Ventral surfaces of propleurae. DS: Shape.* • **Observational criterion:** Propleurae deeply concave, receiving cranium when head tucked posteriorly (*FALSE* = propleurae flat to convex). **• NOTES:** Observed uniquely in Scoliinae (Scoliidae); as for many scoliid characters, the state for *Proscolia* (Proscoliinae) is uncertain.
204 *DC: Prosternum. DS: Size, shape.* • **Observational criterion:** Prosternum massive and broadly exposed (*FALSE* = prosternum smaller, relatively concealed). **• NOTES:** A massively enlarged prosternum has been used to place several Cretaceous fossils in the Scolebythidae, but this state is also observed in Chrysididae (except Amiseginae and Loboscelidiinae; character 13 of Kimsey & Bohart 1990). Putative Mesozoic scolebythids which were difficult to interpret include †*Boreobythus*, †*Mirabythus*, †*Uliobythus*, and †*Zapenesia*; the state of this character for these taxa has been scored as uncertain ("?").
205 *DC: Prosternum. DS: Shape.* • **Observational criterion:** Prosternum almost entirely concealed, exposed portion in form of longitudinally oriented lamella (*FALSE* = prosternum not medially lamellate, not as concealed). **• NOTES:** Observed in Scoliidae.

- *Mesothorax*

206 *DC: Mesoscutum of alate individuals. DS: Proportions.* • **Observational criterion:** Alate: mesoscutum longer than broad in dorsal view (*FALSE* = broader than long, OR as broad as long). **• NOTES:** Most scored taxa have mesoscuta which are shorter than wide. Exceptions include Aulacidae, Gasteruptiidae, †Scolebythidae sp. (CASENT0844570, burmite), Rhopalosomatinae (Rhopalosomatidae), some Vespidae, *Pepsis* (Pompilidae), and various Formicidae (*Discothyrea*, *Leptomyrmex*, *Linepithema*, some Formicinae, and some Myrmicinae).
207 *DC: Mesoscutum of alate individuals. DS: Proportions.* • **Observational criterion:** Alate: mesoscutum width nearly twice length (*FALSE* = width distinctly less than twice length). **• NOTES:** Very wide and short mesoscuta were observed in various fossil Scolebythidae, most Bethylidae, most Chrysididae, †Falsiformicidae, most Dryinidae, *Olixon* (Rhopalosomatidae), and some Formicidae (Leptanillinae, *Apomyrma*).
208 *DC: Mesoscutum of alate individuals. DS: Shape.* • **Observational criterion:** Alate: entire mesoscutum bulgingly produced dorsally, but bulge anterolaterally "shouldered", thus appearing subrectangular in dorsal view (*FALSE* = mesoscutum not bulging, OR if bulging, then bulge not appearing subrectangular). **• NOTES:** Among sampled taxa, uniquely observed Ammoplanidae. Some male Formicidae have similarly bulging mesoscuta. Note that the specimen available for depiction is not as developed as the scored species.
209 *DC: Mesoscutal disc of alate individuals. DS: Shape.* • **Observational criterion:** Alate: median portion of mesoscutum, *i.e.*, that region of the mesoscutum between the notauli and/or parapsides, enlarged and bulging dorsally above mesopectus in profile view (*FALSE* = central portion of mesoscutum not bulging). **• NOTES:** Observed in Aulacidae, extant Gasteruptiidae, and †*Electrobaissa* (†Baissidae). †*Hypselogastrion* (Gasteruptiidae) does not have a bulging median mesoscutal portion, and the kotujellines †*Kotujellites* and †*Kotujisca* could not be evaluated for this state due to their preservation.
210 *DC: Mesoscutum of alate individuals. DS: Shape.* • **Observational criterion:** Alate: median portion of mesoscutum bulging medially and with distinct lateral faces that are offset at an angle from the lateral mesoscutal portions (*FALSE* = mesoscutum not bulging, OR if bulging then medial and lateral portions not meeting at angle, *i.e.*, more-or-less in the same plane). **• NOTES:** Observed uniquely in *Pristaulacus* (Aulacidae).
211 *DC: Mesoscutal disc of alate individuals. DS: Shape.* • **Observational criterion:** Alate: mesoscutum with large but low median bulge between parapsides (*FALSE* = mesoscutum not bulging OR strongly bulging, *i.e.*, bulge high and incorporating most of medial mesoscutal portion or the entire mesoscutum). **• NOTES:** Observed in the scoliids *Trisciloa* and *Campsomeris*. Corresponds to a pair of unusual and unexplained sac-like structures which develop during pupation (see Fig. 9 in Scharfy 2012).
212 *DC: Mesoscutum of alate individuals. DS: Shape, proportions.* • **Observational criterion:** Alate: lateral portions of mesoscutum massively enlarged, such that each is wider than medial portion and the two together with conspicuously greater surface area (*FALSE* = these portions smaller, surface area much less than medial portion). **• NOTES:** Observed in †Falsiformicidae, †*Burmasphex* ("†Angarosphecidae”), and some Dryinidae.
213 *DC: Lateral mesoscutal carina of alate individuals. DS: Expression.* • **Observational criterion:** Alate: parascutal carina present, *i.e.*, mesoscutum with a carina which margins the pit formed by the forewing insertion (*FALSE* = this carina absent, thus the scutum curves evenly from the disc into the foramen). **• NOTES:** Parascutal carinae were observed in Evanioidea, *Bareogonalos* (Trigonalidae), most Chrysididae, †Falsiformicidae, †*Burmasphex* ("†Angarosphecidae), some Dryinidae, most Vespoidea *sensu stricto*, some Pompiloidea *sensu lato*, most Scolioidea (state could not be confirmed for *Proscolia*), most Apoidea (some bees too fuzzy to evaluate), †*Camelomecia*, and many Formicidae. Among the Formicidae, absent in the sampled *Discothyrea*, *Simopelta*, *Hypoponera*, most Dolichoderinae (except *Dolichoderus*), and all Formicinae. In *Rhopalosoma* (Rhopalosomatidae), the carina is absent anteriorly, but is present posteriorly in association with the parapsidal line. Carpenter (2000) in his description of ?*Symmorphus senex* uses the term "pretegular carina", which is here interpreted as a synonym of "parascutal carina".
214 *DC: Parascutal carina of alate individuals. DS: Shape.* • **Observational criterion:** Alate: posterior apex of parascutal carina (as defined above) expanded laterally to posterolaterally as lobate, laminate, or subdigitate process (*FALSE* = parascutal carina absent, OR if present then carina forming more-or-less even curve, not produced posterolaterad tegula). **• NOTES:** Observed uniquely in *Zethus*, *Pachodynerus*, and *Mischocyttarus* (Vespidae). "Parategula" of authors. The lobate expansion of *Mischocyttarus* is here scored as *TRUE*, although this structure is treated as absent in Polistinae (*e.g.*, Goulet & Huber 1993).
215 *DC: Posterolateral mesoscutal carinae of alate individuals. DS: Expression.* • **Observational criterion:** Alate: mesoscutum with "oblique scutal carina", *i.e.*, a transverse to diagonal carina situated on the posterolateral lobe of the mesoscutum which overhangs the tegula (*FALSE* = such a carina absent). **• NOTES:** This state is an apomorphy observed in the apoid groups Nyssonini and Gorytini (Bembicidae, Nyssoninae) as well as Mellinidae (Bohart & Menke 1976). Among sampled taxa, the "oblique scutal carina" only occurs in *Bembix* (Bembicidae).
216 *DC: Mesoscutum of alate individuals. DS: Midline development.* • **Observational criterion:** Alate: mesoscutal anteromedian line externally indicated by ridge or groove (*FALSE* = anteromedian line not indicated externally). **• NOTES:** A distinction is made between the "median mesoscutal sulcus" and "median mesoscutal line" (*e.g.*, Rasnitsyn 1988), as the former is associated with an internal ridge, whereas the latter is apparently not. Because dissection was not employed for extant taxa in the present study, and as the majority of fossils studied were not subjected to µ-CT scanning, we were unable to make this distinction. Here, an anteromedian line of any form was observed on the mesoscutum of most ephialtitoids, most evaniomorphs, †Bethylonymidae, most trigonaloids, and a majority of Aculeata. Among aculeates, the line was not observed in most Chrysidoidea (exceptions include *Plumarius*, *Cleptes*), Embolemidae, Sclerogibbidae, most Pompiloidea *sensu lato* (except Pompilidae, Sapygidae, and *Myrmosa*), some Apoidea (*Dolichurus*, *Heterogyna*, Ammoplanidae, *Dieunomia*), and most Formicidae (except, *e.g.*, *Dolichoderus*, most Formicinae, and most Myrmicinae). Recently this feature has been defined as the "medial mesoscutal suture" (Li *et al*. 2018, character 70), although this is not a true suture.
217 (**Reductive**) *DC: Mesoscutal grooves of alate females. DS: Expression.* • **Observational criterion:** Alate females: notauli present on mesoscutum (*FALSE* = notauli absent). **• NOTES:** The rampant sexual dimorphism of this trait was understood too late to code both sexes for the present study. In general, female notauli were observed in most evaniomorphs, most Chrysidoidea (except Chrysidinae), Dryinidae, †Falsiformicidae, various Vespidae, Fedtschenkia (Sapygidae), Protoscolia and Sinoproscolia (Scoliidae), Ampulicidae, and most Pemphredonidae. No alate female Formicidae were observed to retain notauli, although notaular variation of males is informative for diagnostic purposes. Scored as "-" if alate females absent for a given species.
218 (**Reductive**) *DC: Notauli of alate females. DS: Extent.* • **Observational criterion:** Alate female: notauli complete, *i.e.*, reaching transscutal line posteriorly (*FALSE* = notauli not developed OR incomplete). **• NOTES:** Where female notauli were observed, they were often "complete". Scored as "-" if alate females absent for a given species, or if notauli not observed.
219 (**Additive-reductive**) *DC: Notauli of alate females. DS: Extent, orientation.* • **Observational criterion:** Alate female: notauli complete AND parallel to somewhat convergent, extending from the anterior mesoscutal margin to the posterior, dividing the mesoscutum into three distinct regions but WITHOUT posterior termini close-set (*FALSE* = notauli absent OR strongly converging posteromedially, thus median mesoscutellar portion appearing V shaped, OR notauli converging completely). **• NOTES:** Complete and more-or-less parallel notauli were observed in extant Scolebythidae, some Bethylidae, some Chrysididae, Dryinidae, †Falsiformicidae, †*Burmasphex* ("Angarosphecidae”), †Mutillidae sp. (CASENT0844578), some Ampulicidae, and some fossil apoids. Notauli in Trigonalidae are distinctly convergent, although their posterior termini are distantly separated. Scored as "-" if alate females absent for a given species, or if notauli not observed.
220 (**Reductive**) *DC: Notauli of alate females. DS: Extent, orientation.* • **Observational criterion:** Alate female: notauli meeting posteromedially anterior to transscutal carina, thus forming Y-shaped groove (*FALSE* = notauli absent OR not meeting medially anterad scutoscutellar line). **• NOTES:** Y-shaped female notauli were observed in Aulacidae, Gasteruptiidae, and *Zethus caridei* and †*Priorparagia* (Vespidae). Scored as "-" if alate females absent for a given species, or if notauli not observed. Notaular form is variable in Gasteruptiidae.
221 *DC: Mesoscutellum of alate individuals. DS: Shape.* • **Observational criterion:** Alate: mesoscutellum produced posteriorly, overhanging propodeum in lateral view and obscuring propodeum in dorsal view (*FALSE* = mesoscutellum not produced, OR if produced then not obscuring propodeum in dorsal view nor overhanging propodeum as seen in lateral view). **• NOTES:** Observed in *Bembix* (Bembicidae), some most sampled Apidae, including †*Cretotrigona*.
222 *DC: Mesoscutellum of alate individuals. DS: Shape.* • **Observational criterion:** Alate: mesoscutellum strongly flattened, and with lateromedially broad obtusely triangular laminar processes produced posteriorly over metasoma (*FALSE* = mesoscutum convex OR if flattened then not laminar nor overhanging metasoma). **• NOTES:** Observed uniquely in the spectacular cuckoo bee genus *Thyreus* (Apidae).
223 *DC: Prepectus of alate individuals. DS: Expression.* • **Observational criterion:** Alate: prepectus present (*FALSE* = prepectus absent). **• NOTES:** The prepectus is an intersegmental sclerite between pronotum and mesopectus and bears the origin of the anterior thoracic spiracle occlusor muscle (HAO URI: purl.obolibrary.org/obo/HAO_0000811). Brothers (1975) and Brothers & Carpenter (1993) characterized several variations on prepectal morphology. Because the emphasis of the present work is on the fossil record, and as documentation of fossil prepecta is sparse, a simplified approach was here taken. When the prepectus was clearly visible, it was scored as *TRUE* for the present character, and where the prepectus was clearly free (*i.e.*, not immobile/fused to the mesopectus), it was scored as true for the dependent reductive character which follows.
224 (**Reductive**) *DC: Prepectus of alate individuals. DS: Discreteness.* • **Observational criterion:** Alate: prepectus externally visible and fused to mesopectus (*FALSE* = unfused OR indiscernible without dissection). **• NOTES:** Scored as *TRUE* when clearly present but immobile.
225 *DC: Anteroventral mesopectal lobe. DS: Expression.* • **Observational criterion:** Mesopectus, at posteroventral corner of pronotum, produced as triangular lobe just dorsal to procoxa, with lobe contacting pronotum along posterior ventralmost pronotal margin (*FALSE* = such a lobe absent). **• NOTES:** Observed uniquely in Evaniidae. Among Mesozoic fossils attributed to this family, only three species could be confidently evaluated for this trait: †*Cretevania montoyai* (*TRUE*), †*Lebevania azari* (*FALSE*), and †*Protoparevania lourothi* (*FALSE*).
226 *DC: Mesopectal “subalar pit” of alate individuals. DS: Expression.* • **Observational criterion:** Alate: mesopectus with deep pit subtending flange at base of wing insertion (*FALSE* = such a pit not developed OR pit shallow, not deep and distinct). **• NOTES:** A deep pit subtending the flange below the forewing insertion was only observed in Apoidea, and among apoids was generally *TRUE* except for *Heterogyna*, most Sphecidae, *Sphecius* (Bembicidae), and many Anthophila. This pit has been termed the "subalar fossa" by Bohart & Menke (1976).
227 *DC: Longitudinal mesopectal sulcus of alate individuals. DS: Expression.* • **Observational criterion:** Alate: longitudinal mesopectal sulcus present (*FALSE* = such a sulcus absent). **• NOTES:** Longitudinal sulci on the mesopectus are referred to by many names in the literature (*e.g.*, "epipleural suture" of Bohart & Stange 1965, “oblique mesopleural furrow” of Yoshimura & Fisher 2009, "anapleural sulcus" of Bolton 1994, "mesopleural line" of Buffington & Forshage 2014, *etc.*). In the present study, any longitudinally oriented or more-or-less longitudinal sulci were scored as *TRUE* for this character. That means that, despite the fact that presence of a longitudinal sulcus on the mesopectus is the canonical diagnostic feature of the Pompilidae, this state is scored as *TRUE* for the Formicidae and various other Aculeata. The homology-neutral approach for this particular character taken here means that the putative pompilid †*Bryopompilus* is scored as *TRUE* for this state, having a "transverse mesepisternal groove" (Engel & Grimaldi 2006), but so do the species of †*Burmusculus*, †Mutillidae sp. (CASENT0844578), and other taxa. The considerable variation in form of longitudinal mesopectal sulci may be captured in future study.
228 *DC: Anterior dorsoventral mesopectal sulcus of alate individuals. DS: Expression.* • **Observational criterion:** Alate: dorsoventral mesopectal sulcus present in anterior half of sclerite (*FALSE* = such a sulcus absent). **• NOTES:** This character is equivalent to the “episternal sulcus” of Bohart & Menke (1976). This sulcus was observed in "spheciform" apoids.
229 *DC: “Epicnemial carina” of alate individuals. DS: Expression.* • **Observational criterion:** Alate: mesopectus with dorsoventrally oriented carina situated approximately at the anterolateral margin of the sclerite; this carina delimits the procoxal contact surface of the mesopectus, *i.e.*, the “epicnemium” (*FALSE* = such a carina absent). **• NOTES:** Carinae meeting this description were observed in Evaniidae, Chrysididae, some Vespidae, *Tiphia* (Tiphiidae), some Apoidea, and many Formicidae. This carina is more-or-less equivalent to the epicnemial carina, *i.e.*, that carina which posteriorly delimits the mesopectal scrobe which accommodates the procoxa (see, *e.g.*, the definitions on HAO, URIs: prul.obolibrary.org/obo/HAO_0000294, purl.obolibrary.org/obo/HAO_0000292). In the context of the Apoidea, this carina is referred to as the "omaulus" (*e.g.*, Bohart & Menke 1976), and as scored, no distinction is here made between the "omaulus" and the "postspiracular carina". Because of the taxonomic scope of the present study, it should be noted that "epicnemium" as used by Bohart (*e.g.*, Kimsey & Bohart 1990) refers to the "prepectal carina" (*i.e.*, a carina of the posterior prepectal margin), differing from the sense used here (*i.e.*, as the scrobe, described above).
230 *DC: “Epicnemial carina” of alate individuals. DS: Location.* • **Observational criterion:** Alate: dorsoventral carina of mesopectus situated on anterior surface of anterior mesopectal scrobe, *i.e.*, (*FALSE* = such a carina absent or not situated in the groove of the mesopectus that receives the procoxae OR such a groove not developed). **• NOTES:** In other words, carina of epicnemium ("omaulus") located on anterior surface of broader scrobe. Observed in *Allocoelia*, *Hedychrum*, *Parnopes*, and *Argochrysis* (Chrysididae).
231 *DC: “Epicnemial carina” of alate individuals. DS: Location, shape.* • **Observational criterion:** Alate: Dorsoventral carina situated at anterolateral margin of mesopectus situated between anterior and lateral surfaces AND curving posterodorsally, delimiting dorsal potion of lateral mesopectal face (*FALSE* = such a carina absent OR not situated at margin of anterior and lateral surfaces of mesopectus OR linear, not curving posterodorsally). **• NOTES:** Observed in *Hedychrum*, *Parnopes,* and *Argochrysis* (Chrysididae).
232 *DC: “Epicnemial carina” of alate individuals. DS: Shape.* • **Observational criterion:** Alate: Dorsoventral carina delimiting dorsal potion of lateral mesopectal surfaces curved, rather than angular (*FALSE* = such a carina absent OR distinctly angular). **• NOTES:** Observed in *Parnopes* and *Argochrysis,* while the angular state was observed in *Hedychrum* (all Chrysididae).
233 *DC: Posterior dorsoventral carina of alate individuals. DS: Expression.* • **Observational criterion:** Alate: posterior half of mesopectus with dorsoventral carina which extends from anterior wing process ventrally to mesocoxal articulation (*FALSE* = such a carina absent). **• NOTES:** Observed uniquely in *Allocoelia* (Chrysididae).
234 *DC: Mesopectus of alate individuals. DS: Shape.* • **Observational criterion:** Alate: mesopectus posteriorly scrobiculate, *i.e.*, with a broad, shallow to deep groove or concavity along its dorsoventral length which receives the mesoleg when this leg upflexed (*FALSE* = mesopectus without such a concavity, being flat or convex posteriorly). **• NOTES:** A distinction is here drawn between scrobes which accommodate the mesoleg in primary contact with the mesopectus (the present state), and those which are in primary contact with the metapectus ("metapleural area scrobiform" below). Mesopectal mesoleg scrobes were observed in Evanioidea, most Chrysididae, Sclerogibbidae, and Scoliidae.
235 *DC: Mesopectal disc of alate individuals. DS: Shape.* • **Observational criterion:** Alate: mesopectus in cross-section along the frontal plane broadly V-shaped (*FALSE* = mesopectus without dorsoventrally oriented ridge, OR if with dorsoventrally oriented convexity, convexity broad). **• NOTES:** This is the diagnostic "campsomerine" state of 20th Century scoliid taxonomists (namely, Betrem and Bradley). Among sampled taxa, observed in *Campsomeris* and *Cathimeris.*
236 *DC: Mesopectal posterior margin of alate individuals. DS: Shape.* • **Observational criterion:** Alate: Mesopectus posterior margin, from wing insertion to coxa, broadly angled, with vertex of angle directed posteriorly (*FALSE* = posterior margin straight OR angled with vertex directed anteriorly). **• NOTES:** Observed uniquely in *Hyptia* (Evaniidae).
237 *DC: Mesepimeron of alate individuals. DS: Expression.* • **Observational criterion:** Alate: mesepimeron present as distinct sclerite posterior to "pleural suture", extending from wing base to base of mid coxa (*FALSE* = mesepimeron reduced, not present as such a strip). **• NOTES:** "Pleural suture" being the dorsoventrally oriented sulcus which extends from the mesocoxal insertion to the mesopleural wing process. Although being ancestral in the Pterygota, perhaps independently derived among Aculeata. Observed in *Orthogonalys* (Trigonalidae), Plumariidae, *Clystopsenella* (Scolebythidae), Sclerogibbidae, Rhopalosomatinae (Rhopalosomatidae), *Brachycistis* (Tiphiidae), and *Myrmosa* (Mutillidae *sensu lato*).

- *Metathorax*

238 *DC: Metanotum of alate individuals. DS: Size, medially.* • **Observational criterion:** Alate: metanotum strongly reduced such that mesoscutellum and propodeum contacting or nearly contacting (*FALSE* = metanotum broader). **• NOTES:** Observed in the bethylid subfamilies Bethylidae and Epyrinae (*e.g.*, Terayama 2003). Here scored as *TRUE* for *Lytopsenella* and *Epyris*.
239 *DC: Metanotum of alate individuals. DS: Shape.* • **Observational criterion:** Alate: metanotum with lateral region triangular in cross-section, being produced as angular to triangular processes adjacent to propodeum (*FALSE* = metanotum not produced, *i.e.*, rounded, without projection). **• NOTES:** Traditionally considered as a synapomorphy of Chrysidinae (Chrysididae; *e.g.*, character 12 of Bohart & Kimsey 1982, and character 22 of Kimsey & Bohart 1990, where this is described as part of the "transpleural carina"). Scored as *TRUE* for †Chrysidinae sp. (CASENT0844585, burmite) and †*Auricleptes*; for the latter, the metanotum in Figure 1D of Lucena & Melo (2018) is clearly produced laterally, forming a subtriangular lobe in dorsal view.
240 *DC: Medial metanotal bulge. DS: Degree of production.* • **Observational criterion:** Alate: metanotum with conspicuous (massive) dorsomedian process (*FALSE* = metanotum without such a bulge; medial metanotal bulge much smaller, rounded). **• NOTES:** Observed uniquely in *Bareogonalos* (Trigonalidae).
241 *DC: Metapleural area of alate individuals. DS: Shape.* • **Observational criterion:** Alate: metapleural area scrobiform, *i.e*., surface of metapleural area directed posteriorly due to bulging mesopectus, with capacity for receiving upfolded mesofemur (*FALSE* = this area flat or convex, not concave such that it can cradle the femora when these are upfolded). **• NOTES:** This state was observed in Bethylidae, Chrysididae, Embolemidae, male Sclerogibbidae, *Zethus* (Vespidae), *Tiphia* (Tiphiidae), †Pompiloidea sp. (CASENT0844580, burmite), Ampulicidae, some Sphecidae, some Crabronidae, *Bembix* (Bembicidae), Ammoplanidae, Philanthidae, some Pemphredonidae, and some Anthophila.
242 *DC: Upper metapleural area of alate individuals. DS: Relative size.* • **Observational criterion:** Alate: upper metapleural area posteriorly expanded (thus large) and continuous with often distinctly visible and often quite broad transverse metapostnotum (*FALSE* = this area not expanded AND metapostnotum small, largely unobservable). **• NOTES:** Observed in Pompilidae and †Burmusculidae, but not †*Bryopompilus*. The metapostnotum is highly variable in form, often being sunken, expanded laterally, or expanded medially.
243 *DC: Metapleural areas of alate individuals. DS: Shape, relative size.* • **Observational criterion:** Alate: upper metapleural area strongly reduced, being present as small, longitudinally aligned triangular to rhomboidal area dorsal to proportionally massive lower metapleural area (*FALSE* = upper area not of this specific size and shape, being larger, not in the form of a triangle or rhomboid AND more similar in size relative to lower area). **• NOTES:** As here described, the metapleural area of the metapectus is treated as being divided into "upper" and "lower" areas by the metapleural pit (*i.e.*, the "endophragmal pit", the pit corresponds internally to the metapleural apodeme; HAO URI: purl.obolibrary.org/obo/HAO_0000612). The metapleural area is rich with variation and should be the focus of a specific and detailed anatomical study across the Aculeata; only some clear states were separated in the present study for scoring, and most were not observable in fossil taxa. This particular pattern observed uniquely in Evaniidae and is most pronounced in *Evania*.
244 *DC: Lower metapleural area of alate individuals. DS: Size, shape.* • **Observational criterion:** Alate: lower metapleural area strongly reduced, being present as a dorsoventrally narrow but anteroposteriorly long triangle ventral to propodeum (lower metapleural area = that part of the metapectus ventral to the endophragmal pit and dorsal to the coxal foramen) (*FALSE* = lower area not of this specific form). **• NOTES:** Observed in Thynnidae. Scored as uncertain for *Brachycistis* as the female is without a margined metapleural area.
245 *DC: Metapleural areas of alate individuals. DS: Size, shape.* • **Observational criterion:** Alate: both upper and lower metapleural areas enlarged, being nearly as large together as propodeum in profile view (*FALSE* = these areas smaller, not as large as propodeum in lateral view). **• NOTES:** Observed in Scoliidae. This state conceals gross morphological adaptations of the endoskeleton which have yet to be depicted or described.
246 *DC: Metapleural areas of alate individuals. DS: Size, proportions.* • **Observational criterion:** Alate: lower metapleural area elongated, such that upper metapleural area present as dorsoventrally aligned subtriangular area situated very distant dorsally from mesocoxa with apex directed toward mesocoxal articulation (*FALSE* = lower area not elongated, upper area of variable shape and positioning). **• NOTES:** The description of this character is an effort to capture variation of the form observed in Sphecidae *sensu stricto.*
247 *DC: Metapleural areas of alate individuals. DS: Size, shape.* • **Observational criterion:** Alate: upper and lower metapleural areas with more-or-less parallel anterior and posterior margins, such that metapleural area is in the form of a dorsoventrally long strip (*FALSE* = these areas not aligned and strip-like; either one area enlarged relative to the other OR posterior margin concave or angled). **• NOTES:** Observed in *Euparagia* (Vespidae), Heterogynaidae, *Trypoxylon* (Crabronidae), and Bembicidae.
248 *DC: Upper metapleural carina of alate individuals. DS: Expression.* • **Observational criterion:** Alate: upper metapleural area with longitudinal carina in its dorsal fourth (*FALSE* = such a carina absent). **• NOTES:** Observed in Campsomerini (Scoliidae). In the scoliid literature, referred to as the "carina on transition of metapleuron" (*e.g.*, Betrem 1972).
249 *DC: Metapleural gland. DS: Expression.* • **Observational criterion:** Metapleural gland present in the posterolateral corner of the metapleural area (*FALSE* = this gland absent). **• NOTES:** The metapleural gland it the canonical autapomorphy of the Formicidae (*e.g.*, Wilson *et al*. 1967a,b; Bolton 1994, 2003). The gland is variably present in Camponotini and was confirmed as absent in †*Camelomecia* spp. (ANTWEB1038930, TONG112). It is present in †Haidomyrmecini, †Sphecomyrmini, and †Zigrasimeciini. This structure can be very challenging to evaluate, thus should be subjected to microanatomical study, particularly in fossils.
250 *DC: Metapleural gland orifice. DS: Shape, orientation.* • **Observational criterion:** Metapleural gland orifice dorsoventrally oriented and slit-shaped, not overhung by dorsal carina, orifice may be linear or curved (*FALSE* = metapleural gland absent OR orifice more longitudinally oriented AND/OR more open, elliptical AND/OR overhung by dorsal carina). **• NOTES:** Plausible synapomorphy of Ectatomminae *sensu lato* + Myrmicinae (see Bolton 2003, p. 52).

- *Thoracic venter*

251 *DC: Medioventral mesopectal lobes. DS: Expression.* • **Observational criterion:** Medial lobes of ventral mesopectal surface expanded posteriorly as large, distinct plates which at least partially overlap the coxae (*FALSE* = such lobes not developed OR if lobes developed anterad mesocoxae, these not overlapping/contacting the coxae). **• NOTES:** Expanded ventral mesopectal lobes are generally considered diagnostic of "Tiphiidae", but the traditional concept of the family is void due to paraphyly with the Chyphotidae (Pilgrim *et al*. 2008; Branstetter *et al*. 2017b). Among sampled Thynnoidea, these lobes were observed in Tiphiidae *sensu stricto* and Thynnidae except *Methoca* and are absent in Chyphotidae. Expanded ventral mesopectal lobes of this form were also observed in CASENT0844576, Vespoidea, †*Trichobaissodes*, and "†*Archisphex*" *incertus*. The "small" or "reduced" condition has been postulated to be ancestral in the Hymenoptera, *e.g.*, Johnson (1988). Considerable anatomical work is necessary to resolve the pattern of evolution of these structures. Particular reference should be made to the location of the medial mesocoxal articulations, and whether they are situated on the lobes or not.
252 *DC: Ventromedial mesopectal lobes. DS: Shape.* • **Observational criterion:** Medial lobes of ventral mesopectal surface expanded medially, forming large plates which are at an angle relative to one another (*FALSE* = such lobes not developed OR if lobes present then not angled relative to one another). **• NOTES:** This form of the ventromedial mesopectal lobes was only observed in Chrysididae.
253 *DC: Medial mesopectal process. DS: Expression.* • **Observational criterion:** Medial mesocoxal articulation situated on process, thus migrated distally on coxa with concomitant arm-like expansion of the ventral mesopectal surface (*FALSE* = such a process not developed). **• NOTES:** This is the state of "metapostepisternum elongate, articulating coxa well distant of its base" of Rasnitsyn (*e.g.*, Rasnitsyn 2002) defining his Evanioidea (Evanioidea, Trigonaloidea, Ceraphronoidea, Megalyridae, Stephanidae). Rephrased by Johnson (1988) as "mesal lobes … articulate on the surface of the coxa far from its rim".
254 *DC: Ventral mesopectal region. DS: Shape.* • **Observational criterion:** mesopectus ventrally scrobiculate, with scrobe accommodating mesocoxa when leg extended forward (*FALSE* = this region not concave, not capable of receiving leg when anteriorly pronated). **• NOTES:** Observed uniquely in Thynnoidea.
255 (**Additive**) *DC: Ventral mesopectal scrobe. DS: Shape.* • **Observational criterion:** Mesopectal scrobe deep, associated with relatively elongate mesocoxae (*FALSE* = such a scrobe not developed OR scrobe shallow). **• NOTES:** Observed uniquely in Tiphiidae *sensu stricto*.
256 *DC: Coxal socket of mesopectus. DS: Form.* • **Observational criterion:** Mesocoxae set in ventral thoracic socket, with sternal area nearly reaching ventralmost surface of coxa (*FALSE* = mesocoxae not set in distinct socket OR if coxae somewhat sunken into mesopectus then mesopectal concavity comparatively shallow and broad, not tightly fitting to coxal base). **• NOTES:** Socketed mesocoxae were observed in some Evaniidae, the Bethylidae, most Thynnoidea, Mutillidae *sensu stricto*, and Scolioidea. Although the Apoidea also trend toward socketed mesocoxae, the conformation is distinct, as the ventral mesopectal surface (bearing the medial coxal condyle) is produced ventrally (see, *e.g.*, Michener 1981).
257 *DC: Mesocoxae and mesopectus. DS: Size, shape.* • **Observational criterion:** Mesocoxae set in lateral thoracic socket and dorsoventrally elongate, such that distance between dorsal coxal margin and wing bases is shorter than the dorsoventral height of the coxa (*FALSE* = mesocoxae not socketed OR if socketed, dorsoventrally short). **• NOTES:** Synapomorphy of long-tongued bees (Megachilidae, Apidae) (*e.g.*, Michener 1981). Scored as *TRUE* for *Ceratina* as the coxa is clearly elongate dorsoventrally, although it is shorter than *Thyreus* or *Apis*, for example)
258 *DC: Mesocoxae. DS: Length.* • **Observational criterion:** Mesocoxae extremely foreshortened, being hardly or not at all produced away from the body (*FALSE* = these coxae longer, distinctly produced away from body). **• NOTES:** Foreshortened mesocoxae (see, *e.g.*, Michener 1981) were observed in Scoliidae, Crabronidae, *Sphecius* (Bembicidae), Ammoplanidae, and most Anthophila. Notably, the mesocoxae of Philanthidae are elongate, suggesting either reversal to long or multiple origins of short.
259 *DC: Mesopectal-metacoxal articulation. DS: Form.* • **Observational criterion:** Ventral surface of mesopectus, bearing medial mesocoxal articulation, strongly produced ventrally such that mesocoxal foramen directed laterally and hinge motion of mesocoxa is nearly vertical (*FALSE* = this articulation not of this specific form; medial mesocoxal articulation not produced ventrally OR if produced ventrally, then coxal hinge more horizontal). **• NOTES:** This is a feature of the Apoidea, observed in all sampled families except Ampulicidae and Heterogynaidae. Recorded as *TRUE* For the fossil taxon †*Cretosphecium.*
260 (**Additive**) *DC: Ventromedial mesopectal lobes. DS: Form.* • **Observational criterion:** Ventrally expanded ventral surface of mesopectus lateromedially expanded, forming lobe-like plates which do not widely overlap the mesocoxae (*FALSE* = such lobes not developed OR lobes wide-set, at an angle OR lobes widely concealing bases of coxae). **• NOTES:** This state observed in most spheciform Apoidea except for Ampulicidae, Heterogynaidae, and *Oxybelus*.
261 *DC: Ventral metapectal region. DS: Form.* • **Observational criterion:** Ventral surface of metapectus, bearing medial metacoxal articulation, strongly produced ventrally such that metacoxal foramen directed laterally and hinge of metacoxa is nearly vertical (*FALSE* = ventral metapectus not produced ventrally, metacoxal hinge not nearly vertical). **• NOTES:** Observed in all Apoidea, with some degree of variation. Whether this variation is informative or not was not evaluated.
262 *DC: Ventral metapectal region. DS: Shape.* • **Observational criterion:** Ventral surface of the metapectus flat, without wall-like structures between mesocoxa and ventrally directed metacoxae, thus clear wide space visible between the two pairs of coxae in profile view (*FALSE* = among Thynnoidea, meso- and metacoxae close-set, with such wall-like structures). **• NOTES:** Scored as synapomorphy of *Methoca* and *Aelurus* (Thynnidae). In other Thynnoidea, there is a distinct wall-like process which abuts the metacoxa when the metacoxa is extended forward.
263 *DC: Mesocoxal articulations. DS: Inter-foraminal distance.* • **Observational criterion:** Mesocoxae wide set (*FALSE* = close-set, *i.e.*, contiguous or nearly so in ventral view). **• NOTES:** Wide set mesocoxae occur sporadically among sampled taxa, including *Evania* (Evaniidae), most Chrysidoidea *sensu stricto*, most Thynnoidea, some Mutillidae, all Scolioidea, and the majority of sampled Apoidea.
264 *DC: Metacoxal articulations. DS: Inter-foraminal distance.* • **Observational criterion:** Metacoxae wide set (*FALSE* = close-set, *i.e.*, contiguous or nearly so in ventral view). **• NOTES:** Wide set metacoxae are much less frequent in occurrence than wide set mesocoxae. All Scoliidae have wide-set metacoxae, as well those formicines (Formicidae) which Bolton (2003) classified in his polyphyletic "lasiine tribe group". This condition was also observed in †*Kyromyrma neffi* and the undescribed formicine from Burmite. Despite being classified as an "aneuretine", †*Cananeuretus* was observed to have wide set metacoxae.
265 *DC: Metasternal region. DS: Form.* • **Observational criterion:** Lateromedially broad flat process of metasternal region present between (wideset) metacoxae (*FALSE* = ventral metapectal region lateromedially narrow, nor in the form of large process between wide-set coxae). **• NOTES:** This is a canonical autapomorphic condition of the Scoliidae. Most of the compression fossils attributed to the Scoliidae cannot be evaluated for this trait due to their preservation. However, the broad flat process was confirmed as absent in †*Cretaproscolia asiatica*, †*Cretoscolia laiyangica*, †*Cretoscolia rasnitsyni*, and †*Sinoproscolia yanghuwanziensis*; the first three were scored *FALSE* as the metacoxae are close-set, and the fourth (†*Sinoproscolia*) has a triangular metasternal area.
266 *DC: Ventral meso-metathoracic suture when metasternal region expanded. DS: Shape.* • **Observational criterion:** Lateromedially broad flat process of metasternal region with anterior margin linear (*FALSE* = anteriorly convex OR metasternal region not developed as broad process between metacoxae). **• NOTES:** Osten (1988) clearly illustrates the convex state in *Proscolia*; see also Fig. C266-0.
267 *DC: Posterior ventromedian sulcus of broad metasternal region. DS: Expression.* • **Observational criterion:** Lateromedially broad flat process of metasternal region between metacoxae with transverse sulcus just anterior to posterior margin (*FALSE* = such a sulcus not developed OR metasternal region not expanded lateromedially between metacoxae). **• NOTES:** Observed in *Scolia* and *Megascolia.*
268 *DC: Ventral metapectal region posterior expansion. DS: Expression, shape.* • **Observational criterion:** Metasternal area forming dorsoventrally oriented rectangular sclerite which separates the metacoxal cavities from the propodeal foramen (*FALSE* = this region not strap-shaped, not oriented relatively vertically; *i.e.*, this region lateromedially broader, more-or-less rectangular OR this region miniscule, spiniform OR this region more-or-less in the frontal plane). **• NOTES:** Described as the "propodeal sclerite" by Bohart & Menke (1976). Here observed in Sphecidae plus *Prionyx* (Crabronidae). This state is as scored here regards the conformation of the metasternal area, rather than closure of the metacoxal cavities, which was not evaluated although such closure is informative at least for generic diagnosis.

**-** *Legs*

269 *DC: Procoxotrochanteral articulation. DS: Form.* • **Observational criterion:** Disticoxal foramen of foreleg completely closed and ventrally sutured, with disticoxal foramen situated in sagittal plane (*i.e.*, socket directed laterally) (*FALSE* = foramen open OR incompletely sutured ventrally OR foramen at an oblique angle relative to sagittal plane).
**• NOTES:** This specific conformation is unique to the Formicidae and can be observed in stem taxa. A similar set of modifications has evolved in the Mutillidae *sensu lato* (Myrmosinae) and in Chrysididae (Loboscelidiinae), but the foramen in both is oblique relative to the sagittal plane, and in *Loboscelidia* is extremely torqued. Because of these anatomical distinctions and because these are clearly convergent, the three sets of states are here broken into three characters. In the initial stages of scoring data in the present matrix, characters of the coxal articulations were duplicated between alates (all taxa) and apterous individuals (reductive, with inapplicable when aptery absent). After reviewing the pattern, it is clear that winged and wingless individuals show the same sets of modifications with the sole exception of *Bradynobaenus*, where males have completely open articulations, and females have closed meso- and metacoxal articulations. This implies that closure of these articulations evolved independently between the Scoliidae and Bradynobaenidae. However, for the purpose of the present matrix, the female (apterous) states are scored for this terminal, and all apterous-specific characters of this character complex have been deleted.
270 *DC: Procoxotrochanteral articulation. DS: Form.* • **Observational criterion:** Disticoxal foramen of foreleg completely closed and incompletely sutured ventrally, with disticoxal foramen situated more-or-less in frontal (transverse) plane (*i.e*., socket directed posteriorly to posterolaterally) (*FALSE* = foramen open OR completely sutured ventrally OR foramen oriented in sagittal plane). **• NOTES:** This is the condition observed in some Myrmosinae (Mutillidae). The myrmecophile *Loboscelidia* is scored as *TRUE* for this condition.
271 *DC: Procoxotrochanteral articulation. DS: Form.* • **Observational criterion:** Disticoxal foramen of foreleg completely closed and incompletely sutured ventrally, with disticoxal foramen situated more-or-less in frontal (transverse) plane, with disticoxal foramen rotated (*i.e*., socket directed posteriorly to posterolaterally) (*FALSE* = foramen open OR completely sutured OR foramen in sagittal plane OR coxal apex not torqued). **• NOTES:** This is the condition observed in *Loboscelidia* (Chrysididae). Closure in this case is less likely caused by adaptation for pedal locomotion, but rather as a defense against ants as observed in other myrmecophiles.
272 *DC: Mesothoracicocoxal articulation. DS: Form.* • **Observational criterion:** Mesothoracic coxal cavity closed in lateral view, *i.e.*, membrane either completely concealed or cavity directed ventrally (*FALSE* = this cavity open OR directed laterally). **• NOTES:** Lateral mid coxal closure has evolved independently a number of times across the Apocrita. Among the Aculeata, this condition was observed in Plumariidae, Bethylidae, *Loboscelidia* (Chrysididae), *Sclerogibba* (Sclerogibbidae), *Olixon* (Rhopalosomatidae), Chyphotidae, most Thynnidae, Mutillidae, Scolioidea, Ampulicidae, and stem and crown Formicidae. Because of insufficient anatomical focus on non-aculeates, all taxa except the Trigonalidae and †Maimetshidae are scored as uncertain ("?").
273 *DC: Metathoracicocoxal articulation. DS: Form.* • **Observational criterion:** Metathoracic coxal cavity closed in lateral view, *i.e.*, membrane either completely concealed or cavity directed ventrally (*FALSE* = cavity open OR directed laterally). **• NOTES:** (1) This state is not to be confused with posterior metacoxal cavity closure, *i.e.*, expansion of the ventral metathoracic sclerite around the metacoxal cavity (as used in the myrmecological literature, *e.g.*, Bolton 2002), or expansion of the propodeal sclerite around those cavities (as used in the spheciform literature, *e.g.*, Bohart & Menke 1972).
(1) Hind coxal closure is somewhat less frequent than mid coxal closure across the examined taxa. Taxa with closed metacoxal cavities where the mesocoxal cavities are open include Tiphiidae *sensu stricto*, while closed mesocoxal cavities and open metacoxal cavities were observed in *Sclerogibba*, *Olixon*, *Typhoctes* (Chyphotidae), *Apterogyna*, *Ampulex*, and the apoid fossil CASENT0844594.
274 *DC: Procoxotrochanteral articulation. DS: Relative position.* • **Observational criterion:** Procoxa extended posterior to trochanteral articulation as distinct small to massive lobe (*FALSE* = procoxa not extended distad procoxotrochanteral articulation). **• NOTES:** This state is observed in *Plumarius* and extant Scolebythidae. Among the Mesozoic taxa which have been attributed to the Sclerogibbidae, only †*Libanobythus* clearly displays this state. Evans (1966) illustrated this as small lobe in the female of *Plumarius.* In the Scolebythidae, the lobe is so massive that it appears that the procoxotrochanteral articulation has migrated to the extreme base of the coxa.
275 (**Additive**) *DC: Procoxotrochanteral articulation. DS: Relative position.* • **Observational criterion:** Procoxa extremely extended posteriorly, such that procoxotrochanteral articulation appears to be dorsally migrated and is at base of coxa (*FALSE* = this articulation more distal on coxa). **• NOTES:** Observed uniquely in extant Scolebythidae; fossil “scolebythids” were difficult to evaluate for this specific state.
276 *DC: Anterolateral procoxal spine. DS: Expression.* • **Observational criterion:** Procoxa with a spine located on the anteroventral surface at about 1/3 the length of the coxa from basal to apical; this spine also fitting into a wide notch of the occipital carina (*FALSE* = such a spine not developed). **• NOTES:** Both this spine and the associated notch of the occipital carina are unique to *Santschiella* (Formicinae, Formicidae).
277 *DC: “Basicostal carina” of procoxa. DS: Expression.* • **Observational criterion:** Procoxa with posterolaterally-located dorsoventral carina which extends from lateral pleural articulation (or near to it) down almost to disticoxal foramen (*FALSE* = this carina not developed). **• NOTES:** From a general morphological perspective (Snodgrass 1935), this is probably the "basicostal suture" which divides the basicoxa from the disticoxa, and which, when directed ventrally along the posterolateral coxal surface, delimits posterior "meron" of the coxa. Among sampled taxa, this carina was observed in Aulacidae, Gasteruptiidae, Trigonalidae, some Chrysididae, some Vespidae, *Aelurus* (Thynnidae), Pompilidae, Sapygidae, Scoliidae, Crabronidae, Bembicidae, some Philanthidae, and most Anthophila. Among Formicidae, this was only observed in a single species of †*Zigrasimecia*. The "absent" or *FALSE* state of this character is a composite between “carina lacking entirely”, and “restriction of posterior surface delimited by carina to posterior base of procoxa”; further study is necessary to elucidate the alternative states and associated transformations, including possible associated skeletomuscular modifications.
278 *DC: Protrochanter. DS: Shape.* • **Observational criterion:** Protrochanter with digitate ventroapical elongation (*FALSE* = ventral protrochanteral articulation not produced distally relative to dorsal articulation, *i.e.*, apex of protrochanter appearing more-or-less transversely truncate apically). **• NOTES:** Observed uniquely in the vespid genera *Euparagia* and *Pseudomasaris.*
279 *DC: Protrochanter. DS: Proportions.* • **Observational criterion:** Protrochanter elongate, length ≥ 3 x maximum width (*FALSE* = protrochanter < 3 x maximum width). **• NOTES:** At the outset of the present study, it was expected that this would be a feature unique to a subset of "strange" taxa, such as the †*Hybristodryinus* (Dryinidae) from Burmite. However, as character evaluation progressed, it became apparent that this is a widespread character in the non-aculeate Apocrita, and occurs in Chrysidoidea (*Pristocera*), extant Dryinoidea (*Dryinus gulfensis*), and the Apoidea (Ammoplanidae, Pemphredonidae, Philanthidae), as well as the male of at least one species of Leptanillinae (Formicidae; not scored). A number of marginal cases were observed in the Anthophila, but this information was not captured by the index used here.
280 *DC: Forefemur of female. DS: Proportions.* • **Observational criterion:** Female forefemur grossly swollen along its entire length (*FALSE* = forefemur more than four times as long as wide; dorsal and ventral margins curving well before femorotibial articulation). **• NOTES:** Swollen female forefemora among extant taxa was observed in Chrysidoidea (Plumariidae, Scolebythidae, Bethylidae), Sclerogibbidae, *Olixon* (Rhopalosomatidae), *Aelurus* (Thynnidae), and a number of Apoidea (Crabronidae, Ammoplanidae, Pemphredonidae, Philanthidae, many Anthophila). Among extinct taxa, such femora were additionally observed in a cleptine (Chrysididae), †*Hybristodryinus* (Dryinidae), and †*Camelomecia* sp. (ANTWEB1038930). It is possible that this character may be split in future study based on refinement of the anatomical criteria, but this aggregate state is retained here as a marker of this conformation. Although not sampled in the present study, a number of arboreal Formicidae have inflated-muscular femora.
281 *DC: Protibia of female. DS: Shape.* • **Observational criterion:** Female: protibia swollen (*FALSE* = this segment at least four times as long as wide, ventral margin weakly convex to more-or-less linear). **• NOTES:** Occurring uniquely in Sclerogibbidae, according to Brothers & Carpenter 1993, where this state is the second clause of state 2 of character 111 in Appendix VI. Not observed in the female of †*Sclerogibbodes embioleia.*
282 *DC: Protibial spur. DS: Apex form.* • **Observational criterion:** Protibial calcar bifid (*FALSE* = this spur with a single apical point or rounded apically). **• NOTES:** Here scored as a coarse character aggregating any occurrence of two or more distinct tips of the calcar, including when the second tip is represented by the velum [lamina] of the pectinate portion of the spur. As constituted, this state was observed among Evanioidea, Trigonaloidea, †Falsiformicidae, *Parischnogaster* (Vespidae), most Pompiloidea *sensu lato* (except Mutillidae), *Apterogyna* (Bradynobaenidae), most Apoidea (except *Heterogyna*, Crabronidae, Ammoplanidae), †*Camelomecia*, and stem Formicidae (†*Zigrasimecia*, most †*Gerontoformica*, †*Sphecomyrma freyi*). In the stem Formicidae, the second tip is represented solely by microsetae, rather than the velum.
283 *DC: Protibial spur. DS: Proximodistal curvature.* • **Observational criterion:** Protibial calcar extremely strongly curved inwardly, semicircular in appearance (*FALSE* = this spur weakly curved). **• NOTES:** This condition was observed in Scolioidea and the vespid *Zethus caridei*. This character is modified from Brothers & Carpenter (1993), where it represents states 1 and 2 of character 117 in Appendix VI. An additional unique state of the Scolioidea, that of a hollowed posterior surface of the protibial calcar (states 1 and 2 of character 116 in Appendix VI of Brothers & Carpenter 1993) was excluded. Both states were observed in the present study for *Bradynobaenus* and *Apterogyna.*
284 *DC: Protarsal setae of female. DS: Shape.* • **Observational criterion:** Female protarsus with a row of fossorial setae, these setae long, flattened, and roughly scoopula-shaped (*FALSE* = setae of female protarsus short OR if long then these setae rounded, hair-shaped). **• NOTES:** This state was observed in *Parnopes* (Chrysididae), most Pompiloidea *sensu lato* (except *Sierolomorpha*, Sapygidae, and *Smicromyrmilla*), the Scolioidea, and numerous spheciform apoids. This character was included because it was observed in the enigmatic female †*Camelomecia fossor* (ANTWEB1038930). In Prentice (1998, character 27), this is called the "foretarsal rake" and was observed to be present in most Apoidea, except for Ampulicidae, Heterogynaidae, Sceliphrini, Pemphredonidae, and Apidae, among his other sampled taxa. These setae were not observed in stem or crown Formicidae, nor for the evanioid, trigonaloid, or ephialtitoid outgroups.
285 *DC: Protarsus of female. DS: Length, shape.* • **Observational criterion:** Female foreleg pretarsus modified for clasping, scythe-like, being elongate, curved, and more-or-less pointed apically (*FALSE* = this segment short, not scythe-like, shorter, of variable shape). **• NOTES:** This is a derived feature of the Dryinidae, where this modification is absent in females of the Aphelopinae and Biaphelopinae (*e.g.*, Goulet & Huber 1993).
286 *DC: Pretarsal claws. DS: Symmetry of development.* • **Observational criterion:** Propretarsus with one of the claws reduced or absent (*FALSE* = claws developed symmetrically). **• NOTES:** Observed in Dryinidae and *Tachysphex*. In the latter, this is associated with other modifications of the pretarsal claws. Complete reduction of pretarsal claws also occurs in some male *Temnothorax* (Formicidae, Myrmicinae).
287 (**Additive**) *DC: Propretarsal claw. DS: Symmetry of development.* • **Observational criterion:** Propretarsus with one of the claws completely absent (*FALSE* = both claws developed to some degree). **• NOTES:** Generally known to occur in Anteoninae and Transdryininae, and here specifically observed in *Deinodryinus* and *Burmanteon*.
288 *DC: Basicoxal articulations of mid- and hind legs. DS: Form.* • **Observational criterion:** Basimeso- and basimetacoxae dorsoventrally narrow, constricted as a distinctly small ball-like joint (*FALSE* = basicoxae anteroposteriorly narrow OR round OR not constricted). **• NOTES:** Observed in some Chrysidoidea, most Pompiloidea *sensu lato* (except Pompilidae, Sapygidae), Bradynobaenidae, Formicidae, and the TONG112 specimen of †*Camelomecia* figured by Barden & Grimaldi (2016).
289 *DC: Posterior thorax. DS: Intercoxal distance.* • **Observational criterion:** Mesocoxa situated distantly from metacoxa, the two being separated by ≥ 2 mesocoxal lengths (*FALSE* = these coxae closer, separated by < 2 mesocoxal lengths). **• NOTES:** This remarkable and unique state was observed in *Evania appendigaster* and has been illustrated for *Evaniscus* by Deans & Huben (2003).
290 *DC: Mesocoxal spine. DS: Expression.* • **Observational criterion:** Mesocoxa with distinct acute spine located on ventral surface (*FALSE* = such a spine not developed). **• NOTES:** This is a unique condition observed in †*Cretoecus spinicoxa* (see Budrys 1993).
291 *DC: Metacoxal process. DS: Expression.* • **Observational criterion:** Metacoxa with posterodorsal spiniform or lamellate process (*FALSE* = this coxa without such a process).
**• NOTES:** Metacoxal processes are diagnostic of the Myrmosinae (*e.g.*, Brothers & Lelej 2017), and were also observed to occur in *Cleptes* (Chrysididae), *Dolichurus* (Ampulicidae), *Lioponera* (Dorylinae, Formicidae), and various *Gnamptogenys* (Ectatomminae, Formicidae; not scored).
292 *DC: Metacoxae. DS: Proximodistal curvature.* • **Observational criterion:** Metacoxae with a single longitudinal axis, being elongate and directed ventrally (*i.e*., without lateral bend) (*FALSE* = these coxae bent or curved proximally such that the basicoxal articulation is at a distinct angle relative to the longitudinal axis of the disticoxa). **• NOTES:** This ancestral state of the Hymenoptera was observed among the Evanioidea, Trigonaloidea, most Chrysididae (except Bethylidae), most Vespoidea (except *Olixon*), few Pompiloidea *sensu lato* (Pompilidae, Sapygidae, Sierolomorphidae), Scoliidae, and most Apoidea (except *Dolichurus*). The derived state is an adaptation for pedal locomotion.
293 *DC: Mesotrochanters. DS: Proximodistal length.* • **Observational criterion:** Mesotrochanter elongate, length ≥ 3 x width (*FALSE* = these segments shorter, < 3 x width). **• NOTES:** This condition was observed widely in the Evanioidea and sporadically elsewhere, including some †Maimetshidae, at least one species of Bethylidae, †*Hybristodryinus magnificus* (Dryinidae), and *Ammoplanops* (Ammoplanidae). Characters, such as this one, appear in some sense to be "clocklike", and it is hoped that they contribute to parameterization of the Bayesian clock model formorphological data.
294 *DC: Mesotrochantella. DS: Degree of development.* • **Observational criterion:** Mesotrochantellus distinct and well-developed (*FALSE* = this portion of the mesofemur indistinct, poorly developed). **• NOTES:** This character has been used in a number of previous studies, where it is usually assumed to be ancestral among at least the Apocrita (*e.g.*, Brothers & Carpenter 1993: state 0 of character 196 of Appendix VI; Barden & Grimaldi 2016). The symplesiomorphy assumption may be true, but among the crown Formicidae, there appears to be a complex and functionally driven pattern of gain and loss. Among sampled taxa, the *TRUE* state was recorded for Evanioidea, Trigonaloidea, Plumariidae, Scolebythidae, Rhopalosomatinae (Rhopalosomatidae), some Vespidae, some Thynnoidea, *Pepsis* (Pompilidae), *Sapyga* (Sapygidae), most Mutillidae, some Scolioidea (*Proscolia*, *Apterogyna*), some Apoidea, and various Formicidae, including numerous crown taxa.
295 *DC: Meso- or metafemora of females. DS: Shape.* • **Observational criterion:** Female: meso- and/or metafemora swollen/inflated/fusiform (*FALSE* = one or both of these segments more-or-less linear, not subelliptical or appearing swollen/inflated/fusiform). **• NOTES:** According to Brothers & Carpenter (1993: state 1 of character 111 in Appendix VI), occurring in Chrysidoidea, Sclerogibbidae, and Dryinoidea. Here also observed in Ampulicidae, Anthophila, some Formicidae, and the stephanid-like †Symphytopterinae of the †Ephialtitoidea.
296 *DC: Meso- and metafemora of females. DS: Shape, distal process development.* • **Observational criterion:** Female: meso- and metafemora strongly compressed anteroposteriorly and expanded dorsoventrally, forming discs around a scrobe-like groove which accommodates the respective tibiae (*FALSE* = these segments not compressed and without distal disc-like processes which buttress a longitudinal groove). • **NOTES:** Observed in *Myzinum* and *Aelurus* (Thynnidae). In Tiphiidae *sensu stricto*, the tibia abuts the femur ventrally or is only slightly deflected onto the anterior surface (*Tiphia*) but is not accommodated along the entire anterior surface of the femur. This latter conformation is variably observed across the sampled taxa and was not characterized in time to evaluate consistently.
297 *DC: Denticles on posteroventral metafemoral margin of female. DS: Expression.* • **Observational criterion**: Female: this serration or dentition present (*FALSE* = absent). **• NOTES:** Observed in †Symphytopterinae (†Ephialtitoidea).
298 *DC: Metafemoral apex. DS: Shape.* • **Observational criterion:** Metafemora apically truncate (*FALSE* = this apex rounded apically, not flat and truncate). **• NOTES:** Apical truncation of the metafemora has been previously treated as a functional adaptation for digging in Apoidea. Bohart & Menke (1976) observed that the condition of truncation involved a complex of modifications, including "apicodorsal" and "apicoventral plates" of the metafemur, and a "basoposterior plate of the tibia". These details were not accounted for in the present matrix, however.
299 *DC: Metafemoral lamella. DS: Expression, form.* • **Observational criterion:** Metafemora with posteroapical lamella which is oriented parallel to longitudinal axis of segment (*FALSE* = posteroapical process of metafemur not developed as large lamella). **• NOTES:** Observed in Scoliidae and *Tiphia* (Tiphiidae). In both cases, this is almost certainly an adaptation for digging.
300 *DC: Tibial setation. DS: Form.* • **Observational criterion:** Meso- and/or metatibiae with strong, scattered spurs or chaetae, *i.e.*, tibial “traction setae” present (*FALSE* = pilosity of these segments not of this form, being narrow and hair-like in appearance). **• NOTES:** Tibial chaetae were observed in the female of *Plumarius* (Plumariidae), the female of *Pristocera* (Bethylidae), *Liosphex* (Rhopalosomatidae), *Zethus* (Vespidae), Thynnoidea, Pompilidae, *Fedtschenkia* (Sapygidae), *Myrmosa* (Mutillidae), most Scolioidea, various spheciform Apoidea (some Sphecidae, Crabronidae, Bembicidae, Ammoplanidae, Pemphredonidae, Philanthidae), *Eufriesea* (Apidae), and various Formicidae. Among the ants, the only sampled species with these traction setae is *Feroponera ferox* (Ponerinae), but such chaetae are also observed in the hypogaeic predators *Cryptopone*, *Centromyrmex*, *Buniapone*, *Promyopias*, *Feroponera*, and other Ponerinae. Brothers & Carpenter (1993) recorded this condition as state 1 of characters 119 and 122 in their Appendix VI. In most cases, these setae are associated with digging, but in some (*Liosphex*, *Zethus*, *Methoca*, *Eufriesea*) their function is less certain.
301 *DC: Metatibial pilosity. DS: Spatial distribution.* • **Observational criterion:** Metatibial traction setae/spurs arranged in at least two more-or-less even transverse rows which are buttressed by strong raised cuticular ridges (*FALSE* = pilosity of this form arranged haphazardly, not in even rows subtended by ridges). **• NOTES:** Observed in most Scoliinae, except Scoliini.
302 *DC: Tibiae. DS: Shape.* • **Observational criterion:** Meso- and metatibiae with longitudinal raised ridge or ridges (*FALSE* = tibiae terete, *i.e.*, round in cross-section, or nearly so). **• NOTES:** Observed in *Pepsis* (Pompilidae), and some Formicidae, *e.g.*, *Paraponera, Cephalotes,* and *Gigantiops*. Also scored as *TRUE* for the ephialtitoid †*Karataus daohugouensis*.
303 *DC: External surface of metatibial base. DS: Shape.* • **Observational criterion:** Metatibia truncate on its external face at its base where it articulates with the metafemur (*FALSE* = this area continuous with remainder of tibia, not differentiated in cuticular texture and pilosity). **• NOTES:** Michener (*e.g.*, Michener 1944) considered this condition to be expansion of the "basitibial plate". Among sampled taxa, occurs sporadically in Anthophila, and appears to have limited phylogenetic signal.
304 *DC: Tibiae. DS: Shape.* • **Observational criterion:** Meso- and metatibiae clavate, being grossly swollen (*FALSE* = these segments narrower, not enlarged apically like a club). **• NOTES:** Observed uniquely in most Thynnoidea except *Methoca* (Thynnidae) and Chyphotidae. Associated with digging.
305 *DC: Metatibia. DS: Shape.* • **Observational criterion:** Only metatibia clavate, being grossly swollen, like wide bell-bottom pants (*FALSE* = this segment thin, not massive and clavate). **• NOTES:** This unusual condition is known from *Gasteruption* and †*Zorophratra* [†Maimetshidae]. Here, the *TRUE* state was also applied to some †*Cretevania*, including †*C. alcalai*, †*C. exquisita*, and †*C. minor*.
306 *DC: Metatibia. DS: Shape.* • **Observational criterion:** Metatibia corbiculate, being expanded mediolaterally and with an apical, anterior glabrous region near the tarsal articulation (*FALSE* = anterior surface of this segment convex, without differentiated texture and setation). **• NOTES:** Unique to the corbiculate Apidae.
307 *DC: Apical tibial chaeta row. DS: Expression.* • **Observational criterion:** Meso- and metatibiae with an even to messy row of short, stout chaetae on their anteroapical margins (*FALSE* = tibia without chaetae OR if chaetae present, not restricted to such a row). **• NOTES:** This particular conformation observed in all sampled Vespidae except *Parischnogaster.* Similar but not identically arranged chaetae were observed sporadically, but not scored. Associated with these setae is apical elongation of the anterior surface of the tibiae as a lobe, which was also observed in *Sierolomorpha* and †Sierolomorphidae sp. (CASENT0844589, burmite), but was not characterized in time to evaluate across all sampled taxa.
308 *DC: “Rastellum”. DS: Expression.* • **Observational criterion:** Metatibial posteroapical margin with row of short stiff setae ("rastellum") (*FALSE* = rastellum absent). **• NOTES:** Unique to the corbiculate Apidae.
309 *DC: Posterior mesotibial spur. DS: Expression.* • **Observational criterion:** At least one mesotibial spur present (= posterior mesotibial spur present) (*FALSE* = no spurs developed at the ventral apex of the mesotibia). **• NOTES:** The state of completely absent mesotibial spurs occurs rarely but is observed among multiple clades of in the crown Formicidae, including *Anomalomyrma* (Leptanillinae), *Fulakora* (Amblyoponinae), various Formicinae (*e.g.*, *Cladomyrma*, *Gesomyrmex*, *Oecophylla*, *Aphomomyrmex*, *Lepisiota, Myrmoteras*), and various Myrmicinae (*e.g.*, *Stenamma*, *Wasmannia*, *Crematogaster*).
310 *DC: Anterior mesotibial spur. DS: Expression.* • **Observational criterion:** Two mesotibial spurs present (= anterior mesotibial spur present) (*FALSE* = only one spur developed at the ventral mesotibial apex OR no spurs developed). **• NOTES:** Loss of the second mid tibial spur has evolved independently in the Formicidae numerous times, but elsewhere is considered to be a state of considerable phylogenetic informativeness (*e.g.*, for Apoidea, Bohart & Menke 1976). In the context of the Apoidea, it is curious that the putative stem anthophilan †*Melittosphex* has a second metatibial spur (*e.g.*, Danforth & Poinar 2011), whereas among other apoids only Ampulicidae, Heterogynaidae, and Sphecidae have such a spur. During the scoring procedure of the present study, the bembicid apoid *Sphecius* appeared to have two mesotibial spurs, but upon closer inspection it was observed that the apparent anterior spur arose from the distal rim of the tibia, rather than the tibial-basitarsal membrane, as did the true tibial spur which it closely resembled. This may perhaps also be the case in †*Melittosphex*, but reevaluation will have to be done another time. Among other sampled taxa, loss of the second mesotibial spur was observed in some fossil Scolebythidae, †*Sclerogibba emboleia* (Sclerogibbidae), Dryinidae, CASENT0844576, Vespidae, Tiphiiformes, Scoliidae (diagnostic of Scoliinae), and variably among the Formicidae. Bohart & Menke (1976) list the males of *Didineis* and *Dinetus* as completely lacking tibial spurs.
311 *DC: Posterior metatibial spur. DS: Expression.* • **Observational criterion:** At least one metatibial spur present (= posterior metatibial spur present) (*FALSE* = no spurs developed at the ventral apex of the metatibia). **• NOTES:** As for the mesotibia, complete loss of metatibial spurs is highly unusual, but has been observed in *Apis* (Apidae), and various Formicidae.
312 *DC: Anterior metatibial spur. DS: Expression.* • **Observational criterion:** Two metatibial spurs present (= anterior mesotibial spur present) (*FALSE* = only spur developed at ventral metatibial apex OR no spurs developed). **• NOTES:** Loss of one of two metatibial spurs was observed in some Thynnidae (*Methoca*, *Aelurus*), *Apis* (as noted above), and many Formicidae.
313 *DC: Meso- and/or metatibial spurs. DS: Shape.* • **Observational criterion:** Tibial spurs with wispy, setiform distal extension (*FALSE* = tibial spurs shorter, without long hair-like distal extension OR such spurs not developed). **• NOTES:** This unique conformation was observed in the vespid genus *Parischnogaster.*
314 *DC: Meso- and/or metatibial spurs. DS: Shape, length.* • **Observational criterion:** Tibial spurs extremely elongate, with the posterior spur at least half length of basitarsus (= first or proximalmost tarsomere) (*FALSE* = spurs shorter, posterior spur length < 0.5 x that of the basitarsus). **• NOTES:** Relatively elongate tibial spurs were observed in Rhopalosomatidae, Thynnoidea (Tiphiidae, most Thynnidae), some Scoliinae (Scoliidae), and some Apoidea (*Trypoxylon, Sphecius*).
315 *DC: Meso- and/or metatibial spurs. DS: Shape.* • **Observational criterion:** Tibial spurs flattened and with pronounced large serration along the margins (*FALSE* = these spurs with more even margins, without triangular serration OR these spurs not developed). **• NOTES:** This state was uniquely observed in *Apterogyna* (Bradynobaenidae).
316 *DC: Posterior metatibial spur processes. DS: Expression.* • **Observational criterion:** This spur with one to several long triangular processes on the medial margin (*FALSE* = such processes not developed). **• NOTES:** One such process observed in *Pseudomasaris*, several in *Euparagia* (Vespidae).
317 *DC: Mesobasitarsus. DS: Shape.* • **Observational criterion:** This segment anteroposteriorly flattened (*FALSE* = this segment rounded, neither flattened nor widened). **• NOTES:** An expanded mesobasitarsus was observed among various Anthophila.
318 *DC: Metabasitarsus. DS: Shape.* • **Observational criterion:** This segment anteroposteriorly flattened (*FALSE* = this segment rounded, neither flattened nor widened). **• NOTES:** As for the mesobasitarsus, an expanded metabasitarsus is diagnostic of various Anthophila, and is here scored as *TRUE* for †*Melittosphex*, although this is a marginal case.
319 *DC: Metabasitarsal setation. DS: Distribution.* • **Observational criterion:** Flattened metabasitarsus with posterior surface covered by several rows of evenly aligned transverse setae (*FALSE* = setation of this segment not arranged in such rows). **• NOTES:** This was observed uniquely in *Apis* (Apidae).
320 *DC: Tarsomeres of female. DS: Shape.* • **Observational criterion:** Female: tarsomeres II–V of any leg broadly laminate, appearing swollen lateromedially (*FALSE* = form of these tarsomeres different, not laminate). **• NOTES:** This is, presumably, an adaptation for gripping, and was observed in some †Falsiformicidae and in female Rhopalosomatidae. †*Curiosivespa striata also* appears to have widened foretarsi. The tarsal form of Vespidae (with V-shaped tarsomeres) may be valuable to include in future study.
321 *DC: Tarsomere IV. DS: Symmetry.* • **Observational criterion:** This tarsomere extended apicomedially as asymmetrical lobate process (*FALSE* = this tarsomere symmetrical). **• NOTES:** Observed uniquely in *Olixon* (Rhopalosomatidae).
322 *DC: Tarsomeres I–IV. DS: Distal shape.* • **Observational criterion:** These tarsomeres apically expanded and conical, bell-like in appearance (*FALSE* = these tarsomeres not bell-like distally, form variable being V-shaped or otherwise). **• NOTES:** This condition, reminiscent of various Scarabaeoidea, was observed in *Brachycistis* (Tiphiidae), and most Scolioidea (with the exception of *Apterogyna*).
323 *DC: Tarsomeres IV, V. DS: Form, location of articulation.* • **Observational criterion:** This tarsomere wedge-shaped, with tarsomere V inserting near the base of IV, and with tarsomere IV expanded as triangular lobe covered on its ventral surface by dense pubescence (*FALSE* = tarsomere V not inserting near base of IV, tarsomere IV not triangular). **• NOTES:** Among sampled taxa, observed uniquely in *Ampulex* (Ampulicidae).
324 *DC: Dentition of pretarsal claw ventral margin. DS: Expression.* • **Observational criterion:** Pretarsal claw with at least one median or subapical tooth present (*FALSE* = claws without ventral dentition). **• NOTES:** A number of taxa have more than one tooth on the ventral margin of the pretarsal claw, including, for example, *Prionyx* (Sphecidae) and *Leptogenys* (Formicidae, Ponerinae). Among sampled taxa, absence of the tooth or teeth was observed in *Gasteruption* (Gasteruptiidae), some Chrysidinae, some Dryinidae, Embolemidae, some Vespidae, some Thynnoidea, Mutillidae, Scolioidea, most Apoidea (except Ampulicidae, Heterogynaidae, some Sphecidae), and most Formicidae. Whereas most stem Formicidae have dentate pretarsal claws, a minority of crown lineages have them, and these appear to be unusual reversals to the present state (*e.g.*, *Paraponera*, *Platythyrea*, *Harpegnathos*, myrmeciomorphs, Ectatomminae *sensu lato*).
325 *DC: Pretarsal claws. DS: Orientation.* • **Observational criterion:** Pretarsal claws very strongly reflexed, being directed proximally and in exact line with the apicalmost tarsomere, which itself is ventrally softened and covered with a plush layer of pubescence (*FALSE* = these claws not reflexed in relaxed position, rather being oriented ventrad to distad). **• NOTES:** This modification observed in *Oxybelis*, *Tachysphex*, and *Trypoxylon* (Crabronidae).
326 *DC: Foreleg arolium. DS: Size.* • **Observational criterion:** Pretarsus of foreleg with arolium crossly enlarged as large (sticky) pad spread by wide, paired, ellipsoid sclerites (*FALSE* = these arolia smaller, less conspicuous to inconspicuous or absent). **• NOTES:** This state is distinct of that of the following by the obvious presence of the two ellipsoid sclerites. This condition was observed in †Falsiformicidae, Dryinidae, and Sclerogibbidae. The arolia are otherwise enlarged in some crown Formicidae (*e.g.*, male *Camponotus*), and in various Apoidea. Scored as a "?" for the obligately arboreal formicine genera *Myrmelachista* and *Cladomyrma* as their arolia are enlarged relative to other Formicidae, but the two sclerites were not observable.
327 *DC: Mid, hind leg arolia. DS: Size.* • **Observational criterion:** Pretarsus of meso- and metalegs with arolium crossly enlarged as large (sticky) pad spread by wide, paired, ellipsoid sclerites (*FALSE* = these arolia smaller, less conspicuous to inconspicuous or absent). **• NOTES:** This character is essentially additive relative to the prior, and was observed in †Falsiformicidae and Dryinidae, but not in Sclerogibbidae.
328 *DC: Unguitractor plate. DS: Size.* • **Observational criterion:** Arolia enlarged, with an expanded unguitractor plate (*FALSE* = arolia small, unguitractor plate not expanded). **• NOTES:** Observed in †*Burmusculus* (Pompiloidea), the male of *Bradynobaenus* (Bradynobaenidae), Sphecidae, most Crabronidae, and *Sphecius* (Bembicidae).
329 *DC: Plantar lobes of tarsomeres I–IV. DS: Expression.* • **Observational criterion:** These tarsomeres with plantar lobes (apicomedian ventral lobate processes) (*FALSE* = such lobes absent). **• NOTES:** The scoring here may be considered aggregate, as it includes "plantar lobes" and "pulvilli" *sensu* Schulmeister (2003), Vilhelmsen *et al*. (2010), and is observed sporadically among the sampled taxa, including Trigonalidae, *Aelurus* (Thynnidae), *Pepsis* (Pompilidae), Bradynobaenidae, various Apoidea, stem Formicidae, †*Camelomecia*. There is evidence that the “plantar lobes” of Aculeata are homologous with similarly labeled structures in the symphytan grade (Beutel *et al*. 2020).
330 *DC: Plantar lobes. DS: Size.* • **Observational criterion:** Plantar lobes of tarsomeres massively enlarged, forming broad scale-like structures (*FALSE* = these lobes small and less conspicuous, OR absent). **• NOTES:** Observed in the male of *Bradynobaenus*, and also in *Pepsis formosa.* Such enlarged "plantar lobes" are recorded as a diagnostic trait of the Pompilidae by Gauld & Bolton (1988).
331 *DC: Metafemoral setation. DS: Setal form.* • **Observational criterion:** This segment with dense brush of branched setae, *i.e.*, metafemoral scopa present (*FALSE* = setation of this segment not plumose, rather being hair-like). **• NOTES:** Observed in Anthophila; scopae on hind leg are extremely dense on *Caupolicana.*
332 *DC: Metatibial setation. DS: Setal form.* • **Observational criterion:** This segment covered with dense brush of branched setae, *i.e.*, metatibial scopa present (*FALSE* = setation of this segment not plumose, otherwise variable in form, including hair-like to chaeta-like). **• NOTES:** Observed in Anthophila.
333 *DC: Metabasitarsal setation. DS: Setal form.* • **Observational criterion:** This segment with dense brush of branched setae, *i.e.*, metabasitarsal scopa present (*FALSE* = setation of this segment not plumose, rather being hair-like). **• NOTES:** Observed in Anthophila.

- *Propodeum*

334 *DC: Propodeum. DS: Size, shape.* • **Observational criterion:** Propodeum poorly developed, *i.e.*, short anteroposteriorly/dorsoventrally, without distinct posterior face or posterior declivity, AND without bridge of propodeal sclerite ventral to propodeal foramen, *i.e.*, propodeal foramen must be open ventrally, not enclosed by propodeal sclerite (*FALSE* = this structure well-developed, *i.e.*, with a distinct long surface, often with distinct dorsal and ventral faces AND/OR propodeal foramen enclosed ventrally by an arc of sclerotized cuticle). **• NOTES:** This is the ancestral state for the Hymenoptera. Among sampled taxa, this state was only observed in †Ephialtitoidea and the praeaulacid †*Eonevania*.
335 *DC: Paired propodeal lines. DS: Development, orientation.* • **Observational criterion:** Propodeum with paired lines which are oriented more-or-less longitudinally, and which may or may not contact each other (*FALSE* = such lines absent). **• NOTES:** Among extant taxa, paired longitudinal lines on the propodeum are observed in Apoidea and Scoliidae. These lines in the Apoidea are associated with expansion of the metapostnotum (see Brothers 1976 for characterization), but the latter remains in need of anatomical explanation. Among extinct taxa, paired lines are observed in †*Archaeoscolia*, †*Protoscolia*, †*Cretobestiola tenuipes*, †*Cretospilomena*, and †*Psolimena*. Although the lines of Scoliidae do not meet apically, those lines do meet in †*Protoscolia*, †*Cretobestiola*, and the two pemphredonid genera; because it is unclear whether the U-shaped connection of these lines in †*Protoscolia* or †*Cretobestiola* is homologous, and because the interpretation of homology contradicts their placement in the Scolioidea and Apoidea, respectively, presence of lines is scored as a single homoplastic state, and the forms of the lines themselves are scored as three alternative characters: not meeting, meeting but V-shaped, and meeting but U-shaped.
336 (**Additive**) *DC: Paired propodeal lines. DS: Orientation.* • **Observational criterion:** These lines extending to propodeal foramen without meeting apically, thus propodeum divided into three distinct regions (*FALSE* = lines with a different conformation OR absent). **• NOTES:** This state describes the state observed in Scoliidae. Among the sampled fossil taxa, observed for †*Archaeoscolia hispanica*, †*Archaeoscolia senilis*, and †*Cretoscolia brasiliensis*.
337 (**Additive**) *DC: Paired propodeal lines. DS: Orientation, curvature.* • **Observational criterion:** These lines joining posteroapically via a curve rather than angle, thus middle portion of propodeum U-shaped and lateral portions connecting posteriorly behind U (*FALSE* = lines of a different conformation OR absent). **• NOTES:** Observed in †*Protoscolia* and †*Cretobestiola*; appears to be *TRUE* for †*Pompilopterus corpus* but scored as "?" due to ambiguity.
338 (**Additive**) *DC: Paired propodeal lines. DS: Orientation, curvature.* • **Observational criterion:** These lines, if present, joining posteroapically via an angle rather than curve, thus middle portion of propodeum V-shaped forming "propodeal triangle" (*FALSE* = lines of a different conformation or absent). **• NOTES:** Expansion of the metapostnotum in this manner is a unique feature of Apoidea (*e.g.*, Brothers 1976 as state 1 of character 65 in Appendix VI of Brothers & Carpenter 1993). Kawada *et al*. (2015) describe an elongate metapostnotum of the Bethylidae which forms a rectangular extension on the propodeum, but this character was not added to the present study because of difficulty of evaluation for other "chrysidoid" taxa, given that similar anatomical treatments are not available for them. Although †*Pompilopterus corpus* has been placed in the Apoidea, this does not appear to be the state of the postnotum labeled by Rasnitsyn *et al*. (1998), therefore this taxon scored as "?". The dorsal propodeal surface of †*Burmasphex* is illustrated in Figure 1 of Melo & Rosa (2018), wherein it can be seen that the dorsal surface is evenly smooth and does not have longitudinal lines. In the apoid genus *Sphex* (Sphecidae), the lines were indistinct, but the apex of the "V" was observable as a distinct pit.
339 *DC: Median propodeal line. DS: Expression, length.* • **Observational criterion:** Propodeum with median longitudinal sulcus that extends from metanotum to propodeal foramen (*FALSE* = such a sulcus not developed OR if developed then incompletely spanning propodeum). **• NOTES:** Observed in Vespidae and may also be a form of "postnotal expansion". In some Rhopalosomatidae, two sulci may be seen which extend dorsally from the propodeal foramen, and which are associated with the paired propodeal condyles that articulate with the first metasomal segment.
340 *DC: Dorsomedian propodeal process. DS: Expression.* • **Observational criterion:** Propodeum with ornate dorsomedian process (*FALSE* = such a process not developed). **• NOTES:** Observed in *Parnopes* (Chrysididae) and *Oxybelus* (Crabronidae). Described by Bohart & Menke (1976) as the "propodeal mucro".
341 *DC: Elevated dorsomedian propodeal triangle. DS: Expression.* • **Observational criterion:** Dorsal face of propodeum with medial portion raised and posteriorly elongate relative to the lateral portions, thus the propodeum with a distinct but blunt dorsomedial triangle (*FALSE* = propodeum without differentiated, raised, triangular zone). **• NOTES:** Observed in *Campsomeris* and *Cathimeris* (Scoliidae).
342 *DC: Propodeum. DS: Form.* • **Observational criterion:** Propodeum in form of anteroposteriorly elongate rectangular box (such that length is subequal to or greater than dorsoventral height), with or without distinct dorsolateral and posterolateral margins (*FALSE* = propodeum short, dorsal surface length less than propodeal height). **• NOTES:** Observed in Bethylidae, some Dryinidae, some Embolemidae, *Olixon* (Rhopalosomatidae), some Thynnidae (*Methoca*, *Aelurus*), and some Apoidea (Ampulicidae, Heterogynaidae, Sphecidae). *Methoca* has an elongate propodeum, but the dorsolateral margins are evenly rounded.
343 *DC: Propodeum. DS: Form.* • **Observational criterion:** Propodeum without a dorsal face in profile view (*FALSE* = propodeum with dorsal surface in profile view). **• NOTES:** Observed sporadically among the sampled taxa, including Chrysidinae (Chrysididae), many Vespidae, *Smicromyrmilla* (Mutillidae), *Bradynobaenus* (Bradynobaenidae), and various Apoidea.
344 (**Additive**) *DC: Propodeum. DS: Form.* • **Observational criterion:** Propodeum "wedge-shaped" in profile view, vertical from metanotum to propodeal articulation, with anterior apex of wedge at about midheight of propodeum, *i.e.*, with anterodorsal and anteroventral margins subequal (*FALSE* = shape variable, being boxy AND/OR with distinct dorsal face AND/OR ventral margin distinctly longer than dorsal margin AND/OR posterior propodeal face not vertically aligned ventrad metanotum). **• NOTES:** Synapomorphy of Chrysidinae (Chrysididae) (character 24 of Kimsey & Bohart 1990).
345 *DC: Propodeal carinae. DS: Expression, location.* • **Observational criterion:** Propodeum with dorsolateral and posterolateral margins carinate, *i.e.*, margination present (*FALSE* = such carination not developed). **• NOTES:** The propodeum is rich in sculptural features. Due to time limitation, many sculptural forms were excluded from this study. However, distinct carination of the propodeum was straightforward to score, being observed in some Bethylidae, †Chrysobythidae, †*Protamisega* (Chrysididae), †*Aculeata* sp. (CASENT0844587, burmite) (placement to be determined), some †Falsiformicidae, Embolemidae, *Sclerogibba* (Sclerogibbidae), some Ampulicidae (*Ampulex*, †*Cretampulex*), *Oxybelus* (Crabronidae), †*Camelomecia*.
346 *DC: Dorsolateral propodeal carina. DS: Shape.* • **Observational criterion:** This carina extending posteriorly from juncture between upper and lower metapleural areas and in the form of a broad, downturned curve (*FALSE* = such a carina absent OR carina linear to weakly convex, not broadly curving down to the metathoracicopropodeal corner). **• NOTES:** Observed in Scoliidae, where in *Megascolia* the carina is directed more-or-less dorsally ("carina lateralis" of, *e.g.*, Day *et al*. 1981).
347 (**Additive**) *DC: Broadly curved dorsolateral propodeal carina. DS: Relative length.* • **Observational criterion:** Propodeal lateral longitudinal carina shorter than sulcus dividing dorsal and ventral metapleural areas (*FALSE* = carina as long as or longer than sulcus, OR sulcus absent). **• NOTES:** Observed in *Cathimeris*, which has a short lateral longitudinal carina; other Scoliinae with the carina with longer carina.
348 *DC: Dorsal posterolateral propodeal processes. DS: Expression.* • **Observational criterion:** Propodeum armed, *i.e.*, a propodeal spine, tooth, denticle, angle, or lobe present (*FALSE* = such processes absent). **• NOTES:** Observed variably across the Aculeata. Among sampled extant taxa, observed for Chrysidinae (Chrysididae), *Olixon* (Rhopalosomatidae), *Pseudomasaris* (Vespidae), Ampulicidae, and various Formicidae. Among extinct terminals, *TRUE* for †*Xenodellitha* (†Othniodellithidae), †Chrysobythidae, some Chrysididae (including †*Auricleptes*), †Falsiformicidae, †*Burmasphex*, †*Hybristodryinus resinicolus* (Dryinidae), “vespoid” undescribed taxon (CASENT0844576), some Vespidae, Ampulicidae, and various Pemphredonidae. Notably, propodeal armature is absent in all sampled Mesozoic taxa except for †*Cretomyrma*, which bears a single median tooth, reminiscent of *Dorymyrmex* (Dolichoderinae).
349 *DC: Propodeal spiracle. DS: Location.* • **Observational criterion:** Spiracle situated on lateral propodeal surface, sometimes quite distant posteriorly from metapleuron (*FALSE* = situated on dorsal or dorsolateral propodeal surface, close to or even contiguous with metanotum or metapostnotum). **• NOTES:** A laterally situated propodeal spiracle occurs infrequently among Aculeata and is a Putative synapomorphy of the Formicidae (Bolton 2003). Here, such positioning was observed for Sclerogibbidae, some Chrysididae, *Apterogyna* (Bradynobaenidae), *Plenoculus* (Crabronidae), *Bembix* (Bembicidae), *Apis* (Apidae), and all Formicidae. Spiracle location was difficult to evaluate for the majority of fossils but was confirmed as high and anteriorly situated in †*Camelomecia* (ANTWEB1038930) and two unnamed †*Camelomecia* males.
350 *DC: Propodeal spiracle. DS: Shape.* • **Observational criterion:** Spiracle slit-shaped (*FALSE* = elliptical to circular). **• NOTES:** The propodeal spiracle was observed to be slit-shaped in virtually all sampled taxa except *Clystopsenella* (Scolebythidae), and *Ampulicomorpha* (Embolemidae), although some *Ampulicomorpha* were observed to have slit-shaped spiracles. Within the Formicidae, slit-shaped spiracles were observed in †Haidomyrmecini, †*Zigrasimecia*, †*Gerontoformica*, †*Sphecomyrma freyi*, †*Brownimecia*, *Paraponera*, some ponerines, *Myrmecia*, and some formicines. †*Zigrasimecia* CNU09193, to be recognized as a distinct genus in forthcoming work (“†*Protozigrasimecia chauli*” in Cao *et al*. 2020b), was scored as uncertain but upon reevaluation was determined to have a slit-shaped spiracle.
351 *DC: Propodeal spiracle anterior hood-like projection. DS: Expression.* • **Observational criterion:** Such a projection present (*FALSE* = such a projection absent). **• NOTES:** Observed sporadically among sampled taxa, including *Evania* (Evaniidae), most Trigonalidae, most Mutillidae, Philanthidae, *Thyreus* (Apidae), and some †*Gerontoformica*.
352 *DC: Propodeal spiracle basal process. DS: Expression.* • **Observational criterion:** Propodeal spiracle situated on distinct tubercle (*FALSE* = such a tubercle not developed). **• NOTES:** Observed sporadically in crown Formicidae, but among sampled taxa only *Lepisiota* (Formicinae) has such produced spiracles.
353 *DC: Posteroventral propodeal margin. DS: Length between coxa and metasomal articulation.* • **Observational criterion:** This margin extended posteriorly to posterodorsally, thus separating the propodeal foramen from the metacoxa by about the length of the metacoxa (*FALSE* = this margin not extended, thus metacoxa and metasomal articulation separated by less than one metacoxal length). **• NOTES:** Observed in Rhopalosomatidae, Vespidae, and †Sierolomorphidae sp. (CASENT0844589, burmite).
354 *DC: Ventrolateral propodeal lamina. DS: Expression.* • **Observational criterion:** Propodeum with broad translucent to opaque laminar process just dorsal to the metacoxal articulation which extends posteriorly to shield metasomal insertion in lateral view (*FALSE* = such a lamina not developed). **• NOTES:** Observed in Vespoidea *sensu stricto*, where this has been termed the "valvula of the propodeum" (*e.g.*, Bohart & Stange, 1965). This region of the body deserves very careful, explicit study; many characters here remain undefined in the present study. In particular, a similar-appearing laminate process is observed in *Pluto* and the Anthophila, but this conceals the base of the coxa, although in the spheciform the process is more-posteriorly situated. The propodeum of *Methoca* was observed to have a laminar process which shields the metasomal articulation, so this was scored as *TRUE*.
355 *DC: Propodeal articulation carination. DS: Length, shape.* • **Observational criterion:** Propodeum with distinct but short tubular extension surrounding metasomal articulation (*FALSE* = propodeal articulation carination not extended in a tube-like form). **• NOTES:** This tubular extension was observed in some Sphecidae, *Trypoxylon* (Crabronidae), and Philanthidae.
356 *DC: “Propodeal lobes”. DS: Expression.* • **Observational criterion:** Carina encircling propodeal foramen produced laterally as lobes, with lobes abutting or nearly abutting anterior portion of metasoma (*FALSE* = such lobate processes absent). **• NOTES:** These are the "propodeal lobes" of the myrmecological literature. Bolton (2003, p. 29) considered the lobes to be absent in *Apomyrma*, *Leptanilla*, and Formicinae, with the exception of *Oecophylla*; he observed lobes to be variably present in Dorylinae. Here they were scored as absent for the dolichoderomorphs (Dolichoderinae, Aneuretinae) and most Formicinae); notably, they were present for *Opamyrma*. Propodeal lobes were scored as "?" for the majority of Mesozoic Formicidae.

**-** *Metasoma*

357 *DC: Abdominal segment II (metasomal I). DS: Shape.* • **Observational criterion:** This segment with anterior base broad and sides of segment subparallel in dorsal view (*FALSE* = this segment narrowed anteriorly to some degree, not necessarily forming “wasp waist”). **• NOTES:** This is a fundamental ancestral state of the Hymenoptera and was included specifically to separate †Ephialtitoidea from the "Evaniomorpha" and "Vespomorpha" (*sensu* Rasnitsyn). While scoring, it became clear that the †Symphytopterinae have narrow articulations, and that the praeaulacid †*Eonevania* has a broad articulation. Otherwise, all other sampled taxa have narrow articulations.
358 *DC: Abdominal segment II (metasomal I). DS: Shape.* • **Observational criterion:** This segment dorsoventrally constricted anteriorly (*FALSE* = this segment dorsoventrally broad, *i.e.*, the unmodified plesiomorphic state of the Hymenoptera). **• NOTES:** This is a derived state of the Apocrita, but whether it is indeed a true single origin synapomorphy (*i.e.*, autapomorphy) of the group is unclear when considering the fossil record. For example, Rasnitsyn & Zhang (2010) place †Ephialtitidae as sister to the Stephanidae, which implies three origins of the "wasp waist": Once in Evanioidea, once in the Stephanoidea (Megalyridae + Stephanidae), and once in the remaining Apocrita, at least by their scheme. Although the sampling of non-aculeate non-evanioid Apocrita is too slim to resolve this question, the morphological transformation syndrome is further split by the recognition that, although †Symphytopterinae have lateromedially broad waists, these are also dorsoventrally narrow.
359 *DC: Metasomal articulation with propodeum. DS: Location.* • **Observational criterion:** This articulation separated dorsally away from metacoxae (*FALSE* = this articulation situated close to metacoxae). **• NOTES:** This is the evanioid condition and was observed in most taxa placed in the Evanioidea, except for some †Praeaulacidae, including †*Eosaulacus*, †*Sinaulacogastrinus*, †*Sinevania*, and †*Eonevania*.
360 *DC: Subarticular propodeal bridge. DS: Expression.* • **Observational criterion:** Propodeum forming completely fused and anteroposteriorly/dorsoventrally elongate and sclerotized bridge beneath metasomal articulation (*FALSE* = such a sclerotized bridge not developed). **• NOTES:** This is another characteristic feature of the Evanioidea. Among sampled evanioids, this was only found to be *FALSE* in the praeaulacid genus †*Eosaulacus*; the condition of a number of other extinct evanioids is uncertain.
361 *DC: Metasomal articulation with propodeum. DS: Location.* • **Observational criterion:** This articulation situated above midheight of propodeum (*FALSE* = this articulation situated below propodeal midheight). **• NOTES:** This is another feature of the Evanioidea which is variable in the †Praeaulacidae, being *FALSE* for †*Eosaulacus*,†*Evanigaster*, and †*Praeaulacus orientalis.*
362 *DC: Propodeal articulation projection. DS: Expression.* • **Observational criterion:** Metasomal articulation situated on conical projection of propodeum, this projection not formed by periarticular carinae (*FALSE* = such a projection not developed OR if articulation separated from body of propodeum, this formed by the propodeal carinae, slightly above metacoxae). **• NOTES:** This state is observed in Aulacidae and most of the Mesozoic taxa attributed to this family except for †*Hyptiogastrites*.
363 *DC: Propodeal articulation projection. DS: Distance from metanotum.* • **Observational criterion**: Coniform propodeal projection contacting metanotum (*FALSE* = cone not contacting metanotum OR cone absent). **• NOTES:** Observed in some Mesozoic Aulacidae.
364 *DC: Dorsal propodeal surface. DS: Form, curvature.* • **Observational criterion:** Propodeal surface between elevated propodeal foramen and metanotum convex (*FALSE* = concave to linear OR propodeal foramen not raised). **• NOTES:** This condition is uniquely observed in †*Sinuevania mirabilis* and crown Evaniidae. Because this condition is unique, even among Evanioidea, the "inapplicable" state is aggregated with *FALSE*.
365 *DC: Dorsal propodeal surface. DS: Length.* • **Observational criterion:** Metasomal articulation contacting metanotum (*FALSE* = metasomal articulation not contacting metanotum, OR articulation raised on cone). **• NOTES:** Observed in †Praeaulacidae (†*Sinaulacogastrinus*, †*Sinevania*), some putative Aulacidae (†*Hyptiogastrites)*, and Gasteruptiidae. In †*Archaeofoenus* the metasoma is inserted on a cone and it is the cone which abuts the metanotum, rather than abdominal segment II.
366 *DC: Metasoma. DS: Form, length.* • **Observational criterion:** This tagma long, thin, and tubular, with segments only gradually increasing in dorsoventral width toward abdominal apex (*FALSE* = this tagma shorter, stockier, not clavate in overall appearance). **• NOTES:** Observed in †*Hyptiogastrites* (Aulacidae), and most Gasteruptiidae except for †*Kotujisca*. Also observed in the putative trigonalid †*Albiogonalys*.
367 *DC: Ventral metasomal pilosity. DS: Form.* • **Observational criterion:** Metasoma with dense tufts of plumose setae ("scopa") on ventral surface (*FALSE* = this pilosity not plumose). **• NOTES:** Observed in *Caupolicana*, *Dieunomia*, and *Megachila*. Generally considered diagnostic of the Megachilidae.
368 *DC: Metasoma posterior to segment II. DS: Form.* • **Observational criterion:** This portion of the metasoma very short and lateromedially compressed, in form of oval to tear-drop in lateral view (*FALSE* = metasoma not hatchet-shaped, *i.e.*, posterior portion of metasoma longer AND/OR not compressed side-to-side AND/OR not oval in lateral view). **• NOTES:** Defining feature of Evaniidae (character 111, state 1 in Ronquist *et al*. 1999). Also observed in †*Othniodellitha* (†*Othniodellithidae*). (The condition for †*Xenodellitha* is unknown due to artefactual loss of the metasoma.)
369 *DC: Abdominal segment II tergosternal articulation. DS: Fusion.* • **Observational criterion:** This segment with tergosternal fusion, whether partial or complete (*FALSE* = AII tergum and sternum freely mobile). **• NOTES:** Tergosternal fusion has evolved independently a number of times among the sampled taxa. The state is diagnostic of the Evaniidae, and elsewhere is observed in some Vespidae, some Thynnoidea, *Bradynobaenus* (Bradynobaenidae), *Pluto* (Pemphredonidae), and variably in the Formicidae. Crucially for the ants, Bolton in Fisher & Bolton (2016, p. 483) tabulated occurrence of tergosternal fusion at approximately subfamily level. Among the crown Formicidae, fusion is ABSENT in *Adetomyrma* (Amblyoponinae, a clear reversal), *Probolomyrmex* (Proceratiinae, a clear reversal), Ponerinae, Dorylinae, Myrmeciinae, Pseudomyrmecinae, and Ectatomminae. Unfortunately, it was not possible to evaluate this condition for the stem Formicidae.
370 *DC: Abdominal segment II (“petiole”). DS: Form.* • **Observational criterion:** This segment subsessile to pedunculate; *subsessile* = petiole slightly elongated posterad anterior contact surfaces, with this elongation indistinct relative to the muscular posterior node; *pedunculate* = petiole distinctly elongate posterad anterior contact surfaces, with this anterior region long, narrow, and well differentiated from muscular node (*FALSE* = petiole sessile, *sessile* = without indication of elongation posterad anterior contact surfaces OR petiole completely tubular). **• NOTES:** Relative elongation and narrowing of the second abdominal segment were observed in Loboscelidia (Chrysididae), some †Falsiformicidae, some Dryinidae, Rhopalosomatidae, various Vespidae, most Thynnoidea, most Mutillidae, Bradynobaenidae, various spheciform Apoidea (Ampulicidae, Sphecidae, some Pemphredonidae), †*Camelomecia*, and many Formicidae. Within the Formicidae, scored as *FALSE* for taxa in which have node begins immediately posterior to the anterior articulatory sclerites without any degree of narrowing.
371 *DC: Abdominal segment II (“petiole”) anterior peduncle. DS: Tergosternal composition.* **• Observational criterion:** Peduncle of this segment, *i.e.*, anterior elongation, comprising sternum only, with tergum restricted to posterior portion of segment (*FALSE* = petiole not pedunculate OR if pedunculate, then peduncle comprising tergum and sternum, regardless of whether these sclerites fused). **• NOTES:** Chosen as a character after Bohart & Menke (1976), which they use as a distinguishing feature of Sphecidae (except Ammophilini/inae) and recognize as a general feature of the Pemphredonidae (their Pemphredoninae). Among sampled taxa, this condition also occurs in *Loboscelidia* (Chrysididae), *Chyphotes* (Chyphotidae), *Bradynobaenus* (Bradynobaenidae), and †*Prolemistus apiformis* (Pemphredonidae).
372 *DC: Abdominal segment II. DS: Form.* • **Observational criterion:** This segment in form of perfectly straight, very narrow tube for its entire length (*FALSE* = this segment not in the form of a narrow, straight cylinder, *e.g.*, false if muscular node distinctly developed, whether dorsal or ventral). **• NOTES:** Observed in all sampled extant but not all extinct Evaniidae, and some †Praeaulacidae (†*Evanigaster*, and †*Evaniops*, as illustrated).
373 *DC: Abdominal segment II peduncle. DS: Form.* • **Observational criterion:** This segment with peduncle extremely long and slender, much longer than mesosoma plus head as well as "gaster" (*FALSE* = peduncle shorter than mesosoma plus head). **• NOTES:** Together with wispy setiform tibial spurs, this is another unique state observed in the vespid *Parischnogaster.*
374 *DC: Abdominal segment II. DS: Relative proportions.* • **Observational criterion:** This segment about as broad as remaining metasoma but elongate, such that it is > 1/3 length of all posterior segments (*FALSE* = this segment narrower than remaining metasoma OR if as broad as remaining metasoma, then < 1/3 length of metasoma posterad segment II).
**• NOTES:** This state was uniquely observed in the cleptoparasitic apid *Thyreus*.
375 *DC: Abdominal tergum II posterior constriction. DS: Expression.* • **Observational criterion:** This tergum, in lateral view, with a posterior face distinctly offset from a dorsal face, *i.e.*, petiole with bulging node (*FALSE* = this segment not constricted posteriorly, muscular node without distinct posterior face in lateral view). **• NOTES:** Although "petiolation" is frequently referenced for Aculeata, the condition may be broken into a series of discrete characters which, when observed all together as derived, constitute the "petiolated" condition. Such features include presence of a distinct node (*i.e.*, with distinct posterior face), and very strong posterior constriction of the segment, associated with narrowing of the articulatory sclerites of the subsequent segment. Here, a distinct node of abdominal segment II is observed in *Apterogyna* (Bradynobaenidae), the †Falsiformicidae, the “†Armaniidae”, †*Camelomecia* (including males), all stem Formicidae, and the majority of crown Formicidae with the exception of *Anomalomyrma* (Leptanillinae), and the Amblyoponinae *sensu stricto* (*i.e.*, without *Apomyrma*).
376 (**Reductive**) *DC: Posteriorly offset petiolar node. DS: Proportions.* • **Observational criterion:** Abdominal tergum II in lateral view with node that is anteroposteriorly longer than dorsoventrally tall (*FALSE* = node is dorsoventrally taller than anteroposteriorly long). **• NOTES:** Character applicable when a node is developed, as scored in the previous character. To reiterate, a node is a dorsal muscular expansion which is posteriorly constricted; *Olixon*, for example, does not have a node. This and the following two characters are scored as reductive because we want to do justice to the uncertainty regarding the placement of †*Camelomecia*. By scoring these characters reductively, they are effectively "downweighted" in Bayesian analysis, thus provide less support than a column which is filled with informative states (here, *TRUE* or *FALSE* / 1 or 0). Among sampled taxa for which this character is applicable, both the †Falsiformicidae and *Apterogyna* is *FALSE*, †*Camelomecia* (male, NIGPAS, “Camelosphecia”) is *TRUE*, †*Camelomecia* (both sexes) is *FALSE*, “†Armaniidae” are false, most stem Formicidae are *FALSE*, and variation is observable in the crown of the Formicidae.
377 (**Reductive**) *DC: Posteriorly offset petiolar node. DS: Proportions.* • **Observational criterion:** Abdominal tergum II in dorsal view with node that is lateromedially wider than anteroposteriorly long (*FALSE* = node is anteroposteriorly longer than lateromedially wide Reductively coded, thus "-" = inapplicable). **• NOTES:** Among †Falsiformicidae, only one terminal is *TRUE*, while this state is *TRUE* for “†Armaniidae”, †*Camelomecia*, most stem Formicidae, and most crown Formicidae.
378 (**Reductive**) *DC: Posteriorly offset petiolar node. DS: Form.* • **Observational criterion:** Abdominal tergum II with node that is dorsoventrally tall and very narrow anteroposteriorly, thus squamiform or nearly so (*FALSE* = node not squamiform OR relatively broad anteroposteriorly). **• NOTES:** A squamiform "petiolar node" was observed in †*Falsiformica*, †*Siccibythus*, †*Zigrasimecia*, †*Dolichomyrma longiceps, Hypoponera,* most Dolichoderinae, and most Formicinae.
379 *DC: Abdominal segment II dorsal spine. DS: Expression.* • **Observational criterion:** A single dorsal spine present on petiolar node (*FALSE* = no dorsal spines developed OR two spines present). **• NOTES:** Observed sporadically in Formicidae. Among sampled taxa, *TRUE* for *Odontomachus* and *Acanthoponera*.
380 *DC: Abdominal segment II dorsal spine. DS: Expression.* • **Observational criterion:** Paired of dorsal spines present on node (*FALSE* = no dorsal spines developed OR only one spine present). **• NOTES:** Observed sporadically in Formicidae. Among sampled taxa, *TRUE* for *Diacamma*.
381 *DC: Abdominal tergum II. DS: Shape.* • **Observational criterion:** This tergum in profile view with more-or-less vertical anterior face and posteroventrally-sloping convex dorsal/posterior face (*FALSE* = either anterior surface not vertically oriented AND/OR posterior surface concave to weakly convex OR posterior surface not distinctly developed). **• NOTES:** Among the Formicidae, this state is observed in the Ponerinae. Here scored as *TRUE* for *Neoponera* and the new genus from burmite.
382 *DC: Abdominal sternum II anterior portion. DS: Shape.* • **Observational criterion:** This sternum with anteroventral process which fits between metacoxae when metasoma fully flexed ventrally (*FALSE* = this sternum without such a process, being linear to concave anteriorly, if convex then anterior and posterior margins of the convexity evenly continuous). **• NOTES:** Primarily observed in Formicidae, wherein this structure is frequently termed the "subpetiolar process". Also observed for *Amisega* and †*Auricleptes* (Chrysididae), *Tiphia* (Tiphiidae), *Aelurus* (Thynnidae), *Dolichurus* (Ampulicidae), *Sphecius* (Bembicidae), most Philanthidae, †*Camelomecia*, most sampled Mesozoic Formicidae (quite a few terminals uncertain), and most extant Formicidae. A "subpetiolar process" was observed to be absent in *Pseudomyrmex gracilis*, *Aneuretus*, Dolichoderinae, and most Formicinae.
383 *DC: Abdominal segment II. DS: Proportions, form.* • **Observational criterion:** This segment twice as tall as long, with posterior articulation raised well above anterior articulation (*FALSE* = this segment longer than tall in lateral view OR with posterior articulation situated at about the level of the anterior articulation). **• NOTES:** Uniquely observed in *Tatuidris* (Formicidae, Agroecomyrmecinae). This may to be an autapomorphy of this genus (or species, as stands presently), as the petioles of †*Agroecomyrmex* (Baltic) and †*Eulithomyrmex* (Florissant) do not appear to match, although it is difficult to tell with the latter taxon, which represents compression fossils.
384 *DC: Tergosternal articulation of abdominal segment II. DS: Form.* • **Observational criterion:** Abdominal tergum II very broadly overlying sternum posteriorly but strongly narrowed anteriorly (*FALSE* = this tergum narrowly overlapping sternum posteriorly OR aligned with sternum OR fused to sternum). **• NOTES:** This character is from Appendix VI of Brothers & Carpenter (1993), where this state—with addition of anterior tergosternal fusion—is scored as state 1 of character 152, applying to the Chrysidoidea, Dryinoidea, and Sclerogibbidae.
385 *DC: Abdominal tergum II anterodorsal carina(e). DS: Expression.* • **Observational criterion:** One or two such carinae present; these may be oriented in any manner (*FALSE* = no such carinae developed). **• NOTES:** Modified from character 21 of Prentice (1998). Such carinae occur inconsistently across sampled taxa, here observed for *Olixon* (Rhopalosomatidae), some Vespidae, *Sierolomorpha* (Sierolomorphidae), *Tiphia* (Tiphiidae), *Smicromyrmilla* (Mutillidae), Bradynobaenidae, *Dolichurus* (Ampulicidae), some Crabronidae, Philanthidae, and most Formicidae at the subfamily level. Finer partitioning of this character is desirable but needs further study.
386 *DC: Abdominal tergum II anterior crease. DS: Expression, form.* • **Observational criterion:** This tergum with distinct anterolaterally-situated dorsoventrally oriented margins and broad anteromedian concavity which fits against propodeum (*FALSE* = anterior tergal crease not developed OR if developed then this in the form of a sulcus, rather than wide groove). **• NOTES:** This "scrobiculate" condition of abdominal segment II was observed in a few different forms in Trigonalidae, Scolebythidae, Chrysididae, and Anthophila. As these probably represent convergences, ideally this character should be split taxically, with anatomical definitions refined to match the independent origins of this state. For now, however, this aggregate character is retained. Among fossil scolebythids, clearly visible only in †*Necrobythus* and †Scolebythidae sp. (CASENT0844570, burmite); a groove is observable in the dorsal view figures of †*Aureobythus*, but it is unclear whether this conformation is equivalent, derived from, or independent from the state described here for the Scolebythidae.
387 *DC: Abdominal tergum II pilosity. DS: Form.* • **Observational criterion:** Setae of this sclerite plumose (*FALSE* = this pilosity not plumose). **• NOTES:** "Brachyplumose" setae are a synapomorphy of the Sphaerophthalminae (Mutillidae, *sensu* Brothers & Lelej 2017: state 1 of character 82). Plumose setae are, of course, a defining feature of the Anthophila as well.
388 *DC: Articulation of abdominal segments II and III. DS: Mobility.* • **Observational criterion:** These segments fused tergally, forming cone- to funnel-shaped petiole (*FALSE* = these terga freely mobile, not fused OR this articulation tergosternally fused). **• NOTES:** Observed in Gasteruptiidae, Aulacidae, and various Mesozoic evaniomorphs.
389 *DC: Articulation of abdominal segments II and III. DS: Mobility.* • **Observational criterion:** These segments fused tergosternally (*FALSE* = these terga freely mobile, not fused OR sterna of this articulation unfused). **• NOTES:** Among sampled taxa, uniquely observed in *Anomalomyrma* (see Bolton 1990b).
390 *DC: Abdominal segment II laterotergite. DS: Expression.* • **Observational criterion:** This laterotergite present, defined by crease (*FALSE* = this laterotergite absent, not defined laterally). **• NOTES:** Difficult to evaluate for fossil taxa.
391 (**Reductive**) *DC: Abdominal segment II laterotergite. DS: Size.* • **Observational criterion:** This laterotergite dorsoventrally broad (*FALSE* = this laterotergite narrow; "-" = inapplicable due to absence). **• NOTES:** Broad laterotergites are observed sporadically among those taxa where this character is applicable, including Scolebythidae, Sclerogibbidae, Dryinidae, some Vespidae, most Pompiloidea *sensu lato*, and most Apoidea.
392 *DC: Abdominal segment III laterotergite. DS: Expression.* • **Observational criterion:** This laterotergite present, defined by crease (*FALSE* = this laterotergite not developed or distinctly differentiated). **• NOTES:** Occurs with less frequency than laterotergites of abdominal segment II. Some taxa were observed to have laterotergites on subsequent abdominal segments, but thus distribution was not recorded.
393 (**Reductive**) *DC: Abdominal segment III laterotergite. DS: Size.* • **Observational criterion:** This laterotergite dorsoventrally broad (*FALSE* = this laterotergites narrow; "- " = inapplicable due to absence). **• NOTES:** Observed in Sclerogibbidae, *Euparagia* (Vespidae), *Sierolomorpha* (Sierolomorphidae), most Thynnidae, Pompilidae, and Sapygidae.
394 *DC: Abdominal segment III. DS: Form.* • **Observational criterion:** This segment anteriorly pedunculate, *i.e.*, anteriorly in form of long, straight, very narrow tube, broadening posteriorly (*FALSE* = this segment not pedunculate, *i.e.*, without differentiated tubular anterior portion and broad posterior portion). **• NOTES:** Observed uniquely in the Jurassic praeaulacid †*Nevania*.
395 *DC: Abdominal segment II posterior foramen. DS: Form.* • **Observational criterion:** Posterior foramen of petiole with tergum and sternum fused and forming perfect circle (*FALSE* = either these sclerites unfused OR with the tergal and sternal margins not aligned, not forming complete circle). **• NOTES:** This is a unique and defining state of Myrmicinae (Formicidae; see character 5 of subfamilial treatment Bolton 2003). This state may also occur in Evaniidae, but this represents an obvious convergence, and requires anatomical study (*i.e.*, dissection), to resolve.
396 *DC: Anterior articulatory sclerite of abdominal segment III (“petiolar helcium”). DS: Form.* • **Observational criterion:** This helcium with tergum and sternum in anterior view forming a rough circle, with lateral margins of tergum meeting those of sternum end-to-end (*FALSE* = sternum set within tergum, with tergal and sternal margins not aligned, thus helcium not evenly circular). **• NOTES:** Another defining synapomorphy of the Myrmicinae (see character 6 of Bolton’s 2003 definition of the subfamily). This character, in part, was used as evidence by Bolton to remove *Tatuidris* and *Ankylomyrma* from the Myrmicinae, an action later supported by molecular evidence (Ward *et al*. 2015).
397 *DC: Anterior articulatory sclerite of abdominal segment IV (“postpetiolar helcium”). DS: Location relative to postsclerites.* • **Observational criterion:** This helcium migrated dorsally, thus situated on dorsal surface of abdominal segment IV (*FALSE* = this helcium either not formed OR situated anteriorly on segment). **• NOTES:** This is a unique state of the diverse and complicated myrmicine genus *Crematogaster* (Formicidae).
398 *DC: Abdominal sternum III. DS: Form.* • **Observational criterion:** Disc of postpetiolar sternum in profile view flat to concave (*FALSE* = postpetiole not formed OR sternum bulging or otherwise distinctly convex). **• NOTES:** Among the Myrmicinae (Formicidae), this may be synapomorphy of the Stenammini + Solenopsidini + Attini + Crematogastrini, as Pogonomyrmecini and Myrmicini have ventrally bulging (convex) postpetiolar sterna.
399 *DC: Abdominal segment III tergosternal articulation. DS: Fusion.* • **Observational criterion:** This segment with tergosternal fusion (*FALSE* = this segment without such fusion). **• NOTES:** Observed uniquely in the Formicidae, where the distribution of this condition was tabulated by Bolton in Fisher & Bolton (2016). For Cretaceous taxa, scored as *FALSE* when the tergum and sternum can clearly be seen to overlap. In brief, observed for all Leptanillinae *sensu lato*, most poneroids, Dorylinae, Ectatomminae *sensu lato*, and those dolichoderines and formicines which have "shouldered" third abdominal segments (see below).
400 *DC: Abdominal tergum III anteroposterior differentiation. DS: Expression.* • **Observational criterion:** This tergum anteroposteriorly differentiated, with a distinct line or sulcus separated anterior “pretergite” from posterior “postsclerite” (*FALSE* = such differentiation not developed). **• NOTES:** "Pretergite" is used here *sensu* Bolton (1990a,b), namely, that area of the tergum which is differentiated by a transverse sulcus or other defining line, and which is not necessarily homologous with the acrotergum. This condition is widespread in the "non-chrysidiform" Aculeata, being absent in, for example, *Parischnogaster* (Vespidae), *Methoca* (Thynnidae), Pompilidae, Sapygidae, various spheciform apoids, and some Anthophila.
401 *DC: Abdominal sternum III anteroposterior differentiation. DS: Expression.* • **Observational criterion:** This sternum anteroposteriorly differentiated, with a distinct line or sulcus separated anterior “presternite” from posterior “poststernite” (*FALSE* = such differentiation not developed). **• NOTES:** "Presternite" is used here *sensu* Bolton (1990a,b), namely, that area of the sternum which is differentiated by a transverse sulcus or other defining line, and which is not necessarily homologous with the acrosternum. This condition has a somewhat broader distribution than that of presence of pretergite III, being observed in Evaniidae, *Methoca*, Pompilidae, and many spheciform apoids which lack the tergal division. That said, it should also be noted that whereas pretergite III of the Anthophila is common in frequency, distinction of the presternite is restricted to but two of the sampled Apidae.
402 (**Additive-reductive**) *DC: Abdominal presternite III. DS: Form.* • **Observational criterion:** This presternite (helcial sternite) exposed in lateral view, whether bulging below overlapping tergite or continuous with tergite (*FALSE* = concealed by tergite in lateral view; "-" = inapplicable). **• NOTES:** Initially used by Bolton (1990a,b) in his treatment of the metasomata of Formicidae as the defining synapomorphy of his dorylomorphs (= Dorylinae). Among other Aculeata, observed sporadically among those taxa which have a defined presternite, including most Vespoidea, Tiphiidae, *Apterogyna* (Bradynobaenidae), and a subset of Apoidea (Ampulicidae, some Sphecidae, some Apidae).
403 *DC: Abdominal segment III. DS: Form.* • **Observational criterion:** Anterior articulatory surfaces of this segment narrowed relative to posterior, non-articulatory surfaces (*FALSE* = these surfaces unreduced in circumference, *i.e.*, of more-or-less the same width as remainder of segment). **• NOTES:** Anterior narrowing of the presclerites of abdominal segment III is a condition of the petiolation syndrome. This state was observed in some †Praeaulacidae (†*Evaniops*, †*Habralaucus*, †*Nevania*), †Falsiformicidae (including †*Burmasphex*), Rhopalosomatinae (Rhopalosomatidae), some Vespidae (*Zethus*, *Mischocyttarus*), some Thynnoidea (*Tiphia*, Chyphotidae), *Dasymutilla* (Mutillidae), *Apterogyna* (Bradynobaenidae), *Cerceris* (Philanthidae), †*Camelomecia*, “†Armaniidae”, and all Formicidae (with the single exception of the male of one species of Leptanillinae).
404 (**Additive-reductive**) *DC: Narrowed anterior articulatory surfaces of abdominal segment III. DS: Degree of constriction.* • **Observational criterion:** These surfaces strongly narrowed relative to remainder of segment (*FALSE* = either reduced slightly; "-" = inapplicable). **• NOTES:** This character is additive on the previous but is reductive so as to not overweight independent origins of this condition. Strongly narrowed articulatory sclerites of abdominal segment III was observed in †Praeaulacidae, †*Falsiformica*, some Vespidae, *Apterogyna* (Bradynobaenidae), some “†Armaniidae” (†*Archaeopone*, †*Orapia*, †*Poneropterus*, †*Pseudarmania aberrans*), the male †*Camelomecia*, †*Camelomecia* (BALBukL_41), and most Formicidae (with the exception of *Opamyrma*, *Anomalomyrma*, and most Amblyoponinae).
405 *DC: Abdominal segment III. DS: Form.* • **Observational criterion:** This segment strongly petiolated, *i.e.*, reduced in size, often with distinct posterior face, forming "postpetiole" (*FALSE* = this segment weakly petiolated OR not petiolated). **• NOTES:** "Petiolation" of abdominal segment III was observed in *Apterogyna* (Bradynobaenidae), and some Formicidae (Leptanillinae *sensu lato* except *Opamyrma, Paraponera*, *Tatuidris*, *Myrmecia*, Pseudomyrmecinae, and Myrmicinae; also observed in the putative myrmicine †*Afromyrma petrosa.*
406 *DC: Abdominal segment III. DS: Proportions.* • **Observational criterion:** This segment anteroposteriorly foreshortened, with length ≤ 0.5 x height (*FALSE* = this segment not foreshortened, with length > 0.5 x height). **• NOTES:** Observed in some Evaniidae (including †*Palaeosyncrasis*), some stem Formicidae (†*Gerontoformica*, †*Sphecomyrma freyi*), and a number of crown Formicidae (*Tatuidris*, *Simopelta*, some Dolichoderinae, some Formicinae).
407 *DC: Abdominal poststernite III. DS: Form.* • **Observational criterion:** This poststernite "low shouldered" posterior to narrow helcium, *i.e.*, with tergosternal line strongly curving anteriorly before narrowly curving posterolaterally then posteriorly (*FALSE* = poststernite III tergosternal line directed more-or-less directly posteriorly OR this segment without differentiation between “presternite” and “poststernite”). **• NOTES:** "Shouldering" of abdominal segment III in this sense is observed convergently in Formicinae and Dolichoderinae (Formicidae).
408 (**Additive-reductive**) *DC: Shouldering of abdominal poststernite III. DS: Form.* • **Observational criterion:** This poststernite "high shouldered" posterior to narrow helcium, *i.e.*, with tergosternal line immediately lateral to helcium directed dorsally before curving strongly posteriorly, the tergal area between the raised shouldered concave and concealing the petiole completely in dorsal view when at rest (*FALSE* = "low shouldered"; "-" = inapplicable due to absence of shouldering). **• NOTES:** This state is logically additive on that of the *TRUE* condition of the previous character and is treated as reductive so as to avoid overweighting this condition, which is clearly convergent within the Formicidae.
409 *DC: Anterior articulatory surfaces of abdominal segment III. DS: Relative dorsoventral position.* • **Observational criterion:** Axiality 1: These surfaces set distinctly below segment midheight (= “infraaxial condition”) (*FALSE* = these surfaces either at midheight, *i.e.*, “axial”, OR above midheight, "supraaxial"). **• NOTES:** This and the following character constitute two of the three conditions of axiality, as defined by Keller (2011). As formulated, *FALSE* for "infraaxial" and for "supraaxial" equals the "axial" condition. Because *FALSE* for both states is the predominant state across the phylogeny of the Hymenoptera, it stands to reason that this is the ancestral condition. Therefore, to avoid overweighting symplesiomorphy in the present matrix, the "axial" condition is not scored as an independent character, particularly as this would exacerbate homoplasy observed in the two other forms. Among sampled taxa, infraaxiality was observed in the following taxa: †*Falsiformica* (†Falsiformicidae), *Chyphotes* (Chyphotidae), and a subset of Formicidae, including †*Ceratomyrmex*, †*Haidoterminus*, †*Boltonimecia*, †*Gerontoformica*, †*Sphecomyrma*, †*Cretomyrma*, *Apomyrma* (Amblyoponinae), Ponerinae, *Aneuretus*, Dolichoderinae, most Formicinae, Ectatomminae *sensu lato* (including †*Canapone*), and most sampled Myrmicinae. Bolton (2003) treated *Aneuretus* as having a supraaxial helcium, but here the helcium is interpreted as infraaxial, as posttergite III has a short anterior face dorsal to the helcium and poststernite III is completely flat ventral to the helcium.
410 *DC: Anterior articulatory surfaces of abdominal segment III. DS: Relative dorsoventral position.* • **Observational criterion:** Axiality 2: These surfaces set distinctly above segment midheight (= “supraaxial condition”) (*FALSE* = either at midheight, *i.e.*, “axial condition”, OR below midheight, *i.e.*, "infraaxial"). **• NOTES:** See comments on prior character. Among sampled taxa, relative to the infraaxial condition, the supraaxial condition is on the one hand, more widespread outside of the Formicidae, and on the other less widespread within the Formicidae. Here, supraaxiality was observed for *Sierolomorpha* (Sierolomorphidae), most Thynnoidea (except *Brachycistis*, *Chyphotes*), all Scolioidea, Ampulicidae, *Clypeadon* (Philanthidae), and a subset of Formicidae (†*Afropone oculata*, *Anomalomyrma*, and Amblyoponinae except *Apomyrma*).
411 *DC: Abdominal sternum III anteromedian contact surface margination (“prora”). DS: Expression.* • **Observational criterion:** This sternum with raised ridge or bosses anteriorly, surrounding petiolar contact surface (*FALSE* = this sternum without such margination). **• NOTES:** In the myrmecological literature, this is referred to as the “prora” (*e.g.*, Bolton 1990, Bolton & Fisher 2011). Prorae were observed in *Brachycistis* (Tiphiidae), *Chyphotes* (Chyphotidae), †*Camelomecia*, most stem Formicidae, and most subfamilies of the crown Formicidae (except Aneuretinae, Dolichoderinae, Formicinae, *Opamyrma*, *Apomyrma*, and *Crematogaster*, among sampled taxa). The prora was illustrated as present in "†*Cananeuretus occidentalis*?" by Engel & Grimaldi (2005), but this state needs to be confirmed. Within the Formicidae, there is considerable and informative variation of the prora, but this is aggregated into but three characters (reductively scored; see below).
412 (**Additive-reductive**) *DC: Abdominal sternum III prora. DS: Form.* • **Observational criterion:** This prora in form of longitudinal keel (*FALSE* = prora of another form; "-" = prora absent). **• NOTES:** This conformation was observed in *Brachycistis* (Tiphiidae), †*Zigrasimecia* and some †*Gerontoformica.* This state was absent among all sampled crown ants with a prora.
413 (**Additive-reductive**) *DC: Abdominal sternum III prora. DS: Form.* • **Observational criterion:** This prora in form of transverse lip- or tooth-like projection (*FALSE* = prora of another form; "-" = prora absent). **• NOTES:** This is the predominant state that was observed in crown Formicidae.
414 (**Additive-reductive**) *DC: Abdominal sternum III prora. DS: Form.* • **Observational criterion:** This prora transverse lip-like, and strongly pronounced (*FALSE* = prora of another form; "-" = prora absent). **• NOTES:** This extreme condition was observed in *Rhytidoponera* (Ectatomminae) and †*Canapone occidentalis.*
415 *DC: Abdominal tergum III felt line. DS: Expression.* • **Observational criterion:** This felt line present, *i.e.*, dense setose patch in form of distinct stripe on lateral tergal surface (*FALSE* = such a setose patch absent). **• NOTES:** Felt lines (*e.g.*, Goulet & Huber 1993) have evolved independently in Thynnoidea, Mutillidae, and Bradynobaenidae, being observed among sampled taxa in *Typhoctes*, most mutillids, and *Apterogyna*. Surprisingly, little variation is recorded among the groups anatomically (Debolt 1973).
416 *DC: Metasoma posterior to petiole in females. DS: Count of exposed terga.* • **Observational criterion:** Female: metasomal segments II+ (abdominal segments III+) forming "gaster" (unconstricted tagma) with four exposed terga, segment VI (AVII) being reduced and usually concealed in dorsal view by tergum V (AVI) (*FALSE* = metasomal segments II+ [AIII+] constricted AND/OR metasomal tergum VI [AVII] visible in dorsal view, not retracted beneath tergum of segment V [AVI]). **• NOTES:** Observed in Dolichoderinae; metasomal segment VI exposed in all other Formicidae, regardless of conformation of "gaster", but phrasing intended to clarify state relative to other Aculeata with reduced abdominal segmentation, such as Chrysididae.
417 *DC: Pregenital oviposition complex. DS: Expression.* • **Observational criterion:** Female: metasomal sterna III or III and IV with strong median processes associated with "hole-punching" method of oviposition (*FALSE* = such a complex not developed). **• NOTES:** Observed in some Trigonalidae.
418 *DC: Abdominal segment IV. DS: Constriction.* • **Observational criterion:** Cinctus of this segment (metasomal III) present, defining presclerites whether dorsal or ventral (*FALSE* = such a constriction not developed). **• NOTES:** This condition is correlated with formation of a "postpetiole". Taylor (1978) included this as one of the characters describing the "tubulation syndrome". Here, "cinctus" is used in the sense of Bolton (*e.g.*, Bolton 2003), and is used in an equivalently with "gradulus" of Michener (*e.g.*, Michener 1944). Among sampled taxa, cincti were observed in some Scolioidea, some Apoidea (including most Anthophila), †*Camelomecia*, and numerous stem and crown Formicidae. Among crown taxa, *Opamyrma* has a ventral but no dorsal cinctus for segment IV.
419 *DC: Abdominal segment IV tergosternal articulation. DS: Form.* • **Observational criterion:** Lateral margins of tergum and sternum of this segment (metasomal III) meeting and aligned for their entire length (*FALSE* = this tergum overlapping this sternum, margins not aligned for their length). **• NOTES:** This is the second of the characters describing the tubulation syndrome of Taylor (1978) and is more-or-less equivalent to the condition of tergosternal fusion of abdominal segment IV as tabulated by Bolton in Fisher & Bolton (2016). Observed in †*Camelomecia*, some †Haidomyrmecini, some †*Zigrasimecia*, some †Sphecomyrmini, most poneroids (absent in *Apomyrma*, *Adetomyrma*), and Ectatomminae *sensu lato*. For evaluation of Cretaceous taxa, scored as "0" when the tergum and sternum can clearly be seen to overlap.
420 *DC: Abdominal postsclerites IV. DS: Relative length.* • **Observational criterion:** This poststernite (metasomal III) ventral margin in lateral view distinctly short relative to the isosegmental posttergite dorsal margin or equivalent area (*FALSE* = poststernite AIV/MIII as long as posttergite AIV/MIII OR this segment without transverse constricting sulcus). **• NOTES:** This condition is observed in †*Camelomecia* sp. (ANTWEB1038930), and among some Formicidae, including some †*Zigrasimecia*, some †*Gerontoformica*, †*Sphecomyrma freyi*, *Opamyrma* (Leptanillinae), and most poneroids
421 *DC: Abdominal tergum IV. DS: Relative size.* • **Observational criterion:** This tergum (metasomal III) comparatively large, being longer than petiole, postpetiole, and abdominal segments V+ (metasomal IV+) (*FALSE* = this tergum comparatively smaller, not longer than petiole, postpetiole, and posterior segments). **• NOTES:** This was observed in some Formicidae, including *Anomalomyrma*, *Tatuidris*, *Proceratium*, and the Myrmicinae.
422 *DC: Abdominal tergum IV. DS: Form.* • **Observational criterion:** This tergum (metasomal III) vaulted, *i.e.*, in profile with dorsal margin strongly convex (*FALSE* = this tergum not vaulted, *i.e.*, weakly convex or linear). **• NOTES:** This condition has been homoplastically derived in *Tatuidris*, Proceratiinae, and *Gnamptogenys*, among other Formicidae.
423 *DC: Abdominal tergum IV stridulitrum. DS: Expression.* • **Observational criterion:** Stridulitrum on this tergum present, *i.e.*, anteromedian texture of abdominal pretergite IV distinctly differentiated, forming a fine comb (*FALSE* = this stridulitrum absent). **• NOTES:** Tergal stridulitra were observed in a minority of sampled taxa, including *Olixon* (Rhopalosomatidae), most Mutillidae (see also Brothers & Lelej 2017), and some Formicidae. The stridulitrum was impossible to observe for most fossil taxa. Upon review after analysis, we expect reevaluation of the stridulitrum, especially in Ponerinae, will reveal inadequate scoring (*e.g.*, *Neoponera* definitely has a stridulitrum).
424 *DC: Abdominal sternum IV stridulitrum. DS: Expression.* • **Observational criterion:** Stridulitrum of this sternum present (*FALSE* = this stridulitrum absent). **• NOTES:** Ventral stridulitra are exceedingly rare. Bolton (2003) recorded them for *Nothomyrmecia* and some *Rhytidoponera.* As *Nothomyrmecia* was not included, this is a unique state of *Rhytidoponera* in the present matrix. Examination of scanning electron micrographs of *Nothomyrmecia* show a lack of differentiated stridulatory area on the presternite of abdominal segment IV, but presence of an unusual posterior bulge of abdominal poststernite III.
425 *DC: Abdominal segment V girdling constriction (“cinctus”). DS: Expression.* • **Observational criterion:** This cinctus present, defining presclerites whether dorsal or ventral (*FALSE* = this cinctus absent). **• NOTES:** As above, "cinctus" is used equivalently as "gradulus". Cincti beyond segment IV were observed in some Formicidae that were not sampled (*e.g.*, *Zasphinctus*, Dorylinae). Outside of the Formicidae, these were observed in *Tachysphex* (Crabronidae), some Philanthidae, and most Anthophila.
426 *DC: Abdominal segment VI girdling constriction (“cinctus”). DS: Expression.* • **Observational criterion:** This cinctus present, defining presclerites whether dorsal or ventral (*FALSE* = this cinctus absent). **• NOTES:** See comments for the previous character. Here, scored as *TRUE* for most Philanthidae and Anthophila.
427 *DC: Posteromedian triangular process of female abdominal sternum VII (metasomal VI) of female. DS: Expression.* • **Observational criterion:** Female: this process present, this process has a surface which is distinct relative to the remainder of the sclerite (*FALSE* = such a process absent). **• NOTES:** Observed uniquely in *Clystopsenella* (Scolebythidae).
428 *DC: Paired horn-like processes on lateral margins of abdominal sternum VII (metasomal VI) of female. DS: Expression.* • **Observational criterion:** Female: these processes present (*FALSE* = these processes absent). **• NOTES:** This was observed uniquely in Scoliidae, although the condition in *Proscolia* is uncertain.
429 *DC: Abdominal tergum VII of female and VIII of male (metasomal VI and VII). DS: Form.* • **Observational criterion:** These terga modified as "pygidial plate", *i.e.*, produced posteriorly and/or marginate laterally, often with medial sculpturation (*FALSE* = these terga convex, without lateral margination). **• NOTES:** Observed in some Thynnoidea, Rhopalomutilla (Mutillidae), Scolioidea, some Crabronidae, Sphecius (Bembicidae), some Pemphredonidae, some Philanthidae, †Melittosphecidae, and some Anthophila. Danforth & Poinar (2011) recorded this as synapomorphy of "Crabronidae", †Melittosphecidae, and crown Anthophila. Arguably, based on this definition, some Formicidae (*e.g.*, *Pachycondyla*, "cerapachyine" Dorylinae) could be scored as *TRUE*. Greater anatomical definition may be used to split this character for each approximate origination.
430 *DC: Pilosity of female abdominal tergum VII (metasomal VI). DS: Form.* • **Observational criterion:** Female: This tergum covered in stout, appressed setae which taper to a fine point and resemble scales (*FALSE* = setae of this tergum thinner, hairlike, not squamiform). **• NOTES:** Observed uniquely in the campsomerine scoliids *Cathimeris* and *Campsomeris.*
431 *DC: Abdominal sternum VII (metasomal VI) of male. DS: Form.* • **Observational criterion:** Male: this sternum "unciform", *i.e.*, in the form of upturned stinger- or thorn-like process (*FALSE* = this sternum not in this form, being broader and shield-like, posterior margination variable). **• NOTES:** Observed in Thynnoidea (except *Tiphia*, *Aelurus*), *Apterogyna* (Bradynobaenidae), †*Thanatotiphia*.
432 *DC: Abdominal tergum VIII of female. DS: Size, exposure.* • **Observational criterion:** Female: this tergum fully internalized (*FALSE* = this tergum not reduced, usually visible externally). **• NOTES:** Described as the defining synapomorphy of the Aculeata to the exclusion of the Chrysidoidea *sensu lato* (*e.g.*, , Brothers 1975, Carpenter 1986, Rasnitsyn 1988, Brothers & Carpenter 1993, Ronquist *et al*. 1999). Ronquist *et al*. (1999) call this state 1 of character 121, including the additional, necessary criterion "desclerotized medially except for antecosta, spiracles hypertrophied". Because this additional condition could not be evaluated for fossil forms, the character was simplified to the count of external terga of the female. Consequently, Chrysididae are scored as *TRUE*, despite the fact that this is almost certainly convergent with the vespoid-apoids. Among putative sampled taxa, †*Burmomyrma* Dlussky was illustrated with eight terga, but was scored as "?" as this condition could not be validated. At least some female †Ephialtitoidea have, apparently, nine exposed abdominal terga; an effort was made to characterize this condition for the present matrix but was thrown out during the quality control phase due to the difficulty of evaluating this for the evaniomorphs.
433 *DC: Ovipositor. DS: Relative length.* • **Observational criterion:** Ovipositor long, about as long as or longer than metasoma (*FALSE* = this apparatus shorter, length less than that of the metasoma). **• NOTES:** (1) Elongate ovipositors were observed in many †Ephialtitoidea, †Praeaulacidae, †Baissidae, Aulacidae, Gasteruptiidae (except *Gasteruption kirbii*; other species with long ovipositors), †Othniodellithidae, Evaniidae, and †Maimetshidae. (2) Rasnitsyn (e.g., Rasnitsyn 1988, 2000) used concealment or very short length of the ovipositor as a proxy for the sting, and argued that, for this reason, the †Bethylonymidae were sister to the Aculeata. Because of the difficulty of evaluating this condition for fossils, and because of variability in sting length among aculeates, this character is not scored in the present matrix. (3) Presence or absence of a sting as a character was thrown out of the matrix during the quality control (post data-entry) phase of the present study as too many assumptions are required to score this as an autapomorphy of the Aculeata. From the functional perspective, loss of an ovipositing and gain of an envenomation (sting) function is a behavioral character and cannot be evaluated in fossils, thus all fossils—even those which are clearly aculeates—would have to be assumed to have this function to be scored as *TRUE*, which is untenable. From the anatomical perspective, after study of the literature, it is apparent that the ovipositor apparatus is in dire need of substantial post-genomic anatomical study, with explicit comparisons made across the Aculeata. Moreover, the supposed anatomical synapomorphies of the Aculeata cannot be evaluated in most fossils without application of advanced scanning technologies, which is out of the scope of the present study. These aculeate syn- or autapomorphies include presence of the "postarticulatory incision" of the second valvifer (Oeser 1961), the “basal line” which is a more-or-less filamentous process of the second valvifer (*sensu* Barbosa *et al*. 2021), the furcula (Oeser 1961), and possibly absence of genital membrane muscles (Barbosa *et al*. 2021). In brief, the original study (Oeser 1961) was limited in that only Chrysididae, Bethylidae, and *Apis mellifera* were compared. Subsequent study documented presence of the furcula in Braconidae (Hermann & Morrison 1980), absence of the furcula in Dryinidae, Embolemidae, and Sclerogibbidae (Rasnitsyn 1980; Barbosa *et al*. 2021), and otherwise did not evaluate the postarticulatory incision outside of the Chrysidoidea *sensu lato* (Carpenter 1986). Besides uncertainty about the distribution of the incision among Apocrita, given the genomic phylogeny of Branstetter *et al*. (2017b) and subsequent reanalysis by Tagliacollo & Lanfear (2018), it is unclear whether loss of the furcula is a synapomorphy for the Dryinidae, Embolemidae, and Sclerogibbidae (as proposed by Carpenter 1986 and supported by Barbosa *et al*. 2021), or if the furcula was gained twice in the Aculeata, or, indeed, if the furcula is a symplesiomorphy of the clade. Further sampling is necessary.
434 *DC: Abdominal sternum VII of female. DS: Form.* • **Observational criterion:** Female: this sternum apically folded forming the "acidopore" which is usually lined with a distinct ring of setae, the "coronula" (*FALSE* = this sternum not folded apically). **• NOTES:** This is the unique morphological condition of the Formicinae (Formicidae; *e.g.*, Hung & Brown 1966, Bolton 1994, 2003) which facilitates the spraying of venom (acid) and which is clearly visible in †*Kyromyrma* as well as the new formicine from burmite.
435 *DC: Sting. DS: Development.* • **Observational criterion:** Sting reduced to nonfunctionality AND acidopore and coronula absent (*FALSE* = sting not reduced OR acidopore and coronula present). **• NOTES:** The particular descriptive formulation here provided is intended to separate the Dolichoderinae from the Formicinae, the two major ant subfamilies which have lost the sting. While functional loss of the sting has historically led to the lumping of these two clades (*e.g.*, Wilson *et al*. 1967a,b), it is obvious based on molecular phylogenetic results that this is a parallelism. Technically, the two conditions of can be distinguished by presence of the furcula in Formicinae and fusion of the furcula to the sting base in Dolichoderinae (*e.g.*, Hermann & Chao 1983). In Formicinae, the furcula has been coopted for control of venom spray (Hermann 1983), while the fusion state of the furcula has been observed in Aneuretinae, Dolichoderinae, Dorylinae, and *Simopelta* (Ponerinae) (Hermann & Chao 1983). Although we, the authors, have not been stung by *Simopelta*, it is clear (BEB) that New World dorylines can inveigh with the shaft of their sting, thus experience leads to reason that the distinction between "reduction" and "nonfunctionality" are not equivalent. Because furcular conformation was not here evaluated, the probable synapomorphic condition of fusion for the Aneuretinae and Dolichoderinae is not scored. Notably, reduction of the sting and retention of the furcula was observed in the titanic Eocene †Formiciinae (Lutz 1986), implying a close relationship with the Formicinae or, perhaps more probably, another parallel reduction of the sting shaft from an unknown and well-endowed ancestor.
436 *DC: Metasomal apex of female. DS: Form.* • **Observational criterion:** Female: metasomal apex strongly downcurved, thus the terminal opening is directed more-or-less ventrally, rather than posteriorly (*FALSE* = female metasomal apex directed more-or-less posteriorly). **• NOTES:** Here observed in Trigonalidae.
437 *DC: Abdominal sternum IX of male (metasomal VIII). DS: Form.* • **Observational criterion:** Male: this sternum peglike and not acute apically (*FALSE* = this sternum not peglike, apex variable in form). **• NOTES:** Unique to Sierolomorphidae, from Brothers & Carpenter (1993, state 1 of character 165 in Appendix VI).
438 *DC: Spiniform processes of male abdominal sternum IX (metasomal VIII). DS: Expression.* • **Observational criterion:** Male: these processes present, widely spaced (*FALSE* = such processes absent). **• NOTES:** Observed in Scoliinae (Scoliidae).
439 *DC: Terminal abdominal segments. DS: Internalization, form.* • **Observational criterion:** Abdomen with ≤ 5 exposed segments in females, ≤ 6 in males (≤ 4 and ≤ 5 metasomal) AND remaining segments narrow and telescoping completely into preceding segments (*FALSE* = abdomen with ≥ 6 exposed segments in females, ≥ 7 in males [≥ 5 and ≥ 6 metasomal]). **• NOTES:** This condition is associated with reduction of the sting apparatus to lancets and sheaths, and a distinctly subovoid shape of the first few metasomal segments in dorsal or ventral view (character 18 of Bohart & Kimsey 1982). This modification is a synapomorphy of the Chrysididae including the Cleptinae and Loboscelidiinae (see Day 1979 for abdominal anatomy of latter). Lucena & Melo (2018), drawing from Carpenter (1999), report a count of "terga" in their characters 37 and 38; for reference, these are metasomal, while total abdominal segment counts are recorded here.
440 (**Additive**) *DC: Terminal abdominal segments. DS: Internalization, form.* • **Observational criterion:** Abdomen of female with ≤ 4 exposed segments, ≤ 5 in males (≤ 3 and ≤ 4 metasomal) AND remaining segments of female narrow and telescoping completely into preceding segments (*FALSE* = abdomen with ≥ 5 exposed segments in females, ≥ 6 in males [≥ 4 and ≥ 5 metasomal]). **• NOTES:** Considered a synapomorphy of Chrysidinae (*e.g.*, Kimsey & Bohart 1990).
441 (**Additive**) *DC: Terminal abdominal segments. DS: Internalization, form.* • **Observational criterion:** Abdomen of female and male with 3 exposed segments (2 metasomal) AND remaining segments of female narrow and telescoping completely into preceding segments (*FALSE* = abdomen with ≥ 4 exposed segments in females, ≥ 6 in males [≥ 3 and ≥ 5 metasomal]). **• NOTES:** Derived in Allocoeliini (Chrysidinae, Chrysididae; *e.g.*, Kimsey & Bohart 1990).
442 *DC: Abdominal segment II (metasomal I). DS: Form.* • **Observational criterion:** This segment appearing anteriorly truncate in dorsal view, *i.e.*, anterior margin more-or-less linear for most of its width across the propodeometasomal articulation AND with paired lobate processes lateral to the propodeal articulation which are dorsoventrally compressed ("pinched") (*FALSE* = this segment not appearing truncate AND without such paired lateral lobate processes, *i.e.*, anterior margin of AII/MI angled or arched from the propodeometasomal articulation). **• NOTES:** Observed in Chrysidinae: *Allocoelia*, *Hedychrum*, *Parnopes*, and *Argochrysis.*
443 *DC: Metasomal sternites II+. DS: Sclerotization, form.* • **Observational criterion:** These sternites soft, not convex, thus metasoma after AII/MI concave, *i.e.*, metasoma dome-shaped in cross-section, venter depressed (*FALSE* = these sternites stiff, convex, thus metasoma after AII/MI not depressed, *i.e.*, metasoma elliptical to circular in cross-section). **• NOTES:** Synapomorphy of Chrysidinae (character 32 of Kimsey & Bohart 1990). †*Bohartiura* scored by Lucena & Melo (2018) as having a flat metasomal venter, however the authors noted in the description that the specimen is "apparently convex or flat", therefore as no refinement can be made based on their provided figure 5, and this taxon scored as "?" in the present matrix. Crucially, this condition was confirmed for †Chrysidinae sp. (CASENT0844585, burmite).
444 *DC: Metasomal terga. DS: Relative lengths.* • **Observational criterion:** Metasomal terga I and II longer than III+ (*FALSE* = MI and II not longer than III+). **• NOTES:** Apomorphy occurring in Amiseginae, Loboscelidiinae, and Chrysidinae (Chrysididae); also occurring in some Vespidae and Ampulicidae. The distribution of this condition may be more widespread than here recorded.
445 *DC: Posterior margin of metasomal tergum III (abdominal IV). DS: Form.* • **Observational criterion:** This margin with transverse rim of denticles (*FALSE* = this margin without denticulate rim). **• NOTES:** Observed in Chrysidini and Parnopini (Chrysidinae), although in the former these denticles occur on the distalmost rim of the sclerite and in the latter, the distalmost rim is downfolded, and above this the denticles occur (characters 35 and 36 of Kimsey & Bohart 1990). Absent in †Chrysidinae sp. (CASENT0844585, burmite).
446 *DC: Posterior region of metasomal tergum III (abdominal IV). DS: Sculpture. DS: .* • **Observational criterion:** This region with transverse and linearly-arrange set of foveae (*FALSE* = such foveate line not developed). **• NOTES:** Synapomorphy of the Chrysidini (Chrysididae, character 34 of Kimsey & Bohart 1990, where this is described as the "pit row").
447 *DC: Metasomal tergum III (abdominal IV). DS: Shape.* • **Observational criterion:** This tergum with apical margin downcurved and thickened (*FALSE* = this tergum not thickened and downcurved, being more-or-less evenly curved in lateral view). **• NOTES:** Unique synapomorphy of the Parnopini (Chrysididae, character 36 of Kimsey & Bohart 1990).
448 *DC: Ovipositor tube. DS: Form.* • **Observational criterion:** This tube reduced in size, observable as narrow needle-like structure (*FALSE* = this tube comparatively thick OR absent). **• NOTES:** Scored as *TRUE* for *Amisega* and *Loboscelidia* (Chrysididae) based on Day (1979), and *FALSE* for fossil Chrysididae in which the ovipositor tube is clearly thick and well-developed.

**-** *Forewing*

449 *DC: Tegulum and parascutal area of mesonotum. DS: Size, shape.* • **Observational criterion:** This sclerite massive, covering entire base of fore and hind wings AND locking into a notch of the mesonotum (*FALSE* = this sclerite smaller, axillary sclerites of fore and hind wings exposed to variable degrees AND/OR not capable of locking into mesonotal notch). **• NOTES:** Extremely large tegulae were observed in the present study in the chrysidids *Loboscelidia* (Loboscelidiinae) and *Parnopes* (Chrysidinae). Distinguishing these two taxa is that *Loboscelidia*, a probable myrmecophile, has a locking mechanism in the mesonotum, whereas *Parnopes* does not.
450 *DC: Tegulum and parascutal area of mesonotum. DS: Size, shape.* • **Observational criterion:** This sclerite massive, covering entire base of fore and hind wings but NOT locking into a notch of the mesonotum (*FALSE* = this sclerite smaller, axillary sclerites of fore and hind wings exposed to variable degrees AND/OR capable of locking into mesonotal notch). **• NOTES:** This extreme case observed in the myrmecophilic genus *Loboscelidia*.
451 *DC: Radius distad Sc+R+Rs split. DS: Extent.* • **Observational criterion:** Free radial vein (Rf) reaching wing apex or almost to wing apex, regardless of whether the radial sector reaches the apex (*FALSE* = Rf distinctly separated from wing apex). **• NOTES:** The extent to which tubular wing venation extends to the apex of the wing was observed to be complex and informative across examined taxa. Here, six character-state definitions were developed and evaluated to capture this information: (1) Rf extending to the wing apex (this character), (2) Rsf contacting wing apex, (3) Rsf almost extending to wing apex, (4) Rsf clearly separated from apical region of wing, (5) tubular wing venation more-or-less absent from apical third of wing, and (6) venation restricted to proximal half of wing. A distinction was made between Rf and Rsf contacting the wing apex, as in many taxa Rsf is clearly separated from the wing apex, but Rf extends distally beyond the "marginal cell" to or nearly to the apex. During quality control, the fourth character (Rsf separated from apical region of wing) was deleted, as the information was complementary (and virtually identical) to the prior character (Rsf approaching or reaching wing apex). The sixth character is combined with heavy sclerotization to define the Bradynobaenidae. Among sampled taxa, Rf extended to or nearly to the wing apex in various †Ephialtitoidea (most taxa uncertain), most †Praeaulacidae, most †Anomopterellidae, †Baissidae, Aulacidae, Gasteruptiidae, many extinct Evaniidae, a few †Maimetshidae, †*Albiogonalys* (Trigonalidae), some Vespidae, and some Formicidae.
452 *DC: Radial sector distad Sc+R+Rs split. DS: Extent.* • **Observational criterion:** Free radial sector vein (Rsf) contacting wing apex (*FALSE* = Rsf not reaching wing apex). **• NOTES:** See NOTES for prior character. Here, Rsf was observed to contact the wing apex in a minority of taxa, including †*Thilopterus* (†Ephialtitoidea), some †Praeaulacidae, a few †Anomopterellidae, and some Aulacidae.
453 (**Additive**) *DC: Radial sector distad Sc+R+Rs split. DS: Extent.* • **Observational criterion:** Rsf extending to OR almost to distalmost wing margin (*FALSE* = Rsf clearly separated from distalmost wing margin). **• NOTES:** As this character is formulated, it is additive with respect to the *TRUE* state of the prior character. Slightly more than half of the examined taxa were scored as *FALSE*; taxa for which Rsf was observed to approach or reach the wing apex include †Ephialtitoidea, †Anomopterellidae, †Andreneliidae, most †Baissidae (except †*Humiryssus*), Aulacidae, Gasteruptiidae, †*Mesevania* (Evaniidae), most †Bethylonymidae (exceptions for taxa for which wing apices could not be observed), some †Maimetshidae, Trigonalidae, †Scolebythidae sp. (CASENT0844570, burmite), a †*Burmasphex*, †*Sclerogibba cretacica*, †Vespoidea sp. (CASENT0844576, burmite), Rhopalosomatidae, most Vespidae, †Sierolomorphidae sp. (CASENT0844589, burmite), †*Bryopompilus*, †*Burmusculus*, †*Cretosapyga*, most Thynnidae, *Sapyga* (Sapygidae), Myrmosinae (Mutillidae), most "†Angarosphecidae", *some* Ampulicidae, †Sphecidae sp. (CASENT0844566), †Apoidea sp. (CASENT0844594, burmite), some Pemphredonidae, *Apis* (Apidae), †*Camelomecia*, †*Orapia* (“†Armaniidae”), and some Formicidae (†*Baikuris maximus*, *Dicamba*, *Aneuretus*, *Leptomyrmex*, and *Stenamma*). According to Brothers & Lelej (2017), venation not reaching the distalmost wing margin defines Mutillidae exclusive of Myrmosinae; as noted above, this state occurs homoplastically in various groups, and is extreme in Bradynobaenidae (next character).
454 *DC: Forewing venation, collectively. DS: Extent.* • **Observational criterion:** Tubular to nebulous wing venation absent from most or all of distal 0.20–0.33 of wing (*FALSE* = non-spectral venation extending into distal fifth to third of wing). **• NOTES:** This condition was observed to be *TRUE* for a minority of taxa, including Plumariidae, some Scolebythidae, a bethylid (new genus near Scleroderminae), *Loboscelidia* (Chrysididae), *Sclerogibba* (Sclerogibbidae), †Pompiloidea sp. (CASENT0844580, burmite), *Brachycistis* (Tiphiidae), Chyphotidae, Scolioidea (Scoliidae and Bradynobaenidae, but not fossils attributed to the Scoliidae), †*Apodolichurus*, †*Burmastatus*, *Heterogyna* (Heterogynaidae), and Ammoplanidae.
455 *DC: Forewing venation, collectively. DS: Extent, degree of sclerotization.* • **Observational criterion:** Venation of "bradynobaenid pattern", *i.e.*, very heavily sclerotized veins present in basal half of wing, absent in distal half (*FALSE* = venation with greater longitudinal extent and not so heavily sclerotized). **• NOTES:** (1) Given the molecular phylogenetic and phylogenomic results of Pilgrim *et al*. (2008) and Branstetter *et al*. (2017b), it is clear that the condition here described is a unique condition of the Bradynobaenidae in its restricted sense, *i.e.*, including Bradynobaeninae and Apterogyninae, and excluding the Typhoctinae and Chyphotinae, which together constitute the Chyphotidae in the expanded Thynnoidea, as here treated. (2) Another apparently taxon-restricted character was included during the character development phase of this matrix but was deleted during the quality control phase: "Forewing Rsf and rs-m crossveins spectral or absent proximal to 2-rs, thus Rsf2+ starting from 2r-rs, forming "pterostigmal hook vein’". This condition, *i.e.*, that of the presence of a "pterostigmal hook vein" was observed among the sampled taxa in many Chrysidoidea, the Dryinoidea, †*Cretotrigona* (Apidae), *Anomalomyrma* (Leptanillinae, Formicidae), and *Lioponera* (Dorylinae, Formicidae). This character was deleted because it is both plausible that this condition is homoplastic and, more importantly, because all of the pertinent information was captured by downstream characters.
456 *DC: Longitudinal crease or fold line of forewing membrane (“remigium”). DS: Expression.* • **Observational criterion:** Such a line present (*FALSE* = such a line present). **• NOTES:** This is the vespid "plait" of Carpenter (1982) and Danforth & Michener (1988). Based on the topology of Piekarski *et al*. (2018), this plait is an apparent autapomorphy of Eumeninae + Zethinae + Polistinae + Vespinae within the Vespidae. In the present matrix, scored as *TRUE* for *Pachodynerus*, *Zethus*, and *Mischocyttarus.* The state is uncertain for the New Jersey fossil ?*Symmorphus senex* Carpenter, 2000.
457 *DC: Apical longitudinal wrinkling (serial grooves) of fore- and hindwing membranes. DS: Expression.* • **Observational criterion:** Such wrinkling present (*FALSE* = such wrinkling absent). **• NOTES:** These wrinkles are diagnostic of the Scolioidea *sensu* Pilgrim *et al*. (2008) and Branstetter *et al*. (2017b). Apical wrinkling also observed in *Evania* (Evaniidae), *Pepsis* (Pompilidae), *Dasymutilla* (Mutillidae), and some Anthophila. Among Mesozoic fossils placed in the Scoliidae, only †*Cretaproscolia josai* and †*Sinoproscolia yangshuwanziensis* had evidence of this condition.
458 *DC: Pterostigma. DS: Size.* • **Observational criterion:** This thickening grossly enlarged (*FALSE* = pterostigma not as enlarged OR absent). **• NOTES:** An enlarged pterostigma was observed sporadically among sampled taxa, including †*Westratia* (†Praeaulacidae), most †Baissidae, †Plumalexiidae, Plumariidae, Scolebythidae, some Bethylidae, *Parischnogaster* (Vespidae), most Thynnoidea, *Bradynobaenus* (Bradynobaenidae), Ammoplanidae, †*Psolimena* (Pemphredonidae), *Ceratina* (Apidae), and some Formicidae (including *Anomalomyrma*, most Amblyoponinae, *Proceratium*, *Platythyrea*, *Lioponera*, some Dolichoderinae, and the myrmicine *Stegomyrmex*).
459 *DC: Pterostigma. DS: Size.* • **Observational criterion:** This thickening strongly reduced, being absent or virtually absent (*FALSE* = pterostigma conspicuous, even if small). **• NOTES:** As with gross enlargement, strong reduction was observed sporadically among sampled taxa, including some Bethylidae, various Chrysididae, some †*Burmasphex*, some Embolemidae, Sclerogibbidae, Rhopalosomatidae, some Vespidae, some Thynnoidea, some Mutillidae, *Apterogyna* (Bradynobaenidae), some "†Angarosphecidae", most Sphecidae, most Crabronidae, Bembicidae, some Anthophila, some †*Camelomecia*, and some Formicidae (including *Apomyrma*, *Discothyrea*, and *Leptomyrmex*).
460 *DC: Pterostigma. DS: Location.* • **Observational criterion:** This thickening distinctly situated in the apical third of forewing (*FALSE* = pterostigma situated proximad forewing apical third). **• NOTES:** The peterostigma and associated venation are shifted very distally in extant Vespidae and convergently in *Bembix* (Bembicidae). Among fossil Vespidae, scored as *TRUE* for †*Curiosivespa* and †*Priorparagia*, *FALSE* for †*Protovespa*, and uncertain ("?") for †*Priorvespa* and ?*Symmorphus senex* due to incomplete preservation of their wings.
461 *DC: Costal vein (C). DS: Expression.* • **Observational criterion:** C present (*FALSE* = C incomplete to absent). **• NOTES:** The forewing costal vein is rarely lost, and among sampled taxa was scored as *FALSE* for *Loboscelidia* (Chrysididae), and some Formicidae (*e.g.*, *Opamyrma*, *Lioponera*, *Myrmelachista*). This state, then, has no "grouping information" for the present matrix, but is "clocklike" in transition. See also Brothers & Carpenter (1993, character 92 of Appendix VI, states 1 and 2).
462 *DC: C and SC+R+Rs. DS: Inter-vein distance, i.e., costal cell width.* • **Observational criterion:** These abscissae (= vein “segments”) contacting all or almost all of their length, thus costal cell absent or only present apically (*FALSE* = these veins/abscissae separated, thus costal cell present, long). **• NOTES:** This condition is observed in some Bethylidae and diagnostic of alate Rhopalosomatidae. Sc+R+Rs is the second longitudinal vein posterior to C, and which distally splits into the Radius and Radial Sector; the subcostal vein (Sc) is not expressed in Aculeata, Evanioidea, Trigonaloidea, or †Bethylonymoidea.
463 *DC: Radial vein (R) distad pterostigma. DS: Expression, sclerotization.* • **Observational criterion:** R present and tubular distal to pterostigma for any length (*FALSE* = this abscissa nebulous, spectral, OR absent). **• NOTES:** Loss of R distal to the pterostigma was observed in some Bethylidae, some fossil Scolebythidae, some Chrysididae, †*Falsiformica*, few Dryinidae, *Tiphia* (Tiphiidae), Bradynobaenidae, and some Formicidae (*Opamyrma*, *Anomalomyrma*, *Crematogaster*).
464 *DC: First radial-radial sector crossvein (1r-rs). DS: Expression.* • **Observational criterion:** 1r-rs present at least as a stub, with any orientation (*FALSE* = 1r-rs absent). **• NOTES:** 1r-rs is the crossvein which has an anterior juncture at about the pterostigmal break (*i.e.*, at the proximalmost point of the pterostigma), and a posterior juncture with the radial sector after Rs+M. Although 1r-rs is a symplesiomorphy for the Hymenoptera, among sampled taxa, it occurs in about half of the sampled †Ephialtitoidea, only two of the sampled †Praeaulacidae, one third of the †Anomopterellidae, none of the †Baissidae, none of the Aulacidae, none of the Gasteruptiidae, neither of the †Othniodellithidae, none of the Evaniidae, none of the †Bethylonymidae, none of the Trigonaloidea, and is only present in some Aculeata (†*Thanatotiphia*, *Aelurus*, †*Protoscolia,* †*Sinoproscolia*, *Proscolia*, some †Angarosphecidae, one †*Cretampulex*, one †*Camelomecia*, some “†Armaniidae”, and some *Myrmecia*). Based on this distribution, it is reasonable to infer loss in the ancestor of the extant Evanioidea, all Trigonaloidea (whether plesiomorphic or apomorphic), and of the Aculeata, with several secondary gains. Because informative variation of the orientation and completeness of 1r-rs was observed, four additive characters were scored.
465 (**Additive**) *DC: 1r-rs. DS: Completeness, orientation.* • **Observational criterion:** This abscissa stub-like, *i.e.*, only contacting one abscissa AND longitudinally oriented (*FALSE* = this abscissa complete, *i.e.*, contacting two abscissae OR not longitudinal; “-” = this abscissa absent). **• NOTES:** This condition is observed in †*Acephialtitia* (†Ephialtitoidea) and two of the three †*Protoscolia* species (the third species was scored as uncertain for both 1r-rs presence and orientation).
466 (**Additive**) *DC: 1r-rs. DS: Completeness, orientation.* • **Observational criterion:** This abscissa complete AND longitudinally oriented (*FALSE* = this abscissa incomplete OR not longitudinal). **• NOTES:** This condition was separated from that of "longitudinal and incomplete" as the pattern of occurrence does not indicate homology. Among sampled taxa, a complete and longitudinal 1r-rs was observed in †*Praeproapocritus*, and some "†*Angarosphex*” (those species formerly classified as †*Cretosphex* and †*Calobaissodes*).
467 (**Additive**) *DC: 1r-rs. DS: Completeness, orientation.* • **Observational criterion:** This abscissa complete AND transversely oriented (*FALSE* = this abscissa incomplete OR not transverse). **• NOTES:** Observed in some †Ephialtitoidea (†*Proapocritus atropus*, †*Proapocritus praecursor*, †*Sinephialtites*, †*Karataus*, †*Kuafua*) and in †*Eubaissodes* ("†Angarosphecidae”). †*Praeproapocritus* scored as "?" for transverse because of their intermediate state with respect to †*Proapocritus*. The species of †*Karataus* scored as *TRUE* were rather marginal.
468 (**Additive**) *DC: 1r-rs. DS: Completeness, curvature.* • **Observational criterion:** This abscissa complete AND curved as follows: Anterior juncture with pterostigma directed at an oblique posteroapical angle for about half the anteroposterior width of the "first submarginal cell" (1rrs/2rrs) then abruptly curving distally, ending in Rsf oriented more-or-less parallel to the proximodistal length of the wing (*FALSE* = 1r-rs incomplete OR not curved as described). **• NOTES:** This unique conformation was observed in †*Sinoproscolia yangshuwanziensis*, a putative scoliid.
469 *DC: Second radial-radial sector crossvein (2r-rs). DS: Location.* • **Observational criterion:** Anterior junction of 2r-rs near proximal apex of pterostigma (*FALSE* = this abscissa situated more distally on main axis of pterostigma). **• NOTES:** 2r-rs is more consistently present than 1r-rs and has an anterior juncture with the pterostigma distal to the pterostigmal break, and a posterior juncture with Rs distal to Rs+M. (1) Two characters addressing the angle at which 2r-rs diverges from the pterostigma were thrown out during the quality control phase. These characters were "forewing crossvein 2r-rs, when distinct from Rs, with posterior apex directed more-or-less distally, especially when elongate" and "forewing crossvein 2r-rs perpendicular or nearly perpendicular to long axis of wing". (2) Unlike the angle of 2r-rs, consistently scoring the location of 2r-rs was much easier. The proximal condition (this character) was observed in †*Othniodellitha* (†Othniodellithidae), most extant but few extinct Evaniidae, some Trigonalidae, some †Maimetshidae (†*Zorophratra*), some Scolebythidae, and some Formicidae (*Apomyrma*, *Pogonomyrmex*). (3) There were many states observed that were intermediate between the "proximalmost" and "distalmost" conditions here scored; as for 2r-rs angle, future study would benefit from quantification of 2r-rs angle.
470 *DC: 2r-rs. DS: Location.* • **Observational criterion:** Anterior junction of 2r-rs at distal apex of pterostigma (*FALSE* = this abscissa situated more proximally on main axis of pterostigma). **• NOTES:** Initially included due to description of this condition as a synapomorphy of Scoliidae by Rasnitsyn (1993). Unexpectedly, many Mesozoic Evaniidae had distal 2r-rs, whereas most extant Evaniidae have this crossvein in the proximal position. Some difficulty in consistent scoring is due to variation in form of the pterostigma, and pterostigmal variation was clearly informative but difficult to evaluate discretely.
471 *DC: Rf distad pterostigma and Rsf distad 2r-rs. DS: Distal connection.* • **Observational criterion:** Rs not reaching R or wing margin, *i.e.*, "marginal cell 1" (= 2rrs, or second radial-radial sector cell) distally open (*FALSE* = this cell distally closed). **• NOTES:** The first radial-radial sector cell (1rrs, or “submarginal cell 1”) is proximal to 2r-rs, while the second is distal to that crossvein; 1rrs is subdivided when 1r-rs is present. An open "marginal cell" was observed sporadically among sampled taxa, including some †Praeaulacidae (†*Evanigaster*, several †Cretocleistogastrinae), some Evaniidae, most Chrysidoidea, Dryinidae, †*Falsiformica* (†Falsiformicidae), *Tiphia* (Tiphiidae), and some Formicidae (*Anomalomyrma*, *Tatuidris*, *Proceratium*, several Myrmicinae).
472 *DC: Rf distad pterostigma and Rsf distad 2r-rs. DS: Relative area of enclosure.* • **Observational criterion:** "Marginal cell 1" (= 2rrs) small, approximately equal in size to pterostigma or smaller (*FALSE* = this cell larger than pterostigma). **• NOTES:** This condition certainly captures homoplastic conditions. However, because homologies between various fossil terminals and extant terminals are uncertain, these are lumped as this single aggregate character. Among sampled taxa, small "marginal cells" were observed in some Thynnoidea (*Brachycistis*, *Chyphotes*, CASENT084450), most Scoliidae (except *Trisciloa* and *Campsomeris*), some Apoidea (†*Apodolichurus*, †*Burmastatus*, *Heterogyna*, Ammoplanidae, *Perdita*), some †*Camelomecia*, and some Formicidae (*e.g.*, *Stegomyrmex*).
473 *DC: Rf distad pterostigma, Rsf distad 2r-rs, and 2r-rs. DS: Curvature, lengths.* • **Observational criterion:** "Marginal cell 1" (= 2rrs) triangular AND with more-or-less straight sides AND hypotenuse along anterior margin (*FALSE* = this cell not triangular OR Rf, Rsf, or 2r-rs distinctly curved, excepting the apicalmost length of Rsf OR longest abscissa either Rsf or 2r-rs). **• NOTES:** This particular condition was observed in *Sclerogibba* but not †*Sclerogibba cretacica* (Sclerogibbidae). The forewing of †*Sclerogibbodes* was not illustrated.
474 *DC: Rsf distad 2r-rs. DS: Curvature.* • **Observational criterion:** This abscissa of Rsf curving posteriorly, *i.e.*, "marginal cell 1" (= 2rrs) apically hooked, regardless of whether cell closed (*FALSE* = this abscissa of Rsf not curving posteriorly, being either linear or curving anteriorly). **• NOTES:** The hooked condition was observed sporadically among sampled taxa, including *Plumarius* (Plumariidae), most Chrysididae (except *Loboscelidia*, †*Burmasega*, †*Miracorium*), and several Dryinidae.
475 *DC: Distal juncture of Rf distad pterostigma and Rsf distad 2r-rs. DS: Angle.* • **Observational criterion:** This juncture acute, *i.e.*, "marginal cell 1" (= 2rrs) apex pointed to narrowly rounded, with (*FALSE* = juncture of Rf and Rsf broadly rounded or posteriorly downcurved OR this juncture not formed). **• NOTES:** During the quality control phase, it was determined that the scoring of "marginal cell 1 apex acute and pointed" versus "MC1 apex acute and rounded" was inconsistent, thus these two characters were aggregated. Extant Evaniidae were scored as *FALSE* because of the deep angle at which Rsf meets Rf (nearly perpendicular). In Scoliidae, various fossil species have a clear acute angle where Rsf and Rf meet, whereas these veins meet obtusely in some extinct and all extant species. The minority of terminals were *FALSE*. Some taxa were scored for multiple shape conditions of the "marginal cell" apex.
476 *DC: Distal juncture of Rf distad pterostigma and Rsf distad 2r-rs. DS: Angle, curvature.* • **Observational criterion:** Rsf bent or curving widely anterad to join Rf, *i.e.*, "marginal cell 1" (= 2rrs) apex broadly rounded to nearly perpendicular at its apex (*FALSE* = Rsf not bent anterad, *i.e.*, this cell narrowly rounded apically, posteriorly “downcurved”, OR open). **• NOTES:** Observed sporadically, including in extant Evaniidae (NOT in Mesozoic Evaniidae; apparent transitional states observed in †*Curtevania*, †*Grimaldivania*, †*Iberoevania*), *Myzinum* (Tiphiidae), Pompilidae, *Fedtschenkia* (Sapygidae), *Dasymutilla* (Mutillidae), *Dieunomia* (Halictidae), and some Formicidae (*Martialis*, *Discothyrea*, *Lepisiota*). The condition is most pronounced in Evaniidae. Due to the phylogenetic distribution of this state, the additional scores are likely homoplastic.
477 *DC: Rf distad pterostigma. DS: Proximity to anterior wing margin.* • **Observational criterion:** Rf separating narrowly from anterior wing margin at apex of cell to meet Rsf, which is uncurved, *i.e.*, "marginal cell 1" (= 2rrs) apex slightly "downcurved" (*FALSE* = Rf continuous with anterior wing margin to its distal juncture with Rsf, *i.e.*, apex of MC1/2rrs not downcurved). **• NOTES:** This condition observed in extant Scolebythidae, some Vespidae (including Mesozoic terminals), †*Cretofedtschenkia*, some "†Angarosphecidae", some †*Psolimena* (Pemphredonidae), some Philanthidae, and some Anthophila.
478 *DC: Rsf distad pterostigma. DS: Curvature.* • **Observational criterion:** This abscissa narrowly and abruptly curving anteriorly to its juncture with Rf, *i.e.*, "marginal cell 1" (= 2rrs) apex truncate or oblique, with distalmost point formed by Rsf curve (*FALSE* = Rsf broadly or weakly curving anteriorly OR not curving anteriorly, *i.e.*, MC1/2rrs apex not truncate or oblique OR if oblique then distalmost point formed by juncture of Rf and Rsf OR this cell open). **• NOTES:** Observed in *Lytopsenella* (Bethylidae), some Thynnoidea (Tiphiidae, *Chyphotes*), *Smicromyrmilla* (Mutillidae), all extant Scoliidae, some Apoidea (*Heterogyna*, *Bembix*). Among Mesozoic "Scoliidae", this strong downcurvature was observed in †*Archaeoscolia senilis* and most †*Cretoscolia*; †*Protoscolia* had rounded but not downcurved "marginal cell" apices; †*Sinoproscolia* was *FALSE*; †*Archaeoscolia hispanica* and †*Cretoscolia brasiliensis* were scored as uncertain "?".
479 (**Reductive**) *DC: Rf distad pterostigma. DS: Separation from anterior wing margin.* • **Observational criterion:** This abscissa distinctly separated from anterior wing margin, not just at its tip or by a narrow section of nearly indiscernible membrane (*FALSE* = this abscissa continuous with anterior wing margin OR only separated near its apex; "-" = inapplicable due to absence of Rf distal to pterostigma). **• NOTES:** During the quality control phase a number of marginal cases were rescored from *TRUE* to *FALSE*, leaving very few terminals with the "1" token. Among Formicidae, a distinct, narrow strip of membrane anterior to the distal abscissa of Rf is often visible, but this was considered *FALSE* for separation of Rf in the present sense as Rf does not distinctly curve away from the costal margin being, rather, parallel with it until the very apex of the "marginal" cell. The *TRUE* state is retained in the current matrix for †*Curiosivespa magna* (Vespidae), †Pompiloidea sp. (CASENT0844580, burmite), *Brachycistis* (Tiphiidae), †*Cretoscolia promissiva* (Scoliidae), *Ampulex* (Ampulicidae), and *Dolichoderus* (Formicidae).
480 *DC: Rsf distad rs-m. DS: Curvature.* • **Observational criterion:** Rsf gently distinctly curving anterodistally from distalmost rs-m crossvein at about 45° angle, *i.e.*, "marginal cell 1" (= 2rrs) appearing roughly arrowhead-shaped (*FALSE* = this abscissa without this specific curvature, *i.e.*, Rsf not bent at an ∼45° angle at rs-m). **• NOTES:** Observed nearly uniquely in †Baissidae (†*Humiryssus*, some †*Manlaya*). The condition is similar to that observed in extant Evaniidae but is scored as an independent character as Rsf extends beyond the distal rs-m crossvein before curving in extant Evaniidae, and immediately "kinks" anteriorly in the baissids.
481 *DC: First free abscissa of Rs after branching of R+Rs (= Rsf1). DS: Expression.* • **Observational criterion:** This abscissa present (*FALSE* = Rsf1 absent). **• NOTES:** Rsf1 was observed to be absent (*FALSE*) in a minority of taxa, including *Hyptia* (Evaniidae), *Loboscelidia*, †*Thanatotiphia*, Bradynobaenidae *sensu stricto*, and some Formicidae (*Opamyrma*, *Anomalomyrma*, and *Discothyrea*).
482 *DC: Rsf1 proximal juncture with Rf. DS: Position relative to pterostigma.* • **Observational criterion:** This juncture contacting or very near to pterostigma, *i.e.*, first free abscissa of R (Rf1) length between Rsf1 and pterostigma ≤ 2 x width (*FALSE* = this juncture distant from pterostigma, *i.e.*, Rf1 length > 2 x width OR Rf and/or Rsf1 absent). **• NOTES:** The location of Rsf1 relative to the pterostigma appears to be quite informative across the range of positions, from very distant through contiguous. However, during character construction, only the contiguous state had been noted. Some cases were difficult to evaluate because of the angle at which Rsf diverged from R+Rs, which is to say that in a minority of sampled taxa, Rs did contact the pterostigma distally, but was contiguous with R+Rs proximal to its split. Other difficult cases related to thickening to the juncture of R and the pterostigma. Among sampled taxa, this condition was observed in some †Ephialtitoidea, some †Praeaulacidae, †*Mesepipolaea* (†Baissidae), *Gasteruption* (Gasteruptiidae), many Evaniidae (including some extinct taxa), most †Bethylonymidae, some Bethylidae, most Chrysididae, some Dryinidae, most Vespoidea *sensu stricto*, some †*Mesorhopalosoma* ("†Angarosphecidae"), †*Rasnitsynapus*, some †*Psolimena* (Pemphredonidae), Ammoplanidae, some Anthophila (including †*Cretotrigona*). Among evaniomorphs, Rsf abutting or nearly abutting the pterostigma was used by Li et al. (2013) to transfer †*Anomopterella stenocera* to †*Choristopterella*.
483 (**Reductive**) *DC: Juncture of first free abscissae of Rs and M (Rsf1 and Mf1). DS: Angle.* • **Observational criterion:** Rsf1 and Mf1 meeting at a distinct oblique angle AND the vertex of this angle directed distally (*FALSE* = Rsf1 and Mf1 more-or-less parallel, *i.e.*, forming a single straight line between anterior and posterior junctures of each abscissa OR apex of angle directed proximally; "-" = inapplicable due to absence of Rsf1). **• NOTES:** This is the most frequent condition for the conformation of Rsf1 and Mf1, suggesting that this may be the ancestral condition of at least the Aculeata. Among symphytan Hymenoptera, this condition was observed in Cephidae, Blasticomidae, Xyelidae, Anaxyelidae, and Xiphydriidae. Within the Aculeata, the *FALSE* state generally describes the linear condition of these abscissae. *TRUE* was scored for many †Ephialtitoidea, some †Praeaulacidae, †*Humiryssus specialis* (†Baissidae), †*Othniodellitha* (†Othniodellithidae), most Evaniidae, some †Bethylonymidae, most Trigonaloidea, Plumariidae, most Scolebythidae (all extant, most extinct), †Plumalexiidae, some Bethylidae, most Chrysididae, †Aculeata sp. (CASENT0844587, burmite), some †*Falsiformica*, some Dryinidae, some Embolemidae, *Sclerogibba*, some Vespidae, †Pompiloidea sp. (CASENT0844580, burmite), some Thynnoidea, some Mutillidae, some Scoliidae (a minority of fossil taxa), Ampulicidae, most Sphecidae, most Crabronidae, Ammoplanidae, Pemphredonidae, Philanthidae, most Anthophila, and most Formicidae (except *Apomyrma*, and some Formicinae). Notably, in contrast with the majority of sampled extant Apoidea, the majority of "†Angarosphecidae" were observed to have the linear (*FALSE*) state.
484 (**Reductive**) *DC: Juncture of Rsf1 and Mf1. DS: Angle.* • **Observational criterion:** This juncture distally directed AND acute, *i.e.*, with an inner angle which is ≤ 45° (*FALSE* = this juncture proximally directed OR more oblique; "-" = inapplicable due to absence of Rsf1). **• NOTES:** This extreme condition was scored as *TRUE* for some †Ephialtitidae (†*Proapocritus*, †*Stephanogaster*, *†Thilopterus,* and †*Brigettepterus*).
485 (**Reductive**) *DC: Juncture of Rsf1 and Mf1. DS: Angle.* • **Observational criterion:** Rsf1 and Mf1 meeting at an oblique angle, with the vertex of the angle directed proximally (*FALSE* = Rsf1 and Mf1 more-or-less parallel, *i.e.*, forming a single straight line, between anterior and posterior junctures of each abscissa, OR forming angle with apex directed distally; "-" = inapplicable due to absence of Rsf1). **• NOTES:** The proximally directed angle of Rsf1 and Mf1 was observed in †Cretocleistogastrinae (†Ephialtitidae), †Anomopterellidae, and most †*Manlaya* (†Baissidae).
486 (**Reductive**) *DC: Rsf1 and Mf1. DS: Relative lengths.* • **Observational criterion:** Mf1 long to extremely long, being ≥ 4 x length of Rsf1 (*FALSE* = Mf1 shorter, being < 4 x Rsf1 length; "-" = inapplicable due to absence of Rsf1). **• NOTES:** This character was initially included to capture the condition of most Vespidae (*FALSE* in †*Protovespa haxairei*) but was found to be *TRUE* for numerous taxa which have very distinct venational conformations because of variability in the relative lengths of the Rsf1 and Mf1. As scored, *TRUE* was recorded for extant Evaniidae, †*Newjersevania* (Evaniidae), most Chrysididae, most Vespoidea *sensu stricto*, *Methoca* (Thynnidae), some Sphecidae, some Crabronidae, Bembicidae, some Anthophila (including †*Cretotrigona*), some Formicidae (†*Baikuris mandibularis*, †*Dlusskyidris*, *Stigmatomma*).
487 (**Reductive**) *DC: Rsf1 and Mf1. DS: Expression, conformation.* • **Observational criterion:** Mf1 absent AND Rsf1 (= "1Rs") directly or nearly directly attached to M+Cu *and* Rs+M more-or-less continuous with M+Cu (*FALSE* = Mf1 present OR Mf1 absent *and* Rsf1 not directly attached to M+CU thus Rs+M not continuous with M+Cu; “-” = both abscissae absent). **• NOTES:** This exceptional condition was observed in †*Synaphopterella* (†Anomopterellidae), some Aulacidae (†*Archaeofoenus*, †*Electrofoenus*), and *Gasteruption* (Gasteruptiidae).
488 (**Reductive**) *DC: Mf1. DS: Curvature.* • **Observational criterion:** This abscissa distinctly curved, with the vertex of the curve directed proximally rather than distally (*FALSE* = Mf1 more-or-less linear OR curved such that vertex of curve directed distally; “-” = Mf1 not developed). **• NOTES:** Initially characterized based on the traditional diagnostic condition of Halictidae (Anthophila), but as scoring proceeded it became apparent that this condition is widespread. Here scored as *TRUE* for some †Ephialtitoidea, some †Praeaulacidae, some †Baissidae, some Evaniidae, many †Bethylonymidae, most Trigonaloidea, Plumariidae, some Sclerogibbidae, some Bethylidae, some Chrysididae, some Dryinidae, most †Falsiformicidae (including †*Burmasphex*), Sierolomorpha, most Thynnoidea, most Scoliidae (except *Proscolia*, but including many extinct species), some "†Angarosphecidae", Ampulicidae, Ammoplanidae, Pemphredonidae, some Anthophila in addition to Halictidae, some †*Camelomecia*, some “†Armaniidae” (†*Armania*, †*Poneropterus*), and some Formicidae.
489 (**Reductive**) *DC: Mf1. DS: Curvature and proximity to Sc+R+Rs.* • **Observational criterion:** This abscissa strongly curved and very closely approximating Sc+R+Rs, such that first sectoral cell (1rsm) (= "basal cell", BC) long and very narrow distad M+Cu split (*FALSE* = Mf1 not strongly curved OR if strongly curved then not closely approaching Sc+R+Rs, *i.e.*, 1rsm/BC shorter and/or broader distad M+Cu split; “-” = Mf1 not developed). **• NOTES:** This is an extreme condition of Mf1 curvature and was uniquely observed in Evaniidae.
490 *DC: Medial-cubital vein (M+Cu). DS: Expression.* • **Observational criterion:** This abscissa present (*FALSE* = M+Cu absent). **• NOTES:** M+Cu is the third longitudinal vein posterior to the costal margin of the wing, after C and Sc+R+Rs, and it is proximal to the split of free M (Mf) and free Cu (Cuf). Absence is the exceptional state and was observed in †*Kotephialtites* (†Ephialtitoidea), one species of †*Anomopterellidae*, †*Hyptiogastrites* (Aulacidae), some *Loboscelidia* (Chrysididae), and †*Burmadryinus cenomanianus* (Dryinidae). The character does not have any grouping information, but this is an apomorphy which has evolved independently several times, thus presumably contributing "morphological clock" information. This condition was previously noted by Olmi *et al*. (2014).
491 *DC: Juncture of M+Cu and cubitoanal crossvein (cu-a). DS: Angle.* • **Observational criterion:** This juncture approaching 90°, *i.e.*, M+Cu diverging strongly from cu-a (*FALSE* = angle widely oblique or simply linear). **• NOTES:** This condition is approximated in a number of taxa but was scored *TRUE* for the extreme condition observed in †Maimetshidae, †*Cretogonalys* (Trigonalidae), some Bethylidae, †*Burmasphex*, †*Falsiformica*, some Pemphredonidae, and *Proceratium* (Formicidae).
492 *DC: Radial-sector-media compound abscissa (Rs+M). DS: Expression, sclerotization.* • **Observational criterion:** Rs+M present AND tubular (*FALSE* = this abscissa absent OR nebulous to spectral). **• NOTES:** Rs+M is that abscissa between the distal juncture of Rsf1 and Mf1 and the proximal split of Rsf2 and Mf2; it is a defining feature of Hymenoptera (Ross 1936). Rs+M is sporadically absent among sampled taxa, including †*Tillywhimia* (†Baissidae), †*Hyptia* (Evaniidae), some Scolebythidae, some Bethylidae, most Chrysididae, Dryinidae, †*Falsiformica*, Bradynobaenidae, †*Apodolichurus*, *Oxybelus* (Crabronidae), some Formicidae (*e.g.*, *Opamyrma*, *Anomalomyrma*, *Leptomyrmex*, *Crematogaster*).
493 *DC: Radial sector between first mediocubital crossvein (1m-cu) and 2rs-m (i.e., Rsf2–3). DS: Expression.* • **Observational criterion:** Rsf2–3 absent; consequently, first medial- cubital crossvein (1m-cu) apparently joining Rs+M and the second medial-cubital crossvein (2m-cu), when developed, apparently joining the “second submarginal cell” (SMC2/3rsm) (*FALSE* = Rsf2–3 present; 1m-cu joining Rs+M or not; 2m-cu, when developed, joining SMC2/3rsm or not). **• NOTES:** 1m-cu and 2m-cu are the proximal and distal crossveins joining free M and free Cu distal to M+Cu, respectively; SMC2 is the second cell posterior to SMC1 (= 1rrs) and the second cell distal to 1rm (the cell between Sc+R+Rs and M+Cu). There are several instances of loss of Rsf between Rs+M and 2r-rs, very notably in the Formicidae where a clear transition series may be demonstrated in the Myrmicinae (e.g., Bolton 1982). This condition was observed in extant Scolebythidae (including †*Boreobythus*, †*Libanobythus*), †Chrysobythidae, †Aculeata sp. (CASENT0844587, burmite), some Embolemidae, *Sierolomorpha*, some Thynnoidea, some Crabronidae, some Pemphredonidae, and †*Cretotrigona*. Among sampled Formicidae, this state was observed in *Amblyopone*, *Lioponera*, and *Manica*.
494 (**Reductive**) *DC: Rsf2–3. DS: Curvature.* • **Observational criterion:** Rsf2–3 (= "2Rs" of Engel 2017) kinked or distinctly curved, with vertex of curve directed anteriorly, whether or not 1r-rs present or absent; the curve vertex receives or would receive 1r-rs, pending presence of this crossvein (*FALSE* = Rsf2–3 linear OR curved such that vertex directed posteriorly; "-" = inapplicable due to absence of Rsf between Rs+M and 2r-rs). **• NOTES:** Bending or kinking of Rsf observed sporadically, including in most †Ephialtitoidea, most †Praeaulacidae, most †Anomopterellidae, some †Baissidae, most †Bethylonymidae, some †Maimetshidae, Trigonalidae, Plumalexiidae, †*Burmasphex*, Rhopalosomatidae, some Vespidae, †*Loreisomorpha*, †*Thanatotiphia*, some †Burmusculidae, †*Cretosapyga*, Thynnoidea (where this abscissa present), most Mutillidae, *Proscolia* (Scoliidae), some Mesozoic Scoliidae, many "†Angarosphecidae", *Dolichurus* (Ampulicidae), *Bembix* (Bembicidae), Philanthidae, some Anthophila, †*Camelomecia*, “†Armaniidae”, and some Formicidae.
495 (**Reductive**) *DC: Rsf2–3. DS: Orientation.* • **Observational criterion:** This abscissa perpendicular or nearly perpendicular to longitudinal axis of forewing OR obliquely diverging posteriorly (*FALSE* = Rsf2–3 obliquely diverging anteriorly; "-" = inapplicable due to absence of Rsf2–3). **• NOTES:** This condition was observed in some Evaniidae (†*Grimaldivania*, †*Sinuevania*), †*Mirabythus* (Scolebythidae), *Trypoxylon* (Crabronidae), sampled Bembicidae, some Anthophila, *Proceratium* (Formicidae).
496 (**Additive-reductive**) *DC: Rs+M proximal and distal abscissae. DS: Relative lengths.* • **Observational criterion:** Rs+M with proximal and distal abscissae, *i.e.*, 1m-cu joining Rs+M rather than Rsf AND distal abscissa longer than proximal abscissa (*FALSE* = these abscissae subequal in length or proximal abscissa longer; "-" = inapplicable as Rs+M absent OR not divided into proximal and distal abscissae by 1m-cu). **• NOTES:** A second abscissa of Rs+M is observable when 1m-cu is "antefurcal", *i.e*., when 1m-cu intersects Rs+M prior to the splitting of Rs and M. Therefore, this condition is additive on the "antefurcal" condition of 1m-cu. The elongate condition of the second abscissa of Rs+M was observed sporadically, including in several extinct Aulacidae, some extinct Evaniidae, Gasteruption (Gasteruptiidae), various †Maimetshidae, †*Cretogonalys* (Trigonalidae), †*Eubaissodes* ("†Angarosphecidae"), †*Prolemistus* (Pemphredonidae), †*Camelomecia* sp. (ANTWEB1038930), and some Formicidae (*Lasius* and *Formica*, Formicinae).
497 (**Reductive**) *DC: Rs+M and Mf2, the first free abscissa of M between Rs+M and 1m-cu. DS: Relative lengths.* • **Observational criterion:** Mf2, if present, as long as or longer than Rs+M (*FALSE* = Mf2 shorter than Rs+M; “-” = inapplicable due to absence of Mf2). **• NOTES:** Mf2 cannot be evaluated when 1m-cu is "antefurcal". The *TRUE* condition was scored for some †Praeaulacidae, *Gasteruption* (Gasteruptiidae), †*Maimetsha arctica* (†Maimetshidae), †*Thanatotiphia*, Scoliinae (Scoliidae), many Mesozoic fossils attributed to Scoliidae, some "†Angarosphecidae", and some Formicidae (*Paraponera*).
498 (**Reductive**) *DC: Rs+M and Mf3, the second free abscissa of M after Rs+M, distal to juncture of 1m-cu with M, and proximal to juncture of 2rs-m. DS: Relative lengths.* • **Observational criterion:** Mf3, when developed, as long as or longer than Rs+M (*FALSE* = Mf3 shorter than Rs+M; “-” = inapplicable due to absence of Mf3). **• NOTES:** Mf3 can be evaluated when 1m-cu is "antefurcal". Scored as *TRUE* for various †Ephialtitoidea, most †Praeaulacidae, †Anomopterellidae, †Baissidae, most extinct Aulacidae, *Pristaulacus* (Aulacidae), most extinct Evaniidae, most †Bethylonymidae, †*Mirabythus* (Scolebythidae), †*Burmasphex*, *Pepsis* (Pompilidae), some Scoliidae, †*Pompilopterus worssami*, †*Gallosphex*, *Heterogyna*, †*Camelomecia* sp. (ANTWEB1038930), some “†Armaniidae”, and most Formicidae when present.
499 (**Reductive**) *DC: Split of Rs+M and 2r-rs. DS: Relative location.* • **Observational criterion:** Rs+M splitting at or distad 2r-rs (*FALSE* = this split proximad 2r-rs; “-” = split absent OR insufficient comparative information available to determine precise identity of distalmost abscissa of M). **• NOTES:** In Formicinae (Formicidae), *Myrmelachista* has M splitting from Rs+M proximal to 2r-rs with loss of the rs-m crossveins; free M in other Formicinae is migrated distally, such that its anterior juncture is at or very near 2rs-m. With *Myrmelachista* representing an intermediate state in the subfamily (indeed, the Myrmelachistini is sister to the rest of the Formicinae), most Formicinae are scored as *TRUE* for the present character. Dolichoderinae without a "second submarginal cell" scored as "-" as no *Myrmelachista*-like intermediates have been observed in the subfamily. In dolichoderine taxa with the second section of free M [Mf2+] diverging at or distal to 2rs-m, it is possible that Mf2 between 1m-cu and 2rs-m has been lost with retention of 2rs-m+Mf; see note for "1m-cu antefurcal", below.
500 (**Additive-reductive**) *DC: Split of Rs+M and 2r-rs. DS: Relative location.* • **Observational criterion:** Rs+M splitting distad 2r-rs (*FALSE* = this split at or proximad 2r-rs OR absent; “-” = split absent OR insufficient comparative information available to determine precise identity of distalmost abscissa of M). **NOTE:** This extreme condition observed in a subset of Formicinae (*Gigantiops*, *Gesomyrmex*, *Anoplolepis*, and *Formica*).
501 *DC: M distad Rs+M (= Mf2+ or Mf3+). DS: Sclerotization.* • **Observational criterion:** This abscissa tubular (*FALSE* = this abscissa nebulous, spectral, OR absent). **• NOTES:** During the character development phase, Mf2+ or Mf3+ NOT tubular was treated as a distinct character; during quality control, the number of exceptions was limited, thus the information appeared to be duplicate. The initial segregation of these states was intended to separate extant Evaniidae from many extinct terminals. Among sampled taxa, the *FALSE* state was observed for some Evaniidae (extant taxa, †*Curtevania*, †*Grimaldivania*, †*Iberoevania*, †*Newjersevania,* †*Protoparevania*, †*Sinuevania*), *Orthogonalys* (Trigonalidae), most Chrysidoidea (except Plumariidae, †Plumalexiidae, most Scolebythidae, most Chrysididae), Embolemidae, CASENT0844587, †*Falsiformica*, Dryinidae, Sclerogibbidae, Bradynobaenidae, *Apodolichurus* (Ampulicidae), *Ammoplanops* (Ammoplanidae), and some Formicidae (*Martialis*, *Opamyrma*, *Anomalomyrma*, *Leptomyrmex*, *Wasmannia*, *Crematogaster*). Notably †Aculeata sp. (CASENT0844587, burmite) has most tubular veins *except* for Mf.
502 *DC: Forewing crease network. DS: Expression.* • **Observational criterion:** Distal half of the forewing with 2–4 longitudinal creases between the longitudinal veins; these lines may be tubular to spectral veins or simply creases and include longitudinal and transverse lines; they are labeled “adventitious veins” or “adventitious creases” in the next character definitions for ease of reference (*FALSE* = these creases absent). **• NOTES:** These specific additional veins, creases, or flexion lines were observed in all extant Chrysidoidea and Embolemidae. Extinct chrysidoids which LACK these lines include †*Necrobythus*, †*Plumalexius*, and †*Aureobythus*. The following eight characters address particular conformations of these lines. All characters describing variation of these lines are categorically scored as uncertain ("?") for compression fossils, except for those putative "Scoliidae" for which the apical wrinkles of the forewing were preserved. Two extremely strange and possibly erroneous compression fossils appear to have additional veins: †*Cretavus sibiricus* Sharov, 1957 and †*Limnetus wangyingziensis* Hong, 1983. While the illustration of latter may be a misinterpretation based on the photographic plate included in the original description, the former cannot be reevaluated in present study, thus is excluded. However, it is possible that this fossil represents a chrysidoid, but this question is left for future study. Similar creases are observable in Ichneumonoidea; for this reason, character polarity is *a priori* unclear.
503 *DC: Anteriormost adventitious vein. DS: Proximal articulation.* • **Observational criterion:** This line joined proximally to Rsf, or nearly so (*FALSE* = this crease not joining Rsf proximally OR forewing crease network not developed). **• NOTES:** This line is tubular in *Plumarius* and is very weak in *Clystopsenella* (Scolebythidae), wherein it is not joined to Rs. The line appears to be joined to Rs in *Amisega cooperi* (see, *e.g.*, Krombein 1957), but this taxon was not explicitly evaluated in the present study. Scored as an ancestral state of the Aculeata in Brothers & Carpenter (1993, state 0 of character 201 in Appendix VI), where they considered this the second (posterior) branch of Rs distal to Rs+M.
504 *DC: Second tubular adventitious crossvein. DS: Connections.* • **Observational criterion:** This crossvein, if present, joining 2rs-m (*FALSE* = only one adventitious crossvein present OR if present then not joining 2rs-m OR forewing crease network not developed). **• NOTES:** Observed in *Plumarius*; as defined, also scored as *TRUE* For †*Boreobythus*.
505 *DC: “Crease crossvein”. DS: Expression.* • **Observational criterion:** Adventitious crease system with a transverse crease joining two longitudinal creases, thus forming a "crease crossvein" (*FALSE* = such a crease crossvein not developed OR forewing crease network not developed). **• NOTES:** Observed in Bethylidae and *Ampulicomorpha* (Embolemidae). Additionally, *Ycaploca* (Scolebythidae), appears to have such a crossvein, but it is where the crease crossvein is more proximal than in Bethylidae, being just distal to 1m-cu rather than just posterior to Rsf.
506 *DC: Anteriormost transverse adventitious crease. DS: Connections.* • **Observational criterion:** This crease anteriorly joining a longitudinal adventitious crease (*FALSE* = this crease not joining a longitudinal crease OR forewing crease network not developed). **• NOTES:** Observed in *Lytopsenella* (Bethylidae) and *Ycaploca* (Scolebythidae).
507 *DC: Anteriormost transverse adventitious crease. DS: Connections.* • **Observational criterion:** This crease anteriorly joining Rsf (*FALSE* = this crease not joining Rsf OR forewing crease network not developed). **• NOTES:** Among sampled Bethylidae, observed in *Epyris* and *Goniozus*.
508 *DC: Longitudinal adventitious crease located in expected position of Mf2+. DS: Split, split location.* • **Observational criterion:** This crease with two branches at its distal end (*FALSE* = such a crease not present OR forewing crease network not developed). **• NOTES:** Observed in *Sclerogibba* (Sclerogibbidae), *Pristocera* (Bethylidae) and Chrysididae, the latter with a distinct conformation scored as a separate character.
509 *DC: Longitudinal adventitious crease located in expected position of Mf2+. DS: Split, split location.* • **Observational criterion:** This crease with two branches at its proximal end (*FALSE* = such a crease not present OR forewing crease network not developed). **• NOTES:** Among sampled Bethylidae, observed in *Epyris*, *Goniozus*, and *Lytopsenella*; in the latter, these branches cross a tubular vein, which is either Rsf2+ or the second abscissa of Rs+M.
510 *DC: Crease located at pterostigmal break. DS: Extent, form.* • **Observational criterion:** This crease extending posteroapically to Mf and, where the two meet, the crease divides into two longitudinal creases surrounding the spectral Mf (*FALSE* = such a crease not developed OR not dividing in this way OR forewing crease network not developed). **• NOTES:** Observed uniquely in Chrysididae.
511 *DC: Forewing crossvein 2rs-m. DS: Form.* • **Observational criterion:** This crossvein tubular and distinct from free M (*FALSE* = 2rs-m nebulous, spectral, absent). **• NOTES:** 2rs-m is the proximalmost crossvein joining the free Radial sector (Rsf) and free Media (Mf) distal to Rs+M; the first radial sector-medial crossvein (1rs-m) is traditionally treated as obliterated by the fusion of Rs+M in the Comstock-Needham-Ross system has been lost a number of times across the Hymenoptera. Scored as *FALSE* for †Cretocleistogastrinae (†Ephialtitidae), †Anomopterellidae, most †Baissidae, most Aulacidae, most Evaniidae, Gasteruptiidae, some †Maimetshidae, most Chrysidoidea (except Plumariidae, †Plumalexiidae, some Bethylidae), extant Sclerogibbidae, Dryinoidea, †*Falsiformica*, some Scoliidae (Campsomerini, *Scolia*), Bradynobaenidae, *Apodolichurus* (Ampulicidae), *Heterogyna* (Heterogynaidae), *Oxybelus* (Crabronidae), *Ammoplanops* (Ammoplanidae), †*Cretotrigona*, and various Formicidae (Leptanillinae *sensu lato*, *Tatuidris*, Proceratiinae, *Lioponera*, some Dolichoderinae, all Formicinae, and some Myrmicinae). In Anomopterellidae, 2rs-m is described as absent, and based on comparison across the sampled evaniomorphs, particularly †*Manlaya*, this interpretation is here accepted, although there is some information loss as the relative distance of this crossvein to 2r-rs is variable; †*Cretaproscolia asiatica i*s similarly scored as other Mesozoic "Scoliidae" indicate a transformational loss of 2rs-m. Yamada *et al*. (2020) interpret the abscissa joining the "pterostigmal hook vein" (2r-rs and Rsf4+) as Rsf3.
512 (**Reductive**) *DC: 2rs-m. DS: Curvature.* • **Observational criterion:** This crossvein sinuate, whether tubular, nebulous, or spectral (*FALSE* = 2rs-m linear, regardless of sclerotization; "-" = 2rs-m absent). **• NOTES:** This condition is observed in a minority of taxa, albeit this condition is clearly homoplastic. Scored as *TRUE* for some †Ephialtitoidea, some †Praeaulacidae, some Vespidae, †*Architiphia*, †*Cretofedtschenkia*, †Mutillidae sp. (CASENT0844578), some Thynnoidea (Brachycistis, Chyphotidae), Pompilidae, Scoliidae (including some Mesozoic fossils), various "†Angarosphecidae", *Tachysphex* (Crabronidae), *Sphecius* (Bembicidae), and many Anthophila (including the putative apid from the Crato formation).
513 (**Reductive**) *DC: Juncture between 2rs-m and Rs. DS: Location relative to 2r-rs.* • **Observational criterion:** This juncture distinctly proximad 2r-rs crossvein, resulting in anterior "petiolation" of cell 2rm (*FALSE* = this juncture absent OR juncture close to or distad 2r-rs; "-" = 2rs-m absent). **• NOTES:** State not dependent on development of vein abscissae. Scored as *TRUE* for †*Praeaulacus pachygaster* (†Praeaulacidae), †*Cretevania minor* (Evaniidae), most †Maimetshidae, *Smicromyrmilla* (Mutillidae), *Colpa* (Scoliidae), some fossils attributed to Scoliidae (†*Cretoscolia rasnitsyni*, †*Protoscolia*), †*Ilerdosphex* ("†Angarosphecidae), *Plenoculus* (Crabronidae), Ammoplanidae, *Cerceris* (Philanthidae), the putative Crato apid, †*Camelomecia* sp. (ANTWEB1038930), some Dolichoderinae plus Stegomyrmex (Formicidae). As this is an extreme condition observed in a minority of taxa, the character is not scored additively; however, *FALSE* when any of the following three characters are *TRUE*.
514 (**Reductive**) *DC: Juncture between 2rs-m and Rs. DS: Location relative to 2r-rs.* • **Observational criterion:** This juncture at or very near 2r-rs crossvein (*FALSE* = this juncture distinctly proximad 2r-rs; "-" = 2rs-m absent). **• NOTES:** State not dependent on development of vein. Scored as *TRUE* for some †Ephialtitidae, some †Praeaulacidae, †Othniodellithidae, most †*Cretevania* (Evaniidae), †*Turgonalus* (†Maimetshidae), Trigonalidae, †*Holopsenelliscus* (Bethylidae), *Sclerogibba cretacica* (Sclerogibbidae), †*Protovespa haxairei* (Vespidae), some Scoliidae (*Proscolia*, *Megascolia*), some extinct "Scoliidae", *Heterogyna* (Heterogynaidae), *Manica* (Formicidae). Although the *TRUE* state for the present character is more frequent than that for the prior character, it is much less frequent than the following ("2rs-m distal to 2r-rs"). *FALSE* when at least one of the two following characters is true, and *FALSE* when the prior character is true.
515 (**Additive-reductive**) *DC: Juncture between 2rs-m and Rs. DS: Location relative to 2r- rs.* • **Observational criterion:** This juncture distad 2r-rs by less than one of its lengths (*FALSE* = this juncture at or proximad 2r-rs OR this juncture distad by more than one 2r- rs lengths; "-" = 2rs-m absent). **• NOTES:** State not dependent on development of vein. Scored as *TRUE* for the majority of sampled taxa. The following character is additive on the *TRUE* state of this character. This is the most frequent *TRUE* for the position of 2r-rs among sampled taxa.
516 (**Additive-reductive**) *DC: Juncture between 2rs-m and Rs. DS: Location relative to 2r- rs.* • **Observational criterion:** This juncture distad 2r-rs by one or more than one of its lengths (*FALSE* = this juncture proximad 2r-rs by any lesser distance; "-" = 2rs-m absent). **• NOTES:** State not dependent on development of vein. Additive with respect to the *TRUE* state of the prior character. This extreme condition was observed in some †Ephialtitoidea, †*Archaulacus* (†Praeaulacidae), several †Baissidae, Aulacidae, Gasteruptiidae, extant Evaniidae (among extinct Evaniidae, only observed in †*Mesevania*, †*Newjersevania*, †*Protoparevania*), several †Bethylonymidae, some Vespoidea *sensu stricto*, †*Architiphia*, most †Burmusculidae, †*Cretofedtschenkia*, †Mutillidae sp. (CASENT0844578), *Sierolomorpha*, Thynnidae, Pompilidae, Sapygidae, †*Sinoproscolia* (Scoliidae), various "†Angarosphecidae", some Anthophila, some †*Camelomecia,* and some Formicidae (†*Baikuris maximus*, *Myrmecia*).
517 *DC: Third radial sector-medial crossvein (3rs-m). DS: Sclerotization.* • **Observational criterion:** This crossvein tubular (*FALSE* = 3rs-m nebulous, spectral, or absent). **• NOTES:** 3rs-m is the second crossvein joining the free Radial sector (Rsf) and free Media (Mf) distal to the Rs+M; as noted above (char. 511), a first radial sector-medial crossvein is interpreted as ancestrally obliterated in the Comstock-Needham-Ross system. 3rs-m is homoplastically lost in various groups. Scored as *TRUE* for, i.e., retained as tubular or reversed to presence in most †Ephialtitoidea, †Praeaulacinae (but not †Cretocleistogastrinae, †Praeaulacidae), †Anomopterellidae, few †Baissidae, †Othniodellithidae, †*Mesevania* (Evaniidae), some †Bethylonymidae, some †Maimetshidae, Trigonalidae, †*Mirabythus* (Scolebythidae), †*Burmasphex*, †*Aculeata* sp. (CASENT0844587, burmite), †Vespoidea sp. (CASENT0844576, burmite), most Vespidae, †Sierolomorphidae sp. (CASENT0844589, burmite), †*Loreisomorpha*, †Pompiloidea sp. (CASENT0844580, burmite), †*Architiphia*, †Burmusculidae, †Mutillidae sp. (CASENT0844578), Thynnoidea, Pompilidae, Sapygidae, *Myrmosa* (Mutillidae), Scoliinae (Scoliidae), all fossil "Scoliidae", all "†Angarosphecidae", Ampulicidae, (but not †*Apodolichurus*), Sphecidae, some Crabronidae, Bembicidae, fossil Pemphredonidae, Philanthidae, most Anthophila, and some †*Camelomecia*. Completely absent in “†Armaniidae” and Formicidae.
518 (**Reductive**) *DC: 3rs-m. DS: Curvature.* • **Observational criterion:** This crossvein sinuate, whether tubular, nebulous, or spectral (*FALSE* = 3rs-m linear, regardless of sclerotization; "-" = 3rs-m absent). **• NOTES:** State not dependent on development of vein. Presumably homoplastic across sampled taxa. Scored as *TRUE* for a few †Ephialtitoidea, some †Praeaulacidae, †Othniodellithidae, †*Mesevania* (Evaniidae), †*Afrapia* (†Maimetshidae), one species of †*Burmasphex*, most Vespidae, some Thynnoidea, Pompilidae, †*Bryopompilus*, Sapygidae, some Mutillidae, Scoliinae (Scoliidae), most fossil "Scoliidae", and most Apoidea (most Apidae *FALSE*, several "†Angarosphecidae" *FALSE*).
519 (**Reductive**) *DC: 3rs-m. DS: Relative location.* • **Observational criterion:** This crossvein situated close to apical wing margin, being within one of its lengths from the margin as measured from its posterior juncture with Mf (*FALSE* = this crossvein distant from wing margin by more than one of its lengths; "-" = 3rs-m absent). **• NOTES:** State not dependent on development of vein. Scored as *TRUE* for some †Ephialtitoidea, few †Praeaulacidae, some †Baissidae, †Vespoidea sp. (CASENT0844576, burmite), Vespidae, †*Burmusculus shennongii*, *Methoca* (Thynnidae), Pompilidae, †*Cretoscolia montsecana* and †*Sinoproscolia* ("Scoliidae"), most "†Angarosphecidae", *Ampulex* (Ampulicidae), some Crabronidae, Bembicidae, Philanthidae, and *Apis* (Apidae).
520 (**Reductive**) *DC: 3rs-m. DS: Relative location.* • **Observational criterion:** This crossvein situated close to 2rs-m, such that "submarginal cell 3" (= 3rm) anteroposteriorly wider than proximodistally long (*FALSE* = 3rs-m distant from 2rs-m, such that this cell longer than wide; "-" = 3rs-m absent OR 2rs-m absent). **• NOTES:** State not dependent on development of vein. This minority condition (*TRUE*) was observed in some †Ephialtitoidea, †*Praeaulacops lucidus* (†Praeaulacidae), some †Baissidae, some Evaniidae (†*Lebevania*, †*Mesevania*, †*Sinuevania*), some †*Bethylonymus*, most †Maimetshidae, †*Aculeata* sp. (CASENT0844587, burmite), some †*Burmasphex*, most Vespidae, †*Loreisomorpha*, CASENT0844580, †*Architiphia*, some †Burmusculidae, some Thynnoidea, Pompilidae, Scoliinae (where applicable; Scoliidae), fossil "Scoliidae", most "†Angarosphecidae", *Dolichurus* (Ampulicidae), most Sphecidae, most Crabronidae, *Sphecius* (Bembicidae), most Anthophila, on species of †*Camelomecia*.
521 *DC: Free R between Rsf1 and pterostigma (= Rf1). DS: Relative length.* • **Observational criterion:** Rf1 elongate, such that "submarginal cell 1" (= 1rrs) with prestigmal length greater than half total cell length (*FALSE* = Rf1 shorter, such that SMC1/1rrs with prestigmal length less than half total cell length). **• NOTES:** This minority condition was observed in some †Ephialtitoidea, most †Praeaulacidae, some †Baissidae, some Aulacidae, some Gasteruptiidae, many fossil Evaniidae, †*Albiogonalys* (Trigonalidae), some †*Burmasphex*, †*Cretofedtschenkia*, some Thynnoidea, Pompilidae, Sapygidae, most Mutillidae, *Proscolia* (Scoliidae), *Sinoproscolia* ("Scoliidae"), a few "†Angarosphecidae", Sphecidae, some Crabronidae, some Bembicidae, some Anthophila, †*Camelomecia* sp. (ANTWEB1038930), most †*Camelomecia*, most Formicidae.
522 *DC: Anterior and posterior abscissae of "submarginal cell 1" (= 1rrs). DS: Spacing.* • **Observational criterion:** Cell SMC1/1rrs extremely long and narrow, with anteroposterior width being only slightly wider than costal cell, and with proximodistal length being ≥ 3 x a–p width (*FALSE* = this cell shorter and/or wider, with length < 3 x width). **• NOTES:** Proximal width determined by 1Rsf which is often very short. Observed uniquely in Evaniidae including †*Cretevania*, †*Eovernevania*, and †*Procretevania*. This state is, however, approximated in recorded fossils of Scoliidae, in which case the "submarginal cell 2" is cored as elongate wedge-shaped.
523 (**Reductive**) *DC: Abscissae of "submarginal cell 1" (= 1rrs) and "submarginal cell 2" and "3" (= 3rsm, 4rsm). DS: Spacing.* • **Observational criterion:** Cell SMC1/1rrs as long as or longer than both SMC2/SMC3/3rsm/4rsm when these cells are present OR if SMC3/4rs-m completely absent, then compare to length of SMC2/3rsm alone (*FALSE* = SMC1/1rrs shorter than the two specified cells OR shorter than SMC2/3rsm when SMC3/4rs-m absent; "-" = 3rs-m completely absent). **• NOTES:** Observed in †*Proapocritus elegans* (†Ephialtitoidea), some †Praeaulacidae, †Anomopterellidae, some †Baissidae, Aulacidae, †Kotujellinae (Gasteruptiidae), most extinct Evaniidae, †*Bethylonymus sibiricus* (†Bethylonymidae), †Maimetshidae, Trigonalidae, †Plumalexiidae, Plumariidae, some Vespidae, †*Thanatotiphia*, *Chyphotes*, most Mutillidae, most Scoliidae (except *Trisciloa*), †*Protoscolia* ("Scoliidae"), most Apoidea (except extant Ampulicidae, some Anthophila), some †*Camelomecia*, some “†Armaniidae”, various Formicidae.
524 (**Reductive**) *DC: Abscissae of "submarginal cell 2" (= 2+3rm or 2+3rs). DS: Proportions.* • **Observational** criterion: Length and width of this cell subequal (*FALSE* = proximodistal length distinctly greater than anteroposterior width or width much narrower than length; “-” = abscissae defining this cell not developed). **• NOTES:** The conformation of the "second submarginal cell" is highly variable and appears to be quite informative. Future studies are encouraged to reformulate "SMC2" length and width characters as additive, i.e., "length < width", "length ∼ width", and "length > width". Here scored as *TRUE* for †*Praeproapocritus* (†Ephialtitidae), few †Praeaulacidae, most Aulacidae, Gasteruptiidae, some extinct Evaniidae, many †Maimetshidae, *Orthogonalys* (Trigonalidae), †Vespoidea sp. (CASENT0844576, burmite), most Vespidae, †*Thanatotiphia*, †*Architiphia*, †*Cretosapyga*, many "†Angarosphecidae", some Ampulicidae, Sphecidae, Crabronidae, Bembicidae, Pemphredonidae, Philanthidae, †*Melittosphex*, some Anthophila, †*Sphecomyrma* AMNH_NJ_242.
525 (**Reductive**) *DC: Abscissae of "submarginal cell 2" (= 2+3rm or 2+3rs). DS: Proportions.* • **Observational criterion:** This cell quadrangular to triangular, at least three times as long proximodistally as wide anteroposteriorly (*FALSE* = this cell with length < 3 x width, shape variable; “-” = abscissae defining this cell not developed). **• NOTES:** Observed in most †Ephialtitoidea, most †Praeaulacidae, †Anomopterellidae, most †Baissidae, Aulacidae, †*Hypselogastrion*, †Othniodellithidae, most Evaniidae, *Rhopalosoma* (Rhopalosomatidae), Scoliinae (Scoliidae), most fossil "Scoliidae" (except †*Sinoproscolia*), a few "†Angarosphecidae (†*Angarosphex goldringi*, †*Cretosphecium lobatum*), *Apis* (Apidae), various Formicidae. During quality control the "triangular" and "quadrangular" forms were aggregated.
526 (**Reductive**) *DC: Abscissae of "submarginal cell 2" (= 2+3rm or 2+3rs) and "discal cell 2" (= 2mcu). DS: Relative proportions.* • **Observational criterion:** SMC2/2+3rm/2+3rs nearly equal in size to DC2/2mcu (*FALSE* = these cells of dissimilar proportions OR "submarginal cell 2" not comprising 2+3rm; "-" if "submarginal cell 2" not closed by rs- m crossvein). **• NOTES:** Observed nearly uniquely in †Anomopterellidae. Scored as true for some †Baissidae.
527 (**Additive-reductive**) *DC: Abscissae of "submarginal cell 2" (= 2+3rm or 2+3rs). DS: Conformation.* • **Observational criterion:** SMC2/3rsm/4rsm triangular AND anteriorly "petiolate", *i.e.*, with 2rs-m joining Rsf proximad 2r-rs (*FALSE* = this cell quadrangular, pentangular, or other-angular OR not petiolate, *i.e.*, with 2rs-m joining Rsf distad 2r-rs; “-” = abscissae of this cell not developed). **• NOTES:** This condition is additive on the *TRUE* state of 2rs-m joining Rsf proximal to 2r-rs; these characters separate due to applicability. Scored as *TRUE* for most †Maimetshidae, †*Ilerdosphex* ("†Angarosphecidae"), *Plenoculus* (Crabronidae), *Pulverro* (Ammoplanidae), *Cerceris* (Philanthidae), and the putative apid from the Crato formation.
528 (**Reductive**) *DC: "Submarginal cell 2” and “submarginal cell 3", regardless of precise abscissa definition. DS: Relative shapes, proportions.* • **Observational criterion:** These cells of similar shape and proportions, the two being anteroposteriorly elongate, with their anteroposterior lengths at least twice their width between their anterior junctures with Rsf (*FALSE* = these cells anteroposteriorly shorter OR of different sizes; “-” if one or both cells open OR one or both cells absent). **• NOTES:** (1) Observed uniquely in †*Cretobestiola* ("†Angarosphecidae"). (2) An additional character for relative proportions of "submarginal cells 2 and 3" was deleted during quality control. This character was described as "forewing ’submarginal cells 2 and 3’ of similar shape and proportions, the two being rectangular and small" and was included initially to capture the form of Trigonalidae but was found to be generally *TRUE* in many †Ephialtitoidea and Evanioidea, thus the character was found to be underdefined. For this reason, it was removed.
529 (**Reductive**) *DC*: *"Submarginal cell 3" and “2”, regardless of abscissa composition. DS: Relative sizes.* • **Observational criterion:** SMC3 distinctly smaller than SMC2 (*FALSE* = SMC3 equal in size to or larger than SMC2; “-” = abscissae of one or both of these cells not developed). **• NOTES:** Observed in †*Mesevania* (Evaniidae), *Parischnogaster* (Vespidae), †*Cretosphecium lobatum* ("†Angarosphecidae"), and †*Cirrosphex admirabilis* ("Sphecidae"). Although proportions are convergent, the shapes of these cells are different.
530 *DC: First medial-cubital crossvein (1m-cu). DS: Expression.* • **Observational criterion:** 1m-cu tubular (*FALSE* = this abscissa nebulous, spectral, or absent). **• NOTES:** The minority condition of this character is reduction or loss of 1m-cu (*FALSE*), and was recorded in *Hyptia* (Evaniidae), several fossil Scolebythidae, most Bethylidae, most Chrysididae (except †*Burmasega*, †Chrysididae sp. (CASENT0844579), Chrysidini), Dryinidae, Sclerogibbidae, Bradynobaenidae, and numerous Formicidae (*e.g.*, Leptanillinae *sensu lato*, Proceratiinae, *Simopelta*, *Lioponera*, *Leptomyrmex*, many Formicinae, some Myrmicinae).
531 (**Reductive**) *DC: 1m-cu. DS: Curvature.* • **Observational criterion:** This crossvein distinctly curved or sinuate (*FALSE* = 1m-cu linear; "-" = inapplicable due to absence of 1m-cu). **• NOTES:** State not dependent on development of vein abscissae. The condition of non-linear 1m-cu was observed in some †Ephialtitidae, †*Nevania karatau* (†Praeaulacidae), a few Evaniidae (*Evania*, †*Grimaldivania*), †*Bethylonymus robustus*, Chrysididae, some †*Burmasphex*, *Liosphex* (Rhopalosomatidae), most Vespidae, *Sierolomorpha*, †*Architiphia*, †*Burmusculus shennongii*, †*Cretofedtschenkia*, Tiphiidae, Chyphotidae, *Aelurus* (Thynnidae), Scoliidae, most fossil "Scoliidae" (except †*Archaeoscolia senilis*, †*Cretoscolia brasiliensis*), most "†Angarosphecidae", some Ampulicidae, Sphecidae, Crabronidae, some Pemphredonidae, *Bembix* (Bembicidae), *Cerceris* (Philanthidae), most Anthophila (including the putative Crato apid), a †*Camelomecia*, and few Formicidae (e.g., *Lasiophanes*, *Stegomyrmex*).
532 (**Reductive**) *DC: 1m-cu juncture with Rs+M. DS: Relative location.* • **Observational criterion:** 1m-cu antefurcal, *i.e*., joining Rs+M proximad the distal branching point of Rsf and Mf, thus dividing Rs+M into two abscissae (*FALSE* = 1m-cu anterior juncture with Mf, thus Rs+M undivided; "-" = inapplicable due to absence of 1m-cu). **• NOTES:** (1) "Rs+M[1]" and "Rs+M[2]" in Brown & Nutting (1949) nomenclature, "1Rs+M" and "2Rs+M" in evanioid literature, *e.g.*, Engel (2017); forewing submarginal cell 2 "petiolate" sensu Brothers & Lelej (2017). (2) State not dependent on development of vein abscissae. (3) Scored as *TRUE* for Formicinae where 1m-cu is present based on comparison of formicines with 1m-cu and *Myrmelachista*: In *Myrmelachista*, free M after Rs+M diverges from Rs+M proximal to 2r-rs (this case is unique among Formicinae, even *Brachymyrmex*), while in formicines with 1m-cu (a set which does not include *Myrmelachista*), free M after Rs+M diverges at or distal to Rs+M. (4) Scored as 0 for Dolichoderinae, as not observed in dolichoderines with "second submarginal cell". (5) In future studies, the state of 1m-cu as "interstitial" should be added, i.e., the state of 1m-cu joining M at the distal split of Rs+M. Bolton (1982) provides documentation of a clear transition series for the location of 1m-cu in the Formicidae. It is unclear if any reversals have occurred, although this appears to be unlikely. (6) The "antefurcal" condition is *TRUE* for the minority of taxa, and was observed in some †Cretocleistogastrinae (†Ephialtitidae), most †Baissidae, Aulacidae, Gasteruptiidae, most †*Cretevania* (Evaniidae, plus †*Grimaldivania*), a few †Bethylonymidae, most †Maimetshidae, *Orthogonalys* and †*Cretogonalys* (Trigonalidae), †Plumalexiidae, †*Burmasphex sulcatus*, †*Loreisomorpha*, †*Thanatotiphia*, some "†Angarosphecidae", some †*Cretampulex*, †*Burmastatus* ("Sphecidae"), most Pemphredonidae, some Crabronidae, Ammoplanidae, some †*Camelomecia*, and some Formicidae (*e.g*., *Apomyrma*, *Tatuidris*, *Hypoponera*, *Tetraponera*, *Bothriomyrmex*, Formicinae, and *Stegomyrmex*).
533 (**Reductive**) *DC: 1m-cu. DS: Relative orientation.* • **Observational criterion:** 1m-cu and Mf3 forming a straight line (*FALSE* = Mf3 and 1m-cu forming distinct angle; "-" = inapplicable due to absence of 1m-cu). **• NOTES:** Mf3 is the second free abscissa of M distal to Rs, distal to 1m-cu and proximal to 2rs-m. State not dependent on development of vein abscissae. Observed in Rhopalosomatidae, some †*Curiosivespa* (Vespidae), †*Mesorhopalosoma* ("†Angarosphecidae"), *Trypoxylon* (Crabronidae), *Sphecius* (Bembicidae).
534 (**Reductive**) *DC: Abscissae of medial cell 1 (1mcu = "discal cell 1", “discoidal cell 1”). DS: Shape, proportions.* • **Observational criterion:** This cell rectangular AND small to very small (see figure) AND proximodistal length ∼ 3 x anteroposterior width (*FALSE* = 1mcu/DC1 not rectangular OR this cell larger (see figure) OR proportions dissimilar; “-” = abscissae defining this cell not developed). **• NOTES:** The first medial cell is that cell enclosed by Rs+M anteriorly, Mf1 proximally, 1m-cu distally, and Cuf posteriorly. Observed in †Cretocleistogastrinae (†Ephialtitidae), †Anomopterellidae, most †Baissidae, extinct Aulacidae, Gasteruptiidae, some fossil Scolebythidae, “vespoid genus” CASENT0844576, †*Thanatotiphia*, and †*Camelomecia* sp. (ANTWEB1038930). Scored as *TRUE* for †*Heterobaissa cancellata*, although the cell is rather wide anteroposteriorly. Also observed in the proctotrupomorph family Roprionidae.
535 (**Reductive**) *DC: Abscissae of 1mcu/DC1. DS: Proportions.* • **Observational criterion:** This cell proportionally elongate, with proximodistal length ∼ 4 x its anteroposterior width or longer (*FALSE* = 1mcu/DC1 shorter, length < 4 x width; “-” = abscissae defining this cell not developed). **• NOTES:** Scored as *TRUE* for some †Ephialtitoidea, *Taeniogonalys gundlachi* (Trigonalidae), Plumariidae, Vespoidea *sensu stricto*, †Sierolomorphidae sp. (CASENT0844589, burmite), †*Bryopompilus*, *Sierolomorpha*, most Thynnoidea, Pompilidae, Sapygidae, Mutillidae, most Scoliidae (except *Proscolia*), Ampulicidae, Sphecidae, most Crabronidae, Bembicidae, Philanthidae, most Anthophila. Described as "about" because among species of certain fossil taxa there is variation around this measurement, but otherwise the cells are similar in proportion. Initially included to capture the extremely elongate condition of the first medial cell in Vespidae, but upon critical evaluation, the *TRUE* state for the character as formulated was found to be widespread in the Aculeata, and even occurred in some ephialtitoids, and one species of sampled Trigonalidae. Therefore, two additive states were characterized, forewing first medial cell longer than half forewing length (next character) and forewing first medial cell longer than 2/5 of forewing length.
536 (**Additive**-**reductive**) *DC: Abscissae of 1mcu/DC1. DS: Relative proportions.* • **Observational criterion:** This cell relatively long, with anteroposterior length exceeding maximum anteroposterior width of wing (*FALSE* = 1mcu/DC1 relatively shorter, with length not exceeding wing width). **• NOTES:** This elongate condition was observed in *Rhopalosoma* (Rhopalosomatidae), Vespidae (including Mesozoic fossils, but some scored as uncertain), †*Cretoscolia montsecana*, Sphecidae, Bembicidae, some Philanthidae. "†*Angarosphex*" *beiboziensis* was scored as *TRUE*, but I (BEB) have misgivings about this taxon because it is a Hong name. This condition approached by *Trisciloa* (Scoliidae), *Pluto* (Pemphredonidae), but not exceeded, thus was scored as *FALSE*.
537 (**Additive-reductive**) *DC: Abscissae of 1mcu/DC1. DS: Relative proportions.* • **Observational criterion:** This cell extremely long, with proximodistal length greater than 2/5 maximum wing length (*FALSE* = this cell shorter, length < 2/5 wing length; “-” = abscissae defining this cell not developed). **• NOTES:** This final distinguishing feature of some Apoidea and some Vespidae was observed in Bembicidae, and *Pseudomasaris*, *Pachodynerus*, *Zethus*, *Mischocyttarus*, and all †*Curiosivespa* except †*C. striata*.
538 *DC: Second medial-cubital crossvein (2m-cu). DS: .* • **Observational criterion:** 2m-cu tubular (*FALSE* = this abscissa nebulous, spectral, or absent). **• NOTES:** (1) Loss of 2m- cu (*FALSE*) was observed almost half of the sampled taxa. The *FALSE* state was recorded for †*Parephialtities* (†Ephialtitoidea), †*Miniwestratia* (†Praeaulacidae), some †Baissidae, some extinct Aulacidae, Gasteruptiidae, †Othniodellithidae, all Evaniidae (except †*Mesevania*), †*Bethylonymellus* (†Bethylonymidae), †Maimetshidae, Chrysidoidea, all extant Dryinoidea, †*Aculeata* sp. (CASENT0844587, burmite), †*Falsiformica*, *Sclerogibba* (Sclerogibbidae), Rhopalosomatidae, †*Thanatotiphia*, *Sierolomorpha* (Sierolomorphidae), most Mutillidae, Scoliini (Scoliidae), Bradynobaenidae, †*Apodolichurus* (Ampulicidae), *Heterogyna* (Heterogynaidae), some Crabronidae, some Pemphredonidae, Ammoplanidae, †Melittosphecidae, †*Cretotrigona* (Apidae), † some †*Camelomecia*, “†Armaniidae”, and all Formicidae. (2) During quality control, an additional character relating to 2m-cu was thrown out due to nearly duplicating the information contained in the present character, namely "(Reductive) Abscissae enclosing forewing medial cell 2 (DC2), namely Rsf, 1m-cu, 2m-cu, and Cu, are all tubular (*FALSE* = any one of these veins nebulous, spectral, or absent; "-" scored as inapplicable)".
539 (**Reductive**) *DC: 2m-cu. DS: Curvature.* • **Observational criterion:** 2m-cu curved or sinuate (*FALSE* = this abscissa linear; "-" = 2m-cu absent/undetectable). **• NOTES:** State not dependent on development of the abscissa, inasmuch as 2m-cu must be visible to be applicable. The *TRUE* state was recorded for a minority of †Ephialtitoidea, a few †Bethylonymidae, several †Maimetshidae, a †*Burmasphex*, most Vespidae, †Pompiloidea sp. (CASENT0844580, burmite), †*Cretofedtschenkia*, Tiphiidae, Thynnidae, Pompilidae, Sapygidae, Myrmosinae (Mutillidae), Scoliinae (where applicable; Scoliidae), most fossil "Scoliidae" (exceptions were uncertain), most "†Angarosphecidae", some Ampulicidae, Sphecidae, Crabronidae, Bembicidae, *Philanthus* (Philanthidae), most Anthophila, some †*Camelomecia*.
540 (**Reductive**) *DC: 2m-cu juncture with Mf. DS: Relative location.* • **Observational criterion:** This juncture at or proximad 2rs-m crossvein (*FALSE* = this juncture distad 2rs-m; "-" = 2m-cu absent/undetectable). **• NOTES:** (1) State not dependent on development of the abscissa, inasmuch as 2m-cu must be visible to be applicable. (2) The minority condition (*TRUE*) was observed (where applicable) in *Pristaulacus* (Aulacidae), most Vespidae, Scoliidae, some Sphecidae, Crabronidae, Bembicidae, and most Anthophila. (3) An additional state was scored but deleted due to being, unexpectedly, scored as *TRUE* for all sampled taxa where it was applicable or determinable: "(Reductive) Forewing crossvein 2m-cu joining M proximal to 3rs-m crossvein (*FALSE* = joining M at or distal to 3rs-m; "-" = inapplicable due to absence/undetectability of 2m-cu)".
541 (**Reductive**) *DC: 2m-cu. DS: Relative location.* • **Observational criterion:** 2m-cu close to distal apex of wing, being within one of its length from the wing margin as measured from its posterior juncture with Cuf (*FALSE* = this abscissa distant from wing margin by more than one of its lengths; "-" = 2m-cu absent/undetectable). **• NOTES:** State not dependent on development of the abscissa, inasmuch as 2m-cu must be visible to be applicable. Among taxa for which this character could be evaluated, very nearly half were scored as *TRUE*. These taxa include various †Ephialtitoidea, some †Praeaulacidae, *Pristaulacus* (Aulacidae), one †*Bethylonymellus* and all other †Bethylonymidae, †Maimetshidae, †Vespoidea sp. (CASENT0844576, burmite), most Vespidae (but not Rhopalosomatidae), †Sierolomorphidae sp. (CASENT0844589, burmite), †*Loreisomorpha*, †*Bryopompilus*, most †Burmusculidae, Tiphiidae and Thynnidae (but not Chyphotidae), Pompilidae, Sapygidae, Myrmosinae (Mutillidae), most fossil "Scoliidae" (but not extant Scoliidae), most "†Angarosphecidae", †Sphecidae sp. (CASENT0844566), *Ampulex* (Ampulicidae), *Chalybion* (Sphecidae), *Tachysphex* (Crabronidae), Bembicidae, most Philanthidae, and *Apis* (Apidae).
542 (**Reductive**) *DC: Abscissae defining medial cell 2 (= 2mcu = "discal cell 2"). DS: Relative size.* • **Observational criterion:** This cell distinctly and proportionally large, with a surface area greater than that of "submarginal cells 2 + 3" (= 2+3rm = Rs+Rs1) OR IF "submarginal cell 3" (= 3rm = Rs1) absent THEN medial cell 2 (DC2) area larger than twice that of "submarginal cell 2" (= 2rm = Rs) (*FALSE* = 2mcu/DC2 subequal to 2rm and 3rm together OR this cell smaller than 2rm and 3rm together OR IF 3rm absent THEN this cell smaller than 2rm; "-" = inapplicable due to absence of 2m-cu). **• NOTES:** Medial cell 2 is that cell enclosed anteriorly by Mf2+ (distal to Rs+M), proximally by 1m-cu, distally by 2m-cu, and posteriorly by Cuf. The *TRUE* state described in this character is not dependent on exact development of the abscissae, inasmuch as 2m-cu must be visible (whether tubular, nebular, or spectral) to be applicable. The observed minority condition was *FALSE*, or 2m (= DC2) subequal to or smaller than the second and/or second and third radial-medial cells (= 2rm + 3rm = Rs + Rs1 = SMC2 + SMC3). This condition was scored as *FALSE* for various †Ephialtitidae, various †Praeaulacidae, †Anomopterellidae, most †Baissidae, Aulacidae, †*Xenodellitha* († Othniodellithidae), *Parischnogaster* and †*Curiosivespa zigrasi* (Vespidae), †Sierolomorphidae sp. (CASENT0844589, burmite), †*Burmusculus nuwae* (†Burmusculidae), †*Cretofedtschenkia*, †*Cretosapyga*, *Sierolomorpha* (Sierolomorphidae), *Aelurus* (Thynnidae), Sapygidae, most fossil "Scoliidae" (except †*Cretoscolia laiyangica* and †*Sinoproscolia*), Scoliinae (Scoliidae), some "†Angarosphecidae", †*Gallosphex*, †*Cirrosphex*, †*Melittosphex*, some Anthophila, and some †*Camelomecia*.
543 (**Reductive**) *DC: Abscissae defining medial cell 2 (= mc2 = "discal cell 2"). DS: Proportions, shape.* • **Observational criterion:** This cell approximately longer proximodistally than broad anteroposteriorly and more-or-less rectangular in shape (*FALSE* = mc2/DC2 as long as broad, thus about box-shaped OR shorter proximodistally than broad anteroposteriorly; "-" = inapplicable due to absence of 2m-cu). **• NOTES:** The *FALSE* condition was observed in but a few taxa, including †*Xenodellitha* (†Othniodellithidae), †*Burmusculus fuxii* (†Burmusculidae), some Vespidae, *Aelurus* (Thynnidae), †*Archisphex boothi* and †*Archisphex proximus* ("†Angarosphecidae"), †*Burmastatus*, †*Melittosphex*, a few Anthophila, and all †*Camelomecia* (where applicable).
544 (**Reductive**) *DC: Abscissae defining medial cell 2 (= mc2 = "discal cell 2"). DS: Proportions.* • **Observational criterion:** This cell ≥ 2 x as long proximodistally than broad anteroposteriorly (*FALSE* = mc2/DC2 < 2 x as long proximodistally as broad anteroposteriorly; "-" = inapplicable due to absence of 2m-cu). **• NOTES:** The minority condition was *FALSE* (medial cell 2 less than twice as long proximodistally as broad anteroposteriorly), and was observed in some †Ephialtitoidea, some †Praeaulacidae, †*Xenodellitha* (†Othniodellithidae), *Pristaulacus* (Aulacidae), some Evaniidae (most taxa inapplicable, only †*Cretevania minor TRUE*), some †*Bethylonymus* (†Bethylonymidae), some †Maimetshidae, Trigonalidae, †*Burmasphex* ("†Angarosphecidae"), †Vespoidea sp. (CASENT0844576, burmite), some Vespidae, †Sierolomorphidae sp. (CASENT0844589, burmite), †*Loreisomorpha*, †Pompiloidea sp. (CASENT0844580, burmite), †*Architiphia*, †*Bryopompilus*, †Burmusculidae, fossil "Sapygidae", †Mutillidae sp. (CASENT0844578), *Sierolomorpha* (Sierolomorphidae), *Brachycistis* (Tiphiidae), *Aelurus* (Thynnidae), Chyphotidae, Sapygidae, most Mutillidae, some fossil "Scoliidae", Campsomerini (Scoliidae), most "†Angarosphecidae", †*Cretampulex gracilis*, †Apoidea sp. (CASENT0844594, burmite), †*Burmastatus*, †*Cirrosphex*, some Pemphredonidae, †*Melittosphex*, the putative Crato apid, some Anthophila, †*Camelomecia*.
545 *DC: Abscissae defining medial cell 2 (= mc2 = "discal cell 2"). DS: Conformation.* • **Observational criterion:** This cell with an adventitious tubular vein, whether complete or incomplete (*FALSE* = mc2/DC2 without such an adventitious abscissa). **• NOTES:** This unique adventitious vein was observed in *Trisciloa* and *Campsomeris* (Scoliidae). Not scored reductively, as this condition is an obvious unique state of the Campsomerini, albeit with loss.
546 *DC: Adventitious tubular vein of mc2/DC2. DS: Extent.* • **Observational criterion:** This abscissa complete, joining both 1m-cu and 2m-cu (*FALSE* = this abscissa incomplete, joining only one of the crossveins OR this abscissa absent). **• NOTES:** Among sampled Scoliidae, observed uniquely in *Trisciloa*. Not scored reductively as unique.
547 (**Reductive**) *DC: First cubital-anal crossvein (1cu-a). DS: .* • **Observational criterion:** 1cu-a sinuate (*FALSE* = this abscissa linear; "-" = inapplicable due to absence of 1cu-a). **NOTES:** 1cu-a is the first (and in Aculeata, only) crossvein joining M+Cu or Cuf to 1A. The sinuate condition of 1cu-a was observed sporadically among sampled taxa, including some †*Praeaulacus* (†Praeaulacidae), †*Manlaya pachyura* (†Baissidae), some fossil Aulacidae, *Gasteruption* (Gasteruptiidae), a †*Burmasphex* ("†Angarosphecidae), *Rhopalosoma* (Rhopalosomatidae), some Vespidae (*Euparagia*, †*Curiosivespa curiosa*, †*C. magna*), †*Cretofedtschenkia*, †*Cretaproscolia* ("Scoliidae"), Scoliinae (Scoliidae), some Pemphredonidae, and *Thyreus* (Apidae).
548 (**Reductive**) *DC: 1cu-a. DS: Relative position.* • **Observational criterion:** 1cu-a anterior junction distinctly "postfurcal", *i.e.*, distal to branching of M and Cu from M+Cu (*FALSE* = this abscissa “interstitial” or “prefurcal”, *i.e.*, at or proximad M+Cu split; "-" = inapplicable due to absence of 1cu-a). **• NOTES:** (1) The postfurcal condition (*TRUE*) was observed in about 1/3 of the sampled taxa, including some †Ephialtitinae and all †Symphytopterinae (†Ephialtitidae), some †Kuafuidae (†Ephialtitoidea), a few †Praeaulacidae (but all †Cretocleistogastrinae), many †Anomopterellidae, few †Baissidae, several fossil Aulacidae, †*Othniodellitha* (†Othniodellithidae), all extant and some fossil Evaniidae, a few †Bethylonymidae, †*Zorophratra* (†Maimetshidae), most Trigonalidae (except †*Albiogonalys*), Plumariidae, †Plumalexiidae, some Scolebythidae, some Bethylidae, some Chrysididae, a †*Burmasphex* ("†Angarosphecidae"), some Dryinidae, Rhopalosomatidae, some Vespidae, *Myzinum* (Thynnidae), Pompilidae, *Dasymutilla* (Mutillidae), some fossil "Scoliidae" (†*Cretaproscolia* and †*Protoscolia*), Scoliinae (Scoliidae), some "†Angarosphecidae", Bembicidae, *Thyreus* and †*Cretotrigona* (Apidae), †*Orapia minor* (“†Armaniidae”), and †*Zigrasimecia ferox* and *Apomyrma* (Formicidae). (2) This is one of four characters describing the location of 1cu-a relative to the split of M+Cu; three of these ("postfurcal", "interstitial", "prefurcal") are scored non-additively, while the fourth ("extremely prefurcal") is additive on the *TRUE* condition of "prefurcal". Although the first three conditions are related, it was unclear how to score them additively. In some cases, two of these were scored as *TRUE* simultaneously in order to represent marginal cases. Future study could benefit from inclusion of an additional character, "extremely postfurcal".
549 (**Reductive**) *DC: 1cu-a. DS: Relative position.* • **Observational criterion:** 1cu-a anterior junction "interstitial", *i.e.*, opposite or very nearly opposite Mf1 (*FALSE* = proximal or distal to Mf1; “-” = inapplicable due to absence of 1cu-a). **• NOTES:** Mf1 = "1M" in evanioid nomenclature, e.g., Engel (2017). The *TRUE* state was observed in slightly less than half of all evaluated taxa, including †Ephialtitinae (†Ephialtitidae), †*Symphytogaster* (†Symphytopterinae, †Ephialtitidae), †*Kuafua* (†Kuafuidae), most †Praeaulacidae, various †Anomopterellidae, most †Baissidae, several Aulacidae, Gasteruptiidae, †*Xenodellitha* (†Othniodellithidae), various extinct Evaniidae, most †Bethylonymidae, †*Albiogonalys* (Trigonalidae), most Scolebythidae (except †*Zapenesia*), †Chrysobythidae, several Bethylidae, various Chrysididae, some †*Falsiformica*, most sampled Dryinidae, some Embolemidae, some Sclerogibbidae, several fossil Vespidae plus *Parischnogaster*, †*Architiphia*, most †Burmusculidae, *Brachycistis* (Tiphiidae), *Aelurus* (Thynnidae), Chyphotidae, Sapygidae (including fossil "Sapygidae), most Mutillidae, most fossil "Scoliidae" (except †*Protoscolia*), *Proscolia* (Scoliidae), several "†Angarosphecidae", CASENT0844594, †*Prolemistus* (Pemphredonidae), †*Melittosphex*, the putative Crato apid, *Ampulex* (Ampulicidae), *Prionyx* (Sphecidae), some Anthophila, some †*Camelomecia*, some “†Armaniidae” (†*Archaeopone taylori*, †*Armania curiosa*, †*Pseudarmania rasnitsyni*), and some Formicidae (†*Baikuris maximus*, a †*Gerontoformica*, †*Dlusskyidris*, *Anomalomyrma*, *Diacamma*).
550 (**Reductive**) *DC: 1cu-a. DS: Relative position.* • **Observational criterion:** 1cu-a anterior junction "prefurcal", *i.e.*, proximal to the branching point of M+Cu (*FALSE* = 1m-cu at or distal to branching of M+Cu; "-" = 1cu-a anterior junction opposite Mf1 OR this abscissa absent). **• NOTES:** Scored as *TRUE* in †*Gulgonga* (†Praeaulacidae), †*Electrofoenops diminuta* (Aulacidae), some †Bethylonymidae, most †Maimetshidae (except †*Zorophratra*), †*Libanobythus* (Scolebythidae), †*Holopsenelliscus* (Bethylidae), *Parnopes* (Chrysididae), *Ampulicomorpha* (Embolemidae), *Sclerogibba* (Sclerogibbidae), †*Aculeata* sp. (CASENT0844587, burmite), most †Burmasphex, some †Falsi*formica*, †*Branstetteria*, some Vespidae (†*Priorparagia*, †*Symmorphus senex*, *Euparagia*, *Pseudomasaris*), †Sierolomorphidae sp. (CASENT0844589, burmite), †Loreisomorpha, *Sierolomorpha* (Sierolomorphidae), †Pompiloidea sp. (CASENT0844580, burmite), †*Thanatotiphia*, †*Bryopompilus*, †*Burmusculus nuwae*, *Tiphia* (Tiphiidae), *Methoca* (Thynnidae), Bradynobaenidae, some "†Angarosphecidae", most Apoidea, some †*Camelomecia*, some “†Armaniidae”, most Formicidae.
551 (**Additive-reductive**) *DC: 1cu-a. DS: Relative position.* • **Observational criterion:** 1cu- a considerably proximal to branching of M+Cu, being about its length from the branching or even further proximal (*FALSE* = 1cu-a less than one of its own lengths from the split of M+Cu; "-" = inapplicable due to absence of 1cu-a). **• NOTES:** This character is additive on the *TRUE* state of the prior character ("1cu-a prefurcal"). During quality control, it was decided that this character should remain applicable even when 1cu-a was not prefurcal as this will provide further "clock" information. The *TRUE* state was observed for a small minority of taxa, including †*Thanatotiphia*, Bradynobaenidae, †*Apodolichurus*, *Dolichurus* (Ampulicidae), *Tachysphex* (Crabronidae), Ammoplanidae, †*Camelomecia* sp. (ANTWEB1038930), and various Formicidae (*e.g.*, *Martialis*, *Fulakora*, *Tatuidris*, Proceratiinae, *Simopelta*, Pseudomyrmecinae, *Aneuretus*, most Dolichoderinae, all Formicinae, all Myrmicinae).
552 (**Reductive**) *DC: Abscissae defining "subdiscal cell 1" (= 2cua = SDC1). DS: Proportions.* • **Observational criterion:** This cell anteroposteriorly wider than proximodistally long (*FALSE* = cell 2cua/SDC1 proximodistally longer than anteroposteriorly wide; "-" = inapplicable due to incomplete closure of cell 2cua). **• NOTES:** 2cua is the second cubital-anal cell, enclosed anteriorly by M+Cu or Cuf, proximally by 1cu-a, distally by Cu (whether Cuf, Cu1, or Cu2), and posteriorly by 1A. The *TRUE* state was observed in a minority of taxa, and never in Aculeata. Scored as *TRUE* for some †Ephialtitoidea, some †Praeaulacidae, †Anomopterellidae, most †Baissidae, most fossil Aulacidae, †*Hypselogastrion* (Gasteruptiidae), †*Grimaldivania* (Evaniidae), and †*Maimetshasia kachinensis* (†Maimetshidae).
553 *DC: Free Cubitus, i.e., that abscissa distad M+Cu (= Cuf). DS: Expression.* • **Observational criterion:** Cuf tubular (*FALSE* = this abscissa absent or poorly developed). **• NOTES:** Loss of Cuf was the minority condition and was observed in †*Bethylonymellus oligocerus* (†Bethylonymidae), Hyptia (Evaniidae), †*Uliobythus* (Scolebythidae), some Bethylidae, various Chrysididae, Dryinidae, Embolemidae, fossil Sclerogibbidae, one species of sampled †*Falsiformica*, *Olixon*, Bradynobaenidae, †*Apodolichurus* (Ampulicidae), and *Odontomachus meinerti* (Formicidae). Scored as absent for Sapygidae and Myrmosidae by Brothers & Carpenter (1993; their character 104).
554 (**Reductive**) *DC: M+Cu, Cuf1, cu-a. DS: Conformation.* • **Observational criterion:** These abscissae in forming the “vespid triangle”, *i.e.*, cu-a postfurcal, short, and subequal in length to the first free abscissa of Cu (Cuf1), which itself obliquely angles posteroapically from M+Cu to meet cu-a at a similar angle (*FALSE* = Cuf1 and cu-a dissimilar in length AND/OR not meeting at an oblique angle AND/OR Cuf1 directed distad or anterodistad from its juncture with M+Cu AND/OR cu-a interstitial to prefurcal AND/OR these abscissae long (see figure)). **• NOTES:** Observed primarily in Vespidae, but also scored as *TRUE* for *Allocoelia* (Chrysididae). Among Vespidae, scored as *TRUE* for *Pachodynerus*, *Zethus*, and *Mischocyttarus*.
555 (**Reductive**) *DC: Juncture of first (Cu1) and second (Cu2) cubital abscissae. DS: Angle.* • **Observational criterion:** These abscissae diverging at approximately a right angle (*FALSE* = diverging more-or-less obliquely or no divergence occurring; "-" = inapplicable due to absence of divergence of Cu1 and Cu2). **• NOTES:** About 2/5 of the sampled taxa were scored as *TRUE*, including some †Ephialtitoidea, some †Praeaulacidae, some †Baissidae, most fossil Aulacidae, some fossil Evaniidae, some †Bethylonymidae, some †Maimetshidae, †*Burmasphex*, some †*Eorhopalosoma*, †*Symmorphus senex* (Vespidae), †*Bryopompilus*, †*Burmusculus shennongii*, †Mutillidae sp. (CASENT0844578), *Brachycistis* (Tiphiidae), Pompilidae, Sapygidae, some fossil "Scoliidae"*, Proscolia* (Scoliidae), various fossil "†Angarosphecidae", CASENT0844566, *Ampulex* (Ampulicidae), Bembicidae, Ampulicidae, some Pemphredonidae, most Philanthidae, some Anthophila, some †*Camelomecia*, and †*Orapia rayneri* (“†Armaniidae”).
556 (**Reductive**) *DC: Posterior branch of Cu (CuPf1). DS: Orientation.* • **Observational criterion:** This abscissa directed proximad from its proximal split with the anterior branch of Cu (CuAf1) (*FALSE* = this abscissa directed posterad OR distad; “-” = inapplicable due to absence of CuPf1). **• NOTES:** This "typically" vespid condition was observed in a minority of taxa, for which the *TRUE* state was recorded for †*Westratia curtipes* (†Praeaulacidae), †Anomopterellidae, †*Electrofoenops rasnitsyni* (Aulacidae), *Gasteruption* (Gasteruptiidae), †*Cretevania cyrtocera* (Evaniidae), Vespidae, most fossil "Scoliidae" (except †*Cretoscolia laiyangica*, †*Protoscolia*), Scoliinae (Scoliidae), some "†Angarosphecidae", Bembicidae, *Clypeadon* (Philanthidae), and the putative Crato apid. Initially ?*Symmorphus senex* (Vespidae) was scored as *FALSE*, but during quality control this was changed to uncertain ("?").
557 (**Additive-reductive**) *DC: Proximal and distal abscissae of free Cubitus (Cuf) relative to 1m-cu. DS: Orientation.* • **Observational criterion:** These abscissae set at a narrow acute angle relative to one another, with the distal abscissa directed proximad (*FALSE* = these abscissae at a broader angle AND/OR distal abscissa not directed proximad; “-” = inapplicable due to absence of one or both abscissae). **• NOTES:** This is the "apically produced" state of the "subdiscal cell" which has been adduced as synapomorphy of Euparagiinae in Vespidae by Perrard *et al*. (2017, their character 49), where they recorded this condition as true for *Euparagia*, †*Curiosivespa*, and †*Priorparagia,* and scored as polymorphic for *Gayella eumenoides.* Here, the *TRUE* state was scored for *Euparagia*, all †*Curiosivespa*, and for †*Priorparagia*, as done by Perrard *et al*. (2017). Among Vespidae, scored as *FALSE* for †*Priorvespa*, †*Protovespa*, †*Symmorphus senex*, and all extant taxa except *Euparagia*.
558 *DC: Posterior branch of Cu (CuPf1) and first anal vein (1A). DS: Connection.* • **Observational criterion:** CuPf1 joining 1A as tubular abscissa, even if CuPf1 fenestrated Second cubital vein (Cu2, posterior branch of free Cu) reaching 1A vein (*FALSE* = Cu2 not reaching 1A OR reduced or absent, OR A absent distal to cu-a. **• NOTES:** (1) This is state 0 of character 215 in Appendix VI of Brothers & Carpenter (1993), wherein contact of Cu2 and 1A is recorded as generally true for Vespoidea, Pompiloidea *sensu lato*, Scolioidea, Formicoidea, and Apoidea, in contrast to the Chrysidoidea, Dryinoidea, and Sclerogibbidae. (2) Here, the minority state was *FALSE*, and was recorded for †*Miniwestratia* (†Praeaulacidae), †*Baissa anomala* and †*Tillywhimia spectra* (†Baissidae), most fossil Aulacidae, *Gasteruption* (Gasteruptiidae), †*Sorellevania* and *Hyptia* (Evaniidae), †*Bethylonymellus* (†Bethylonymidae), various †Maimetshidae, †*Albiogonalys* (Trigonalidae), most Chrysidoidea (except †*Unicobythus*, some fossil Bethylidae, †Chrysididae sp. CASENT0844579), †Aculeata sp. (CASENT0844587, burmite), †*Burmasphex*, Dryinoidea (except †*Hybristodryinus resinicolus*), Sclerogibbidae, †*Loreisomorpha* (Sierolomorphidae), Bradynobaenidae, †*Pompilopterus corpus* ("†Angarosphecidae"), †*Pittoecus pauper* (Pemphredonidae), †*Cretotrigona* (Apidae), †*Pseudarmania rasnitsyni* (“†Armaniidae”), and most Formicidae.
559 *DC: Second anal vein (2A). DS: Expression.* • **Observational criterion:** This abscissa present (*FALSE* = 2A absent). **• NOTES:** Presence of 2A is ancestral in the Hymenoptera, and among sampled taxa was observed in most †Ephialtitoidea, †*Eonevania* and †*Nevania* (†Praeaulacidae), and †*Cretevania cyrtocera* (Evaniidae). It is possible that the state observed in †*Nevania* and †*Eonevania* represent reversals to presence from absence and is almost certainly the case in the †*Cretevania* species.
560 *DC: First anal-anal crossvein (1a-a), i.e., transverse abscissa joining 1A and 2A. DS: Expression.* • **Observational criterion:** This abscissa present (*FALSE* = 1a-a absent). **• NOTES:** As with the prior character (2A presence), presence of crossvein 1a-a is ancestral in the Hymenoptera. Here it was scored as *TRUE* for various †Ephialtitoidea and †*Cretevania cyrtocera* (Evaniidae). The *TRUE* state in †*Cretevania certocera* is certainly a reversal.

**-** *Hind wing*

561 *DC: Basal hamuli. DS: Expression.* • **Observational criterion:** These proximal wing- linking hooks present (*FALSE* = these hamuli absent). **• NOTES:** Hind wing basal hamuli were observed primarily in extant taxa, including Plumariidae, Bethylidae, Embolemidae, Rhopalosomatidae, Chyphotidae, some Mutillidae, *Proscolia* (Scoliidae), Heterogynaidae, and *Chalybion* (Sphecidae). Most fossils were scored as uncertain ("?").
562 *DC: Costal vein (C). DS: Expression.* • **Observational criterion:** This vein present (*FALSE* = C absent). **• NOTES:** Observed in †Ephialtitoidea, some †Praeaulacidae, †Maimetshidae, Plumariidae, some Chrysididae, vespoid genus CASENT0844576, many Vespidae, most Thynnoidea, Pompilidae, †Burmusculidae, Sapygidae, some Mutillidae, Scolioidea, and most Apoidea. For Evanioidea, scored as character 52 in Li *et al*. (2018); for Apoidea, scored as character 12 by Prentice (1998).
563 *DC: Free Radial sector (Rsf). DS: Expression.* • **Observational criterion:** This abscissa present, even if partially nebulous (*FALSE* = Rsf absent). **• NOTES:** Rsf is the abscissa of Rs which is "free" from R+Rs, *i.e.*, Rf after the split of R+Rs. Observed in †Ephialtitoidea, †Praeaulacidae, †Anomopterellidae, †Othniodellithidae, †Bethylonymidae, some Evaniidae, most †Maimetshidae (except †*Zorophratra*), Trigonalidae, Vespoidea, Pompiloidea *sensu lato*, Scoliidae, most Apoidea, and most Formicidae. For Evanioidea, scored as character 54 in Li *et al*. (2018).
564 *DC: Rsf. DS: Curvature.* • **Observational criterion:** This abscissa enclosing distal cell of wing by curving anteriorly to meet anterior wing margin (*FALSE* = Rsf not curving anteriorly to meet anterior wing margin). **• NOTES:** This state was observed in †Ephialtitidae, most †Praeaulacidae, and a minority of †Bethylonymidae.
565 *DC: First radial-medial crossvein (1rs-m). DS: Expression.* • **Observational criterion:** This abscissa tubular (*FALSE* = 1rs-m nebulous, spectral, or absent). **• NOTES:** 1rs-m is easiest to recognize by focusing on the Radial sector, as in all sampled taxa it is the only crossvein connected to Rf; development of free M (Mf) is variable and thus difficult to use as a consistent identifying juncture. Crossvein 1r-m was observed in a majority of taxa and was present in most but not all taxa which have Rsf.
566 (**Reductive**) *DC: Juncture of 1rs-m and R+Rs or Rs. DS: Relative location.* • **Observational criterion:** 1rs-m antefurcal, *i.e.*, joining R+Rs prior to branching of Rf and Rsf (*FALSE* = this abscissa interstitial or postfurcal). **• NOTES:** An antefurcal anterior juncture of crossvein r-m was observed among sampled taxa only in Formicidae (*Paraponera*, *Aneuretus*, *Linepithema*, some Formicinae, and some Myrmicinae; note that this list is not exhaustive).
567 *DC: Second free abscissa of M (Mf2), 1rs-m, Rsf. DS: Conformation.* • **Observational criterion:** These abscissae in the following conformation: Mf2 present AND 1rs-m meeting Rsf at an oblique angle AND 1r-sm strongly curved or V-shaped thus "broken" in appearance, *i.e.*, crossvein with distinct angle, with the abscissa anterior to the angle’s point oriented more-or-less in parallel with longitudinal axis of wing and the abscissa posterior to the angle’s point directed more-or-less posteriorly (*FALSE* = Mf2 absent OR 1rs-m and Rsf forming a more-or-less straight line OR 1rs-m weakly or not curved). **• NOTES:** Observed uniquely in Rhopalosomatidae, scored as *TRUE* for *Rhopalosoma*, †*Eorhopalosoma*, and the contentious fossil †*Mesorhopalosoma* which is currently placed in "†Angarosphecidae".
568 *DC: Mf2. DS: Expression.* • **Observational criterion:** This abscissa tubular (*FALSE* = Mf2 nebulous, spectral, or absent). **• NOTES:** Mf2 is the second abscissa of M, distal to the juncture of Mf1 with 1rs-m. Mf2 was frequently observed as absent. Mf2 was observed to be retained, as it were, in most †Ephialtitoidea, †Praeaulacidae, †Anomopterellidae, †Othniodellithidae, †Bethylonymidae, some †Maimetshidae, some Trigonalidae, some Evaniidae, †Plumalexiidae, Plumariidae, some †*Burmasphex*, †Vespoidea sp. (CASENT0844576, burmite), Rhopalosomatidae, most Vespidae, most Pompiloidea *sensu lato* (except Sierolomorphidae and some Mutillidae), most Scoliidae, many Apoidea, †*Camelomecia*, and a number of Formicidae.
569 *DC: Free Cubitus (Cuf). DS: Expression.* • **Observational criterion:** This abscissa present, tubular at least at its very base (*FALSE* = Cuf completely nebulous or spectral or absent). **• NOTES:** Cuf is the abscissa of Cu which is distal to the split of M+Cu and is retained more frequently than Mf2.
570 *DC: Cubital-anal crossvein (cu-a). DS: Expression.* • **Observational criterion:** This abscissa present with any development (*FALSE* = cu-a absent). **• NOTES:** Retained with about equal frequency of Cuf.
571 (**Reductive**) *DC: Juncture of Cu and cu-a. DS: Relative location.* • **Observational criterion:** cu-a postfurcal, *i.e.*, junction with Cu distal to split of M and Cu from M+Cu (*FALSE* = present proximal to split; "-" = inapplicable, *i.e.*, Cuf and cu-a absent). **• NOTES:** This character used with some frequency in prior studies (*e.g.*, Brothers & Carpenter 1993 as their character 103 of their Appendix VI). Among the taxa where this state is applicable, *antefurcal* cu-a junctures were observed in some †Ephialtitoidea, some †Praeaulacidae, †Othniodellithidae (inapplicable in other Evanioidea), some †Bethylonymidae, †Plumalexiidae, Plumariidae (inapplicable in Chrysidoidea, Dryinoidea), Rhopalosomatidae, some Vespidae, Sierolomorphidae, some Thynnoidea (*Myzinum*, Chyphotidae), some Mutillidae, Scolioidea, most Apoidea (except Ampulicidae, Bembicidae, Ammoplanidae, some Philanthidae), some †*Camelomecia*, and all Formicidae.
572 (**Reductive**) *DC: Abscissae defining the “basal cell” (= BC = 1rsm). DS: Conformation.* **Observational criterion:** This cell distally elongate (*FALSE* = BC/1rsm not distally elongate; "-" = inapplicable due to complete absence of Rsf and Mf). **• NOTES:** The “basal cell” is the first cell between R+Rs and M+Cu and is enclosed distally by 1rs-m; a second “basal cell” is not observable in Aculeata, †Bethylonymoidea, Trigonaloidea, or Evanioidea. During the character development phase, it became apparent that Formicidae consistently differ from other Aculeata (Apoidea–Vespoidea). At first it appeared this may be a synapomorphy of the Apoidea–Vespoidea, but upon evaluation of all Hymenoptera at more-or-less the family level, this condition was also observed in Xiphydriidae, Orussidae, various Ichneumonidae (although without curvature of Rsf and Mf), many †Ephialtitoidea, some †Praeaulacidae, a few †Bethylonymidae (†*Meiagaster*, †*Allogaster*), †Vespoidea sp. (CASENT0844576, burmite), Vespidae, many Pompiloidea *sensu lato* (except *Sierolomorpha*, Chyphotidae, some Mutillidae), some Scolioidea (Scoliini, Bradynobaenidae), most Apoidea (except *Heterogyna*, *Oxybelus*), and some †*Camelomecia*. Some †*Bethylonymus* are marginal cases which have been scored as *FALSE*, but a clear case of presence is observed in †*Meiagaster*. The condition in Trigonalidae is similar, but without the distinct r-m between Rsf and Mf, thus the apex of the cell is pointed, rather than squared off. Among Aculeata, this character is inapplicable for the majority of Chrysidoidea, and the Dryinoidea.
573 *DC: Jugal lobe. DS: Expression.* • **Observational criterion:** This lobe present (*FALSE* = jugal lobe absent). **• NOTES:** Despite the generally accepted importance of jugal lobes, they are exceedingly difficult to evaluate for fossils, even specimens in amber. In Formicidae, several distinct reduction grades are observable, including complete presence with transition to complete loss within a single genus (*Thaumatomyrmex*, Ponerinae).
574 (**Reductive**) *DC: Jugal lobe and anal area. DS: Relative size.* • **Observational criterion:** Jugal lobe large, nearly as long as anal area (*FALSE* = this lobe smaller, distinctly shorter than anal area; “-” = inapplicable due to absence of the jugal lobe). **• NOTES:** Future studies may improve the informativeness of this character by defining clear gradients of relative length, as some jugal lobes are extremely large.
575 *DC: Plical lobe (the area posterior to the plical fold, which is between M+Cu and 1A). DS: Distal apex.* • **Observational criterion:** Plical lobe apically marked by an incision at the wing margin (*FALSE* = this incision absent). **• NOTES:** Plica here used *sensu* Hamilton (1971, 1972a,b,c), corresponding to states 0 and 1 of character 107 in Appendix VI of Brothers & Carpenter (1993).
576 (**Reductive**) *DC: Plical incision. DS: Depth.* • **Observational criterion:** This incision shallow (*FALSE* = moderate to deep; “-” = inapplicable due to absence of incision). **• NOTES:** Used *sensu* Brothers & Carpenter (1993), states 0 (= *FALSE*) and 1 (= *TRUE*) of character 107 in Appendix VI. Shallow scored by Brothers & Carpenter (1993) for: Dryinidae, Thynnoidea, "mutillids", Sierolomorphidae, Pompilidae, Formicidae, Vespidae; moderate to deep scored by them for: Chrysidoidea, Sclerogibbidae, Scolebythidae, Embolemidae, Apoidea, Anthoboscinae, Rhopalosomatidae, and Eotillini.

### Character state illustrations

All images were manually selected to best represent the state or states under consideration. An effort has been made to capture variation within and among taxa, but some characters could not be illustrated as certain loaned material was returned prior to the assembly of these figures (Table 1). These exceptions are listed below, prior to the figure legends. Refer to the character state definitions for explanation of the figures. To ease the process of character evaluation, all images were processed in Photoshop, most were cropped to fit the panels, and many were reflected or rotated such that cranial is left and caudal right. To decrease clutter in the legends, image sources are provided in Supplemental File S1. Tables and references are provided after the legends.

**Table 13.**
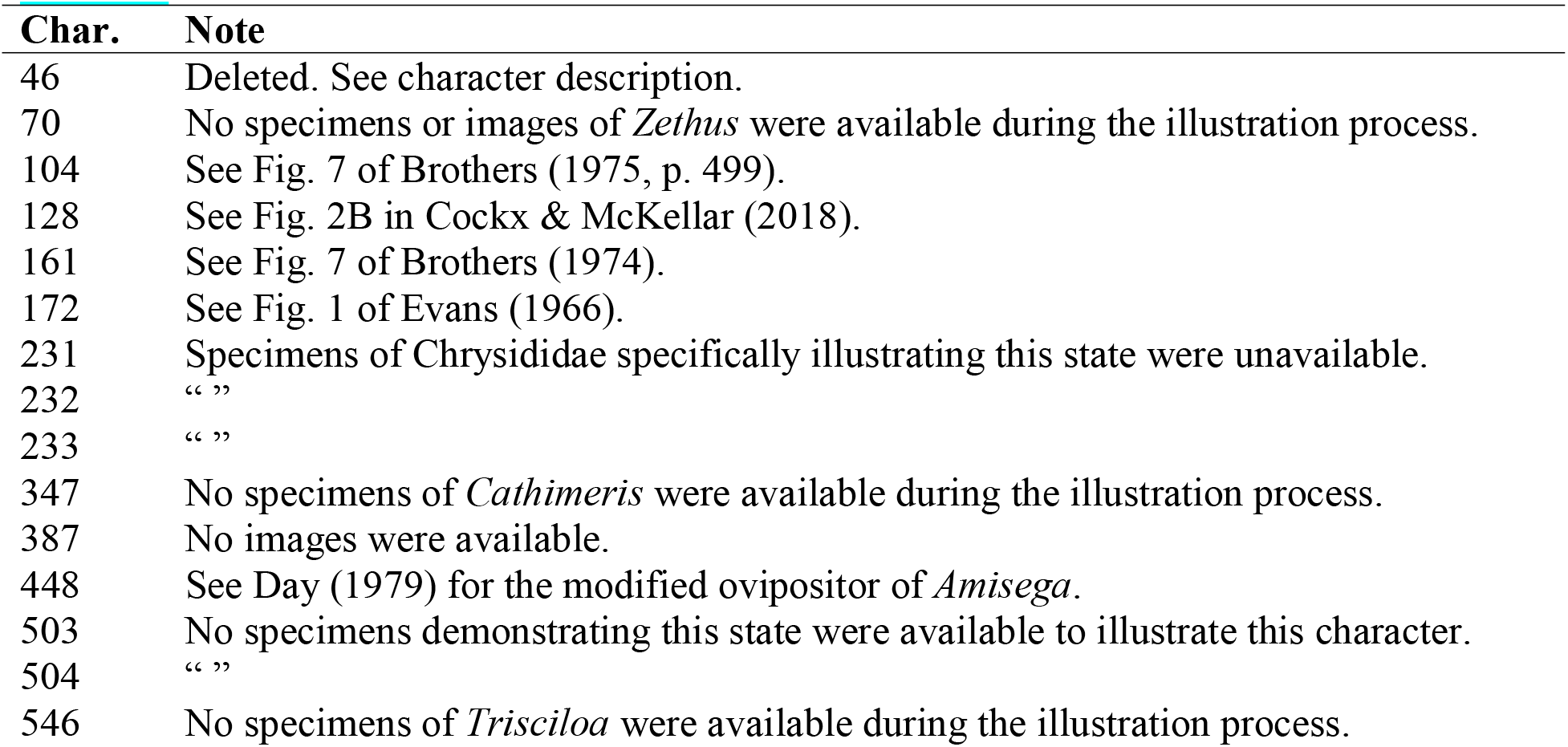
Fifteen of the 576 described characters are not illustrated here.

| Char. | Note |
| --- | --- |
| 46 | Deleted. See character description. |
| 70 | No specimens or images of *Zethus* were available during the illustration process. |
| 104 | See Fig. 7 of Brothers (1975, p. 499). |
| 128 | See Fig. 2B in Cockx & McKellar (2018). |
| 161 | See Fig. 7 of Brothers (1974). |
| 172 | See Fig. 1 of Evans (1966). |
| 231 | Specimens of Chrysididae specifically illustrating this state were unavailable. |
| 232 | “ ” |
| 233 | “ ” |
| 347 | No specimens of *Cathimeris* were available during the illustration process. |
| 387 | No images were available. |
| 448 | See Day (1979) for the modified ovipositor of *Amisega*. |
| 503 | No specimens demonstrating this state were available to illustrate this character. |
| 504 | “ ” |
| 546 | No specimens of *Trisciloa* were available during the illustration process. |

## Acknowledgments

We thank: Dave Grimaldi (AMNH), Brian Fisher (CASC), Martin Hauser and Kevin Williams (CDFA), Taiping Gao and Huijia Cao (CNUC), Bo Wang (NIGPAS), Lynn Kimsey, Steve Heydon, and Phil Ward (UCDC), and many others for access to specimens and specimen data; Jason Bond for sharing computational resources; Mike May for RevBayes and MCMC guidance; Jéssica Gillung for alternative UCE assemblies; Waring (Buck) Trible for providing comments on an earlier draft; Rolf Beutel, Marek Borowiec, Jim Doyle, Daniel Friedman, J. Gao, Steve Heydon, Roberto Keller, Lynn Kimsey, Seraina Klopfstein, Ziv Lieberman, Jack Longino, István Mikó, Corrie Moreau, Jill Oberski, Oliver Niehuis, Ralph Peters, Matt Prebus, Alex Rasnitsyn, and Phil Ward for numerous discussions; Jim Carpenter for clarifying authority names of Vespidae; and Simon van Noort for graciously allowing us to use his images to illustrate our characters. For thoughtful and exceedingly valuable reviews of a prior version of this work, we thank Fred Ronquist, Corrie Moreau, and two anonymous reviewers; these reviews provided both substantial action items to address and the time to improve the manuscript. BEB is grateful to Jacobus (Koos) Boomsma for clarifying the distinction between Wheeler’s and Wilson’s notions of eusociality; BEB thanks Jon Shik for initiating this interaction. We acknowledge the KIT light source for provision of instruments at their beamlines; we thank the Institute for Beam Physics and Technology (IBPT) for operation of the storage ring, the Karlsruhe Research Accelerator (KARA), and we are grateful to the IPS staff for assistance at their beamline. We dedicate this work to Barry Bolton for his epoch-defining work on the comparative morphology and systematics of ants, and to the memory of Christian Peeters, who emphasized the importance of anatomical function in explaining the evolution of ant morphology. RIP Ed Wilson.

## Author contributions

B.E.B. conceived the study. B.E.B., P.B., V.P. provided fossils for study. B.E.B., A.R., T.v.d.K. performed µ-CT and digital data reconstruction. B.E.B. generated the morphological data matrix. A.R., Z.G., P.B., V.P. contributed to character interpretation. B.E.B., Z.K. generated new sequence data. Z.K. processed sequence data. B.E.B., Z.K. performed phylogenetic analysis. B.E.B. interpreted the results, wrote the entire original daft, and made the figures. Z.K., A.R., T.v.d.K., Z.G., P.B. participated in pre-submission draft revision.

## Data and materials availability

Digital morphological models, trees, data matrices, custom scripts, list of specimen repositories, and additional phylogenetic output are available at Zenodo (doi: [ZENODO_XXX]). Undescribed fossils are imaged and databased on AntWeb.org. Raw UCE reads have been submitted to the NCBI SRA (BioProject PRNJA637110; XXX). ZooBank Life Science Identifiers (LSID) for each new name recognized in the present study are as follows: †*Renymus* (urn:lsid:zoobank.org:act:252AF945-F7E6-47B3-BACE-A7622546249D), †*Apisphex* (urn:lsid:zoobank.org:act:572DFFF2-0E80-4B3A-A79A-AFA4096EF746), †@@@idae (urn:lsid:zoobank.org:act:AD2C2819-A25E-4FC5-BBF2-0EFA456D3C91), †Aquilomyrmecini (urn:lsid:zoobank.org:act:73EFE029-7478-4754-A467-5F15F983A534), Opamyrmini (urn:lsid:zoobank.org:act:6939AB2D-B428-4025-8D6C-B6327743AFE8).

---

**Chars. 1-11.**
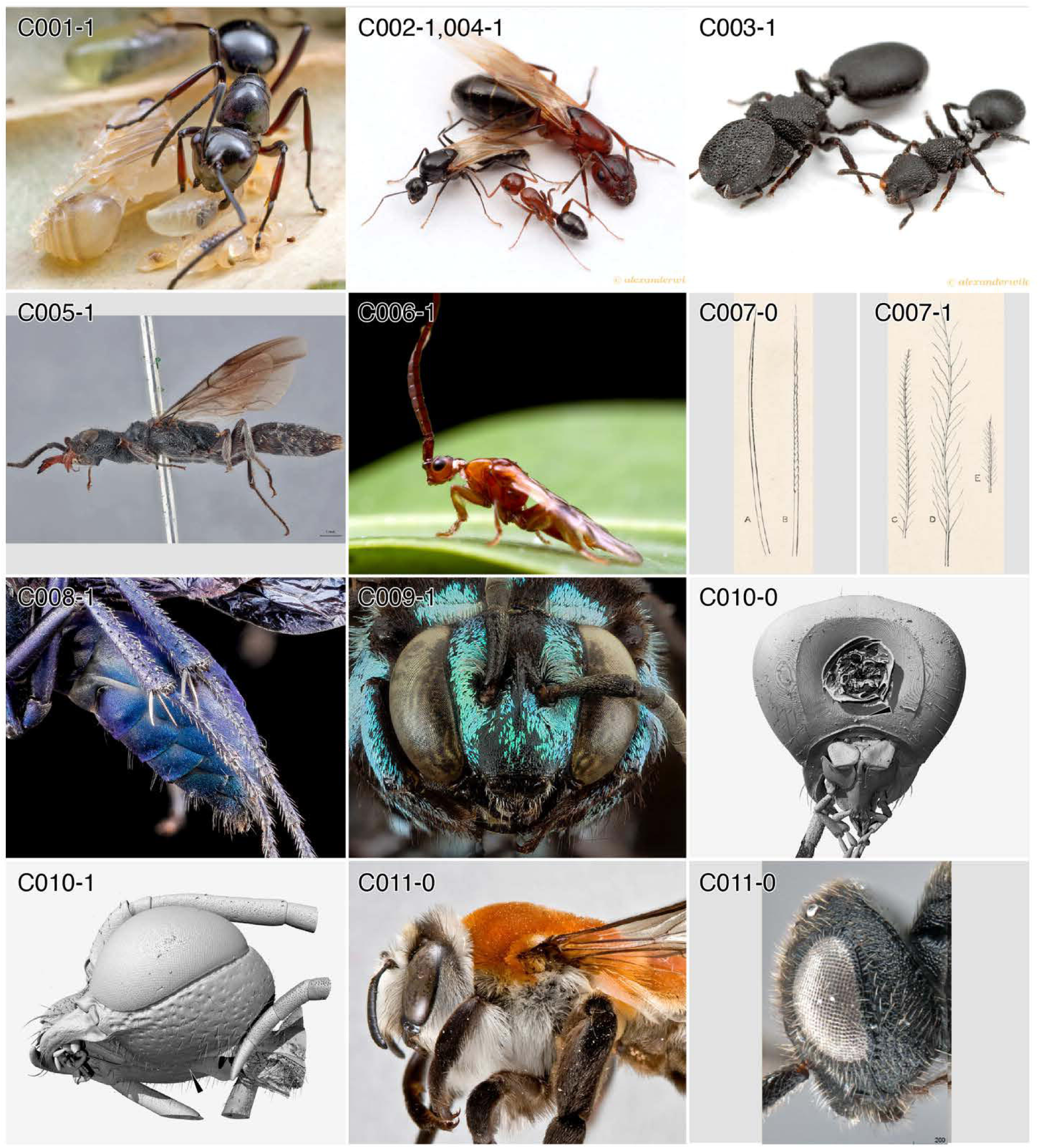
Character-states: (C1-1) Reproductive division of labor; *Polyrhachis robsoni.* **(C2-1)** Winglessness; *Camponotus discolor.* **(C3-1)** Discrete soldiers; *Cephalotes rohweri.* **(C4-1)** Winged-wingless diphenism; *Ca. discolor.* **(C5-1)** Body flattened; *Pristocera communis.* **(C6-1)** Setal tufts; Loboscelidiinae. **(C7-0, -1)** Simple and plumose setae. **(C8-1)** Pubescence iridescent; *Pepsis ruficornis.* **(C9-1)** Iridescent squamiform setae; *Thyreus.* **(C10-0)** Postgenal bridge short; *Anoplius* (*Arachnophronctus*). **(C10-1)** Postgenal bridge long; *Ampulex* sp. **(C11-0a)** Head “hypognathous”; *Caupolicana fulvicollis.* **(C11-0b)** Head “hypognathous”; *Bocchus forestalls.* **Families: (C1-4)** Formicidae. **(C5)** Bethylidae. **(C6)** Chrysididae. **(C8)** Pompilidae. **(C9)** Apidae. **(C10-0)** Pompilidae. **(C10-1)** Ampulicidae. **(C11-0a)** Colletidae. **(C11-0b)** Dryinidae.

**Chars. 11-15.**
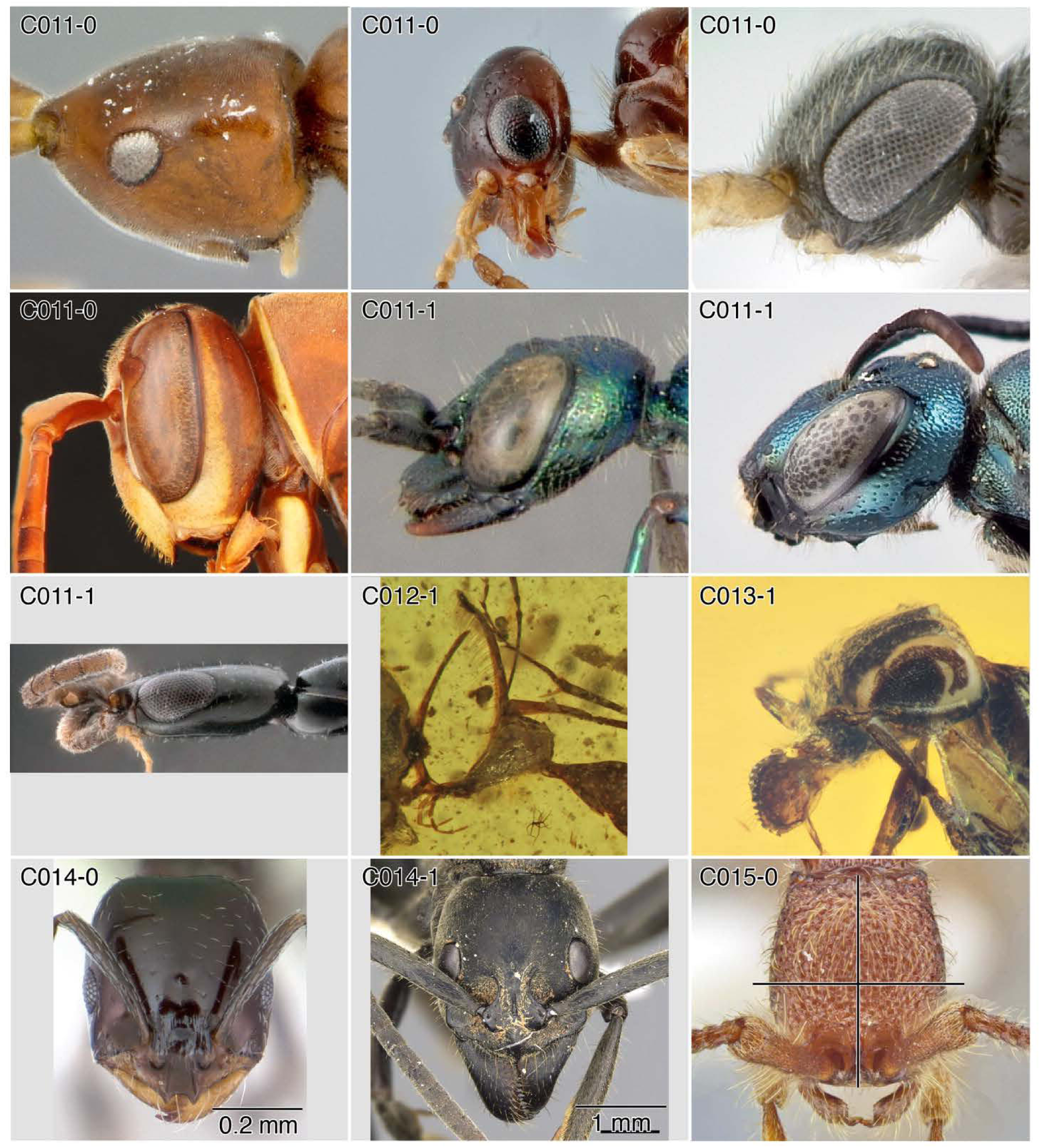
Character-states: (C11-0) Head “hypognathous”; (a) *Embolemus africanus*; (b) *Myrmecopterinella okahandja*; (c) *Probethylus callani*; (d) *Polistes bellicosus.* **(C11-1)** Head “hypognathous”; (a) *Ampulex bredoi*; (b) *Ceratina calcarata*; (c) *Tuberepyris nihilus.* **(C12-1)** Head orally narrowed; †*Ceratomyrmex ellenbergeri.* **(C13-1)** Face profile orally concave; †*Camelosphecia fossor.* **(C14-0)** Head width < 1 mm; *Monomorium minimum.* **(C14-1)** Head width ≥ 1 mm; *Megaponera analis.* **(C15-0)** Head longer than wide; *Syscia borowieci.* **Families: (C11-0a)** Embolemidae. **(C11-0b)** Plumariidae. **(C11-0c)** Sclerogibbidae. **(C11-0d)** Vespidae. **(C11-1a)** Ampulicidae. **(C11-1b)** Apidae. **(C11-1c)** Bethylidae. **(C12, 14, 15)** Formicidae. **(C13)** †@@@idae.

**Chars. 15–25.**
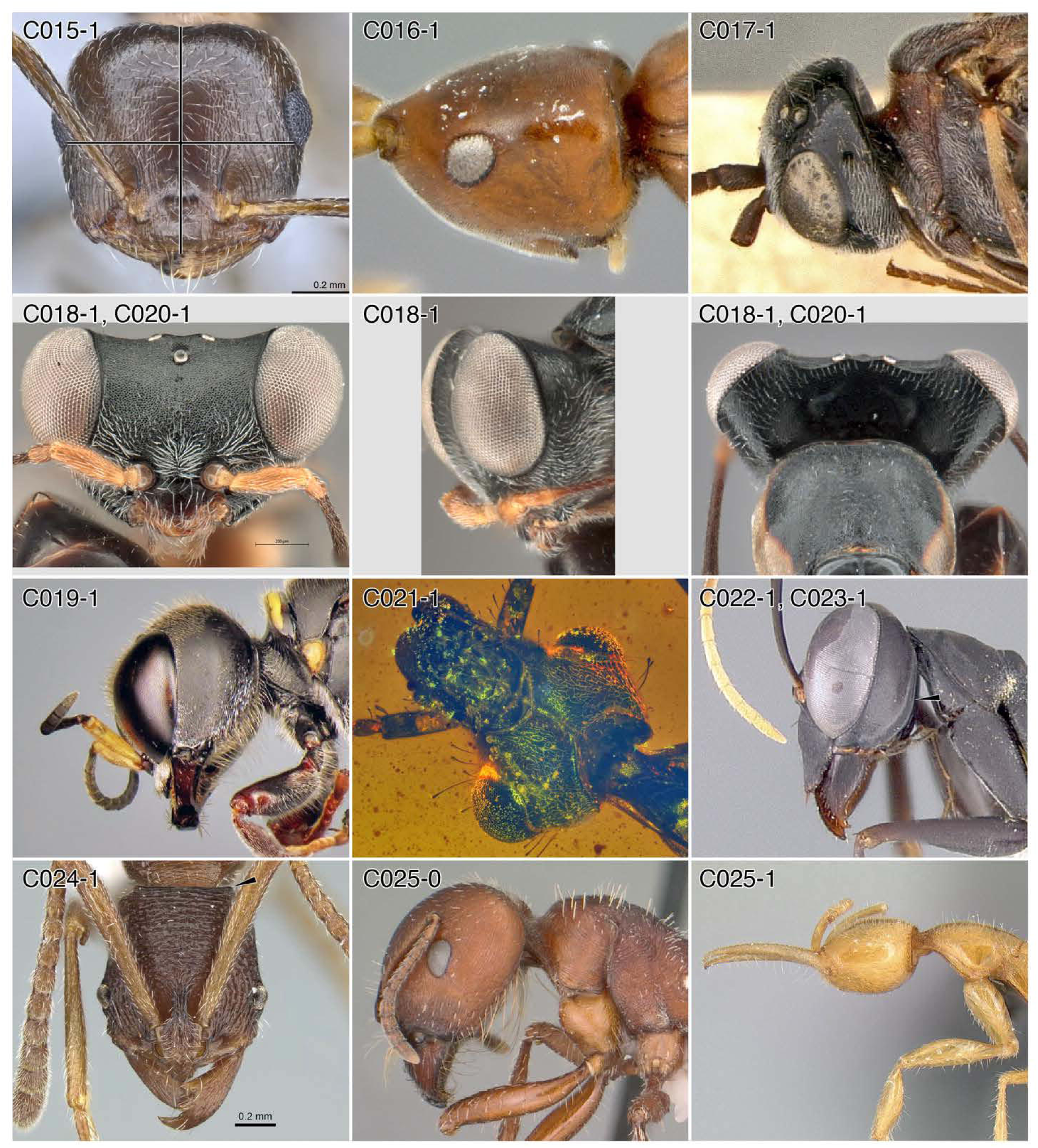
Character-states: (C15-1) Head wider than long; *Crematogaster amita matabele.* **(C16-1)** Head conical; *Embolemus africanus.* **(C17-1)** Head posteriorly truncate; *Zeuxevania* sp. **(C18-1)** Head wedge-shaped; *Dryinus dentatiforceps.* **(C19-1)** Head cuboidal; *Dasyproctus braunsii.* **(C20-1)** Aboral head margin concave; *Dr. dentatiforceps.* **(C21-1)** Occipital carina present; †*Camelomecia* sp. **(C22-1)** Occipital carina on hypostoma; *Gigantiops destructor.* **(C23-1)** Occipital carina flared; *G. destructor.* **(C24-1)** Occipital carina frontally visible; *Simopelta andersoni.* **(C25-0)** Postocciput below head apex; *Pogonomyrmex barbatus.* **(C25-1)** Postocciput at head apex; *Martialis heureka.* **Families: (C15, 22–25)** Formicidae. **(C16)** Embolemidae. **(C17)** Evaniidae. **(C18, 20)** Dryinidae. **(C19)** Crabronidae. **(C21)** †@@@idae.

**Chars. 26–36.**
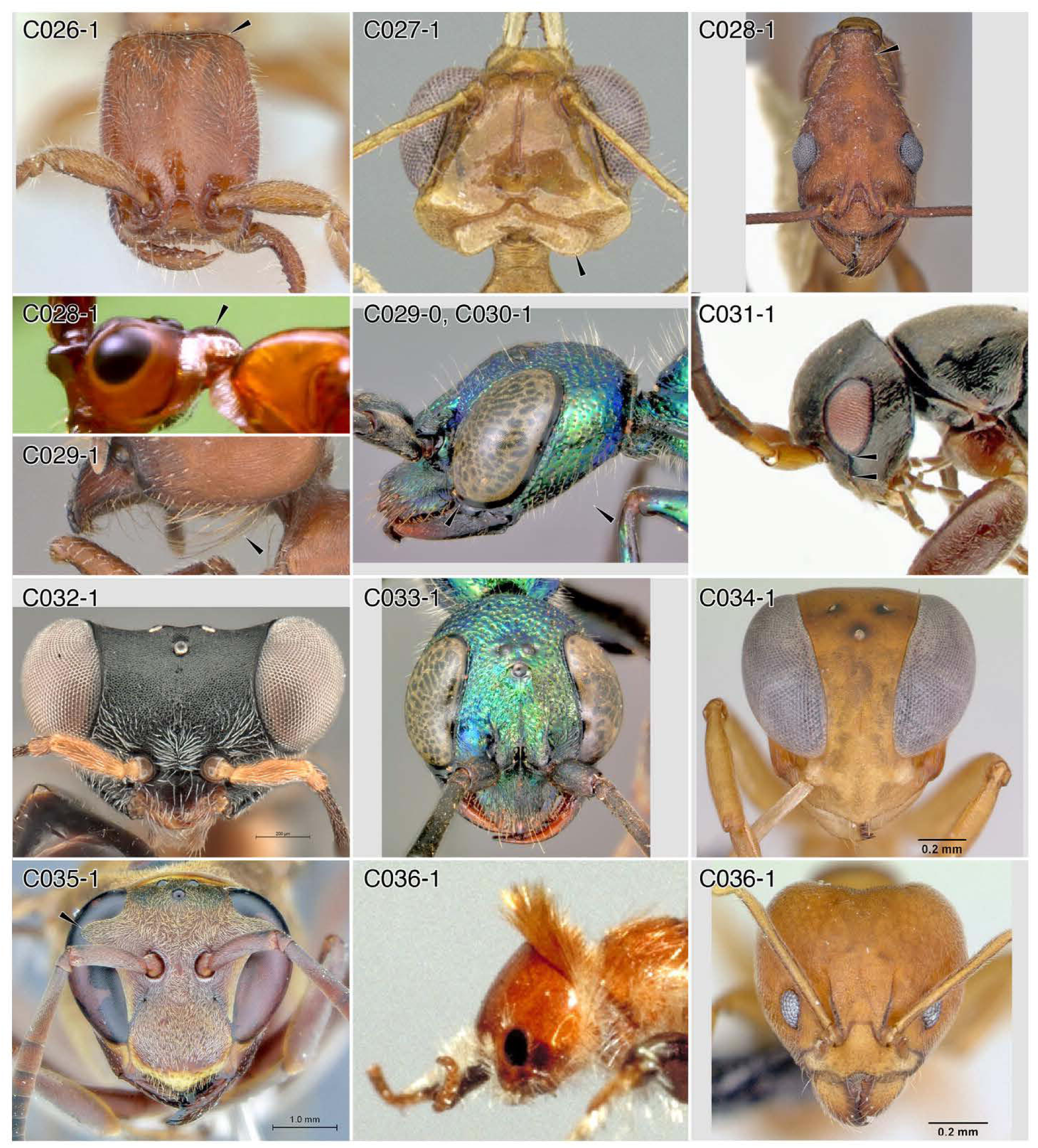
Character-states: (C26-1) Postocciput visible frontally; *Opamyrma hungvuong.* **(C27-1)** Aboral transverse lobe; *Myrmoteras bakeri.* **(C28-1)** Head posteriorly elongated; (a) *Aphaenogaster araneoides*; (b) Loboscelidiinae. **(C29-1)** Psammophore present; *Pogonomyrmex barbatus.* **(C29-0)** Psammophore absent; *Ampulex bredoi.* **(C30-1)** Eyes abutting pleurostoma; *Am. bredoi.* **(C31-1)** Eyes distant from pleurostoma; *Olixon* sp. **(C32-1)** Eyes nearly stalked; *Dryinus dentatiforceps.* **(C33-1)** Eyes converging aborally; *Am. bredoi.* **(C34-1)** Eyes converging orally; *Santschiella kohli.* **(C35-1)** Eyes emarginate; *Polistes smithii.* **(C36-1)** Eyes small; (a) *Bradynobaenus chubutinus*; (b) *Aneuretus simoni.* **Families: (26, 27, 28-1a, 29-1, 34, 36-1b)** Formicidae. **(C28-1b)** Chrysididae. **(C29-0, 30, 33)** Ampulicidae. **(C31)** Rhopalosomatidae. **(C32)** Dryinidae. **(C35)** Vespidae. **(C36-1a)** Bradynobaenidae.

**Chars. 37–44.**
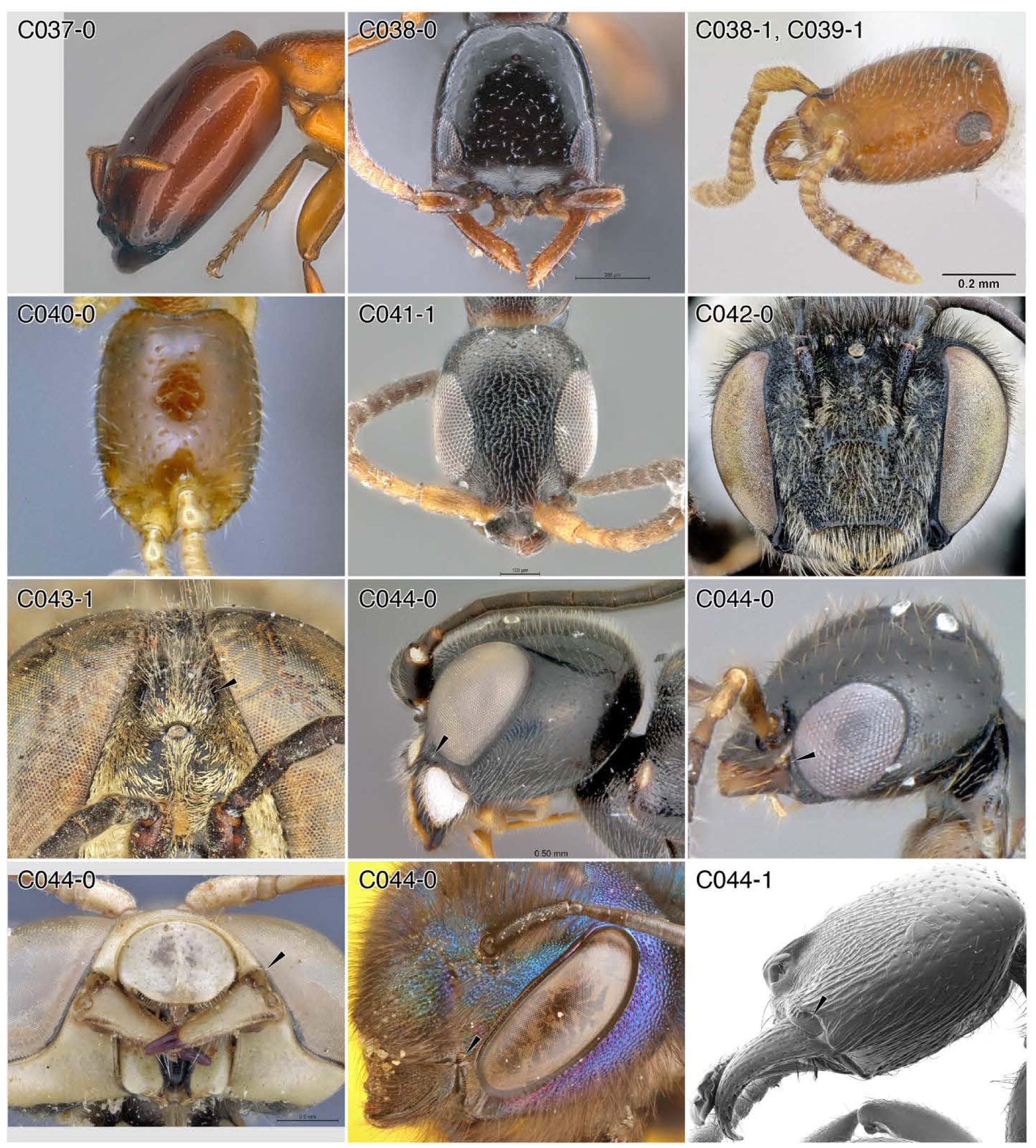
Character-states: (C37-0) Eyes not developed; *Dorylus conradti.* **(C38-0)** Eyes anterior to midlength; *Glenosema ebazileya.* **(C38-1)** Eyes oral to head midlength; *Apomyrma stygia.* **(C39-1)** Eyes extremely aboral; *Ap. stygia.* **(C40-0)** Ocelli not developed; *Dissomphalus* sp. (female). **(C41-1)** Ocelli far aboral; *Holepyris gaigherae.* **(C42-0)** Ocelli not at about midface; *Anthophora terminalis.* **(C43-1)** Ocellus/ocelli deformed; *Tachytes rarus.* **(C44-0)** Cranial condyle small; (a) *Trigonalys* sp. (female); (b) *Dissomphalus* sp. (male); (c) *Stizus somalicus*; (d) *Osmia rubifloris.* **(C44-1)** Cranial condyle enlarged; *Amblyopone australis.* **Families: (C37, 38-1, 39, 44-1)** Formicidae. **(C38-0, 40-0, 41, 44-0b)** Bethylidae. **(C42)** Apidae. **(C43)** Crabronidae. **(C44-0a)** Trigonalidae. **(C44-0c)** Bembicidae. **(C44-0d)** Megachilidae.

**Chars. 44–52.**
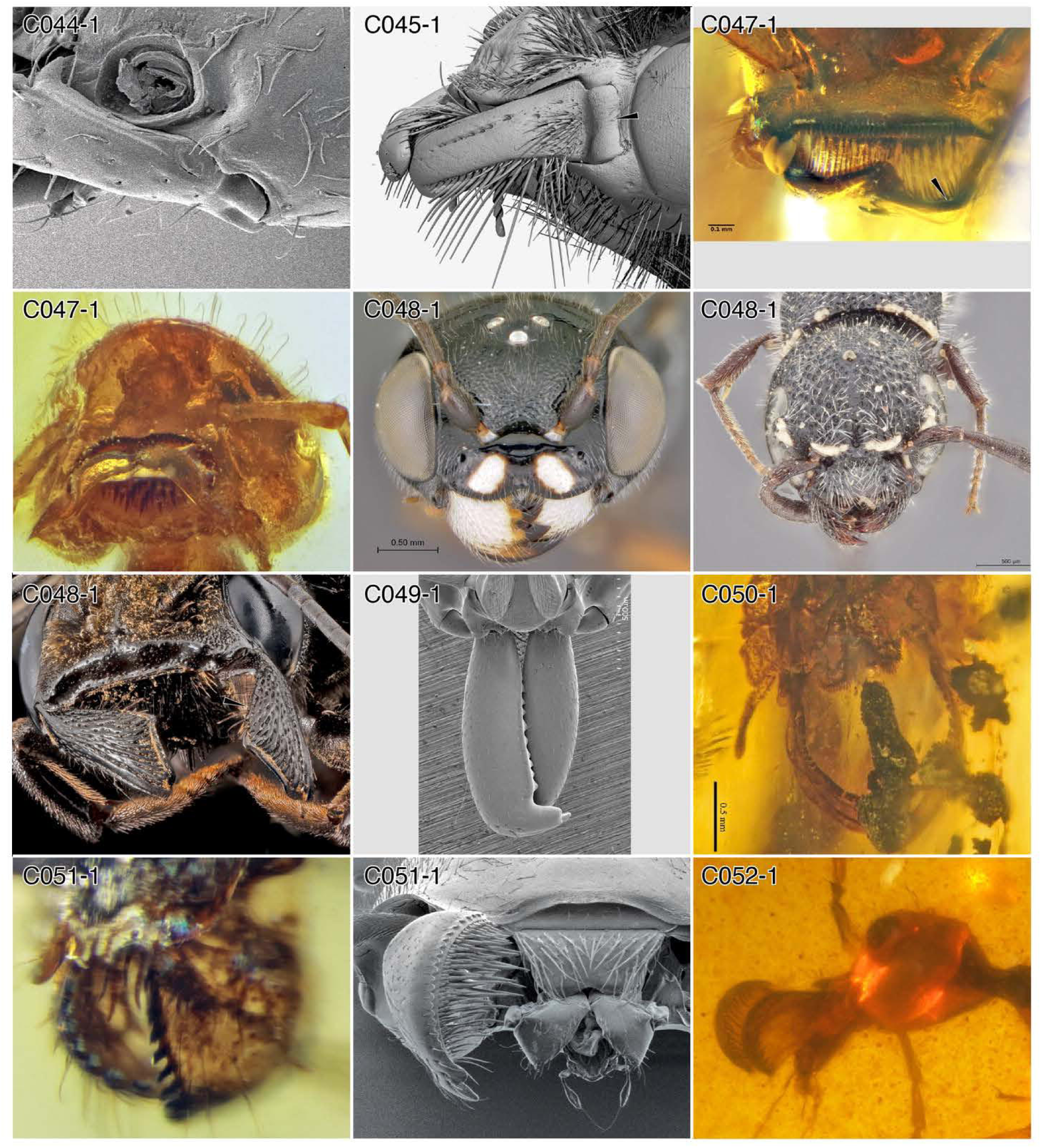
Character-states: (C44-1) Cranial condyle enlarged; *Leptanilla swani.* **(C45-1)** Lateral mandibular base cylindrical; *Scolia* sp. **(C47-1)** Mandible strongly torqued; †*Zigrasimecia.* **(C48-1)** Mandible triangular in form; (a) *Trigonalys* sp. (female); (b) *Sapygina simillina·,* (c) *Megachile* sp. **(C49-1)** Mandibles linear; *Odontomachus bauri.* **(C50-1)** Mandible elongate-curved; †*Myanmyrma gracilis.* **(C51-1)** Mandible strongly bowed; (a) †*Camelosphecia fossor,* (b) *Tatuidris tatusia.* **(C52-1)** Mandible massively enlarged; camelomeciid genus and species undefined. **Families: (C44, 47, 49, 50, 51-1b)** Formicidae. **(C45)** Scoliidae. **(C48-1a)** Trigonalidae. **(C48-1b)** Sapygidae. **(C48-1c)** Megachilidae. **(C51-1a, 52)** †@@@idae.

**Chars. 53-61.**
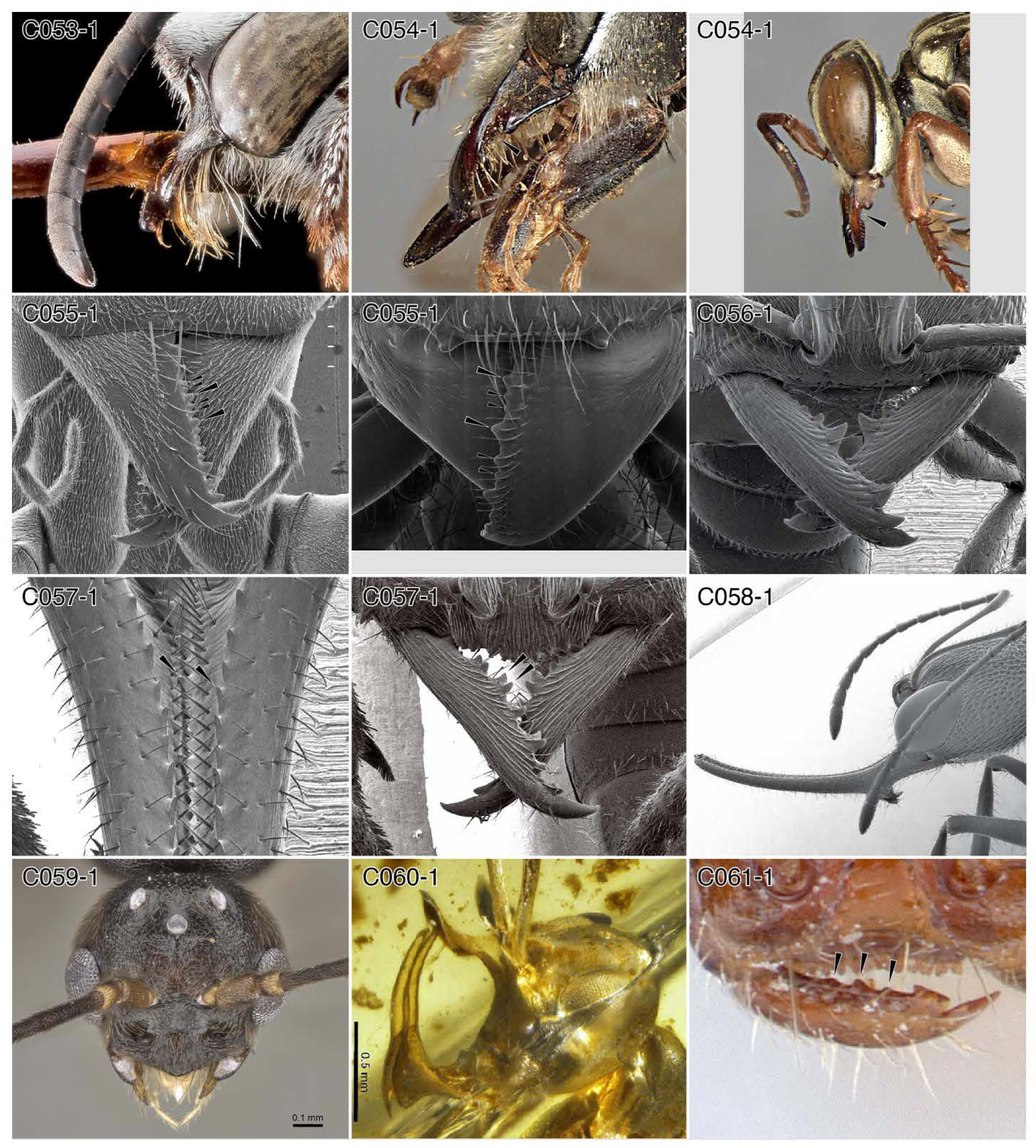
Character-states: (C53-1) Mandible digitiform in lateral view; *Thyreus* sp. **(C54-1)** Mandible abfrontally notched; (a) *Tachytes rhodesianus*; (b) *Liris bembesiana.* **(C55-1)** Mandibular teeth alternating in size; (a) *Leptomyrmex pollens*; (b) *Paraponera clavata.* **(C56-1)** Mandibular teeth fang-like; *Amblyopone australis.* **(C57-1)** Teeth in double-rows; (a) *Harpegnathos saltator,* (b) *Stigmatomma pallipes.* **(C58-1)** Mandible hastate; *Harpegnathos saltator.* **(C59-1)** Male mandible rudimentary; *Promyopias silvestrii.* **(C60-1)** Mandibular apex directed toward face; †*Linguamyrmex vladi.* **(C61-1)** Teeth on basal margin; *Opamyrma hungvuong.* **Families: (C53)** Apidae. **(C54)** Crabronidae. **(C55-61)** Formicidae.

**Chars. 62-73.**
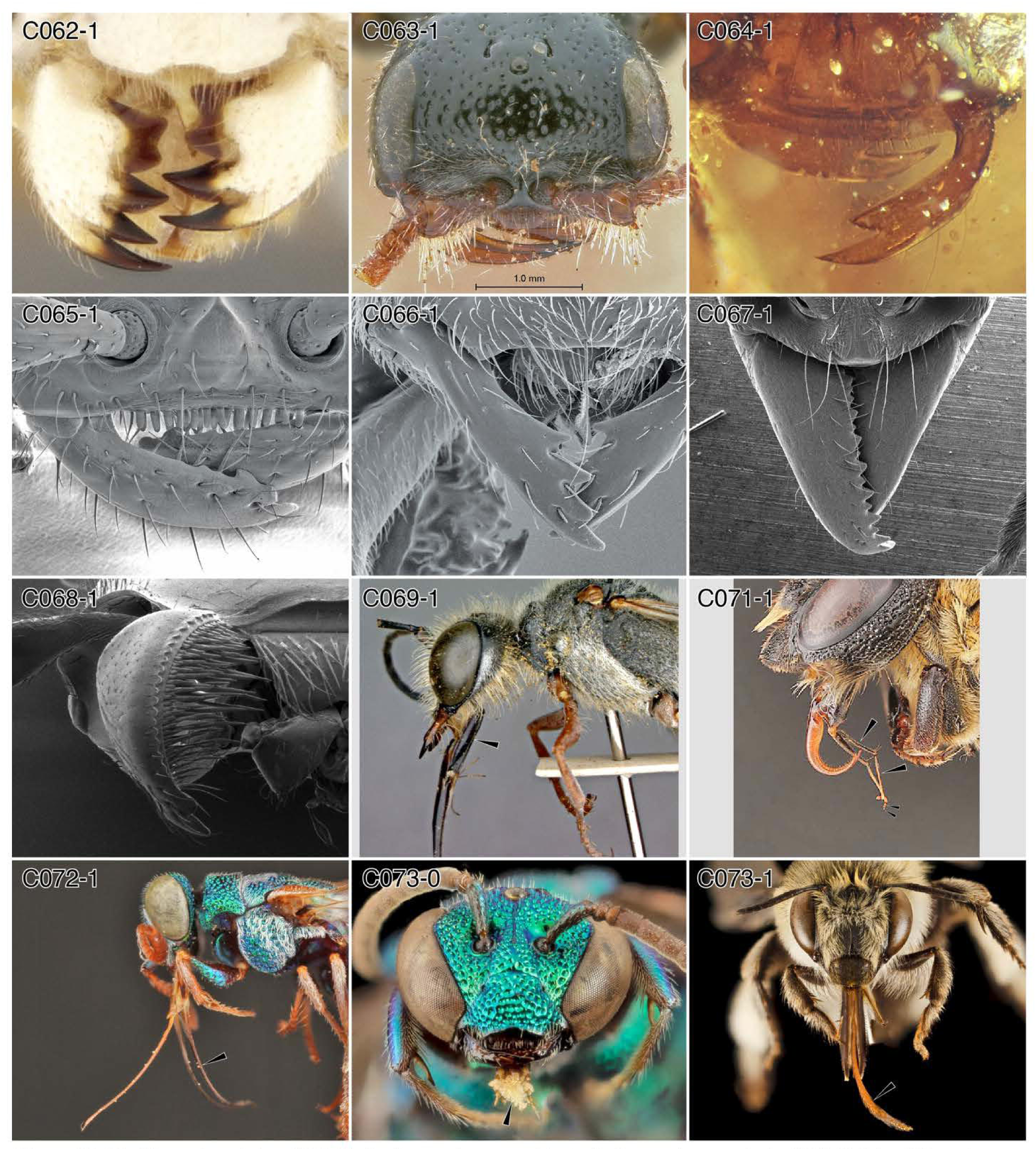
Character-states: (C62-1) Tooth count asymmetrical; *Orthogonalys maculata.* **(C63-1)** Mandible uni- or e-dentate; *Tiphia alecto* (female). **(C64-1)** Mandible bidentate; †*Gerontoformica* (male). **(C65-1)** Mandible ≥ 3-dentate; *Apomyrma stygia.* **(C66-1)** Mandible 4-9-dentate; *Acropyga* sp. **(C67-1)** Mandible ≥ 10-dentate; *Neoponera villosa.* **(C68-1)** Traction seta brush on aboral mandible surface; *Tatuidris tatusia.* **(C69-1)** Prementum elongate; *Ammophila insignis.* **(C71-1)** Proximal two labial palpomeres elongate; *Megachile sculpturalis.* **(C72-1)** Galea greatly elongate; *Parnopes* sp. **(C73-0)** Glossa short; *Temnosoma* sp. **(C73-1)** Glossa long; *Anthophora dalmatica.* **Families: (C62)** Trigonalidae. **(C63)** Tiphiidae. **(C64–68)** Formicidae. **(C69)** Sphecidae. **(C71)** Megachilidae. **(C72)** Chrysididae. **(C73-0)** Halictidae. **(C73-1)** Apidae.

**Chars. 74-86.**
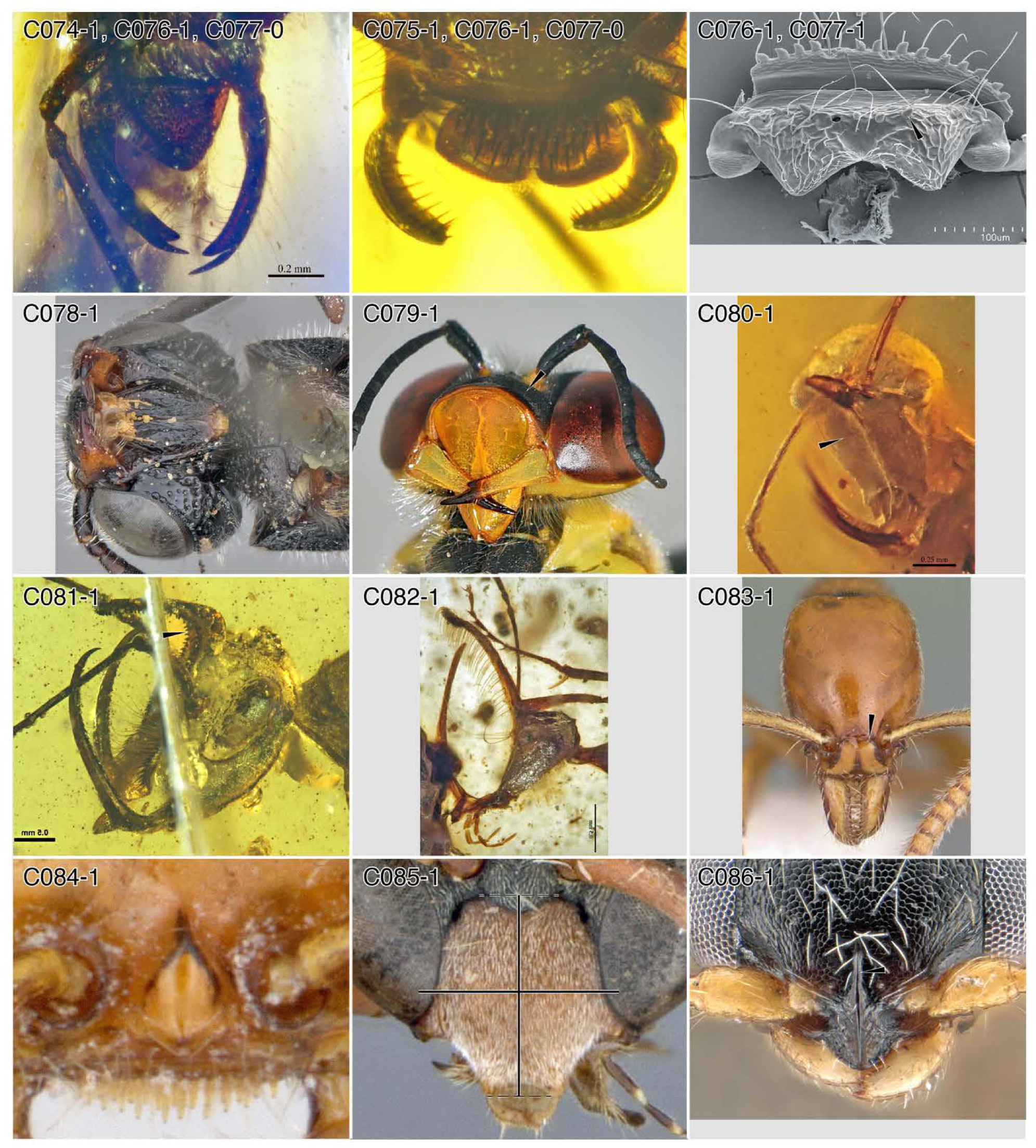
Character-states: (C74-1) Labrum massive, arc-shaped; †*Myanmyrma mauradera.* **(C75-1)** Labrum massive, bilobed; †*Zigrasimecia hoelldobleri.* **(C76-1)** Labral traction setae; (a) †*M. mauradera,* (b) †Z *hoelldobleri’,* (c) *Onychomyrmex doddi.* **(C77-0)** Labral traction setae across sclerite; (a) †*M. mauradera*; (b) †*Z. hoelldobleri.* **(C77-1)** Labral traction setae restricted in location; *O. doddi.* **(C78-1)** Oral foramen elongate; *Sapygina enderleini.* **(C79-1)** Clypeus bulbous; *Bembix* sp. **(C80-1)** Front of face concave; †*Ceratomyrmex ellenbergeri.* **(C81-1)** Oral clypeal margin migrated aborally; †*Dhagnathos autokrator.* **(C82-1)** Oral clypeal margin on long stalk; †*Ceratomyrmex ellenbergeri.* **(C83-1)** Medioclypeus raised, marginate; *Protanilla wardi.* **(C84-1)** Medioclypeus pyramidal; *Apomyrma stygia.* **(C85-1)** Medioclypeus longer than wide; *Eumenes amoldi.* **(C86-1)** Median longitudinal carina on clypeus; *Goniozus kiefferi.* **Families: (C74–77, 80–84)** Formicidae. **(C78)** Sapygidae. **(C79)** Bembicidae. **(C85)** Vespidae. **(C86)** Bethylidae.

**Chars. 87-95.**
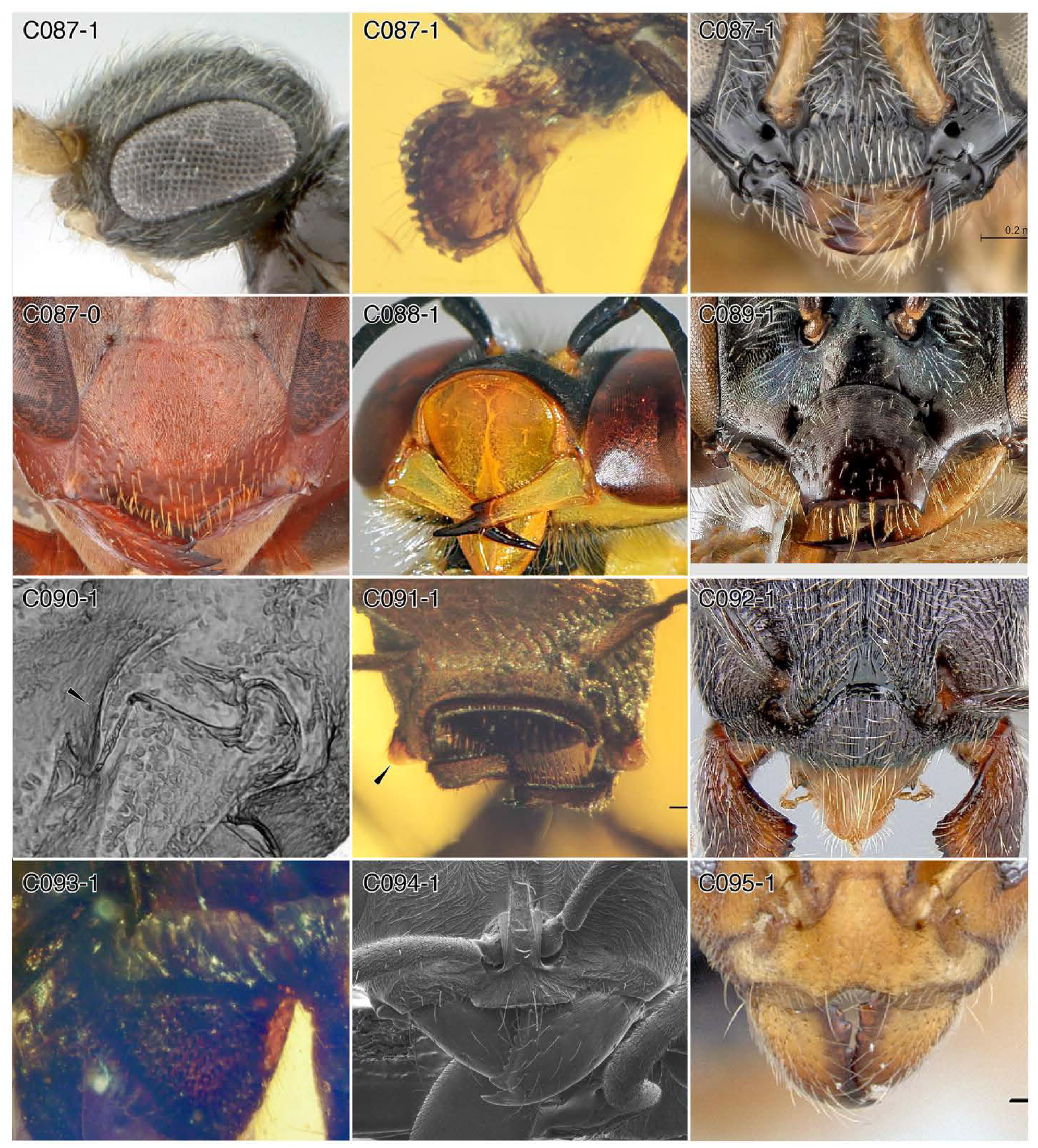
Character-states: (C87-1) Medioclypeus produced orally; (a) *Sclerogibba turneri*; (b) †*Camelosphecia fossor,* (c) *Probethylus callani.* **(C87-0)** Medioclypeus not produced orally; *Polistes metricus.* **(C88-1)** Clypeus curved around labral base; *Bembix* sp. **(C89-1)** Clypeus and face produced orally; *Perdita halictoides.* **(C90-1)** Lateroclypeal lobes; †*Gerontoformica pilosa.* **(C91-1)** Lateroclypeal lobes shield-like; †*Zigrasimecia ferox.* **(C92-1)** Oral clypeal margin convex; *Monica bradleyi.* **(C93-1)** Oral clypeal margin linear; †*Myanmyrma gracilis.* **(C94-1)** Oral-median clypeal lobe; *Pseudomyrmex gracilis.* **(C95-1)** Oral-median clypeal notch with lateral lamellae; *Aneuretus simoni.* **Families: (C87-1a, c)** Sclerogibbidae. **(C87-1b)** †@@@idae. **(C87-0)** Vespidae. **(C88)** Bembicidae. **(C89)** Andrenidae. **(C90-95)** Formicidae.

**Chars. 96–106.**
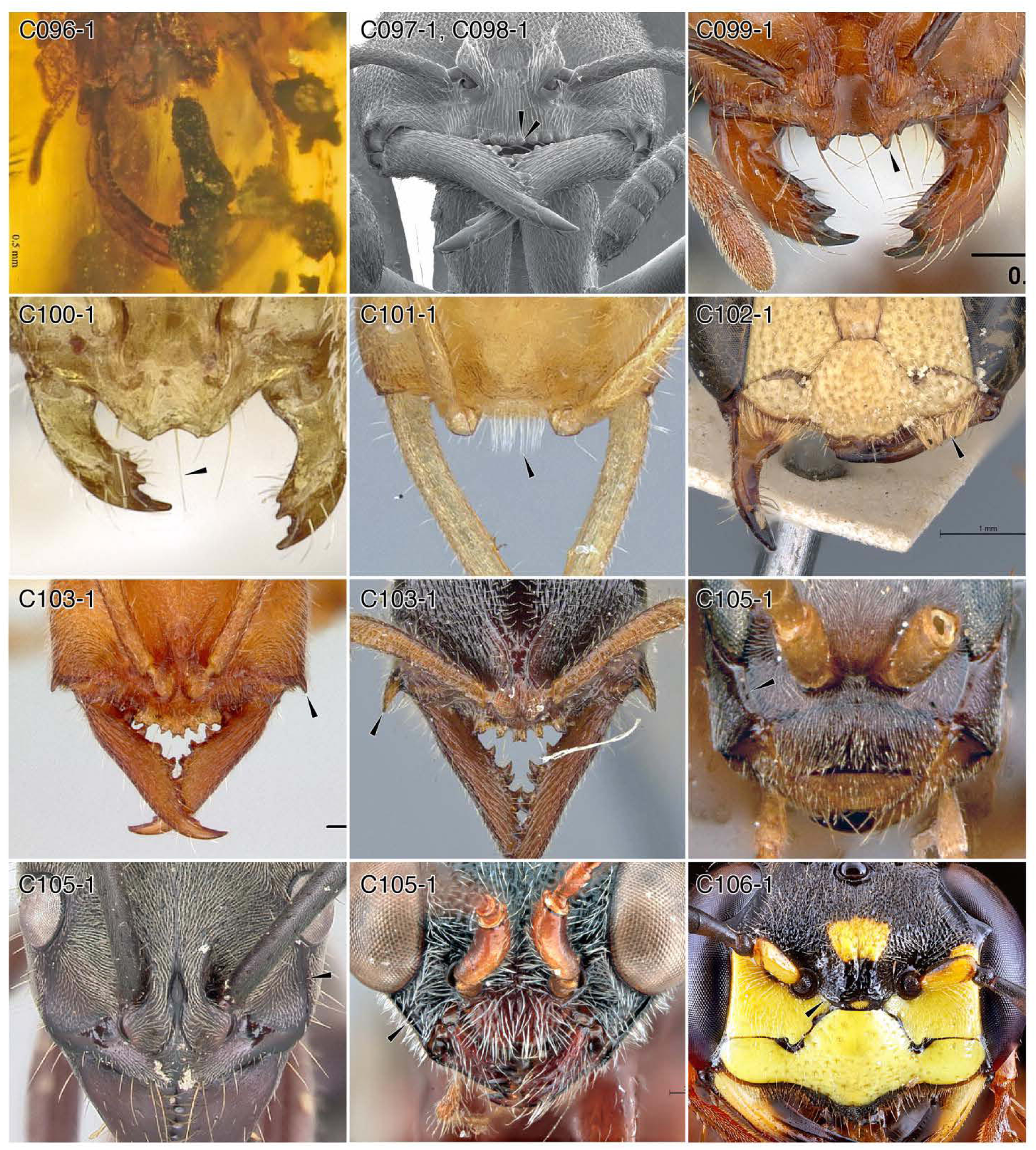
Character-states: (C96-1) Clypeus and labrum medially notched; †*Myanmyrma gracilis.* **(C97-1)** Clypeal traction setae; *Stigmatomma pallipes.* **(C98-1)** Clypeal traction setae on raised prominences; *St pallipes.* **(C99-1)** Medioclypeus with paired denticles; *Solenopsis xyloni.* **(C100-1)** Unpaired median clypeal seta; *Monomorium anderseni.* **(C101-1)** Median clypeal seta brush; *Martialis heureka.* **(C102-1)** Male lateroclypeal brushes; *Cerceris amoldi.* **(C103-1)** Each gena armed with small to large spiniform process; (a) *Fulakora armigera*; (b) *F. gracilis.* **(C105-1)** Malar ridge/sulcus; (a) *Olixon saltator,* (b) *Neoponera villosa,* (c) *Lonchodryinus madagascolus.* **(C106-1)** “Subantennal line”; *Philanthus gibbosus.* **Families: (C96–101, 103, 105-1b)** Formicidae. **(C102, 106)** Philanthidae. **(C105-1a)** Rhopalosomatidae. **(C105-1c)** Dryinidae.

**Chars. 106–117.**
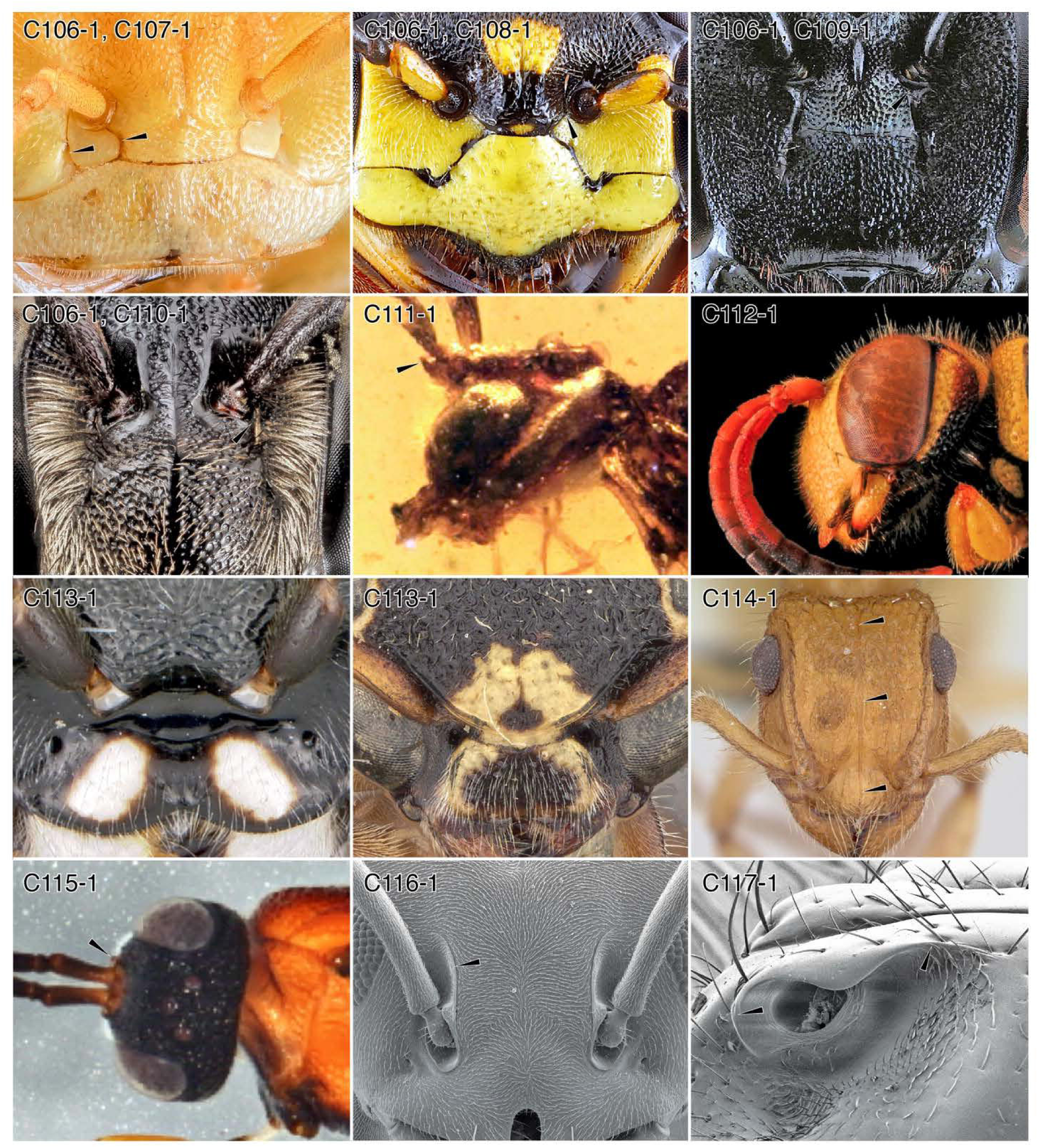
Character-states: (C106-1) “Subantennal sutures” present. **(C107-1)** “Subantennal sutures” paired; *Arhysosage* sp. **(C108-1)** “Subantennal suture” aboral terminus mediad torulus mid-width; *Philanthus gibbosus.* **(C109-1)** “Subantennal suture” aboral terminus at torulus mid-width; *Xylocopa virginica.* **(C110-1)** “Subantennal suture” aboral terminus laterad torulus mid-width *Euaspis* sp. **(C111-1)** Medial facial process; †*Xenodellitha preta.* **(C112-1)** Face broadly convex; *Cerceris triangulata.* **(C113-1)** Face raised aborad toruli; (a) *Trigonalys* sp. (female); (b) *Sapygina undulata.* **(C114-1)** Medial facial carina; *Acanthoponera minor.* **(C115-1)** Lateral frontal carinae; Evaniidae: *Evaniella semaeoda.* **(C116-1)** Medial frontal carinae; *Tapinoma simrothi.* **(C117-1)** Torulus and frontal carina undifferentiated; †*Centromyrmex brachycola.* **Families: (C107)** Andrenidae. **(C108, 112)** Philanthidae. **(C109)** Apidae. **(C110)** Megachilidae. **(C111)** †Othniodellithidae. **(C113-1a)** Trigonalidae. **(C113-1b)** Sapygidae. (015) Evaniidae. **(C114, 116 117)** Formicidae.

**Chars. 118-129.**
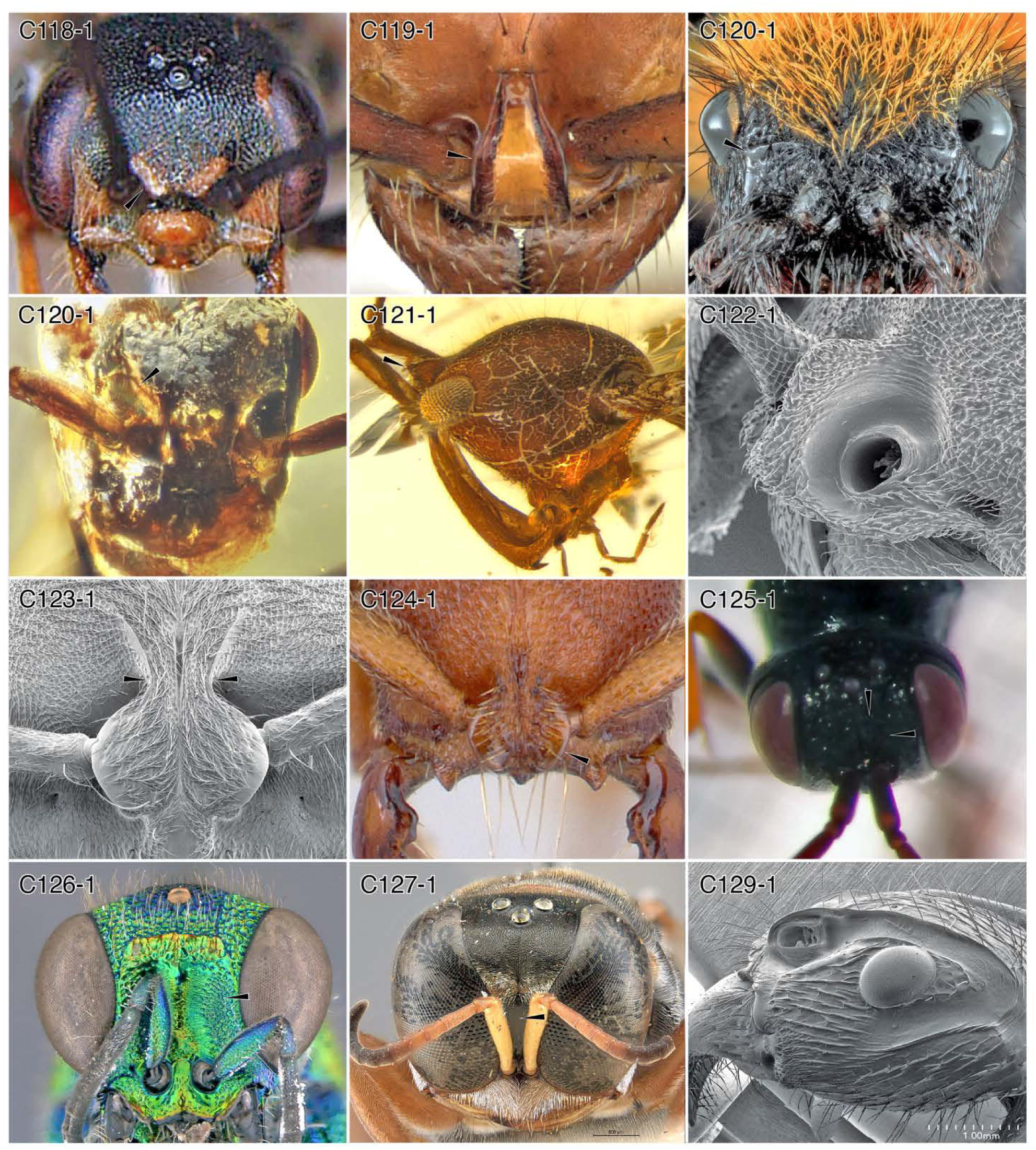
Families: (C118) Thynnidae. **(C119, 120-1b, 121–124, 129)** Formicidae. **(C120-1a)** Mutillidae. **(C125)** Evaniidae. **(C126)** Chrysididae. **(C127)** Crabronidae. Character-states: **(C118-1)** Medial frontal carinae projecting frontad, more-or-less lateromedially oriented; *Upa porteri.* **(C119-1)** Medial frontal carinae projecting frontad, more-or-less anteroposteriorly oriented; *Lioponera clara.* **(C1**20-1) Medial frontal carinae semicircular; (a) mutillid genus and species indet.; (b) †*Gerontoformica orientalis* group. **(C121-1)** Medial pyramid; †*Haidomyrmex davidbowiei.* **(C122-1)** Medial frontal carinae medially fused; *Discothyrea testacea.* **(C123-1)** Medial frontal carinae aborally “pinched”; *Pseudoponera stigma.* **(C124-1)** Medial frontal carinae on clypeus; *Feroponera ferox.* **(C125-1)** Aboral-medial antennal scrobe on face; genus and species indet. **(C126-1)** Aboral-medial antennal scrobe longer than wide, large; *Chrysis eximia.* **(C127-1)** Aboral-medial antennal scrobe longer than wide, small; *Crossocerus glabricornis.* **(C129-1)** Antennal scrobe receiving scape and flagellum separately; *Paraponera clavata.*

**Chars. 130-138.**
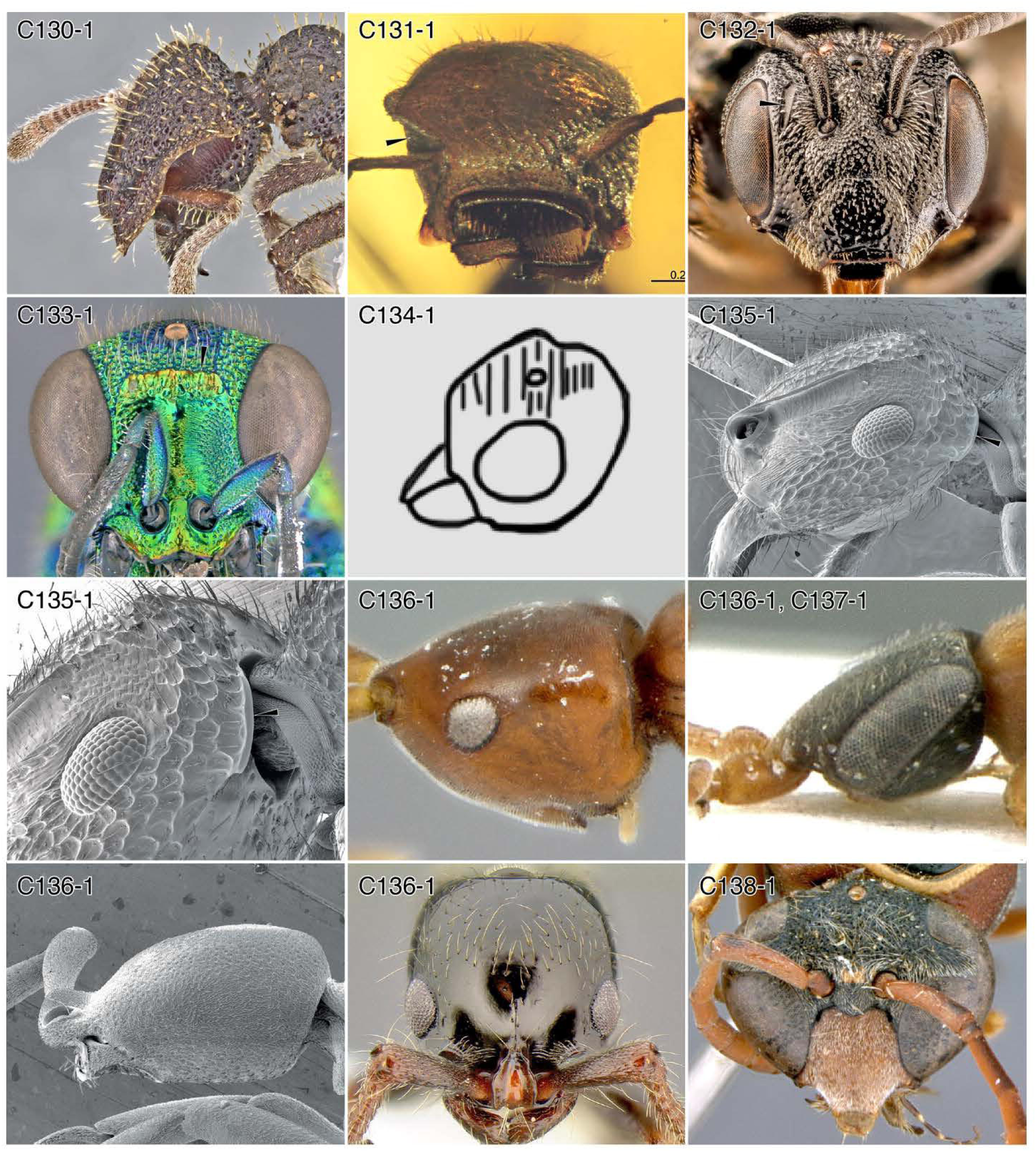
Character-states: (C130-1) Antennal scrobe receiving scape and flagellum together; *Stegomyrmex olindae.* **(C131-1)** Antennal scrobes extending to eyes; †*Zigrasimecia ferox.* **(C132-1)** Facial foveae; *Pseudopanurgus rugosus.* **(C133-1)** Transverse carina orad ocelli; *Chrysis eximia.* **(C134-1)** Face with transverse rugosity; †*Andrenelia pinnata* after Rasnitsyn & Martínez-Delclòs (2000). **(C135-1)** Posterolateral head lobes; *Acanthoponera minor.* **(C136-1)** Antennal toruli produced anterad past mandibles; (a) *Embolemus africanus;∖* (b) *Sclerogibba talpiformis;* (c) *Probolomyrmex guineensis;* (d) *Lioponera larvata.* **(C137-1)** Face produced anterad, toruli directed aborad; *Sc. talpiformis.* **(C138-1)** Toruli at about middle of face; Vespidae: *Eumenes amoldi.* **Families: (C130, 131, 135, 136)** Formicidae. **(C132)** Andrenidae. **(C133)** Chrysididae. **(C134)** †Andreneliidae. **(C136-1a)** Embolemidae. **(C136-1b, 137)** Sclerogibbidae. **(C138)** Vespidae.

**Chars. 139-147.**
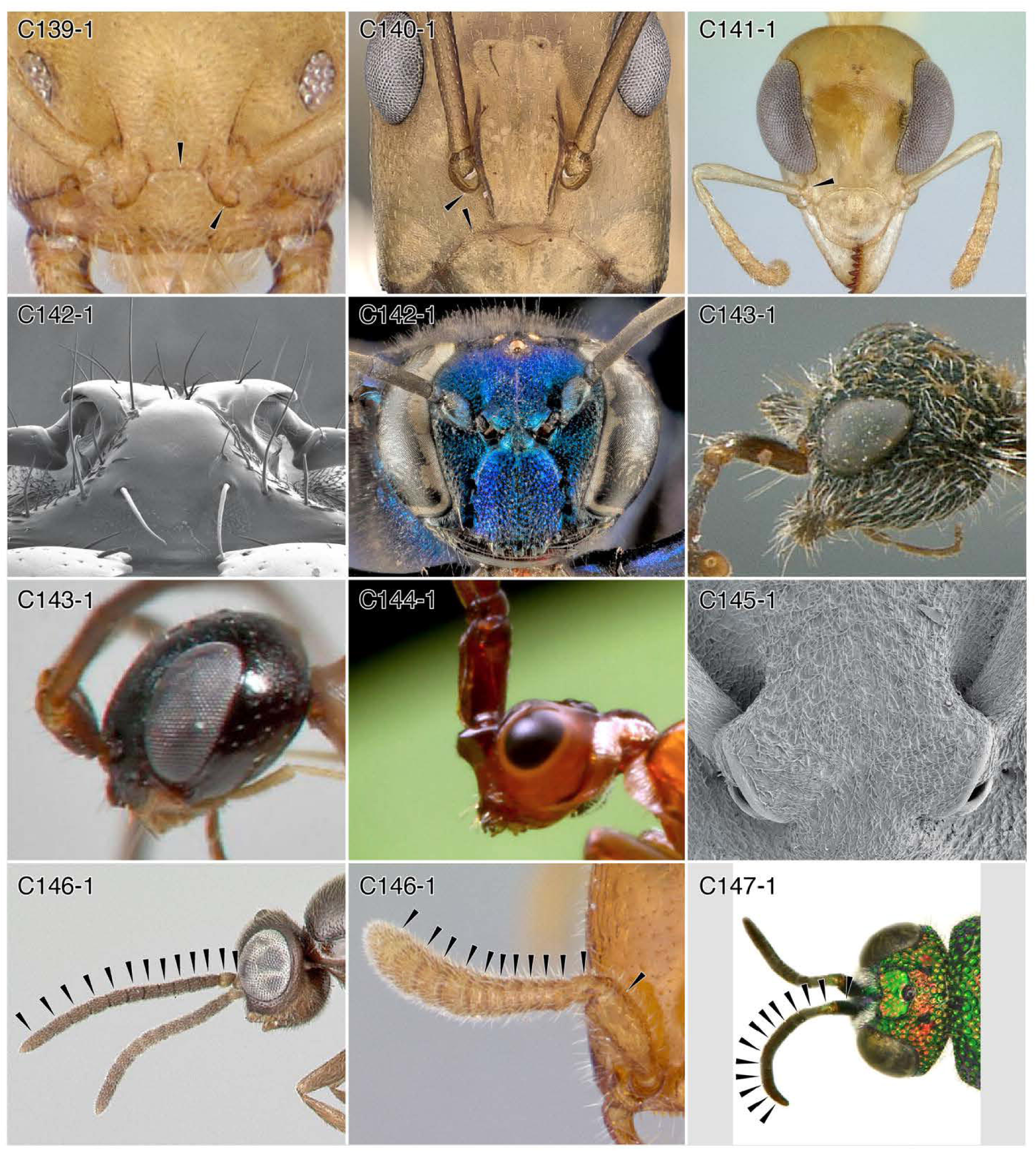
Families: Character-states: (C139-1) Clypeus extending between toruli; *Nebothriomyrmex majeri.* **(C140-1)** Toruli distant from clypeus; *Camponotus aegyptiacus.* **(C141-1)** Clypeus orad eyes; *Gesomyrmex* th0l. **(C142-1)** Toruli directed laterally; (a) *Centromyrmex brachycola;* (b) Sphecidae: *Chalybion californicum.* **(C143-1)** Toruli directed aborad; (a) *Amoldtia bischoffi,* (b) *Heterogyna* sp. **(C144-1)** Toruli directed relatively aborad; *Loboscelidia* sp. **(C145-1)** Toruli largely concealed by “frontal lobes”; *Platythyrea tumeri.* **(C146-1)** Antennomere count sexual dimorphism (13 male, 12 female); (a) *Apomyrma* cf0l (male); (b) *Ap. stygia* (female). **(C147-1)** Antennomere count ≥ 13 in both sexes; *Odontochrydium bicristatum* (female). **(C139-141, 142-1a, 145, 146)** Formicidae. **(C143-1a)** Mutillidae. **(C134-1b)** Heterogynaidae. **(C144, 147)** Chrysididae.

**Chars. 148-153.**
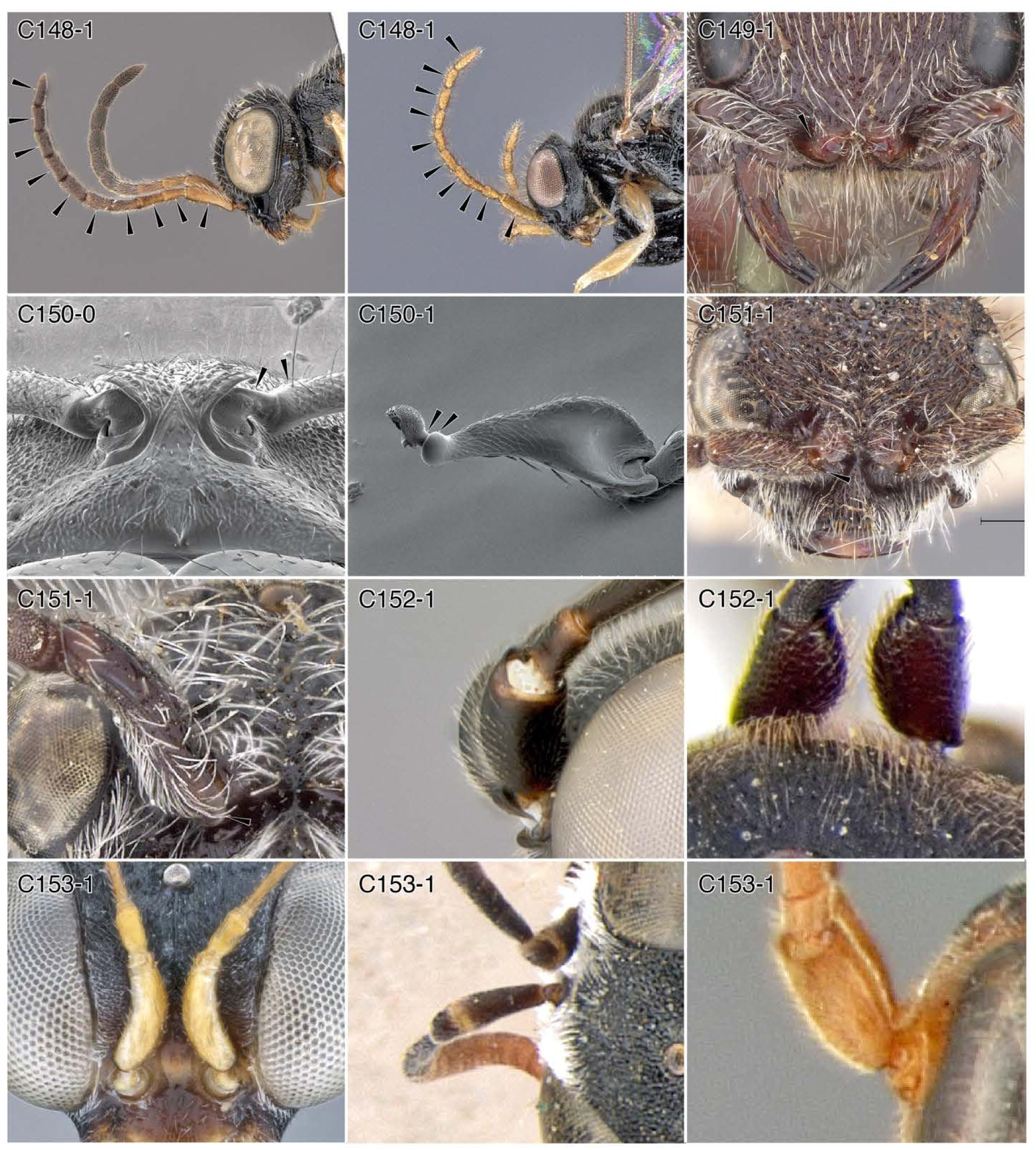
Character-states: (C148-1) Antennomere count 10; (a) *Anteon alteri* (female); (b) *A. malagasy* (male). **(C149-1)** Radicle and scape angled; *Cephalotilla ceratophora.* **(C150-0)** Radicle and scape slightly angled; *Ponera alpha.* **(C150-1)** Radicle and scape strongly angled; *Tatuidris tatusia.* **(C151-1)** Scape with flange; (a) *Mutilla katanga;* (b) *Bisulcotilla umhalali.* **(C152-1)** Scape bulbous medially; (a) Trigonalidae: *Trigonalys* sp.; (b) Aulacidae: *Pristaulacus africanus.* **(C153-1)** Scape length ≥ 2 x width; (a) *Dryinus basilewskyi;* (b) Rhopalosomatidae: *Paniscomima bekilyi;* (c) Crabronidae: *Oxybelus acutissimus.* **Families: (C148, 153)** Dryinidae. **(C149, 151-a, b)** Mutillidae. **(C150)** Formicidae. **(C152-1a)** Trigonalidae. **(C152-1b)** Aulacidae. **(C153-1b)** Rhopalosomatidae. **(C153-1c)** Crabronidae.

**Chars 154–162.**
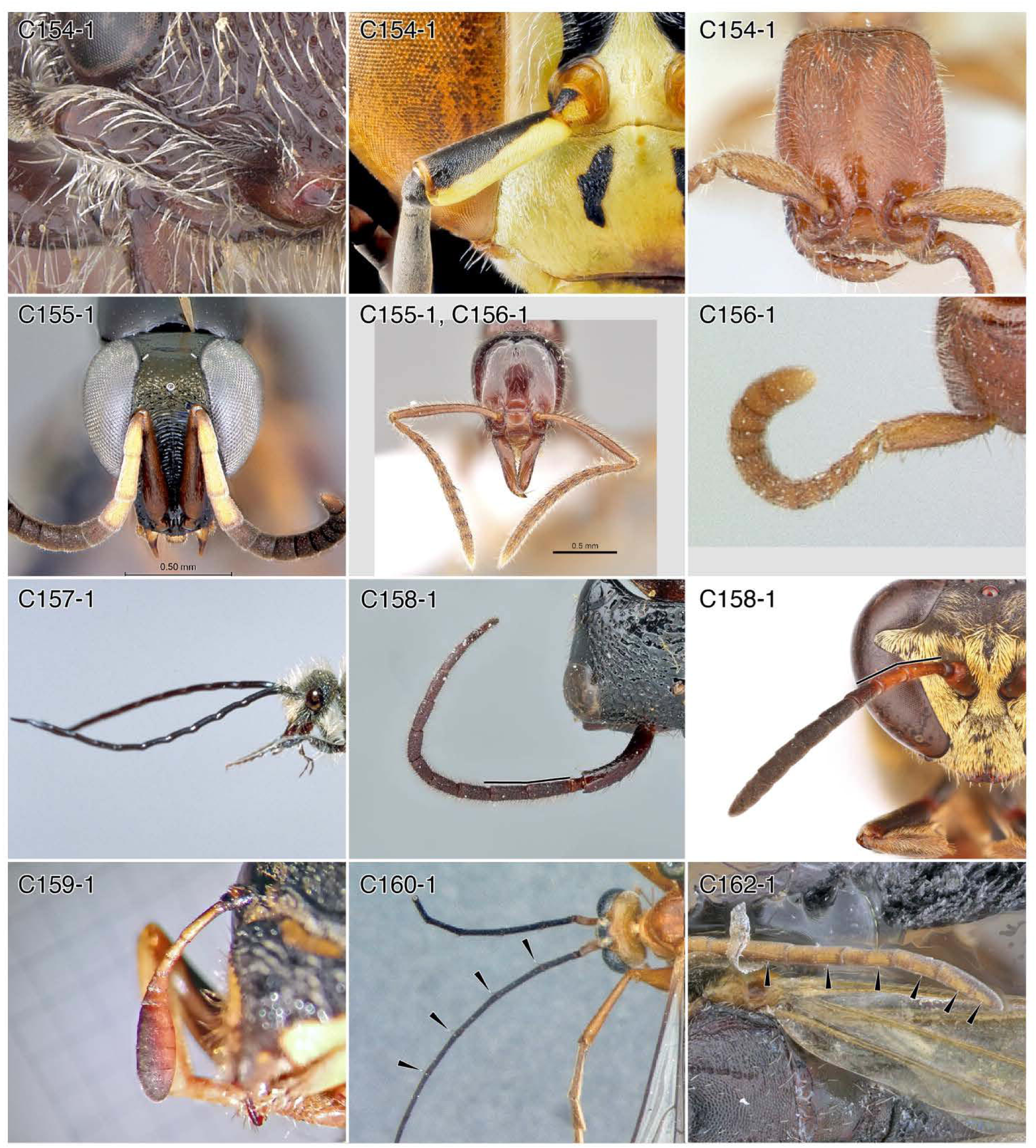
Character-states: (C154-1) Scape length ≥ 4 x width; (a) *Cephalotilla ceratophora;* (b) *Bembix* sp.; (c) *Opamyrma hungvuong.* **(C155-1)** Scape length > 1/2 head width; (a) *Obenbergerella aenigmatica;* (b) *Anomalomyrma helenae.* **(C156-1)** Scape length > funiculus (pedicel and flagellum) length; (a) *A. helenae;* (b) *Opamyrma hungvuong;* note that the scape appears slightly yet proportionally long because of flagellar-scapal angle of this specific preserved specific. **(C157-1)** Flagellum enlarged with shining smooth surface; bradynobaenid genus and species indet. **(C158-1)** Antennomere III length > IV; (a) *Pristocera maximum-,* (b) *Trypoxylon texense.* **(C159-1)** Flagellum with ovate club; *Pseudomasaris vespoides.* **(C160-1)** Flagellomeres with apical seta pairs; *Paniscomima bekilyi.* **(C162-1)** “Antennal dorsal organs”; *Dryinus basilewskyi.* **Families: (C154-1a)** Mutillidae. **(C154-1b)** Bembicidae. **(C154-1c, 155-1b, 156)** Formicidae. **(C155-1a)** Chrysididae. **(C157)** Bradynobaenidae. **(C158-1a)** Bethylidae. **(C158-1b)** Pemphredonidae. **(C159)** Vespidae. **(C160)** Rhopalosomatidae. **(C162)** Dryinidae.

**Chars. 163-170.**
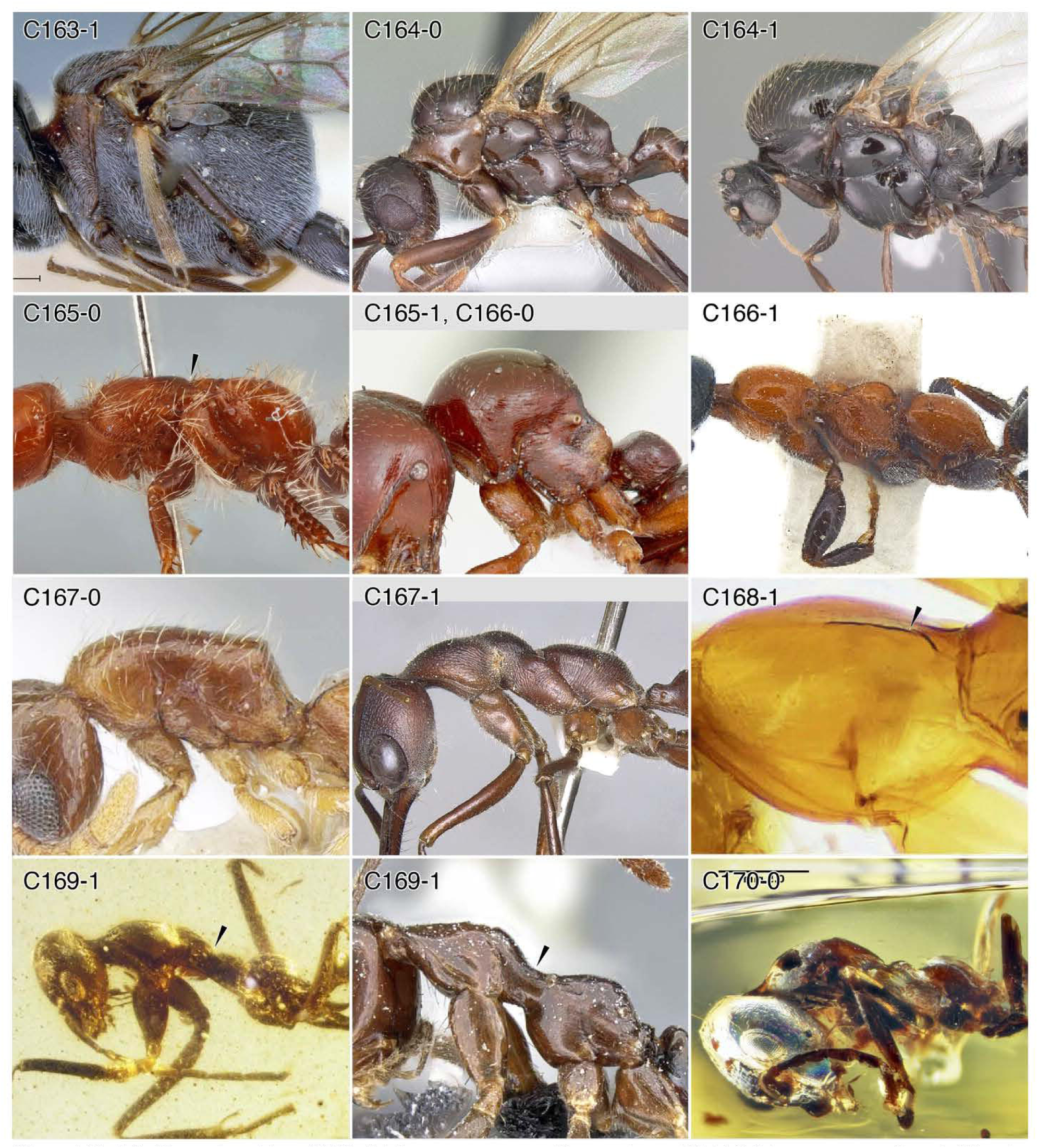
Character-states: (C163-1) Mesosoma compact; *Zeuxevania* sp. **(C164-0)** Mesosoma not enlarged; *Myrmica rubra.* **(C164-1)** Mesosoma enlarged, swollen-appearing; *Solenopsis invicta.* **(C165-0)** Promesonotal articulation mobile; *Braunsomeria albokirta.* **(C165-1)** Promesonotal articulation immobile; *Tatuidris tatusia.* **(C166-0)** Mesosomal profile not diagonal; *T. tatusia.* **(C166-1)** Mesosomal profile diagonal; *Methocha litoralis.* **(C167-0)** Promesonotum not raised above propodeum; *Lioponera longitarsus.* **(C167-1)** Promesonotum raised above propodeum; *Myrmecia browningi.* **(C168-1)** Pronotum with median carina; *Heterogyna nocticola.* **(C169-1)** Mesothorax constricted, “strangled”; (a) †*Gerontoformica gracilis;* (b) *Prenolepis jerdoni.* **(C170-0)** Mesoscutum without keel; †*Gerontoformica orientalis group.* **Families: (C163)** Evaniidae. **(C164, 165-1, 166-0, 167, 169, 170)** Formicidae. **(C165-0)** Thynnidae. **(C166)** Tiphiidae. **(C168)** Heterogynaidae.

**Chars. 170-177.**
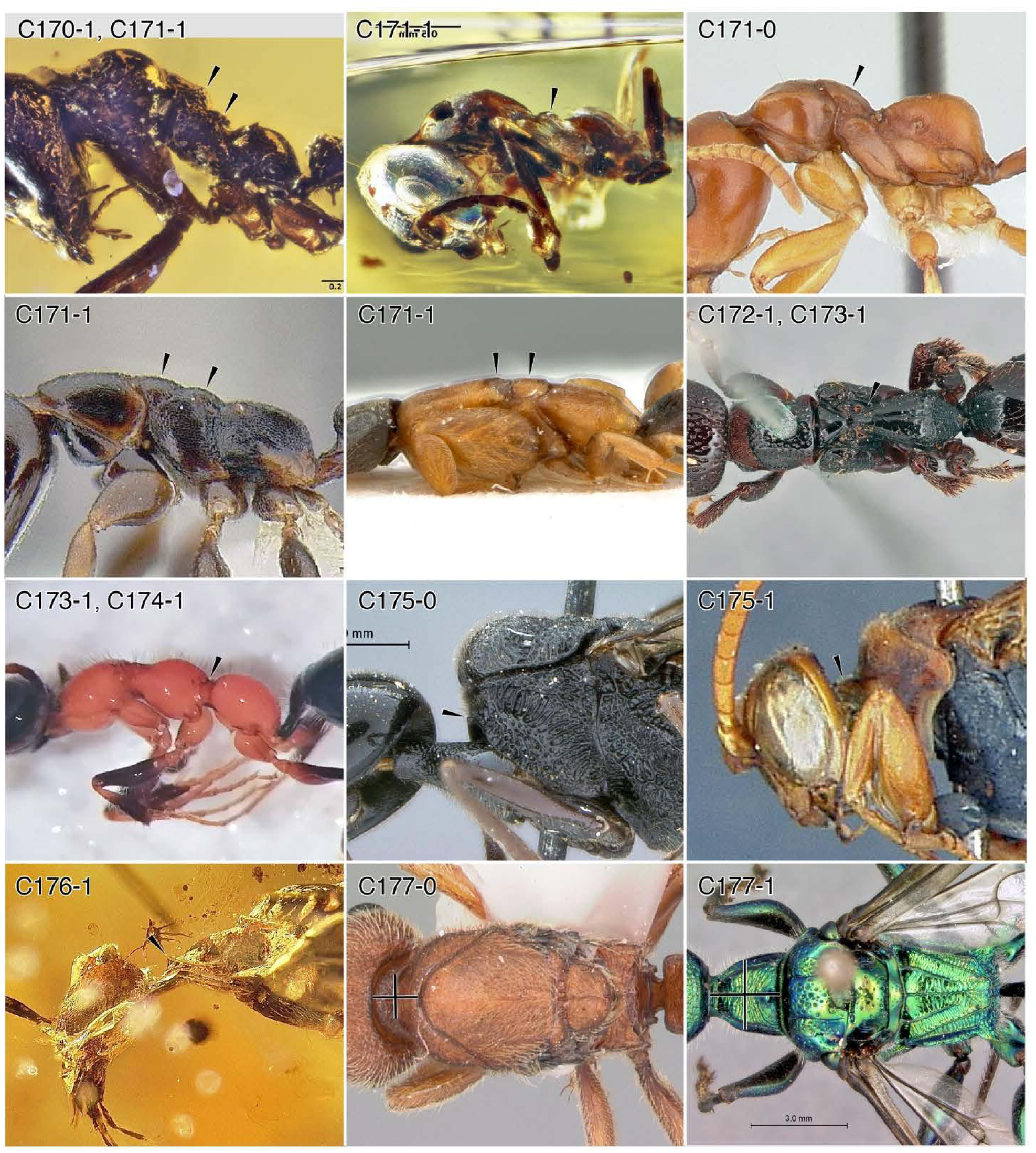
Character-states: (C170-1) Mesoscutum transversely keeled; †*Gerontoformica pilosa* group. **(C171-0)** Mesonotum with single bulge; *Tetraponera merita.* **(C171-1)** Mesoscutum with two bulges; (a) †G. *pilosa* group; (b) †G. *orientalis* group. (c) *Tetraponera continua;* (d) *Sclerogibba berlandi.* **(C172-1)** Mesometanotal articulation mobile. **(C173-1)** Mesometanotal articulation, taxic (Bethylidae); *Pristocera incerta.* **(C174-1)** Mesometanotal articulation, taxic (Tiphiidae); *Methocha*(*Dryinopsis*) sp. **(C175-0)** Pronotal collar/neck absent; *Pristaulacus pilotoi.* **(C175-1)** Pronotal collar/neck present; *Ceropales scobinifera.* **(C176-1)** Pronotum anterior rim upturned; †*Camelomecia?* (male). **(C177-0)** Pronotum small, weak; *Proceratium croceum.* **(C177*-***1) Pronotum large, strong; *Ampulex bredoi.* **Families: (C170, 171-0, la, 177)** Formicidae. **(C171-1d)** Sclerogibbidae. **(C173)** Bethylidae. **(C174)** Tiphiidae. **(C175-0)** Aulacidae. **(C175-1)** Pompilidae. **(C176)** †@@@idae. **(C177)** Ampulicidae.

**Chars. 177-187.**
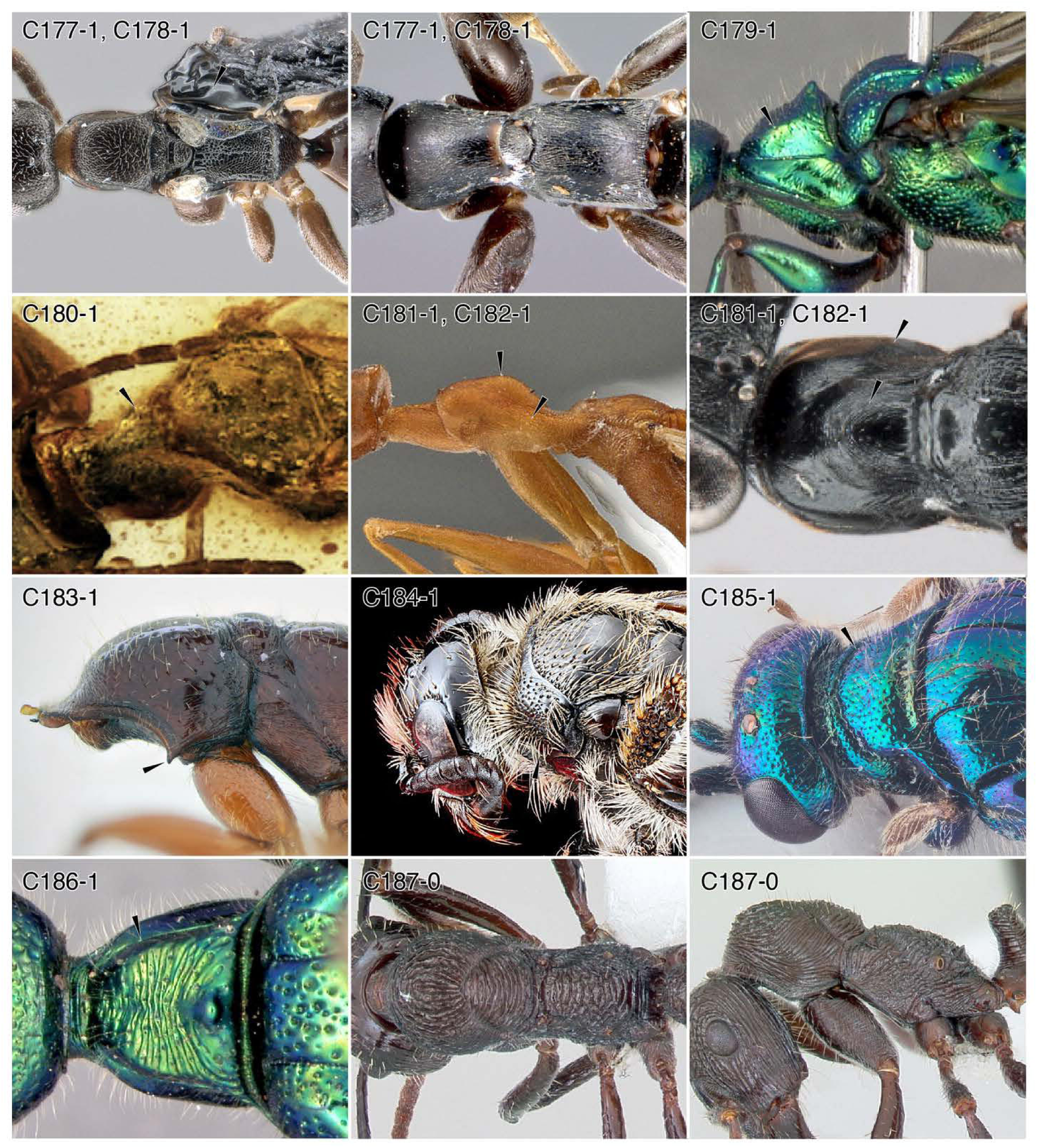
Character-states: (C177-1) Pronotum large, strong; (a) *Holepyris gaigherae;* (b) *O. dentatum.* **(C178-1)** Pronotum rectangular; (a) *Ho. gaigherae;* (b) *O. dentatum.* **(C179-1)** Pronotum with dorsolateral margination; *Ampulex bredoi.* **(C180-1)** Pronotal disc triangular; †*Hybristodryinus shan.* **(C181-1)** Pronotum with median bulge; (a) *Dryinus spangleri;* (b) *D. nigrithorax.* **(C182-1)** Pronotum with expanded lateral collar; (a) *D. spanglerr,* (b)*D*. *nigrithorax.* **(C183-1)** Anteroventral pronotal spiniform process; *Amblyopone australis.* **(C184-1)** Pronotum transverse line/groove; scoliid genus and species indet. **(C185-1)** Deep pronotal sulcus completely arcing around pronotum; *Cleptes semiauratus.* **(C186-1)** Shallow pronotal sulcus completely arcing around pronotum; *Amp. bredoi.* **(C187-0)** Pronotum without scrobe receiving leg; (a) *Ectatomma ruidum,* (b) *E. ruidum.* **Families: (C177-1a, 178-1a)** Bethylidae. **(C177-1b, 178-1b)** Rhopalosomatidae. **(C179, 186)** Ampulicidae. **(C180-182)** Dryinidae. **(C183, 187)** Formicidae. **(C184)** Scoliidae. **(C185)** Chrysididae.

**Chars. 187-195.**
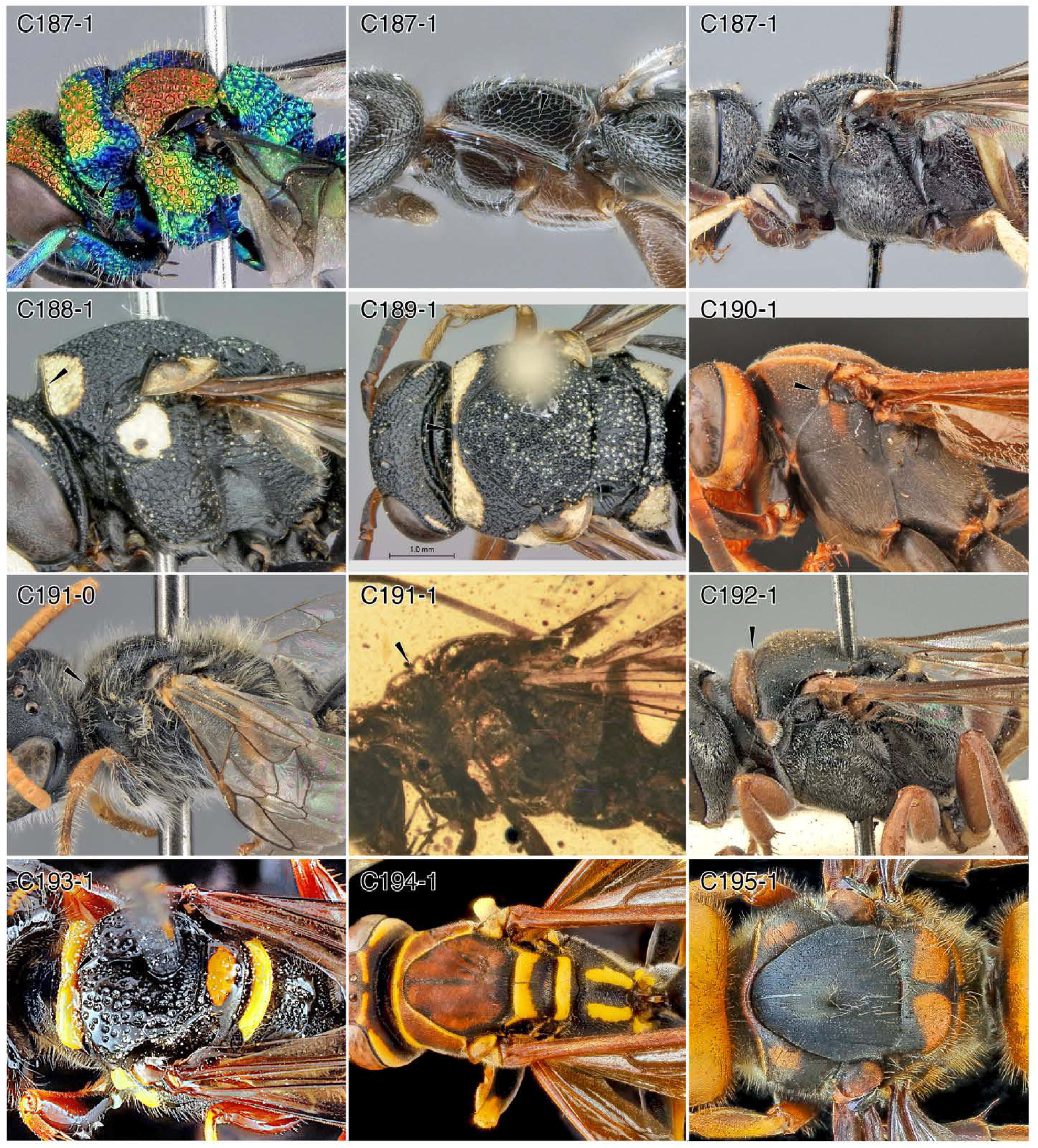
Character-states: (C187-1) Pronotum laterally scrobiculate, receiving profemur; (a) *Chrysis splendens;* (b) *Holepyris gaigherae;* (c) *Cerceris amatoria.* **(C188-1)** Pronotum with lateral, dorsoventrally oriented carina; *Knemodynerus* sp. **(C189-1)** Pronotum with transverse, dorsal carina; *Knemodynerus* sp. **(C190-1)** Pronotum with dorsoventrally oriented carina just anterad tegulum; *Mischocyttarus flavitarsis.* **(C191-0)** Pronotum not constricted posteriorly; *Scrapter luridus.* **(C191-1)** Pronotum posteriorly constricted; †*Camelosphecia venator.* **(C192-1)** Pronotum separated from mesoscutum by apparent incision; *Crossocerus glabricornis.* **(C193-1)** Posterior pronotal portion with dorsomedian notch; philanthid genus and species indet. **(C194-1)** Posterior pronotal margin long and narrowly arcuate; *Polistes* sp. **(C195-1)** Lateral pronotal portions broad; *Vespa mandarina.* **Families: (C187-1a)** Chrysididae. **(C187-1b)** Bethylidae. **(C187-1c, 193)** Philanthidae. **(C188-190, 194, 195)** Vespidae. **(C191)** Colletidae. **(C191)** †@@@idae. **(C192)** Crabronidae. **(C193)** Philanthidae.

**Chars. 196-206.**
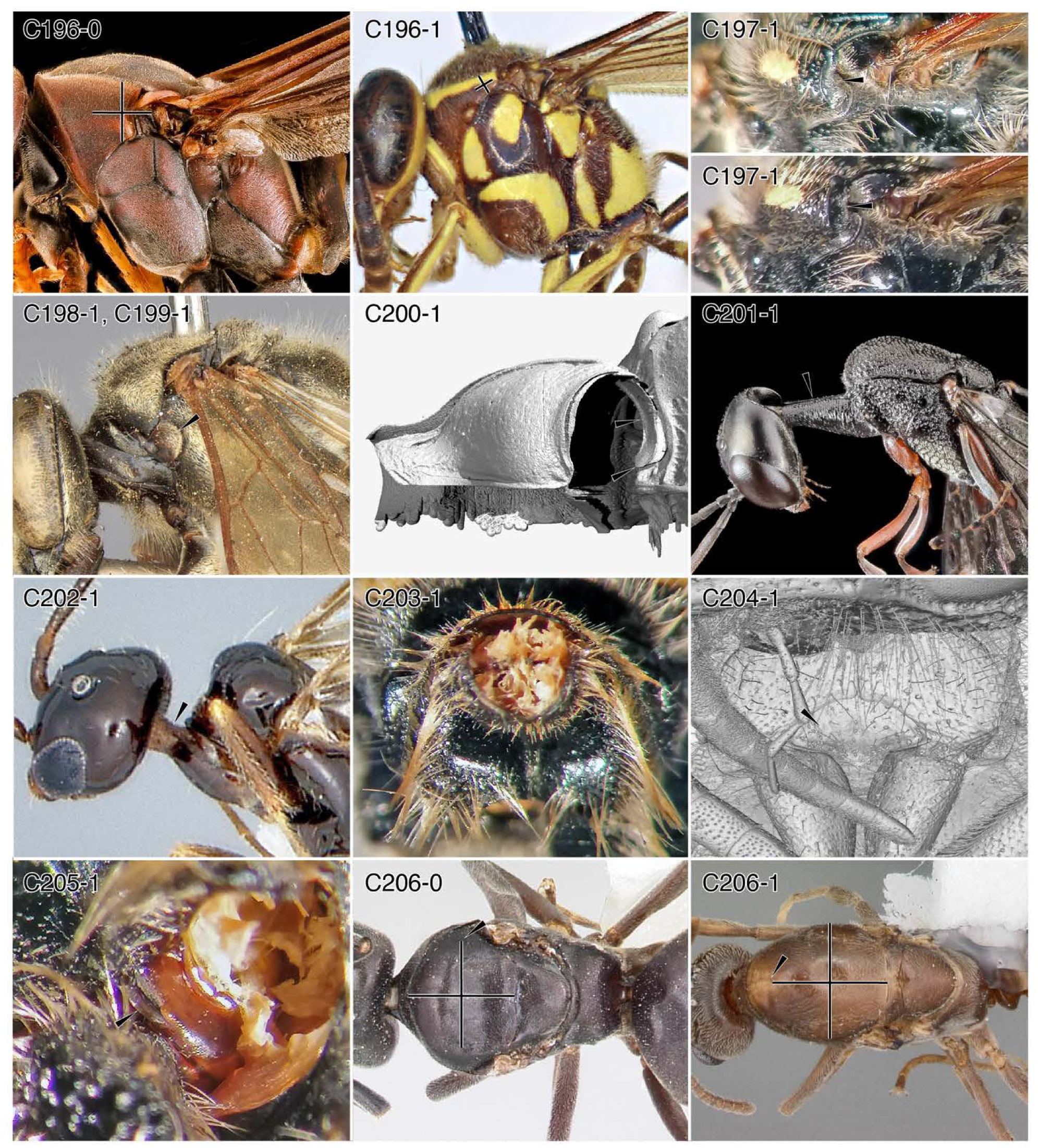
Character-states: (C196-0) Lateral pronotal portions broad; *Polistes metricus.* **(C196-1)** Lateral pronotal portions narrow; *Liostenogaster maculiceps.* **(C197-1)** Lateral pronotal portions broad and posteriorly notched; *Campsomeris* sp. **(C198-1)** Pronotal lobe grossly enlarged; *Tachytes rarus.* **(C199-1)** Pronotum separated from tegulum; *Tac. rarus.* **(C200-1)** Pronotum elongated posterad procoxae, ring-like; *Ampulex compressa.* **(C201-1)** Propleurae forming tube-like collar; *Gasteruption* sp. **(C202-1)** Propleural tube fused dorsally and ventrally; *Myrmecopterina* sp. **(C203-1)** Propleurae concave, receiving head; *Ca. pilipes.* **(C204-1)** Prosternum massive, broadly exposed. *Chrysura* sp. **(C205-1)** Prosternum external small, laminae; *Ca. pilipes.* **(C206-0)** Mesoscutum wider than long; *Tapinoma sessile.* **(C206-1)** Mesoscutum longer than wide; *Linepithema humile.* **Families: (C196)** Vespidae. **(C197, 203, 205)** Scoliidae. **(C198, 199)** Crabronidae. **(C200)** Ampulicidae. **(C201)** Gasteruptiidae. **(C202)** Plumariidae. **(C204)** Chrysididae. **(C206)** Formicidae.

**Chars. 207-216.**
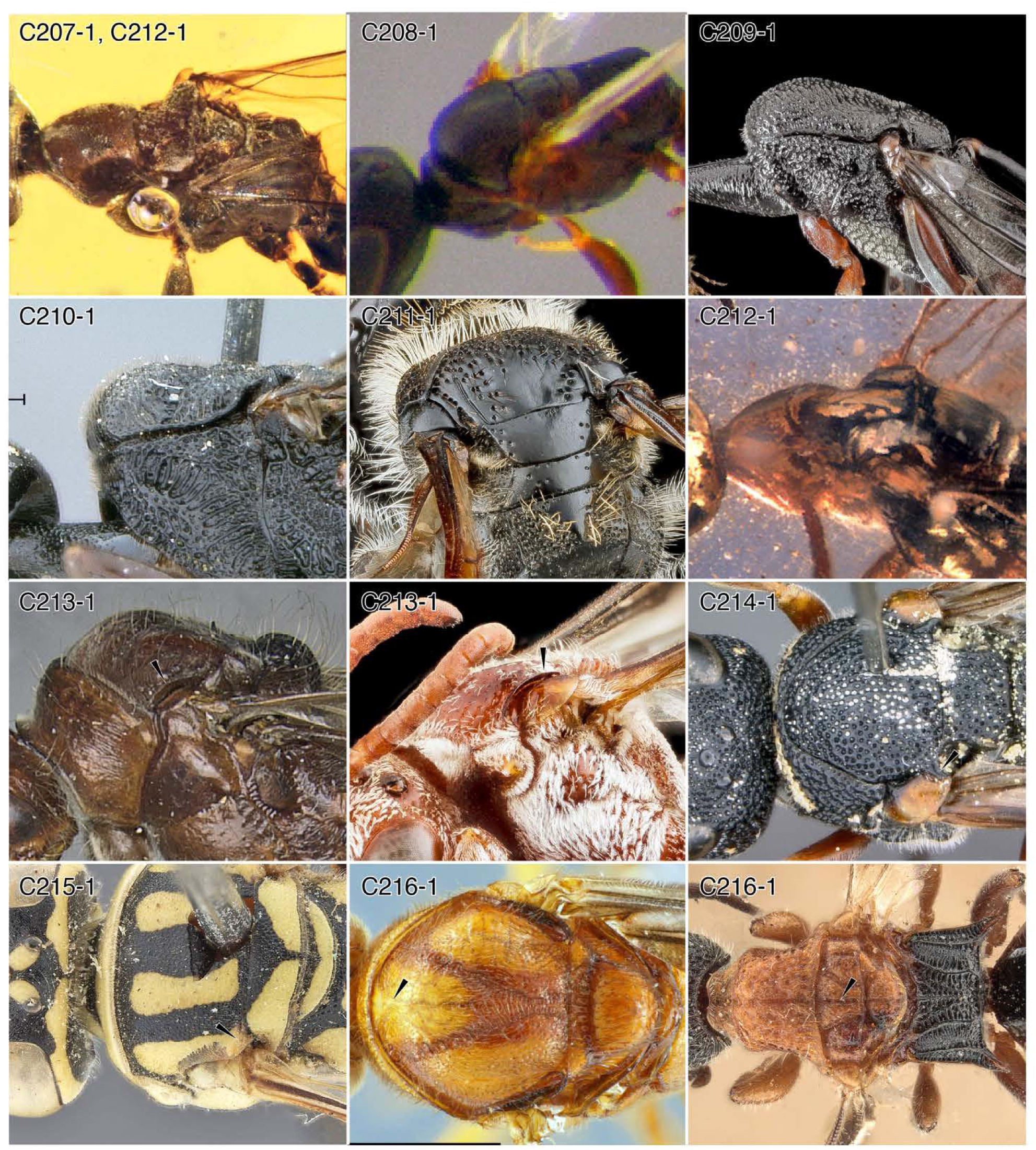
Families: Character-states: (C207-1) Mesoscutum width ≥ 2 x length; †*Falsiformica* sp. **(C208-1)** Mesoscutum anteriorly bulging, with lateral “shoulders”; *Ammoplanus hofferi.* **(C209-1)** Mesoscutum anteriorly bulging, without “shoulders”; Gasteruptiidae: *Gasteruption* sp. **(C210-1)** Median bulge of pronotum with distinct lateral faces; Aulacidae: *Pristaulacus pilotoi.* **(C211-1)** Mesoscutum with low, median raised bulge; Scoliidae: *Campsomeris* sp. **(C212-1)** Mesoscutal lateral portions enlarged; (a) †*Falsiformica* sp.; (b) Aculeata: †*Burmasphex sulcatus.* **(C213-1)** Parascutal carina present; (a) *Paraponera clavata’,* (b) Apidae: *Pasites maculatus.* **(C214-1)** Posterolateral mesoscutal digitate processes present; Vespidae: *Allepipona species* (male). **(C215-1)** Mesoscutum with “oblique scutal carina”; Bembicidae: *Bembecinus somalicus.* **(C216-1)** Mesoscutum with anteromedian line that is raised or sunken; (a) *Anochetus mixtus,* (b) Bethylidae: *Mesitius capensis.* **(C207, 212)** †Falsiformicidae. **(C208)** Ammoplanidae. **(C209)** Gasteruptiidae. **(C210)** Aulacidae. **(C211)** Scoliidae. **(C213-1b)** Apidae. **(C214)** Vespidae. **(C215)** Bembicidae. **(C216-1b)** Bethylidae. **(C213-1a, 216-1a)** Formicidae.

**Chars. 217-228.**
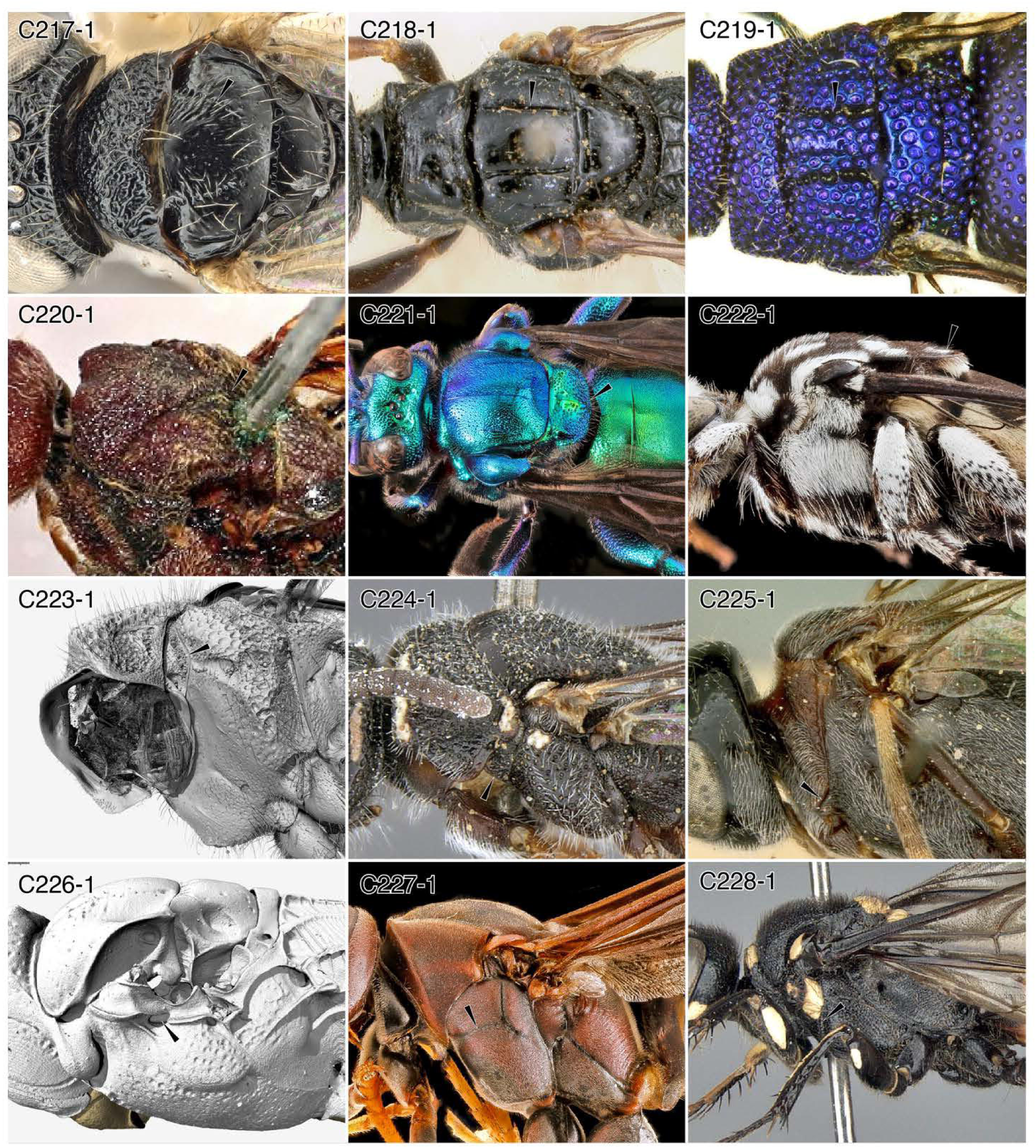
Character-states: (C217-1) Female notauli present; *Anteon elongation.* **(C218-1)** Female notauli completely traversing mesoscutum; *Dolichurus basuto.* **(C219-1)** Female notauli complete, parallel; *Odontochrydium bicristatum.* **(C220-1)** Female notauli meeting medially; *Pristaulacus irenae.* **(C221-1)** Mesoscutellum overhanging propodeum; *Exaerete smaragdina.* **(C222-1)** Mesoscutellum flat, sheet-like posteriorly; *Thyreus* sp. **(C223-1)** Prepectus present; *Chrysura* sp. **(C224-1)** Prepectus distinct, fused to mesopectus; *Sapygina enderleini.* **(C225-1)** Mesopectus with anteroventral process fitting with pronotum; *Zeuxevania* sp. **(C226-1)** Deep pit in mesopectus ventrad forewing insertion; *Ampulex compressa.* **(C227-1)** Longitudinal mesopectal sulcus; *Polistes metricus.* **(C228-1)** Mesopectus with dorsoventral sulcus in anterior half; *Philanthus loeflingi.* **Families: (C217)** Dryinidae. **(C218, 226)** Ampulicidae. **(C219, 223)** Chrysididae. **(C220)** Aulacidae. **(C221, 222)** Apidae. **(C224)** Sapygidae. **(C225)** Evaniidae. **(C277)** Vespidae. **(C288)** Philanthidae.

**Chars. 228-241.**
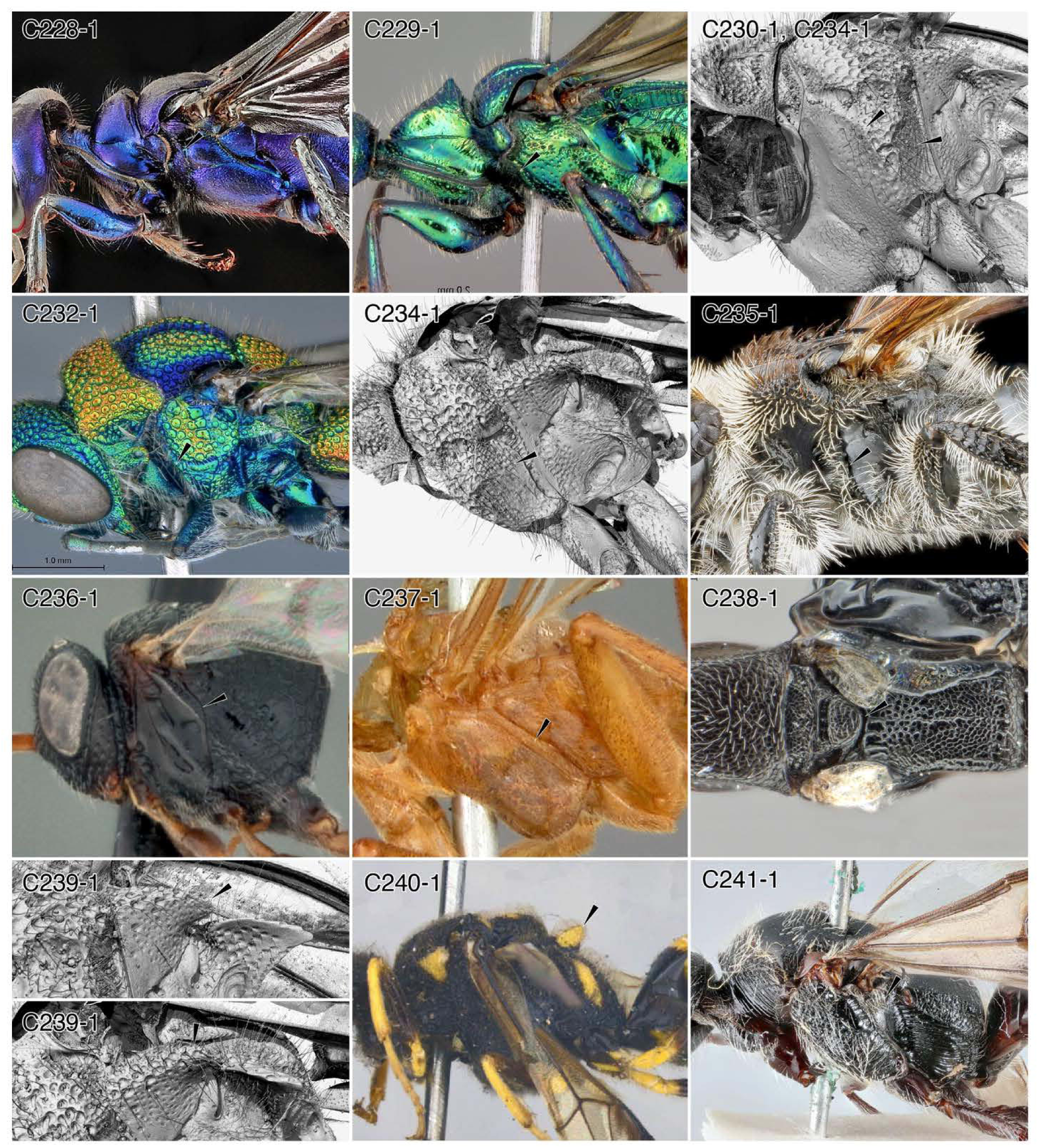
Character-states: (C228-1) Mesopectus with dorsoventral sulcus in anterior half; *Chlorion aerarium.* **(C229-1)** Mesopectus with dorsoventral carina in anterior half; *Ampulex bredoi.* **(C230-1)** Dorsoventral carina on anterior surface of mesopectal scrobe; *Chrysura* sp. **(C232-1)** Dorsoventral mesopectal carina curved, rather than angular; *Chrysis eximia.* **(C234-1)** Mesopectus posteriorly scrobiculate, receiving femur; *Chrysura* sp. **(C235-1)** Mesopectus triangular in the frontal plane; *Campsomeris* sp. **(C236-1)** Posterior mesopectal margin convex; *Hyptia floridana.* **(C237-1)** Mesepimeron developed; *Paniscomima bekilyi.* **(C238-1)** Metanotum reduced, not visible externally; *Holepyris gaigherae.* **(C239-1)** Metanotum dorsolaterally produced as shark-fin-like flange; *Chrysura* sp. **(C240-1)** Metanotum with dorsomedian process; *Bareogonalos canadensis.* **(C241-1)** Metapleural area scrobiculate, receiving femur; *Kathepyris nyassica.* **Families: (C228)** Sphecidae. **(C229)** Ampulicidae. **(C230, 232, 234, 239)** Chrysididae. **(C245)** Scoliidae. **(C236)** Evaniidae. **(C237)** Rhopalosomatidae. **(C238, 241)** Bethylidae. **(C240)** Trigonalidae.

**Chars. 241-249.**
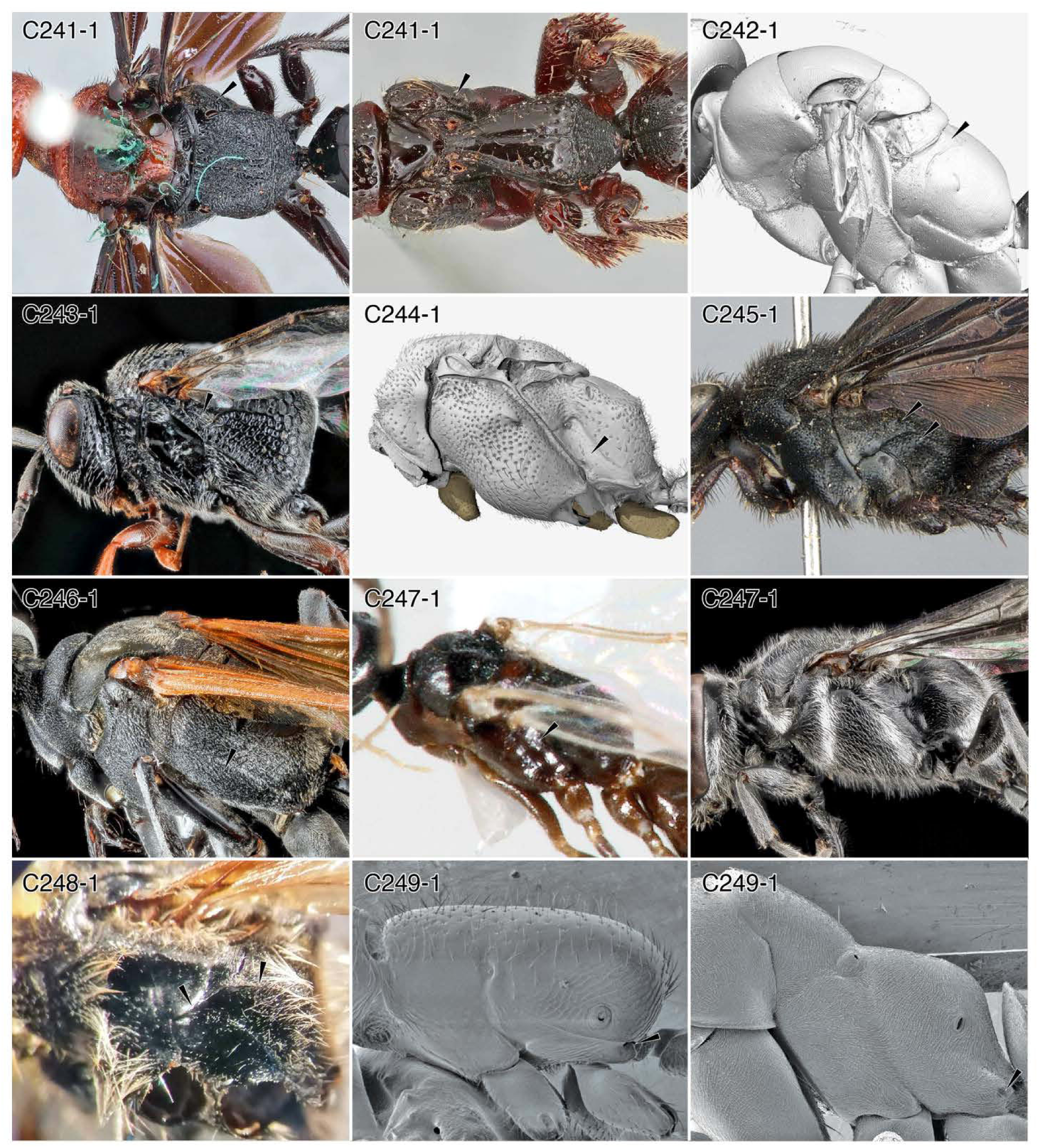
Character-states: (C241-1) Metapleural area scrobiform, receiving femur; (a) *Pristocera atopogamia* (male); (b) *P. incerta* (female). **(C242-1)** Upper metapleural area large, metapostnotum broad; *Anoplius* (*Arachnophronctus’*). **(C243-1)** Upper metapleural area small; likely *Hyptia harpyoides.* **(C244-1)** Lower metapleural area small; *Colocistis* sp. **(C245-1)** Upper and lower metapleural areas large; *Scolia erythropyga.* **(C246-1)** Lower metapleural area elongated, subtriangular; *Ammophila apicalis.* **(C247-1)** Anterior and posterior margins of the metapleural area parallel; (a) *Heterogyna* (male); (b) *Trypoxylon subimpressum.* **(C248-1)** Upper metapleural area with longitudinal carina in dorsal fourth; *Campsomeris pilipes.* **(C249-1)** Metapleural gland present; (a) *Apomyrma stygia;* (b) *Formica fusca* group. **Families: (C241)** Bethylidae. **(C242)** Pompilidae. **(C243)** Evaniidae. **(C244)** Tiphiidae. **(C245, 248)** Scoliidae. **(C246)** Sphecidae. **(C247-1a)** Heterogynaidae. **(C247-1b)** Pemphredonidae. **(C249)** Formicidae.

**Chars. 250-259.**
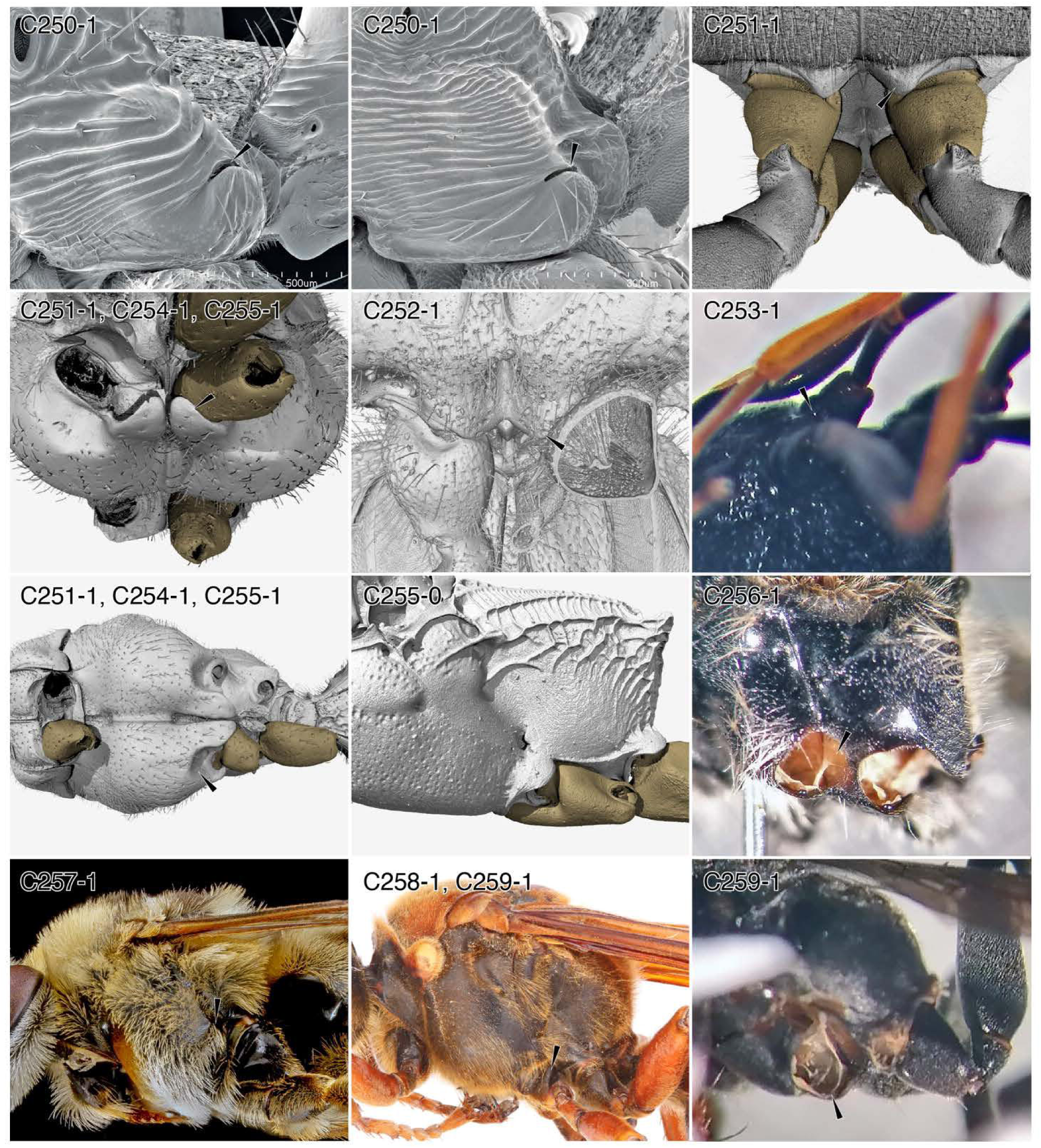
Character-states: (C250-1) Metapleural gland orifice slit-shaped, exposed, oriented dorsad; (a) *Ectatomma tuberculatum;* (b) *Manica rubida.* **(C251-1)** Mesopectus with lobes anterad mesocoxae, especially large in Thynnoidea; (a) *Pseudomasaris vespoides;* (b) *Colocistis* sp. **(C252-1)** Mesopectus with lobes anterad mesocoxae, lateromedially expanded; *Chrysura* sp. **(C253-1)** Medial mesocoxal articulation situated mediad coxa on digitate, ventral process; genus and species indet. **(C254-1)** Mesopectus with ventral scrobes anterior to and receiving coxae; *Colocistis* sp. **(C255-0)** Mesopectus without coxal scrobes; *Ampulex compressa.* **(C255-1)** Mesopectal coxal scrobes deep; *Colocistis* sp. **(C256-1)** Mesocoxae set in large, deep socket; *Campsomeris pilipes.* **(C257-1)** Mesocoxae dorsoventrally elongate; *Svastra petulca.* **(C258-1)** Mesocoxae foreshortened; *Sphecius speciosus.* **(C259-1)** Medial mesocoxal articulation produced ventrolaterally; *Tachysphex* sp. **Families: (C250)** Formicidae. **(C251-1a)** Vespidae. **(C251-1b, 254)** Tiphiidae. **(C252)** Chrysididae. **(C253)** Evaniidae. **(C255)** Ampulicidae. **(C256)** Scoliidae. **(C257)** Apidae. **(C258)** Bembicidae. **(C259)** Crabronidae.

**Chars. 260-269.**
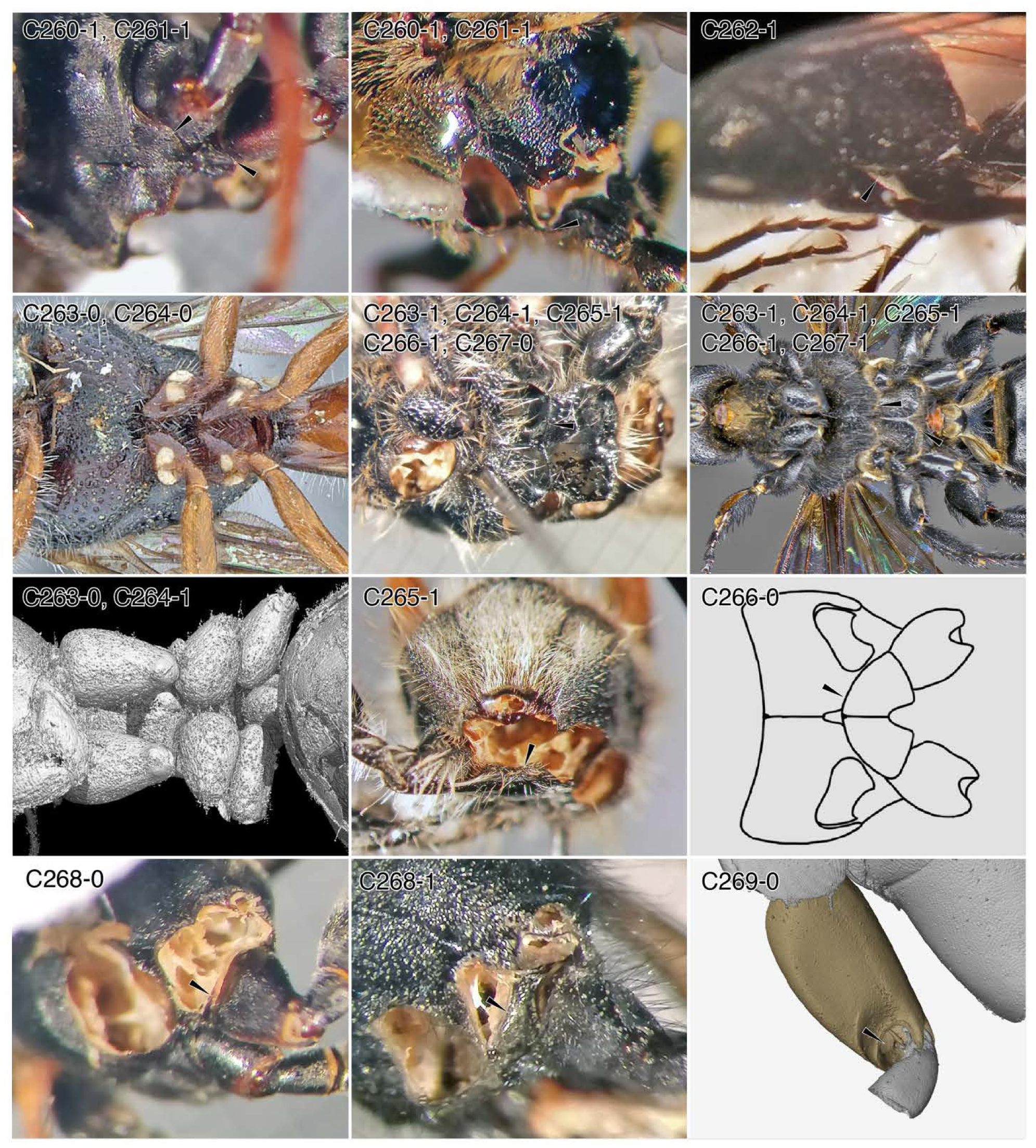
Character-states: (C260-1) Mesopectus expanded lateromedially as lobes not overlapping mesocoxae; (a) *Tachysphex* sp.; (b) *Dianthidium* sp. **(C261-1)** Medial metacoxal articulations produced ventrally; (a) *Tachysphex* sp.; (b) *Dianthidium* sp. **(C262-1)** Metapectal ventral surface flat; *Methocha* sp. **(C263-0)** Mesocoxae close-set; (a) *Sapygina braunsi;* (b) burmite genus near *Lasius.* **(C263-1)** Mesocoxae wide-set; (a) *Campsomeris pilipes;* (b) *Megascolia procer.* **(C264-0)** Metacoxae close-set; *Sa. braunsi.* **(C264-1)** Metacoxae wide-set; (a) *C. pilipes;* (b)*Meg. procer,* (c) burmite genus near *Lasius.* **(C265-1)** Broad flat process between metacoxae; *C. pi lipes.* **(C266-0)** Anterior margin of metasternal area strongly curved; *Proscolia spectator,* after Osten (1988). **(C266-1)** Anterior margin of metasternal area more-or-less linear; *C. pilipes;* (b) *Meg. procer.* **(C267-0)** Transverse sulcus absent anterad intermetacoxal process; *C. pilipes.* **(C267-1)** Transverse sulcus present anterad intermetacoxal process *Meg. procer.* **(C268-0)** Metasternal area more-or-less dorsoventrally oriented, lateromedially narrow; *Tachysphex* sp. **(C268-1)** Metasternal area more-or-less dorsoventrally oriented, lateromedially broad; *Sceliphron* sp. **(C269-0)** Prodisticoxal foramina open; *Anoplius* (*Arachnophronctus*) sp. **Families: (C260-1a, 261-1 a, 268-0)** Crabronidae. **(C260-1b, C261-1b)** Megachilidae. **(C262)** Tiphiidae. **(C263, 264)** Sapygidae. **(C263-0b, 264-1c)** Formicidae. **(C263-1, 264-267)** Scoliidae. **(C268-1)** Sphecidae. **(C269)** Pompilidae.

**Chars. 269-279.**
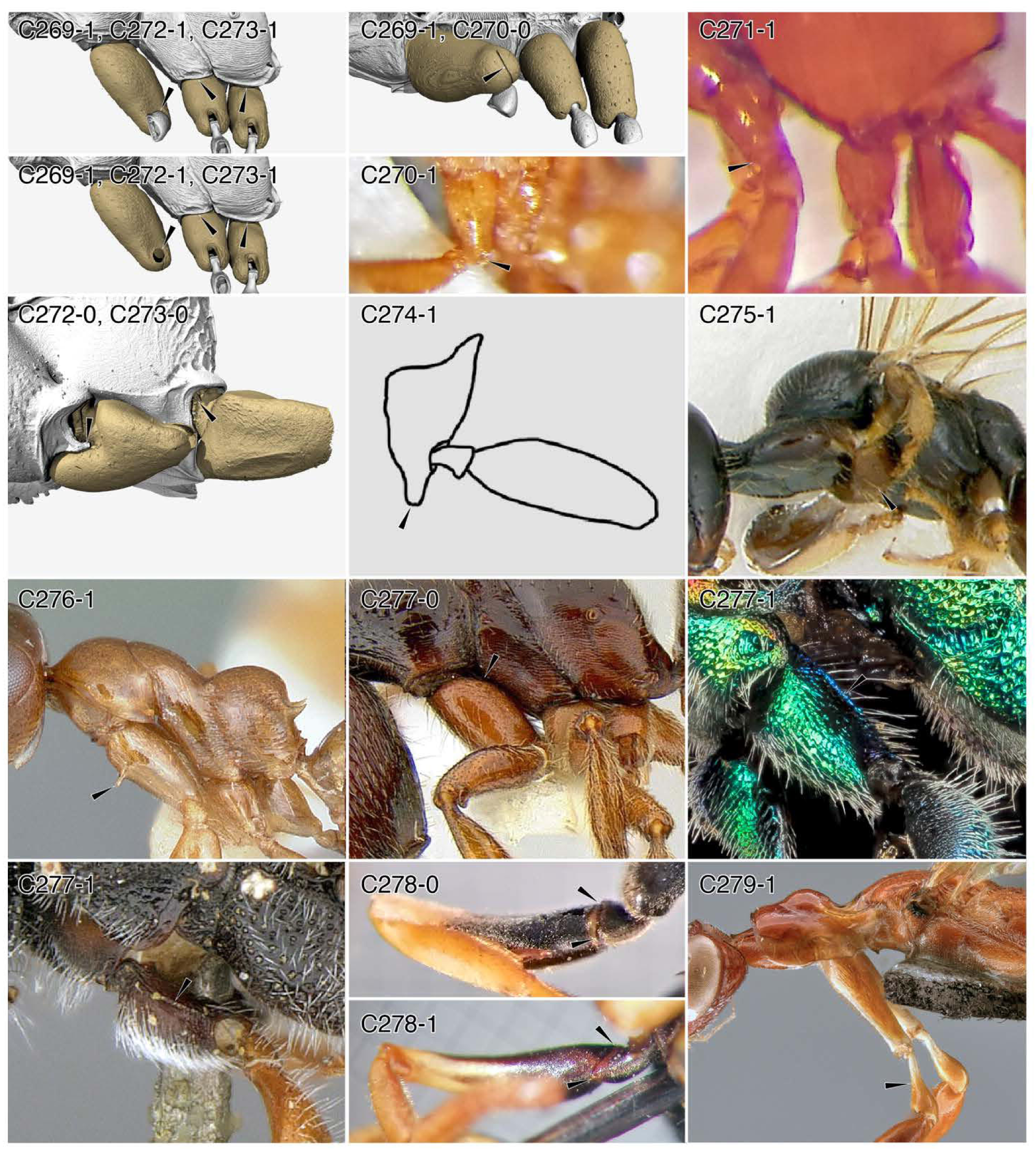
Character-states: (C269-1) Prodisticoxal foramina closed laterally and ventrally; *Amblyopone australis.* **(C270-0)** Mesocoxal foramina closed; *Amb. australis.* **(C270-1)** Prodisticoxal foramina incompletely closed ventrally; *Myrmosa* sp. **(C271-1)** Prodisticoxa curved; *Loboscelidia* sp. **(C272-0)** Mesocoxal foramina opened; *Ampulex compressa.* **(C272-1)** Mesocoxal foramina closed; *Amb. australis.* **(C273-0)** Metacoxal foramina open; *Amp*. *compressa.* **(C273-1)** Metacoxal foramina closed; *Amb. australis.* **(C274-1)** Procoxa produced ventrally; *Plumarius* sp. **(C275-1)** Prodisticoxal foramina migrated anteriorly; *Ycaploca evansi.* **(C276-1)** Procoxae with anterior processes; *Santschiella kohli.* **(C277-0)** Procoxa without posterior dorsoventral carina; *Amb. australis.* **(C277-1)** Procoxa with posterior dorsoventral carina; (a) *Chrysura* sp. (b) *Sapygina enderleini.* **(C278-0)** Protrochanter ventroapically truncate; *Polistes* sp. **(C278-1)** Protrochanter ventroapically produced, triangular; *Pseudomasaris vespoides.* **(C279-1)** Protrochanter length > 3 x width; *Dryinus bellicosus.* **Families: (C269, 270-0, 272-1, 273-1, 276-1, 277-0)** Formicidae. **(C270-1)** Mutillidae. **(C271-1, 277-1a)** Chrysididae. **(C272-0, 273-0)** Ampulicidae. **(C274)** Plumariidae. **(C275)** Scolebythidae. **(C277-1b)** Sapygidae. **(C278)** Vespidae. **(C279)** Dryinidae.

**Chars. 280-289.**
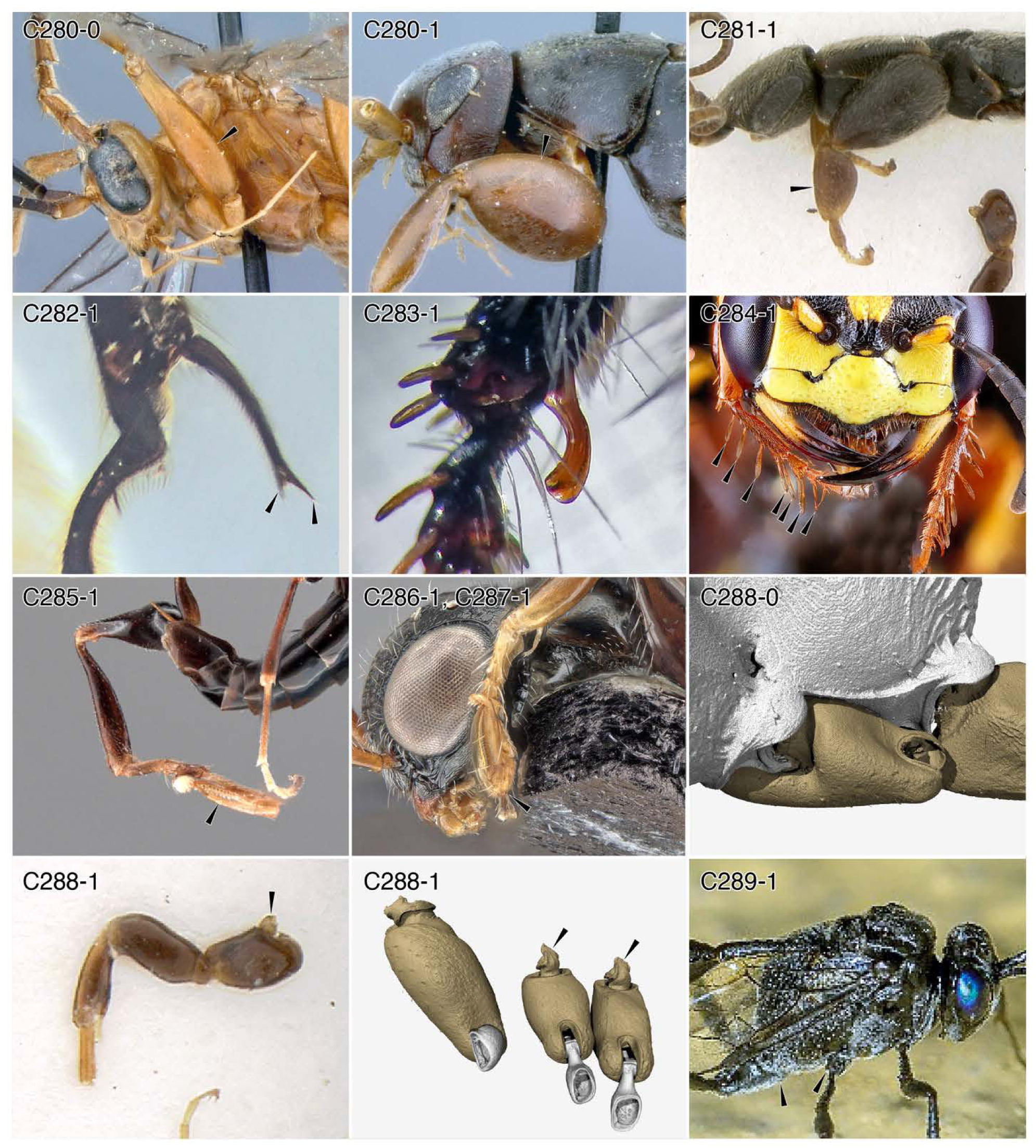
Character-states: (C280-0) Female profemur thin; *Paniscomima bekilyi.* **(C280-1)** Female profemur grossly thickened; *Olixon saltator.* **(C281-1)** Female protibia grossly thickened; *Sclerogibba crassifemorata.* **(C282-1)** Protibial calcar bifid; †*Gerontoformica* sp. **(C283-1)** Protibial calcar laminate, strongly curved; *Campsomeris pilipes.* **(C284-1)** Protarsi with psammochaetae; *Philanthus gibbosus* (female). **(C285-1)** Protarsus modified for clasping; *Gonatopus costalis.* **(C286-1)** One propretarsal claw reduced; *Anteon beankanum.* **(C287-1)** One propretarsal claw completely absent; *An. beankanum.* **(C288-0)** Basimeso- and basimetacoxae broad; *Ampulex compressa.* **(C288-1)** Basim eso- and basimetacoxae narrowed, ball-like; (a) *S. crassifemorata;* (b) *Amblyopone australis.* **(C289-1)** Meso- and metacoxae wide-set; *Evania appendigaster.* **Families: (C280)** Rhopalosomatidae. **(C281, 288-1a)** Sclerogibbidae. **(C282, 288-1b)** Formicidae. **(C283)** Scoliidae. **(C284)** Philanthidae. **(C285-287)** Dryinidae. **(C288)** Ampulicidae. **(C289)** Evaniidae.

**Chars. 290-299.**
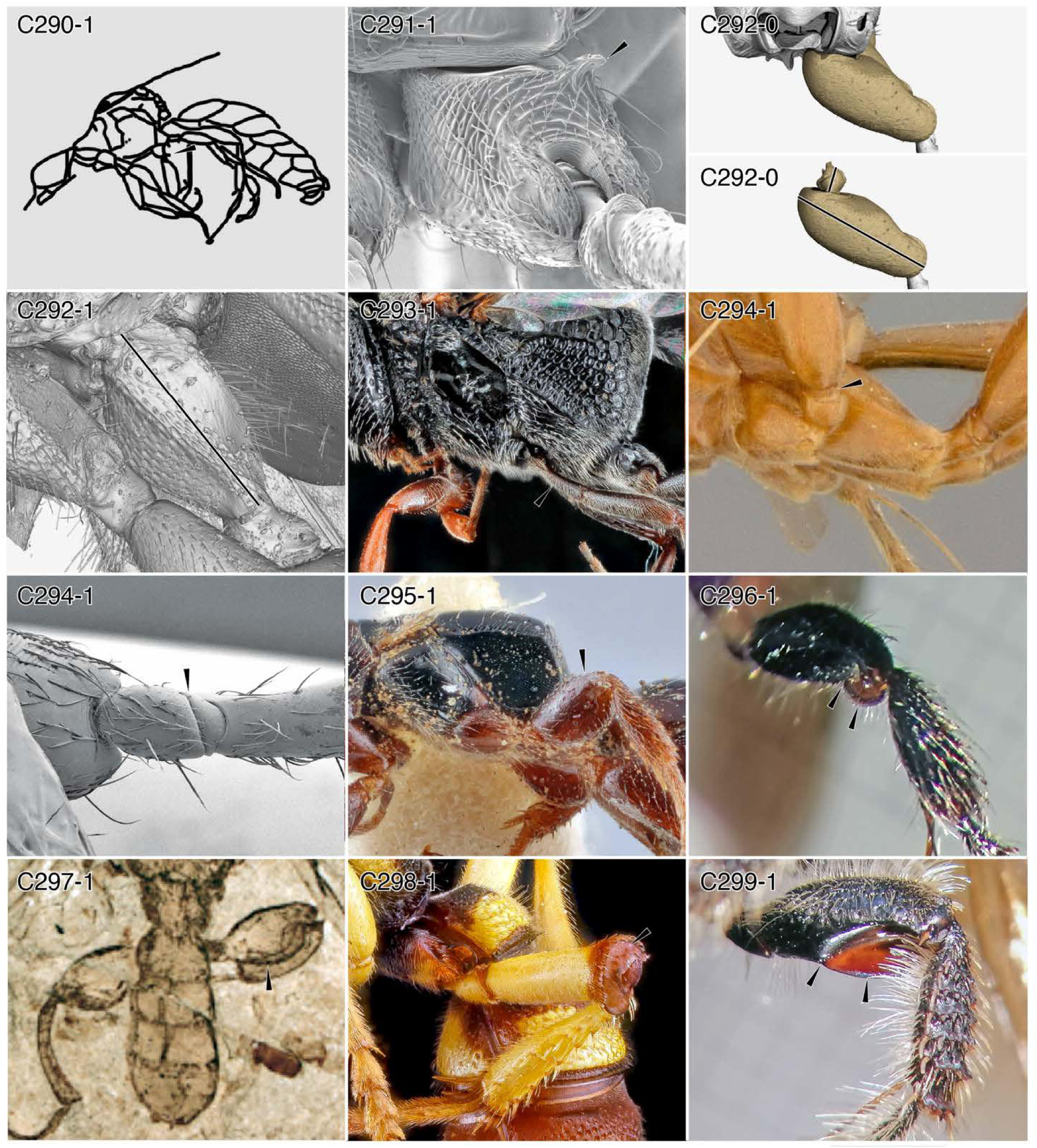
Character-states: (C290-1) Mesocoxa with ventral spine; †*Cretoecus spinicoxa,* after Budrys (1993). **(C291-1)** Metacoxa with posterodorsal process; *Gnamptogenys striatula.* **(C292-0)** Metacoxa with two distinct longitudinal axes; *Amblyopone australis.* **(C292-1)** Metacoxa with single longitudinal axis; *Chrysura* sp. **(C293-1)** Mesotrochanter length > 3 x width; likely *Hyptia harpyoides.* **(C284-1)** Mesotrochantellus well-developed; (a) *Paniscomima bekilyi;* (b) *G. annulata.* **(C295-1)** Female meso- and/or metafemora grossly thickened; *Epyris viduatus.* **(C296-1)** Female meso- and metafemora compressed, with apical discs; genus and species indet. **(C297-1)** Female metafemora with serrate/dentate posteroventral margin; †*Karataus daohugouensis.* **(C298-1)** Metafemora apically truncate; *Cerceris triangulata* (female). **(C299-1)** Metafemora with posteroapical lamella that is not disc-like; *Campsomeris pilipes.* **Families: (C290)** Pemphredonidae. **(C291, 292-0, 294-1b)** Formicidae. **(C292-1)** Chrysididae. **(C293)** Evaniidae. **(C284-1a)** Rhopalosomatidae. **(C295)** Bethylidae. **(C296)** Tiphiidae. (297) †Ephialtitoidea. **(C298)** Philanthidae. **(C299)** Scoliidae.

**Chars. 300-307.**
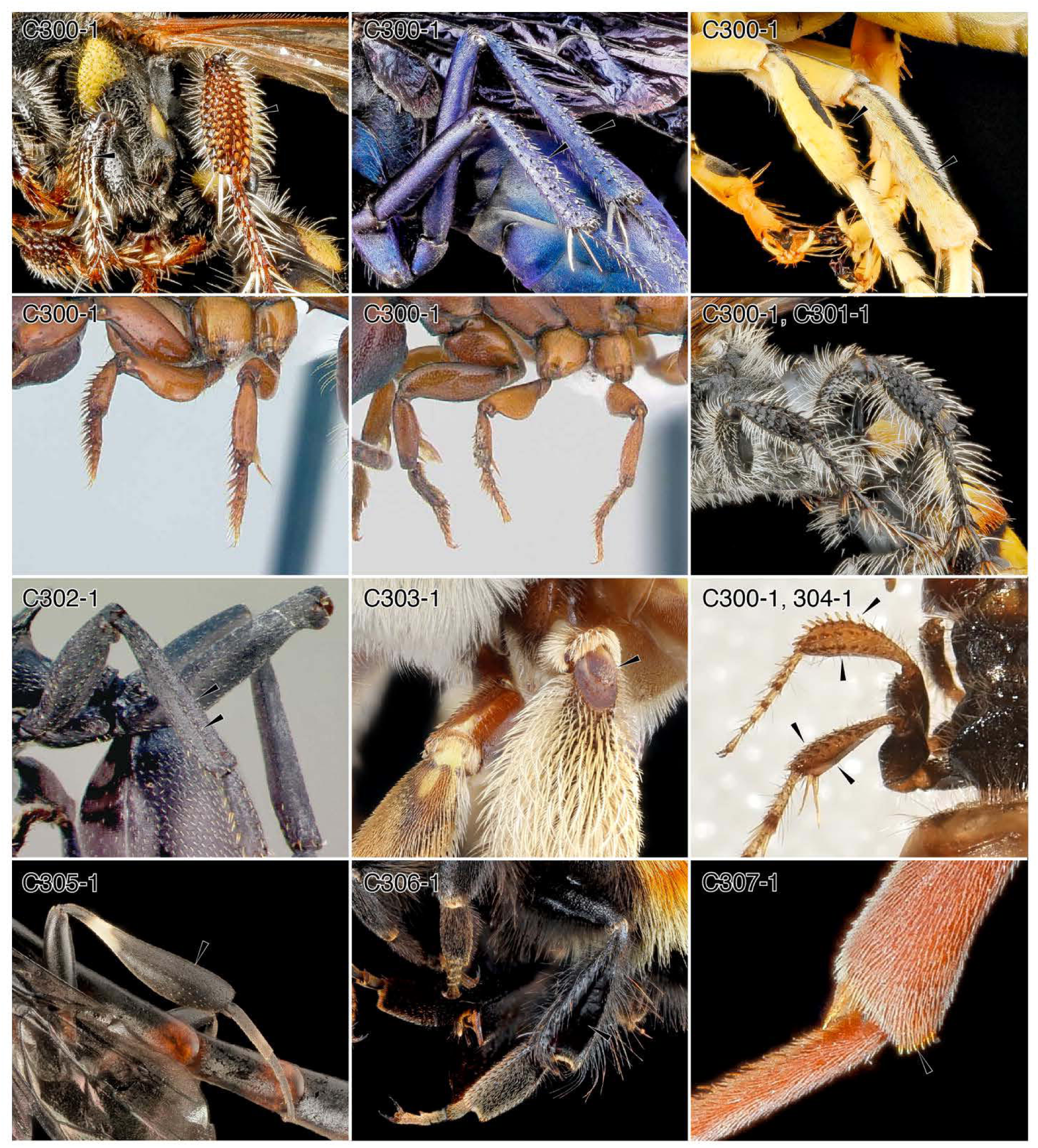
Character-states: (C300-1) Meso- and metatibiae with traction setae/spurs; (a) *Myzinum maculatum;* (b) Pompilidae: *Pepsis ruficomis;* (c) genus and species indet.; (d) *Centromyrmex decessor,* (e) *Promyopias silvestrii;* (f) *Campsomeris* sp.; (g) genus and species indet. **(C301-1)** Metatibial traction spurs in at least two transverse rows; *Campsomeris* sp. **(C302-1)** Meso- and metatibiae with longitudinal ridge; *Cephalotes atratus.* **(C303-1)** Metatibia proximodorsally truncate; *Centris aethyctera.* **(C304-1)** Meso- and metatibiae clavate; thynnid genus and species indet. **(C305-1)** Only metatibia clavate; *Gasteruption* sp. **(C306-1)** Metatibia corbiculate; *Bombus huntii.* **(C307-1)** Meso- and metatibial anteroapical margins with row of short, stout setae/chaetae; *Leptochilus acolhuus.* **Families: (C300-1a, g, 304)** Thynnidae. **(C300-1b)** Bembicidae. **(C300-1d, e, 302)** Formicidae. **(C300-1f, 301)** Scoliidae. **(C303, 306)** Apidae. **(C305)** Gasteruptiidae. **(C307)** Vespidae.

**Chars. 307-318.**
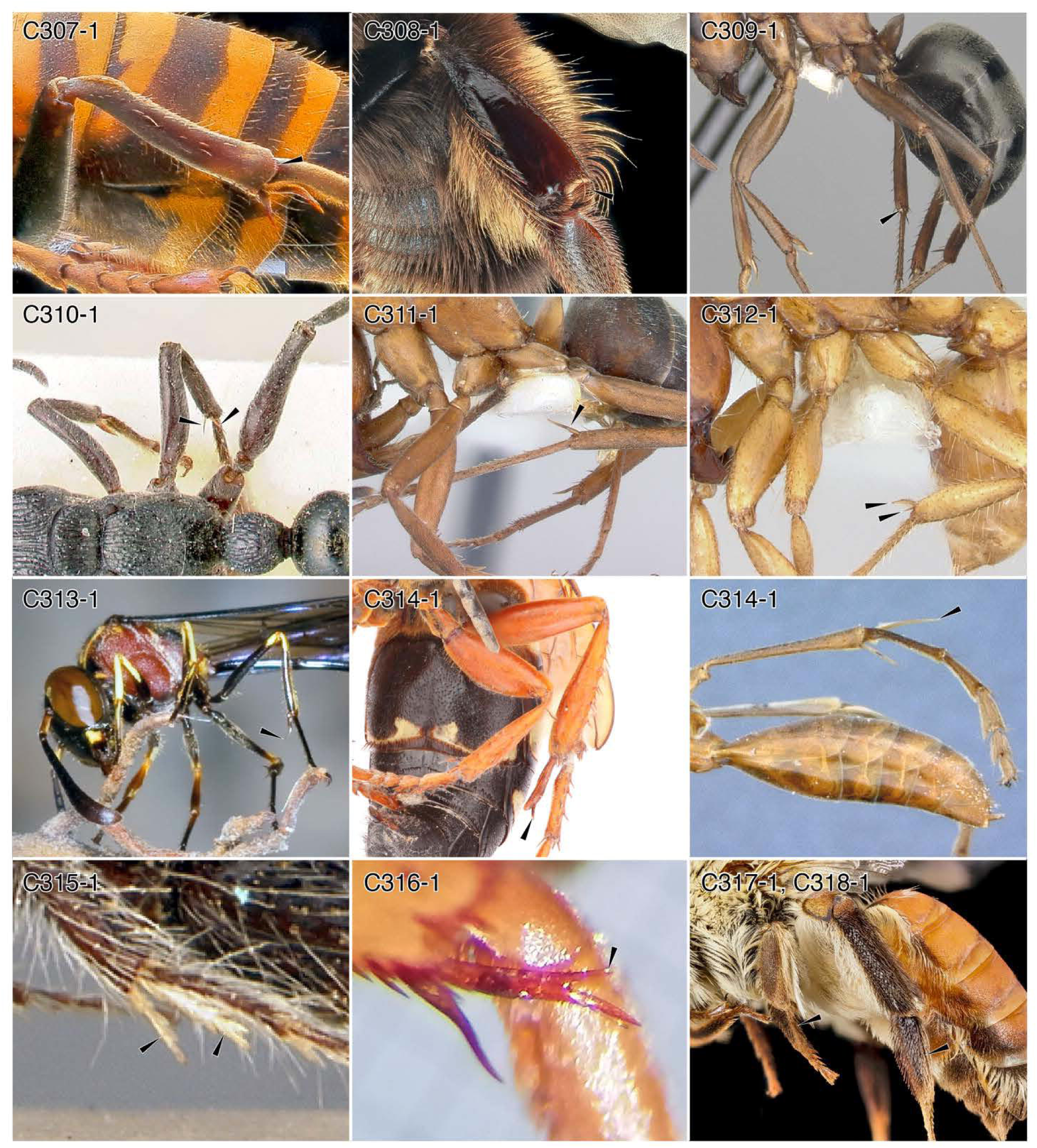
Character-states: (C307-1) Meso- and metatibial anteroapical margins with row of short, stout setae/chaetae; *Vespa manderinia.* **(C308-1)** Metatibial posteroapical margin with rastellum; *Bombus terricola.* **(C309-1)** Mesotibia with at least one ventroapical spur; *Formica browni.* **(C310-1)** Mesotibia with two ventroapical spurs; *Myrmecia queenslandica.* **(C311-1)** Metatibia with at least one ventroapical spur; *F. adamsi alpina.* **(C312-1)** Metatibia with two ventroapical spurs; *Labidus mars.* **(C313-1)** Tibial spurs elongate, wispy; *Parischnogaster mellyi.* **(C314-1)** Meso- and/or metatibial spur or spurs with length ≥ 1/2 that of their respective basitarsi; (a) *Sphecius speciosus;* (b) *Paniscomima bekilyi.* **(C315-1)** Meso- and metatibial spurs flattened with large serrations; *Apterogyna globularia.* **(C316-1)** Posterior metatibial spur with one or more preapical processes; *Pseudomasaris vespoides.* **(C317-1)** Mesobasitarsus flattened; *Notoxaea ferruginea.* **(C318-1)** Metabasitarsus flattened; *N ferruginea.* **Families: (C307, 313, 316)** Vespidae. **(C308)** Apidae. **(C309-312)** Formicidae. **(C314-1a)** Bembicidae. **(C314-b)** Rhopalosomatidae. **(C315)** Bradynobaenidae. **(C317, 318)** Andrenidae.

**Chars. C319-326.**
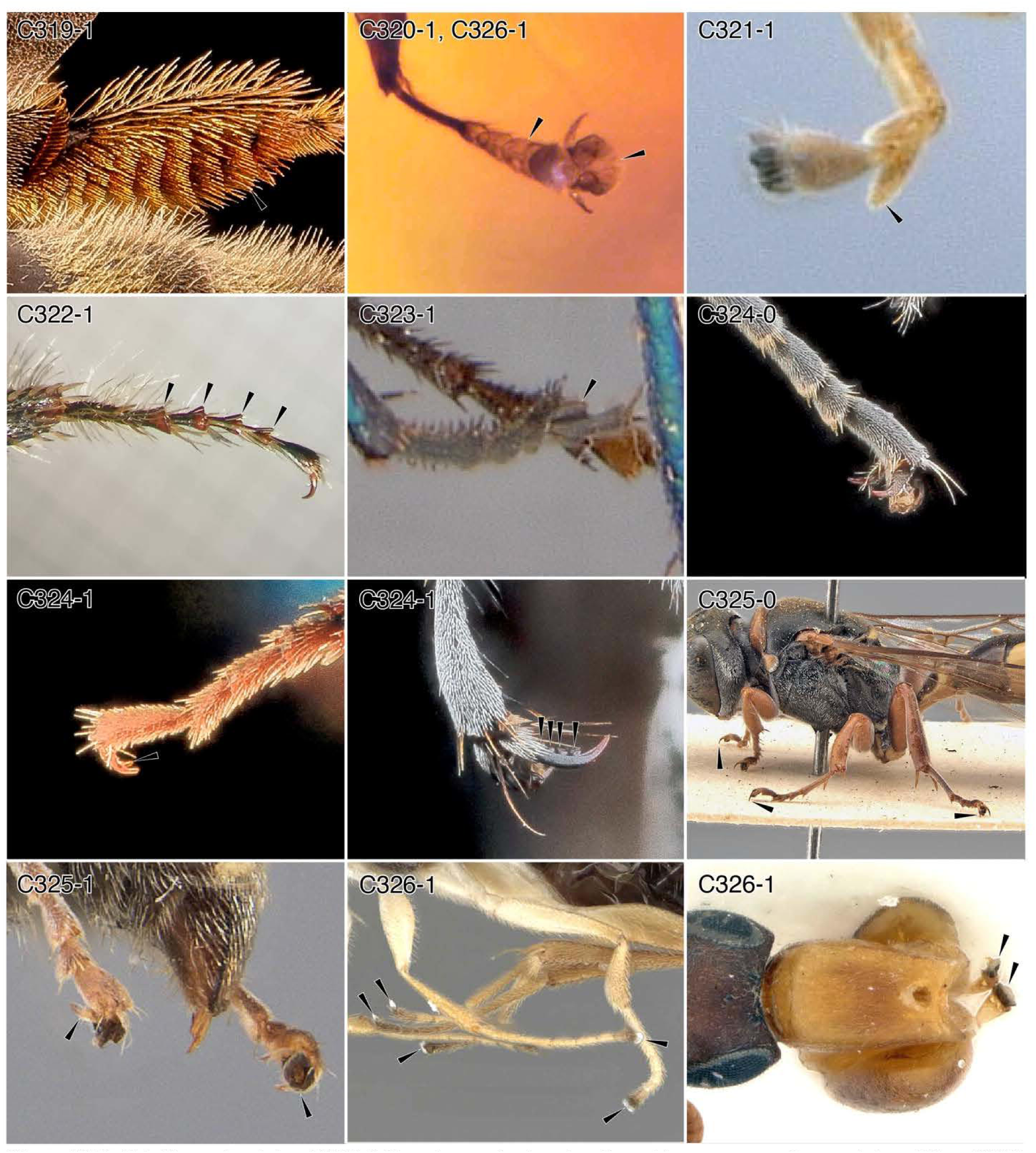
Character-states: (C319-1) Posterior metabasitarsal surface with transverse setal rows; *Apis mellifera.* **(C320-1)** Female tarsomeres II-V of any leg lateromedially expanded; †*Falsiformica* sp. **(C321-1)** Tarsomere IV asymmetrically lobate; *Olixon saltator.* **(C322-1)** Tarsomeres I-IV with apices expanded, conical; *Campsomeris pilipes.* **(C323-1)** Tarsomere IV wedge-shaped, receiving V near its base; *Ampulex bredoi.* **(C324-0)** Pretarsal claws edentate; *Trypoxylon subimpressum.* **(C324-1)** Pretarsal claws uni- to multidentate; (a) *Hedychrum parvum;* (b) *Prionyx thomae.* **(C325-0)** Pretarsal claws not reflexed ventrad; *Crossocerus glabricornis.* **(C325-1)** Pretarsal claws reflexed ventrad; *Oxybelus acutissimus.* **(C326-1)** Protarsal aroliae expanded with well-developed sclerites; (a) †*Falsiformica* sp.; (b) *Aphelopus witter,* (c) *Sclerogibba madegassa.* **Families: (C319)** Apidae. **(C320, 326-1a)** †Falsiformicidae. **(C321)** Rhopalosomatidae. **(C322)** Scoliidae. **(C323)** Ampulicidae. **(C324-0)** Pemphredonidae. **(C324-1a)** Chιysididae. **(C324-1b)** Sphecidae. **(C325)** Crabronidae. **(C326-1b)** Dryinidae. **(C326-1c)** Sclerogibbidae.

**Chars. C327-338.**
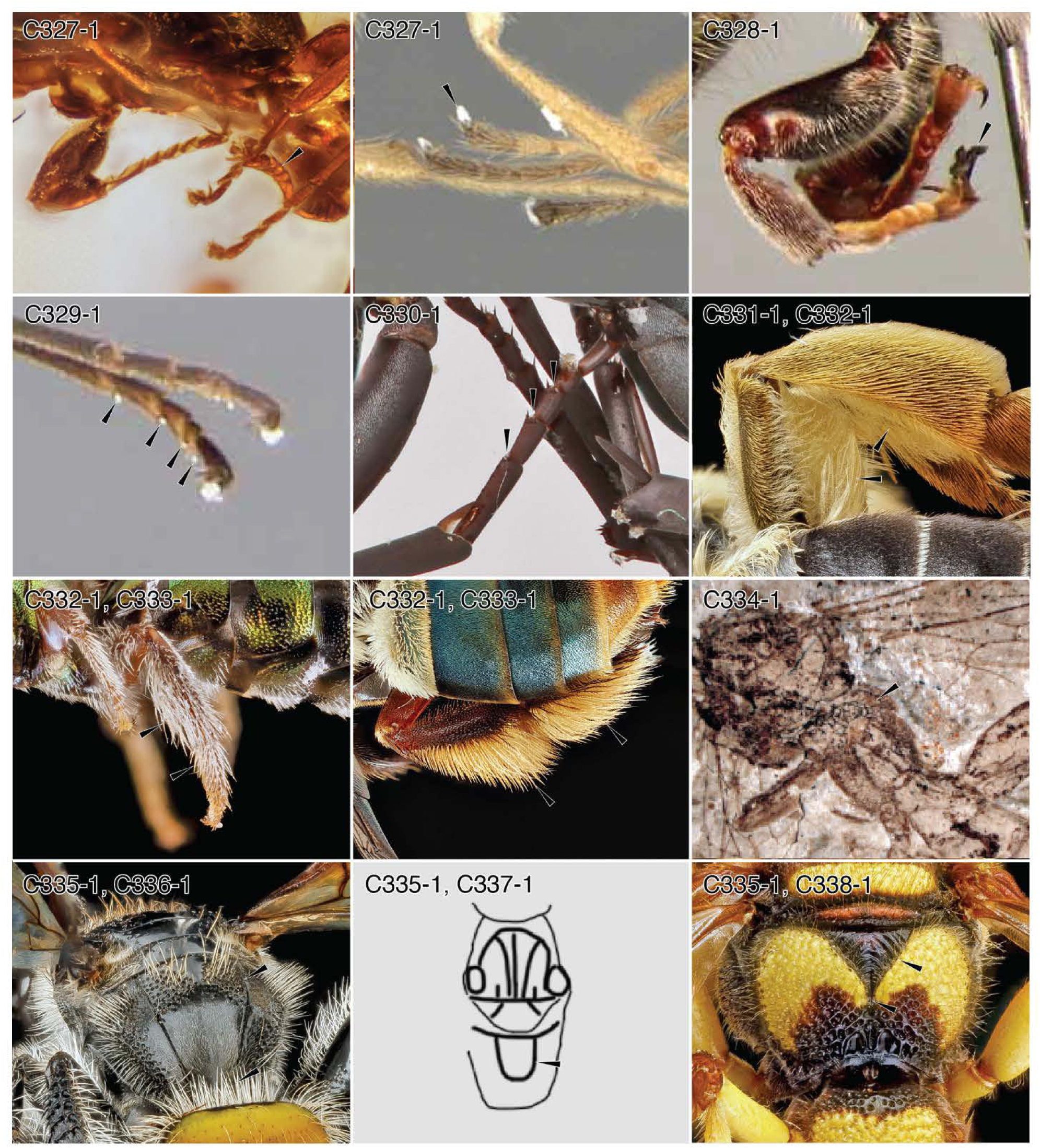
Character-states: (C327-1) Meso- and/or metaaroliae expanded; (a) † *Siccibythus ohmkuhnlei;* (b) *Aphelopus wittei.* **(C328-1)** Proarolia expanded, with large unguitractor plate; *Dasyproctus braunsii.* **(C329-1)** Tarsomeres I-IV with plantar lobes; *Trigonalys* sp. **(C330-1)** Plantar lobes relatively large; *Allochares azureus.* **(C331-1)** Metafemur with scopa; *Caupolicana electa.* **(C332-1)** Metatibia with scopa; (a) *Cau. electa,* (b) *Ceratina buscki;* (c) *Centris fasciata.* **(C333-1)** Metabasitarsus with scopa; (a) *Cera. buscki;* (b) *Cen. fasciata.* **(C334-1)** Abdominal segment I not forming propodeum; †*Karataus exilis.* **(C335-1)** Propodeum with paired paramedial lines, regardless of form; (a) *Campsomeris* sp.; (b) †*Protoscolia sinensis;* (c) *Cerceris triangulata.* **(C336-1)** Paramedial propodeal lines subparallel for their length; *Campsomeris* sp. **(C337-1)** Paramedial propodeal lines joined posteroapically through an even curve; †*P. sinensis,* after Zhang *et al.* (2002). **(C338-1)** Paramedial propodeal lines joined posteroapically, forming triangle; *Cerc. triangulata.* **Families: (C327-1a)** †Falsiformicidae. **(C327-1b)** Dryinidae. **(C328)** Crabronidae. **(C329)** Trigonalidae. **(C330)** Pompilidae. **(C331, C332-1a)** Colletidae. **(C332-1b, c, 333)** Apidae. **(C334)** †Ephialtitoidea. **(C335-1a, 337)** Scolioidea. **(C335-1b, 336)** Scoliidae. **(C335-1a, 338)** Philanthidae.

**Chars. C339-343.**
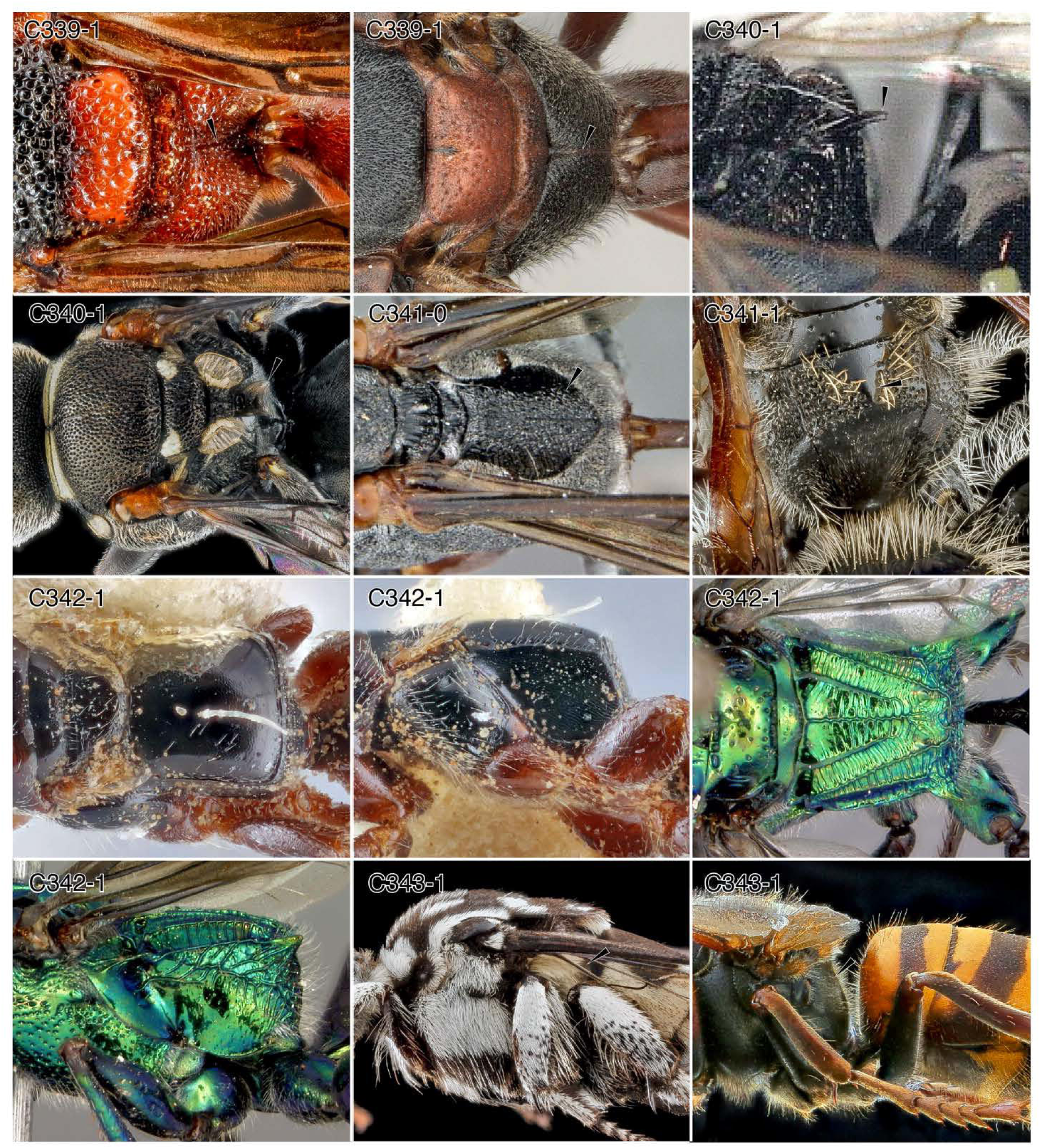
Character-states: (C339-1a) Propodeum with median longitudinal sulcus that reaches propodeal foramen; (a) *Leptochilus acolhuus;* (b) *Belonogaster juncea colonialis.* **(C340-1a)** Propodeum with ornate dorsomedian process; (a) *Oxybelus bipunctatus;* (b) *O. analis.* **(C341-0)** Propodeum without raised dorsomedian process, even if with median triangle; *Ammophila bechuana.* **(C341-1)** Propodeum with raised dorsomedian triangular/conical process; (a) *Campsomeris* sp. **(C342-1a)** Propodeum in form of long box; (a) *Epyris viduatus;* (b) *E. viduatus;* (c) *Ampulex bredor,* (d) *Amp. bredoi.* **(C343-1)** Propodeum without a dorsal surface in lateral view; (a) *Thyreus* sp.; (b) *Vespa manderinia.* **Families: (C339, 343-1b)** Vespidae. **(C340)** Crabronidae. **(C341-0)** Sphecidae. **(C341-1)** Scoliidae. **(C342-1a, b)** Bethylidae. **(C343-1a)** Apidae.

**Chars. C344-349.**
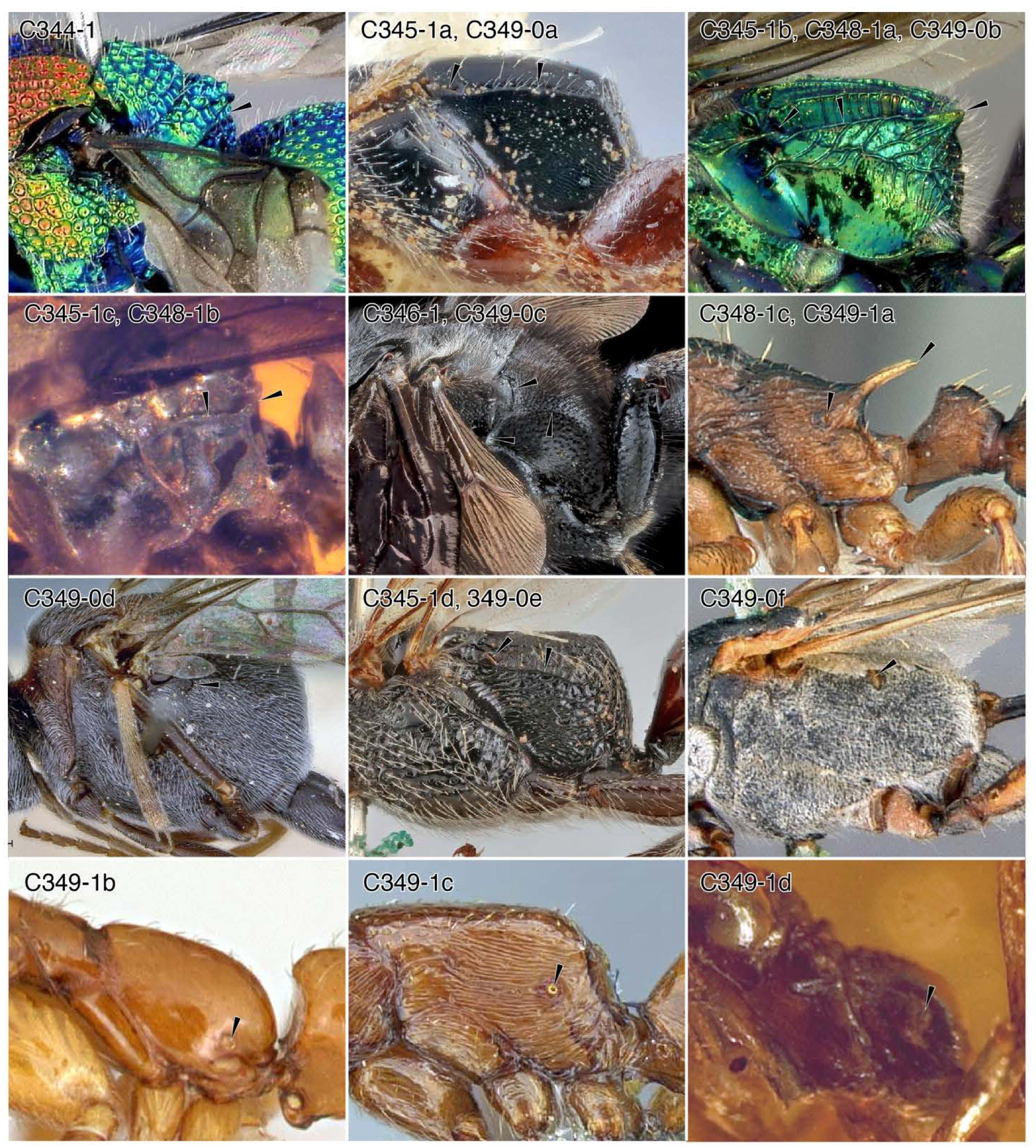
Character-states: (C344-1) Propodeum without dorsal surface and wedge-shaped in profile view; *Chrysis splendens.* **(C345-1)** Propodeal dorsolateral margins carinate; (a) *Epyris viduatus;* (b) *Ampulex bredoi;* †*Falsiformica* sp.; (c) *Pristocera armaticeps.* **(C346-1)** Propodeum with dorsolateral carina that has its anterior surface at the juncture of the upper and lower metapleural areas; *Scolia bicincta.* **(C348-1)** Dorsal posterolateral comers armed with processes of variable shape; (a) *A. bredoi;* (b) †*Falsiformica* sp.; (c) *Myrmica alaskensis.* **(C349-0)** Propodeal spiracle situated high and anteriorly, near the metanotum and/or at about posterodorsal comer of metapleural area; (a) *E. viduatus;* (b) *A. bredoi;* (c) *S. bicincta,* (d) *Zeuxevania* sp.; (e) *Pri. armaticeps;* (f) *Ammophila peringueyi.* **(C349-1)** Propodeal spiracle situated low and posteriorly, distant from metanotum and upper metapleural area; (a) *M. alaskensis;* (b) *Protanilla wardi;* (c) *Boloponera ikemkha,* (d) †*Haidomyrmex zigrasi.* **Families: (C344)** Chrysididae. **(C345-1a, d, 349-0a, e)** Bethylidae. **(C345-1b, 348-1a, 349-0b)** Ampulicidae. **(C345-1c, 348-1b)** †Falsiformicidae. **(C346, 349-0c)** Scoliidae. **(C348-1c, 349)** Formicidae. **(C349-0d)** Evaniidae. **(C349-0f)** Sphecidae.

**Chars. C350-357.**
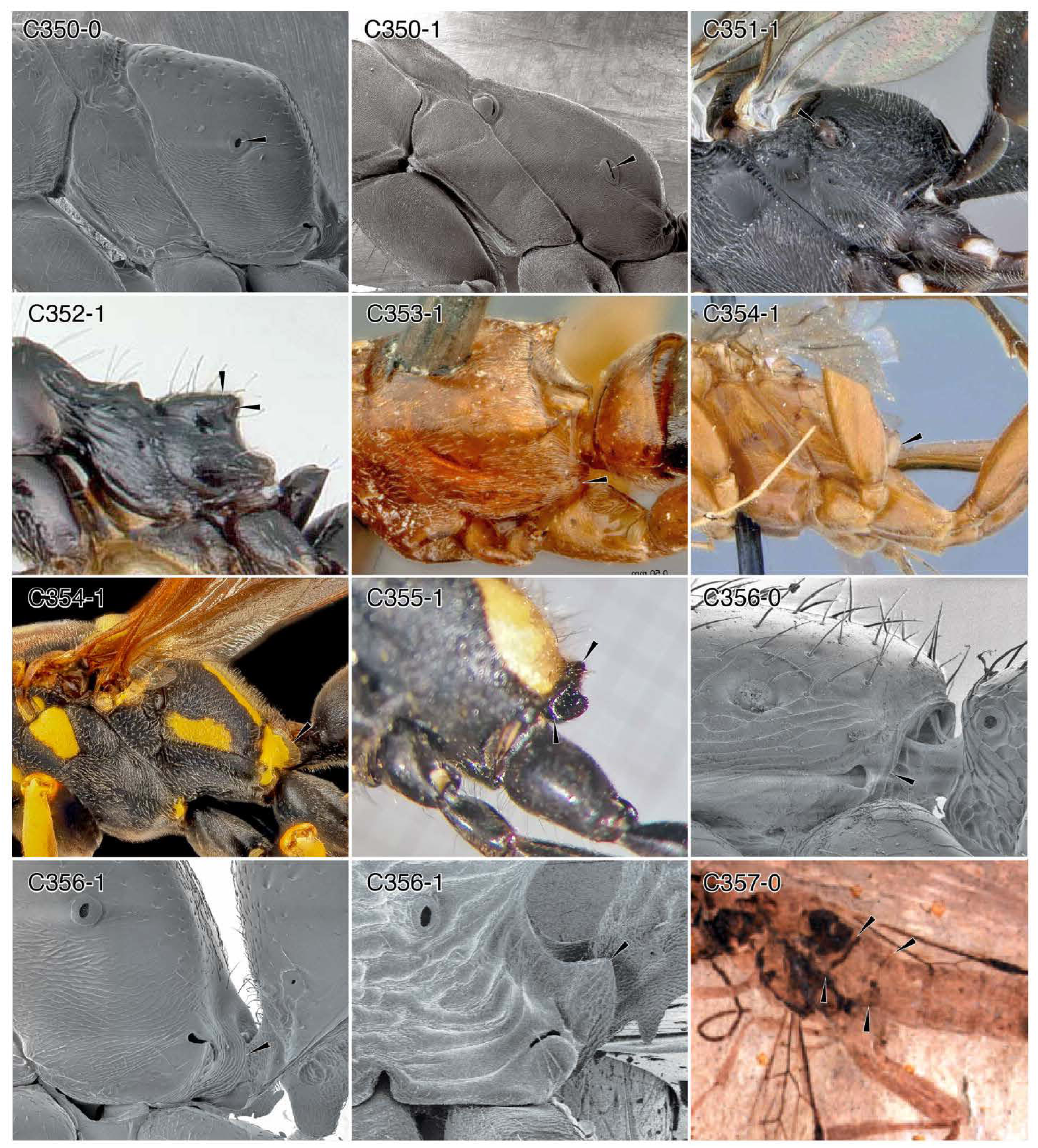
Character-states: (C350-0) Propodeal spiracle circular to elliptical; *Amblyopone australis.* **(C350-1)** Propodeal spiracle slit-shaped; *Neoponera apicalis.* **(C351-1)** Propodeal spiracle with anterior hood; *Trigonalys* sp. **(C352-1)** Propodeal spiracles on posterolateral propodeal processes; *Lepisiota canescens.* **(C353-1)** Ventrolateral propodeal margin expanded; *Olixon myrmosaeforme.* **(C354-1)** Ventrolateral propodeal margin laminate; (a) *Paniscomima bekilyi;* (b) *Polistes* sp. **(C355-1)** Propodeal foramen on short, distinct tube; *Sceliphron* sp. **(C356-0)** Propodeal foraminal carina without lobes; *Leptanilla swani.* **(C356-1)** Propodeal foraminal carina laterally lobate; (a) *A. australis;* (b) *Myrmica americana.* **(C357-0)** Anterior base of abdominal segment II as broad as I; †*Acephialtitia colossa.* **Families: (C350, 352, 356)** Formicidae. **(C351)** Trigonalidae. **(C353, 354-1a)** Rhopalosomatidae. **(C354-1b)** Vespidae. **(C355)** Sphecidae. **(C357)** †Ephialtitoidea.

**Chars. C357-361.**
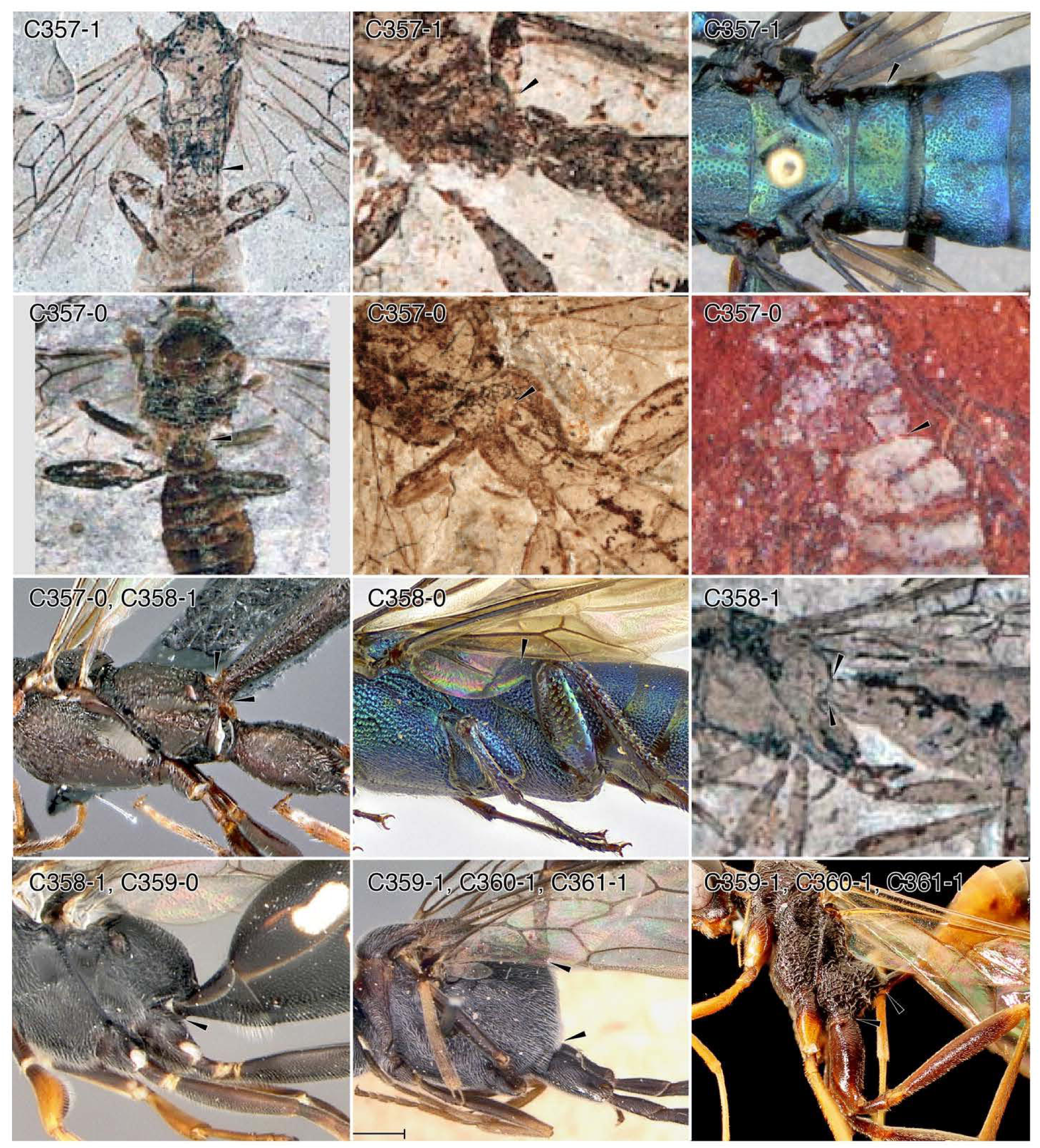
Character-states: (C357-1) Abdominal segment II as broad as I; (a) †*Praeproapocritusflexus;* (b) †*Eonevania robusta;* (c) *Chalinus haugi.* **(C357-0)** Abdominal segment II narrowed relative to I; (a) †*Symphytopterus graciler,* (b) †*Karataus exilis;* (c) †Praeaulacidae: †*Gulgonga beattiei;* (d) *Foenatopus* sp. **(C358-0)** Abdominal segment II not dorsoventrally constricted anteriorly; *C. timnaensis.* **(C358-1)** Abdominal segment II dorsoventrally constricted anteriorly; (a) *Foenatopus* sp.; (b) †*Kuafua polyneura,* (c) *Trigonalys* sp. **(C359-0)** Metasomal articulation close to metacoxae; *Trigonalys* sp. **(C359-1)** Metasomal articulation separated from metacoxae; (a) *Zeuxevania* sp.; (b) *Pristaulacus stigmaticus.* **(C360-1)** Propodeum forming bridge ventrad metasomal articulation; (a) *Zeuxevania* sp.; (b) *Pri. stigmaticus.* **(C361-1)** Metasomal articulation above midheight of propodeum; (a) *Zeuxevania* sp.; (b) *Pri. stigmaticus.* **Families: (C357-1a, b, 357-0a, b, 358-1b)** †Ephialtitoidea. **(C357-1c, 358-0)** Orussidae. **(C357-0, 358-1a)** Stephanidae. **(C358-1c, 359-0)** Trigonalidae. **(C359-1a, 360-1a, 361-1a)** Evaniidae. **(C359-1b, 360-1b, 361-1b)** Aulacidae.

**Chars. C362-370.**
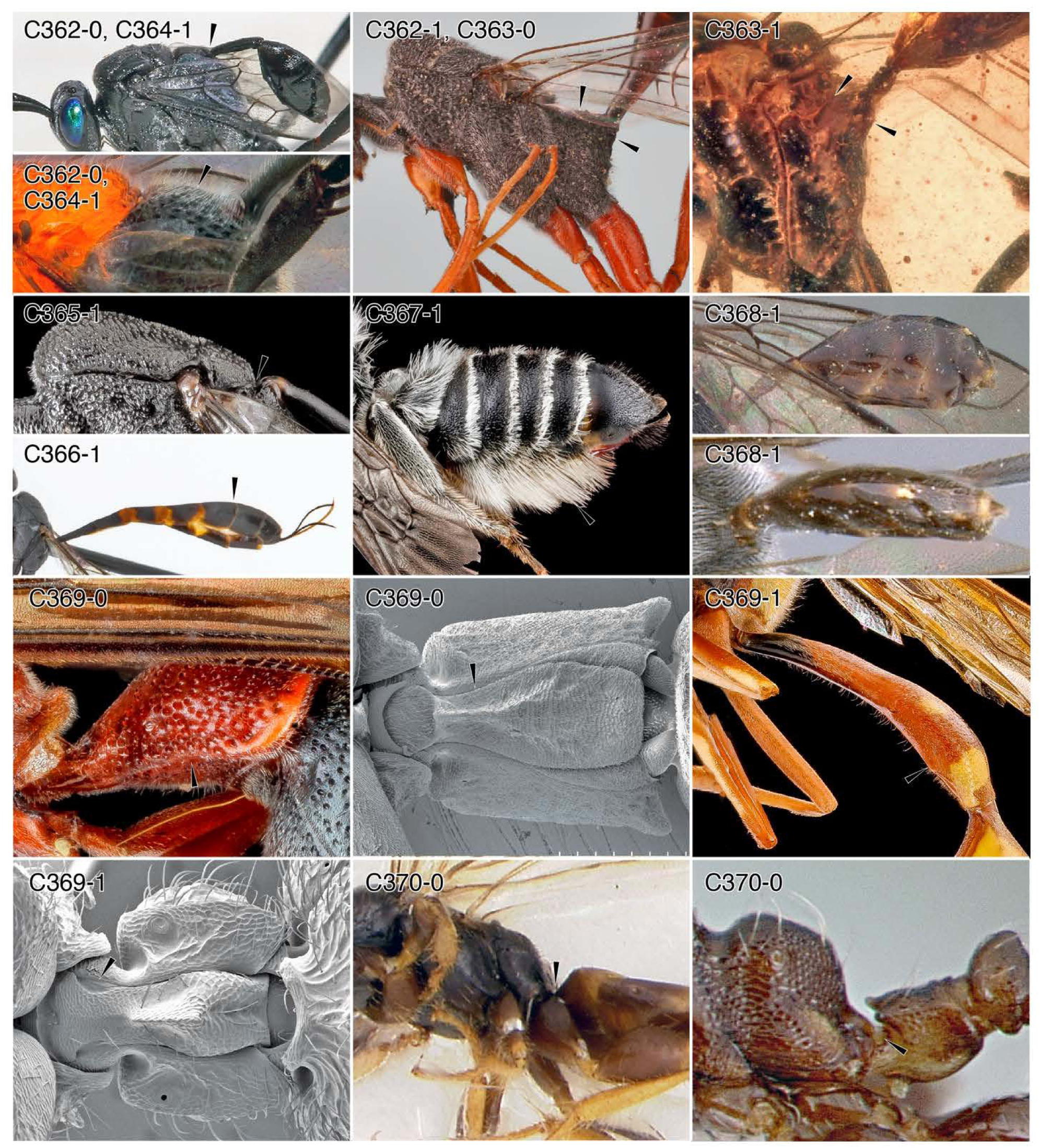
Character-states: (C362-0) Metasomal articulation not situated on conical projection; (a) *Evania appendigaster,* (b) evaniid genus and species indet. **(C362-1)** Metasomal articulation situated on conical projection; *Pristaulacus* sp. **(C363-0)** Propodeal cone distant from metanotum; *Pristaulacus* sp. **(C363-1)** Propodeal cone contacting metanotum; †*Archeofoenus tartaricus.* **(C364-1)** Propodeum convex between metanotum and elevated metasomal articulation; (a) *Evania appendigaster,* (b) evaniid genus and species indet. **(C365-1)** Metasomal articulation contacting metanotum; *Gasteruption* sp. **(C366-1)** Metasoma long, thin, tubular, with segments gradually broadening apically; *Gasteruption* cf. *kirbii* or *assectator.* **(C367-**1) Metasoma with ventral scopa; *Megachile pseudobrevis.* **(C368-1)** Metasoma after segment II short and lateromedially compressed, oval in form; *Zeuxevania* sp. **(C369-0)** Abdominal segment II without tergosternal fusion; (a) *Leptochilus acolhuus;* (b) *Platythyrea punctata.* **(C369-1)** Abdominal segment II with tergosternal fusion; (a) vespid genus and species indet.; (b) *Proceratium croceum.* **(C370-0)** Abdominal segment II (petiole) anterior foramen sessile; (a) *Ycaploca evansï,* (b) *Xenomyrmex panamanus.* **Families: (C362, 364, 368)** Evaniidae. **(C362-1, 363)** Aulacidae. **(C365, 366)** Gasteruptiidae. **(C367)** Megachilidae. **(C369-0a, C369-1a)** Vespidae. **(C369-0b, C369-1b, C370-0b)** Formicidae. **(C370-0a)** Scolebythidae.

**Chars. 370-375.**
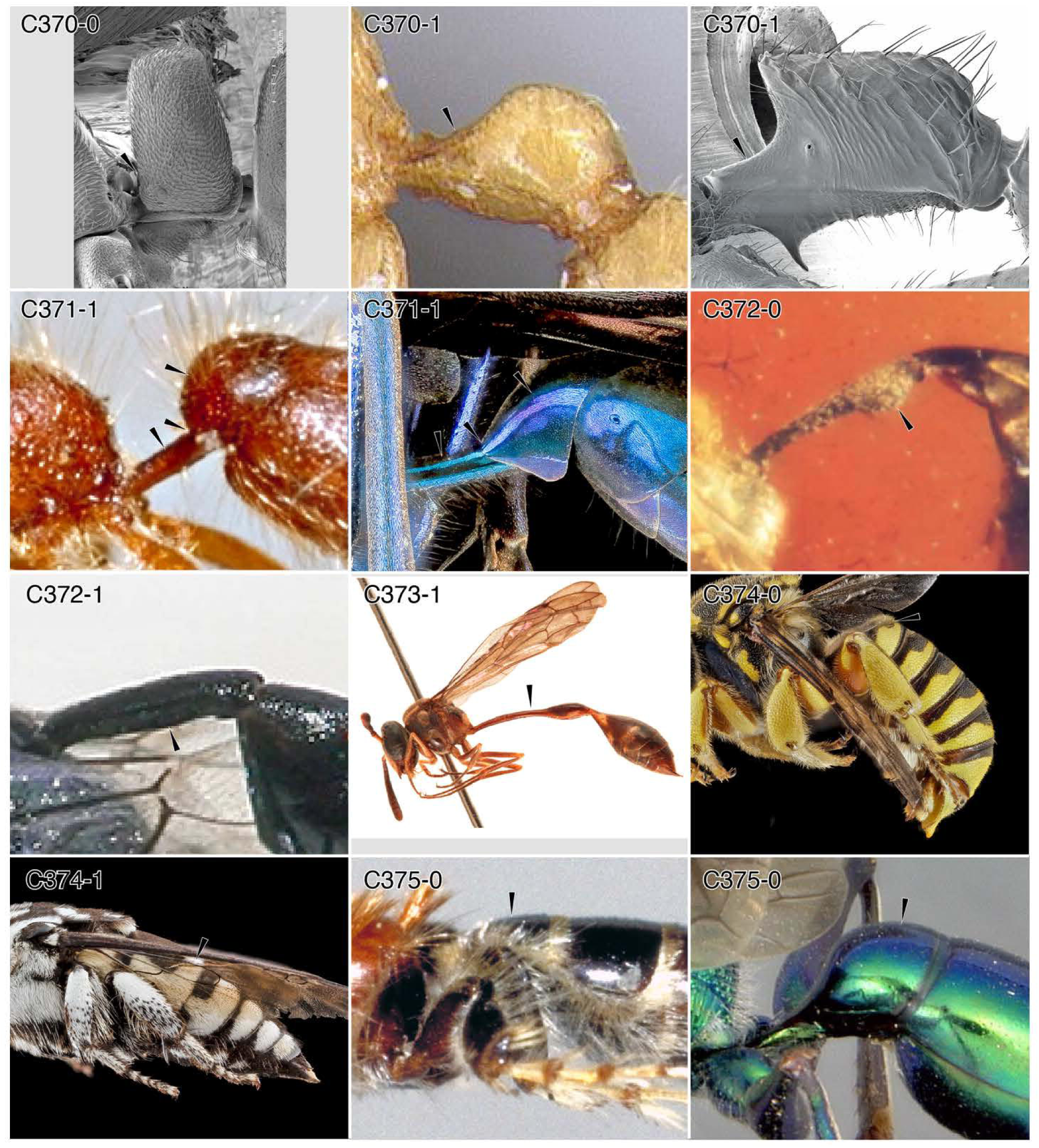
Character-states: (C370-0) Abdominal segment II (ASII, petiole) anterior foramen sessile; *Hypoponera* mx0l. **(C370-1)** ASII anterior foramen subsessile to pedunculate; (a) *Martialis heureka;* (b) *Paraponera clavata.* **(C371-1)** ASII peduncle comprising sternum only, tergum restricted posteriorly; (a) *Chyphotes* (female); (b) *Chalybion californicum.* **(C372-0)** ASII not in form of straight, narrow tube; †*Cretevania bechlyi.* **(C372-1)** ASII in form of straight, narrow tube; *Evania appendigaster.* **(C373-1)** ASII longer than head and mesosoma, as well as “gaster”; *Parischnogaster alternata.* **(C374-0)** ASII short; *Trachusa dorsalis.* **(C374-1)** ASII longer than at least following two segments; *Thyreus* sp. **(C375-0)** ASII without node, *i.e.,* without distinct posterodorsal surface; (a) *Bradynobaenus sp.;* (b) *Ampulex bredoi.* **Families: (C370)** Formicidae. **(C371-1a)** Chyphotidae. **(C371-1b)** Sphecidae. **(C372)** Evaniidae. **(C373)** Vespidae. **(C374-0)** Megachilidae. **(C374-1)** Apidae. **(C375-0a)** Bradynobaenidae. **(C375-0b)** Ampulicidae.

**Chars. 375-382.**
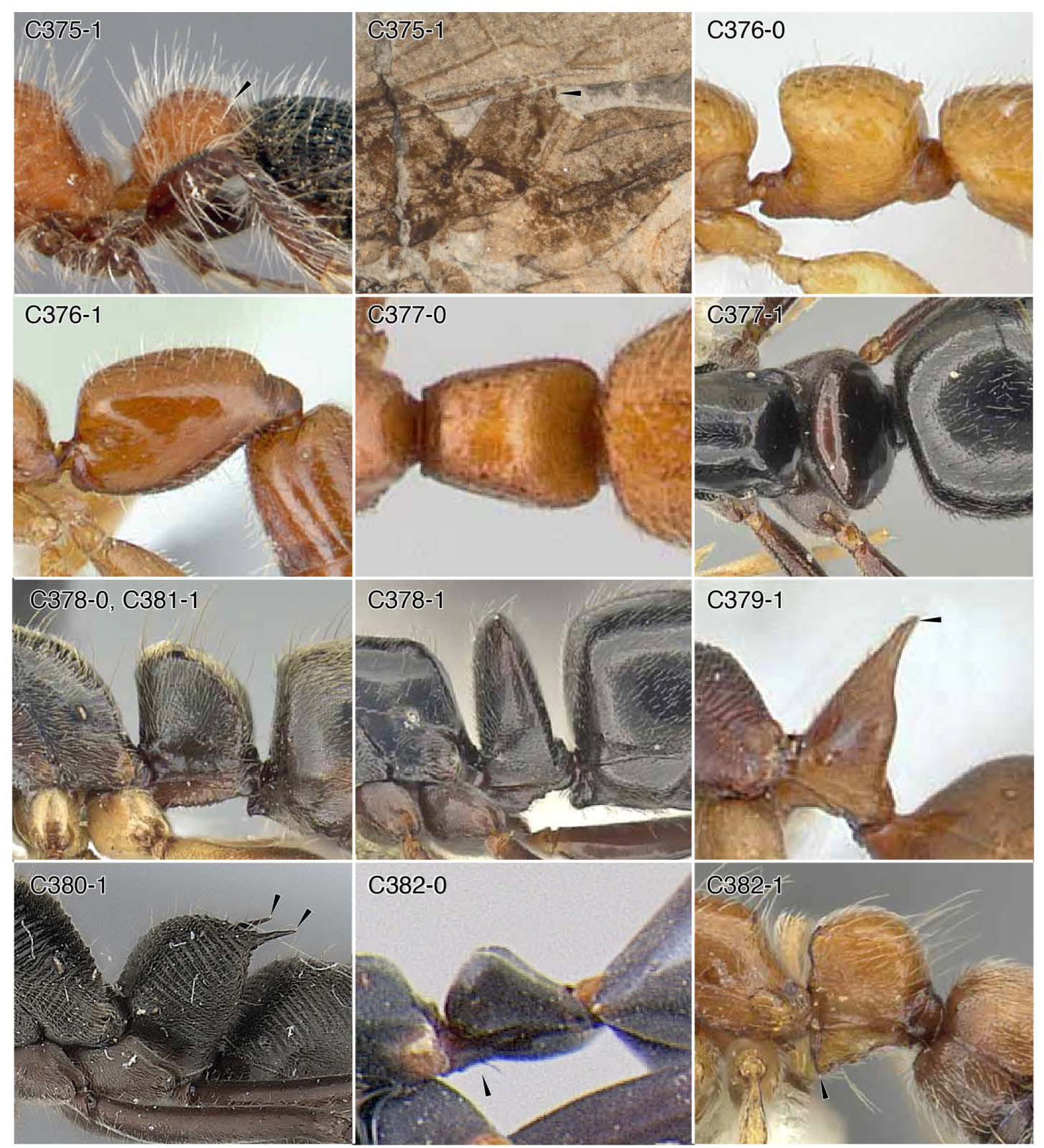
Character-states: (C375-1) Abdominal segment II (ASII) with node, *i.e.,* with distinct posterodorsal surface; (a) *Apterogyna globularia;* (b) †*Armania robusta.* **(C376-0)** ASII node in lateral view as tall as long or taller than long; *Apomyrma stygia.* **(C376-1)** ASII node in lateral view longer than tall; *Opamyrma hungvuong.* **(C377-0)** ASII node in dorsal view longer than wide or as wide as long; *Feroponera ferox.* **(C377-1)** ASII node in dorsal view wider than long; *Hypoponera nitidula.* **(C378-0)** ASII node not squamiform; *Neoponera unidentata.* **(C378-1)** ASII node squamiform; *H. nitidula.* **(C379-1)** ASII node with single dorsal spine; *Odontomachus meinerti.* **(C380-1)** ASII node with pair of dorsal spines; *Diacamma rugusom.* **(C381-1)** ASII node in lateral view with vertical anterior margin and broadly curving dorsal/posterior margin; *N. unidentata.* **(C382-0)** ASII sternum without anteroventral process; *Leptomyrmex erythrocephalus.* **(C382-1)** ASII sternum with anteroventral process; *Lioponera longitarsus.* **Families: (C375-1a)** Bradynobaenidae. **(C375-1b, C376-382)** Formicidae.

**Chars. 383-390.**
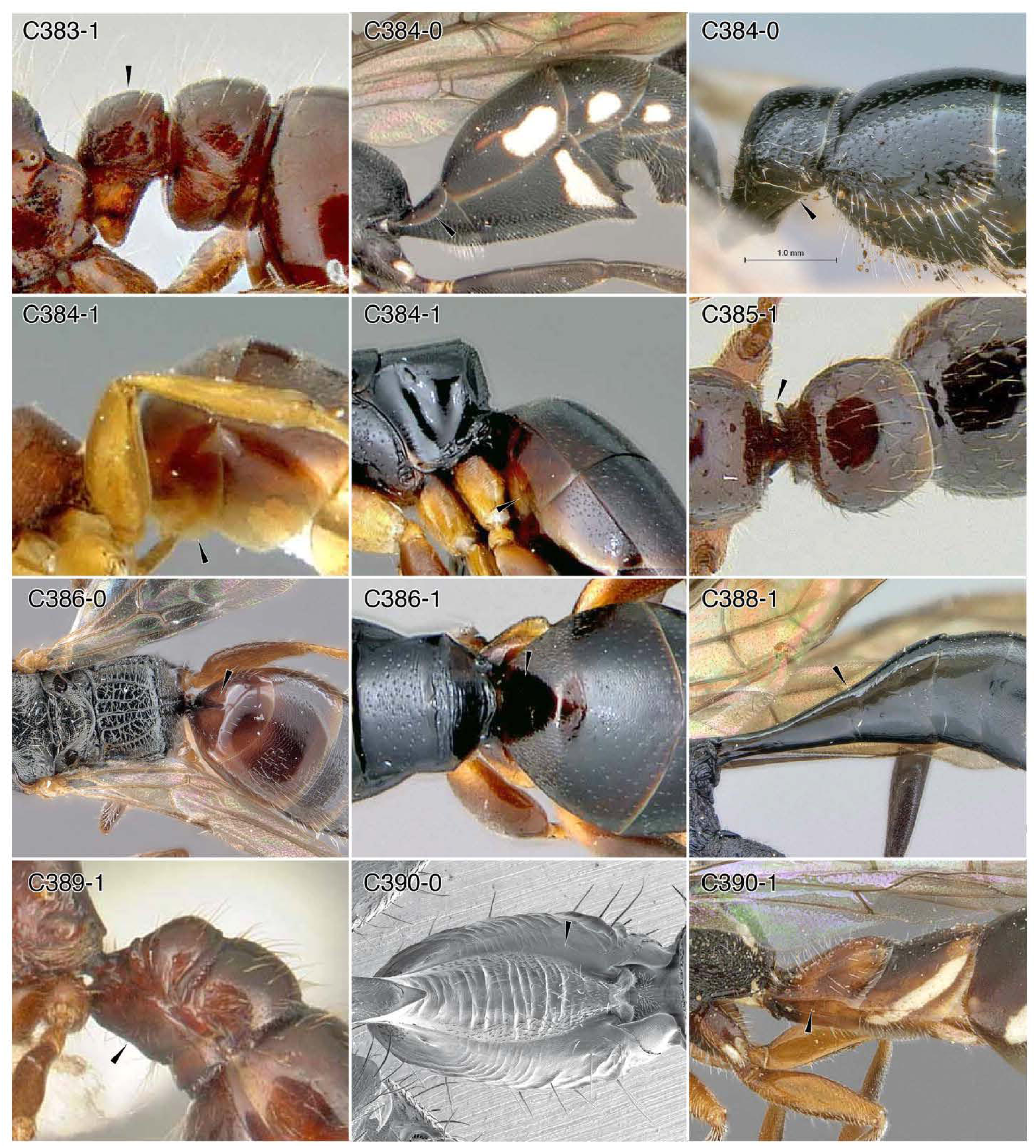
Character-states: (C383-1) Abdominal segment II (ASII) twice as tall as long, posterior articulation raised above anterior articulation; *Tatuidris tatusia.* **(C384-0)** ASII tergum not broadly overlying sternum; (a) *Trigonalys* sp.; (b) *Tiphia alecto.* **(C384-1)** ASII tergum broadly overlying sternum posteriorly, narrowed anteriorly; (a) *Embolemus africanus;* (b) *Obenbergerella aenigmatica.* **(C385-1)** ASII tergum with one or two anterior carinae/processes; *Amblyopone australis.* **(C386-0)** ASII tergum without anteromedian concavity (sulcus shown); *Disepyris semiruber.* **(C386-1)** ASII tergum with anteromedian concavity that fits against propodeum; *O. aenigmatica.* **(C388-1)** ASII-III terga fused, forming conical or funnel-shaped petiole; *Pristaulacus pilotoi.* **(C389-1)** ASII-III sterna fused; *Anomalomyrma taylori.* **(C390-0)** ASII without distinct laterotergite; *Paraponera clavata.* **(C390-1)** ASII with distinct laterotergite; *Sapygina enderleini.* **Families: (C383, 385, 389, 390-0)** Formicidae. **(C384-0a)** Trigonalidae. **(C384-0b)** Tiphiidae. **(C384-1a)** Embolemidae. **(C384-1b, 386-1)** Chrysididae. **(C386-0)** Bethylidae. **(C388)** Aulacidae. **(C390-1)** Sapygidae.

**Chars. 390-397.**
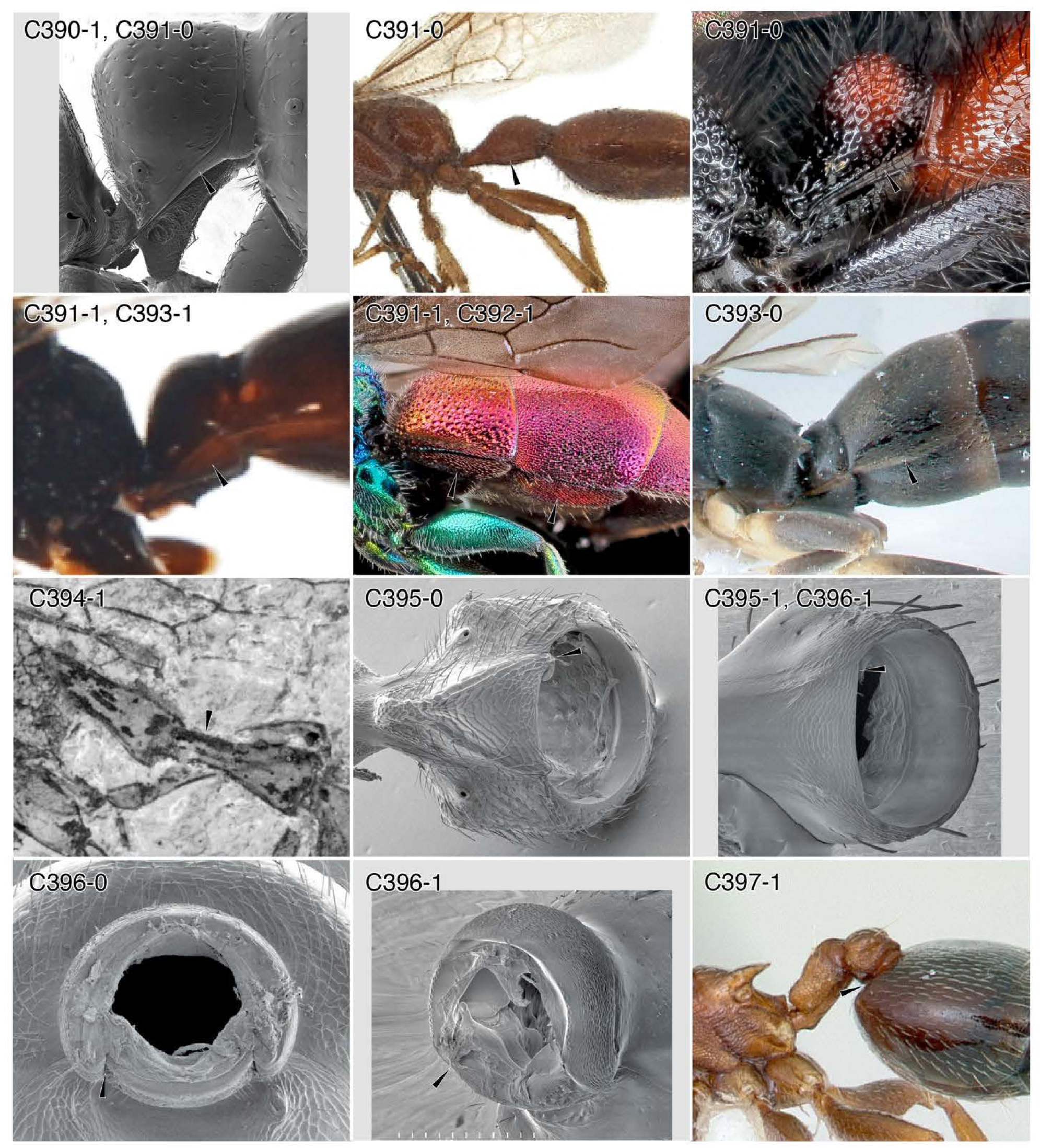
Character-states: (C390-1) Abdominal segment II (ASII) laterotergite present, defined by crease; *Amblyopone australis.* **(C391-0)** ASII laterotergite narrow; (a) *Amblyopone australis;* (b) *Brachycistis alcanor,* (c) mutillid genus and species indet. **(C391-1)** ASII laterotergite broad; (a) *Sierolomorpha canadensis;* (b) *Chrysura* sp. **(C392-1)** ASIII laterotergite present, defined by crease; *Chrysura* sp. **(C393-0)** ASIII laterotergite narrow; *Olixon toliaraensis.* **(C393-1)** ASIII laterotergite broad; *S. canadensis.* **(C394-1)** ASIII narrow, tubular; †*Nevania ferocula.* **(C395-0)** Posterior petiolar foramen not forming even circle, outline interrupted; *Tatuidris tatusia.* **(C395-1)** Posterior petiolar foramen forming even circle; *Pogonomyrmex barbatus.* **(C396-0)** Postpetiolar helcium (ASIV) pretergite overlapping presternite; *Tatuidris tatusia.* **(C396-1)** Postpetiolar helcium pretergite not overlapping presternite, forming more-or-less even circle; *Pogonomyrmex barbatus.* {C397-1) Postpetiolar helcium migrated dorsally; *Crematogaster pinicola.* **Families: (C390, 391-0a, 395-397)** Formicidae. **(C391-0b)** Tiphiidae. **(C391-0c)** Mutillidae. **(C391-1a, C393-1)** Sierolomorphidae. **(C391, 392)** Chrysididae. **(C393-0)** Rhopalosomatidae. (393-0) Rhopalosomatidae. **(C394)** †Praeaulacidae.

**Chars. 397-403.**
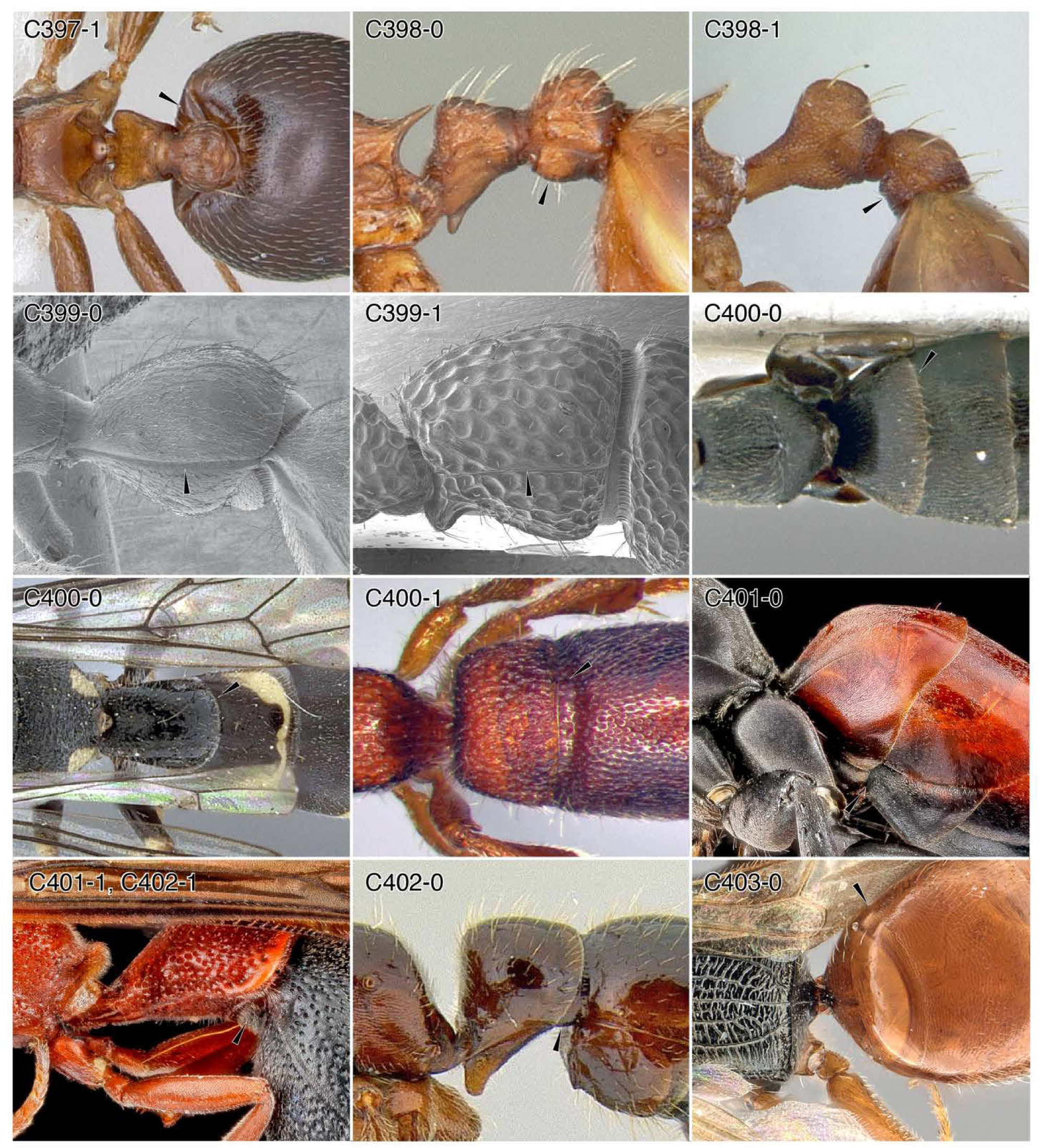
Character-states: (C397-1) Postpetiolar helcium migrated dorsally; *Crematogaster pinícola.* **(C398-0)** Postpetiolar sternum bulging ventrally; *Myrmica brevispinosa.* **(C398-1)** Postpetiolar sternum flat or concave; *Wasmannia lutzi.* **(C399-0)** Postpetiole without tergosternal fusion; *Pseudomyrmex gracilis.* **(C399-1)** Postpetiole with tergosternal fusion; *Parasyscia nitidula.* **(C400-0)** Abdominal segment III (ASIII) tergum without distinct line or sulcus dividing pre- and posttergite; (a) *Sclerogibba crassifemorata,* (b) *Sapygina lobengulae.* **(C400-1)** ASIII tergum with distinct line or sulcus dividing pre- and posttergite; *Rhopalomutilla carinaticeps.* **(C401-0)** ASIII sternum without distinct line or sulcus dividing pre- and poststernite; *Drepanaporus collaris.* **(C401-1)** ASIII sternum with distinct line or sulcus dividing pre- and poststernite; *Leptochilus acolhuus.* **(C402-0)** ASIII presternite exposed in lateral view; *Amblyopone australis.* **(C402-1)** ASIII presternite concealed in lateral view; *Leptochilus acolhuus.* **(C403-0)** ASIII anterior articulatory surfaces not narrowed; *Disepyris semiruber.* **Families: (C397-399, 402)** Formicidae. **(C400-0a)** Sclerogibbidae. **(C400-0b)** Sapygidae. **(C400-1)** Mutillidae. **(C401-0)** Pompilidae. **(C401-1, 402-1)** Vespidae. **(C403)** Bethylidae.

**Chars. 403-408.**
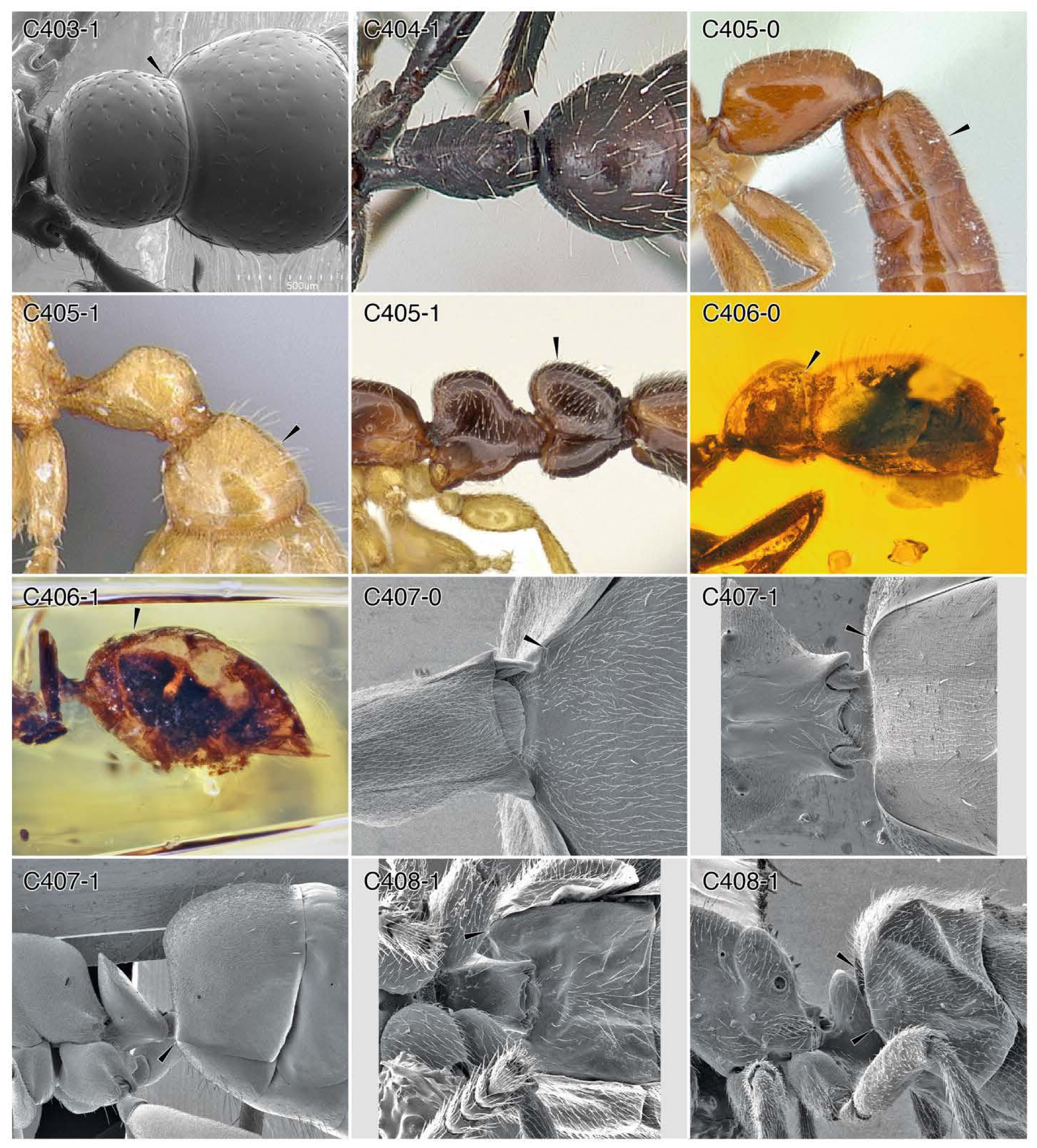
Character-states: (C403-1) Abdominal segment III (ASIII) anterior articulatory surfaces narrowed; *Amblyopone australis.* **(C404-1)** ASIII anterior articulatory surfaces strongly narrowed; *Paraponera clavata.* **(C405-0)** ASIII slightly reduced in size; *Opamyrma hungvuong.* **(C405-1)** ASIII strongly reduced in size, completely petiolated; (a) *Martialis heureka,* (b) *Protanilla bicolor.* **(C406-0)** ASIII not foreshortened; †*Gerontoformica pilosa.* **(C406-1)** ASIII foreshortened; †G. *orientalis* group. **(C407-0)** ASIII poststernite not or only very weakly “shouldered”; *Oecophylla smaragdina.* **(C407-1)** ASIII poststernite “low shouldered”; *Formica fusca* group. **(C408-1)** ASIII poststernite “high shouldered”, forming anteromedian groove which receives petiole; *Acropyga* sp. **Families: (C403-408)** Formicidae.

**Chars. 409-417.**
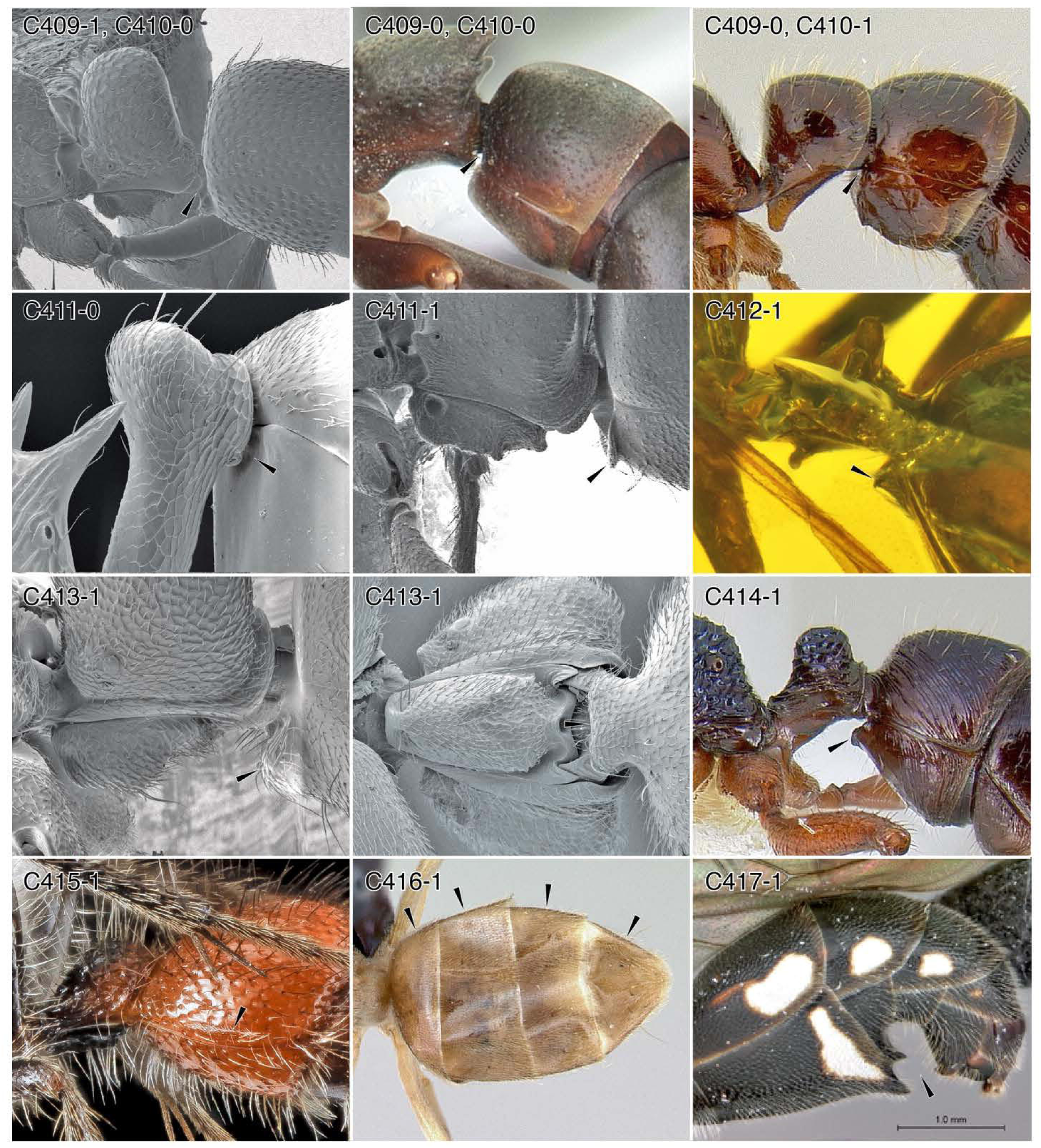
Character-states: (C409-1; C410-0) Helcium infraaxial (below segment midheight); *Emeryopone buttelreepeni.* **(C409-0; C410-0)** Helcium axial (at segment midheight); *Platythyrea mocquerysi.* **(C409-0; C410-1)** Helcium supraaxial (above segment midheight); *Amblyopone australis.* **(C411-0)** Abdominal sternum III (ASIII) without anteroventral processes (prora); *Aneuretus simoni.* **(C411-1)** ASIII with prora; *Ponera alpha.* **(C412-1)** Prora in form of keel; †*Zigrasimecia tonsora.* **(C413-1)** Prora transverse, lip-like; *Pseudoponera stigma.* **(C414-1)** Prora transverse, lip-like, strongly produced; *Rhytidoponera chalybaea.* **(C415-1)** AIII tergum with felt line; mutillid genus and species indet. **(C416-1)** “Gaster” with only four exposed terga; *Bothriomyrmex saundersi.* **(C417-1)** “Gaster” with sternal processes; *Trigonalys* sp. **Families: (C409-414, 416)** Formicidae. **(C415)** Mutillidae. **(C417)** Trigonalidae.

**Chars. 418-430.**
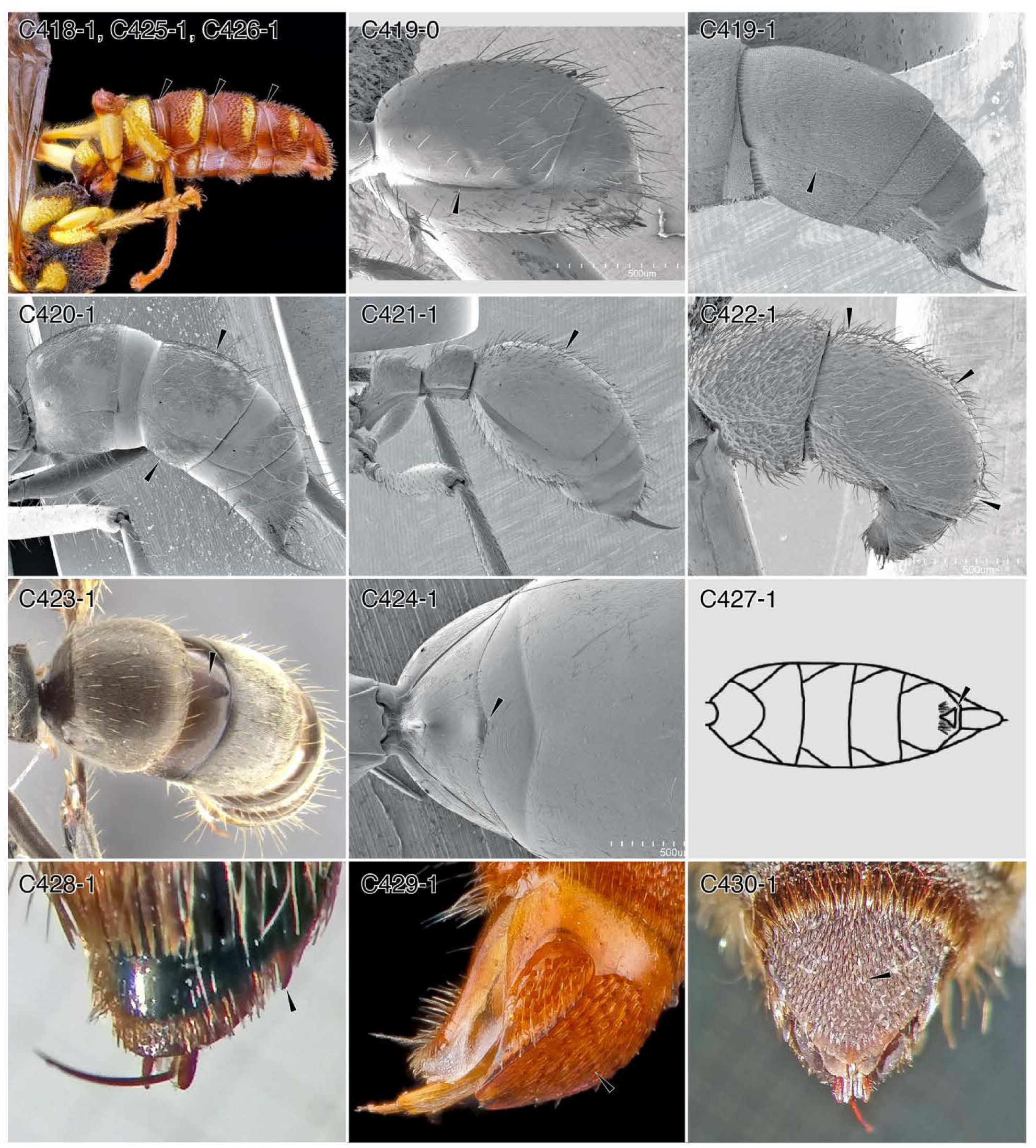
Character-states: (C418-1) Abdominal segment IV (ASIV) with transverse sulcus (cinctus); *Cerceris triungulata.* **(C419-0)** ASIV tergum (ASIV-T) overhanging sternum; *Labidus coecus.* **(C419-1)** ASIV-T aligned laterally with sternum; *Platythyrea punctata.* **(C420-1)** ASIV-T longer than sternum; *Neoponera villosa.* **(C421-1)** ASIV-T large, longer than both ASII-III and ASV-VII; *Manica rubida.* **(C422-1)** ASIV-T vaulted; *Proceratium croceum.* **(C423-1)** ASIV-T with anteromedian stridulitrum; *Ne. antecurvata.* **(C424-1)** ASIV sternum (ASIV-S) with anteromedian stridulitrum; *Nothomyrmecia macrops.* **(C425-1)** ASV with cinctus; *Ce. triungulata.* **(C426-1)** ASVI with cinctus; *Ce. triungulata.* **(C427-1)** ASVII-S with posteromedian triangular process; *Clystopsenella longiventris.* **(C428-1)** ASVII-S with paired horn-like processes laterally; *Campsomeris pilipes.* **(C429-1)** ASVII-T modified as “pygidial plate”; *Oxybelus analis.* **(C430-1)** ASVII-T covered with appressed, stout, squamiform setae; *Ca. pilipes.* **Families: (C418, 425, 426)** Philanthidae. **(C419-424)** Formicidae. **(C427)** Scolebythidae. **(C428, 430)** Scoliidae. **(C429)** Crabronidae.

**Chars. 431-438.**
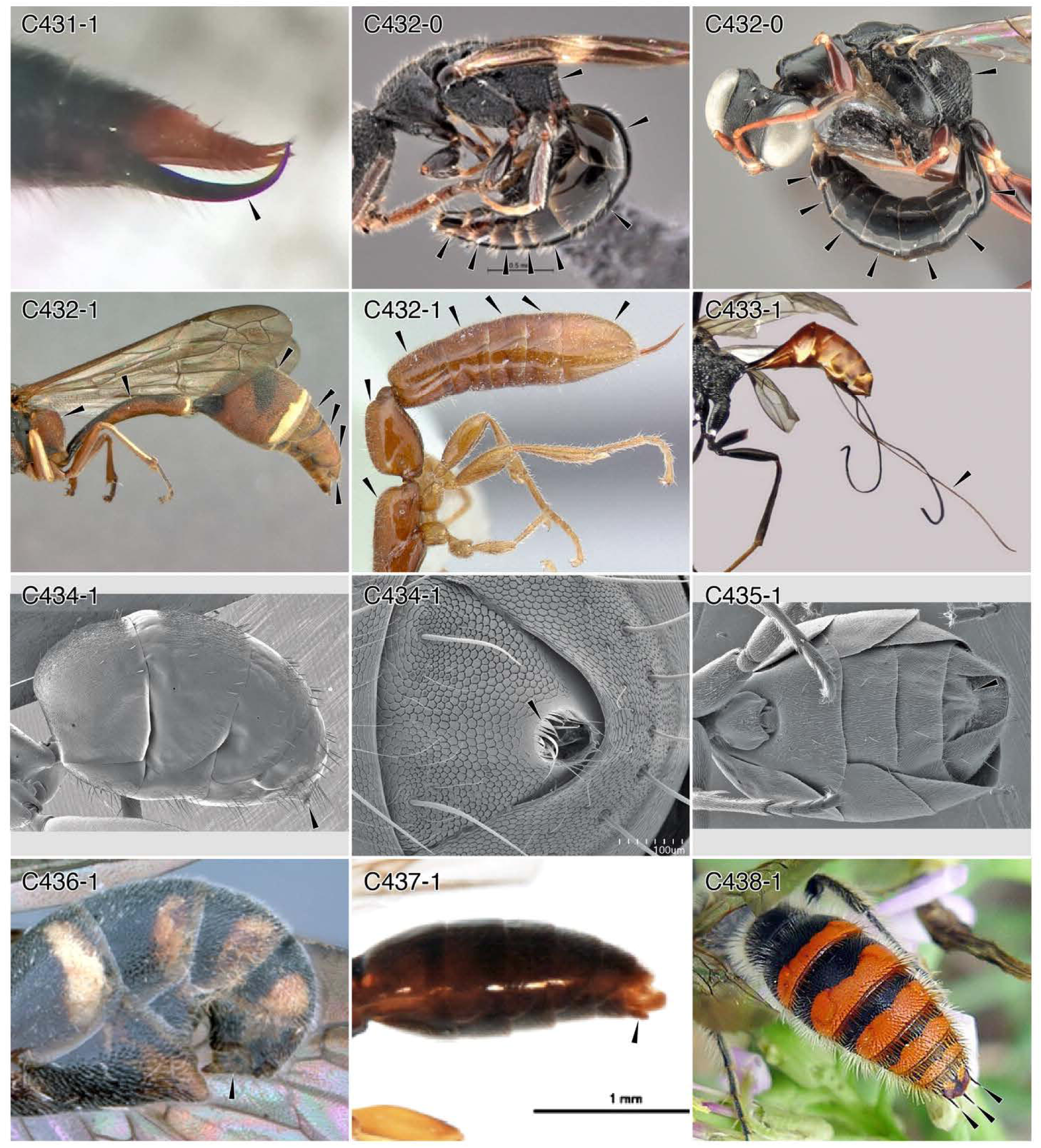
Character-states: (C431-1) Abdominal sternum VII (ASVII-S) of male unciform; Methochinae genus and species indet. **(C432-0)** ASVIII tergum (ASVIII-T) of female not completely internalized; (a) *Incertosulcus* sp.; (b) *Dryinus mobotensis.* **(C432-1)** ASVIII-T of female completely internalized; (a) *Eumenes amoldr,* (b) *Opamyrma hungvuong.* **(C433-1)** Ovipositor long, being as long or longer than metasoma; *Pristaulacus compressus.* **(C434-1)** ASVII-S apically curved and lined with hair, forming acidopore and coronula; *Formica fusca* group. **(C435-1)** Sting reduced and acidopore absent; *Tapinoma simrothi.* **(C436-1)** Female metasoma apex downcurved; *Taeniogonalys gracilicornis.* **(C437-1)** ASIX-S of male peg-like; *Sierolomorpha canadensis.* **(C438-1)** ASIX-S of male with three wide-spaced spiniform processes; *Campsomeriella aureola.* **Families: (C431)** Tiphiidae. **(C432-0a)** Bethylidae. **(C432-0b)** Dryinidae. **(C432-1a)** Vespidae. **(C432-1b, 434, 435)** Formicidae. (433) Aulacidae. **(C436)** Trigonalidae. **(C437)** Sierolomorphidae. **(C438)** Scoliidae.

**Chars. 439-450.**
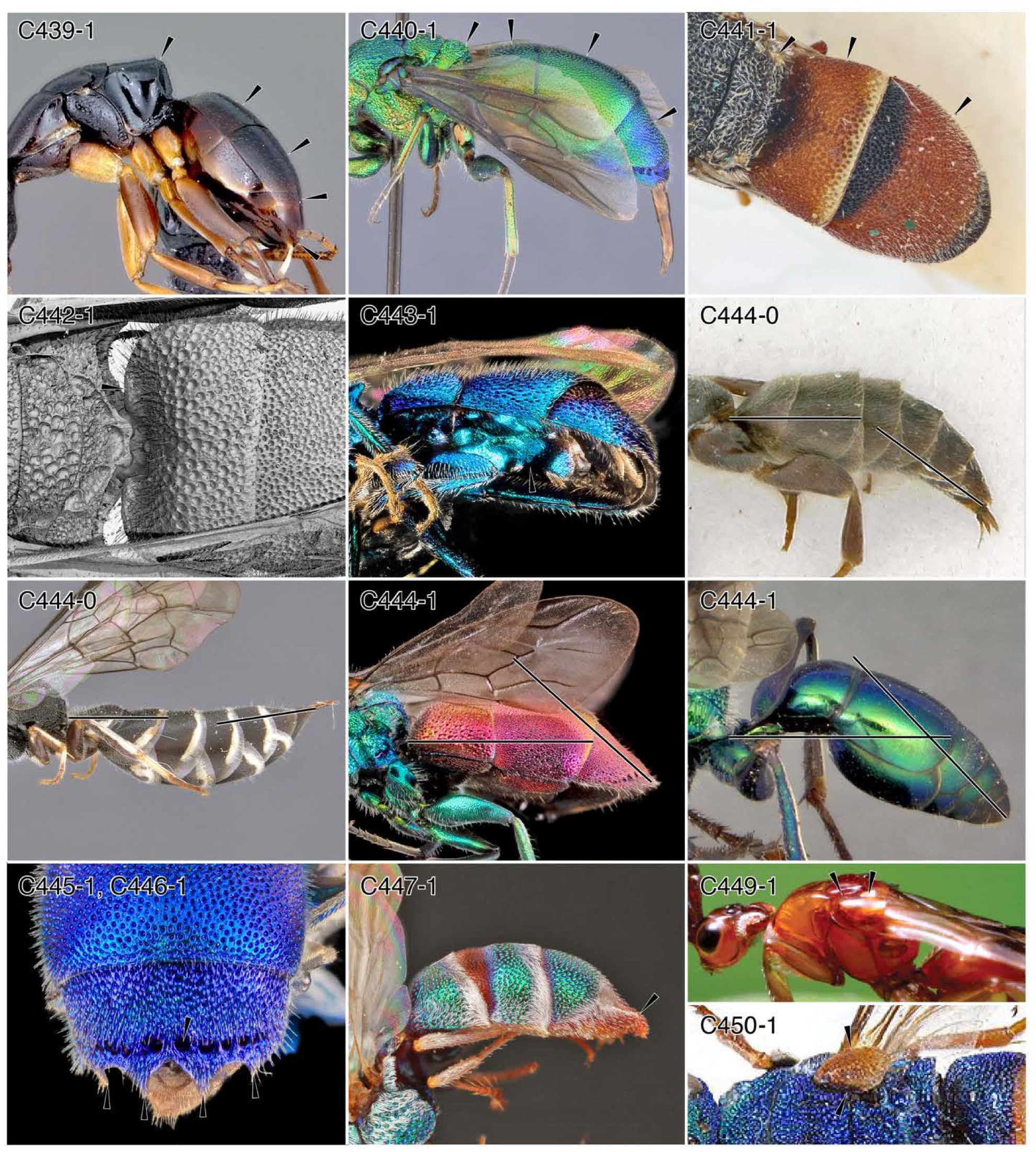
Character-states: (C439-1) Abdomen of female with ≤ 5 exposed segments, ≤ 6 in male; *Obenbergerella aenigmatica.* **(C440-1)** Abdomen of female with ≤ 4 exposed segments, ≤ 5 in male; *Chrysis eximia.* **(C441-1)** Abdomen of female with 3 exposed segments; *Allocoelia bidens.* **(C442-1)** Abdominal segment II with dorsoventrally compressed processes lateral to the propodeal articulation; *Chrysura* sp. **(C443-1)** Metasoma concave ventrally; *Coenochrysis doriae.* **(C444-0)** Abdominal terga II and III not longer than IV+; (a) *Sclerogibba crassifemorata,* (b) *Sapygina manica.* **(C444-1)** Abdominal terga II and III longer than IV+; (a) *Chrysura* sp.; (b) *Ampulex bredoi.* **(C445-1)** Abdominal tergum IV with distal rim of denticles; *Chrysis propria.* **(C446-1)** Abdominal tergum IV with transverse row of foveae; *Chrysis propria.* **(C447-1)** Abdominal tergum IV apical margin downcurved and thickened; *Parnopes fulvicornis.* **(C449-1)** Tegulum massive, locking into mesonotal notch; *P. grandior.* **(C450-1)** Tegulum massive, not locking into mesonotal notch; *Loboscelidia* sp. **Families: (C439-443, C444-1a, 445-450)** Chrysididae. **(C440-0a)** Sclerogibbidae. **(C440-0b)** Sapygidae. **(C444-1b)** Ampulicidae.

**Chars. 451-457.**
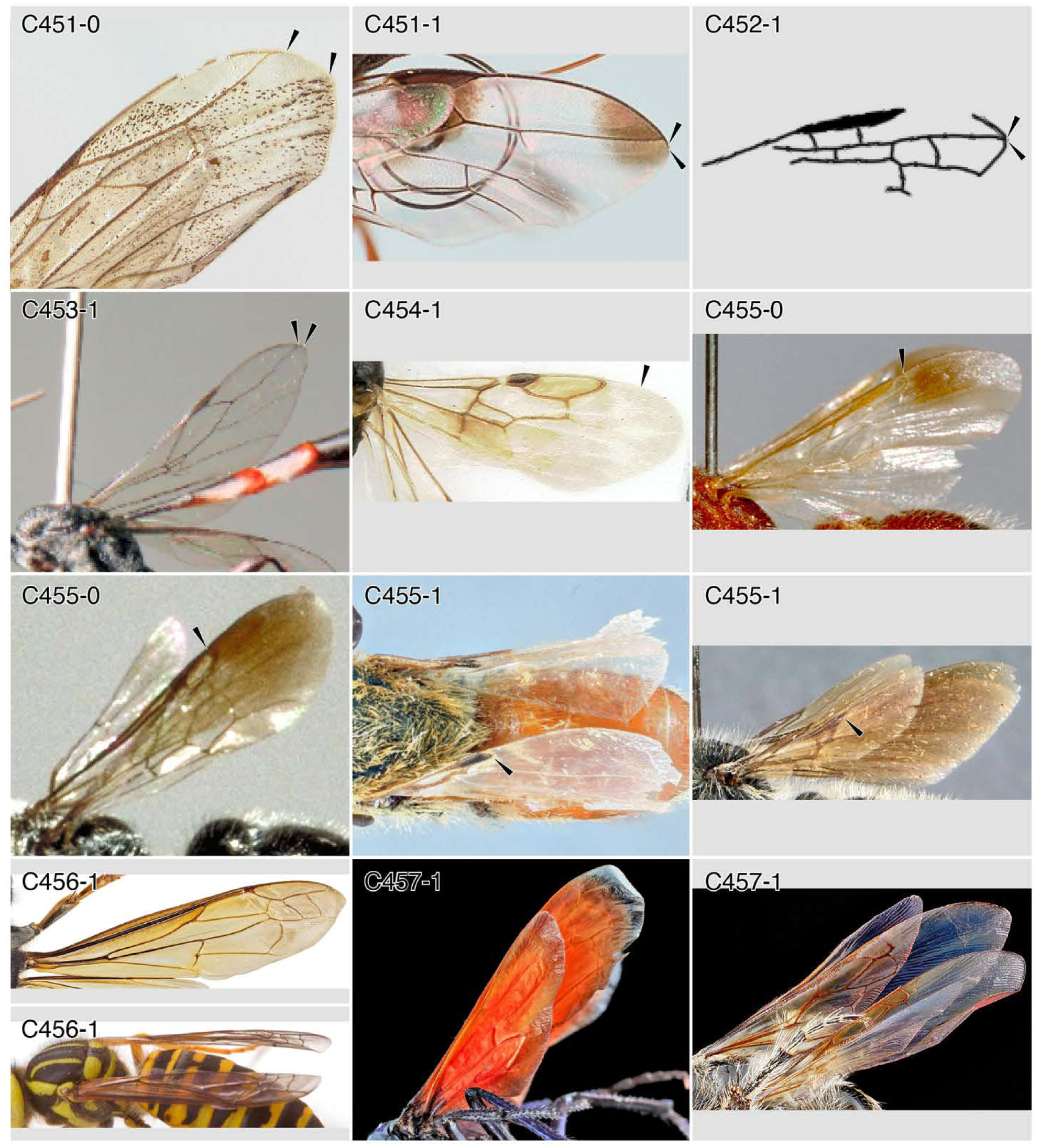
Character-states: (C451-0) Free radial vein (Rf) not nearing wing apex; *Paniscomima bekilyi.* **(C451-1)** Rf nearing or reaching wing apex; *Pristaulacus* sp. **(C452-1)** Free radial sector vein (Rsf) contacting wing apex; †*Proapocritus elegans.* **(C453-1)** Rsf nearing or reaching wing apex; *Gasteruption caucasicum.* **(C454-1)** Distal 1/4 to 1/3 of wing without tubular or nebulous venation; *Ycaploca evansi.* **(C455-0)** Venation not in “bradynobaenid” pattern”; (a) *Chyphotes* (male); (b) *Typhoctes peculiaris* (male). **(C455-1)** Venation in “bradynobaenid” pattern”, with heavily sclerotized veins restricted to proximal half; (a) *Bradynobaenus gayr,* (b) *Apterogyna globularia.* **(C456-1)** Fore wing with longitudinal fold line; (a) *Vespa velutina nigrithorax;* (b) *Vespula squamosa.* **(C457-1)** Wings with apical wrinkles; (a) *Pepsis rubra,* (b) scoliid genus and species indet. **Families: (C451-0)** Rhopalosomatidae. **(C451)** Aulacidae. **(C452)** †Ephialtitoidea. **(C453)** Gasteruptiidae. **(C454)** Scolebythidae. **(C455-0)** Chyphotidae. **(C455)** Bradynobaenidae. **(C456)** Vespidae. **(C457-1a)** Pompilidae. **(C457-1b)** Scoliidae.

**Chars. 458-468.**
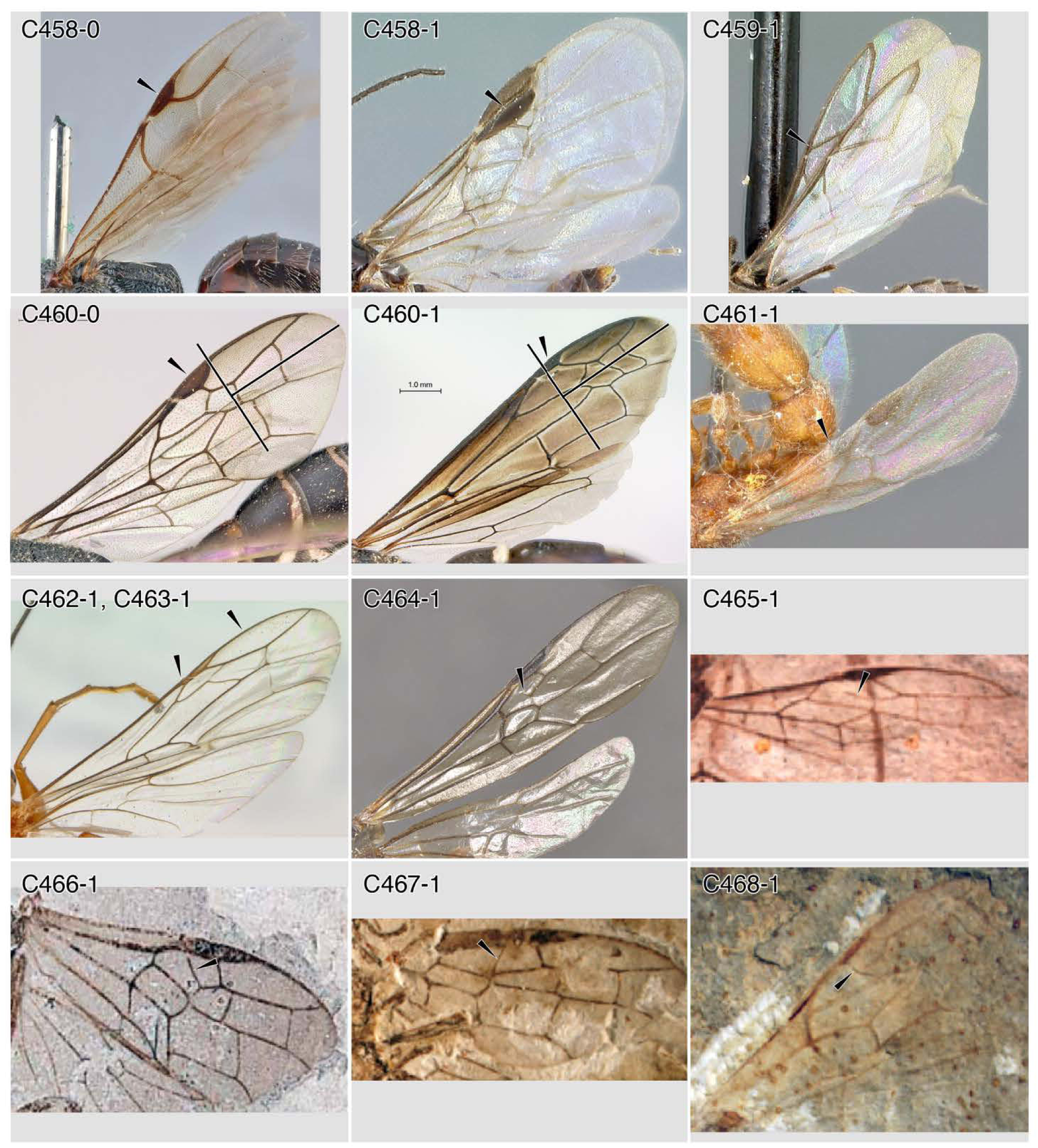
Character-states: (C458-0) Pterostigma (Ptstg.) not enlarged; *Pristocera armaticeps.* **(C458-1)** Ptstg. grossly enlarged; *Myrmecopterina* sp. **(C459-1)** Ptstg. strongly reduced to absent; *Sclerogibba tumeri.* **(C460-1)** Ptstg. not in apical third; *Sapygina enderleini.* **(C460-1)** Ptstg. in apical third; *Allepipona* sp. **(C461-1)** Costa **(C) incomplete or absent;** *Lioponera reticulata.* (C462-1) C and Sc+R+Rs contacting for most of their length; *Paniscomima bekilyi.* **(C463-1)** R present, tubular distal to ptstg.; *Pa. bekilyi.* **(C464-1)** 1r-rs present, even if as stub; *Myrmecia auriventris.* **(C465-1)** 1r-rs more-or-less (+/-) longitudinally oriented; †*Acephialtitia colossa.* **(C466-1)** 1r-rs +/- longitudinal and complete; †*Praeproapocritus flexus.* **(C467-1)** 1r-rs +∕-transverse and complete; †*Karataus orientalis.* **(C468-1)** 1r-rs complete and uniquely curved; †*Sinoproscolia yangshuwanziensis.* **Families: (C458-0)** Bethylidae. **(C458-1)** Plumariidae. **(C459)** Sclerogibbidae. **(C460-1a)** Sapygidae. **(C460-1b)** Vespidae. **(C461, 464)** Formicidae. **(C462, 473)** Rhopalosomatidae. **(C465-467)** †Ephialtitoidea. **(C468)** Scolioidea.

**Chars. 469-476.**
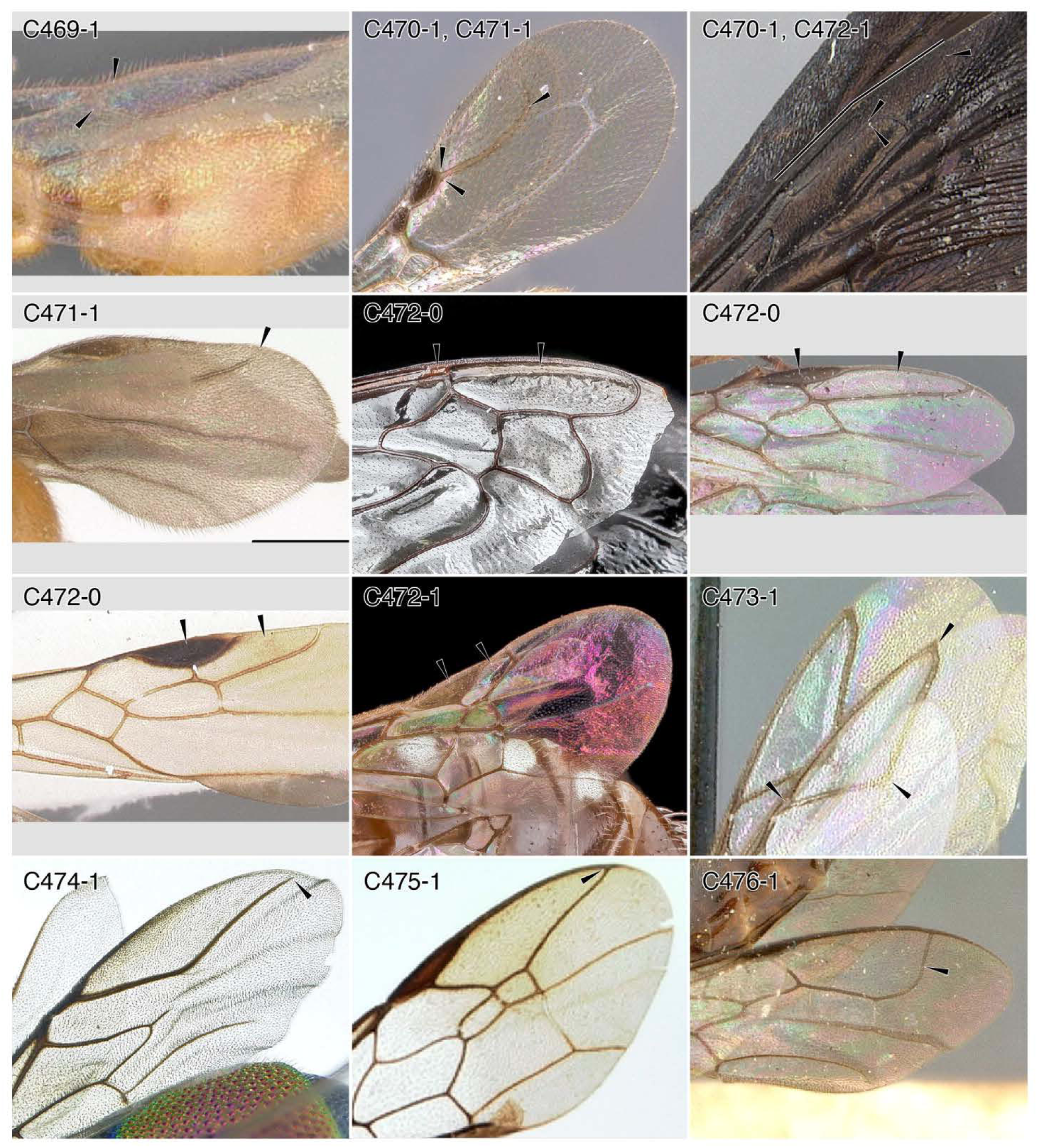
Character-states: (C469-1) 2r-rs anterior junction at or near proximal apex of pterostigma (ptstg.); *Apomyrma stygia.* **(C470-1)** 2r-rs anterior junction at or near distal apex of ptstg.; (a) *Goniozus kiefferi;* (b) *Scolia erythropyga.* **(C471-1)** Rs not reaching R or wing apex; (a) *Goniozus kiefferi;* (b) *Tatuidris tatusia.* **(C472-0)** Marginal cell 1 (2rrs) larger ptstg.; (a) bembicid. genus and species indet.; (b) *Pseudomyrmex gracilis;* (c) *Platythyrea mocquerysi.* **(C472-1)** 2rrs shorter or smaller than ptstg.; (a) *Scolia erythropyga,* (b) *Perdita sexmaculata.* **(C473-1)** 2rrs triangular; *Sclerogibba tumeri.* **(C474-1)** Rsf curving posteriorly; *Chrysis eximia.* **(C475-1)** 2rrs pointed to narrowly rounded; *Trigonalys micanticeps.* **(C476-1)** Rs widely curving anterad to Rf; *Zeuxevania* sp. **Families: (C469, 471-1b, 472-0b, c)** Formicidae. **(C470-1a, 471-1a)** Bethylidae. **(C470-1b, 472-1a)** Scoliidae. **(C472-0a)** Bembicidae. **(C472-1b)** Andrenidae. **(C473)** Sclerogibbidae. **(C474)** Chrysididae. **(C475)** Trigonalidae. **(C476)** Evaniidae.

**Chars. 476-482.**
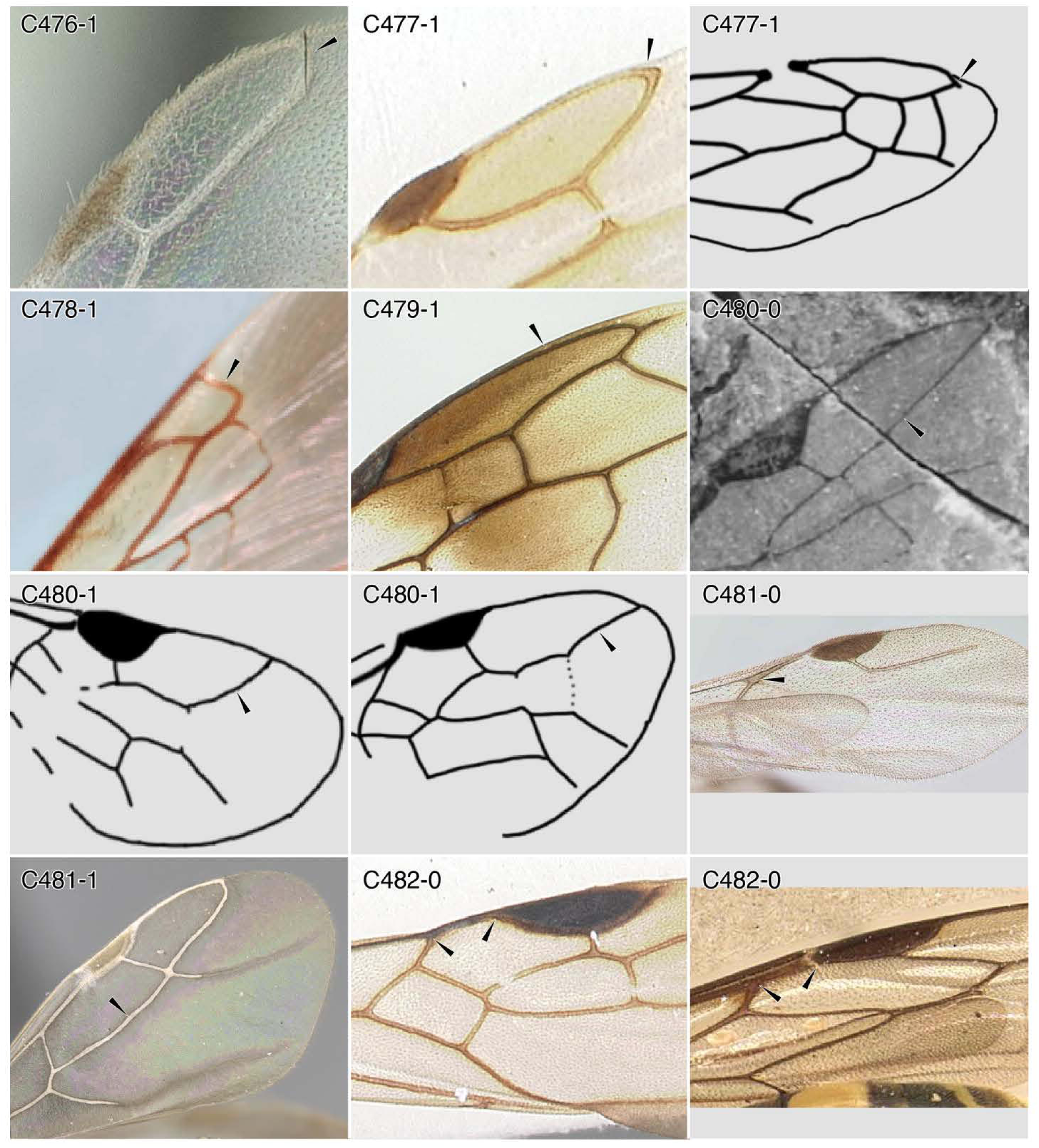
Character-states: (C476-1) Marginal cell 1 (2rrs) apex broadly rounding to nearly perpendicular; *Discothyrea mixta.* **(C477-1)** Rf apex separating at its from anterior wing margin; (a) *Ycaploca evansi;* (b) †*Curiosivespa derivata.* **(C478-1)** Rsf curving narrowly to R, nearly perpendicular; *Campsomeriella minutalis.* **(C479-1)** Rf apex separated for considerable length from the anterior wing margin; *Ampulex bredoi.* **(C480-0)** Distal Rsf abscissa not curving to R at about 45° from 3rs-m; †*Manlaya capelensis.* **(C480-1)** Distal Rsf abscissa curving to R at about 45° from 3rs-m; (a) †*Humiryssus cancellatus*; (b) †*Manlaya ockleyensis.* **(C481-0)** First free abscissa of Rs (Rsf1) absent; *Anomalomyrma* sp. **(C481-1)** Rsf1 present; *Lepisiota canescens.* **(C482-0)** Rsf1 separated widely from pterostigma; (a) *Crossocerus glabricomis;* (b) *Platythyrea mocquerysi.* **Families: (C476, 481, 482-0b)** Formicidae. **(C477-1a)** Scolebythidae. **(C477-1b)** Vespidae. **(C478)** Scoliidae. **(C479)** Ampulicidae. **(C480)** †Baissidae. **(C482-0a)** Crabronidae.

**Chars. 482-485.**
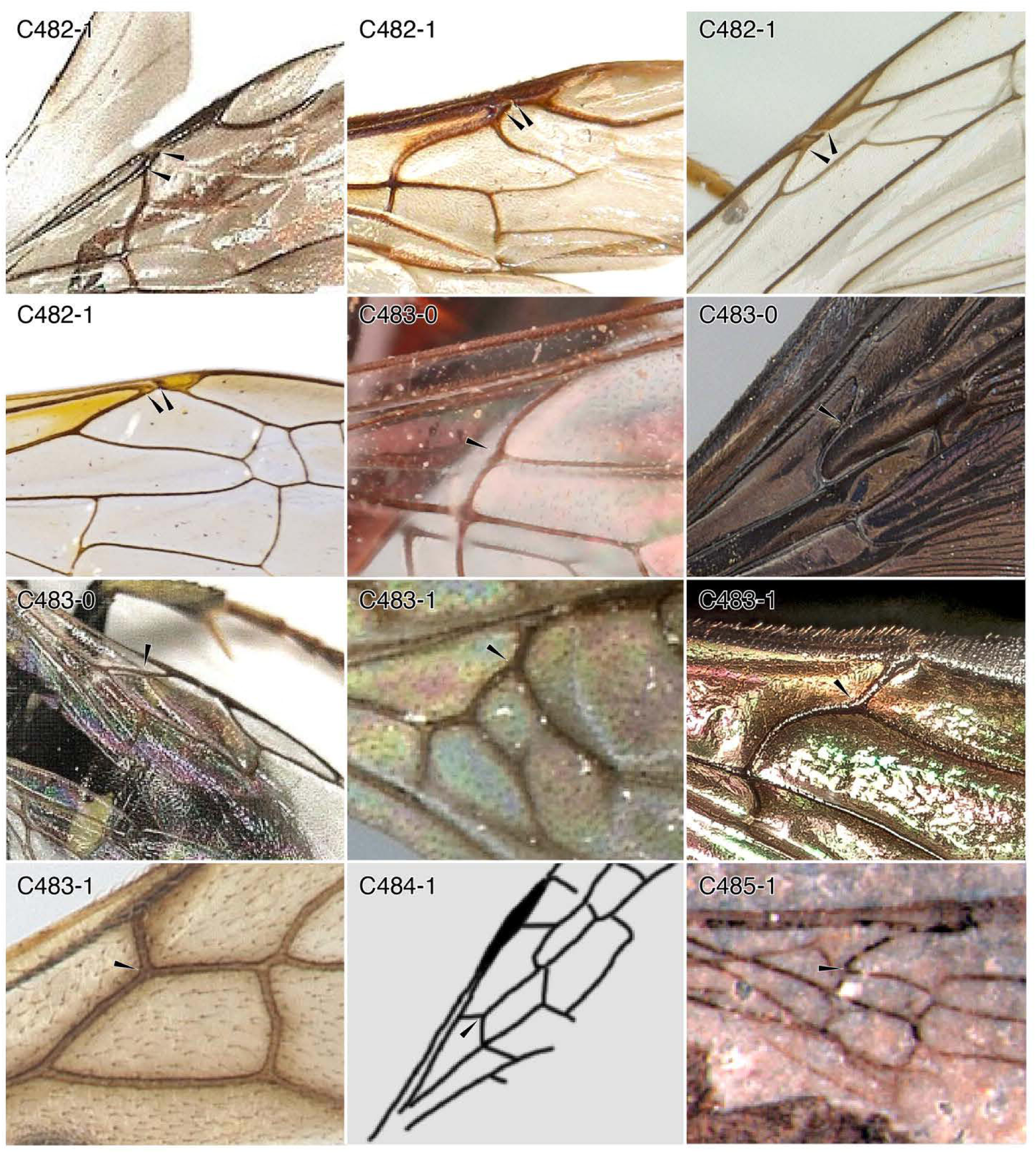
Character-states: (C482-1) Rsf1 and pterostigma closely approximated; (a) *Cleptes nitidulus;* (b) *Chrysis zetterstedtï,* (c) *Paniscomima bekilyi;* (d) Vespidae: *Polistes dominula.* **(C483-0)** Rsf1 and first free abscissa of M (Mf1) aligned; (a) *Pristaulacus* sp.; (b) *Scolia erythropyga,* (c) *Oxybelus* sp. **(C483-1)** Rsf1 and Mf1 at angle, apex directed distally; (a) *Ampulicomorpha magna,* (b) *Ceratina* sp.; (c) *Stegomyrmex vizottoi.* **(C484-1)** Angle between Rsf1 and Mf1 narrow; †*Proapocritus elegans.* **(C485-1)** Rsf1 and Mf1 at angle, apex directed proximally; †*Anomopterella ampla.* **Families: (C481-1a, b)** Chrysididae. **(C482-1c)** Rhopalosomatidae. **(C482-1d)** Vespidae. **(C483-0a)** Aulacidae. **(C483-0b)** Scoliidae. **(C483-0c)** Crabronidae. **(C483-1a)** Embolemidae. **(C483-1b)** Apidae. **(C483-1c)** Formicidae. **(C484)** †Ephialtitoidea. **(C485-1)** †Anomopterellidae.

**Chars. 486-490.**
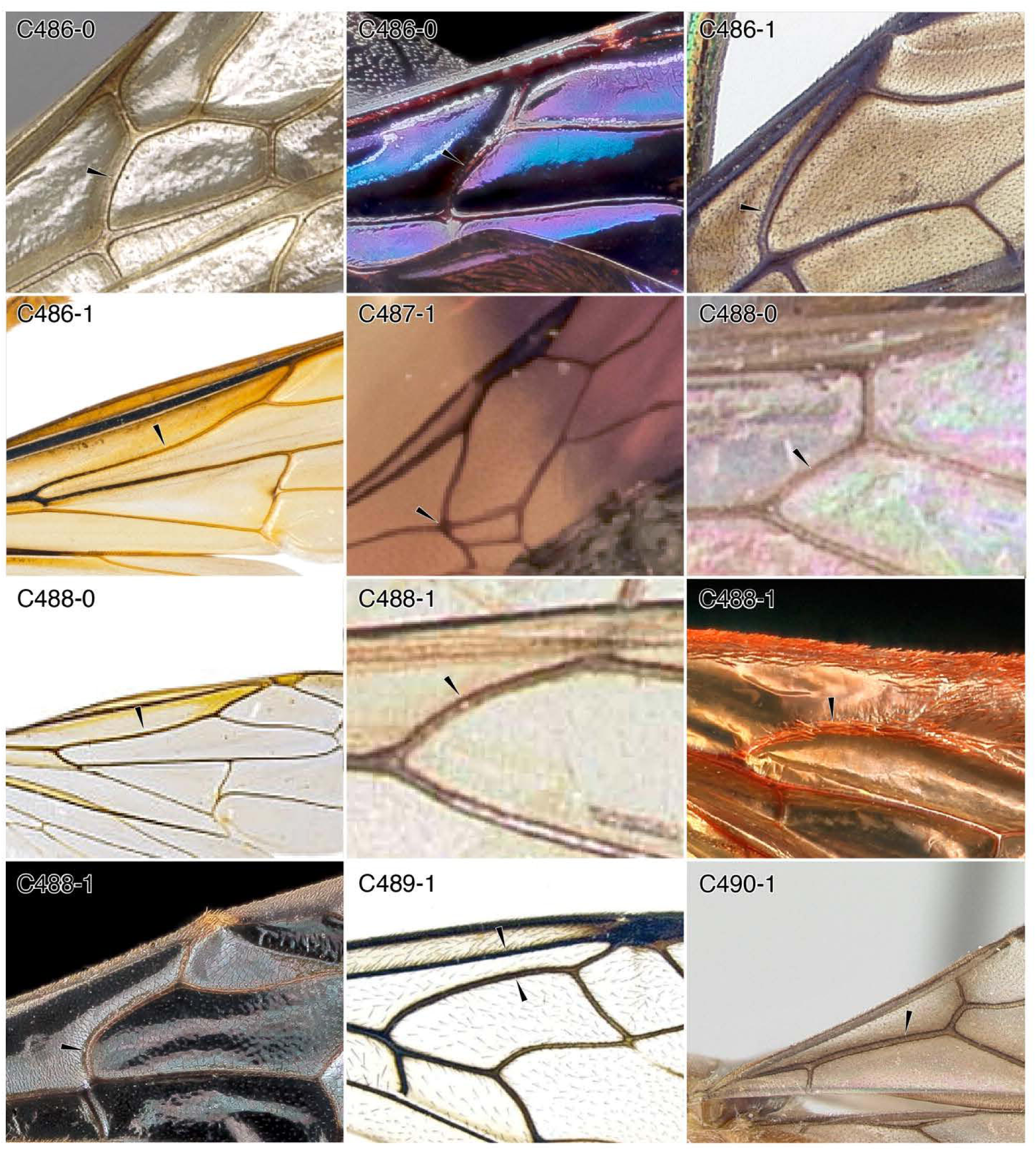
Character-states: (C486-0) Mf1 < 4 x length of Rsf1; (a) *Paraponera clavata,* (b) *Xylocopa* sp. **(C486-1)** Mf1 ≥ 4 x length of Rsf1; (a) *Chrysis pachysoma,* (b) *Vespa velutina nigrithorax.* **(C487-1)** Mf1 absent; †*Archeofoenus engeli.* **(C488-0)** Mf1 straight or curved with the vertex directed posterad; (a) *Pseudomyrmex gracilis;* (b) *Polistes dominula.* **(C488-1)** Mf1 curved with the vertex directed anterad; (a) *Tetraponera nigra,* (b) scoliid genus and species indet.; (c) *Lasioglossum aberrans.* **(C489-1)** Mf1 curved and closely approaching Sc+R+Rs; *Evania appendigaster.* **(C490-1)** M+Cu tubular; *Cladomyrma petalae.* **Families: (C486-0a, 4880-0a, 488-1a, 490)** Formicidae. **(C486-0b)** Apidae. **(C486-1a)** Chrysididae. **(C486-1b, 488-0)** Vespidae. **(C487)** Aulacidae. **(C488-1b)** Scoliidae. **(C488-1c)** Halictidae. **(C489)** Evaniidae.

**Chars. 490-495.**
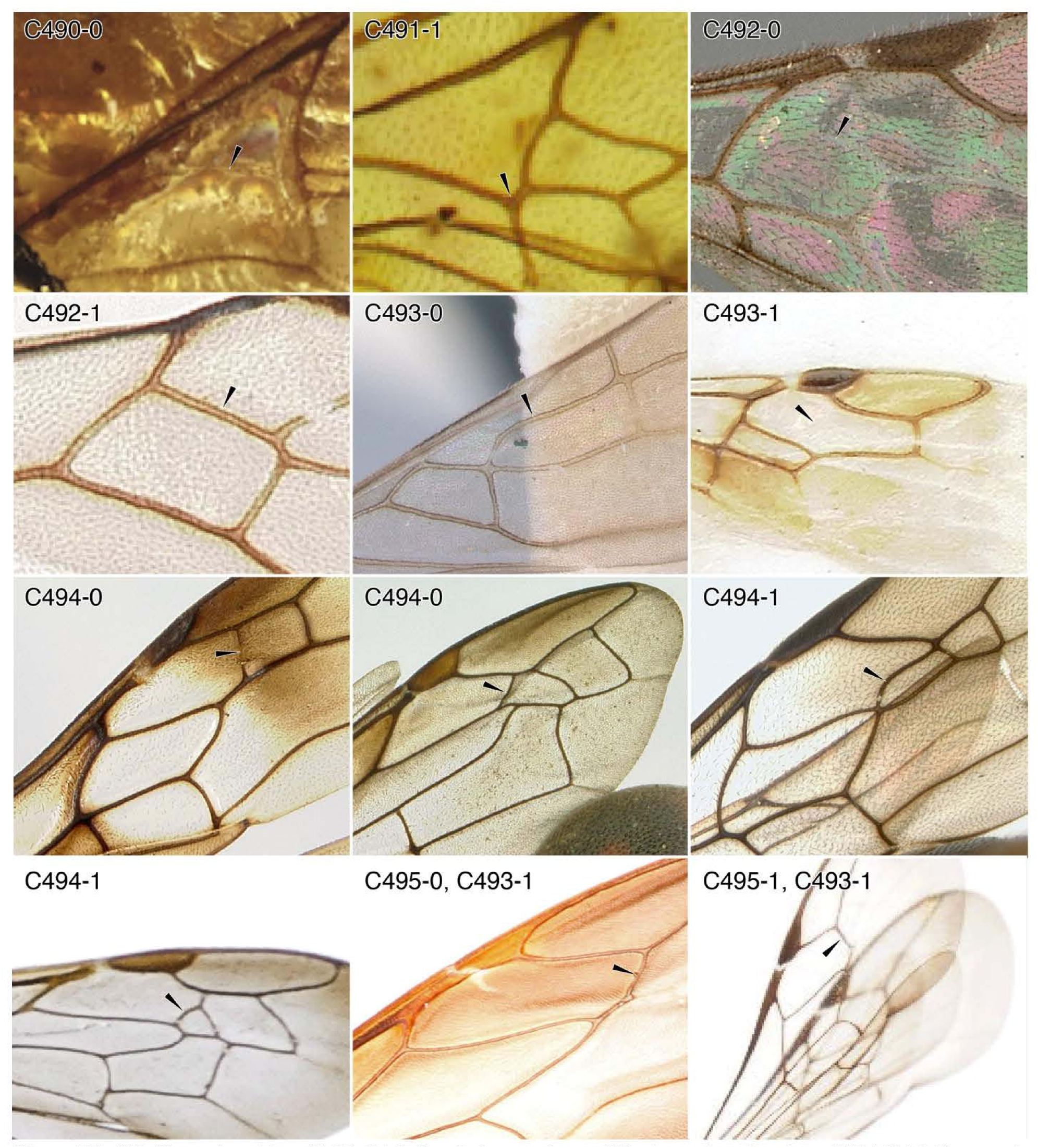
Character-states: (C490-0) M+Cu nebulous to absent; †*Hyptiogastrites electrinus.* **(C491-1)** M+Cu strongly angled from cu-a; †*Maimetshasia kachinensis.* **(C492-0)** Rs+M nebulous to absent; *Apenesia tagala.* **(C492-1)** Rs+M tubular; *Platythyrea mocquerysi.* **(C493-0)** Rsf2 present; *Aneuretus simoni.* **(C493-1)** Rsf2 absent; (a) *Ycaploca evansr,* (b) *Trypoxylon texense;*(c) *Try. clavicerum.* **(C494-0)** Rsf2 straight; (a) *Ampulex bredor;* (b) *Eumenes amoldi.* **(C494-1)** Rsf2 kinked or curved; (a) *Trigonalys* sp.; (b) *Cerceris sabulosa.* **(C495-0)** Rsf2 not perpendicular to long axis of wing; *Trypoxylon texense.* **(C495-1)** Rsf2 perpendicular or nearly so; *Try. clavicerum.* **Families: (C490-0)** Aulacidae. **(C491-1)** Trigonaloidea. **(C492-0)** Bethylidae. **(C492-1, 493-0)** Formicidae. **(C493-1a)** Scolebythidae. **(C494-0a)** Ampulicidae. **(C494-0b)** Vespidae. **(C494-1a)** Trigonalidae. **(C494-1b)** Philanthidae. **(C495, C493-1b, c)** Pemphredonidae.

**Chars. 495-502.**
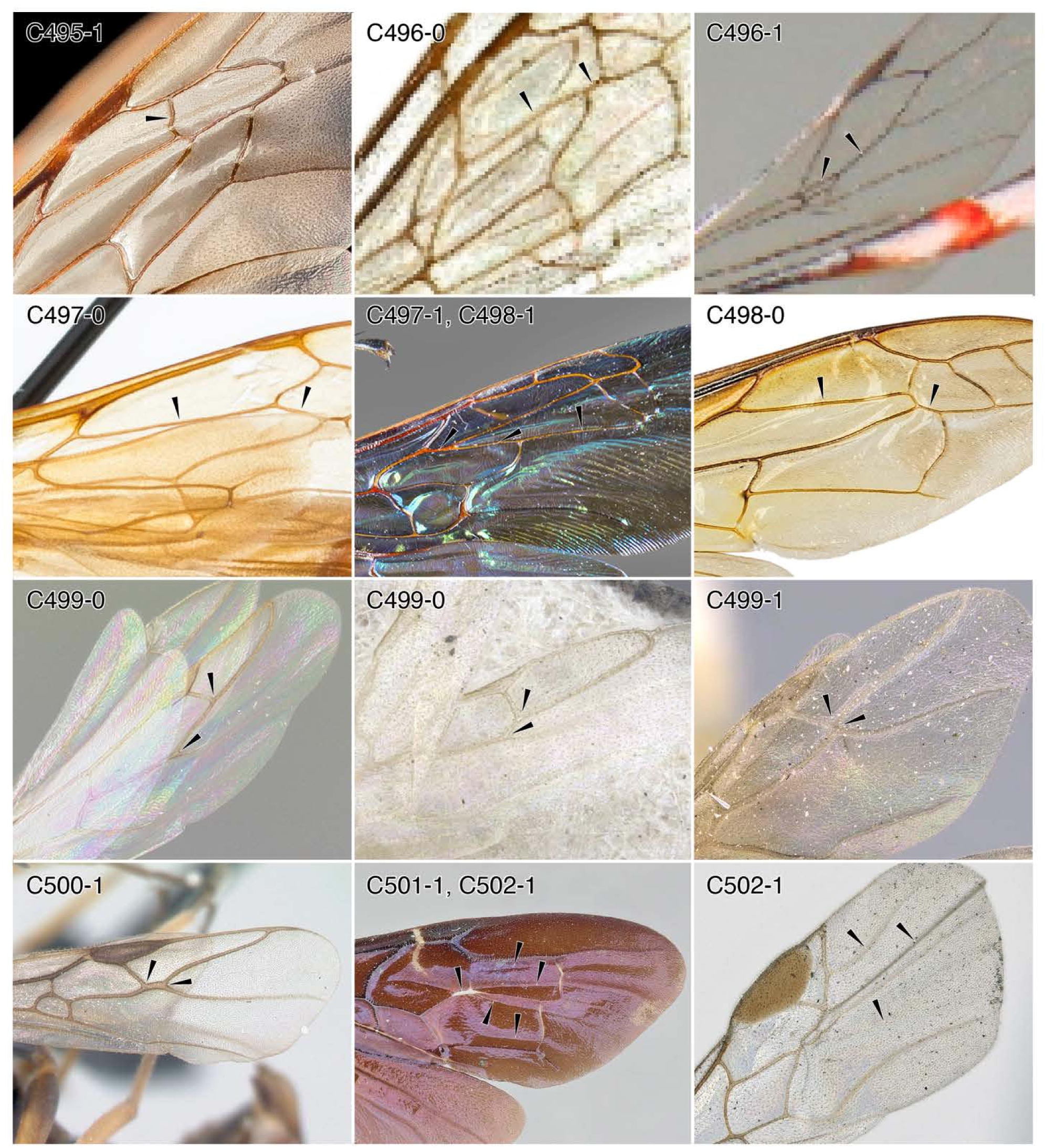
Character-states: (C495-1) Rsf2 perpendicular or nearly so; *Dasypoda* sp. **(C496-0)** Second abscissa of Rs+M, if present, shorter than first; *Plenoculus propinquus.* **(C496-1)** Rs+M[2] longer than first; *Gasteruption caucasicum.* **(C497-0)** Mf2 shorter than Rs+M; *Sceliphron caementarium.* **(C497-1)** Mf2 as long as or longer than Rs+M; *Megascolia procer.* **(C498-0)** Mf3 shorter than Rs+M; *Vespa velutina nigrithorax.* **(C498-1)** Mf3 as long or longer than Rs+M; *Me. procer.* **(C499-0)** Mf2 diverging from Rs+M proximal to 2r-rs; (a)*Myrmelachista ramulorum* (female); (b) *Myrmel. gallicola* (male). **(C499-1)** Mf2 diverging from Rs+M at or distal to 2r-rs; *Lasius californicus.* **(C500-1)** Mf2 diverging from Rs+M distal to 2r-rs; *Formica fusca.* **(C501-1)** Mf distal to Rs+M tubular; *Pristocera atopogamia.* **(C502-1)** Fore wing with 2-4 adventitious veins or creases; (a) *Pr. atopogamia,* (b) *Pr. atopogamia;* (c) *Myrmecopterina minor.* **Families: (C495)** Melittidae. **(C496-0)** Crabronidae. **(C496-1)** Gasteruptiidae. **(C497-0)** Sphecidae. **(C497-1, 498-1)** Scoliidae. **(C498-0)** Vespidae. **(C499, 500)** Formicidae. **(C501, 502-1a, b)** Bethylidae. **(C502-1c)** Plumariidae.

**Chars. 505-516.**
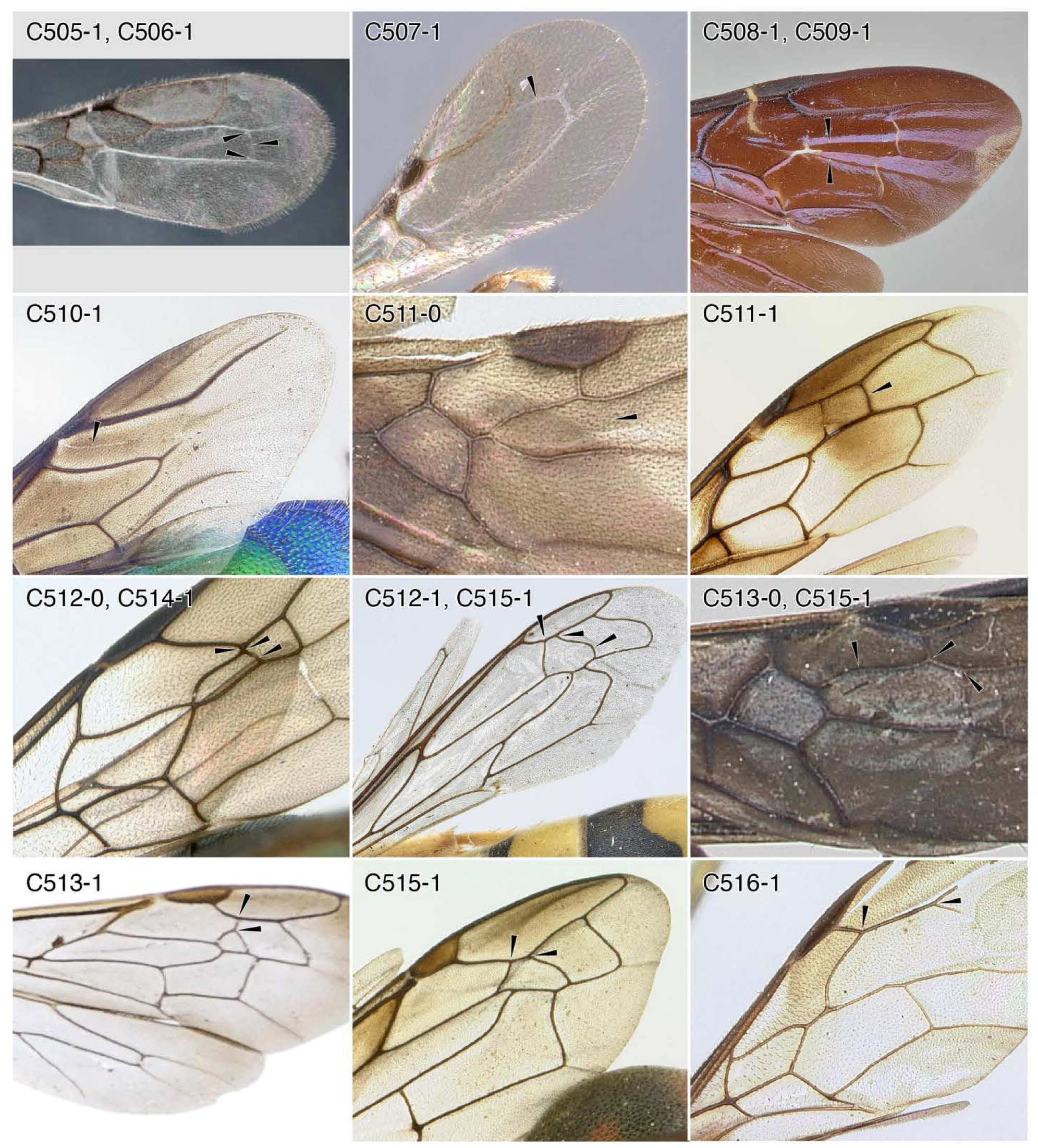
Character-states: (C505-1) Crease crossvein present; *Lytopsenella* sp. **(C506-1)** Crease crossvein joining anterior crease; *Lytopsenella* sp. **(C507-1)** Anterior crease joining Rsf; *Goniozus kiefferi.* **(C508-1)** Crease in location of Mf2 forked distally; *Pristocera atopogamia.* **(C509-1)** Mf present as spectral vein between forked crease; *Pristo. atopogamia.* **(C510-1)** Pterostigmal break crease extending to Mf; *Chrysis pachysoma.* **(C511-0)** 2rs-m nebulous to absent; *Tatuidris tatusia.* **(C511-1)** 2rs-m tubular; *Ampulex bredoi.* **(C512-0)** 2rs-m straight; *Trigonalys* sp. **(C512-1)** 2rs-m sinuate; *Stizus quadristrigatus.* **(C513-0)** 2rs-m joining Rs at or distal to 2r-rs; *Odontomachus bauri.* **(C513-1)** 2rs-m joining Rs distinctly proximad 2r-rs; *Cerceris sabulosa.* **(C514-1)** 2rs-m joining Rs at or near 2r-rs; *Trigonalys* sp. **(C515-1)** 2rs-m joining Rs distal to 2r-rs by less than one of its lengths; (a) *Stizus quadristrigatus;* (b) *Odontomachus baurr,* (c) *Eumenes amoldi.* **(C516-1)** 2rs-m joining Rs distal to 2r-rs by more than one of its length; *Pristaulacus africanus.* **Families: (C505-509)** Bethylidae. **(C510)** Chrysididae. **(C511-0, 513-0, 515-1b)** Formicidae. **(C511-1)** Ampulicidae. **(C512-0)** Trigonalidae. **(C512-1, 515-1a)** Bembicidae. **(C513-1)** Philanthidae. **(C514)** Trigonalidae. **(C515-1c)** Vespidae. **(C516)** Aulacidae.

**Chars. 517-523.**
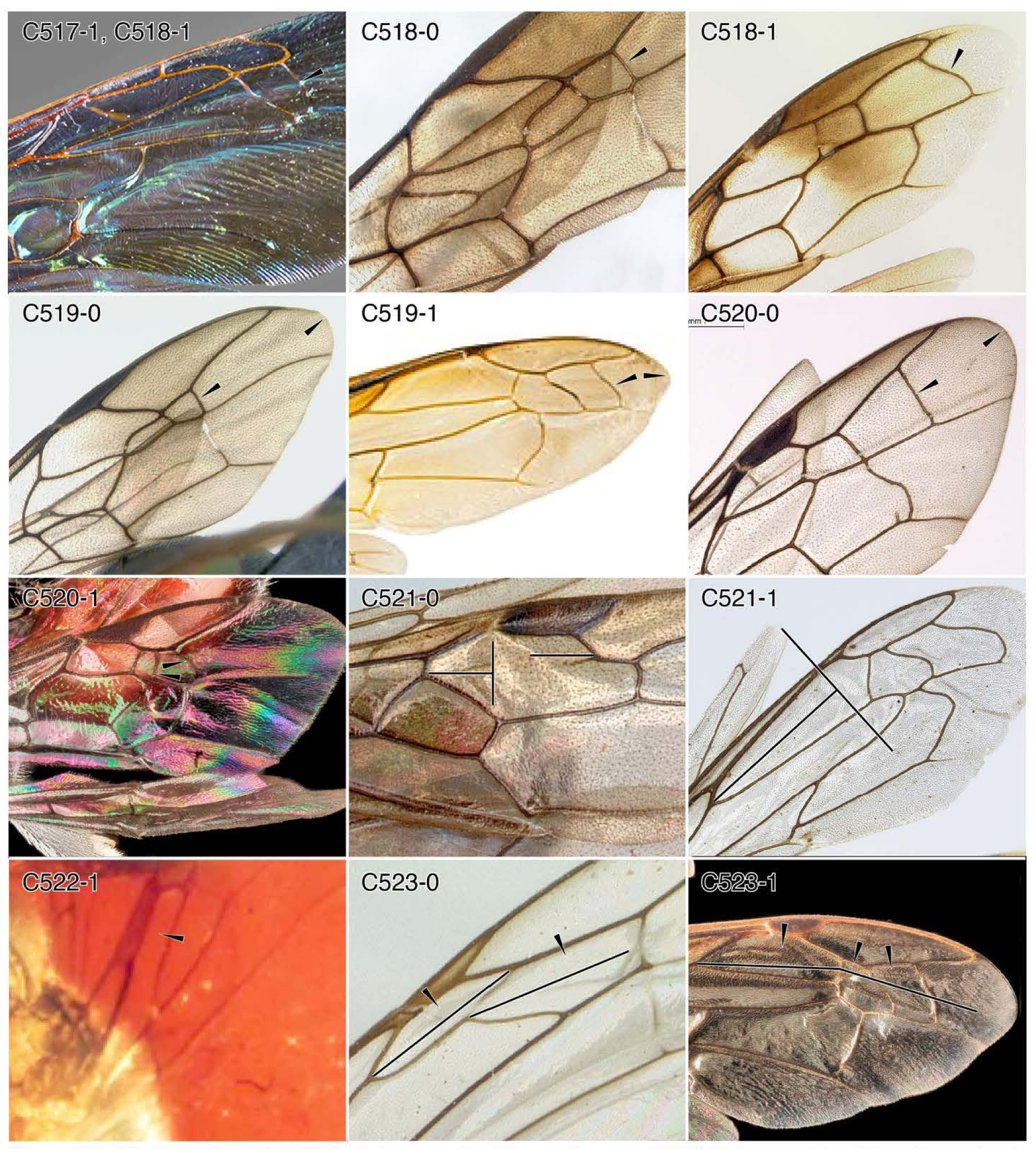
Character-states: (C517-1) 3rs-m tubular; *Megascolia procer.* **(C518-0)** 3rs-m straight; *Trigonalys natalensis.* **(C518-1)** 3rs-m sinuate; (a) *M. procer,* (b) *Ampulex bredoi.* **(C519-0)** 3rs-m distant from the distal wing margin; *T. natalensis.* **(C519-1)** 3rs-m close to distal wing margin by one or less than one of its lengths; *Vespa velutina nigrithorax.* **(C520-0)** 3rs-m distant from 2rs-m thus submarginal cell 3 (3rm) longer than wide; *Sapygina manica.* **(C520-1)** 3rs-m close to 2rs-m thus cell 3rm wider than long; *Sphecodes* sp. **(C521-0)** Prestigmal length of submarginal cell 1 (1rrs) less than half total cell length; *Ambly opone australis.* **(C521-1)** Prestigmal length of 1rrs greater than half total cell length; *Stizus quadristrigatus.* **(C522-1)** 1rrs extremely long and narrow; †*Cretevania bechlyi.* **(C523-0)** 1rrs as long as or shorter than submarginal cells 2 and 3 (3rsm, 4rsm); *Paniscomima bekilyi.* **(C523-1)** 1rrs longer than 3rsm, 4rsm; *Vespula squamosa.* **Families: (C517, C518-0a)** Scoliidae. **(C518-0b, 519-0)** Trigonalidae. **(C518-1)** Ampulicidae. **(C519-1, 523-1)** Vespidae. **(C520-0)** Sapygidae. **(C520-1)** Apidae. **(C521-0)** Formicidae. **(C521-1)** Bembicidae. **(C522-1)** Evaniidae. **(C523-0)** Rhopalosomatidae.

**Chars. 524-528.**
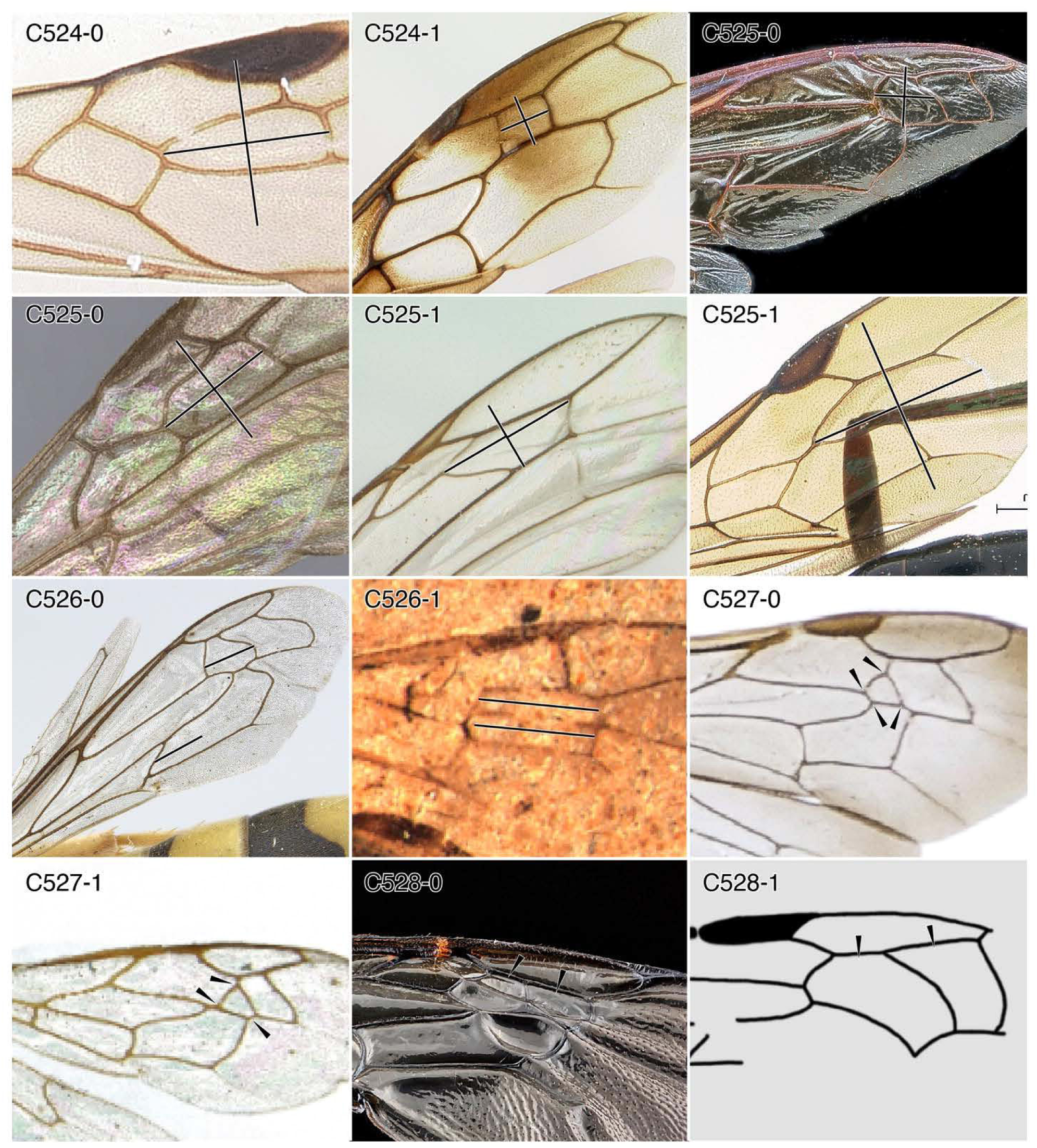
Character-states: (C524-0) 3rsm distinctly longer than wide; *Platythyrea mocquerysi.* **(C524-1)** 3rsm nearly as wide as long or wider than long; *Ampulex bredoi.* **(C525-0)** 3rsm length < 3 x width; (a) *Vespula mandarinia;* (b) *Diacamma pallidum.* **(C525-1)** 3rsm length ≥ 3 x width; (a) *Paniscomima bekilyi;* (b) *Pristaulacus pilotoi.* **(C526-0)** 3rsm not equal in size to subdiscal cell 1 (2mcu); *Stizus quadristrigatus.* **(C526-1)** 3rsm equal in size and shape 2mcu; †*Anomopterella coalita.* **(C527-0)** 3rsm with > 3 sides or not anteriorly “petiolate”; *Cerceris sabulosa.* **(C527-1)** 3rsm with three sides and anteriorly petiolate; *Plenoculus davisi.* **(C528-0)** 3rsm and 4rsm dissimilar in shape and proportions or not wider than long and curved; *Habropoda laboriosa.* **(C528-1)** 3rsm and 4rsm similar in shape and proportions, wider than long and curved; †*Cretobestiola communis.* **Families: (C524-0)** Sapygidae. **(C524)** Ampulicidae. **(C525-0a)** Vespidae. **(C525-0b)** Formicidae. **(C525-1a)** Rhopalosomatidae. **(C525-1b)** Aulacidae. **(C526-0)** Bembicidae. **(C526-1)** †Anomopterellidae. **(C527-0)** Philanthidae. **(C527-1)** Crabronidae. **(C528-0)** Apidae. **(C528-1)** †Angarosphecidae.

**Chars. 529-534.**
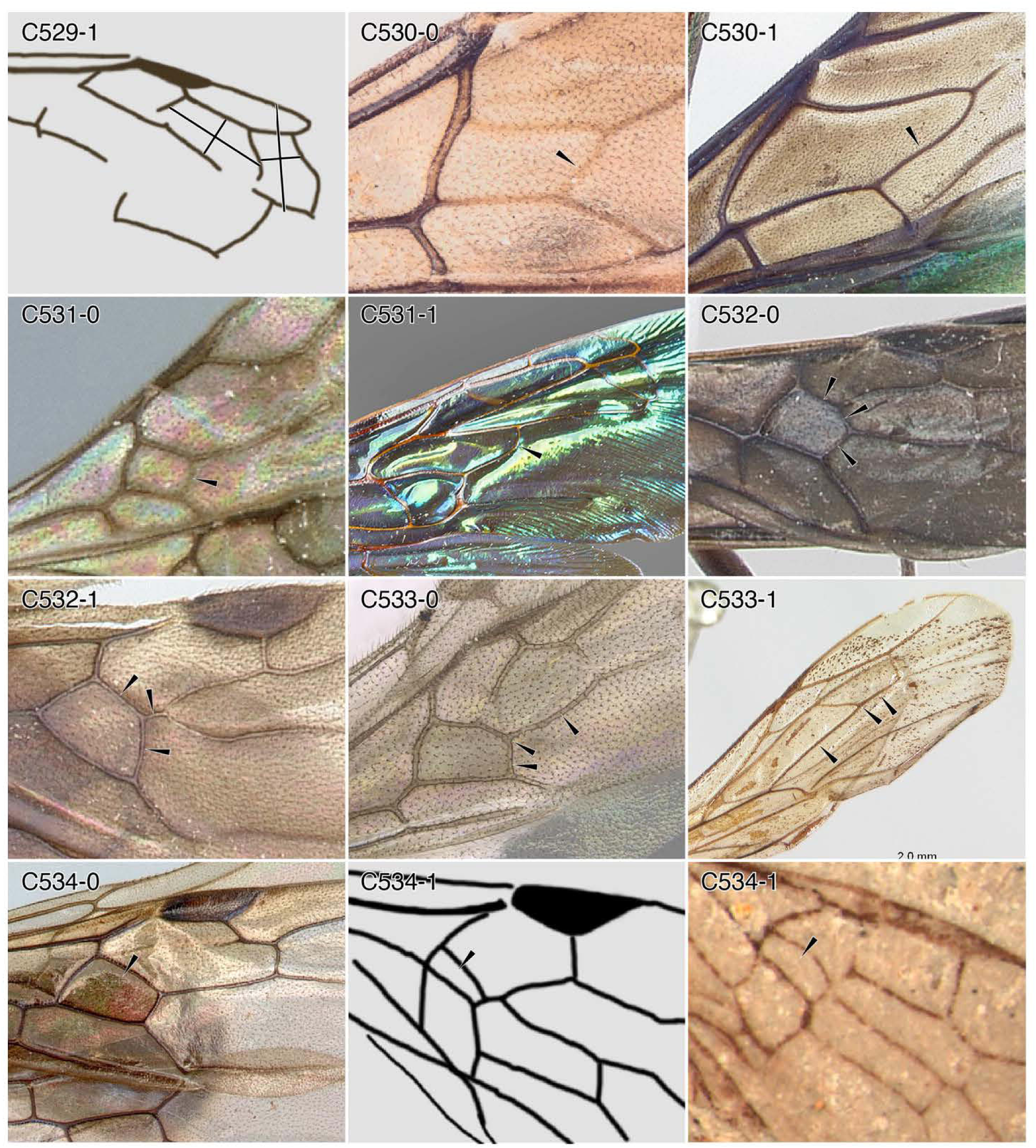
Character-states: (C529-1) 4rsm smaller than 3rsm; †*Cretosphecium lobatum.* **(C530-0)** 1m-cu nebulous to absent; *Cleptes semicyaneus.* **(C530-1)** 1m-cu tubular; *Chrysis pachysoma.* **(C531-0)** 1m-cu straight to weakly curved; *Ampulicomorpha magna.* **(C531-1)** 1m-cu distinctly curved; *Megascolia procer.* **(C532-0)** lm-cu anterior apex distal to distal Rs+M split; *Odontomachus bauri.* **(C532-1)** lm-cu anterior apex proximal to distal Rs+M split; *Tatuidris tatusia.* **(C533-0)** lm-cu and Mf3 not forming straight line; *Fulakora orizabana.* **(C533-1)** lm-cu and Mf3 forming straight line; *Paniscomima bekilyi.* **(C534-0)** lmcu cell not small, elongate; *Amblyopone australis.* **(C534-1)** lmcu very small, rectangular, and with length > 3 x width; (a) †*Mesepipolaea nanligezhuangica;* (b) †*Anomopterella brachystelis.* **Families: (C529-1)** †Angarosphecidae. **(C530)** Chrysididae. **(C531-0)** Embolemidae. **(C531-1)** Scoliidae. **(C532, 533-0, 543-0)** Formicidae. **(C533-1)** Rhopalosomatidae. **(C534-1a)** †Baissidae. **(C534-1b)** †Anomopterellidae.

**Chars. 535-540.**
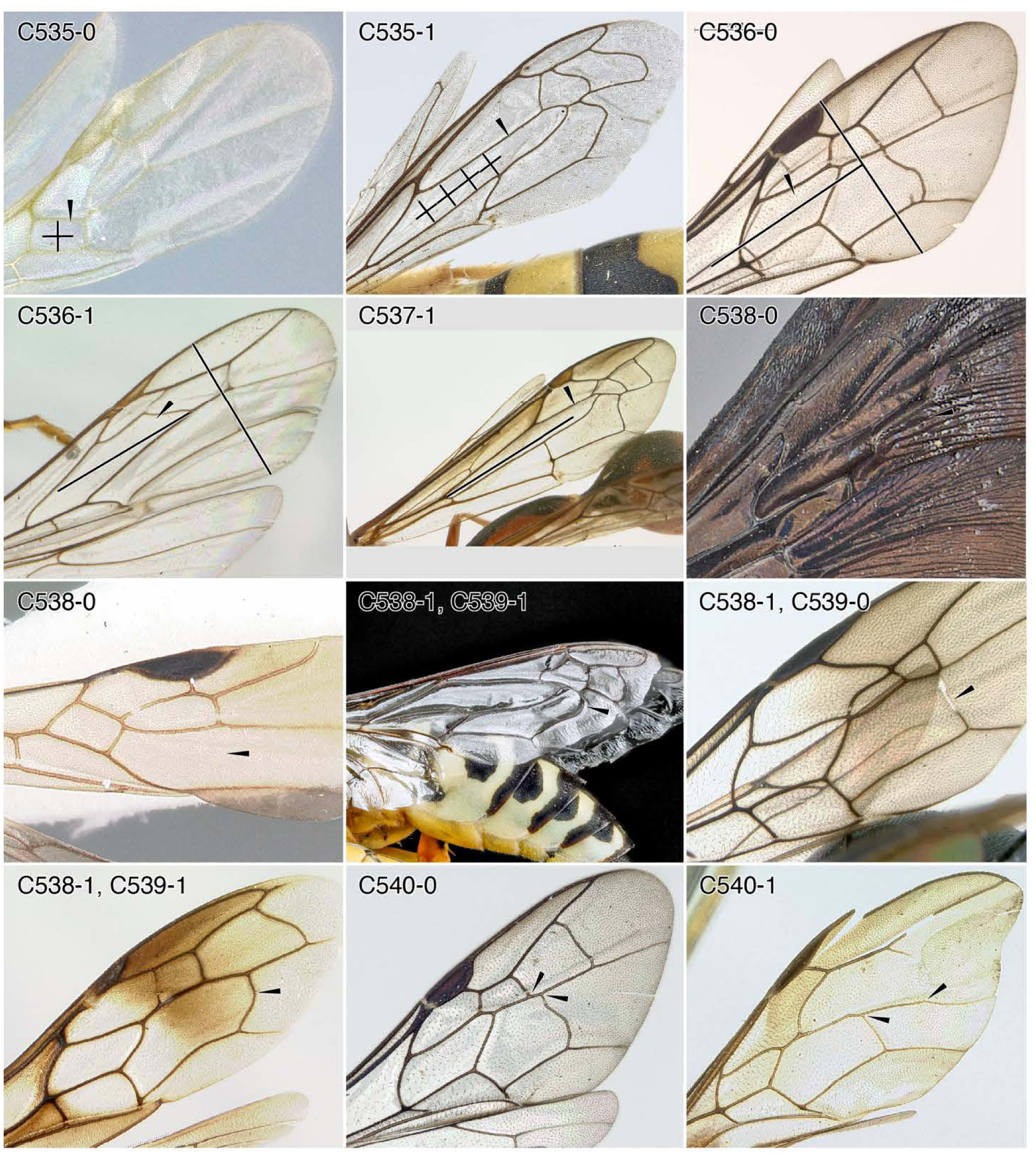
Character-states: (C535-0) lmcu length < 4 x width; *Hypoponera distinguenda.* **(C535-1)** lmcu length ≥ 4 x width; *Stizus quadristrigatus.* **(C536-0)** lmcu shorter than maximum width of wing; *Sapygina manica.* **(C536-1)** lmcu longer than maximum width of wing; *Paniscomima bekilyi.* **(C537-1)** lmcu extremely long, length > 2/5 x that of the wing; *Eumenes amoldi.* **(C538-0)** 2m-cu nebulous to absent; (a) *Scolia erythropyga;* (b) *Platythyrea mocquerysi.* **(C538-1)** 2m-cu tubular; (a) bembicid genus and species indet.; (b) *Trigonalys* sp.; (c) *Ampulex bredoi.* **(C539-0)** 2m-cu straight to weakly curved; *Trigonalys* sp. **(C539-1)** 2m-cu sinuate; (a) bembicid genus and species indet; (b) *A. bredoi.* **(C540-0)** 2m-cu joining M distad 2r-sm; *Sa lugens.* **(C540-1)** 2m-cu joining M at or proximad 2r-sm; *Pristaulacus africanus.* **Families: (C535-0, 538-0b)** Formicidae. **(C535-1, 538-1a, 539-1a)** Bembicidae. **(C536-0, 540-0)** Sapygidae. **(C536-1)** Rhopalosomatidae. **(C537-1)** Vespidae. **(C538-0a)** Scoliidae. **(C538-1b, 539-0)** Trigonalidae. **(C538-1c, 539-1b)** Ampulicidae. **(C540-1)** Aulacidae.

**Chars. 541-549.**
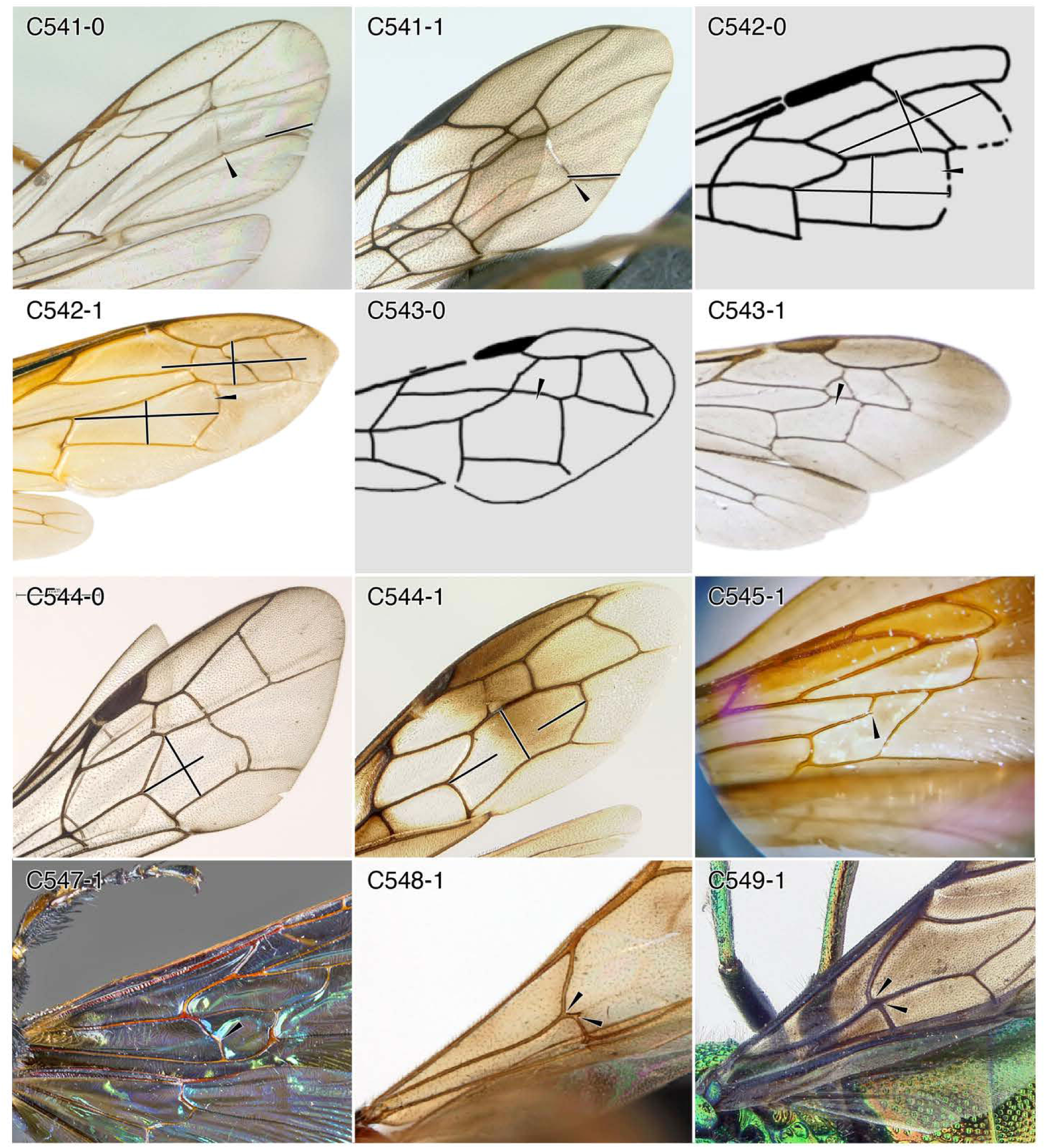
Character-states: (C541-0) 2m-cu distant from wing apex; *Paniscomima bekilyi.* **(C541-1)** 2m-cu close to wing apex, within one of its lengths; *Trigonalys* sp. **(C542-0)** 2mcu cell about same size are 3rsm and 4rsm or if 4rsm absent, then 2mcu size < 2 x that of 3rsm; †*Cretoscolia laiyangica.* **(C542-1)** 2mcu distinctly larger than 3rsm and 4rsm or if 4rsm absent, then 2mcu size > 2 x that of 3rsm; *Vespa velutina nigrithorax.* **(C543-0)** 2mcu as wide as long or wider than long; †*Archisphex boothi.* **(C543-1)** 2mcu longer than wide; *Cerceris sabulosa.* **(C544-0)** 2mcu length < 2 x width; *Sapygina manica.* **(C544-1)** 2mcu length ≥ 2 x width; *Ampulex bredoi.* **(C545-1)** 2mcu with adventitious vein or veinlet; *Campsomeris pilipes.* **(C547-1)** lcu-a sinuate; *Megascolia procer.* **(C548-1)** lcu-a anterior junction distal to M+Cu split; *Dryinus bellicosus.* **(C549-1)** lcu-a anterior junction at M+Cu split; *Chrysis pachysoma.* **Families: (C541-0)** Rhopalosomatidae. **(C541-1)** Trigonalidae. **(C542-0)** Scolioidea. **(C545, 547)** Scoliidae. **(C542-1)** Vespidae. **(C543-0)** †Angarosphecidae. **(C543-1)** Philanthidae. **(C544-0)** Sapygidae. **(C544-1)** Ampulicidae. **(C548)** Dryinidae. **(C549)** Chrysididae.

**Chars. 550-562.**
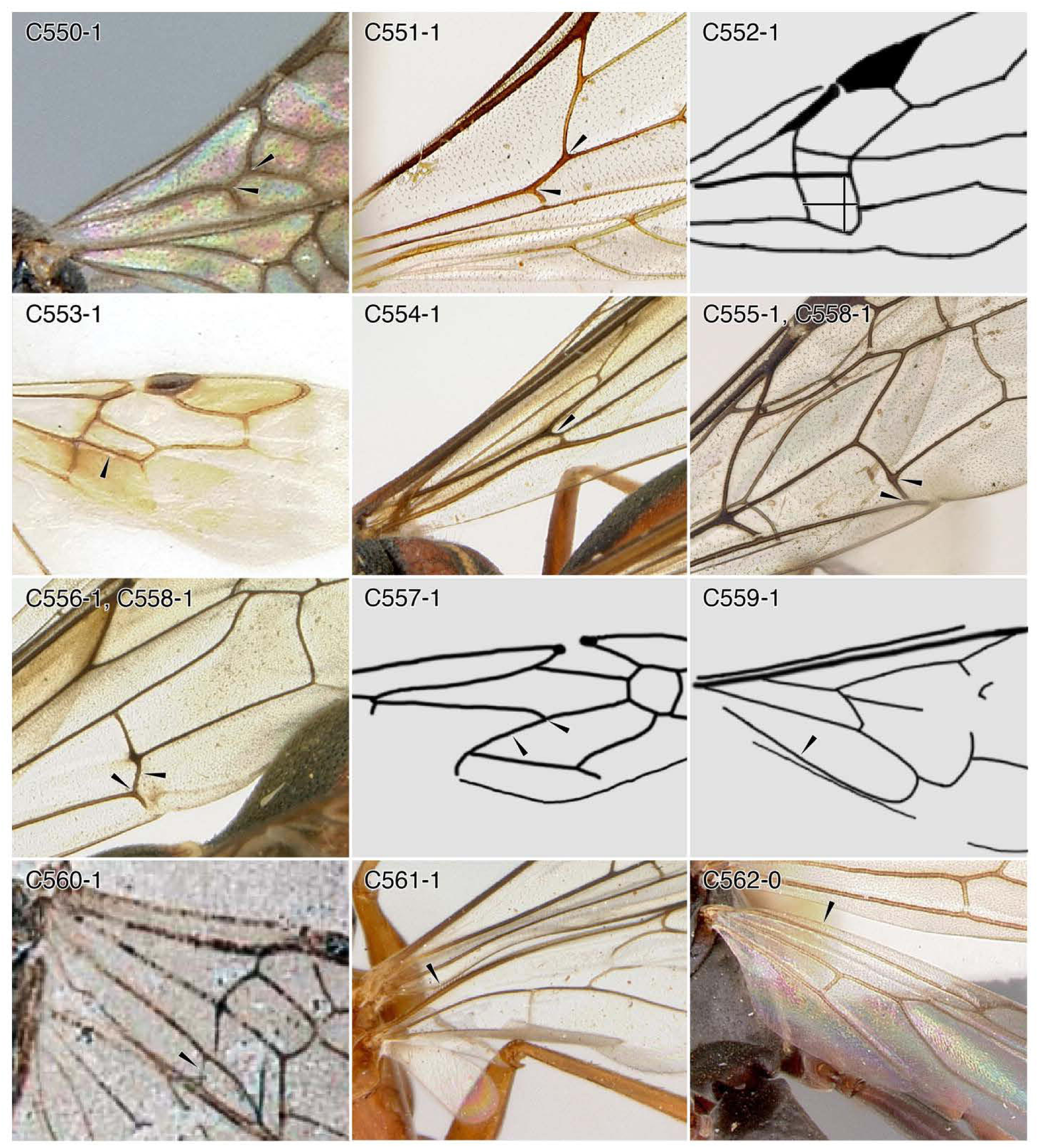
Character-states: (C550-1) lcu-a anterior junction proximal to M+Cu split; *Ampulicomorpha magna.* **(C551-1)** 1 cu-a anterior junction very proximal to M+Cu split; *Dolichurus basuto.* **(C552-1)** 2cua wider than long; †*Manlaya anglica.* **(C553-1)** Free cubitus **(Cuf)** tubular; *Ycaploca evansi.* **(C554-1)** Cuf1 angled posteroapically from M+Cu, length subequal to cu-a; *Eumenes amoldi.* **(C555-1)** Distal split of Cu at approximate right angle; *Sapygina lobengulae.* **(C556-1)** Posterior branch of Cu directed postero-proximally; *E. amoldi.* **(C557-1)** Second abscissa of Cuf **(Cuf2)** directed posteriorly and at acute angle relative to Cuf1 ; †*Curiosivespa derivata.* **(C558-1)** Posterior branch of Cu reaching 1 A; (a) *Sapygina lobengulae;*(b) *E. arnoldi.* **(C559-1)** Second anal vein (2A) present; †*Nevania delicata.* **(C560-1)** la-a crossvein present; †*Praeproapocritus flexus.* **(C561-1)** Hind wing (HW) basal hamuli present; *Paniscomima bekilyi.* **(C562-0)** HW costal vein **(C) absent;** *Platythyrea mocquerysi.* **Families: (C550)** Embolemidae. **(C551)** Ampulicidae. **(C552)** †Baissidae. **(C553)** Scolebythidae. **(C554, 556, 557, 558-1b)** Vespidae. **(C558-1a)** Sapygidae. **(C559-1)** †Praeaulacidae. **(C560)** †Ephialtitoidea. **(C561)** Rhopalosomatidae. **(C562)** Formicidae.

**Chars. 562-576.**
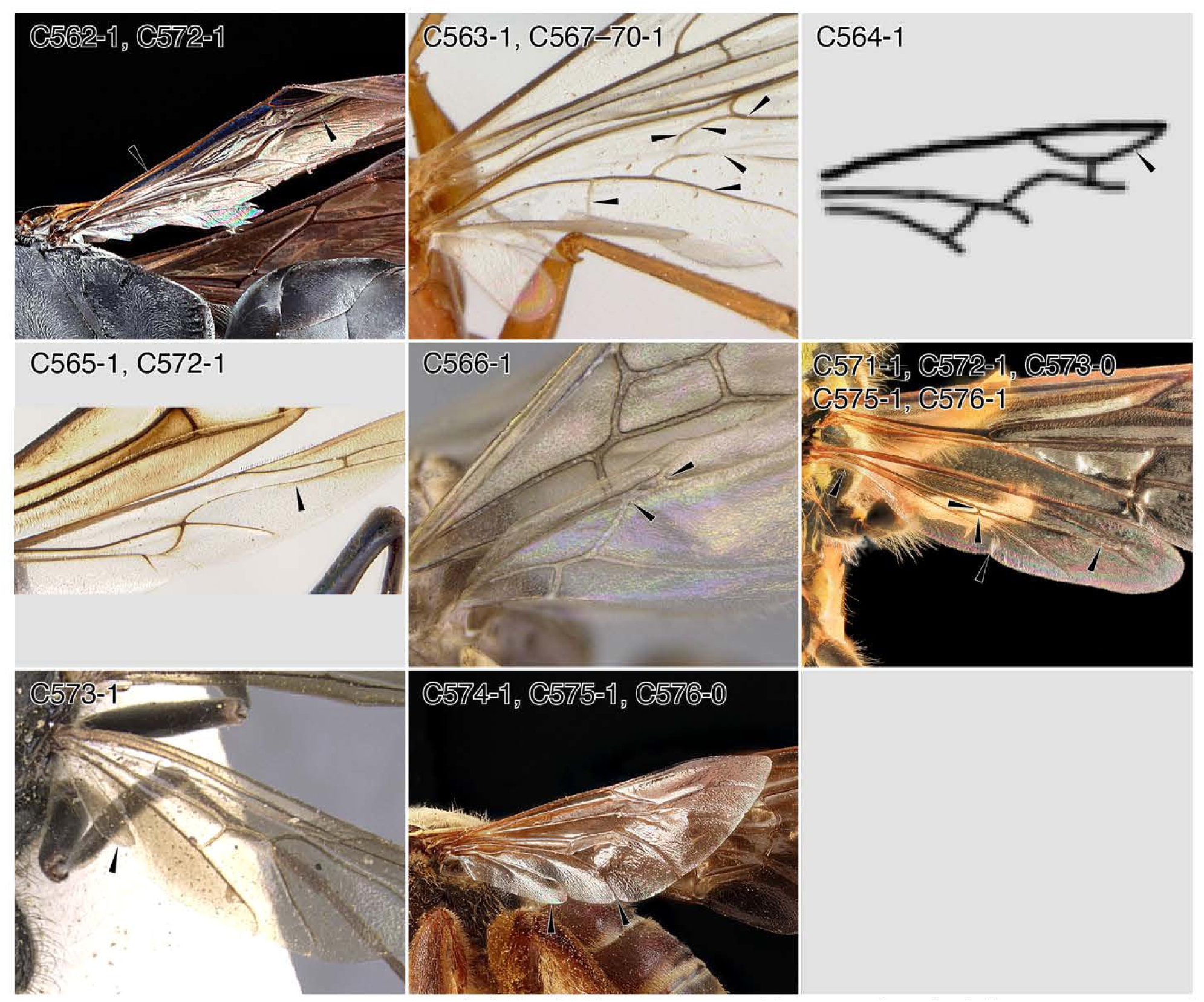
Character-states: (C562-1) Hind wing (HW) C present; Larrini genus and species indet. **(C563-1)** HW Rsf present; *Paniscomima bekilyi.* **(C564-1)** HW Rsf curving anteriorly and enclosing distal cell; †*Leptephialtites pallidus.* **(C565-1)** HW lrs-m tubular; *Ampulex bredoi.* **(C566-1)** HW lrs-m joining R+RS proximal to Rf and Rsf split; *Linepithema humile.* **(*C567-****1*) *HW lrs-m curving strongly* (*or angled*) *between Mf2 and Rsf and meeting Rsf at oblique angle; P. bekilyi.* **(C568-1)** HW Mf2 tubular; *P. bekilyi.* **(C569-1)** HW Cuf tubular; *P. bekilyi.* **(C570-1)** HW cu-a present; *P. bekilyi.* **(C571-1)** HW cu-a joining Cu distal to M+Cu split; *V. squamosa.* **(C572-1)** “Basal cell” (lrsm) distally elongate and narrow; (a) Larrini genus and species indet; (b) *Ampulex bredor,* (c) *V. squamosa.* **(C573-0)** Jugal lobe absent; *V. squamosa.* **(C573-1)** Jugal lobe present; *Neoponera villosa.* **(C574-1)** Jugal lobe large, nearly as long as anal area; *Dieunomia xerophila.* **(C575-1)** Plical lobe marked by apical incision; (a) *V. squamosa.* (b) *D. xerophila.* **(C576-0)** Apical incision of plica moderate to deep; *V. squamosa.* **(C576-1)** Apical incision of plica shallow; *D. xerophila.* **Families: (C562, 572-1b)** Crabronidae. **(C563, 567-570)** Rhopalosomatidae. **(C564)** †Ephialtitoidea. **(C565, 572)** Ampulicidae. **(C566, 573-1)** Formicidae. **(C571, 572, 573-0, 575-1a, 576-0)** Vespidae. **(C574, 575-1b, 576-1)** Halictidae.

